# Variant-Specific Rewiring of Lysosomal Positioning by STARD9 Produces Opposite Cholesterol Trafficking Outcomes

**DOI:** 10.64898/2026.09.25.754437

**Authors:** Mehdi Dianatpour, Dipanjan Banerjee, Hyun Kyung Kim, Jiyoung Moon, Rui Zhu, Hang Zhou, Arpita Neogi, Trojan Rugira, Shrikant Mane, Adebowale Adeniran, Brisa Palikuqi, William Lee, Lucila Ohno Machado, Arya Mani

## Abstract

Residual cardiovascular risk persists despite lipid-lowering therapy, independent of LDL and Lp(a), reflecting cholesterol dysregulation. We identify STARD9, a lysosomal cholesterol-sensing kinesin, as a causal gene for divergent familial dyslipidemias and premature atherosclerosis. Its START domain binds cholesterol to govern lysosomal positioning. Rare variants segregate with autosomal dominant hypercholesterolemia and premature coronary disease, while common variants associate with reduced HDL and elevated triglycerides. Both converge on lysosomal cholesterol sequestration, ER depletion, mTORC1–SREBP2 activation, impaired autophagy, and NF-κB inflammation, yet diverge through opposite lysosomal positioning: the rare variant disperses lysosomes peripherally, phenocopying START-domain loss across cholesterol sequestration, lysosomal positioning, and TFEB activation, triggering LDLR degradation and CASM/TFEB-dependent efflux that preserves HDL, whereas the common variant causes perinuclear retention that suppresses TFEB-dependent lipid-handling transcripts, yielding hypertriglyceridemia and low HDL. In Stard9-knockout mice, enterocyte cholesterol sequestration drives SREBP2-dependent NPC1L1 upregulation and apical localization, increasing intestinal cholesterol absorption. These findings position lysosomal cholesterol trafficking as a targetable node in statin-refractory cardiovascular disease.

**Graphical abstract:** (Left panel) STARD9 mutations p.V2821L and p.I2772V drive divergent cholesterol uptake but converge on a shared lysosomal pathology. While p.V2821L reduces LDL uptake and p.I2772V increases it, both mutations cause lysosomal cholesterol sequestration. This accumulation leads to lysosomal membrane damage and nuclear SREBP2 localization, exacerbating cholesterol dysregulation and cellular stress. Lysosomal damage also disperses lysosomes and activates mTOR, suppressing canonical autophagy while driving NF-κB inflammatory signaling and pro-inflammatory cytokine secretion. (Right panel) Concurrently, lysosomal stress engages CASM specifically in p.V2821L lines, driving paradoxical TFEB activation and increased cholesterol efflux, whereas failure to engage this pathway in p.I2772V lines is associated with reduced TFEB activation and reduced cholesterol efflux.

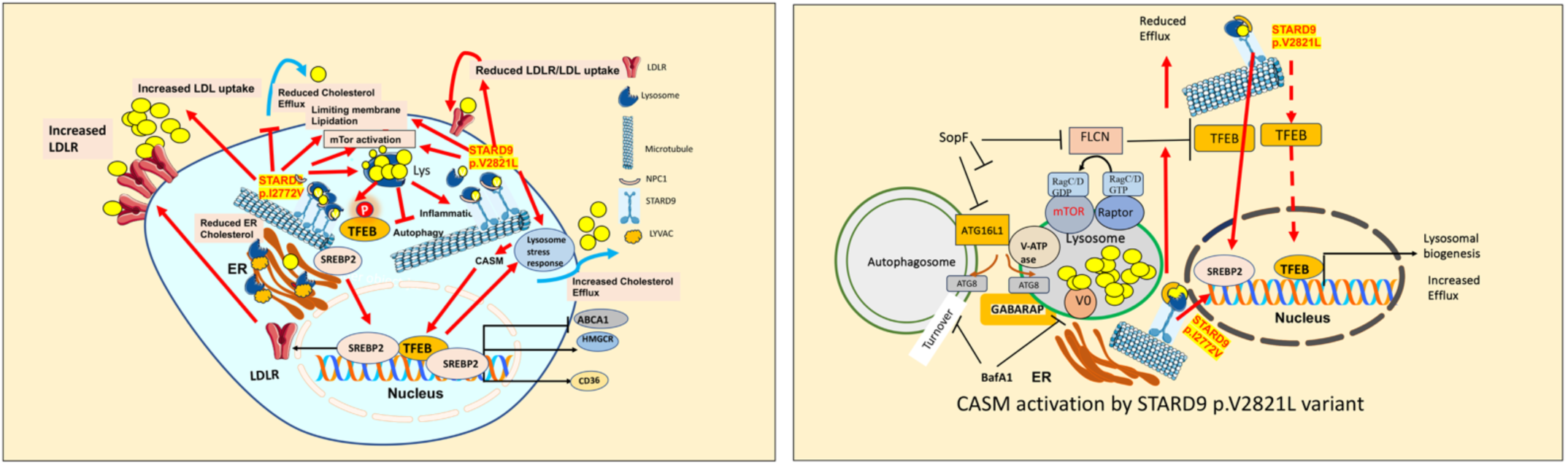

## Introduction

Atherosclerotic cardiovascular disease (ASCVD) remains the leading cause of death worldwide despite widespread use of statins and other lipid-lowering therapies^1^. A substantial proportion of treated patients continue to experience myocardial infarction and stroke, a phenomenon termed residual cardiovascular risk^2^, which cannot be explained by persistent LDL cholesterol elevation alone and is only partially caused by elevated Lp(a). Familial dyslipidemias confer markedly elevated risk of premature ASCVD and contribute substantially to residual risk^3,4^, yet the genetic basis and molecular mechanisms for the majority remain unknown; classical monogenic forms account for only ∼5–10% of cases^5^, suggesting the existence of undiscovered pathways.

To identify novel genetic contributors to familial dyslipidemia, we performed exome sequencing in multiplex kindreds presenting with unexplained hypercholesterolemia and premature atherosclerosis in whom classical monogenic causes had been excluded. Rare variant burden analysis identified *STARD9* as the top candidate gene segregating with autosomal dominant hypercholesterolemia and premature coronary artery disease across independent kindreds. Complementary genome-wide association analyses in large population cohorts independently identified common variants at the *STARD9* locus associating with atherogenic dyslipidemia at genome-wide significance, and cross-cohort colocalization confirmed convergence of these signals on *STARD9*.

*STARD9* encodes is a largely uncharacterized kinesin-3 family member^6,7^ containing a StAR-related lipid-transfer (START) domain, that functions as a kinesin. It NPC1 to regulate lysosomal position and tubulation^8^. Using an NBD-cholesterol binding assay, we verified that it also binds cholesterol and contributes to STARD9–NPC1 complex formation and proper lysosomal positioning: deleting the START domain redistributed lysosomes to the cell periphery, a finding that is consistent with an earlier report that the C-terminal tail domain of kinesin-1 autoinhibits the motor head and its removal unmasks processive, plus-end-directed motility^9^.

Recessive mutations in *STARD9* cause a rare neurodevelopmental syndrome with intellectual disability^10^, but whether common or rare coding variants in *STARD9* contribute to dyslipidemia or atherosclerosis, and by what mechanism, has been unknown. Here, integrating rare-variant segregation analysis, population genetics, CRISPR–Cas9 isogenic knock-in modeling, single-cell transcriptomics of patient-derived immune cells, and in vivo mouse models, we establish STARD9 as a lysosomal cholesterol sensor coupling sterol content to kinesin motor activity and spatial organization. A rare familial variant (p.V2821L) and a GWAS variant rs202077402 (p.I2772V) converge on lysosomal cholesterol sequestration and mTORC1–SREBP2 dysregulation but diverge in their effects on lysosomal positioning. The p.V2821L variant phenocopies the loss of the START domain by producing the same peripheral lysosomal dispersion phenotype, which suggests that this variant disrupts the START-domain autoinhibitory function and drives unopposed movement of lysosomes toward the microtubule plus end.

This subsequently results in CASM (conjugation of ATG8 proteins to single membranes) engagement, TFEB activation, reduced cholesterol uptake and normal cholesterol efflux, producing opposing plasma lipid phenotypes compared to the GWAS-associated variant, which exhibits minus-end activity, resulting in normal LDL uptake but reduced TFEB-dependent transcript regulation and diminished cholesterol efflux. In *Stard9* knockout mice, we identify a previously unrecognized intestinal axis of dyslipidemia driven by the STARD9– NPC1L1 interaction and SREBP2-mediated NPC1L1 upregulation, explaining the observation that rare variant carriers selectively respond to ezetimibe. Together, these findings demonstrate how distinct functional alterations within the same gene produce heterogeneous lipid phenotypes, reveal a previously unrecognized lysosomal cholesterol trafficking pathway underlying familial dyslipidemia and statin-refractory intracellular inflammation, and identify STARD9 as a mechanistically actionable therapeutic target.

## Results

### Identification of *STARD9* Variants in Familial Dyslipidemia

Exome sequencing of the extended kindred of an index case with familial hypercholesterolemia (kindred A, Fig. 1A), who was intolerant to high-dose statin, sustained a myocardial infarction due to acute plaque rupture in the LAD, and had no variants identified in known familial hypercholesterolemia (FH) genes, revealed a rare heterozygous STARD9 missense variant (chr15:42690039, G>C, GRCh38) causing substitution of a highly conserved valine with leucine (p.V2821L). Strikingly, he subsequently showed a dramatic response to ezetimibe, normalizing his LDL cholesterol. The variant segregated perfectly with hypercholesterolemia (LDL-C ≥190 mg/dL on no drugs, or >130 mg/dL on statins) across three generations, with no other shared deleterious variants remaining after stringent filtering. The variant is extremely rare (AF <10^-5^), and is predicted to be damaging (PolyPhen-2^11^, SIFT 4G ^12^; CADD^13^ 22.7). ENCODE annotation revealed that this variant lies within a predicted HNF1α binding site (ENCODE Consortium, 2012)^14^. All family members had Lp(a) less than 20 mg/dl.

**Figure 1:**
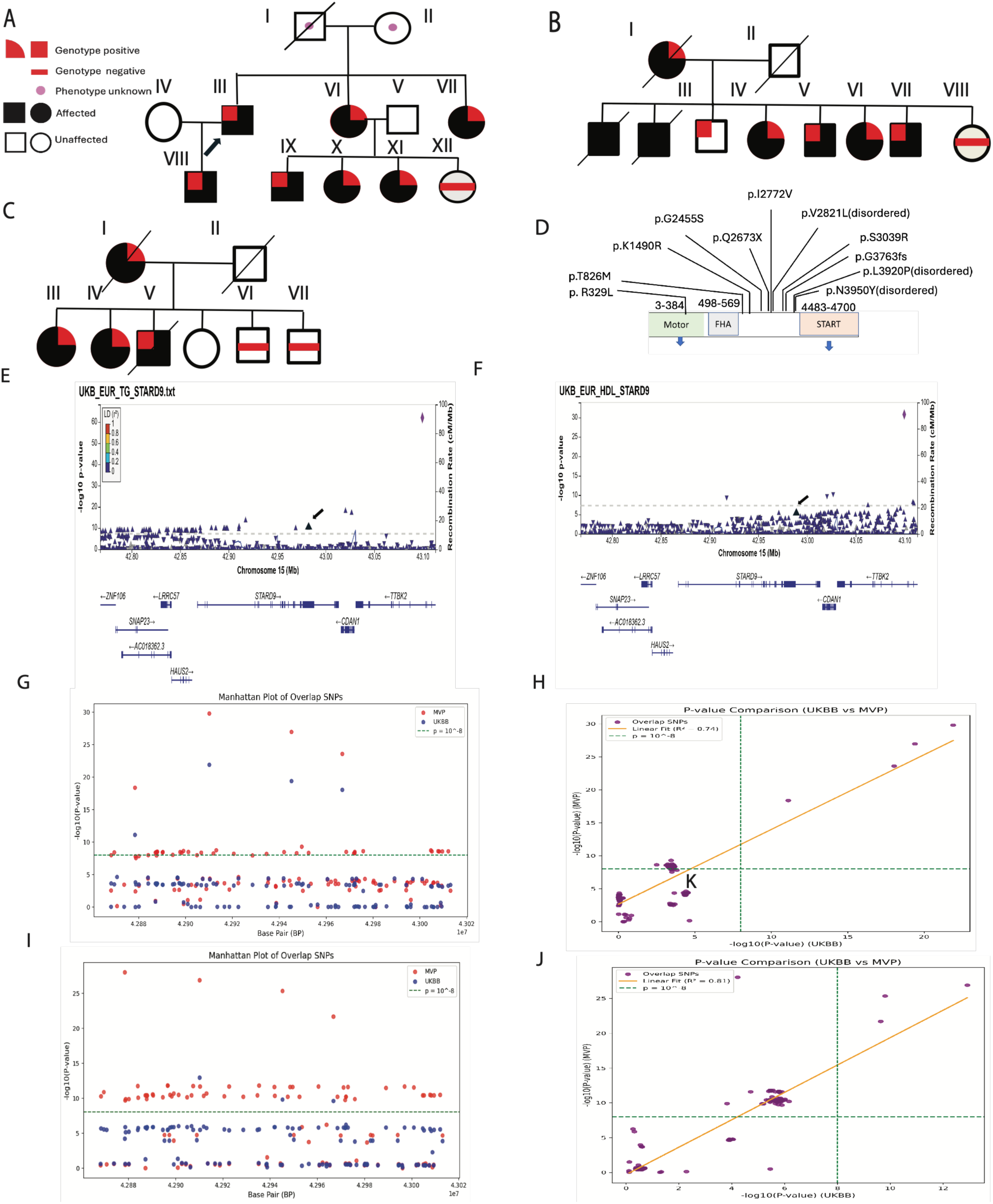
Identification and Genetic Association of *STARD9* Variants: Rare *STARD9* variants co-segregate with elevated LDL and atherosclerosis in multiplex families, and common variants are associated with elevate plasma triglyceride and reduced HDL cholesterol levels in population cohorts. **(A)** Pedigree of kindred 1 showing segregation of the p.V2821L variant, most mutation carriers are female and have elevated LDL. **(B-C)** Segregation of p.Lys1490Arg and p.Leu3920Pro variants in independent kindreds with premature CAD and elevated LDL. **(D)** Spectrum of rare *STARD9* variants identified in 69 unrelated index cases. **(E-F)** Manhattan plots from UK Biobank show genome-wide significant associations (P < 5 × 10⁻⁸) between common *STARD9* variants and plasma triglycerides (TG) and HDL cholesterol (HDL-C). The arrowheads denote the location of nonsynonymous GWAS variant p.I2772V in the UK Biobank database. **(G-J)** Cross-cohort colocalization demonstrates concordant association signals for TG and HDL-C between UK Biobank and the Million Veteran Program.

Unexpectedly, exome sequencing of nine additional multiplex kindreds with inherited hypercholesterolemia (LDL-C >180 mg/dL), normal Lp(a) (<50 mg/dL), and premature coronary artery disease (CAD), after excluding variants in known FH genes and filtering for allele frequency <0.001, identified two independent missense variants in STARD9 (p.Lys1490Arg and p.Leu3920Pro) affecting conserved residues, in two of these kindreds. Both variants segregated completely with premature CAD, defined as myocardial infarction or angiographically confirmed obstructive disease before age 55 in men or 65 in women, and were absent in unaffected relatives (Fig. 1B–C, Table 1). These variants are ultra-rare (allele frequencies 3×10⁻⁵ and 1.6×10⁻⁴, respectively).

In a separate cohort of 69 unrelated outlier index cases with extreme hyperlipidemia (LDL-C >200 mg/dL and/or triglycerides >200 mg/dL, Lp(a) <50 mg/dL), drawn from a larger cohort of 450 patients with early-onset CAD meeting the same definition of premature disease (myocardial infarction or angiographically confirmed obstructive disease before age 55 in men or 65 in women), the same filtering strategy identified eight additional rare STARD9 variants, including a nonsense mutation and a frameshift allele (G3763fs) (Fig. 1D; Table 2). Structural mapping showed that, apart from p.R329L in the motor domain, all identified variants fall outside the annotated motor and START domains (Fig. 1D). Variant enrichment was assessed using a two-sided Fisher’s exact test, comparing variant carrier frequencies in affected individuals versus the gnomAD v2.1.1 non-neuro, non-cardiac subset as the control population. This analysis revealed a significant excess of *STARD9* variant carriers among cases (P = 3.2 × 10⁻⁴), corresponding to an odds ratio of 4.6(95% CI, 2.1–9.8), supporting *STARD9* as a disease gene for familial dyslipidemia and premature atherosclerosis.

### Common Variant Association and Cross-Cohort Colocalization at the *STARD9* Locus

To complement rare-variant analyses, we evaluated genetic variation at the *STARD9* locus in two large population cohorts: the UK Biobank imputed genome-wide association study (GWAS) dataset ^15^ (UKB; Table 3) and the Million Veteran Program^16^ (MVP; Table 4).

In UKB, Variant data for the 20-kb genomic interval spanning *STARD9* was extracted and subjected to stringent quality control, including filtering for common variants (minor allele frequency [MAF] > 0.05) and applying standard UK Biobank array criteria, including variant call rate ≥95%, Hardy–Weinberg equilibrium P > 1 × 10⁻⁶, minor allele frequency filters appropriate for single-variant testing, and imputation quality thresholds (INFO ≥0.8). Analyses were restricted to individuals of White British ancestry to minimize population stratification, with adjustment for age, sex, and the first 10 principal components. Lipid phenotypes underwent harmonized quality control, including exclusion of outliers, biologically implausible values, and individuals with missing covariates. Single-variant association testing used linear mixed models to evaluate HDL-C (n = 404,121), LDL-C (n = 404,741), and triglycerides (n = 391,626). Common variants at the *STARD9* locus were significantly associated with plasma triglycerides (Fig. 1E) and HDL-C (Fig. 1F), surpassing genome-wide significance thresholds (P < 5 × 10⁻⁸) and false discovery rate–adjusted criteria (P < 0.05). Based on summary statistics from the Common Cardiovascular Disease Knowledge Portal, the P values are 4.37 × 10⁻¹⁴ for HDL and 6.53 × 10⁻⁸ for triglycerides, with HuGE scores of 20 for both (reflecting rare and common variant evidence), demonstrating a significant association between common variants and each trait.

Complementary ancestry-stratified analyses in MVP used imputed genotypes filtered for INFO ≥0.8 and cohort-specific genotype QC metrics. Phenotype harmonization and covariate adjustment mirrored UKB analyses. Cross-cohort colocalization was assessed by comparing −log₁₀(P) values for overlapping variants, revealing strong concordance for triglycerides (Fig. 1G–H) and HDL-C (Fig. 1I–J). Despite differences in SNP coverage and imputation structure, lipid-associated signals converged on shared candidate causal alleles at the *STARD9* locus with consistent effect direction across cohorts. Together, these findings support the presence of bona fide lipid-regulating variants at *STARD9*.

Several common variants demonstrated genome-wide significant associations and functioned as expression quantitative trait loci (eQTLs) for *STARD9*, including rs11070374 (Chr15:42559266, C:T; P = 3.25 × 10⁻⁸), rs10851411 (Chr15: 42567037, T:G; P = 4.23 × 10⁻⁸), rs11631193 (Chr15: 42575192, C:G; P = 4.25 × 10⁻⁸), and rs10454039 (Chr15: 42540336, C:T; P = 1.36 × 10⁻⁸). These variants were associated with reduced triglyceride levels and increased *STARD9* expression in different tissues, including the lung and tibial artery. Conversely, eQTLs such as rs10851410 (Chr15: 42552590, A:G; P = 5.30 × 10⁻⁸) and rs12324717 (Chr15: 42554355, C:A; P = 5.25 × 10⁻⁸) were associated with increased triglyceride levels and reduced *STARD9* expression. Notably, a rare protein-altering variant, rs202077402 (Chr15:42,689,892; p.Ile2772Val; arrows in Fig. 1E–F indicate its position in the UK Biobank whole-genome sequencing dataset), is strongly associated with elevated triglyceride levels (P = 1.44 × 10⁻¹²), triglyceride-to-HDL ratio (P = 2.38 × 10⁻⁸), and low HDL cholesterol (P = 1.41 × 10⁻⁷) across independent cohorts, based on summary statistics from the Common Cardiovascular Disease Knowledge Portal. ^17^. One individual with this variant was in our cohort with very similar lipid traits (table 2). Interestingly, this variant was recently reported to be associated with HDL (p = 5.5 × 10⁻¹⁴) and TG (p = 5.0 × 10⁻²⁵) in an exome-wide association study of blood lipids^18^. ENCODE annotation revealed that this variant lies within a predicted RXRβ binding site, a nuclear receptor central to lipid metabolism^14^. Integration of rare familial variants with convergent common-variant associations and cross-cohort colocalization establishes *STARD9* as a genetically validated regulator of lipid homeostasis and dyslipidemia risk.

### *STARD9* Mutation Causes Cholesterol Sequestration in Lysosomes

*STARD9* is expressed in nucleated peripheral blood cells. Isolation of peripheral monocytes from individuals carrying the p.V2821L *STARD9* variant revealed pronounced expression of STARD9 within CD68-positive cells, while total STARD9 protein levels were reduced in these primary monocytes compared with controls. (Fig. 2A). Staining for BODIPY^19^ demonstrated a significant increase in intracellular lipid content relative to wild-type (WT) controls (Fig. 2B). A substantial fraction of this lipid accumulation consisted of free cholesterol assayed by Filipin staining (Fig. 2C). Co-staining with LysoTracker showed that cholesterol predominantly colocalized with lysosomes, which appeared more abundant and enlarged in p.V2821L carriers compared with noncarriers (Fig. 2D).

**Figure 2:**
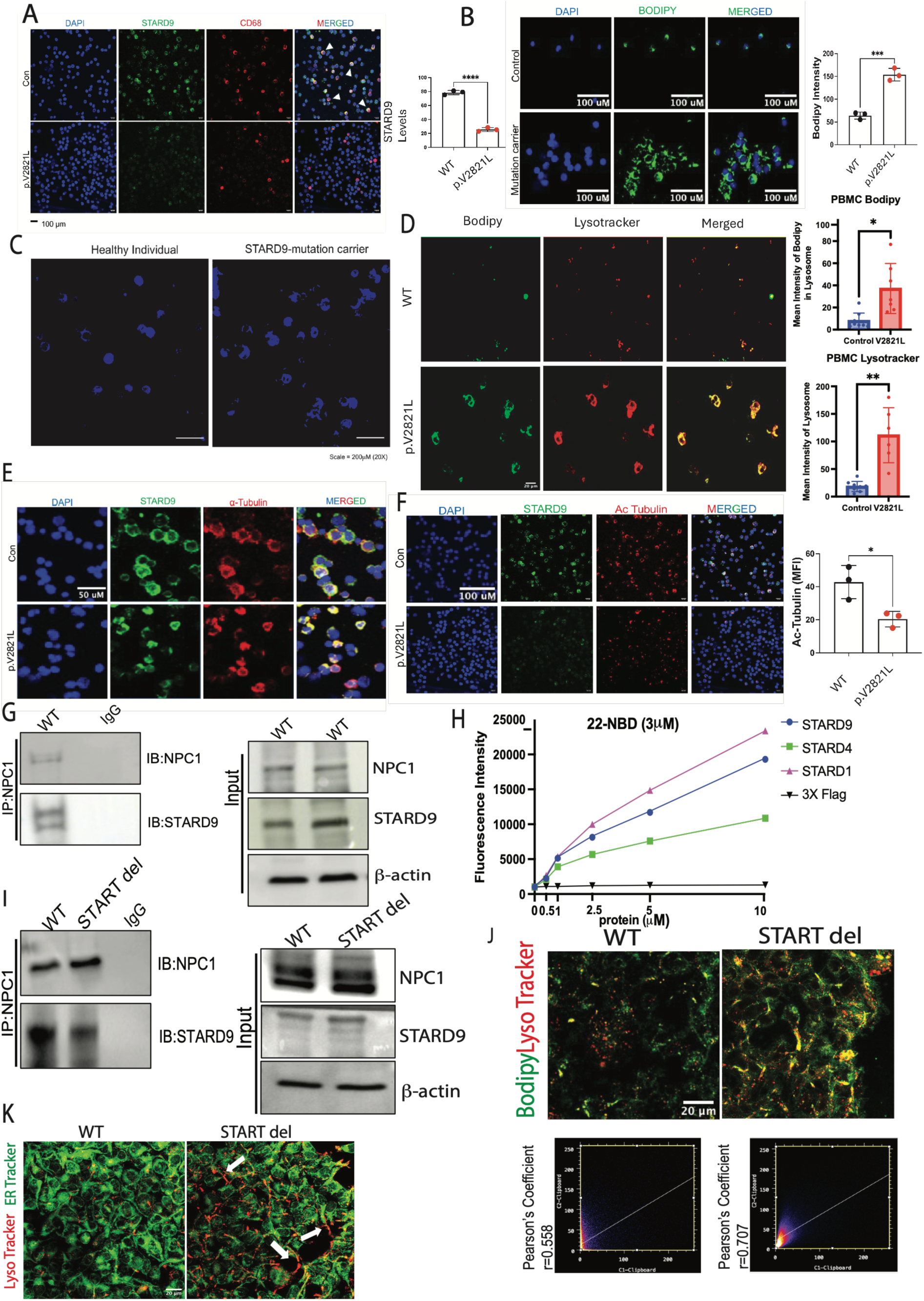
*STARD9* Mutation Causes Lysosomal Cholesterol Sequestration: The rare p.V2821L variant leads to lysosomal cholesterol accumulation and lysosomal defects in patient-derived cells. **(A)** Immunofluorescence shows STARD9 clustering in CD68+ monocytes from a p.V2821L carrier. **(B)** BODIPY staining reveals increased intracellular neutral lipid content in carrier cells (quantified, mean ± SEM, unpaired t-test). **(C)** Filipin staining shows accumulation of free cholesterol **(D)** Co-staining with LysoTracker demonstrates cholesterol sequestration within lysosomes. **(E)** α-tubulin staining shows preserved microtubule architecture. **(F)** Acetylated tubulin levels are significantly reduced in mutation carriers. (G) Co-immunoprecipitation confirms STARD9–NPC1 complex formation in THP-1 cells. (H) NBD-cholesterol binding assay of the isolated STARD4 and START domain of STARD9, STARD1 show concentration-dependent cholesterol binding. (I) CRISPR-mediated deletion of the START domain (STARD9-ΔSTART) reduces co-immunoprecipitation with NPC1. (J) Bodipy and LysoTracker co-staining shows that STARD9-ΔSTART cells accumulate cholesterol within lysosomes, comparable to p.V2821L knock-in cells. (K) LysoTracker and ER-Tracker co-staining shows that loss of the START domain shifts lysosomes from a pericentric to a peripheral, ER-associated distribution in HEK cells. *P ≤ 0.05, **P ≤ 0.01, ***P ≤ 0.001, **\*\*\*\***P ≤ 0.0001.

Because STARD9 is a kinesin family member implicated in microtubule-dependent trafficking, we assessed cytoskeletal integrity. α-Tubulin staining showed preserved microtubule architecture (Fig. 2E); however, acetylated tubulin levels were significantly reduced in mutation carriers (Fig. 2F), consistent with impaired microtubule stability and transport efficiency^20,21^.

### STARD9 binds cholesterol, forms a complex with NPC1 and regulates lysosomal location

STARD9 has previously been shown to colocalize with NPC1 and to regulate lysosomal movement within the cell¹⁰. To confirm and extend this interaction, we performed co-immunoprecipitation in THP-1 cells and confirmed that STARD9 forms a complex with NPC1 (Fig. 2G). Whether STARD9 itself binds cholesterol had not previously been established. Because full-length STARD9 is a large, technically difficult protein to express, we expressed its START domain in isolation and performed an NBD-cholesterol binding assay using increasing concentrations of the expressed domain; this demonstrated concentration-dependent binding of the STARD9 START domain to cholesterol (Fig. 2H). We next used CRISPR-mediated deletion to generate STARD9 lacking the START domain (STARD9-ΔSTART); this construct modestly reduced co-immunoprecipitation with NPC1 (Fig. 2I), indicating that the loss of START domain does not inhibit but may reduce STARD9–NPC1 complex formation. Bodipy and LysoTracker co-staining showed that STARD9-ΔSTART cells accumulate cholesterol within lysosomes to a degree comparable to p.V2821L knock-in cells, indicating that loss of the START domain alone is sufficient to reproduce the lysosomal cholesterol sequestration phenotype (Fig. 2J). Strikingly, using LysoTracker and ER-Tracker co-staining in HEK cells, we noted that loss of the START domain redistributed lysosomes to the cell periphery along the ER, in contrast to the pericentric localization observed in cells expressing full-length STARD9 (Fig. 2K).

### STARD9 loss impairs Cholesterol Trafficking from Lysosomes to the ER and activates mTOR–NF-κB– driven Inflammatory Reprogramming

To directly test the role of STARD9 in intracellular cholesterol trafficking, we silenced *STARD9* in THP-1 cells using a *STARD9*-specific shRNA, achieving near-complete knockdown (suppl. Fig. 1A). Mirroring findings in patient-derived cells, *STARD9* knockdown following 6 hours of exposure to 2μM BODIPY-cholesterol containing medium resulted in robust lipid accumulation (suppl. Fig. 1B), with the majority of accumulated lipid identified by Filipin staining as free cholesterol ^22^ (suppl. Fig. 1C). In contrast, esterified cholesterol levels were reduced, as measured by Oil Red O staining (suppl. Fig. 1D), indicating impaired cholesterol delivery to sites of esterification.

To examine cholesterol trafficking dynamics, WT and *STARD9*-knockdown PMA-differentiated THP-1 cells were treated with OxLDL (100 µg/mL) and analyzed over time. In WT cells, lysosomal cholesterol levels peaked at approximately 20 minutes and then gradually declined, consistent with efficient cholesterol egress (Suppl. Figs. 1E–J). In *STARD9*-knockdown cells, BODIPY signal started to increase after 30 minutes, indicating that *STARD9* deficiency does not significantly alter initial cholesterol uptake kinetics in these cells. However, in contrast to WT cells, *STARD9*-deficient cells exhibited persistent lysosomal cholesterol retention at later time points. These findings demonstrate that STARD9 is required for efficient late endosomal/lysosomal rather than early cholesterol trafficking.

Consistent with this trafficking defect, time-course analysis of ER cholesterol demonstrated that cholesterol failed to efficiently reach the ER in *STARD9*-knockdown cells following OxLDL treatment (suppl. Fig. 2A). Impaired ER delivery would be expected to disrupt sterol sensing and dysregulate sterol-responsive pathways. Accordingly, nuclear SREBP2 levels remained persistently elevated in *STARD9*-knockdown cells despite BODIPY-cholesterol treatment, whereas nuclear localization declined after 2 hours in most wild-type THP-1 cells (suppl. Fig. 2B).

Collectively, these alterations resulted in upregulation of CD36 and canonical SREBP2 target genes, including *LDLR* (suppl. Fig. 2C). CD36 contributes to innate immune activation through Toll-like receptor signaling^23^. Consistent with increased CD36 expression, *STARD9* knockdown was associated with elevated pro-inflammatory cytokines, including IL-1β and TNF-α (suppl. Fig. 2C).

### Single-cell RNA-seq identifies cholesterol metabolism, microtubule cytoskeleton, and lysosomal stress as dominant altered pathways

Since *STARD9* is expressed in peripheral blood cells, we performed an unbiased interrogation of disease-relevant cellular programs using single-cell RNA sequencing (scRNA-seq) of peripheral blood mononuclear cells (PBMCs) from two p.V2821L variant carriers (father and son) and 20 age- and sex-matched affected non-carriers. Following quality control, cells were projected onto the Azimuth reference atlas, and supervised label transfer was used to assign cell identities. The resulting UMAP embedding (Fig. 3A) displays 30 predicted canonical immune populations, including classical (CD14⁺CD16⁻), intermediate (CD14⁺⁺CD16⁺), and non-classical (CD14⁺CD16⁺⁺) monocytes, CD4⁺ and CD8⁺ T cells, B cells, and NK cells. Comparative compositional analysis revealed selective immune remodeling in STARD9 carriers, marked by expansion of CD16⁺ NK cells, and mucosal-associated invariant T (MAIT) cells, with a concomitant reduction in classical monocytes (suppl. Fig. 3A). This shift suggests altered immune homeostasis consistent with chronic metabolic stress.

**Figure 3:**
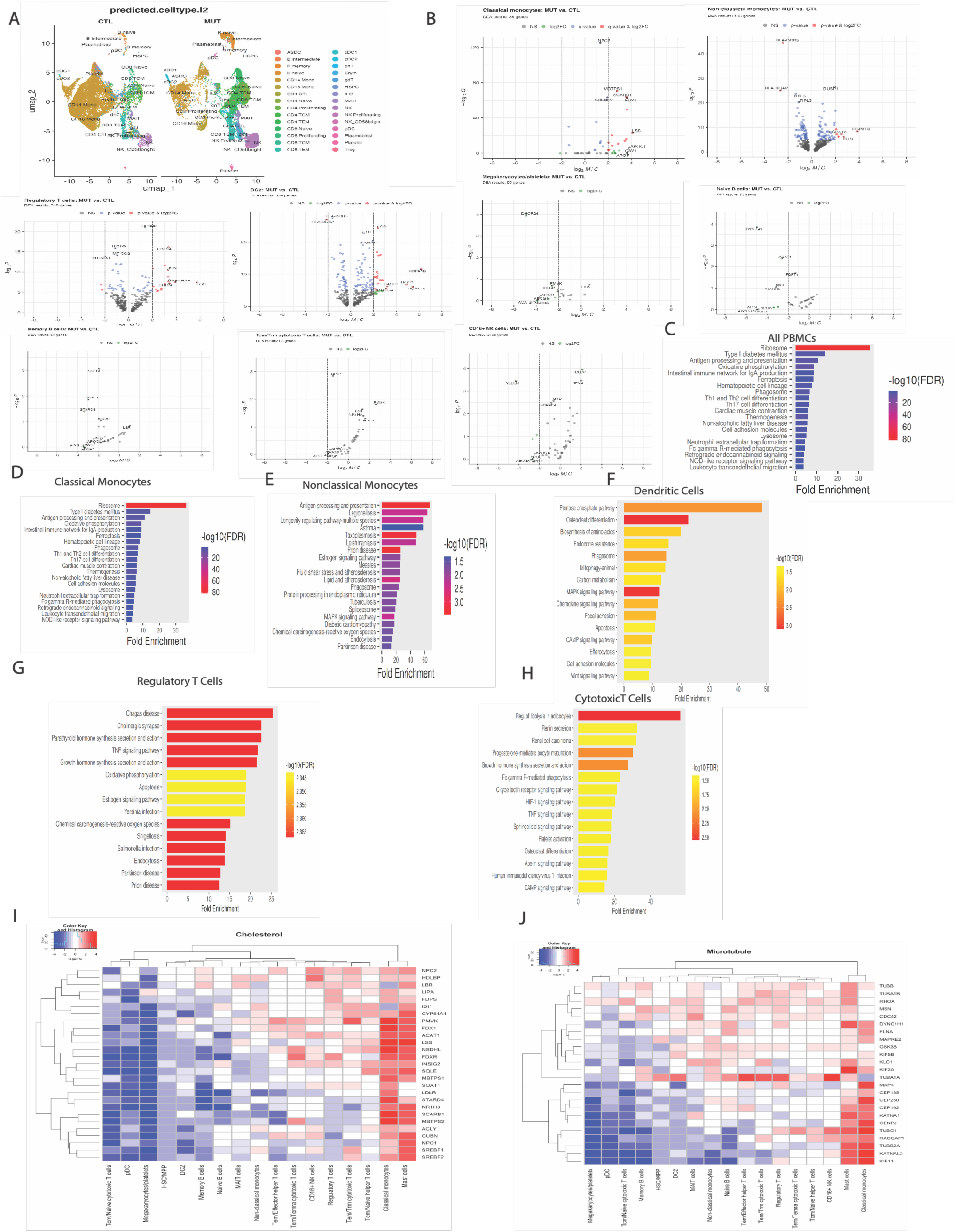
Single-cell RNA sequencing of peripheral blood mononuclear cells from p.V2821L carriers identifies coordinated dysregulation of cholesterol metabolism, lysosomal programs, and cytoskeletal transport pathways. (A) UMAP with cell types predicted by the Azimuth method in Seurat (B) Volcano plots illustrating differential gene expression across distinct blood cell populations. (C–H) KEGG pathway enrichment analysis demonstrating upregulation of ribosomal, lysosomal, inflammatory, and mTOR-related pathways in carrier versus non-carrier blood cells. (I) Heatmap summarizing coordinated dysregulation of cholesterol metabolism genes across immune subsets. (J) Heatmap showing altered expression of microtubule and vesicular transport genes linking cytoskeletal remodeling to lysosome positioning and sterol trafficking.

### Cell type–specific transcriptional consequences of STARD9 dysfunction

Transcriptional perturbations were most particularly pronounced in classical monocytes, key drivers of atherogenesis (Table 5), which exhibited coordinated upregulation of sterol regulatory genes, including *SREBF2, FDX1, SCARB1, LSS (Lanosterol synthase),* and *MBTPS1(Membrane-bound transcription factor site-1 protease),* encoding S1P (Site-1 Protease) (Fig. 3B). This signature indicates activation of a sterol-deprivation response despite intracellular cholesterol accumulation, consistent with defective lysosome-to-ER cholesterol trafficking. Functionally, ER cholesterol depletion promotes S1P-dependent SREBP2 processing^24^, induction of de novo sterol biosynthesis (*LSS*)^25^, altered HDL flux (*SCARB1*)^26^, and compensatory mitochondrial sterol metabolism (*FDX1*)^27^, collectively reflecting a maladaptive attempt to restore sterol homeostasis.

Parallel lipid-handling defects were observed across immune subsets (Suppl Fig. 3B). Megakaryocytes displayed suppression of cholesterol synthesis and transport genes (*HDLBP, ACAT1, ABCG1,* and *LBR*) (Fig. 3B), while naïve B cells showed reduced expression of cholesterol esterification and efflux components (*ACAT1, APOA1,* and *APOE*). Cytotoxic T cells demonstrated coordinated downregulation of mevalonate pathway and cholesterol trafficking genes (*IDI1, NPC2, ABCG8,* and *APOC1*) (Fig. 3B), indicating reduced sterol flux. In regulatory T cells, upregulation of lysosomal stress-associated genes (*GABARAP, FOS, JUN, H3F3A,* and *SRGN*) alongside decreased mitochondrial *MT-CO2* suggests a metabolic shift toward stress-adaptive lysosomal programs. Across all populations, cholesterol metabolism emerged as a unifying axis of dysregulation (HeatMap Fig. 3I), highlighted by altered expression of genes governing esterification (*ACAT1*), HDL handling (*HDLBP, SCARB1*), and biosynthesis (*NSDHL, PMVK*).

### Coordinated and paradoxical dysregulation of lysosomal programs, and autophagy

Pathway enrichment analyses using KEGG (Fig. 3C–H) identified recurrent signatures involving ribosomes, lysosomes, neutrophil extracellular trap formation, phagosomes, and efferocytosis. Reactome analysis of classical and nonclassical monocytes revealed translation initiation, translation elongation, and cellular responses to starvation among the most significantly altered pathways. Together, these processes intersect with mTOR signaling, a central regulator of anabolic metabolism, lysosomal function, and autophagy. In cytotoxic T cells and dendritic cells (DC2), Reactome analysis identified Innate and Adaptive Immune System pathways, interferon-γ signaling, and interleukin-12 signaling among the most significantly altered pathways (Table 5). Consistent with altered mTOR pathway activity, monocytes from variant carriers exhibited differential expression of multiple ribosomal genes, including several RPS and RPL family members, notably *RPS6*, a canonical downstream target of mTOR signaling. Paradoxically, broad activation of lysosomal and hydrolase genes (CTSB, CTSD, CTSS, CTSZ, ASAH1, and *PSAP*) and lysosomal membrane-associated components (*CD68, LAPTM5, ATP6V0B*) was observed across multiple cell types. Autophagy/CASM-related genes (*ATG7, GABARAP*) and cholesterol trafficking factors (*NPC2*) were also differentially expressed (Bonferroni-adjusted *P* < 0.001; Table 5). These changes may represent either compensatory responses to cellular stress or contributors to disease pathogenesis.

Interrogation of transcriptional programs governing lysosomal biogenesis and autophagy revealed a mechanistically informative divergence. TFEB/TFE3-regulated lysosomal and degradative genes, including cathepsins, *ATG7, ATP6V1G,* and *ATP6V0B*, were upregulated, whereas several ribosomal genes, including *RPS29*, were suppressed across multiple cell types.

### Microtubule cytoskeleton and vesicular transport pathways are disrupted in *STARD9* variant carriers

Beyond metabolic rewiring, scRNA-seq analysis revealed dysregulation of genes governing microtubule-based trafficking (SFig. 3C), including *KIF5B, KLC1,* and *TUBA1A* (Heatmap Fig. 3J). Because lysosome positioning, sterol transport, and endolysosomal maturation depend on intact kinesin and microtubule dynamics, these changes provide a structural framework linking STARD9 dysfunction to impaired intracellular cholesterol routing.

### CRISPR–Cas9 knock-in of STARD9 variants reveals shared and divergent effects on cholesterol trafficking and LDL handling

To determine the functional consequences of STARD9 coding variation, we generated heterozygous CRISPR/Cas9 knock-in HEK293T cell lines harboring either the autosomal dominant p.V2821L variant, or the rare GWAS-associated variant rs202077402 (p.I2772V). We also generated a heterozygous knockout (KO) line by deleting portions of exons 5 and 7 together with the entirety of exon 6, resulting in a frameshift mutation. These three isogenic lines, together with wild-type controls, allowed us to systematically dissect the shared and variant-specific consequences of STARD9 dysfunction on intracellular cholesterol trafficking and receptor-mediated endocytosis.

Before characterizing the cellular phenotypes of these lines, we first sought to understand the regulatory context of the p.V2821L variant. Interrogation of ENCODE ChIP-seq datasets revealed that this locus resides within a predicted binding site for HNF1α, which plays established roles in hepatic cholesterol metabolism^28^. Electrophoretic mobility shift assays (EMSA) demonstrated markedly reduced HNF1α binding to the p.V2821L allele compared with the wild-type sequence, indicating disruption of a canonical HNF1α recognition motif (Fig. 4A). HNF1α overexpression in HEK293T cells, which do not express this transcription factor, did not alter STARD9 transcript levels in wild-type cells but significantly increased STARD9 expression in p.V2821L mutant cells (Fig. 4A). These findings are consistent with a model in which HNF1α binding at the wild-type locus imposes transcriptional repression, a context-dependent transcriptional mechanism previously described for this factor^29^. Collectively, these data suggest that p.V2821L shifts the regulatory balance at the STARD9 locus from a repressed to a permissive state.

**Figure 4:**
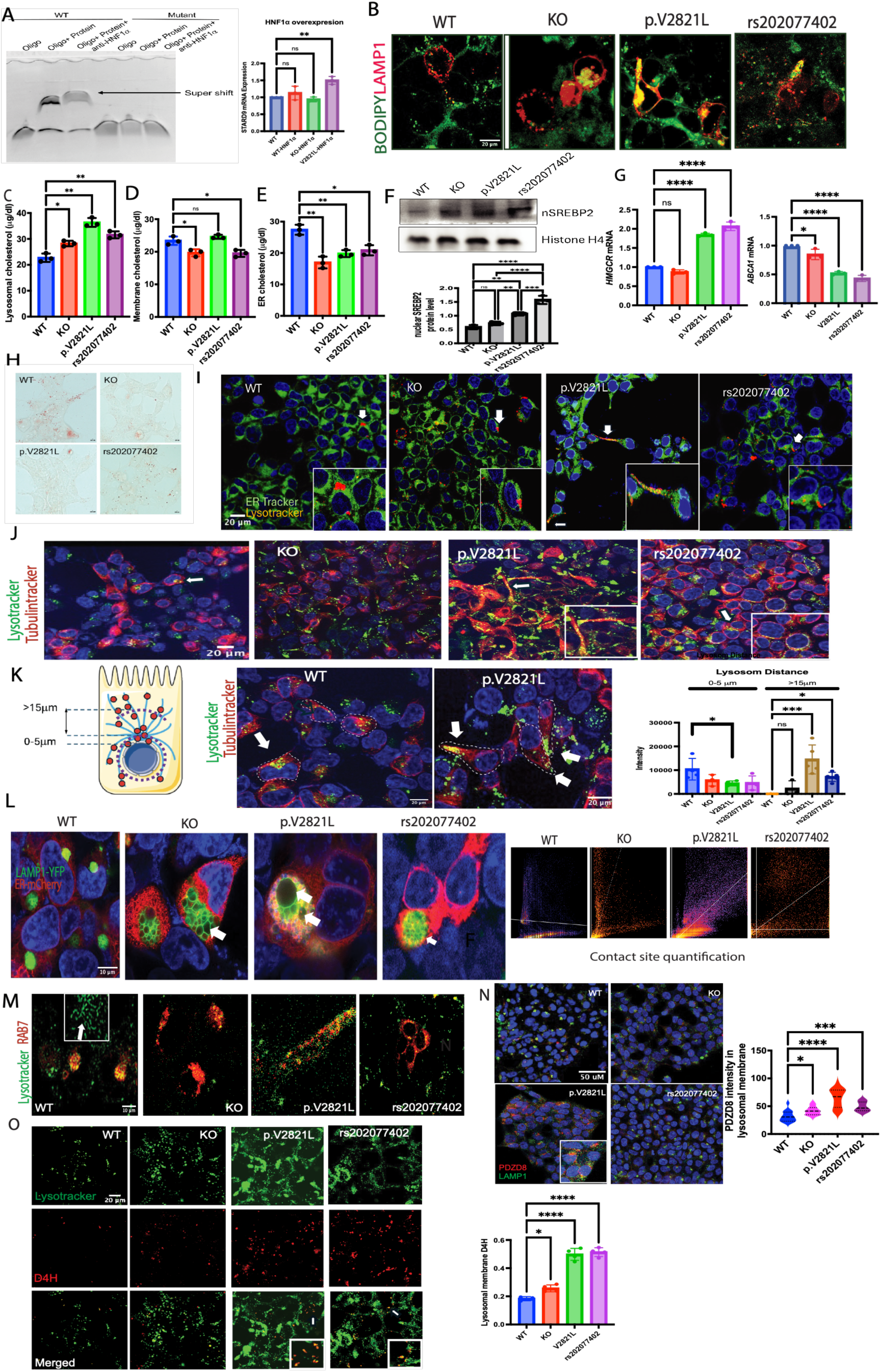
*STARD9* variants converge on lysosomal cholesterol sequestration but diverge in lysosomal positioning and ER–lysosome contact site expansion. (A) Electrophoretic mobility shift assay (EMSA) demonstrating reduced HNF1α binding to the p.V2821L allele relative to wild type, accompanied by increased *STARD9* transcript levels following HNF1α overexpression specifically in p.V2821L knock-in cells. (B) Persistent lysosomal cholesterol accumulation in all *STARD9* mutant lines following BODIPY-cholesterol exposure. (C) Lysosomal cholesterol content measured by LysoIP followed by cholesterol extraction and Folch quantification. (D) Plasma membrane cholesterol content following methyl-β-cyclodextrin/lovastatin treatment in p.V2821L KI, p.I2772V KI, and KO cell lines. (E) ER cholesterol is reduced in all three mutant lines relative to controls following oxysterol treatment. (F) Sustained nuclear localization of SREBP2 in mutant lines after 12 hours of cholesterol exposure, indicating failure of normal sterol-dependent feedback re-sequestration. (G) Upregulation of HMGCR expression consistent with chronic SREBP2 activation and downregulation of ABCA1, a key cholesterol efflux mediator suppressed by SREBP2. (H) Oil red O staining of all 3 mutant cells lines shows reduced esterified cholesterol (I) Peripheral redistribution of lysosomes in p.V2821L KI cells, with increased colocalization with ER-Tracker and alignment along cellular extensions. (J) Enhanced colocalization of dispersed lysosomes with Tubulin-Tracker in p.V2821L KI cells. (K) Quantitative analysis confirming a significant shift toward lysosomes located more than 15 μm from the nucleus in p.V2821L KI cells. (L) Expansion of ER–lysosome contact sites and vacuolization and (M) Loss of lysosomal membrane tubulation in all mutant lines. (N) Increased PDZD8/LYVAC colocalization with lysosomes in both missense lines, most prominently in p.V2821L KI cells. (O) Increased accessible cholesterol on the lysosomal limiting membrane detected by the D4H biosensor in mutant cells. Data are presented as mean ± SEM, with each point indicating an independent biological replicate. For multi-group comparisons, one-way ANOVA with Holm–Šidák correction was used for normally distributed data; non-parametric comparisons used the Mann–Whitney U test. *P ≤ 0.05, **P ≤ 0.01, ***P ≤ 0.001, ****P ≤ 0.0001.

We next characterized the intracellular cholesterol trafficking phenotype of all three mutant lines. Cells were exposed to BODIPY-cholesterol, after 4h, Bodipy containing media was removed and the cells kept for 24h. All the mutant lines exhibited persistent lysosomal cholesterol accumulation compared to controls, as demonstrated by increased BODIPY-cholesterol/LAMP1 colocalization (Fig. 4B). To directly quantify this sequestration, we performed LysoIP followed by cholesterol extraction and quantification by the Folch method, which confirmed significantly elevated lysosomal cholesterol content in all three mutant lines relative to controls (Fig. 4C).

Despite this shared defect, the two knock-in lines diverged in how lysosomal cholesterol sequestration affected downstream compartments. To assess cholesterol distribution across cellular compartments, we isolated plasma membrane and ER fractions from cells pretreated with methyl-β-cyclodextrin (1 mM, 1 hour) followed by lovastatin (5 μg/ml) to standardize membrane cholesterol conditions, and quantified cholesterol by a modified Folch procedure. Plasma membrane cholesterol was normal in p.V2821L KI cells but significantly reduced in KO and p.I2772V KI cells (Fig. 4D). In contrast, ER cholesterol was reduced in all three mutant lines relative to controls following oxysterol treatment (Fig. 4E). These findings indicate that while all mutant lines share impaired lysosome-to-ER cholesterol delivery, the p.I2772V variant and KO, additionally impairs cholesterol transport to the plasma membrane, a distinction that, as we describe below, has important consequences for downstream signaling and endocytic function.

Reduced ER cholesterol content in all mutant lines was associated with a failure of normal sterol-dependent feedback regulation. Both knock-in lines exhibited sustained nuclear localization of SREBP2 following 12 hours of cholesterol exposure, in contrast to the efficient cytoplasmic re-sequestration observed in controls (Fig. 4F). Consistent with chronic SREBP2 activation, expression of its canonical target *HMGCR* was upregulated (Fig. 4G), while *ABCA1*, a key mediator of cholesterol efflux was correspondingly downregulated. Accordingly, there was reduced esterified cholesterol assessed by oil red-O in all 3 mutant lines (Fig. 4H). This pattern of SREBP2-driven transcriptional dysregulation mirrors that observed in NPC1, a prototypical lysosomal cholesterol storage disorder, and reflects the inability of sequestered lysosomal cholesterol to reach the ER sensing machinery that normally restrains SREBP2 processing^30^.

### Differential effects of p.V2821L vs. p.I2772V variants on lysosomal positioning

*STARD9* variants caused pronounced differences in lysosomal positioning, a key determinant of organelle function, since lysosomal cytoplasmic trafficking along the cytoskeleton governs their ability to locate and fuse with appropriate compartments for substrate processing^31^. In wild-type controls, KO, and p.I2772V KI cells, lysosomes were predominantly perinuclear, consistent with their canonical role in autophagy, where central positioning facilitates autophagosome–lysosome fusion and efficient cargo degradation³¹, while in p.V2821L cells they were shifted toward the cell periphery (Fig. 4I), colocalizing with microtubules (Fig. 4J). confirmed by quantitative analysis showing a significant distance from the nucleus (Fig. 4K). This peripheral, ER-aligned lysosomal distribution closely resembled the lysosomal mislocalization observed following CRISPR-mediated deletion of the START domain (Fig. 2K), suggesting that the p.V2821L variant phenocopies loss of START domain function with respect to lysosomal positioning. This lysosomal redistribution was accompanied by expansion of ER–lysosome contact sites and lysosomal vacuolation, most significantly in p.V2821L KI cells (Fig. 4L). Contact sites were quantified by measuring spatial colocalization between LAMP1-positive lysosomes and ER signal, with overlapping voxels serving as a proxy for physical membrane apposition. This expansion of ER–lysosome contacts likely represent a lysosomal stress, analogous to the signaling hubs formed at these contacts in NPC1 disease for mTORC1-mediated cholesterol sensing^32^. In addition, compared to lysosomes in wildtype cells, there was loss of tubulation (arrow) in all mutant lines in response to treatment with cholesterol for 2h after 1 h starvation, required for proper lysosomal movement on microtubules ^8,31^(Fig. 4M).

Expansion of ER–lysosome membrane apposition has been shown to be associated with PDZD8/LYVAC recruitment and lipid transfer from the ER to lysosomes, leading to lysosomal vacuolation^33^. Accordingly, there were increased levels of PDZD8/LYVAC colocalizing with lysosomes in both missense lines and primarily in p.V2821L KI cells (Fig. 4N). Accordingly, there was increased accessible cholesterol on the lysosomal limiting membrane, as detected by the D4H biosensor, a genetically encoded sensor derived from the fourth domain of Perfringolysin O toxin that specifically reports accessible cholesterol on the cytosolic leaflet of cellular membranes^34^ (Fig. 4O).

### Activation of mTORC1 pathway by accessible cholesterol, and paradoxical activation of TFEB, associated with increased lysosomal biogenesis and normal cholesterol efflux caused by p.V2821L variant

Limiting membrane cholesterol accumulation drives constitutive mTORC1 activation via the SLC38A9–Rag GTPase–Ragulator signaling complex on the lysosomal surface^32^. Consistent with this, all mutant lines exhibited elevated phospho-mTOR and phospho-S6 (Fig. 5A), most apparent in p.V2821L KI cells. The peripheral lysosomal dispersion observed in p.V2821L KI cells is consistent with the mTORC1 hyperactivation ^35^, and closely mirrors findings reported in NPC1 disease, where lysosomal cholesterol sequestration similarly drives mTORC1 hyperactivation^32^.

**Figure 5:**
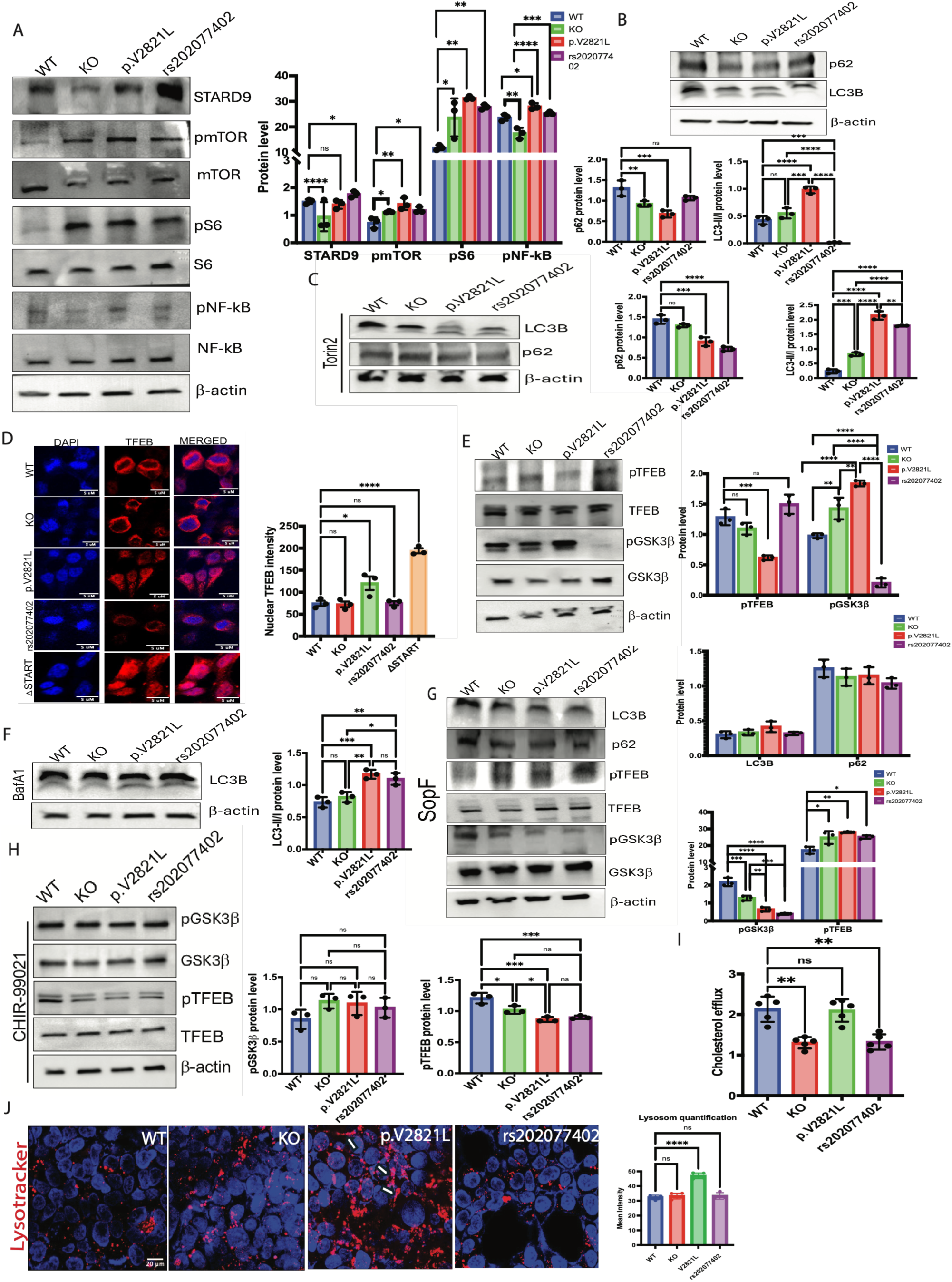
Loss of *STARD9* function drives mTORC1 hyperactivation and engages CASM to paradoxically activate TFEB and enhance cholesterol efflux. (A) Elevated phospho-mTOR (Ser2448) and phospho-S6 (Ser235/236) in all STARD9 mutant lines, most pronounced in p.V2821L KI cells, confirming mTORC1 hyperactivation downstream of lysosomal cholesterol accumulation. (B) Paradoxical elevation of LC3B-II in p.V2821L KI cells despite mTORC1 activation, pointing toward engagement of noncanonical ATG8 lipidation. (C) Torin treatment further increases LC3B-II levels in all three mutant lines, demonstrating that basal LC3B lipidation is partially suppressed by mTOR. (D) Immunofluorescence showing increased nuclear localization of TFEB in p.V2821L KI cells, reproduced in STARD9-ΔSTART cells. (E) Western blot confirming increased nuclear TFEB and reduced TFEB phosphorylation in p.V2821L KI cells, associated with reduced GSK3β activation. (F) Bafilomycin A1 treatment further increases LC3B lipidation in both KI lines (G) SopF-mediated CASM inhibition markedly reduces LC3B-II accumulation, restores TFEB phosphorylation, and normalizes GSK3β signaling specifically in p.V2821L KI cells. (H) CHIR-99021–mediated GSK3β inhibition reduces TFEB phosphorylation across all genotypes. (I) Reduced HDL-mediated cholesterol efflux in p.I2772V KI cells. (J) Expanded lysosomal number in p.V2821L KI cells. Data are presented as mean ± SEM. Each dot represents an independent biological replicate. For comparisons among multiple groups, one-way ANOVA with Holm–Šidák post hoc test was used when data were normally distributed; otherwise, the Mann–Whitney U test was used for non-normal data. *P ≤ 0.05, **P ≤ 0.01, ***P ≤ 0.001, **\*\*\*\***P ≤ 0.0001.

The shared mTORC1 hyperactivation across all mutant lines, combined with the peripheral lysosomal dispersion unique to p.V2821L cells, raised the question of whether these variants engage distinct lysosomal stress response programs downstream of a common mTOR signal. Canonical autophagy is initiated by mTORC1 inhibition and requires perinuclear lysosomal positioning for autophagosome–lysosome fusion. The peripheral lysosomal dispersion in p.V2821L cells would therefore be expected to impair autophagic flux, and mTORC1 hyperactivation in these cells predicts suppression of canonical LC3B lipidation. Paradoxically, LC3B-II levels were elevated and p62 levels were decreased in p.V2821L KI cells (Fig. 5B), which in the context of persistent mTORC1 activity, pointed toward engagement of CASM (conjugation of ATG8 proteins to single membranes), a pathway in which ATG8-family proteins are conjugated via the ATG12–ATG5– ATG16L1 complex to single-membrane compartments, as opposed to the double-membrane autophagosomes associated with canonical autophagy; it is a pathway triggered by lysosomal membrane damage, altered lipid composition, or V-ATPase dysfunction^36^. To test this, we treated cells with the mTOR inhibitor Torin (HY-13002, MedChemExpress, 100 nM, 2 hours), which further increased LC3B-II levels and decreased p62 levels across all three mutant lines (Fig. 5C), suggesting that mTOR partially suppressed basal LC3B lipidation. We then examined activation of TFEB, a lysosomal biogenesis transcription factor phosphorylated and inactivated by mTOR as well as by kinases like ERK^37^ and GSK3β^38^. Surprisingly, there was increased nuclear localization of TFEB in p.V2821L KI cells, as assessed by IF (Fig. 5D), a finding reproduced in STARD9-ΔSTART cells (Fig. 5D), and reduced phosphorylation assessed by western blot (Fig. 5E), findings consistent with scRNAseq in p.V2821L carriers. Accordingly, these changes were associated with increased GSK3β phosphorylation and inactivation. We then treated cells with bafilomycin A1 (BafA1; HY-100558, MedChemExpress, 100 nM, 2 hours), a V-ATPase inhibitor commonly used to assess autophagosome turnover^39^, which further increased LC3B lipidation in both KI lines (Fig. 5F). This BafA1-induced increase in LC3B-II is consistent with ongoing ATG8 lipidation, which in the presence of sustained mTORC1 activity raised the possibility that it reflects CASM rather than canonical autophagic flux, a question we addressed directly in subsequent experiments (Fig. 5G).To explore CASM involvement, cells were transfected with pEGFP-C1-SopF, a bacterial effector that selectively inhibits CASM by disrupting V-ATPase–ATG16L1 coupling without affecting canonical autophagosome formation^40^. SopF treatment markedly reduced LC3B-II accumulation and restored TFEB phosphorylation specifically in p.V2821L KI cells (Fig. 5G), demonstrating that both the noncanonical LC3B lipidation and the anomalous TFEB activation in these cells are CASM-dependent.

The mechanistic link between CASM and TFEB activation in p.V2821L cells is provided by previous studies, which have shown that GABARAP:CASM-driven lipidation of GABARAP onto the lysosomal limiting membrane sequesters the FLCN–FNIP1/2 complex away from the Rag GTPase machinery, preventing FLCN-dependent RagC/D GAP activity, thereby relieving mTORC1-dependent TFEB phosphorylation and enabling nuclear translocation despite sustained bulk mTORC1 activity^41,42^. Additionally, the elevated inhibitory phosphorylation of GSK3β observed in p.V2821L KI cells was also reduced following SopF treatment (Fig. 5G), indicating that CASM-dependent lysosomal signaling coordinately suppresses both mTORC1-dependent and GSK3β-dependent TFEB phosphorylation to achieve full TFEB nuclear accumulation. To confirm GSK3β as a causal regulator of TFEB phosphorylation in this system, pharmacologic inhibition of GSK3β with CHIR-99021 (HY-10182, MedChemExpress) reduced TFEB phosphorylation across all cell lines (Fig. 5H), establishing a direct functional role for this kinase. Increased TFEB activation has been associated with increased cholesterol efflux and lysosomal biogenesis^37,43,44^. Accordingly, there was normal cholesterol efflux (Fig. 5I) and increased lysosomal number (Fig. 5J) in p.V2821L KI cells compared to wildtype and p.I2772V KI cells. Together, these data reveal that the p.V2821L variant drives a CASM-dependent lysosomal stress response that paradoxically co-opts mTORC1 signaling to activate TFEB, culminating in enhanced lysosomal biogenesis and cholesterol efflux, potentially underlying the elevated HDL phenotype observed in this carrier.

### Variants produce opposing effects on LDLR surface expression and clathrin-mediated endocytosis

The two STARD9 variants also produced strikingly divergent effects on receptor-mediated endocytosis. Live-cell imaging with fluorescent transferrin, a readout of clathrin-mediated endocytosis, revealed that p.V2821L KI cells had significantly reduced transferrin uptake at 20 minutes, whereas p.I2772V KI cells exhibited increased transferrin uptake relative to controls (Fig. 6A). To confirm that this increase reflected a gene dosage-dependent effect rather than a variant-specific gain of function, we assessed transferrin uptake following STARD9 knockdown by shRNA, which reduced mRNA expression by >75%. Knockdown similarly increased transferrin uptake in HEK293T cells (Fig. 6B).

**Figure 6:**
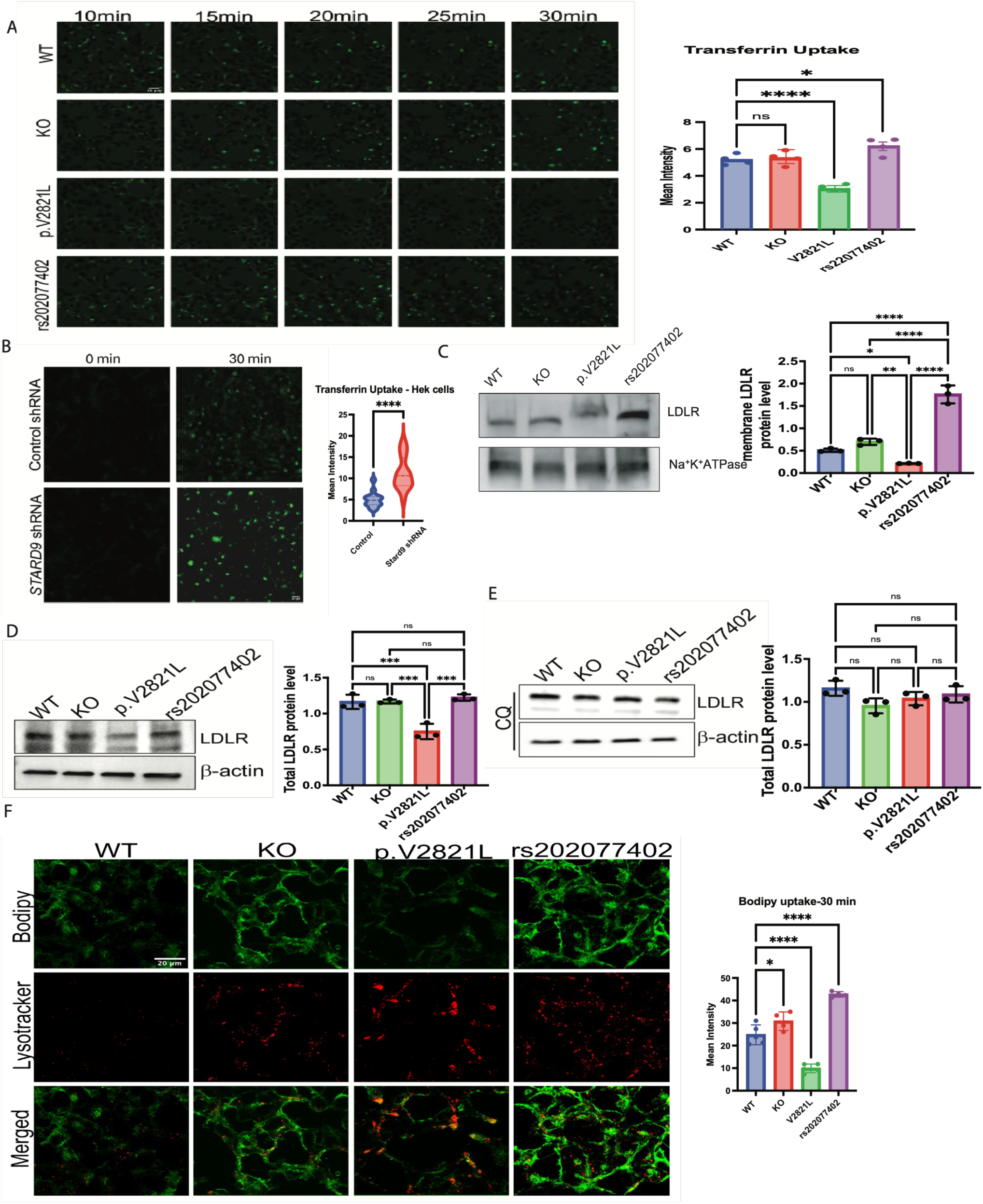
STARD9 variants produce opposing effects on clathrin-mediated endocytosis, LDLR abundance, and LDL uptake. (A) Live-cell imaging of fluorescent transferrin uptake showing significantly reduced clathrin-mediated endocytosis in p.V2821L KI cells and increased uptake in p.I2772V KI cells relative to controls at 20 minutes. (B) Increase transferrin uptake in HEK293T cells upon shRNA-mediated STARD9 knockdown (C) Western blotting of plasma membrane LDLR surface in mutant cell lines. (D) Total LDLR protein is dramatically reduced in p.V2821L KI cells. (E) Chloroquine treatment restores total LDLR to wild-type levels in p.V2821L KI cells. (F) BODIPY-LDL uptake is increased in p.I2772V KI and heterozygous KO cells and significantly reduced in p.V2821L KI cells. Data are presented as mean ± SEM. Each dot represents an independent biological replicate. For comparisons among multiple groups, one-way ANOVA with Holm– Šidák post hoc test was used when data were normally distributed; otherwise, the Mann–Whitney U test was used for non-normal data. *P ≤ 0.05, **P ≤ 0.01, ***P ≤ 0.001, ****P ≤ 0.0001.

These divergent endocytic phenotypes were accompanied by differences in LDLR surface and total expression. Western blotting of plasma membrane fractions, isolated after methyl-β-cyclodextrin and lovastatin treatment as described, revealed significantly increased LDLR protein in p.I2772V KI cells and reduced in p.V2821L KI cells compared with controls (Fig. 6C). Total LDLR was also reduced in p.V2821L KI cells (Fig. 6D), consistent with the carrier’s clinical trait of hypercholesterolemia but despite increased nuclear localization of SREBP2. The apparent increase in RAB7-positive intermediate multivesicular bodies in p.V2821L KI cells (Fig. 4M) suggested that LDLR might undergo facilitated turnover through a chloroquine-sensitive route^45^. Consistent with this, chloroquine treatment restored total LDLR levels in mutant cells to those seen in WT (Fig. 6E). This indicated that reduced LDLR abundance is caused by increased turnover and likely underlies reduced LDL clearance in carriers of this variant.

Consistent with altered endocytic trafficking, the p.I2772V variant and heterozygous KO both increased BODIPY-cholesterol uptake, whereas the p.V2821L variant significantly reduced it (Fig. 6F). To assess this in real time, cells were transfected with mCherry-Rab7 and imaged live during BODIPY-cholesterol exposure. Cholesterol delivery to Rab7-positive late endosomes was increased in p.I2772V KI cells but reduced in p.V2821L KI cells at 30 minutes (Video 1-4), consistent with the opposing uptake phenotypes above: p.I2772V is associated with accelerated late endosomal sterol delivery, while p.V2821L shows impaired delivery. While STARD9 colocalization with EEA1 was negligible (suppl. Fig. 4B), its colocalization with the late endosomal marker Rab7 was enhanced in p.I2772V KI cells (suppl. Fig. 4C), consistent with STARD9’s reported binding to NPC1, a late endosomal resident^46^. The functional consequence of this enhanced association warrants further investigation.

Taken together, these findings reveal that while all STARD9 mutant lines share a common defect in lysosomal cholesterol egress and downstream ER cholesterol depletion, they diverge fundamentally in their effects on plasma membrane cholesterol content, endocytic activity, and LDLR surface expression.

The localization of STARD9 to late endosomes and lysosomes, and its proposed role in lysosomal cholesterol transport and positioning, prompted analysis of its interaction with NPC1 across the mutant lines. Co-immunoprecipitation (CoIP) demonstrated STARD9–NPC1 complex formation in all three mutant lines (suppl. Fig. 4D), indicating that the observed differences in lysosomal redistribution are not due to loss of STARD9–NPC1 binding. AlphaFold-Multimer structural models visualized in PyMOL^47^ predicted multiple stable STARD9–NPC1 interfaces with low inter-chain predicted alignment error (PAE < 4 Å) and multiple atomic contacts within 3.5 Å (suppl. Fig. 4E–F): On interface involvies two hydrogen bonds between the NPC1 C-terminus and the STARD9 motor domain. Since neither interface involves the STARD9 START domain, this modeling is consistent with retained NPC1 binding despite loss of START domain function.

### STARD9 Global Knockout Mice Exhibit Enhanced Intestinal Cholesterol Absorption

STARD9 is highly expressed across all human tissues, including the intestine (suppl. Fig. 4A). Given the dramatic clinical response to ezetimibe in the p.V2821L carrier, a variant that reduces STARD9 expression in vitro, and the critical role of the intestinal epithelium as the primary site of dietary cholesterol absorption, we sought to determine whether STARD9 deficiency alters intestinal cholesterol homeostasis in vivo.

STARD9 expression was most prominent at the apical brush border of epithelial cells in the jejunum and terminal ileum, where it colocalized with NPC1L1 (distinct from the nonspecific autofluorescence of the muscularis layer). (Fig. 7A). To directly visualize intestinal cholesterol handling, we administered BODIPY-cholesterol by oral gavage to STARD9 global knockout (KO) and wild-type (WT) mice. Confocal imaging of dissected jejunal segments and isolated intestinal epithelial cells revealed significantly greater BODIPY-cholesterol fluorescence in KO mice compared with WT controls, with prominent signal accumulation in villus enterocytes lining the intestinal lumen (Fig. 7B). Quantification of BODIPY-cholesterol intensity per villus confirmed a marked increase in cholesterol uptake in KO intestinal epithelium, consistent with STARD9 deficiency augmenting cholesterol internalization at the apical enterocyte surface.

**Figure 7:**
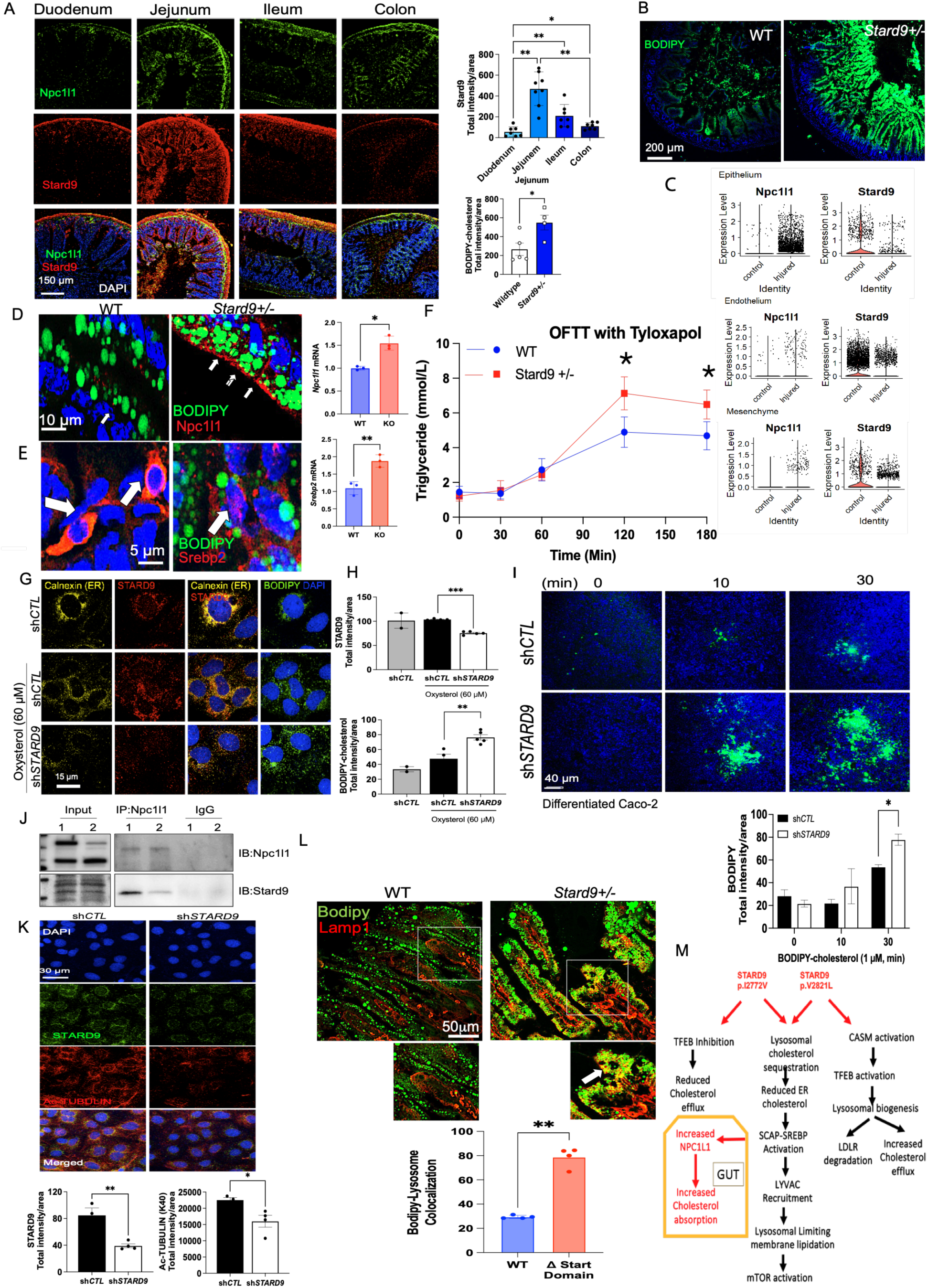
*STARD9* deficiency drives SREBP2 activation, NPC1L1 upregulation, and enhanced intestinal cholesterol absorption *in vivo*. (A) Stard9 tissue expression profiles in mouse intestine and its colocalization with NPC1L1 (B) Confocal imaging of dissected jejunal segments and isolated intestinal epithelial cells from *STARD9* global KO and WT mice following oral BODIPY-cholesterol gavage; KO mice show markedly greater BODIPY-cholesterol fluorescence in villus enterocytes. Quantification of BODIPY-cholesterol intensity per villus is shown (mean ± SEM, unpaired t-test). (C) scRNAseq data demonstrating the expression of NPC1L1 and STARD9 in different gut layer before and after injury(D)NPC1L1 apical membrane protein expression in WT and KO enterocytes (E) increased nuclear SREBP in epithelial cells of STARD9 KO vs WT mice (F) Elevated SREBP2 and NPC1L1 mRNA in the jejunum of STARD9 KO mice. (G) Oral fat tolerance test: plasma triglyceride levels measured at 0, 30, 60, 120 and 180 min following olive oil gavage in *STARD9 KO* vs. WT mice pre-treated with Tyloxapol (0.5 g/kg i.p.) to inhibit lipoprotein lipase; KO mice exhibit significantly higher plasma triglycerides after 60 min of gavage. (H) Increased BODIPY-cholesterol uptake in Caco-2 cells transfected with *STARD9*-specific shRNA compared to WT cells. (I) Time course of BODIPY-cholesterol uptake in differentiated Caco-2 cells showing accelerated uptake kinetics as early as 10-minute following STARD9 knockdown. (J) Co-immunoprecipitation demonstrating physical interaction between STARD9 and NPC1L1 in mouse small intestine. (K) Reduced lysine 40 acetylation of tubulin in Caco-2 cells following *STARD9* knockdown. (L) Bodipy and LysoTracker co-staining of isolated jejunal enterocytes from STARD9 KO mice shows lysosomal cholesterol sequestration comparable to that observed in patient-derived cells. (M) A schematic of disease pathways caused by rare and common STARD9 variants. Data are presented as mean ± SEM. Each dot represents an independent biological replicate. For comparisons among multiple groups, one-way ANOVA with Holm–Šídák post hoc testing was applied when data were normally distributed; otherwise, the Mann–Whitney U test was used for non-normally distributed data. *P ≤ 0.05, **P ≤ 0.01, ***P ≤ 0.001

NPC1L1 is a major protein for the uptake of cholesterol in the gut^48^. scRNA-seq analysis of mouse jejunum before and after injury with 5-FU, which eradicates proliferating cells, showed that reduced STARD9 expression in epithelial cells was associated with a rise in NPC1L1 (Fig. 7C). STARD9 was also expressed in endothelial and mesenchymal cells. Based on our prior data, this raised the question of whether impaired lysosomal cholesterol transport results in SREBP2 activation and a consequent rise in NPC1L1 in enterocytes. Most importantly, NPC1L1 protein was markedly upregulated in the apical membranes of KO enterocytes by immunostaining (Fig. 7D). It has been shown that SREBP2 binds the NPC1L1 promoter to increase its expression^49,50^. Accordingly, nuclear SREBP2 localization was increased in KO versus WT mice (Fig. 7E), and consistent with these findings, NPC1L1 mRNA levels were increased in the jejunum of KO mice (Fig. 7F). These findings reflect a global cholesterol-sensing defect^51^, consistent with prior work showing that intestine-specific (but not liver-specific) SREBP2 overexpression in mice raises jejunal and systemic cholesterol^52^. In summary, our findings in the gut show replication of our in vitro data in vivo, indicating a defective pathway whereby abnormal cholesterol sensing results in SREBP2-dependent NPC1L1 upregulation. To assess the functional consequence of enhanced intestinal cholesterol uptake on systemic lipid absorption, we performed a fat tolerance test using Tyloxapol to inhibit lipoprotein lipase (LPL) activity, thereby preventing peripheral triglyceride clearance and enabling direct quantification of intestinal lipid secretion into the circulation. Following intraperitoneal injection of Tyloxapol (0.5 g/kg), mice received an oral olive oil gavage (10 μl/g bw), and plasma triglycerides were measured serially at 0, 30, 60, 120 and 180 min. STARD9 KO mice exhibited considerably higher plasma triglyceride levels after 120 minutes of gavage, compared with WT controls (Fig. 7G), indicating substantially greater intestinal fat absorption and chylomicron secretion in the absence of STARD9.

To further interrogate the cell-autonomous role of STARD9 in enterocyte cholesterol handling, we used Caco-2 cells, a well-established intestinal epithelial model. Under basal conditions, STARD9 localized to the perinuclear region and co-stained with the ER marker calnexin, consistent with its association with perinuclear late endosomes/lysosomes and ER-proximal membrane contact sites. Treatment with oxysterol (60 μM) mobilized STARD9 from this perinuclear compartment to a more dispersed cytoplasmic distribution, suggesting that STARD9 localization is dynamically regulated by the extracellular sterol environment (Fig. 7H).

BODIPY-cholesterol staining demonstrated increased cholesterol uptake in cells transfected with STARD9-specific shRNA compared to shCTL cells (Fig. 7H), and a time course of Caco-2 cells treated with BODIPY-cholesterol (1 μM) confirmed increased uptake kinetics at 10 and 30 min, following STARD9 knockdown (Fig. 7I). These findings establish that STARD9 suppresses cholesterol absorption in a cell-autonomous manner, independent of systemic factors. Mechanistically, in mice small intestine STARD9 co-immunoprecipitated with NPC1L1 (Fig. 7J), the apical sterol transporter responsible for intestinal cholesterol absorption and the molecular target of ezetimibe. NPC1L1 mediates cholesterol uptake at the brush border membrane through clathrin/AP2-mediated vesicular endocytosis^48^, a process that is blocked by ezetimibe. The physical interaction between STARD9 and NPC1L1 suggests that STARD9-driven lysosomal tubulation may directly regulate NPC1L1 trafficking or recycling at the apical membrane, providing additional mechanistic basis for the observed upregulation of cholesterol absorption upon STARD9 loss.

Finally, STARD9 knockdown was associated with reduced acetylated tubulin levels (Fig. 7K). Kinesin motor proteins preferentially recognize microtubule tracks enriched in acetylated tubulin, and this modification facilitates kinesin-1-mediated cargo transport. The reduction in acetylated tubulin observed upon STARD9 loss likely reflects impaired stabilization of the kinesin-associated microtubule subset as shown for kinesin-1^53^, pointing to a broader role of STARD9 in maintaining the microtubule infrastructure required for organelle motility beyond cholesterol trafficking alone.

Collectively, these in vivo and in vitro data establish that STARD9 deficiency in the intestinal epithelium drives a self-amplifying cycle of SREBP2 activation, NPC1L1 upregulation, and enhanced cholesterol absorption and provides a mechanistic explanation for the selective and robust clinical response to ezetimibe observed in the index patient. The convergence of lysosomal cholesterol trapping, ER sterol sensing failure, and transcriptional upregulation of the principal intestinal cholesterol importer defines a previously unrecognized intestinal axis of dyslipidemia driven by STARD9 dysfunction. Direct imaging confirmed lysosomal cholesterol sequestration within enterocytes of STARD9 KO mice, paralleling the cellular phenotype observed in patient-derived cells (Fig. 7L). (See the schematic of disease pathways caused by rare and common STARD9 variants Fig. 7M).

## Discussion

This study identifies *STARD9* as a lysosomal cholesterol-sensing kinesin and establishes *STARD9* as a causal gene for divergent familial dyslipidemias converging on a shared mechanism of intracellular cholesterol dysregulation. By integrating rare variant segregation, population genomics^15,16^, isogenic CRISPR modeling, single-cell transcriptomics, and in vivo mouse studies, we define a pathway in which *STARD9*, a START domain containing plus-end kinesin-family motor protein^6,7^ regulates lipid handling through differential lysosomal positioning. Although its START domain lacks the conserved residues that form the canonical cholesterol-binding tunnel of STARD1 and STARD4 ^54,55^, we show directly, using an NBD-cholesterol binding assay, that this domain nonetheless binds cholesterol like STARD1 and STARD4 and is required for pericentric lysosomal positioning.

All *STARD9* mutant lines share a dissociation between lysosomal cholesterol content and ER sterol sensing. When *STARD9* is dysfunctional, cholesterol accumulates in late endosomes and lysosomes rather than reaching the ER, triggering a sterol-deprivation transcriptional response in cells that are, paradoxically, cholesterol-laden. This paradox precisely recapitulates the hallmark biochemistry of Niemann–Pick type C (NPC) disease, ^56^ in which NPC1 mutations impair lysosomal cholesterol egress and drive constitutive SCAP-mediated SREBP2 activation. In both NPC disease^32^ and *STARD9* mutant cells, the expanded ER– lysosome contact sites that form because of this defect facilitate cholesterol transfer onto the lysosomal limiting membrane, constitutively activating mTORC1 through the SLC38A9–Rag GTPase–Ragulator complex and suppressing autophagy^57^. The question that follows is why this shared upstream lesion produces opposite plasma lipid phenotypes in different variant carriers.

The answer begins at the structural level. The divergent plasma lipid phenotypes, monogenic hypercholesterolemia versus atherogenic hypertriglyceridemia and low HDL, reflect a context-dependent effect of *STARD9* dysfunction rather than a simple loss-versus gain-of-function dichotomy. The p.V2821L variant phenocopies both the lysosomal cholesterol sequestration and the positioning defect caused by loss of the START domain, in contrast to the perinuclear localization seen with p.I2772V. In conventional kinesin-1, the C-terminal tail domain acts as an intramolecular inhibitor of the motor head by folding back onto the head to suppress microtubule-stimulated ATPase activity, and either cargo engagement or genetic removal of the tail is sufficient to relieve this autoinhibition and unmask processive, plus-end-directed motility^9^. We propose that p.V2821L disrupts the cholesterol-sensing function, relieving this autoinhibition in a functionally equivalent manner, unmasking unopposed anterograde kinesin activity and redistributing lysosomes toward the microtubule plus end at the cell periphery. This variant-specific disruption of lysosomal positioning in turn produces mechanistically divergent downstream phenotypes that mirror the opposing lipid profiles observed in human carriers.

The perinuclear lysosomal retention caused by the GWAS variant p.I2772V is, under this minus-end-directed transport model, predicted to favor delivery of internalized cholesterol toward the perinuclear/ER compartment rather than recycling back to the plasma membrane, offering a structural rationale for its failure to support efflux and for the reduced HDL phenotype of carriers who nonetheless maintain normal LDL cholesterol. This model is inferential and built on an analogy to a well-established paradigm of kinesin regulation; a direct testing, for example plasma membrane cholesterol pool labeling, will be required to confirm it.

This divergence in lysosomal positioning also dictates whether a CASM-dependent compensatory stress response is engaged, explaining the opposing HDL phenotypes of the two variants. In p.V2821L cells, peripheral lysosomal dispersion triggers CASM through V-ATPase–ATG16L1 coupling,^36,40^ driving lipidation of GABARAP onto single lysosomal membranes. CASM-lipidated GABARAP sequesters the FLCN–FNIP1/2 complex away from the Rag GTPase machinery, relieving mTORC1-dependent TFEB phosphorylation and enabling nuclear translocation despite sustained bulk mTORC1 activity^58,59^. Concurrent mTORC1-mediated inhibitory phosphorylation of GSK3β provides a second convergent route to TFEB activation^38,59^. SopF-mediated CASM inhibition that abolishes both LC3B lipidation and TFEB nuclear localization^40^ establishes a causal mechanistic link in p.V2821L cells. TFEB-driven lysosomal exocytosis then delivers lysosomal cholesterol to the plasma membrane^43^, creating a compensatory efflux pathway that preserves HDL cholesterol despite profound lysosomal dysfunction, while the accompanying increase in lysosomal number and peripheral dispersion promotes lysosomal degradation of LDLR, depleting surface receptor abundance and driving the hypercholesterolemic phenotype in rare variant carriers. In p.I2772V cells, by contrast, perinuclear lysosomal retention fails to engage CASM, TFEB remains phosphorylated and inactive, and lysosomal exocytosis is suppressed, depleting plasma membrane cholesterol; instead, p.I2772V enhances *STARD9* association with Rab7-positive late endosomes, increase LDLR levels and augments endocytic cholesterol delivery, a phenotype also produced by heterozygous *STARD9* loss and by *STARD9* knockdown. Together, suppressed TFEB-dependent efflux and enhanced Rab7-mediated cholesterol delivery likely explain the reduced HDL efflux and elevated triglycerides^60^ associated with this GWAS locus. This mechanistic architecture, in which the same upstream lysosomal cholesterol defect produces opposite clinical phenotypes depending on lysosomal spatial organization and the CASM–TFEB axis, represents a previously unrecognized principle of dyslipidemia pathogenesis. Because chronic lysosomal cholesterol sequestration would be expected to reprogram immune cell function as well, we next asked whether this signature was detectable in circulating immune cells from carriers.

Indeed, single-cell transcriptomic profiling of peripheral blood cells from p.V2821L carriers revealed immune–metabolic remodeling consistent with these cellular phenotypes. Classical monocytes, principal drivers of atherogenesis, showed coordinated upregulation of sterol-regulatory genes, alongside expansion of CD16+ NK cells, and MAIT cells, a compositional shift consistent with chronic metabolic and inflammatory stress. These data identify *STARD9* as a kinesin that functions as a central regulator of lysosome-to-ER cholesterol trafficking in human immune cells, although the correlative nature of patient sampling cannot establish causality. To test whether this signature reflects a direct, cell-autonomous consequence of *STARD9* loss, we turned to a controlled cellular model.

*STARD9* knockdown in THP-1 macrophages directly activated the mTOR–NF-κB axis^20,21^, with increased phospho-mTOR (Ser2448), phospho-S6, and phospho-NF-κB p65 (Ser536), establishing that lysosomal cholesterol sequestration is sufficient to engage innate immune pathway activation in a cell-autonomous manner. This inflammatory state, driven by intracellular events rather than circulating lipoproteins, provides a mechanistic basis for atherogenic risk that is constitutively active and not corrected by statin-mediated LDL reduction. Beyond this immune phenotype, the clinical course of the index kindred pointed to a second, tissue-specific consequence of *STARD9* loss: an unusually strong response to intestinal cholesterol absorption inhibition.

Our in vivo intestinal studies provide a molecular explanation for ezetimibe sensitivity of the p.V2821L carriers. In classical familial hypercholesterolemia due to LDLR loss of function, ezetimibe provides only modest additive LDL lowering beyond statins, because the primary defect is hepatic receptor-mediated clearance. The index patient’s outsized ezetimibe response implicated the intestine as a dominant site of pathological cholesterol dysregulation, and our data in *Stard9* knockout mice now provide the mechanism. *STARD9* deficiency in enterocytes triggers constitutive SCAP-mediated SREBP2 processing and subsequent transcriptional upregulation of NPC1L1, a direct SREBP2 target gene^50^, at the apical brush border membrane, creating a feed-forward loop that amplifies luminal cholesterol absorption and chylomicron secretion. Importantly, *STARD9* physically interacts with NPC1L1 in enterocytes, and in KO mice is apically localized, suggesting that beyond transcriptional regulation, *STARD9* may directly modulate NPC1L1 trafficking or recycling at the apical membrane, a mechanistic layer that warrants further investigation.

More broadly, this work positions intracellular cholesterol compartmentalization as a mechanistically distinct and underappreciated axis of cardiovascular risk, providing genetic proof-of-concept that correcting lysosomal cholesterol egress, rather than further lowering circulating lipoproteins, may be required to address the residual inflammatory cardiovascular risk that persists despite optimal medical management. *STARD9* and its interactors, LYVAC/PDZD8, NPC1, NPC1L1, and the CASM–GABARAP–TFEB axis, represent candidate therapeutic nodes for this purpose.

### Ethics Statement (Human Subjects)

Human samples used in this study were obtained from patients under protocols approved by the Institutional Review Board of Yale University All participants or their legal guardians provided written informed consent prior to enrollment. Peripheral blood and/or tissue samples were collected for the purpose of DNA extraction and establishment of patient-derived cells for genetic and functional studies. All procedures involving human subjects were conducted in accordance with institutional guidelines and the principles of the Declaration of Helsinki.

### Animals

All animal procedures were approved by the Institutional Animal Care and Use Committee (IACUC) at Yale University. *Stard9* knockout mice were generated by deleting exons 2–4. Mice were housed in a specific-pathogen-free facility at Yale University on a 12-hour light/dark cycle with ad libitum access to water and standard chow.

## Data Availability

The datasets generated during this study will be deposited in the Gene Expression Omnibus (GEO) repository upon acceptance of the manuscript. Processed data underlying the figures and analyses are included as Supplementary Tables provided with the manuscript.

## Acknowledgment

We thank all patients and their relatives for their participation. We appreciated Dr. Shawn M Ferguson, Dr. Derek Toomre, and Dr. Felix Rivera-Molina, from Yale University for helpful discussions and insights. This work was supported by awards R01HL171054 and R01DK134329 to A.M.

## Methods and Materials

### Study Subjects and Genetic Analysis Kindred Recruitment and Phenotyping

Two independent kindreds presenting with familial dyslipidemia and premature atherosclerosis were recruited from our institutional lipid clinics and cardiovascular disease registries. Kindred 1 consisted of a multi-generational family with documented early-onset ASCVD (myocardial infarction or coronary revascularization before age 55 in males, age 65 in females) and elevated LDL cholesterol (LDL-C >190 mg/dL) or elevated triglycerides (TG >200 mg/dL) in the absence of secondary causes. Kindred 2 was identified from a population-based ASCVD cohort with enriched dyslipidemia phenotypes.

For all subjects, fasting lipid panels were obtained after a 12-hour fast, with quantification of total cholesterol, LDL-C, HDL cholesterol, and triglycerides using standardized laboratory methods. Apolipoprotein B (ApoB), apolipoprotein A-I (ApoA-I), and lipoprotein(a) were measured using immunoassays. Cardiovascular phenotyping included detailed personal and family history of ASCVD, age of first cardiovascular event, and imaging studies (coronary computed tomography angiography, carotid ultrasound) where available. Sequencing of genes associated with monogenic dyslipidemia (LDLR, APOB, PCSK9, APOE, LCAT, CETP) was performed to exclude classical genetic forms.

All subjects provided written informed consent, and the study was approved by the Institutional Review Board. Blood samples were collected and peripheral blood mononuclear cells (PBMCs) were isolated by density gradient centrifugation for downstream analyses.

### Exome and Whole-Genome Sequencing

Whole exome sequencing (WES) was performed on index cases from each kindred using commercially available exome capture kits (SureSelect, Agilent Technologies, Santa Clara, CA) followed by paired-end sequencing on an Illumina platform (HiSeq 4000 or NovaSeq 6000). Sequencing reads were mapped to the human reference genome (GRCh38/hg19) using the Burrows-Wheeler Aligner (BWA). Variants were called using GATK (Genome Analysis Toolkit) best practices pipelines. Variant annotation was performed using ANNOVAR and VEP (Variant Effect Predictor). Rare variants (minor allele frequency <0.01 in gnomAD and ExAC databases) with predicted loss-of-function or damaging missense effects (CADD score >20, REVEL score >0.5) were prioritized.

Familial segregation of identified variants was confirmed by Sanger sequencing in available family members. For Kindred 1, a rare heterozygous missense variant in *STARD9* (NM_001009178.2) was identified in the index case and co-segregated with dyslipidemia and ASCVD phenotypes in affected family members. Unaffected family members were either homozygous for the reference allele or heterozygous carriers with mild lipid abnormalities.

### Genome-Wide Association Study Analysis

A genome-wide association study (GWAS) was conducted in a separate dyslipidemia cohort (n=5,000) recruited from cardiovascular and metabolic disease registries, genotyped using a customized SNP array (Illumina CoreExome) covering approximately 550,000 SNPs. Standard GWAS quality control filters were applied: SNP call rate >99%, Hardy-Weinberg equilibrium p-value >1×10−6, minor allele frequency >1%. Lipid traits (LDL-C, HDL-C, triglycerides, and total cholesterol) were adjusted for age, sex, and ancestry principal components. Association testing was performed using linear regression with additive genetic models. Manhattan plots were generated using standard bioinformatic approaches.

A GWAS-identified common variant in *STARD9* (rs202077402) demonstrated significant association with low HDL-C and elevated triglycerides (p<5×10−8). This variant was a missense change affecting a conserved residue (p.I2772V). The variant frequency in the dyslipidemic cohort was approximately 15-20% heterozygotes and 2-3% homozygotes.

### Cell culture

For extended culture and differentiation experiments, ^61^THP-1 cells (human acute monocytic leukemia cell line, ATCC) were cultured in the same conditions. For monocyte-to-macrophage differentiation, THP-1 cells were treated with 100 nM phorbol 12-myristate 13-acetate (PMA) for 48 hours followed by culture in fresh medium for an additional 24-48 hours.

Human embryonic kidney 293T (HEK293T) cells were cultured in DMEM supplemented with 10% FBS, 100 U/mL penicillin, and 100 μg/mL streptomycin. All cells were maintained at 37°C in humidified air with 5% CO₂ and subcultured at 70-80% confluence.

Human colorectal adenocarcinoma Caco-2 cells (ATCC) were cultured in DMEM containing 10% FBS and 100 U/mL penicillin and 100 µg/mL streptomycin. Cells were seeded onto Transwell plates (Corning, 12 mm diameter inserts, 0.4 µm pore size) for differentiation. Upon reaching 100% confluency on the inserts, the complete medium in both the apical and basolateral compartments was replaced every two days for 21 days. All cells were maintained at 37°C in humidified air with 5% CO₂ and subcultured at 70-80% confluence.

### Cell Culture and CRISPR-Mediated Gene Editing Primary Cell Isolation and Culture

Patient-derived peripheral blood mononuclear cells (PBMCs) were isolated from whole blood drawn into EDTA-coated vacutainer tubes by density gradient centrifugation using Ficoll-Paque (GE Healthcare, Chicago, IL). Cells were cultured in RPMI 1640 medium supplemented with 10% fetal bovine serum (FBS), 100 U/mL penicillin, and 100 μg/mL streptomycin at 37°C in 5% CO₂.

### CRISPR/Cas9 Gene Editing Strategy

Heterozygous knock-in HEK293T cell lines carrying *STARD9* p.V2821L or p.I2772V variants and a heterozygous knockout line (exons 5–7 deletion causing frameshift) were generated using CRISPR/Cas9. All guides and donor sequences were designed and obtained from Integrated DNA Technology (IDT, USA). For comparison, wild-type control cell lines (STARD9-WT) were generated with the same CRISPR construct but with inactivating mutations in the Cas9 nuclease domain.

For making KI cell lines, the guide RNA sequences were: TGTAGAATTCTGAACATCAC for p.V2821L and ACTGCAGAGGGCATACCCCC for p.I2772V. Guide RNAs, Donors and GFP conjugated Cas9 protein were transfected into HEK293T cells using CRISPRMAX lipofectamine (Invitrogen, USA) according to the manufacturer’s protocol. Forty-eight to seventy-two hours post-transfection, GFP positive cells were isolated using a BD FACS Aria flow cytometer. Single cells were collected in 96 well plates. Single-cell colonies were then expanded for the selection of clones with desired mutations.

For making *STARD9* knockout, the guide RNA sequences were: CCGTGGTGTCAACCCAACAG for Exone5 and CGGGATCTGTTGAAGCAATC for exon 7.

*STARD9* knockout was confirmed by: (1) DNA sequencing to document insertions/deletions mutations introduced by CRISPR; (2) quantitative reverse-transcription PCR (qRT-PCR) confirming reduction of *STARD9* mRNA; and (3) Western blotting demonstrating reduction of STARD9 protein.

Similarly, START domain deletion was performed by using guide RNA sequences:

Guide 1: CGAACAAGCAGCCATTACCT and Guide 2: ACGGGAGAGGTGAACCCAAC and confirmed by sequencing (suppl. Fig 5 A-B).

### NBD-Cholesterol Binding Experiment

A fluorescent sterol binding assay was performed as previously described^62^, with some modifications. Briefly, purified START proteins were added at different concentrations in PBS to a 96-well black flat-bottom plate. 22-NBD-cholesterol (3 μM, HY-W020012, MedChemExpress) was then added and equilibrated at 37 °C for 30 min. Measurements were made using a BioTek Synergy H1 microplate reader. The NBD excitation and emission wavelengths were 470 nm and 525 nm, respectively, and the resulting data were plotted using Prism software.

### Cloning, Protein Expression, and Purification

Human START domains of STARD9 (4483–4700), STARD4 (1–205) and STARD1 (67-280) were cloned into the p3×FLAG-CMV backbone (Plasmid #20011) (suppl. Fig 6 A-B). HEK293 cells were transfected with vectors for each START domain protein separately using Lipofectamine 3000 (Invitrogen). After 48 hours, cells were harvested and lysed in IP lysis buffer (Invitrogen) with 1% protease inhibitor cocktail (Invitrogen). Proteins were purified using anti-FLAG magnetic beads (HY-K0207, MedChemExpress) according to the manufacturer’s protocol. Briefly, cell lysates containing FLAG-tagged proteins were incubated with anti-FLAG beads overnight at 4 °C with gentle rotation. Tubes were then placed on a magnetic stand to collect the beads against the side of the tube, and the supernatant was removed and discarded. Beads were washed four times with wash buffer (TBST: 50 mM Tris-HCl, 150 mM NaCl, 0.5% Tween-20, pH 7.4) and eluted with 1 mg/mL 3×FLAG peptide in elution buffer (50 mM Tris, 0.15 M NaCl, pH 7.4; MedChemExpress).

### Lipid accumulation and cholesterol trafficking assays

Cells were exposed to BODIPY-cholesterol (2 µM) or oxidized LDL (OxLDL, 100 µg/ml) to assess intracellular lipid handling. Neutral lipid accumulation was evaluated by BODIPY staining, and cholesterol distribution by filipin staining. Oil Red O staining quantified esterified cholesterol.

Time-course experiments assessed lysosomal retention versus clearance. For cholesterol egress assays, PMA-differentiated THP-1 cells were treated with OxLDL and analyzed over 60 minutes.

### Subcellular Fractionation and measurement of ER cholesterol content

ER fraction was isolated using Minute^TM^ ER Enrichment Kit (ER-036, Invent Biotechnologies, Inc.). Briefly, 30 X 10^6^ cells were collected through low-speed centrifugation (600 X *g* for 5 min) washed once with cold PBS, and pellets were snap-frozen at −80°C for 10 min. Cells were resuspended in 550 µl Buffer A, vortexed vigorously for 20–30s and passed through a filter cartridge. The filtrate was centrifuged at 2,000 × *g* for 3 min to remove nuclei and large debris. The supernatant was transferred to a fresh tube and centrifuged at 8,000 × *g* for 10 min at 4°C to remove mitochondria, lysosomes, and membrane debris. The cleared supernatant (400 µl) was mixed with Buffer B (1:10, v/v) and incubated at 4°C for 20–30 min, followed by centrifugation at 8,000 × *g* for 10 min. The resulting pellet was resuspended in 400 µl cold Buffer A and incubated with Buffer C (1:10, v/v) for 10–15 min at room temperature with intermittent mixing. After centrifugation (8,000 × *g*, 5 min), the supernatant was collected and mixed with Buffer D (1:1, v/v), incubated at 4°C for 20 min, and centrifuged at 10,000 × *g* for 10 min. The final pellet was washed briefly, residual supernatant was removed, and the ER-enriched fraction was resuspended in 100 µl detergent-containing buffer.

Lipids were extracted from isolated ER fractions using a modified Folch procedure^63^. Briefly, ER pellets were homogenized on ice in chloroform: methanol (2:1, v/v) for cell pellets using a glass homogenizer. The homogenate was transferred to glass tubes for phase separation. Water was added to achieve a final chloroform: methanol: water ratio of approximately 8:4:3 (v/v/v), followed by gentle mixing. Samples were incubated on ice for 10–15 min and centrifuged at 2,000 × *g* for 10 min at 4°C to promote clear phase separation. The lower organic phase containing lipids was carefully collected without disturbing the interphase. The aqueous phase was re-extracted with additional chloroform to maximize lipid recovery, and the organic phases were pooled. Solvents were evaporated under nitrogen and the dried lipid extracts were resuspended in assay buffer and total cholesterol was quantified using Cholesterol/ Cholesteryl Ester Assay Kit (ab65359, Abcam).

### Membrane fraction isolation for LDLR protein expression and cholesterol quantification

Cells in all four groups were treated with methyl-β-cyclodextrin (MβCD) (HY-101461, MedChemExpress) at a concentration of 1mM for 1 h, the media was washed and incubated with lovastatin (HY-N0504, (5μg/ml, MedChemExpress) for 3 h. Plasma membrane fraction was isolated using Minute™ Plasma Membrane/Protein Isolation and Cell Fractionation Kit (SM-005, Invent Biotechnologies, Inc.). Briefly, 30 X 10^6^ cells were collected through low-speed centrifugation (600 X *g* for 5 min) washed once with cold PBS, resuspended the pellet in 500μl of buffer A, vortexed vigorously for 20–30s and passed through a filter cartridge. The filtrate was then centrifuged at 16,000 X g for 30 seconds. Pellet was resuspended by vigorously vertexing for 10 seconds and centrifuged at 700 X g for one min (the pellet contains intact nuclei). Supernatant was transferred to a fresh 1.5 ml microcentrifuge tube and centrifuged at 4°C for 30 min at 16,000 X g. The supernatant was removed and resuspended in 200 µl buffer B by repeatedly pipetting up and down or vertexing and then centrifuged at 7,800 X g for 20 min at 4°C. Supernatant was carefully transferred to a fresh 2.0 ml microcentrifuge tube and 1.6 ml cold PBS was added to it. The mixture was then centrifuged at 16,000 X g for 30 min to obtain pellet contains plasma membrane fraction.

Total cholesterol was extracted from plasma membrane fraction as described in previous section using a modified Folch procedure^63^.

To check the LDLR protein expression in plasma membrane the membrane fraction was subjected to 4-15% SDS-PAGE and detected by using anti-LDLR (1:5000, 66414-1-Ig, Proteintech) antibody.

### Immunofluorescence Microscopy and Imaging Analysis Fixed-Cell Imaging

Cells were cultured on poly-D-lysine or collagen-coated coverslips, then fixed with 4% paraformaldehyde in PBS for 15 minutes at room temperature. After permeabilization with 0.1% Triton X-100 in PBS for 10 minutes and blocking in 3% bovine serum albumin (BSA) for 30 minutes, cells were incubated with primary antibodies overnight at 4°C. Primary antibodies were all previously validated and included: anti-LAMP1^64^ (1:200, 65051-1-Ig, Proteintech), anti-calnexin^65^ (1:2000, 66903-1-Ig, Proteintech), anti-α-tubulin^66^ (1:1000, #3873, Cell Signaling Technology) or anti-α-tubulin acetyl K40 antibody^67^ (1:100, EPR16772, abcam), anti-STARD9 (1:200, PA5-113470, Thermo Scientific), anti-SREBP-2^68^ (1:200, sc-13552, Santa Cruz Biotechnology), Anti-EEA1^69^ (1:1500, 68065-1-Ig, Proteintech), Anti-Rab7^70^ (1:100, 9^16^46, Cell Signaling Technology), Anti-PDZD8 (1:1000, 25512-1-AP, Proteintech)

Following primary antibody incubation, cells were washed with PBS containing 0.1% Tween-20 and incubated with appropriate secondary antibodies conjugated to Alexa Fluor fluorophores (1:500, Molecular Probes/Thermo Fisher) for 1 hour at room temperature. Coverslips were mounted using ProLong™ Gold Antifade Mountant with DNA Stain DAPI (P36935, Invitrogen) and imaged on a laser-scanning confocal microscope (Leica SP8) using appropriate laser lines. Z-stacks of 0.5-1.0 µm optical sections were collected for three-dimensional reconstruction

### Quantitative Image Analysis

All quantifications including Lysosomal number, size, and clustering, bodipy and transferrin uptake and ER-lysosome contact sites were performed using Fiji (version 2.15.0) image analysis. ER-lysosome contact sites were defined as regions of colocalization (Pearson correlation coefficient >0.5).

### Live-Cell Imaging

For dynamic imaging, cells were cultured in glass-bottom 35-mm imaging dishes and maintained at 37°C in 5% CO₂ during imaging using an incubation chamber. Cells were imaged every 1 minute for 20-60 minutes using an inverted confocal microscope (Leica TCS SP8, Germany).

Lysosomes were labeled with LysoTracker Red DND-99 (Apex Bio B8814, 100 nM), and Lysotracker Green DND-26 (Thermo scientific, L7526, 75 nM). Endoplasmic reticulum and microtubules were visualized with ER tracker (Thermo scientific T-34250, 100nM) and Tubulin Tracker green (Thermo scientific T-34075,100nM) respectively. Before adding trackers, cells cultured for 24 hours, starved in LPDS containing media for 1 hour then oxycholesterol was added in 60 μM final concentration for 4 hours. For transferrin uptake cells cultured for 24 hour and then starved in LPDS containing media for 1 hour. Cells were placed on ice for 10 minute to inhibit endocytosis. Then Alexa 488 conjugated transferrin (Thermo scientific T-13342, 25 μg/ml) was added and live image were taken for 30 minutes. To check BODIPY-cholesterol uptake, cells cultured for 24 hour and starved in LPDS containing media for 1 hour then BODIP-cholesterol (Med Chem Express HY-125746, 2μM) was added and live image were taken for 60 minutes. ER-lysosome contact sites were visualized by co-transfection of LAMP1-YFP (#1816, addgene) and mCherry-ER-3 (#55041, addgene) vectors for calnexin (ER marker) and LAMP1 (lysosomal marker) respectively. Transfections were performed using lipofectamine 3000 reagents (Invitrogen, USA), and cells were analyzed 24–48 h later.

### Lyso-IP and lysosomal cholesterol quantification

We performed Lyso-IP following previously described method^71^. Briefly, approximately 35 million cells were cultured in 15 cm plates for each experimental sample. Cells were transfected with TMEM192-3xHA (pLJC5-Tmem192-3xHA, #102930, addgene) using lipofectamine 3000 reagent. 36h of post-transfection, cells were incubated with 60μM Oxysterol for 16 h. On termination of incubation cells were rinsed twice with ice-cold PBS before scraping them into 1 mL of KPBS (136 mM KCl, 10 mM, pH 7.25 adjusted with KOH). The suspension was then centrifuged at 1000 x *g* for 2 minutes at 4°C. Following centrifugation, the cell pellet was resuspended in 1 ml of KPBS. The cell suspension was subjected to mechanical lysis via 20 strokes in a pre-chilled 2 ml Dounce homogenizer. This homogenate was then centrifuged at 1000 x *g* for 2 minutes at 4°C to pellet unlysed cells and nuclei. The resulting supernatant, containing the cellular organelles and lysosomes, was collected and immediately incubated with 150 μL of KPBS-prewashed anti-HA magnetic beads. This incubation was carried out on a gentle rotator shaker for 15 minutes at 4°C. Finally, the lysosome-bound immunoprecipitates were sequestered using a DynaMag Spin Magnet and gently washed three times with ice-cold KPBS before proceeding to cholesterol extraction by using a modified Folch procedure^63^.

### Gene Expression Analysis

#### Quantitative Reverse-Transcription PCR (qRT-PCR)

Total RNA was extracted from cells using TRIzol reagent (Invitrogen, Carlsbad, CA). RNA concentration was determined by spectrophotometry and RNA integrity verified using an Agilent 2100 Bioanalyzer. cDNA was synthesized from 1-2 μg total RNA using iScript cDNA synthesis kit (Bio Rad, USA). Quantitative PCR was performed in triplicate using SYBR Green qPCR Master Mix (Bio Rad, USA) on a C1000 Touch Real-Time PCR System (Bio Rad, USA). Primer sequences for genes of interest are listed in Table 7. Target genes included: *STARD9, SREBP-2 (SREBF2), HMGCOA* reductase (*HMGCR*), *LDLR, ABCA1, CD36, LDLR, TNF-α, IL-1β, S100S8*. Normalization was performed using geometric mean of housekeeping genes *GAPDH*. Relative gene expression was calculated using the ΔΔCt method. Fold-change in mRNA levels was expressed relative to wild-type control cells (STARD9-WT), with statistical significance determined by one-way ANOVA (p<0.05).

#### shRNA mediated *STARD9* gene silencing

Differentiated THP-1 macrophage was cultured in RPMI media supplemented with 10% FBS and HEK293T and Caco-2 cells were cultured in DMEM supplemented with 10% FBS and transfected at 70%-80% confluence with *STARD9* specific shRNA vector (sc-63083-SH, Santa Cruz, USA) using lipofectamine 3000 (Invitrogen, USA). Scrambled shRNA was used as a control. 48 hours post transfection cells were used for experiments. Knock out efficiency was validated via RT-qPCR.

#### Single-Cell RNA Sequencing (scRNAseq)

Fresh PBMCs from two *STARD9* p.V2821L variant carriers (father and son) and 20 age- and sex-matched non-carrier controls were isolated as described above. Cell viability was confirmed to be >90% by trypan blue exclusion. scRNAseq libraries were prepared using the 10X Genomics Chromium Single Cell 3’ GEM Kit v3.1 according to manufacturer’s protocols, targeting ∼5,000-8,000 cells per sample. cDNA amplification and sequencing library preparation were performed as specified. Libraries were sequenced on an Illumina NovaSeq 6000 to a minimum depth of 50,000 reads per cell.

Raw sequencing data were demultiplexed and analyzed using Cell Ranger pipeline (10X Genomics). Quality control filtering included removal of cells with <500 genes detected, >5% mitochondrial reads, or potential doublets identified by DoubletFinder. Downstream analysis was performed using Seurat (v3.2) in R. Dimensionality reduction (UMAP), clustering, and cell type identification were performed using standard workflows. Cell-type annotation was further refined using Azimuth (v0.5.0), which maps query cells onto a curated reference atlas via supervised anchor-based label transfer; resulting predicted labels and mapping scores were used to assign high-confidence cell identities. Cell types were annotated based on canonical marker genes (CD14+CD16− for classical monocytes, CD4+ for T cells, CD8+ for cytotoxic T cells, CD19+ for B cells, CD56+ for NK cells, etc.).

Differential expression analysis between patient and control samples was performed using the MAST statistical framework, comparing lipid metabolism and autophagy pathway genes across matched cell types. Gene set enrichment analysis (GSEA) was performed using KEGG to identify dysregulated pathways.

#### Western Blotting

Cells were lysed in ice-cold RIPA buffer (89900, Thermo Scientific™) with protease and phosphatase inhibitor cocktails (Halt™ Protease Inhibitor Cocktail, EDTA-Free (100X), 87785, Thermo Scientific™). Protein concentration was determined using the BCA protein assay kit (A65453, Thermo Scientific™). Equal amounts of protein (50 μg) were separated by SDS-PAGE on 4-15% polyacrylamide gradient gels and transferred to PVDF membranes at 100V for 90 minutes.

#### Antibody Incubations and Detection

Membranes were blocked with 5% non-fat milk or 5% BSA in Tris-buffered saline containing 0.1% Tween-20 (TBST) for 1 hour at room temperature, then incubated with primary antibodies overnight at 4°C. Primary antibody used in this study were validated by prior publication or by us. Primary antibodies and their dilutions included: anti-STARD9 (1:1000, PA5-113470, Thermo Fischer Scientific), anti-CD68^72^ (1:1000, 14-0688-82, Invitrogen), anti-SREBP-2^73^ (1:250, 557037, BD Pharmingen), anti-Histone H4^74^ (1:500, 16047-1-AP, Proteintech), anti-phospho-mTOR Ser2448^75^ (1:1000, 2971, Cell Signaling Technology), anti-mTOR^76^ (1:1000, 2983, Cell Signaling Technology), anti-phospho-S6 Ser235/236^77^ (1:2000, #4858, Cell Signaling Technology), anti-S6^78^ (1:1000, #2217, Cell Signaling Technology), anti-phospho-NF-kappaB Ser536^79^ (1:1000, #3033, Cell Signaling Technology), anti-NF-kappaB^77^ (1:1000, #8242, Cell Signaling Technology), anti-phospho-TFEB Ser211^80^ (1:1000, #37681, Cell Signaling Technology), anti-TFEB^81^ (1:1000, A7311, Abclonal), anti-LC3B^82^ (1:1000, 2775, Cell Signaling Technology), anti-p62/SQSTM1^83^ (1:1000, A11483, Abclonal), anti-NPC1^56^ (1:500, sc-271335, Santa Cruz Biotechnology), Anti-NPC1L1^84^ (1:100, sc-166802, Santa Cruz Biotechnology), and anti-β-Actin^85^ (1:1000, #4970, Cell Signaling Technology)

After washing in TBST, membranes were incubated with HRP-conjugated anti-mouse or anti-rabbit secondary antibodies (1:1000, Cell Signaling Technologies)) for 1 hour at room temperature. Protein detection was performed using enhanced chemiluminescence (ECL) substrate (32209, Pierce™ ECL Western Blotting Substrate) and imaged on a G:BOX, Syngene imaging system (Cambridge, UK). Densitometric quantification was performed using ImageJ software, with intensities normalized to β-Actin.

#### Co-immunoprecipitation

Cells from WT and START domain deleted STARD9 (STARD9-ΔSTART) and in parallel WT, KO, p.V2821L, rs202077402 were starved in lipoprotein deficient medium for 2h and then incubated with 25-HC (60μM) for 4h and lysed in ice-cold Pierce IP lysis buffer (Thermo Fisher Scientific, 87788) containing protease and phosphatase inhibitors. The small intestine from wildtype mice was lysed in ice-cold IP lysis buffer containing protease and phosphatase inhibitors and homogenized with beads. Immunoprecipitation was performed according to manufacturer’s protocol using Pierce™ Protein A/G Magnetic Beads (88802). Briefly, 500μg protein lysate was incubated with 10μg anti-NPC1 antibody (sc-271335, Santa Cruz Biotechnology), 10 µg anti-NPC1L1 antibody (sc-166802, Santa Cruz Biotechnology), or normal mouse IgG (#68860, Cell Signaling Technology) for overnight at 4°C. Next day 25μl of beads were washed in TBST (TBS/Tween20, 0.05% v/v) and incubated for 60 mins with protein samples/antibody mixture at RT on a rocking platform. After incubation, beads were washed 2 times with TBST, and the supernatant was carefully removed. Then proteins were eluted from the beads with 4X Laemmli sample buffer (Bio-Rad) followed by heated at 90°C for 10 mins and analyzed by western blot. A measure of 10% of the IP homogenate (50 μg) was taken for examination of the input and loading control.

#### Antibody validation

All antibodies were validated either by prior publications or by the manufacturer. According to the manufacturer’s datasheets, Anti-CD68 (14-0688-82, Invitrogen) was validated using THP-1 monocyte differentiation. The anti–acetyl-α-tubulin (K40) antibody (EPR16772, Abcam) was validated by increased acetylation following Trichostatin A treatment. Anti-TFEB (A7311, Abclonal) was validated by TFEB knockdown. Anti-phospho-mTOR (#2971, Cell Signaling Technology) showed increased signal following EGF treatment, and anti-mTOR (#2983, Cell Signaling Technology) was validated using siRNA-mediated knockdown. Anti-phospho-S6 (#4858, Cell Signaling Technology) showed increased signal after hIGF-1 treatment, and anti-phospho-NF-κB (#3033, Cell Signaling Technology) was validated by increased signal following TNF-α stimulation. Histone H4 (16047-1-AP, Proteintech), HNF1α (22426-1-AP, Proteintech), EEA1 (68065-1-Ig, Proteintech), and Calnexin (66903-1-Ig, Proteintech) were validated by shRNA-mediated knockdown according to the datasheets. Anti-STARD9 (PA5-113470, Thermo Fisher Scientific) was validated in our laboratory using a CRISPR-generated *STARD9* knockout HEK cell line.

#### Nuclear protein fractionation and isolation

Cell nuclear extracts were prepared from WT, KO, p.V2821L, rs202077402 cells using NE-PER Kit (Thermo 78833). Cells were harvested and washed with ice-cold PBS, then resuspended in ice-cold CER I buffer, vortexed and incubated on ice for 10 minutes. Ice-cold CER II was added, vortexed. After 1 minute on ice, cells were centrifuged at 16000 x *g* for 5 minutes. Pellets were suspended in ice-cold NER buffer vortex vigorously in every 10 minutes till 40 minutes. After centrifugation at 16,000×*g* for 10 minutes, the supernatant (nuclear extract) was recovered and western blot was performed for SREBP2 protein expression analysis.

#### Electrophoretic Mobility Shift Assay (EMSA)

An electrophoretic mobility shift assay (EMSA) was performed to determine whether HNF1α binds to a sequence in the *STARD9* gene flanking the p.V2821L variant. A 65-bp FAM-labeled double-stranded oligonucleotide spanning the predicted *HNF1α* motif was synthesized (W.M. Keck Oligonucleotide Synthesis Facility, Yale University) and incubated with nuclear extract from HepG2 cells for 20 minutes in RT. Reactions (20 μL) contained 10nM probe, 10 μg nuclear extract, 10mM Tris-HCL, pH 7.5, 10mM NaCl, 40mM KCl, 1mM MgCl2, 1mM EDTA pH8.0, 1mM DTT, 10mg/ml BSA and ddH2O). For supershift assays, 1 μLanti-HNF1α antibody^86^ (22426-1-AP, Proteintech) was pre-incubated with the extract before adding the probe. Samples were resolved on a 5% native polyacrylamide gel in 0.5x TBE at 80V for 30, gel was visualized in FAM detection system (G:BOX, Syngene).

#### SopF-mediated inhibition of CASM

To assess the contribution of conjugation of ATG8 to single membranes (CASM) to lysosomal signaling, cells were transiently transfected with a plasmid encoding the bacterial effector SopF (pEGFP-C1-SopF; Addgene plasmid #137734). SopF selectively disrupts the interaction between the V-ATPase complex and ATG16L1, thereby inhibiting CASM without impairing canonical autophagosome formation. Cells were plated to achieve ∼60–70% confluence at the time of transfection and transfected with pEGFP-C1-SopF using Lipofectamine 3000 (Invitrogen, USA), according to the manufacturer’s protocol. Control cells received the corresponding empty vector where indicated. After transfection, cells were incubated for 24–48 h to allow expression of SopF prior to downstream analyses.

CASM activity was evaluated by measuring LC3B lipidation and TFEB signaling. Whole-cell lysates were prepared in RIPA buffer supplemented with protease and phosphatase inhibitors, resolved by SDS–PAGE, and immunoblotted for LC3B and phospho-TFEB. SopF expression was verified by GFP fluorescence and immunoblotting where required. Changes in LC3B-II accumulation and TFEB phosphorylation were interpreted as readouts of CASM-dependent lysosomal signaling.

#### Cholesterol efflux assay

Cells were seeded at a density of 1 x 10^6^ cells per well one day prior to loading with Bodipy-cholesterol (2μM) for 24 h. Then, cells were washed twice with PBS and incubated in RPMI supplemented with 5% lipoprotein deficient medium for 4 h prior to addition of 35 μg/ml of HDL. Supernatants were collected after 6 h and Bodipy fluorescence intensity was measured using fluorometer. Similarly, cells were lysed to measure the intracellular Bodipy-cholesterol. Cholesterol efflux was calculated as: [(medium Bodipy− background)/ (intracellular Bodipy)] × 100%. Results were expressed as percentage of total cellular Bodipy that was released into the medium. For each cell type and efflux condition, n=5 biological replicates were performed.

#### Generation and Housing of STARD9 Global Knockout Mice

STARD9 global knockout (KO) mice were generated by JAX using CRISPR/Cas9-mediated deletion of exons 5 through 7, introducing a frameshift mutation that results in nonsense-mediated mRNA decay and absence of detectable STARD9 protein, confirmed by RT-qPCR and Western blotting. Mice were maintained on a C57BL/6J background. For all intestinal absorption experiments, age-matched (8–12 weeks), sex-matched STARD9 KO and wild-type (WT) littermate controls were used. Mice were housed in a specific pathogen-free facility under standard light–dark cycles (12h/12h) with ad libitum access to standard chow and water unless otherwise specified. All animal procedures were conducted in accordance with the guidelines of the Yale University Institutional Animal Care and Use Committee (IACUC) under approved protocols.

#### In Vivo Intestinal Cholesterol Uptake Assay (BODIPY-Cholesterol Gavage)

To visualize intestinal cholesterol uptake in vivo, STARD9 KO and WT mice were fasted for 16 hours and then administered BODIPY-cholesterol (BODIPY Cayman Chemical; 3μg/gr body weight in olive oil) by oral gavage. Mice were sacrificed 1 hour post-gavage by CO₂ asphyxiation followed by cervical dislocation. The proximal jejunum (5 cm segment starting 2 cm distal to the ligament of Treitz) was harvested, flushed with ice-cold PBS, and fixed with 4% paraformaldehyde in ice for 2h in dark followed by overnight incubation with 30% sucrose solution at 4°C. For whole-mount or cryo-section imaging, fixed tissue was embedded in OCT compound, cryosectioned at 5 μm, counterstained with DAPI (1 μg/ml), and imaged by confocal microscopy (Leica SP8) using appropriate BODIPY excitation/emission parameters (488/505–540 nm). BODIPY-cholesterol fluorescence intensity per villus unit or per cell was quantified using Fiji (v2.15.0) with at least 20 villi or 100 cells per animal analyzed. Data are expressed as mean integrated fluorescence intensity ± SEM, with n ≥ 6 mice per genotype. Statistical significance was assessed by unpaired t-test.

#### Oral Fat Tolerance Test with Tyloxapol–Mediated LPL Inhibition

Intestinal lipid absorption capacity was quantified using an oral fat tolerance test with lipoprotein lipase inhibition. STARD9 KO and WT mice were fasted overnight prior to the experiment and orally gavaged with olive oil (Sigma-Aldrich, 01514, 10 µL/g body weight). Twenty minutes after lipid administration, mice received an intraperitoneal injection of Tyloxapol (Triton WR-1339; GLPBio, Cat# GC30017) at a dose of 0.5 g/kg body weight to inhibit peripheral triglyceride lipolysis. Tyloxapol was dissolved in sterile 0.9% sodium chloride solution at a concentration of 100 mg/mL before use. Blood samples were collected via submandibular bleeding at the indicated time points, and plasma triglyceride levels were measured using a commercially available triglyceride assay kit (ELABSCIENCE BIONOVATION INC, Cat#E-BC-K109-S) according to the manufacturer’s instructions.

#### Intestinal SREBP2 and NPC1L1 Expression Analysis

Immunohistochemistry of jejunal segments was performed by embedding the Jejunum from WT and KO groups in OCT molds and sectioning at 5 μm. Sections were permeabilized using 0.1% triton in PBS for 10 minutes in room temperature and blocked with 5% normal goat serum, 30 min, and incubated overnight at 4°C with anti-SREBP2 (1:200) or anti-NPC1L1 (1:200) antibodies. Signal was detected using HRP-conjugated secondary antibody, counterstained with DAPI and imaged by confocal microscopy (Leica SP8) using appropriate laser. For each marker, n ≥ 4 mice per genotype were analyzed; statistical comparisons used the unpaired t-test or Mann–Whitney U test as appropriate.

#### Statistical Analysis

All quantitative data are expressed as mean ± standard error of the mean (SEM) or standard deviation (SD) as specified. For normally distributed data, statistical comparisons between two groups were performed using unpaired Student’s t-test (two-tailed). Comparisons among more than two groups were performed using one-way or two-way ANOVA followed by Bonferroni post-hoc tests. For non-normally distributed data, Mann-Whitney U test or Kruskal-Wallis test was used. Correlation analyses were performed using Pearson or Spearman correlation coefficients as appropriate.

A p-value <0.05 was considered statistically significant. All analyses were performed using GraphPad Prism v8.0 (GraphPad Software, San Diego, CA) or R statistical software (R Foundation for Statistical Computing, Vienna, Austria). Multiple testing corrections (false discovery rate, Benjamini-Hochberg procedure) were applied where appropriate for transcriptomic and proteomic analyses involving multiple comparisons.

---

**Suppl. Fig. 1:**
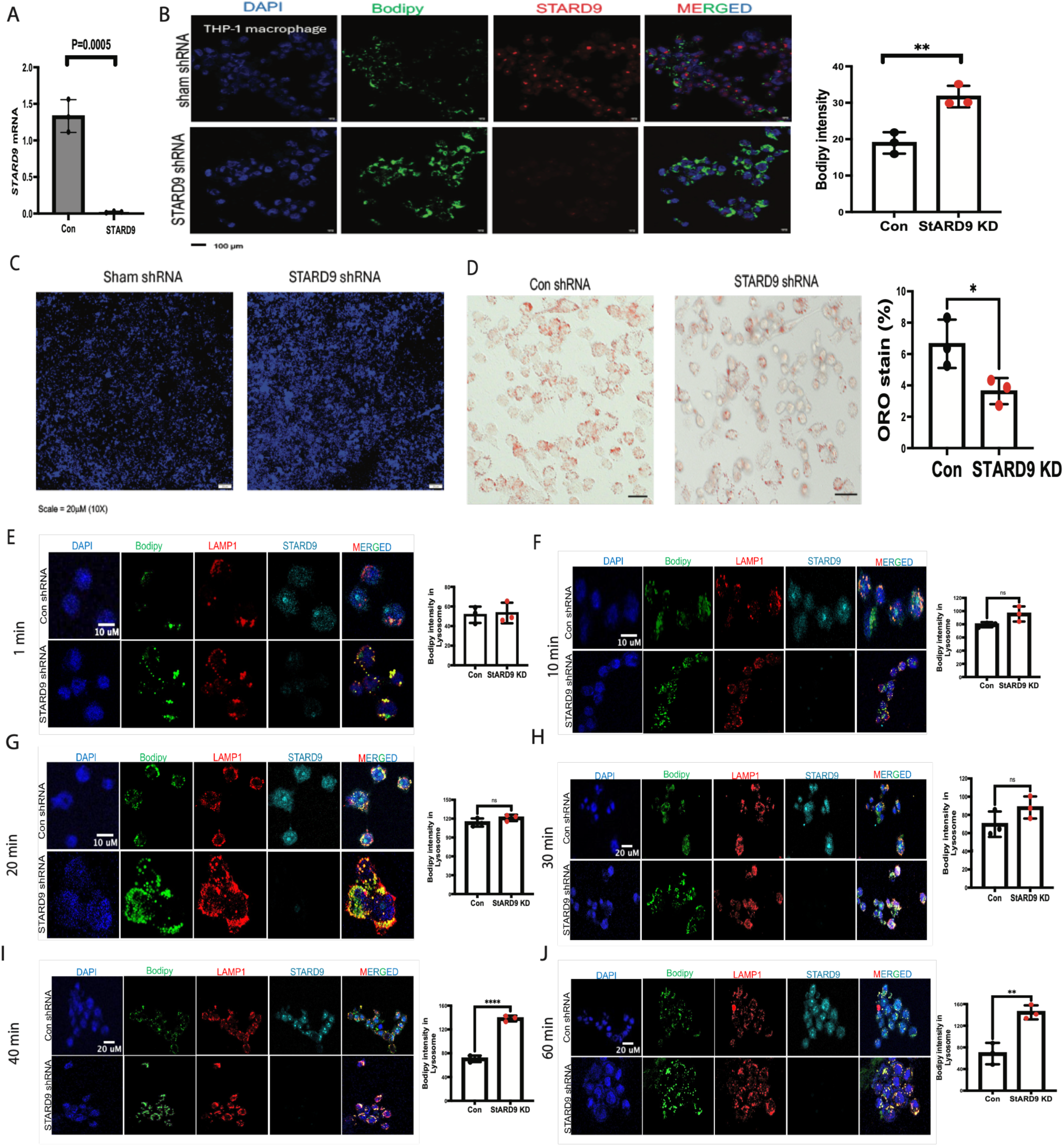
*STARD9* Knockdown in THP-1 Cells Recapitulates Lysosomal Cholesterol Accumulation and Activates mTOR–NF-κB Inflammatory Signaling. (A) Near-complete STARD9 mRNA knockdown in THP-1 cells using specific shRNA (B) Increased intracellular neutral lipid accumulation (BODIPY staining) in *STARD9*-knockdown cells following 6 hours of BODIPY-cholesterol exposure. (C) Increased free cholesterol accumulation in knockdown cells, assessed by filipin staining. (D) Reduced esterified cholesterol content in knockdown cells, assessed by Oil Red O staining. (E–J) Time-course analysis of lysosomal cholesterol clearance following OxLDL (100 µg/mL) treatment in PMA-differentiated THP-1 cells: wild-type cells clear lysosomal cholesterol by 60 minutes, whereas *STARD9*-deficient cells exhibit persistent lysosomal retention

**Suppl. Fig. 2:**
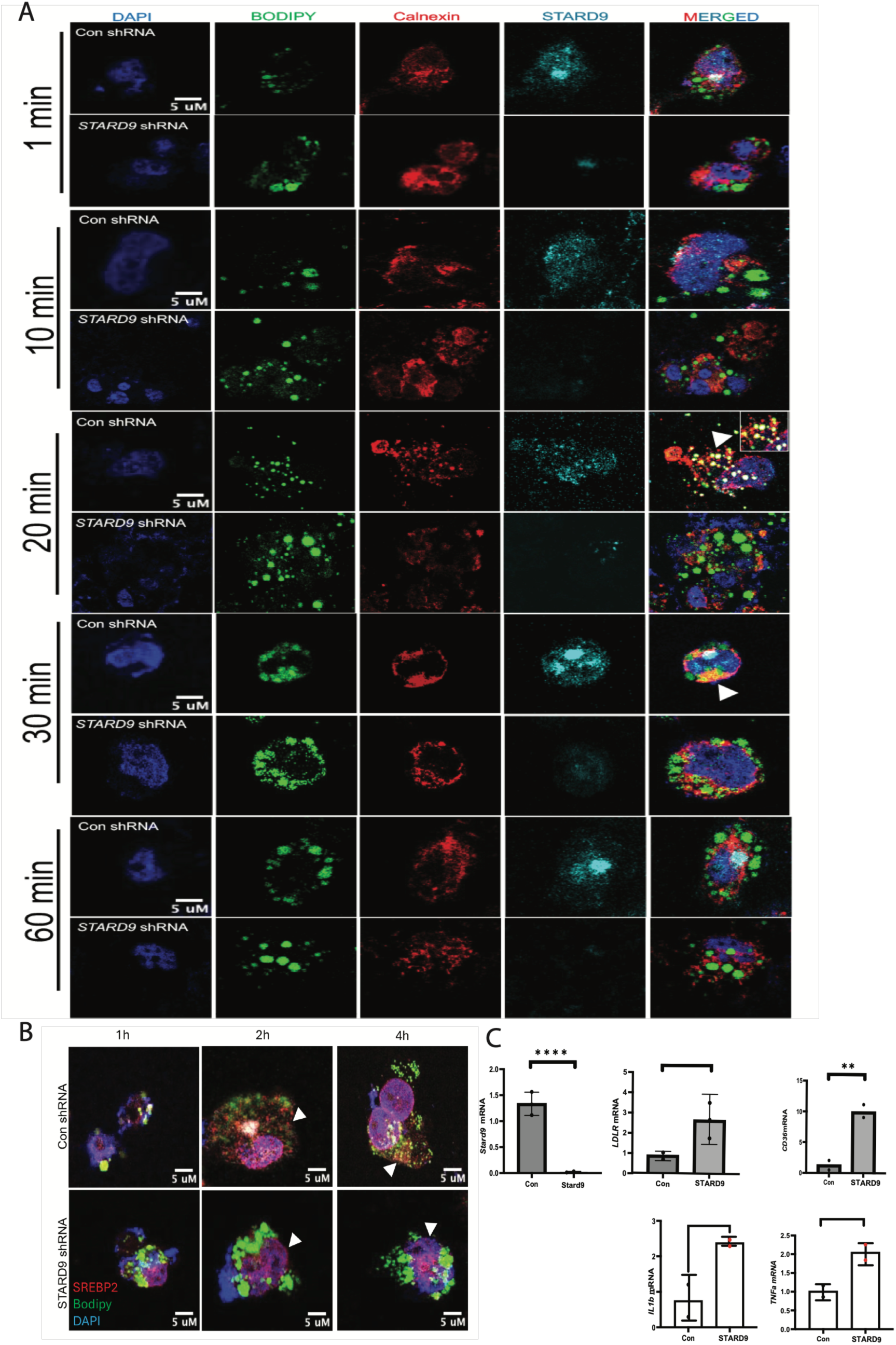
*STARD9* Deficiency Disrupts ER Cholesterol Delivery and Activates Pro-Inflammatory Pathways. (A) Impaired delivery of cholesterol to the ER in *STARD9*-knockdown THP-1 vs. WT cells following oxysterol treatment. (B) Persistent nuclear localization of SREBP in *STARD9*-knockdown cells despite BODIPY-cholesterol treatment. (C) Altered expression of lipid and inflammatory genes in *STARD9*-deficient THP-1 cells: upregulation of SREBP2 (LDLR) and LXR (CD36) targets, and elevated pro-inflammatory cytokines (IL-1β, TNF-α) expression levels by qRT-PCR. Data are presented as mean ± SEM. Each dot represents an independent biological replicate. For comparisons among multiple groups, one-way ANOVA with Holm– Šidák post hoc test was used when data were normally distributed; otherwise, the Mann–Whitney U test was used for non-normal data. *P ≤ 0.05, **P ≤ 0.01, ***P ≤ 0.001, **\*\*\*\***P ≤ 0.0001.

**Suppl. Fig. 3:**
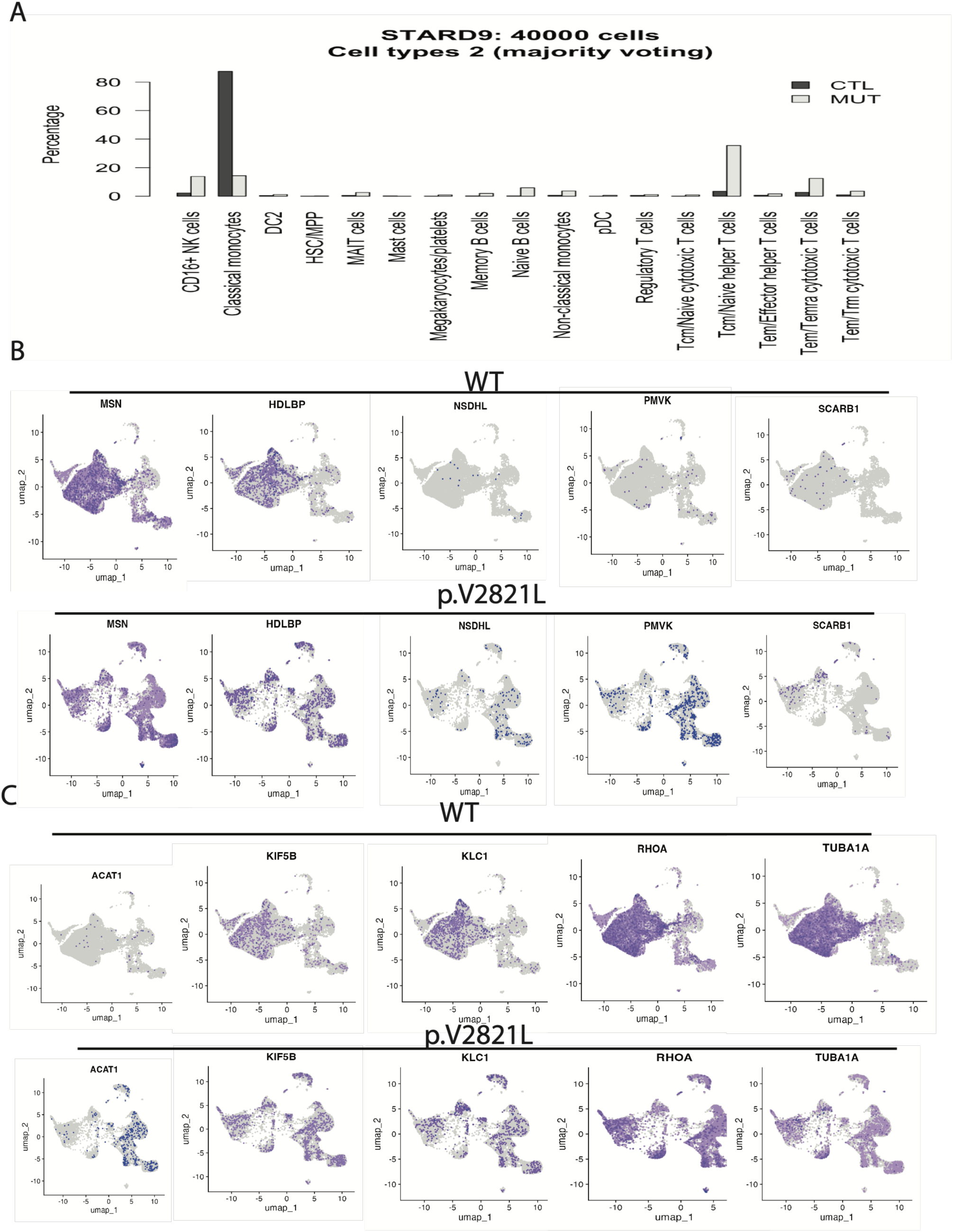
Immune cell composition and lipid-metabolic transcriptional changes in STARD9 p.V2821L carriers. (A) Azimuth-based cell type deconvolution showing altered immune composition in carriers, marked by expansion of CD16⁺ NK cells, TCM/Naive helper T cells, Tem/Temra cytotoxic T cells, with reduced classical CD14⁺CD16⁻ monocytes. (B) Differential expression of lipid and cholesterol-metabolism and (C) microtubule-associated genes across immune subsets.

**Suppl. Fig. 4:**
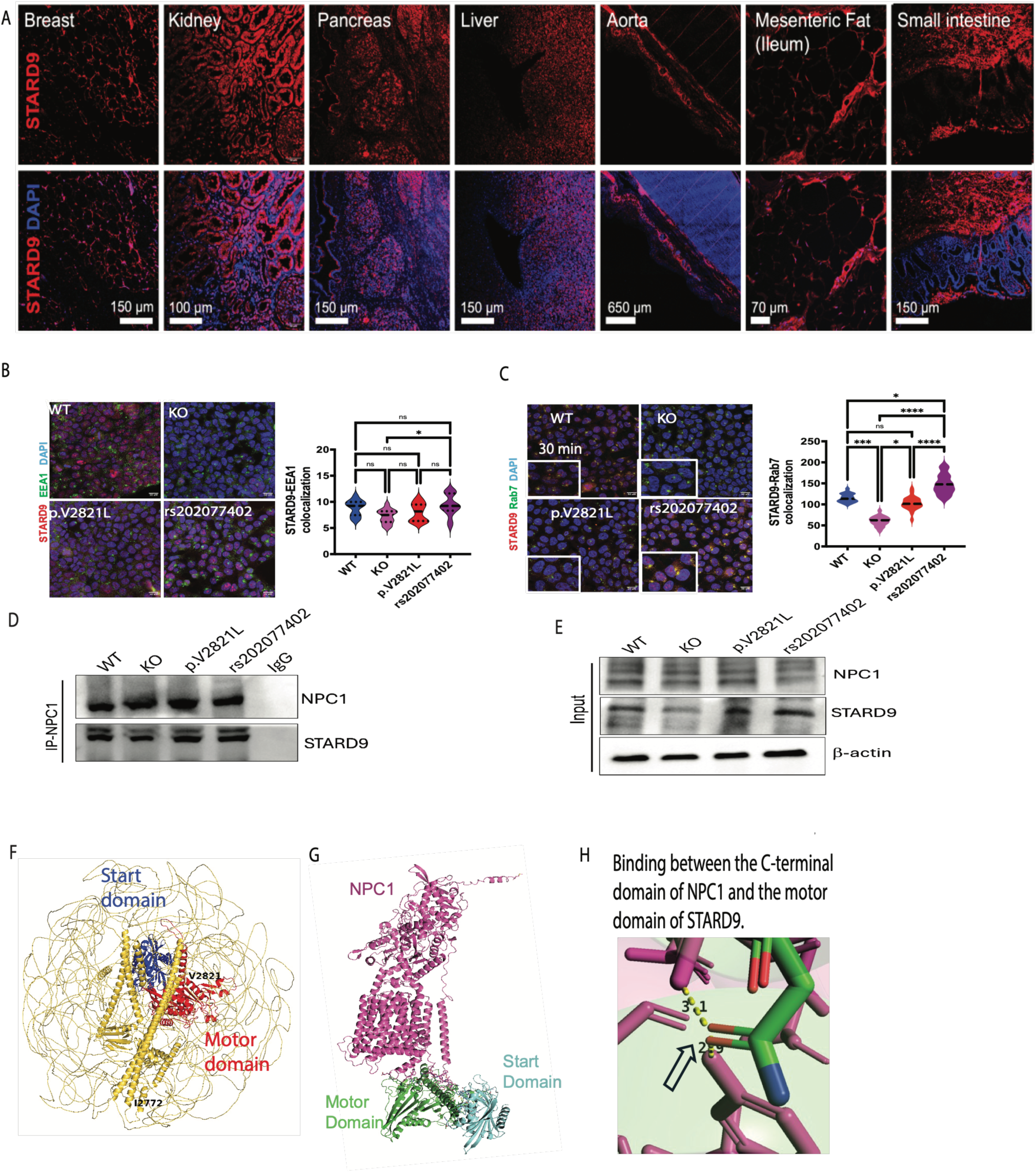
STARD9 localizes to late endosomes, forms a complex with NPC1. (A) STARD9 tissue expression profiles in human datasets, confirming broad expression including breast, kidney, pancreas, liver, aorta, mesenteric adipose tissue, jejunum. (B) colocalization of STARD9 with the early endosomal marker (EEA1) following oxysterol treatment, (C) Colocalization of STARD9 with late endosomal marker Rab7. (D) Co-immunoprecipitation confirming preserved STARD9–NPC1 complex formation across all three mutant lines. (E–F) AlphaFold-Multimer and PyMOL structural modeling of the STARD9–NPC1 complex, illustrating multiple stable binding interfaces characterized by low inter-chain predicted alignment error (PAE < 4 Å) and multiple atomic contacts within 3.5 Å. One interface involves two hydrogen bonds between the NPC1 C-terminus and the motor domain of STARD9. Data are presented as mean ± SEM. Each dot represents an independent biological replicate. *P ≤ 0.05, **P ≤ 0.01, ***P ≤ 0.001, ****P ≤ 0.0001.

**Suppl. Fig. 5:**
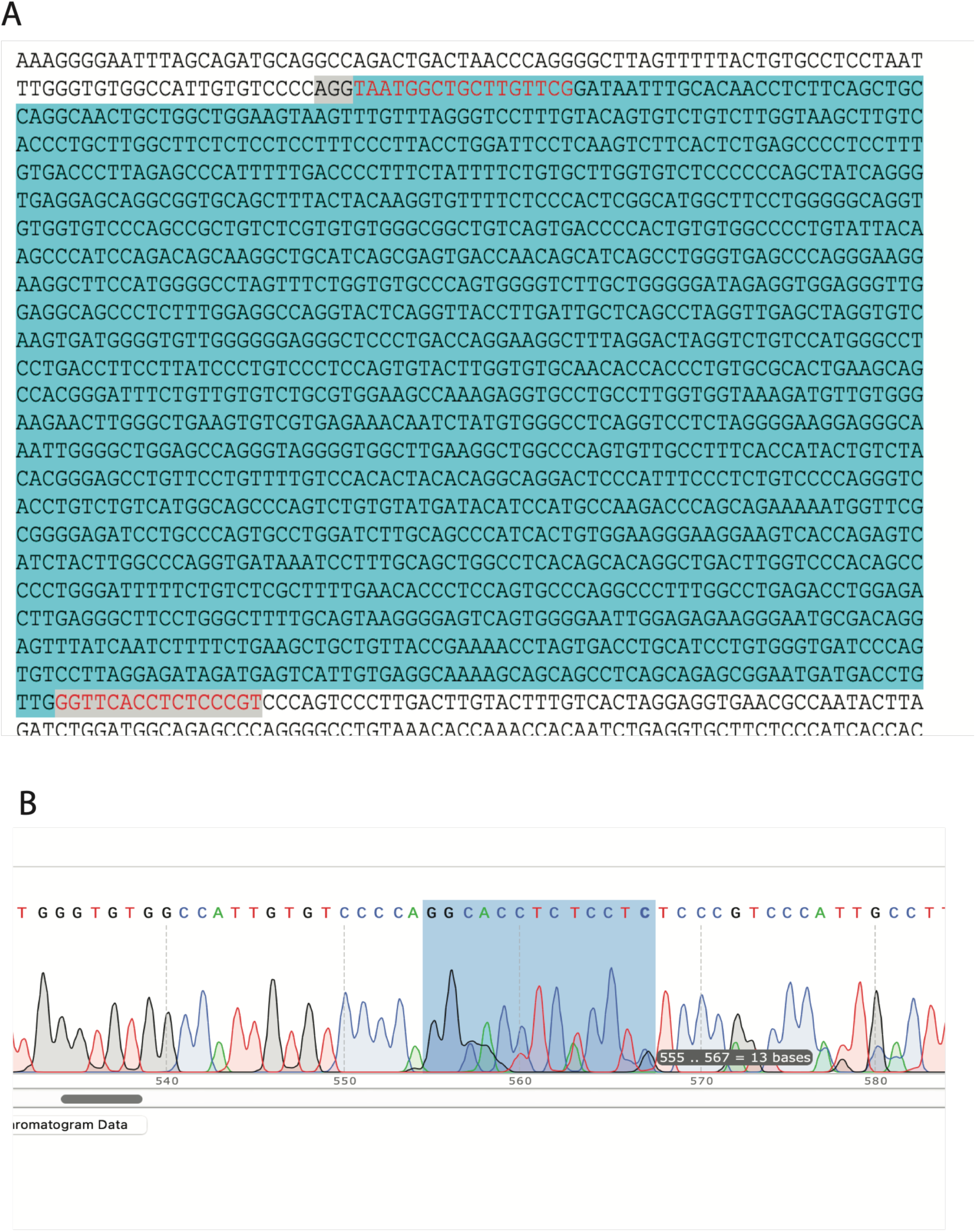
Sequencing of CRISPR/Cas9 mediated START domain deletion of STARD9 in HEK293T cells. (**A**) Highlighted area of nucleotide sequences is the START domain (deleted part) of *STARD9* gene. (B) Sanger sequencing showed homozygous deletion of START domain (blue color).

**Suppl. Fig. 6:**
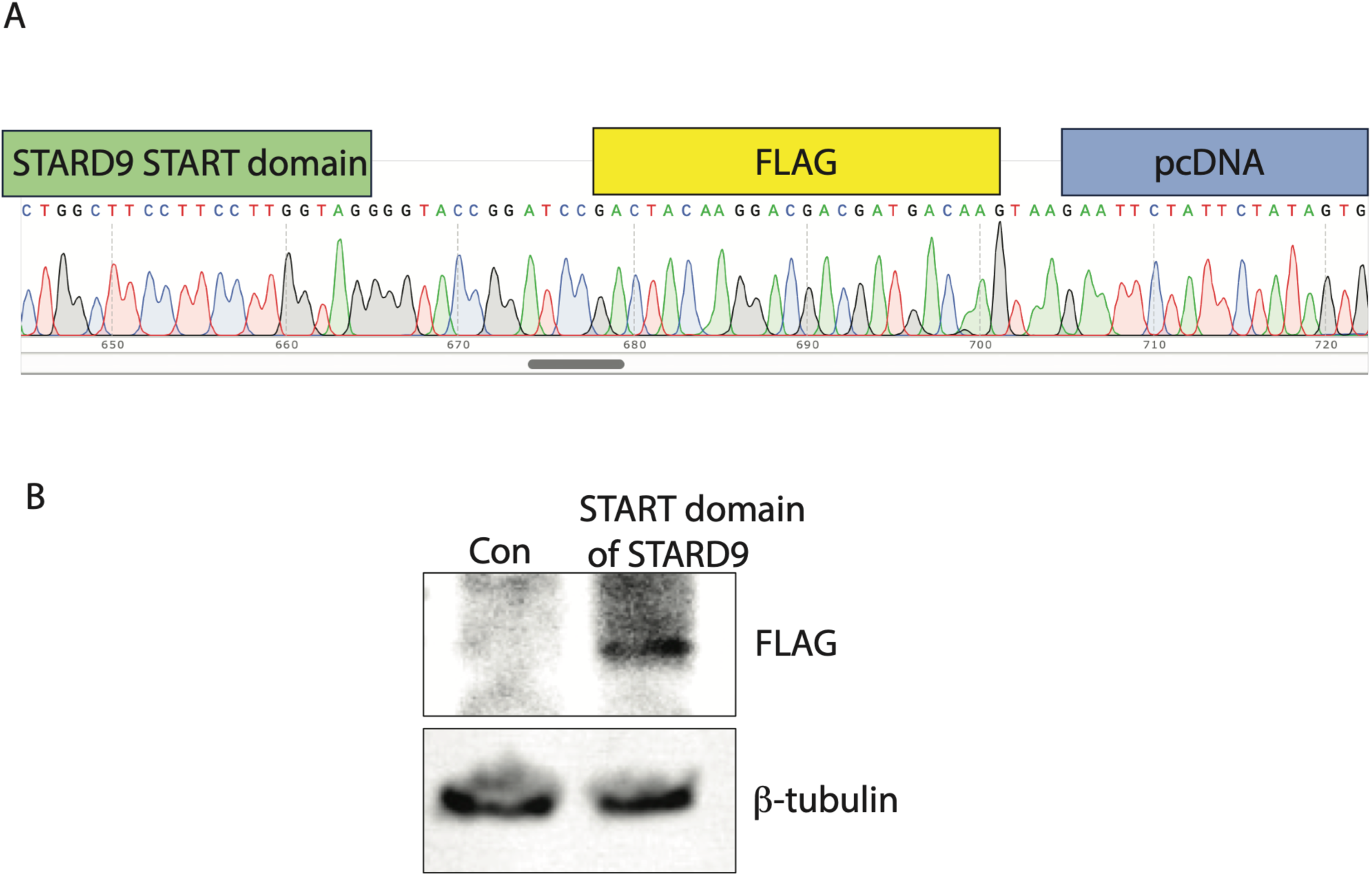
Overexpression of START domain in HEK293T cells by cloning. (A) Sanger sequencing showed START domain cloning in pcDNA-cFLAG plasmid. (B) FLAG protein expression by western blot confirms the positive cloning of START domain in HEK293T cells.

