## Supplementary material for "Variant-Specific Rewiring of Lysosomal Positioning by STARD9 Produces Opposite Cholesterol Trafficking Outcomes": Table 1

| Sample code | sex | Age | Ae of PCI/MI | status | relationship to patient |
| --- | --- | --- | --- | --- | --- |
| A-I | male | Unknown | Unknown | Unknown | Father |
| A-II | female | Unknown | Unknown | Unknown | Mother |
| A-III | male | 64 | 61 | carrier | Index case |
| A-IV | female | 58 | No CAD | noncarrier | spouse |
| A-V | male | 63 | No CAD | noncarrier | unnrelated |
| A-VIII | female | 67 | NA | carrier | sister |
| A-VIII | female | 61 | NA | carrier | sister |
| A-VIII | male | 13 | NA | carrier | son |
| A-IX | male | 41 | NA | carrier | nephew |
| A-X | female | 39 | NA | carrier | niece |
| A-XII | female | 37 | NA | carrier | niece |
| A-XII | female | 32 | NA | noncarrier | niece |
| B-I | female | 63 | <50 | carrier | Mother |
| B-II | male | 82 | NA | Unknown | Father |
| B-III | male | 56 | No CAD | carrier | brother |
| B-IV | female | 42 | 56 | carrier | sister |
| B-V | male | 45 | 41 | carrier | Index case |
| B-VI | female | 67 | 54 | carrier | sister |
| B-VII | male | 47 | 43 | carrier | brother |
| B-VIII | female | 72 | NA | noncarrier | sister |
| C-I | male | Unknown | Unknown | Unknown | father |
| C-II | female | Unknown | Unknown | Unknown | mother |
| C-III | female | 52 | 52 | carrier | index case |
| C-IV | Male | 38 | 36 | carrier | brother |
| C-V | female | 64 | 61 | carrier | sister |
| C-VI | male | 56 | Unknown | Unaffected | Mother |
| C-VII | male | 38 | Unknown | Unaffected | Mother |

| H/O elevated LDL | total cholesterol | LDL | HDL | TG | Notes |
| --- | --- | --- | --- | --- | --- |
|  |  | (mg/dl) | (mg/dl) | (mg/dl) |  |
| Unknown | Unknown | Unknown | Unknown | Unknown | Deceased |
| Unknown | Unknown | Unknown | Unknown | Unknown | Deceased |
| Yes | NA | 208 | 51 | 57 | *On ezetimibe and statin |
| No | Unknown | Unknown | Unknown | Unknown |  |
| No | Unknown | Unknown | Unknown | Unknown |  |
| Yes | 192 | 160* | Unknown | NA | *On statin |
| Yes | 200 | 147* | 48 | NA | *On ezetimibe |
| Yes | NA | 192 | 65 | NA | on Statin |
| Yes | 279 | 199 | 62 | NA | on Statin |
| Yes | 219 | 160 | 67 | NA | on Statin |
| Yes | 183 | 130* | 55 | NA | on Statin |
| No | 192 | 102 | 39 | NA | ? |
| Yes | Unknown | Unknown | Unknown | Unknown | Deceased |
| No | Unknown | Unknown | Unknown | Unknown | Deceased |
| Yes | NA | 188 | 43 | NA | *On statin |
| Yes | NA | 189 | 47 | NA | *On statin |
| Yes | NA | 183 | 53 | 168 | *On statin |
| Yes | NA | 202 | 59 | 145 | *On statin |
| Yes | 276 | 190 | 38 | 180 |  |
| No |  |  |  |  |  |
| Unknown | Unknown | Unknown | Unknown | Unknown | Deceased |
| Unknown | Unknown | Unknown | Unknown | Unknown | Deceased |
| Yes | NA | NA | NA | NA | *On statin |
| Yes | NA | >180 | 37 | NA | deceased |
| Yes | NA | 185 | 49 | NA | *On statin |
| No | 180 | 106 | Unknown | Unknown | *On statin |
| No | 198 | 110 | Unknown | Unknown |  |
