## Supplementary material for "Variant-Specific Rewiring of Lysosomal Positioning by STARD9 Produces Opposite Cholesterol Trafficking Outcomes": Table 2

| NO | Variant | Age at baseline | Sex | CAD/PCI | SBP | DBPYES |
| --- | --- | --- | --- | --- | --- | --- |
| 19 | p.R329L | 69.00 | Male | LAD | 150.00 | 80.00 |
| 18 | p.T826M | 43.00 | Female | RCA | 120.00 | 80.00 |
| 13 | G2455S | 38.00 | Female | RCA | 110.00 | 80.00 |
| 17 | Q2673X | 60.00 | Female | LCX | 137.50 | 80.00 |
| 11 | I2772V | 44.00 | Male | 3VD | 120.00 | 80.00 |
| 15 | S3039R | 53.00 | Male | 3VD | 190.00 | 90.00 |
| 27 | p.G3763fs | 56.00 | Female | LAD,LCX | 150.00 | 100.00 |
| 24 | p.N3950Y | 61.00 | Male | LAD, RC | 140.00 | 90.00 |

| **FBS** | **HDL** | **LDL** | **TG** | **T-Ch** | **waist circumference** |
| --- | --- | --- | --- | --- | --- |
| 64.00 | 33.00 | 70.00 | 208.00 | 164.60 | 96.00 |
| 85.00 | 32.00 | 102.00 | 260.00 | 166.00 | 75.00 |
| 97.00 | 57.00 | 200.00 | 146.00 | 276.20 | 121.00 |
| 121.00 | 55.00 | 218.00 | 256.00 | 314.20 | 104.00 |
| 83.00 | 36.84 | 128.85 | 429.00 | 261.50 | 93.00 |
| 69.00 | 49.00 | 145.00 | 236.00 | 241.20 | 85.00 |
| 78.00 | 37.00 | 87.00 | 307.00 | 205.40 | 104.00 |
| 120.00 | 38.00 | 201.00 | 250.00 | 289.00 | 96.00 |

| Body Mass index | HTN | DM2 | Smoking | Panel Diagnostic |
| --- | --- | --- | --- | --- |
| 20.08 | Yes | 0.00 | past smoker | Unstable angina |
| 19.56 | 0.00 | 0.00 | never smoker | Non Fatal MI |
| 37.42 | 0.00 | 0.00 | never smoker | Non Fatal MI |
| 21.93 | Yes | Yes | never smoker | Unstable angina |
| 29.67 | 0.00 | 0.00 | never smoker | Non Fatal MI |
| 23.13 | Yes | 0.00 | Current smoker | Unstable angina |
| 30.13 | Yes | 0.00 | never smoker | Unstable angina |
| 28.23 | Yes | 0.00 | never smoker | Unstable angina |

| Age | Event | Note |
| --- | --- | --- |
| 79.25 |  |  |
| 53.50 |  |  |
| 46.08 |  |  |
| 65.42 |  |  |
| 47.33 |  | Low CADD, but GWAS locus for TG/HDL |
| 58.67 |  |  |
| 64.33 |  |  |
| 65.50 |  |  |
