## Supplementary material for "Variant-Specific Rewiring of Lysosomal Positioning by STARD9 Produces Opposite Cholesterol Trafficking Outcomes": Table 3

| rsID | CHROM | POS_b37 | REF | ALT | N |
| --- | --- | --- | --- | --- | --- |
| rs62019326 | 15 | 42917387 | G | A | 1236482 |
| rs532800789 | 15 | 42966538 | A | G | 831181 |
| rs538041733 | 15 | 42938029 | C | G | 790122 |
| rs187892402 | 15 | 42994453 | G | A | 746545 |
| rs67391479 | 15 | 43004351 | G | A | 1088086 |
| rs12914570 | 15 | 42970899 | A | G | 1244376 |
| rs2412740 | 15 | 43009448 | T | C | 1244490 |
| rs2280048 | 15 | 43011297 | C | A | 1244492 |
| rs3199486 | 15 | 43012195 | T | C | 1244518 |
| rs12914539 | 15 | 42910624 | C | A | 1244367 |
| rs1995939 | 15 | 43012517 | A | G | 1244518 |
| rs62019322 | 15 | 42897374 | T | C | 1244539 |
| rs12912064 | 15 | 43004170 | C | T | 1244499 |
| rs62019320 | 15 | 42895389 | C | T | 1244540 |
| rs17767270 | 15 | 42898612 | C | T | 1244542 |
| rs17767264 | 15 | 42886656 | C | T | 1244541 |
| rs12102055 | 15 | 42997546 | T | A | 1244499 |
| rs28562951 | 15 | 42998365 | T | C | 1244500 |
| rs2136900 | 15 | 42993105 | A | G | 1244498 |
| rs79044457 | 15 | 42895208 | G | T | 1244516 |
| rs9269 | 15 | 43012274 | A | G | 1244519 |
| rs11070375 | 15 | 42870587 | A | G | 1244450 |
| rs78366241 | 15 | 42895209 | C | T | 1244518 |
| rs28479391 | 15 | 42963400 | A | G | 1244435 |
| rs9919994 | 15 | 43011537 | G | T | 1244490 |
| rs7169202 | 15 | 42973053 | G | C | 1244462 |
| rs112840248 | 15 | 42950885 | C | T | 1244426 |
| rs8043472 | 15 | 42948862 | A | G | 1244412 |
| rs4924690 | 15 | 42950219 | C | G | 1244426 |
| rs7163030 | 15 | 42971620 | T | C | 1244456 |
| rs16957020 | 15 | 42964426 | T | C | 1244447 |
| rs12910538 | 15 | 42910357 | G | C | 1244350 |
| rs6493059 | 15 | 42977526 | A | C | 1244454 |
| rs28823029 | 15 | 42952109 | G | C | 1244447 |
| rs10400824 | 15 | 42951277 | C | T | 1244451 |
| rs1197551 | 15 | 42931717 | T | A | 1230260 |
| rs28811114 | 15 | 42952112 | C | T | 1244452 |
| rs9920163 | 15 | 42942381 | G | T | 1244439 |
| rs28798437 | 15 | 42952005 | A | C | 1244450 |
| rs4924691 | 15 | 42950237 | T | C | 1244443 |
| rs10162939 | 15 | 42930777 | A | T | 1230262 |
| rs56172155 | 15 | 42882182 | C | T | 1059633 |
| rs1197549 | 15 | 42929154 | C | T | 1230251 |
| rs1197548 | 15 | 42928764 | G | A | 1230250 |
| rs623271 | 15 | 42928208 | G | A | 1231937 |
| rs1212814 | 15 | 42933603 | C | G | 1230258 |
| rs3099758 | 15 | 42927984 | C | T | 1230249 |
| rs3106286 | 15 | 42927717 | T | C | 1230243 |
| rs28744617 | 15 | 42981022 | A | G | 1244404 |
| rs1705356 | 15 | 42922656 | C | T | 1230222 |
| rs3106285 | 15 | 42927668 | T | C | 1230245 |

|  |  |  |  |  |
| --- | --- | --- | --- | --- |
| rs398039432 | 15 | 43000191 G | GA | 1037153 |
| rs550777038 | 15 | 42969019 C | T | 1216997 |
| rs60284149 | 15 | 42870823 GT | G | 1028875 |
| rs34868427 | 15 | 42995565 GT | G | 1037110 |
| rs2136901 | 15 | 42993266 G | A | 1244517 |
| rs7182479 | 15 | 42999565 G | A | 1244517 |
| rs202077402 | 15 | 42982090 A | G | 1225248 |
| rs35616173 | 15 | 43001799 T | C | 1244518 |
| rs34191150 | 15 | 43005718 A | G | 1244515 |
| rs12905592 | 15 | 43002536 G | A | 1244517 |
| rs6493061 | 15 | 42988412 C | A | 1244511 |
| rs9944248 | 15 | 42965191 A | G | 1244471 |
| rs7166373 | 15 | 42954888 C | T | 1244471 |
| rs4923950 | 15 | 42959743 A | G | 1244474 |
| rs7176860 | 15 | 42998858 G | A | 1244519 |
| rs12905187 | 15 | 43002534 A | T | 1244520 |
| rs35738819 | 15 | 42966288 G | A | 1244469 |
| rs4924694 | 15 | 42991066 T | A | 1244516 |
| rs6493060 | 15 | 42977676 G | A | 1217186 |
| rs2136899 | 15 | 42990386 A | G | 1244513 |
| rs12903762 | 15 | 43006385 C | A | 1226082 |
| rs8030587 | 15 | 42981806 G | A | 1244488 |
| rs7183487 | 15 | 42999553 T | A | 1244514 |
| rs58938096 | 15 | 43005804 G | T | 1244514 |
| rs3742993 | 15 | 42983923 A | G | 1244498 |
| rs34603230 | 15 | 42958216 A | G | 1244476 |
| rs8024902 | 15 | 42957532 T | G | 1244491 |
| rs4545789 | 15 | 42958928 A | C | 1244489 |
| rs2175370 | 15 | 42960322 G | A | 1244491 |
| rs8039711 | 15 | 42957180 G | A | 1244491 |
| rs8031218 | 15 | 42982340 C | T | 1244494 |
| rs57298545 | 15 | 42956620 A | G | 1244486 |
| rs938047 | 15 | 42955967 A | G | 1244486 |
| rs12902830 | 15 | 42997382 C | T | 1244520 |
| rs4924692 | 15 | 42950297 G | A | 1244482 |
| rs8039765 | 15 | 42957373 C | A | 1244489 |
| rs1058846 | 15 | 43012024 C | T | 1244539 |
| rs1197550 | 15 | 42930891 T | A | 994566 |
| rs61122145 | 15 | 42991887 G | A | 1244518 |
| rs28664781 | 15 | 42953809 T | G | 1210383 |
| rs672054 | 15 | 42918017 C | T | 1230237 |
| rs12911585 | 15 | 42934545 T | C | 1244416 |
| rs34070722 | 15 | 42938962 G | A | 1244393 |
| rs71474496 | 15 | 42918141 A | G | 1244368 |
| rs398043203 | 15 | 42933807 CT | C | 1042115 |
| rs28882725 | 15 | 42924027 A | T | 1207782 |
| rs11638835 | 15 | 42918341 G | A | 1230246 |
| rs67470909 | 15 | 43010435 T | C | 1244448 |
| rs12594855 | 15 | 42939904 G | A | 1214876 |
| rs28498538 | 15 | 43011353 G | C | 1239870 |
| rs1018744820 | 15 | 42918643 T | C | 554145 |
| rs111810960 | 15 | 43007276 CA | C | 1023404 |

|  |  |  |  |  |
| --- | --- | --- | --- | --- |
| rs61683241 | 15 | 43007718 A | AT | 1029405 |
| rs34117505 | 15 | 42925825 C | T | 1165809 |
| rs28650274 | 15 | 42944829 C | T | 1239878 |
| rs2241998 | 15 | 42869028 G | A | 1230268 |
| rs35392023 | 15 | 43001660 G | A | 1085327 |
| rs2136903 | 15 | 42941469 A | G | 1089029 |
| rs1667493 | 15 | 42889170 T | C | 1223180 |
| rs35006192 | 15 | 42935698 G | A | 1244397 |
| rs6493053 | 15 | 42886074 A | G | 1232114 |
| rs7168835 | 15 | 42876014 C | T | 1232111 |
| rs9783683 | 15 | 42875276 A | C | 1230380 |
| rs114824872 | 15 | 42920562 G | A | 850560 |
| rs12903726 | 15 | 42954739 A | C | 1237692 |
| rs530032660 | 15 | 42952212 G | C | 784285 |
| rs35915660 | 15 | 42873866 TCAG | T | 1041002 |
| rs61750788 | 15 | 42953382 T | G | 1239880 |
| rs12903416 | 15 | 42881039 T | C | 1077634 |
| rs143379946 | 15 | 42974907 C | T | 1229231 |
| rs559353595 | 15 | 43009409 G | A | 369744 |
| rs12901400 | 15 | 42942262 C | T | 1239866 |
| rs548861293 | 15 | 42918759 C | T | 366390 |
| rs67741694 | 15 | 42940334 A | T | 1239850 |
| rs117066630 | 15 | 42953718 T | C | 1239862 |
| rs55959547 | 15 | 42976518 A | G | 1228580 |
| rs180854628 | 15 | 42988525 C | A | 365285 |
| rs12904592 | 15 | 42890198 G | A | 1025481 |
| rs145113087 | 15 | 43005947 G | C | 803846 |
| rs12594951 | 15 | 42934631 A | C | 1195404 |
| rs117954128 | 15 | 42974055 G | A | 1239866 |
| rs569220966 | 15 | 42943178 A | G | 370638 |
| rs74345837 | 15 | 43005264 T | C | 1232852 |
| rs28431056 | 15 | 43010138 C | T | 1188995 |
| rs73410940 | 15 | 43010574 G | A | 1188996 |
| rs186697829 | 15 | 42878575 C | A | 1009646 |
| rs191917476 | 15 | 42988641 G | A | 477755 |
| rs556780600 | 15 | 42925880 C | T | 584480 |
| rs12591415 | 15 | 43009399 C | T | 1187363 |
| rs150746990 | 15 | 42932515 A | T | 925169 |
| rs542649755 | 15 | 42995728 AT | A | 706841 |
| rs143805766 | 15 | 43009642 G | A | 815065 |
| rs538352673 | 15 | 42980127 A | G | 360811 |
| rs537835505 | 15 | 42915072 G | A | 366231 |
| rs185294220 | 15 | 43001221 G | C | 1212629 |
| rs557974103 | 15 | 42955918 T | G | 365285 |
| rs1197547 | 15 | 42905340 T | G | 1244439 |
| rs111730580 | 15 | 42907211 AT | A | 640154 |
| rs368247671 | 15 | 42870824 T | G | 774072 |
| rs11632169 | 15 | 42903115 A | G | 1235530 |
| rs1197546 | 15 | 42902246 G | A | 1235532 |
| rs528091580 | 15 | 42925476 AG | A | 425688 |
| rs376095487 | 15 | 42953467 C | T | 230552 |
| rs544507849 | 15 | 42999666 CT | C | 1001834 |

|  |  |  |  |  |
| --- | --- | --- | --- | --- |
| rs4447398 | 15 | 42904904 C | A | 1235526 |
| rs565991513 | 15 | 42909088 CA | C | 922779 |
| rs764967488 | 15 | 42949280 C | T | 460197 |
| NA | 15 | 42948935 C | T | 376296 |
| rs56801471 | 15 | 42896146 A | G | 1185620 |
| rs577581215 | 15 | 42885609 T | C | 544906 |
| rs561421938 | 15 | 42937189 T | A | 577971 |
| rs535068320 | 15 | 42949070 G | T | 577970 |
| rs535127983 | 15 | 42889684 G | A | 54951 |
| rs150192451 | 15 | 42931427 C | G | 566539 |
| rs563833340 | 15 | 42977303 G | A | 432332 |
| rs916413970 | 15 | 42946588 A | G | 420554 |
| rs573466110 | 15 | 42933777 A | G | 369062 |
| rs545253441 | 15 | 42970464 C | T | 66360 |
| rs567894069 | 15 | 43011180 T | A | 401914 |
| rs146582932 | 15 | 42897448 G | T | 1228797 |
| rs913429324 | 15 | 42999523 G | A | 27798 |
| rs147950526 | 15 | 42913198 C | T | 678996 |
| rs188508115 | 15 | 43003063 A | C | 668014 |
| rs141297638 | 15 | 42893808 G | A | 508989 |
| rs192196303 | 15 | 42909620 G | A | 986940 |
| rs529710891 | 15 | 42935792 G | T | 868413 |
| rs55773330 | 15 | 42956987 AAC | A | 1017802 |
| rs746547270 | 15 | 42897884 G | A | 599403 |
| rs181167607 | 15 | 42906448 G | T | 1215957 |
| rs550441855 | 15 | 42956924 A | G | 710223 |
| rs150402119 | 15 | 42894680 G | C | 972019 |
| rs569388978 | 15 | 42953642 C | T | 366232 |
| rs117100467 | 15 | 43008831 G | A | 1223712 |
| rs556362354 | 15 | 42918499 C | T | 379016 |
| rs112165067 | 15 | 42906924 G | A | 1186559 |
| rs202074007 | 15 | 42967130 G | T | 1202654 |
| rs564244442 | 15 | 43008093 C | CTGG | 357680 |
| rs544930217 | 15 | 42991793 A | G | 818028 |
| rs138641635 | 15 | 42998558 A | C | 1228232 |
| rs868042277 | 15 | 43008884 A | G | 6702 |
| NA | 15 | 43007040 G | A | 6702 |
| rs559872238 | 15 | 42900330 G | C | 577971 |
| rs191282954 | 15 | 42915084 G | T | 371320 |
| rs970534725 | 15 | 43012893 T | G | 382447 |
| rs557984924 | 15 | 42869457 T | C | 365285 |
| rs113206636 | 15 | 43010551 G | GGGGA | 1037160 |
| rs149873427 | 15 | 42893027 C | T | 679810 |
| rs117112275 | 15 | 42887565 T | C | 1230405 |
| rs543357875 | 15 | 42975078 C | T | 764676 |
| rs569786942 | 15 | 42875501 T | C | 708760 |
| rs565472380 | 15 | 42979114 T | C | 378928 |
| rs142114926 | 15 | 42991816 T | C | 1222378 |
| rs546585224 | 15 | 42979250 C | T | 378508 |
| rs568003335 | 15 | 42894191 C | G | 967107 |
| rs79313070 | 15 | 43002191 G | T | 1167801 |
| rs138157695 | 15 | 42993425 G | A | 497976 |

|  |  |  |  |  |
| --- | --- | --- | --- | --- |
| rs76175552 | 15 | 42887079 A | T | 1225070 |
| rs74475031 | 15 | 42979146 A | G | 1175281 |
| rs147473758 | 15 | 42883617 C | T | 1163365 |
| rs184219872 | 15 | 42988314 G | A | 589136 |
| NA | 15 | 42954809 C | A | 513928 |
| rs184122994 | 15 | 42990730 T | G | 589136 |
| rs542299362 | 15 | 42953963 A | C | 31575 |
| rs544743742 | 15 | 42938738 A | T | 370110 |
| rs77080857 | 15 | 42928937 T | C | 756917 |
| rs546203561 | 15 | 42923268 C | A | 577971 |
| rs561457101 | 15 | 43011558 A | G | 421885 |
| rs180867650 | 15 | 43008642 C | A | 1211504 |
| rs764650416 | 15 | 42993496 G | T | 514463 |
| rs931528965 | 15 | 42918444 T | C | 509907 |
| rs145400823 | 15 | 42986274 C | T | 374849 |
| rs77298618 | 15 | 42876902 T | G | 1238398 |
| rs548121692 | 15 | 42915911 G | A | 369744 |
| rs762517984 | 15 | 42945917 C | A | 428483 |
| rs138819618 | 15 | 42891380 C | T | 1222743 |
| rs534725679 | 15 | 42985477 A | G | 369871 |
| NA | 15 | 42903471 G | A | 363054 |
| rs397722507 | 15 | 42953149 GT | G | 357298 |
| rs537604330 | 15 | 43003401 G | A | 357298 |
| rs555394489 | 15 | 42895336 C | A | 63184 |
| rs187995678 | 15 | 42990326 C | T | 526744 |
| rs1056030401 | 15 | 42917567 C | T | 551926 |
| rs4923949 | 15 | 42892748 A | G | 1014824 |
| rs571444516 | 15 | 42879849 C | G | 604731 |
| rs564955678 | 15 | 42965237 G | A | 361757 |
| rs550730817 | 15 | 42964752 G | A | 18745 |
| NA | 15 | 42918707 T | C | 382447 |
| rs11070376 | 15 | 42879757 T | G | 1230401 |
| rs138147686 | 15 | 42911919 G | A | 581748 |
| rs769089032 | 15 | 42910077 C | T | 448839 |
| rs574691968 | 15 | 42935706 T | TA | 613259 |
| rs1044735653 | 15 | 43005905 C | T | 586831 |
| rs12904010 | 15 | 42881210 A | G | 1230399 |
| rs540950607 | 15 | 42952786 A | C | 259077 |
| rs191874576 | 15 | 42881098 C | T | 370110 |
| rs544111850 | 15 | 42920659 T | C | 534733 |
| rs114786341 | 15 | 42953446 G | A | 259077 |
| rs756135004 | 15 | 42904832 G | C | 464867 |
| rs780449628 | 15 | 42944948 A | C | 628242 |
| rs116577053 | 15 | 42941664 C | G | 259077 |
| rs759558730 | 15 | 42978435 A | C | 399382 |
| rs187388117 | 15 | 43011625 C | T | 901950 |
| rs141270837 | 15 | 42998672 A | G | 1190573 |
| rs370116985 | 15 | 42918624 T | C | 48795 |
| rs1024511261 | 15 | 42942332 A | T | 409148 |
| rs142591139 | 15 | 42988901 G | A | 1228461 |
| rs557266111 | 15 | 42996335 C | G | 894841 |
| rs147330525 | 15 | 42999707 G | A | 749124 |

|  |  |  |  |  |
| --- | --- | --- | --- | --- |
| rs11399920 | 15 | 42917845 AT | A | 1009087 |
| rs559676742 | 15 | 42994563 G | A | 756917 |
| rs77003671 | 15 | 42896234 T | C | 1238397 |
| rs779017439 | 15 | 42970028 G | A | 398402 |
| rs559750005 | 15 | 42893677 G | A | 619607 |
| rs1007016522 | 15 | 42910442 T | A | 382447 |
| rs570992508 | 15 | 42941522 G | A | 227674 |
| rs113910756 | 15 | 42965461 G | A | 577971 |
| rs564597643 | 15 | 42930718 CA | C | 590415 |
| rs539428789 | 15 | 42946624 G | T | 230552 |
| rs560093737 | 15 | 42979758 T | G | 370638 |
| rs577245802 | 15 | 42901367 AAG | A | 368189 |
| rs201119685 | 15 | 42994073 T | TC | 582430 |
| rs12442544 | 15 | 42995418 C | G | 681890 |
| rs1050331778 | 15 | 42891501 C | T | 375575 |
| rs533443278 | 15 | 42994547 G | A | 601143 |
| rs185196126 | 15 | 42965455 C | T | 253475 |
| rs3742994 | 15 | 42983742 C | A | 391999 |
| rs574547539 | 15 | 42920203 T | G | 719652 |
| rs572690483 | 15 | 42880406 A | G | 635769 |
| rs563158318 | 15 | 42932912 G | A | 360129 |
| rs549319649 | 15 | 42967398 C | G | 977091 |
| rs543084498 | 15 | 42990117 T | A | 468622 |
| rs751457103 | 15 | 42931670 A | G | 697348 |
| rs548604256 | 15 | 42948022 A | G | 298985 |
| rs192615557 | 15 | 42954183 G | T | 360129 |
| rs567867043 | 15 | 42964780 C | T | 372647 |
| rs187216270 | 15 | 42904022 A | T | 624730 |
| rs111612014 | 15 | 42874007 A | G | 647306 |
| rs539858496 | 15 | 42904674 C | T | 602609 |
| rs184215065 | 15 | 43008342 G | C | 361757 |
| rs777870405 | 15 | 42993947 C | A | 590365 |
| rs114539185 | 15 | 42914152 T | G | 599106 |
| rs398027029 | 15 | 42870304 A | AT | 1017141 |
| rs768035777 | 15 | 42919268 G | A | 426067 |
| rs868202680 | 15 | 42868358 C | T | 469886 |
| rs538472001 | 15 | 42918459 TTCCTCTTCCTC T |  | 888013 |
| rs569851918 | 15 | 42988920 GGAGA | G | 360129 |
| rs1042575965 | 15 | 42956237 G | C | 540202 |
| rs189193514 | 15 | 42928407 T | C | 785433 |
| rs551025057 | 15 | 42902777 A | G | 422095 |
| rs563451455 | 15 | 42968307 A | G | 769500 |
| rs563742211 | 15 | 42992049 C | T | 773558 |
| rs553736195 | 15 | 43011409 A | C | 360811 |
| rs78994161 | 15 | 42975022 C | A | 1174333 |
| rs756943396 | 15 | 42947980 T | A | 468941 |
| rs202017657 | 15 | 42981101 C | G | 1171254 |
| rs542874709 | 15 | 42949334 C | T | 570051 |
| rs144271727 | 15 | 42876084 T | G | 416928 |
| rs534940697 | 15 | 42966921 G | GTA | 362315 |
| rs557130886 | 15 | 42987901 G | T | 716651 |
| rs201416428 | 15 | 42890742 A | AT | 581748 |

|  |  |  |  |  |
| --- | --- | --- | --- | --- |
| rs148181870 | 15 | 42911523 A | G | 1206581 |
| rs117478364 | 15 | 42959003 A | T | 1203248 |
| rs1017461242 | 15 | 42950058 C | A | 526743 |
| rs116737895 | 15 | 42986042 C | T | 526743 |
| rs742444935 | 15 | 42871706 A | G | 434684 |
| rs528882270 | 15 | 42932699 C | G | 548103 |
| NA | 15 | 42913750 A | ATACAGAAAAC/ | 29487 |
| rs1814518 | 15 | 42873672 G | A | 1230419 |
| rs777242940 | 15 | 42944170 CTG | C | 425687 |
| rs181391693 | 15 | 42923116 A | G | 450180 |
| rs767904319 | 15 | 42941881 C | G | 627628 |
| rs577253212 | 15 | 42922291 T | C | 379474 |
| rs537314110 | 15 | 42954801 C | T | 533662 |
| rs778329983 | 15 | 43009275 C | T | 398741 |
| rs778682289 | 15 | 42927059 C | T | 512001 |
| rs539519966 | 15 | 42918498 C | T | 379016 |
| rs552741284 | 15 | 43010039 T | A | 373521 |
| rs535171426 | 15 | 42946687 C | A | 365285 |
| rs566366165 | 15 | 42929197 G | A | 796804 |
| rs117143870 | 15 | 42938044 C | T | 1225946 |
| rs535667563 | 15 | 42918555 C | T | 772557 |
| rs145525359 | 15 | 42911771 G | A | 1206857 |
| NA | 15 | 42953936 T | A | 363282 |
| rs778181709 | 15 | 42999548 A | G | 509421 |
| rs542175917 | 15 | 42916758 T | C | 794909 |
| rs141192915 | 15 | 42897985 C | G | 365285 |
| rs183904150 | 15 | 43011179 G | A | 1228569 |
| rs145054253 | 15 | 42932805 G | A | 242754 |
| rs542927420 | 15 | 42971913 A | G | 726418 |
| rs140243268 | 15 | 43000117 A | G | 1231296 |
| rs180869067 | 15 | 42877641 G | T | 448723 |
| rs541890756 | 15 | 42931425 G | A | 582047 |
| rs148405956 | 15 | 43008230 T | C | 829750 |
| rs61750914 | 15 | 42984825 C | T | 714180 |
| rs77876077 | 15 | 42869977 G | C | 769279 |
| rs534886977 | 15 | 42910451 C | T | 365285 |
| rs78617381 | 15 | 42912689 A | T | 400447 |
| NA | 15 | 42902586 A | G | 376277 |
| rs150051531 | 15 | 42952133 T | C | 630579 |
| rs565298 | 15 | 42893192 C | A | 1235607 |
| rs550992626 | 15 | 42900445 A | G | 596300 |
| NA | 15 | 42991741 G | A | 94193 |
| rs571385368 | 15 | 42872500 T | TC | 754869 |
| rs187191059 | 15 | 42941341 C | T | 425687 |
| rs547585065 | 15 | 42920939 T | C | 843699 |
| NA | 15 | 42988850 G | A | 20757 |
| rs140771911 | 15 | 42882036 C | T | 771193 |
| rs537743332 | 15 | 42886644 C | T | 366231 |
| rs150686043 | 15 | 42921661 A | G | 365285 |
| rs563033349 | 15 | 42959907 T | C | 365285 |
| rs765034095 | 15 | 42997643 T | G | 398727 |
| rs12594724 | 15 | 42999650 A | G | 1205143 |

|  |  |  |  |  |
| --- | --- | --- | --- | --- |
| rs182845723 | 15 | 42924574 T | C | 1159162 |
| rs758857318 | 15 | 42989972 T | A | 542504 |
| rs4334272 | 15 | 42994169 T | C | 1205143 |
| rs778633745 | 15 | 42874075 A | G | 426323 |
| rs549395191 | 15 | 42911679 G | A | 529921 |
| rs149176049 | 15 | 42982891 A | C | 1202816 |
| rs150784222 | 15 | 42985224 A | G | 365285 |
| rs117015041 | 15 | 42912674 T | A | 655665 |
| rs754291307 | 15 | 42894385 A | T | 421306 |
| rs182260499 | 15 | 42956967 T | C | 685370 |
| rs113915421 | 15 | 42991684 C | T | 1237844 |
| rs12594837 | 15 | 42977290 C | T | 1231293 |
| NA | 15 | 43002106 T | C | 48826 |
| rs566288410 | 15 | 42892849 G | A | 365286 |
| rs201349756 | 15 | 42978424 C | T | 422304 |
| rs111336295 | 15 | 42984179 C | G | 245757 |
| rs530299717 | 15 | 42922023 A | C | 365285 |
| rs147801000 | 15 | 42963833 AC | A | 577971 |
| rs28680600 | 15 | 42979105 A | G | 1234629 |
| rs528267195 | 15 | 42990112 C | T | 808892 |
| rs563268103 | 15 | 42925722 A | G | 360811 |
| rs8023743 | 15 | 42957542 C | G | 1235457 |
| rs11637248 | 15 | 42882016 G | C | 1231284 |
| rs6493054 | 15 | 42886504 A | G | 1235611 |
| rs1705360 | 15 | 42889047 A | G | 1235607 |
| rs185487313 | 15 | 42905168 C | A | 809693 |
| rs16957061 | 15 | 42984182 A | G | 1233937 |
| rs766671995 | 15 | 42966902 C | T | 415113 |
| rs1052463329 | 15 | 42930144 A | G | 379922 |
| rs112210775 | 15 | 42939130 G | A | 245757 |
| rs73410931 | 15 | 43003939 G | C | 1234631 |
| rs116719583 | 15 | 42937465 T | G | 648191 |
| rs766573983 | 15 | 42941572 A | G | 551729 |
| rs143465521 | 15 | 42997507 G | A | 373521 |
| rs78261202 | 15 | 43003113 G | C | 715112 |
| rs143155792 | 15 | 42921007 C | T | 230552 |
| rs111607613 | 15 | 43012888 G | A | 241298 |
| rs144580936 | 15 | 42870421 G | A | 549335 |
| rs60723187 | 15 | 42997347 C | T | 1234632 |
| rs73410923 | 15 | 42997586 C | T | 1234632 |
| rs551035769 | 15 | 42937527 G | A | 582857 |
| rs73408619 | 15 | 42949817 G | C | 1235458 |
| NA | 15 | 43006385 C | T | 463254 |
| rs73410937 | 15 | 43006957 T | G | 1234630 |
| rs150250448 | 15 | 42878578 G | T | 764296 |
| rs150454654 | 15 | 42898683 G | A | 240083 |
| rs192352355 | 15 | 42959921 C | T | 1044312 |
| rs12594725 | 15 | 42999641 C | G | 1234632 |
| rs75988797 | 15 | 43001332 G | A | 1234632 |
| rs1197545 | 15 | 42893485 A | G | 1068699 |
| rs567681136 | 15 | 42913727 T | C | 22048 |
| rs12594696 | 15 | 42999540 A | G | 1234631 |

|  |  |  |  |  |
| --- | --- | --- | --- | --- |
| rs73410927 | 15 | 43000977 A | G | 1234631 |
| rs768060909 | 15 | 42962501 A | G | 534677 |
| rs549644430 | 15 | 43005010 G | A | 997531 |
| rs571684887 | 15 | 42956687 G | A | 381504 |
| rs3742992 | 15 | 42983943 C | G | 1233938 |
| rs549133434 | 15 | 42960894 G | A | 603036 |
| rs576444632 | 15 | 42923161 C | T | 496209 |
| rs530272524 | 15 | 42873680 A | G | 276985 |
| rs530835353 | 15 | 42919320 T | C | 357298 |
| rs560029646 | 15 | 43003736 T | C | 630428 |
| rs557162342 | 15 | 42888744 G | A | 1024341 |
| rs115420503 | 15 | 42909687 A | T | 1060842 |
| rs7178483 | 15 | 42956626 A | G | 1235457 |
| rs370894209 | 15 | 42918444 TTCTTC | T | 694850 |
| rs58237318 | 15 | 42984858 G | T | 1233939 |
| rs755438104 | 15 | 42936547 C | T | 467372 |
| rs9944270 | 15 | 42962209 A | T | 1235457 |
| rs12440556 | 15 | 42874506 C | G | 881207 |
| rs8028699 | 15 | 42993469 T | C | 1234631 |
| rs75442140 | 15 | 42868643 T | A | 250556 |
| rs8039881 | 15 | 42935849 A | G | 1233049 |
| rs8043143 | 15 | 42948674 A | G | 1235457 |
| rs16957063 | 15 | 42984288 A | G | 1233938 |
| rs149462255 | 15 | 42903026 T | G | 232613 |
| rs114191253 | 15 | 42924026 T | A | 1179773 |
| rs143692879 | 15 | 42977628 T | C | 1187147 |
| rs530440643 | 15 | 42899141 A | C | 376781 |
| rs74477058 | 15 | 43001334 C | G | 1230477 |
| rs1211642 | 15 | 42890450 C | T | 1068700 |
| NA | 15 | 42957127 G | T | 94193 |
| rs752599758 | 15 | 42964444 A | T | 477772 |
| rs16957055 | 15 | 42980390 C | T | 1233936 |
| rs61218673 | 15 | 42981978 G | A | 672690 |
| rs8028365 | 15 | 42993930 C | T | 1234631 |
| rs534891696 | 15 | 42868588 C | T | 366232 |
| rs574188738 | 15 | 42900199 C | G | 32855 |
| rs146394639 | 15 | 42927581 G | A | 1227473 |
| rs3742995 | 15 | 42982819 A | G | 1233938 |
| rs565169604 | 15 | 42936375 C | T | 362434 |
| rs56286330 | 15 | 42950486 G | GGT | 994619 |
| rs8036733 | 15 | 42998776 G | T | 1237996 |
| rs145239633 | 15 | 42982455 A | G | 365728 |
| rs72711776 | 15 | 42916041 T | G | 1218020 |
| rs550231775 | 15 | 42925881 G | A | 379381 |
| rs876992 | 15 | 42878456 C | T | 1221490 |
| rs376374890 | 15 | 43009962 C | T | 571225 |
| rs533630312 | 15 | 42903619 C | T | 399220 |
| rs553415415 | 15 | 42998513 G | A | 370404 |
| rs770679768 | 15 | 43005828 T | G | 515107 |
| rs189296046 | 15 | 42906614 C | A | 365285 |
| rs750624051 | 15 | 42954133 C | A | 547936 |
| rs144334043 | 15 | 42873927 G | GT | 578803 |

|  |  |  |  |  |
| --- | --- | --- | --- | --- |
| rs778050361 | 15 | 42885190 T | G | 404433 |
| rs551482565 | 15 | 42905800 C | T | 680959 |
| rs574436638 | 15 | 42996347 G | A | 524530 |
| rs111805819 | 15 | 42947077 A | G | 741563 |
| rs188846832 | 15 | 42988440 G | T | 365728 |
| rs138677372 | 15 | 42931457 C | T | 1201285 |
| rs7164196 | 15 | 42869986 G | T | 682988 |
| rs630620 | 15 | 42909082 C | T | 879237 |
| rs538996660 | 15 | 42929482 CT | C | 373346 |
| rs559499569 | 15 | 42869508 T | C | 428908 |
| rs28736688 | 15 | 42945022 G | A | 1238928 |
| rs71474497 | 15 | 42951427 T | C | 1218659 |
| rs752837000 | 15 | 43002061 T | C | 484786 |
| rs557261358 | 15 | 42988192 A | G | 379380 |
| rs150895411 | 15 | 42889015 T | G | 1189594 |
| rs560512058 | 15 | 42992562 C | G | 607865 |
| rs141247863 | 15 | 42916512 CAG | C | 371945 |
| rs769686927 | 15 | 42926127 C | T | 502631 |
| rs145395424 | 15 | 42904938 T | C | 1205018 |
| rs112280084 | 15 | 42945127 C | T | 588102 |
| rs182241507 | 15 | 42967326 C | G | 370638 |
| rs12911561 | 15 | 42868357 C | T | 775896 |
| rs866926503 | 15 | 43005024 C | T | 707617 |
| rs572371078 | 15 | 43007480 T | C | 381504 |
| rs370811683 | 15 | 42964861 A | AG | 611354 |
| rs578201778 | 15 | 42970456 T | TTG | 712793 |
| rs1052365338 | 15 | 42897631 G | T | 29487 |
| rs574256734 | 15 | 42898994 A | G | 381329 |
| rs1056686032 | 15 | 42907056 G | A | 29487 |
| rs138121440 | 15 | 42974713 C | T | 1202618 |
| rs201049061 | 15 | 42907744 CAT | C | 624814 |
| rs568429587 | 15 | 42934240 AT | A | 853910 |
| rs539464370 | 15 | 43008093 C | T | 365539 |
| rs74009101 | 15 | 42895210 T | C | 645609 |
| rs544930217 | 15 | 42991793 A | T | 483132 |
| rs200569969 | 15 | 42906593 TAGG | T | 593455 |
| rs563855481 | 15 | 42889884 C | T | 423588 |
| rs541065612 | 15 | 42924612 T | C | 369744 |
| NA | 15 | 42892387 C | T | 360570 |
| rs113551877 | 15 | 42918582 C | T | 885777 |
| rs564907467 | 15 | 42952039 G | A | 602706 |
| rs530185347 | 15 | 42923671 G | T | 605863 |
| rs113103629 | 15 | 42987592 C | T | 1105875 |
| rs553132117 | 15 | 42956733 C | CA | 773210 |
| rs777344032 | 15 | 42872371 G | A | 6702 |
| rs536362302 | 15 | 42977617 T | C | 863618 |
| rs779195925 | 15 | 42892283 A | G | 642419 |
| rs74626491 | 15 | 42970123 C | A | 823492 |
| rs535962355 | 15 | 42960676 T | G | 828138 |
| rs753994390 | 15 | 42964301 C | G | 440454 |
| rs529269398 | 15 | 42972145 A | G | 545438 |
| rs565180029 | 15 | 43004581 C | T | 373979 |

|  |  |  |  |  |
| --- | --- | --- | --- | --- |
| rs1030428778 | 15 | 42929667 C | A | 26547 |
| rs574731559 | 15 | 42941284 A | G | 360699 |
| rs543049266 | 15 | 42919059 T | G | 366613 |
| rs189916861 | 15 | 42934339 G | A | 901307 |
| rs147055430 | 15 | 43005504 T | A | 361271 |
| rs555203230 | 15 | 42994269 C | A | 270377 |
| rs577736918 | 15 | 42995463 T | C | 373797 |
| rs556127396 | 15 | 42998583 A | G | 365285 |
| rs573571482 | 15 | 42932937 T | G | 369985 |
| rs113245184 | 15 | 42933898 G | A | 662302 |
| rs183125787 | 15 | 42934233 T | A | 655839 |
| rs577186934 | 15 | 42957491 T | C | 492332 |
| rs183640181 | 15 | 42963391 C | T | 619804 |
| rs185865927 | 15 | 42884511 C | T | 580131 |
| rs551937058 | 15 | 42893450 C | T | 365048 |
| rs576049319 | 15 | 42894935 T | C | 909790 |
| rs12323990 | 15 | 42971515 C | T | 1235138 |
| rs530762141 | 15 | 43009056 G | A | 403205 |
| rs143643681 | 15 | 42903812 C | T | 588784 |
| rs555231317 | 15 | 42952781 A | C | 141318 |
| rs747908929 | 15 | 43012424 C | T | 147900 |
| rs757758595 | 15 | 42959406 G | A | 401275 |
| rs541657557 | 15 | 42888732 A | G | 712920 |
| rs546152162 | 15 | 42929192 A | T | 541403 |
| rs144738245 | 15 | 42957109 A | G | 817899 |
| rs1027710403 | 15 | 42925755 A | C | 394043 |
| rs191655165 | 15 | 42904217 A | G | 588102 |
| rs558809042 | 15 | 42922202 T | C | 713877 |
| rs61753412 | 15 | 42986471 A | T | 1099429 |
| rs531993169 | 15 | 42916749 G | T | 382071 |
| rs575362514 | 15 | 42989281 A | G | 462437 |
| rs479519 | 15 | 42892809 C | T | 383889 |
| rs545086800 | 15 | 42972524 G | A | 677069 |
| rs568687731 | 15 | 42992682 A | T | 360129 |
| rs191154656 | 15 | 42874605 A | T | 365048 |
| rs373520291 | 15 | 42991781 T | C | 633569 |
| rs145590620 | 15 | 42933274 T | C | 1223535 |
| rs11314415 | 15 | 42916751 CT | C | 780619 |
| rs117775549 | 15 | 42973997 C | T | 1196541 |
| rs536329199 | 15 | 42991579 T | C | 606011 |
| rs79130229 | 15 | 42896228 G | C | 631021 |
| rs568813363 | 15 | 42952486 G | A | 374467 |
| rs772067408 | 15 | 42982047 G | T | 94193 |
| NA | 15 | 42981870 T | C | 394400 |
| rs1037588084 | 15 | 42875428 G | A | 170504 |
| rs186674983 | 15 | 42911925 G | A | 1221617 |
| rs74890375 | 15 | 42958672 A | G | 263494 |
| rs184720517 | 15 | 42991842 T | C | 812942 |
| rs191328807 | 15 | 42897371 A | G | 369743 |
| rs571391144 | 15 | 42904356 T | C | 365285 |
| rs117731492 | 15 | 42934853 A | G | 1223536 |
| rs200378215 | 15 | 42955816 A | G | 1232173 |

|  |  |  |  |  |
| --- | --- | --- | --- | --- |
| rs555851843 | 15 | 42908243 T | C | 877479 |
| rs563962097 | 15 | 42924629 A | G | 900288 |
| rs184949186 | 15 | 42954225 T | A | 21517 |
| rs539879268 | 15 | 42961387 A | G | 371779 |
| rs781419142 | 15 | 42923300 C | T | 558421 |
| rs533049646 | 15 | 42924634 A | G | 376238 |
| rs566075663 | 15 | 42963258 A | C | 371779 |
| rs10162744 | 15 | 42879383 A | C | 622708 |
| rs367880689 | 15 | 42880211 TTG | T | 578803 |
| rs147646076 | 15 | 42929139 G | C | 1223680 |
| rs150157757 | 15 | 42919458 T | C | 718433 |
| rs182258163 | 15 | 42936560 G | A | 1223537 |
| rs536290738 | 15 | 42947564 A | G | 436454 |
| rs567147466 | 15 | 43004874 A | G | 424123 |
| rs9652443 | 15 | 42892631 T | C | 628893 |
| rs752932916 | 15 | 42895952 T | C | 1215 |
| rs151012902 | 15 | 42899817 G | A | 1233997 |
| rs756077457 | 15 | 42920689 A | G | 219558 |
| rs529322354 | 15 | 42968801 A | T | 876427 |
| rs142987541 | 15 | 43006485 C | T | 831512 |
| rs78325423 | 15 | 43012549 G | C | 1211944 |
| rs541204716 | 15 | 42889085 C | T | 365285 |
| rs74715782 | 15 | 42903428 G | A | 670456 |
| rs192212691 | 15 | 42929560 G | A | 437961 |
| rs184321765 | 15 | 42881661 A | G | 377849 |
| rs75939632 | 15 | 42907791 A | T | 715955 |
| NA | 15 | 42981031 C | G | 389557 |
| rs546146720 | 15 | 42945342 T | A | 591649 |
| NA | 15 | 42956315 T | C | 125769 |
| rs558670205 | 15 | 42948234 C | T | 897714 |
| rs570784384 | 15 | 42876851 A | C | 369743 |
| rs117522776 | 15 | 42963841 C | T | 605731 |
| rs942809430 | 15 | 42996893 G | C | 535411 |
| rs116565691 | 15 | 42888132 G | A | 595629 |
| rs138864471 | 15 | 42936564 A | G | 1222844 |
| rs943592293 | 15 | 42950524 C | G | 29487 |
| rs78562640 | 15 | 43002759 A | G | 1208570 |
| rs939140502 | 15 | 42869541 T | C | 30186 |
| rs12324135 | 15 | 42942696 C | T | 1238923 |
| rs12592374 | 15 | 42991299 A | G | 1235478 |
| rs75617784 | 15 | 42914904 G | A | 578803 |
| rs767836352 | 15 | 42894878 A | G | 682814 |
| rs541664623 | 15 | 42961778 C | T | 357928 |
| rs79994978 | 15 | 42902639 T | C | 762175 |
| rs539091007 | 15 | 42893886 G | A | 577971 |
| rs57133852 | 15 | 42923960 GA | G | 986176 |
| rs538247556 | 15 | 42943741 C | T | 897676 |
| rs202031252 | 15 | 42935804 TCGGCTGAGG | T | 29487 |
| rs112745231 | 15 | 42982708 A | G | 643295 |
| rs546043408 | 15 | 42881894 T | G | 365221 |
| rs748457096 | 15 | 42957287 G | C | 156631 |
| rs572610576 | 15 | 42969282 C | G | 230552 |

|  |  |  |  |  |
| --- | --- | --- | --- | --- |
| rs79331266 | 15 | 42953161 T | G | 252111 |
| rs143499451 | 15 | 42953711 T | C | 600212 |
| rs199940227 | 15 | 42984727 T | G | 382407 |
| rs188267149 | 15 | 42891458 C | G | 172859 |
| rs199708541 | 15 | 42896064 TC | T | 1080825 |
| rs547575932 | 15 | 42896069 G | A | 1082128 |
| rs187403913 | 15 | 42909578 G | A | 490356 |
| rs750109325 | 15 | 42934843 C | T | 357298 |
| rs140413699 | 15 | 42924271 C | T | 401727 |
| rs78196628 | 15 | 42921854 C | T | 1203758 |
| rs770699990 | 15 | 42901032 G | A | 810835 |
| rs563630307 | 15 | 42924465 G | T | 484464 |
| rs537715420 | 15 | 42911798 G | A | 492732 |
| rs576555473 | 15 | 42919375 T | C | 591648 |
| rs533865727 | 15 | 42963902 AG | A | 236906 |
| rs111312181 | 15 | 42898767 C | T | 644482 |
| rs560294077 | 15 | 42933084 CAT | C | 413860 |
| rs183583949 | 15 | 42959536 C | T | 722760 |
| rs114356937 | 15 | 42974042 T | C | 623444 |
| rs148448542 | 15 | 42911342 A | G | 1144737 |
| rs910876412 | 15 | 42934648 A | G | 21412 |
| rs12594858 | 15 | 42939916 G | A | 1231495 |
| rs561702892 | 15 | 42992719 A | G | 497282 |
| rs185252821 | 15 | 42868239 G | C | 422111 |
| rs187967591 | 15 | 42948375 G | A | 364588 |
| rs79365604 | 15 | 42976440 G | A | 1213947 |
| rs141542733 | 15 | 42911557 C | T | 364588 |
| rs150050265 | 15 | 42917181 T | A | 1191878 |
| rs572054086 | 15 | 42890464 C | T | 222451 |
| rs139864533 | 15 | 42889447 C | T | 755166 |
| rs531067051 | 15 | 43011162 C | T | 448859 |
| rs142075513 | 15 | 42923651 G | A | 655408 |
| rs114793100 | 15 | 42932355 A | T | 1229643 |
| rs59900882 | 15 | 43002685 C | G | 1158009 |
| rs372097263 | 15 | 42879057 G | T | 222451 |
| rs149877181 | 15 | 42972932 G | A | 1145721 |
| rs77922743 | 15 | 42993355 C | G | 1145447 |
| rs371614413 | 15 | 43011821 A | G | 839476 |
| rs776350968 | 15 | 43011503 G | A | 19925 |
| rs147572892 | 15 | 42868482 CG | C | 592827 |
| rs114831533 | 15 | 42885803 C | A | 236946 |
| NA | 15 | 42944185 G | A | 103038 |
| rs753886624 | 15 | 42938027 A | G | 412767 |
| rs540055435 | 15 | 42873773 AGC | A | 222451 |
| rs541928952 | 15 | 42902878 G | A | 577971 |
| rs562642851 | 15 | 42944671 C | A | 371762 |
| rs189800151 | 15 | 42960589 T | C | 1194625 |
| rs117842035 | 15 | 42978426 T | A | 820720 |
| rs770269074 | 15 | 42898276 A | G | 575553 |
| rs16956958 | 15 | 42944616 G | A | 280097 |
| rs58903190 | 15 | 42946958 C | T | 840051 |
| rs115058341 | 15 | 42901551 A | T | 668879 |

|  |  |  |  |  |
| --- | --- | --- | --- | --- |
| rs12910030 | 15 | 42868286 A | G | 1182976 |
| rs116267701 | 15 | 42876795 C | T | 236946 |
| rs535464309 | 15 | 42946008 T | TA | 359120 |
| rs542582189 | 15 | 42979085 G | A | 60911 |
| rs183344754 | 15 | 42985129 A | G | 46613 |
| rs571668597 | 15 | 43003370 C | T | 392931 |
| rs771981477 | 15 | 42893470 G | A | 29487 |
| rs187407014 | 15 | 42931263 A | G | 863674 |
| rs12324838 | 15 | 42940790 C | T | 1230002 |
| rs114784267 | 15 | 42996974 G | C | 258284 |
| rs550166365 | 15 | 42869382 C | G | 408191 |
| rs143222805 | 15 | 42889798 G | C | 1237981 |
| rs569984356 | 15 | 42869414 C | T | 539886 |
| rs117170089 | 15 | 42997618 G | T | 966885 |
| rs182223038 | 15 | 42941333 T | C | 369744 |
| rs558891400 | 15 | 42956104 C | T | 657500 |
| rs114533051 | 15 | 42980487 T | A | 1012981 |
| rs551993514 | 15 | 42992228 C | T | 523163 |
| rs543662054 | 15 | 42890537 A | G | 365285 |
| rs116820442 | 15 | 42921051 C | T | 740487 |
| rs114690791 | 15 | 42937403 A | G | 741040 |
| rs530364055 | 15 | 42880494 A | G | 365285 |
| rs6493056 | 15 | 42913470 G | A | 1229302 |
| rs541093919 | 15 | 42897691 T | G | 445326 |
| rs537910025 | 15 | 42875968 T | C | 404755 |
| rs552708360 | 15 | 42953454 A | C | 539093 |
| rs557922265 | 15 | 42988143 A | G | 588392 |
| NA | 15 | 42874103 CTT | C | 29487 |
| rs771776965 | 15 | 42908221 C | T | 82056 |
| rs553969178 | 15 | 42915402 A | C | 633406 |
| rs546340248 | 15 | 42935820 G | C | 1053552 |
| rs77096807 | 15 | 42982124 A | G | 818460 |
| rs538838662 | 15 | 42900649 C | G | 370831 |
| rs773669948 | 15 | 42883556 C | T | 855802 |
| rs552844514 | 15 | 42950749 T | G | 70132 |
| NA | 15 | 42965470 G | A | 223174 |
| rs13380302 | 15 | 42900391 G | C | 619245 |
| rs527501558 | 15 | 42935817 CTA | C | 1052209 |
| rs187495317 | 15 | 42963968 G | A | 983333 |
| rs116233593 | 15 | 42986987 G | A | 1018020 |
| NA | 15 | 42897419 G | A | 29487 |
| rs145552657 | 15 | 42996251 A | G | 1238365 |
| rs115595492 | 15 | 42888436 C | G | 626485 |
| rs28699522 | 15 | 42946935 G | C | 1238665 |
| rs528684067 | 15 | 42896224 G | C | 1082919 |
| rs186001502 | 15 | 42903616 T | G | 927144 |
| rs548883046 | 15 | 42921116 T | G | 802184 |
| rs185441627 | 15 | 42937866 T | G | 605893 |
| rs139275104 | 15 | 42988857 C | T | 838775 |
| rs566475856 | 15 | 42902894 G | A | 941776 |
| NA | 15 | 42908353 T | C | 96813 |
| rs1006058718 | 15 | 43012832 G | A | 532081 |

|  |  |  |  |  |
| --- | --- | --- | --- | --- |
| rs141596381 | 15 | 42898603 T | C | 222451 |
| rs187732802 | 15 | 42905119 T | G | 222451 |
| rs73408611 | 15 | 42945580 C | T | 1238923 |
| rs182571354 | 15 | 42870794 A | G | 222451 |
| rs80319409 | 15 | 42921919 T | C | 1216445 |
| rs56839966 | 15 | 42945683 G | T | 1238924 |
| rs774605196 | 15 | 42987180 CA | C | 98822 |
| rs144945764 | 15 | 42999341 T | TA | 1078501 |
| rs114416020 | 15 | 42981421 C | T | 1012720 |
| rs116318862 | 15 | 42989917 A | G | 1018765 |
| rs148013951 | 15 | 42906336 C | T | 1141318 |
| rs778806352 | 15 | 42938861 G | C | 677029 |
| NA | 15 | 42963392 G | A | 22669 |
| NA | 15 | 42908082 G | C | 6702 |
| rs60484860 | 15 | 43008972 G | A | 907870 |
| rs186670557 | 15 | 42893536 A | G | 627317 |
| rs201923100 | 15 | 42944469 TA | T | 1032676 |
| NA | 15 | 42966323 A | G | 381177 |
| rs986243537 | 15 | 42967543 C | T | 70372 |
| rs539402884 | 15 | 43001119 T | A | 19194 |
| NA | 15 | 42879347 CCCAG | TCCAG | 222364 |
| rs113162826 | 15 | 42897162 C | T | 644482 |
| rs1005109579 | 15 | 42941159 T | G | 415736 |
| rs190857140 | 15 | 42870763 G | A | 1174064 |
| rs569250173 | 15 | 42904514 CAGTG | C | 720724 |
| rs139197926 | 15 | 42905546 C | T | 492432 |
| rs963090177 | 15 | 42915175 C | T | 381177 |
| rs139264905 | 15 | 42908568 G | A | 1232808 |
| rs572358923 | 15 | 43000193 A | G | 379986 |
| rs79749780 | 15 | 42963806 T | A | 280098 |
| rs114876104 | 15 | 42921192 A | G | 607776 |
| rs114642146 | 15 | 42922785 T | G | 607776 |
| rs139898805 | 15 | 42951125 G | A | 596435 |
| rs889746332 | 15 | 42875974 C | T | 418179 |
| rs1013279760 | 15 | 42926283 G | A | 381177 |
| rs572736554 | 15 | 42918591 T | C | 787538 |
| rs550361351 | 15 | 42942246 C | G | 591858 |
| rs550103536 | 15 | 42939466 T | C | 810686 |
| rs115593137 | 15 | 42970388 G | A | 1015935 |
| rs746489852 | 15 | 43002363 A | G | 485223 |
| rs185405546 | 15 | 42885977 T | C | 661683 |
| rs549994482 | 15 | 42924359 T | G | 236030 |
| rs751302053 | 15 | 42871640 G | A | 861619 |
| rs60111534 | 15 | 42868169 CCG | C | 1054768 |
| rs1036190902 | 15 | 42959670 C | T | 361954 |
| rs376229251 | 15 | 42978141 A | C | 290954 |
| rs565841251 | 15 | 42910911 C | T | 450106 |
| rs114435521 | 15 | 42938902 G | A | 711275 |
| rs142916862 | 15 | 42913461 A | G | 566370 |
| rs7342639 | 15 | 42925904 C | T | 677408 |
| rs777092034 | 15 | 42982568 C | T | 613914 |
| rs569941620 | 15 | 42937337 C | T | 467040 |

|  |  |  |  |  |
| --- | --- | --- | --- | --- |
| rs141336135 | 15 | 42891358 G | C | 1128408 |
| rs10163044 | 15 | 42903456 G | C | 630963 |
| rs974904581 | 15 | 42918564 T | C | 626037 |
| rs141161691 | 15 | 42992007 C | T | 589524 |
| rs181986747 | 15 | 42995783 C | T | 615702 |
| rs566344796 | 15 | 42878328 G | A | 511046 |
| rs150824141 | 15 | 42885513 AT | A | 895130 |
| rs924868347 | 15 | 42871846 A | G | 421596 |
| NA | 15 | 42896073 C | G | 29487 |
| rs578228153 | 15 | 42911780 A | G | 717935 |
| rs554716041 | 15 | 42922198 C | T | 341483 |
| rs148437368 | 15 | 42938858 T | G | 376231 |
| rs751191269 | 15 | 43011782 G | C | 183579 |
| rs6493055 | 15 | 42912620 A | G | 1224625 |
| rs747499658 | 15 | 42935069 T | C | 681831 |
| rs559982616 | 15 | 42936475 A | G | 356895 |
| rs60753136 | 15 | 42915693 G | A | 1224992 |
| rs554337611 | 15 | 42881817 T | C | 549819 |
| rs555489047 | 15 | 42886831 G | GT | 368626 |
| rs184696803 | 15 | 42910131 T | A | 360129 |
| rs182501865 | 15 | 42948152 G | A | 593104 |
| rs113206433 | 15 | 42893203 A | G | 1162129 |
| rs116333487 | 15 | 42977839 G | C | 1012980 |
| rs565525729 | 15 | 42983143 C | A | 371400 |
| rs557439861 | 15 | 42926132 G | A | 510943 |
| rs149230374 | 15 | 42986336 A | G | 380000 |
| rs143081363 | 15 | 42991837 C | T | 1016726 |
| rs566924050 | 15 | 42991956 G | A | 586032 |
| rs537457295 | 15 | 42942607 C | T | 365221 |
| rs79533437 | 15 | 42948926 A | G | 1145319 |
| rs543405408 | 15 | 42886259 G | A | 700019 |
| rs142645685 | 15 | 42899443 C | G | 630131 |
| rs543217741 | 15 | 42940658 G | C | 225966 |
| rs150729042 | 15 | 42950747 C | G | 670816 |
| rs181125969 | 15 | 42977568 G | A | 377496 |
| rs16973396 | 15 | 42915785 A | T | 739740 |
| rs138873449 | 15 | 42891145 C | T | 222451 |
| rs137999320 | 15 | 42868678 T | G | 222451 |
| rs151310546 | 15 | 42881602 C | T | 222451 |
| rs562232008 | 15 | 42936049 A | G | 835082 |
| rs190028056 | 15 | 42941943 G | A | 224113 |
| rs138423727 | 15 | 43005653 T | C | 1229497 |
| rs114643417 | 15 | 42892578 G | A | 222451 |
| NA | 15 | 42983932 G | A | 27798 |
| rs188025826 | 15 | 42870741 A | G | 222451 |
| rs16957031 | 15 | 42969114 G | A | 438910 |
| rs564756186 | 15 | 42959022 A | AT | 597035 |
| rs142543234 | 15 | 42873991 A | G | 222451 |
| rs145036897 | 15 | 42896996 C | T | 1203300 |
| rs185606555 | 15 | 42902671 A | G | 222451 |
| rs7167971 | 15 | 42912455 A | G | 596435 |
| rs145521822 | 15 | 42914681 T | C | 406236 |

|  |  |  |  |  |
| --- | --- | --- | --- | --- |
| NA | 15 | 42982744 G | C | 27798 |
| rs551210946 | 15 | 42991368 G | C | 531003 |
| rs144114925 | 15 | 42903341 A | G | 222451 |
| rs187974547 | 15 | 42979226 C | G | 1011260 |
| rs567833296 | 15 | 42910039 C | T | 594595 |
| rs189324154 | 15 | 42868078 G | A | 222451 |
| rs116259624 | 15 | 42975260 C | T | 1014980 |
| rs759661465 | 15 | 42985854 G | T | 357298 |
| rs10163179 | 15 | 42879555 G | A | 827563 |
| rs77836884 | 15 | 42986787 C | T | 788517 |
| NA | 15 | 42915711 G | T | 19723 |
| rs529590315 | 15 | 42941017 T | C | 425687 |
| rs116745790 | 15 | 42980023 G | A | 1012980 |
| rs536362397 | 15 | 42984299 T | G | 370007 |
| rs78904754 | 15 | 42994141 C | T | 577971 |
| rs575816891 | 15 | 42961747 C | G | 875138 |
| rs115105685 | 15 | 42874825 T | A | 259116 |
| rs8039217 | 15 | 42987137 C | A | 1147653 |
| rs772774346 | 15 | 42980278 A | C | 544839 |
| rs144291102 | 15 | 42912619 C | T | 230552 |
| NA | 15 | 42993022 T | G | 20912 |
| rs529923027 | 15 | 42957022 A | C | 835027 |
| rs117366041 | 15 | 42914185 A | G | 567148 |
| rs112765931 | 15 | 42931212 GAAAC | G | 969372 |
| rs113697440 | 15 | 42970776 C | T | 1018205 |
| rs186290703 | 15 | 42932876 C | T | 794232 |
| rs200058353 | 15 | 42983771 C | G | 827375 |
| rs181335046 | 15 | 43001629 A | G | 1205076 |
| rs192217223 | 15 | 42899883 C | A | 925412 |
| rs779112444 | 15 | 42992926 C | A | 443401 |
| rs546974729 | 15 | 42963858 G | A | 906071 |
| rs754784581 | 15 | 42982890 G | GAGC | 89993 |
| rs185424711 | 15 | 43009737 C | T | 355834 |
| NA | 15 | 42975628 T | G | 38048 |
| rs672150 | 15 | 42918082 C | T | 277370 |
| rs142538022 | 15 | 42911994 A | G | 577971 |
| rs141935808 | 15 | 42905231 AG | A | 603047 |
| rs573580149 | 15 | 42911336 A | C | 366164 |
| rs560437326 | 15 | 42872620 T | G | 967019 |
| rs562182558 | 15 | 42880682 G | T | 586904 |
| rs562465519 | 15 | 42894211 G | A | 372414 |
| rs372595609 | 15 | 42918728 C | T | 432538 |
| rs76712048 | 15 | 42925075 G | A | 1233871 |
| rs185512143 | 15 | 42943200 C | T | 428884 |
| rs148183291 | 15 | 42991986 C | T | 596435 |
| rs546611281 | 15 | 42985711 T | C | 786831 |
| rs756126576 | 15 | 42869616 G | A | 589238 |
| rs57460884 | 15 | 43008756 C | T | 1158252 |
| rs9652442 | 15 | 42891654 T | C | 630131 |
| rs150492102 | 15 | 42999937 TTG | T | 630200 |
| rs777539145 | 15 | 43010965 A | G | 574614 |
| rs150519590 | 15 | 42907387 T | C | 958800 |

|  |  |  |  |  |
| --- | --- | --- | --- | --- |
| rs7163324 | 15 | 42988175 C | T | 1143742 |
| rs574639690 | 15 | 42923049 T | G | 556437 |
| rs576866690 | 15 | 42945758 G | A | 369744 |
| rs147795392 | 15 | 42953329 A | G | 934291 |
| rs576086194 | 15 | 42910995 A | ATGTTGCCAGC | 586903 |
| rs77757620 | 15 | 42953597 C | T | 1150170 |
| rs188518430 | 15 | 42895715 C | A | 606566 |
| rs569316332 | 15 | 42901679 C | G | 732477 |
| rs189176833 | 15 | 42906239 G | A | 818035 |
| NA | 15 | 42975732 T | C | 75008 |
| rs765294767 | 15 | 42876417 T | G | 504525 |
| rs62019357 | 15 | 42948374 C | T | 1222839 |
| rs114426631 | 15 | 42871821 C | T | 258284 |
| rs187131404 | 15 | 42911059 C | T | 987887 |
| rs540343022 | 15 | 42915699 T | C | 364587 |
| rs574542771 | 15 | 42922405 G | A | 790687 |
| rs72711781 | 15 | 42952175 A | G | 1241262 |
| rs572298610 | 15 | 42869451 A | G | 557237 |
| NA | 15 | 42873849 G | C | 373248 |
| rs2899060 | 15 | 42899437 G | T | 243860 |
| rs8039346 | 15 | 42987070 G | C | 1145864 |
| rs544510397 | 15 | 42938358 T | TCAA | 69589 |
| rs550555857 | 15 | 42987196 T | TA | 951145 |
| rs57568065 | 15 | 42945796 CT | C | 1068032 |
| rs563830686 | 15 | 42960390 A | AC | 236906 |
| rs543953990 | 15 | 42965920 G | A | 1223702 |
| rs9806554 | 15 | 42876753 T | A | 258284 |
| rs183399424 | 15 | 42941468 C | T | 445071 |
| rs16957043 | 15 | 42975053 G | A | 1236333 |
| rs182806011 | 15 | 42930784 A | G | 240083 |
| rs545016214 | 15 | 42940515 C | T | 1186440 |
| rs539697836 | 15 | 42889573 A | T | 311465 |
| rs146948676 | 15 | 42906046 C | A | 630131 |
| rs116232111 | 15 | 42994170 G | A | 1013652 |
| rs887317340 | 15 | 42958646 G | A | 626191 |
| rs561210836 | 15 | 42871263 G | A | 366230 |
| rs769158133 | 15 | 42992247 T | C | 549294 |
| rs529988191 | 15 | 42956480 C | G | 838775 |
| rs191115724 | 15 | 42884735 C | T | 611109 |
| rs575135631 | 15 | 42980297 A | G | 357680 |
| rs148235732 | 15 | 42916043 G | A | 676279 |
| rs79114507 | 15 | 42930636 G | A | 700219 |
| rs773809476 | 15 | 42938556 G | A | 841541 |
| rs112297283 | 15 | 42886367 G | T | 815437 |
| rs564661828 | 15 | 42928648 G | A | 733481 |
| rs542480757 | 15 | 42884760 C | T | 365285 |
| rs564939782 | 15 | 42905529 A | G | 369062 |
| rs150908758 | 15 | 43006166 G | A | 647306 |
| rs541693905 | 15 | 42920232 A | T | 620098 |
| rs551060551 | 15 | 43004638 T | C | 647306 |
| rs113567591 | 15 | 42921313 G | C | 1231114 |
| rs114982552 | 15 | 42913679 G | A | 606200 |

|  |  |  |  |  |
| --- | --- | --- | --- | --- |
| rs184939172 | 15 | 42913788 A | T | 621050 |
| rs898531013 | 15 | 42956882 T | A | 547584 |
| rs188499850 | 15 | 42974646 G | A | 562565 |
| rs778466650 | 15 | 42903801 A | ATTGTCC | 76061 |
| rs566759400 | 15 | 42942150 C | T | 631892 |
| rs535004528 | 15 | 42977084 C | T | 373521 |
| rs114736360 | 15 | 42969940 C | T | 1014592 |
| rs183222293 | 15 | 42973350 G | A | 838775 |
| rs72711791 | 15 | 43007646 A | G | 1244487 |
| rs115660182 | 15 | 42889346 T | G | 270133 |
| rs140074516 | 15 | 42903809 T | C | 365285 |
| rs776432879 | 15 | 42905068 G | C | 357298 |
| rs764114296 | 15 | 42964236 A | G | 714268 |
| rs75079841 | 15 | 42923403 T | C | 606200 |
| rs560294808 | 15 | 42943755 T | C | 602730 |
| rs145276351 | 15 | 42914910 A | G | 577971 |
| rs116507617 | 15 | 42915653 A | G | 606200 |
| rs757526455 | 15 | 42919492 T | G | 387974 |
| rs1010566444 | 15 | 42933566 T | G | 602730 |
| rs148793160 | 15 | 42882955 C | CGA | 1073082 |
| rs556120932 | 15 | 42932790 C | T | 369744 |
| rs528635271 | 15 | 42935706 T | A | 460109 |
| rs79774962 | 15 | 43005092 C | T | 1149941 |
| rs373035305 | 15 | 42867925 G | A | 443776 |
| rs941905115 | 15 | 42899346 A | G | 29487 |
| rs539280750 | 15 | 42929787 T | C | 562587 |
| rs16957011 | 15 | 42961901 T | C | 662230 |
| rs117486394 | 15 | 42984892 A | C | 691384 |
| rs28461422 | 15 | 42883883 G | A | 266356 |
| rs1009186043 | 15 | 42989671 A | G | 458212 |
| rs151101516 | 15 | 42913469 C | T | 439646 |
| rs761481866 | 15 | 42902640 G | T | 635068 |
| rs761882822 | 15 | 42954006 T | G | 614321 |
| rs139919621 | 15 | 42946665 G | T | 706369 |
| rs76108347 | 15 | 42955118 A | C | 707000 |
| rs186941746 | 15 | 42933630 T | C | 613636 |
| rs532830494 | 15 | 42968254 T | G | 851864 |
| rs193223759 | 15 | 42902603 C | T | 956859 |
| rs190551061 | 15 | 42907287 A | T | 273999 |
| rs188552931 | 15 | 42901005 G | A | 230552 |
| rs187890018 | 15 | 42973471 G | C | 838775 |
| rs541758310 | 15 | 42918606 T | C | 748298 |
| rs563034338 | 15 | 42924040 T | A | 659853 |
| rs146015002 | 15 | 42929887 A | G | 600965 |
| rs78859522 | 15 | 42952782 C | A | 607068 |
| rs114844882 | 15 | 42909326 T | C | 619245 |
| rs558776330 | 15 | 42929884 T | C | 365285 |
| rs116088885 | 15 | 42879631 G | A | 596435 |
| rs766798280 | 15 | 42949754 A | G | 369518 |
| rs547550289 | 15 | 42982862 A | G | 360811 |
| rs574063567 | 15 | 42893636 G | A | 270965 |
| rs17767439 | 15 | 42927182 A | G | 1231379 |

|  |  |  |  |  |
| --- | --- | --- | --- | --- |
| rs539366399 | 15 | 42973867 C | T | 360811 |
| rs571609256 | 15 | 42996263 G | A | 1072896 |
| rs545657563 | 15 | 42997366 C | T | 780776 |
| rs545694138 | 15 | 42998053 C | T | 803408 |
| rs148023580 | 15 | 42960312 G | A | 1137166 |
| rs760584528 | 15 | 42966962 G | T | 937029 |
| rs73406557 | 15 | 42868515 C | G | 1123244 |
| rs567987273 | 15 | 42973305 T | C | 403038 |
| rs565324621 | 15 | 43011460 C | T | 360699 |
| rs188303137 | 15 | 42888701 G | A | 938303 |
| rs77736048 | 15 | 42909596 T | G | 1050486 |
| rs753974961 | 15 | 42992817 T | C | 683741 |
| rs140656467 | 15 | 43012692 G | A | 709190 |
| rs10162619 | 15 | 42879686 T | G | 267188 |
| rs28378282 | 15 | 42884072 A | T | 267188 |
| rs76594962 | 15 | 42979456 T | G | 595721 |
| rs370295021 | 15 | 42867875 C | G | 262239 |
| rs766696626 | 15 | 42955169 A | G | 786972 |
| rs570832687 | 15 | 42976712 G | C | 29487 |
| rs888907858 | 15 | 42900841 A | C | 508181 |
| rs562486786 | 15 | 42958834 A | G | 487065 |
| rs115889964 | 15 | 42964282 T | C | 698188 |
| rs570858258 | 15 | 43008007 A | C | 257047 |
| rs920583154 | 15 | 42906931 G | A | 26317 |
| rs567595733 | 15 | 42936635 G | A | 708292 |
| rs765261127 | 15 | 42954125 A | C | 780852 |
| rs561497228 | 15 | 42979405 A | T | 853277 |
| rs143110514 | 15 | 42892381 C | T | 1213984 |
| rs11273757 | 15 | 42964854 GGGACAGAGGC | G | 969443 |
| rs2412739 | 15 | 42972955 T | C | 709111 |
| rs147737228 | 15 | 42881116 G | A | 267188 |
| NA | 15 | 42960106 A | T | 43581 |
| rs569336802 | 15 | 42964263 T | A | 547819 |
| rs115044157 | 15 | 42891021 C | T | 752886 |
| rs553534463 | 15 | 42993913 C | T | 635585 |
| rs139768798 | 15 | 42879951 C | T | 361075 |
| rs550580422 | 15 | 42885935 C | T | 369062 |
| rs183773847 | 15 | 42915614 A | C | 885394 |
| rs573499753 | 15 | 42931951 A | G | 25863 |
| rs111315571 | 15 | 42915042 A | G | 614690 |
| rs528276071 | 15 | 42979359 CAGCACA | C | 168339 |
| rs771882568 | 15 | 42990748 T | C | 737779 |
| rs575749928 | 15 | 42893616 C | T | 593818 |
| NA | 15 | 42949475 T | C | 46120 |
| rs111656044 | 15 | 42888606 A | G | 593818 |
| rs181739281 | 15 | 42874183 A | C | 586041 |
| rs552485294 | 15 | 42982602 G | A | 806531 |
| rs202070432 | 15 | 42983192 A | G | 936532 |
| rs565914422 | 15 | 42868570 G | A | 371132 |
| rs189531502 | 15 | 42951301 A | C | 1162216 |
| rs192447262 | 15 | 42889845 C | T | 577971 |
| rs147095009 | 15 | 42910982 A | T | 410615 |

|  |  |  |  |  |
| --- | --- | --- | --- | --- |
| rs562322333 | 15 | 42992185 G | A | 369062 |
| rs116577253 | 15 | 42873997 A | C | 240083 |
| rs60051108 | 15 | 42920507 C | T | 948072 |
| rs577549839 | 15 | 42940429 C | T | 111544 |
| rs138565386 | 15 | 42970310 T | C | 1059176 |
| rs144343572 | 15 | 42871691 C | T | 753566 |
| rs1029083837 | 15 | 43006299 C | T | 367267 |
| rs143440000 | 15 | 42879189 G | A | 240083 |
| rs7174847 | 15 | 42913750 A | T | 687935 |
| rs1045397885 | 15 | 42916252 C | A | 365725 |
| rs150313767 | 15 | 42958792 G | A | 233468 |
| rs568016106 | 15 | 42959617 G | A | 233468 |
| rs141553512 | 15 | 43006043 T | G | 372166 |
| rs150536629 | 15 | 42895591 G | A | 1156355 |
| rs185205489 | 15 | 42961844 C | T | 233468 |
| rs761953576 | 15 | 42886839 T | C | 499509 |
| rs144363278 | 15 | 42968140 G | A | 318759 |
| rs558081449 | 15 | 43005954 G | C | 979061 |
| rs544454539 | 15 | 42964412 C | T | 604082 |
| rs190287691 | 15 | 42989886 C | T | 950326 |
| rs529032170 | 15 | 42960888 T | G | 239820 |
| rs369454740 | 15 | 42983848 T | C | 71751 |
| rs138770997 | 15 | 42879489 C | G | 240083 |
| rs78263141 | 15 | 42880383 G | T | 757344 |
| rs760057838 | 15 | 42869202 G | A | 363054 |
| rs112369865 | 15 | 42910650 G | C | 614690 |
| rs112267992 | 15 | 42895540 G | C | 573566 |
| rs151217518 | 15 | 42923871 G | A | 577971 |
| rs577438946 | 15 | 42939829 T | C | 369744 |
| rs139999040 | 15 | 42883642 C | T | 656514 |
| rs538590478 | 15 | 43004069 G | A | 577971 |
| rs566567441 | 15 | 42965971 G | GCCATAATCT | 748992 |
| rs148176549 | 15 | 43009749 G | A | 1239828 |
| rs62019359 | 15 | 42976706 C | T | 1229811 |
| rs186343429 | 15 | 42891962 G | A | 824843 |
| rs572370881 | 15 | 42902097 C | G | 481285 |
| rs142703729 | 15 | 42915717 A | G | 1221197 |
| rs780107990 | 15 | 42913907 G | A | 388798 |
| rs895643189 | 15 | 42889532 A | G | 19723 |
| rs543442379 | 15 | 42920839 T | G | 399115 |
| rs74009211 | 15 | 43008971 C | A | 714806 |
| rs764365243 | 15 | 42918606 TTCC | T | 357298 |
| rs556801807 | 15 | 42932670 A | G | 832498 |
| rs1036113951 | 15 | 42919970 A | G | 385858 |
| rs767425815 | 15 | 42935673 C | A | 252417 |
| rs938046 | 15 | 43010364 C | T | 1022377 |
| rs751263954 | 15 | 43006721 G | A | 417238 |
| rs554039449 | 15 | 42880312 G | T | 370287 |
| rs563666899 | 15 | 42944936 C | T | 946832 |
| rs12438056 | 15 | 42959540 G | A | 1193486 |
| rs577202617 | 15 | 43000362 A | T | 439129 |
| rs193064675 | 15 | 42882083 G | A | 910047 |

|  |  |  |  |  |
| --- | --- | --- | --- | --- |
| rs182572446 | 15 | 42878337 G | C | 632410 |
| rs895701914 | 15 | 42924428 A | G | 522702 |
| rs75891360 | 15 | 42891957 A | C | 658090 |
| rs561020009 | 15 | 42941197 G | C | 373521 |
| rs6493058 | 15 | 42962576 T | C | 1121716 |
| rs189545083 | 15 | 42992904 A | G | 1079458 |
| rs188271346 | 15 | 42910848 G | A | 642861 |
| rs577807047 | 15 | 42889101 C | T | 427198 |
| rs536364283 | 15 | 42977319 T | C | 691995 |
| rs201643568 | 15 | 42984156 C | G | 1190479 |
| rs116760018 | 15 | 42887668 T | G | 613440 |
| rs188599187 | 15 | 42991957 T | G | 385485 |
| rs189832961 | 15 | 42922492 C | T | 367966 |
| rs188267549 | 15 | 42939749 C | G | 1225224 |
| rs559047960 | 15 | 43005859 GTCTTT | G | 588102 |
| NA | 15 | 42884547 A | G | 361954 |
| rs183580304 | 15 | 42891278 T | C | 656514 |
| rs527471622 | 15 | 42938913 T | C | 670659 |
| rs114984341 | 15 | 42879064 C | T | 757344 |
| rs143689566 | 15 | 42897434 A | G | 1156354 |
| rs1020271692 | 15 | 42972418 T | C | 501159 |
| rs373300237 | 15 | 42918621 CTCT | C | 1029930 |
| rs745743014 | 15 | 42930019 C | T | 503500 |
| rs547725013 | 15 | 42953580 G | A | 398393 |
| rs564355348 | 15 | 42933848 A | G | 581748 |
| rs557425991 | 15 | 42950657 A | G | 372285 |
| rs569234592 | 15 | 42907158 G | T | 365285 |
| rs151016272 | 15 | 43008947 A | G | 370639 |
| rs187036267 | 15 | 42947859 C | T | 374468 |
| rs774795729 | 15 | 42945865 A | G | 715906 |
| rs183794720 | 15 | 42935957 T | C | 596435 |
| rs761344257 | 15 | 42935771 C | T | 513744 |
| rs186959564 | 15 | 42870931 T | C | 1083826 |
| rs576093418 | 15 | 42931389 G | A | 686503 |
| rs755384675 | 15 | 42967612 A | G | 394400 |
| rs535583406 | 15 | 42893630 C | T | 666506 |
| rs551970799 | 15 | 42929307 C | T | 405840 |
| rs115969538 | 15 | 42896702 T | C | 596435 |
| rs59881302 | 15 | 42930419 G | A | 982399 |
| rs146007972 | 15 | 42886739 A | G | 630579 |
| rs546146429 | 15 | 42961977 G | A | 859954 |
| rs576523100 | 15 | 42992120 G | T | 360129 |
| rs544593985 | 15 | 42909007 C | T | 1057210 |
| rs552552047 | 15 | 42874020 C | T | 793235 |
| rs143562438 | 15 | 42893927 A | G | 230552 |
| rs778100847 | 15 | 42986370 C | T | 472252 |
| rs547182963 | 15 | 43011781 C | T | 366627 |
| rs148150965 | 15 | 42924487 CT | C | 921412 |
| rs539637890 | 15 | 42970185 C | G | 930161 |
| rs532786415 | 15 | 42902395 G | T | 544487 |
| rs141930037 | 15 | 42901502 C | G | 577971 |
| rs201346447 | 15 | 42967124 A | G | 1086155 |

|  |  |  |  |  |
| --- | --- | --- | --- | --- |
| rs547275377 | 15 | 42872870 C | T | 452773 |
| rs190820260 | 15 | 42972022 A | T | 1008313 |
| rs185622584 | 15 | 42940650 C | T | 366231 |
| rs748249971 | 15 | 42888005 T | A | 608303 |
| rs79710890 | 15 | 42953617 G | T | 1148134 |
| rs768291062 | 15 | 42973213 A | G | 652297 |
| rs549260871 | 15 | 42973275 G | A | 835720 |
| rs144400192 | 15 | 42933225 A | G | 233468 |
| rs551791019 | 15 | 42941985 C | G | 694595 |
| rs115134206 | 15 | 42943048 A | G | 269552 |
| rs544718627 | 15 | 42911991 G | T | 24984 |
| rs750361625 | 15 | 42935674 G | C | 260474 |
| rs527386875 | 15 | 42923528 C | T | 686503 |
| rs139534250 | 15 | 42928165 C | A | 1227406 |
| rs187488302 | 15 | 42941594 C | T | 910070 |
| rs374381156 | 15 | 42946538 AGAAACACACC A | A | 592782 |
| rs148223590 | 15 | 42874204 G | A | 577971 |
| rs188271981 | 15 | 42919241 A | G | 901806 |
| rs138657899 | 15 | 42919534 T | C | 663209 |
| rs562269594 | 15 | 42933234 T | A | 573113 |
| rs577240569 | 15 | 42959807 G | A | 563502 |
| rs555742898 | 15 | 42994775 T | C | 373520 |
| rs115582718 | 15 | 42996987 A | G | 802943 |
| rs73406601 | 15 | 42938683 T | C | 978590 |
| rs775814927 | 15 | 42953874 C | T | 121623 |
| rs763967566 | 15 | 42968068 A | T | 499788 |
| rs149026782 | 15 | 42959209 C | T | 578716 |
| rs151166224 | 15 | 42999982 A | G | 1132760 |
| rs144819529 | 15 | 42958167 A | G | 578716 |
| rs76473483 | 15 | 42975557 G | C | 1140679 |
| rs532387773 | 15 | 42899343 A | G | 510347 |
| rs370550265 | 15 | 42891689 C | T | 360129 |
| rs57481952 | 15 | 42992989 A | G | 746653 |
| rs79158309 | 15 | 42877590 A | G | 753566 |
| rs61707019 | 15 | 42918962 G | A | 980237 |
| rs142562373 | 15 | 42965599 C | CA | 958699 |
| rs181506969 | 15 | 42971266 G | A | 1006060 |
| rs747065117 | 15 | 43000785 C | T | 425306 |
| rs549801198 | 15 | 42897008 ATTTT | A | 442607 |
| NA | 15 | 42918201 C | T | 60123 |
| rs564632546 | 15 | 42896159 C | A | 528705 |
| NA | 15 | 42901140 C | T | 8824 |
| rs182390722 | 15 | 42930168 A | G | 1008091 |
| rs367902230 | 15 | 42935270 TATAAA | T | 230552 |
| rs577168290 | 15 | 42953432 G | A | 370390 |
| rs546990102 | 15 | 42894164 C | G | 528705 |
| rs763038765 | 15 | 42908885 C | T | 29487 |
| rs61997204 | 15 | 42986065 G | A | 309418 |
| rs140235780 | 15 | 42995538 C | T | 447288 |
| rs533124511 | 15 | 42883247 G | C | 360129 |
| rs899259002 | 15 | 42900771 A | G | 614690 |
| rs1050706910 | 15 | 42899088 A | G | 445881 |

|  |  |  |  |  |
| --- | --- | --- | --- | --- |
| rs559447218 | 15 | 42978245 A | T | 369744 |
| rs144424767 | 15 | 42884767 C | T | 631033 |
| rs529629900 | 15 | 42887224 G | A | 365285 |
| rs139196909 | 15 | 42922603 T | C | 1138226 |
| rs187874698 | 15 | 42934546 C | T | 577971 |
| rs572679906 | 15 | 42877298 G | T | 550548 |
| rs541248519 | 15 | 42917177 G | A | 594767 |
| rs534149147 | 15 | 42973156 A | G | 490227 |
| NA | 15 | 42879024 A | C | 361954 |
| rs567027783 | 15 | 42900648 T | A | 371319 |
| rs745796005 | 15 | 43001017 A | G | 431764 |
| rs566582680 | 15 | 42886495 G | A | 577971 |
| rs754491580 | 15 | 42898085 A | T | 826504 |
| rs140540731 | 15 | 42943177 C | T | 691042 |
| rs986169146 | 15 | 42950972 G | A | 462150 |
| rs114128771 | 15 | 42934908 C | T | 691142 |
| rs74012616 | 15 | 42970814 A | G | 637169 |
| rs748629198 | 15 | 42910271 G | A | 446470 |
| rs199897680 | 15 | 42975162 T | C | 430158 |
| rs545079807 | 15 | 42964221 A | G | 369062 |
| rs755722649 | 15 | 42874521 G | C | 598904 |
| rs562709108 | 15 | 42909133 A | G | 365285 |
| rs144624716 | 15 | 42959424 C | T | 1229053 |
| rs76506589 | 15 | 42990064 A | G | 929763 |
| NA | 15 | 42989408 C | T | 26547 |
| rs781370951 | 15 | 43001949 C | T | 421787 |
| rs138717526 | 15 | 42902159 T | C | 749971 |
| rs114129956 | 15 | 43008736 C | T | 594976 |
| NA | 15 | 42931548 C | T | 6702 |
| rs565034967 | 15 | 42963671 C | T | 531130 |
| rs865821954 | 15 | 42905124 G | T | 458523 |
| rs748384434 | 15 | 42989599 T | G | 578357 |
| rs535188924 | 15 | 42991899 TA | T | 259451 |
| rs149348057 | 15 | 42998848 A | G | 1130800 |
| rs149335396 | 15 | 42882419 T | G | 1161131 |
| rs781346857 | 15 | 42916524 A | G | 642491 |
| rs72711786 | 15 | 42994336 C | T | 1174000 |
| rs115532536 | 15 | 42879312 C | T | 658090 |
| rs181359744 | 15 | 42941161 C | G | 1006061 |
| rs539954662 | 15 | 42992689 T | C | 701484 |
| rs147213920 | 15 | 42984783 T | C | 583186 |
| rs572409548 | 15 | 43009738 G | A | 97025 |
| rs547797147 | 15 | 42947514 C | T | 686010 |
| rs115363065 | 15 | 42977266 G | C | 715141 |
| rs183414402 | 15 | 42978417 T | C | 875207 |
| rs570478377 | 15 | 42936300 A | G | 577971 |
| rs550267054 | 15 | 42933960 G | A | 535984 |
| rs138442121 | 15 | 42948357 C | T | 1229046 |
| rs573215252 | 15 | 42985549 A | G | 42378 |
| rs572428417 | 15 | 42948976 A | C | 370390 |
| rs569045927 | 15 | 42959642 A | G | 687262 |
| rs189692396 | 15 | 42879140 T | C | 591731 |

|  |  |  |  |  |
| --- | --- | --- | --- | --- |
| NA | 15 | 42991942 T | C | 26547 |
| rs779133096 | 15 | 43013154 C | T | 600630 |
| rs188252842 | 15 | 42872751 C | T | 752358 |
| rs573448038 | 15 | 42893064 T | C | 584325 |
| rs73406561 | 15 | 42873793 C | A | 1109122 |
| rs149354576 | 15 | 42931464 C | A | 1225395 |
| rs7162040 | 15 | 42999994 T | C | 1158457 |
| rs7179256 | 15 | 42962258 T | A | 1125491 |
| rs548248907 | 15 | 42974169 C | A | 219032 |
| rs149018034 | 15 | 42948125 C | T | 734592 |
| rs754988459 | 15 | 42871305 C | T | 202625 |
| rs759560583 | 15 | 42914523 T | C | 260504 |
| rs116602451 | 15 | 42932722 T | C | 596435 |
| rs148022476 | 15 | 42882970 G | A | 586904 |
| rs9972577 | 15 | 42910951 C | T | 661504 |
| rs1001761276 | 15 | 42919787 C | T | 589422 |
| rs751297558 | 15 | 42992954 A | G | 425687 |
| rs376018737 | 15 | 42983973 C | T | 450926 |
| rs142520634 | 15 | 42890090 G | A | 658090 |
| rs569918428 | 15 | 42982934 G | A | 361954 |
| rs150370280 | 15 | 42993191 A | G | 240083 |
| rs149616782 | 15 | 42898801 G | A | 234329 |
| NA | 15 | 42911211 A | C | 44747 |
| NA | 15 | 42955996 A | T | 60123 |
| rs142549145 | 15 | 42996712 C | T | 586788 |
| rs144697433 | 15 | 42891968 C | G | 1155478 |
| rs576914901 | 15 | 42919272 C | T | 455089 |
| NA | 15 | 42923365 A | C | 553348 |
| rs756488144 | 15 | 42911941 G | A | 542143 |
| rs765405244 | 15 | 42993620 G | A | 425687 |
| rs142508976 | 15 | 42952718 G | A | 570225 |
| rs61192504 | 15 | 42989060 C | A | 1150479 |
| rs150750050 | 15 | 43011145 C | A | 1171697 |
| rs577962095 | 15 | 42908997 G | A | 373520 |
| rs562680697 | 15 | 43001428 T | TAC | 83699 |
| rs140183846 | 15 | 42916970 C | A | 618032 |
| rs115549874 | 15 | 42958108 G | A | 691043 |
| rs116067069 | 15 | 42888213 A | G | 664007 |
| rs189150192 | 15 | 42944420 C | T | 739340 |
| rs749066345 | 15 | 42880005 C | T | 467805 |
| rs140192774 | 15 | 42881857 C | T | 671143 |
| rs551812998 | 15 | 42887789 T | C | 236023 |
| rs770886983 | 15 | 42894934 A | G | 652962 |
| rs545235526 | 15 | 42985103 G | A | 589809 |
| rs116809323 | 15 | 42868836 G | C | 596435 |
| rs78594152 | 15 | 42901445 G | A | 688657 |
| rs549202838 | 15 | 42918182 C | T | 605394 |
| rs77336982 | 15 | 42999952 C | T | 1137791 |
| rs112377990 | 15 | 42928463 C | T | 1143239 |
| rs555887680 | 15 | 42954944 T | C | 506474 |
| rs555144614 | 15 | 42980924 A | G | 366613 |
| rs570354852 | 15 | 42884271 G | GTGATC | 685275 |

|  |  |  |  |  |
| --- | --- | --- | --- | --- |
| rs1038998860 | 15 | 42872719 A | T | 421470 |
| rs578043321 | 15 | 42942579 A | C | 396470 |
| rs76787264 | 15 | 42983155 G | A | 704711 |
| rs140017728 | 15 | 42921751 A | G | 574002 |
| rs762148444 | 15 | 42929474 G | A | 632732 |
| rs60472854 | 15 | 42969243 G | A | 605368 |
| rs574640812 | 15 | 42972264 A | G | 30094 |
| rs61732534 | 15 | 42985873 C | T | 1191546 |
| rs114743946 | 15 | 42896913 T | C | 746284 |
| rs563800652 | 15 | 42995026 T | C | 542401 |
| rs114547623 | 15 | 42940503 G | A | 693537 |
| rs530454876 | 15 | 42952699 G | A | 517915 |
| rs866924731 | 15 | 42869204 T | G | 29487 |
| rs576898675 | 15 | 42900931 A | G | 360129 |
| rs545539486 | 15 | 42893768 C | T | 712039 |
| rs191342069 | 15 | 42968129 C | T | 674466 |
| rs114863961 | 15 | 42967544 G | A | 389416 |
| rs533919230 | 15 | 42922349 G | A | 613654 |
| rs763317493 | 15 | 42905383 ACTC | A | 425687 |
| rs202110998 | 15 | 42984712 G | A | 743032 |
| rs541403134 | 15 | 42933835 A | C | 577971 |
| rs151260934 | 15 | 42951968 T | G | 577971 |
| rs141349705 | 15 | 42970404 C | T | 629334 |
| rs763159117 | 15 | 42949738 A | C | 430804 |
| rs75021400 | 15 | 42981377 C | T | 881030 |
| rs554263361 | 15 | 42991801 C | T | 635779 |
| rs534849153 | 15 | 42914548 C | CCTCT | 813311 |
| rs141012846 | 15 | 42931366 A | G | 384351 |
| rs190352047 | 15 | 42878577 C | G | 382088 |
| rs967418990 | 15 | 42893416 G | T | 494170 |
| rs771779746 | 15 | 42982372 C | G | 644458 |
| rs561481835 | 15 | 42979721 A | G | 389249 |
| rs115491632 | 15 | 42982583 G | T | 320412 |
| NA | 15 | 42939984 T | G | 117036 |
| rs533481642 | 15 | 42947115 A | G | 23657 |
| rs543709938 | 15 | 42941372 C | T | 897265 |
| rs114829780 | 15 | 42940955 T | C | 718707 |
| rs146774151 | 15 | 42982996 G | C | 587286 |
| rs118027222 | 15 | 42908213 G | A | 1158949 |
| rs545978244 | 15 | 42868026 C | A | 374919 |
| NA | 15 | 42887096 A | G | 19925 |
| rs541589109 | 15 | 42900851 A | G | 1155601 |
| rs544284211 | 15 | 42957625 T | A | 384109 |
| rs189297009 | 15 | 42955222 A | G | 697408 |
| rs140924205 | 15 | 42977810 T | G | 318351 |
| rs140746565 | 15 | 42978102 C | T | 318351 |
| rs761997441 | 15 | 42900978 T | C | 766119 |
| rs180793543 | 15 | 42997839 G | A | 578803 |
| NA | 15 | 42930850 A | T | 27798 |
| rs780649253 | 15 | 42933552 A | G | 436478 |
| rs539460492 | 15 | 42936407 C | G | 577971 |
| rs565647014 | 15 | 42947538 A | G | 369744 |

|  |  |  |  |  |
| --- | --- | --- | --- | --- |
| rs78373841 | 15 | 42989002 A | G | 1153633 |
| rs148382999 | 15 | 42942895 A | G | 684520 |
| rs534058064 | 15 | 42881762 G | T | 596047 |
| rs183088570 | 15 | 42908063 T | C | 192801 |
| rs541989351 | 15 | 42993931 G | A | 627043 |
| rs75548024 | 15 | 42887849 A | G | 455954 |
| rs117807571 | 15 | 42932609 C | T | 379753 |
| rs531332837 | 15 | 42907087 C | T | 645170 |
| rs561000481 | 15 | 42919070 C | CGTCT | 577971 |
| rs564833722 | 15 | 42919265 G | A | 290954 |
| rs138257631 | 15 | 42921324 C | A | 1205551 |
| rs553920574 | 15 | 42924012 G | C | 290954 |
| rs145900049 | 15 | 42978144 A | T | 577971 |
| NA | 15 | 42987552 G | A | 33317 |
| rs184935062 | 15 | 42887479 T | G | 440562 |
| NA | 15 | 42977820 C | A | 29487 |
| rs143587129 | 15 | 42982912 A | G | 577971 |
| rs201252548 | 15 | 43011726 G | A | 601147 |
| rs564138574 | 15 | 43007719 T | A | 383473 |
| rs552109201 | 15 | 42885029 T | C | 360128 |
| rs76465418 | 15 | 42887258 C | A | 715957 |
| rs140413699 | 15 | 42924271 C | G | 1142640 |
| rs9806648 | 15 | 43007688 G | A | 677372 |
| rs181996302 | 15 | 42887884 T | C | 733513 |
| rs185791709 | 15 | 42888179 C | T | 290954 |
| rs936199949 | 15 | 42893312 C | T | 453239 |
| NA | 15 | 42973934 G | T | 501254 |
| rs142780671 | 15 | 42910004 T | C | 231384 |
| rs973227743 | 15 | 42943691 C | T | 456974 |
| rs555233136 | 15 | 42960377 T | C | 744809 |
| rs200568949 | 15 | 42978892 A | G | 940611 |
| rs569639072 | 15 | 42895430 C | G | 290954 |
| rs187519157 | 15 | 42904780 T | C | 290954 |
| rs143568497 | 15 | 42920277 C | T | 1189764 |
| rs551667710 | 15 | 42960145 A | C | 582003 |
| rs547010092 | 15 | 42893819 G | A | 842093 |
| rs568463978 | 15 | 42932447 C | T | 290954 |
| rs556659632 | 15 | 42933766 C | T | 370942 |
| rs535923283 | 15 | 42869430 C | T | 938610 |
| rs557804844 | 15 | 42883641 G | A | 360128 |
| rs149080202 | 15 | 42966135 G | A | 234794 |
| rs571444533 | 15 | 42918107 T | C | 830781 |
| rs142940493 | 15 | 42931797 C | T | 596435 |
| rs1033036112 | 15 | 42933713 C | T | 245371 |
| rs182392365 | 15 | 42989697 G | A | 374658 |
| rs80347429 | 15 | 42875246 A | G | 713519 |
| rs531519065 | 15 | 42909557 A | G | 906847 |
| rs539505032 | 15 | 42937661 G | T | 524835 |
| rs201393731 | 15 | 42979099 C | T | 752348 |
| rs8036993 | 15 | 43010887 G | A | 644559 |
| rs191659987 | 15 | 42874061 A | G | 564752 |
| rs570880299 | 15 | 42883424 G | A | 512573 |

|  |  |  |  |  |
| --- | --- | --- | --- | --- |
| rs535717612 | 15 | 42909006 A | G | 981968 |
| rs188648617 | 15 | 42969278 G | A | 358448 |
| rs544793721 | 15 | 42979703 A | G | 379754 |
| rs763235317 | 15 | 42966791 T | C | 640932 |
| rs146027773 | 15 | 43009197 C | T | 875368 |
| rs190350231 | 15 | 42892087 G | A | 834573 |
| rs775556799 | 15 | 42902599 T | G | 367080 |
| rs185998838 | 15 | 42971310 T | G | 290954 |
| rs544175764 | 15 | 42971579 G | T | 290954 |
| rs765293647 | 15 | 42921511 C | A | 508497 |
| rs114621323 | 15 | 42924567 C | T | 687367 |
| rs553674710 | 15 | 43004059 T | C | 586043 |
| rs553908774 | 15 | 43005841 C | T | 361456 |
| rs147888154 | 15 | 42877151 G | A | 649875 |
| rs114641497 | 15 | 42972198 C | T | 318351 |
| rs561303050 | 15 | 42932406 G | A | 498142 |
| rs148862329 | 15 | 42953372 A | G | 1176813 |
| rs535868330 | 15 | 42916538 TATAAC | T | 484086 |
| rs1054031892 | 15 | 42910298 A | G | 375177 |
| rs189020874 | 15 | 42954642 C | T | 397197 |
| rs532648900 | 15 | 42992572 G | A | 87954 |
| rs115569250 | 15 | 42994390 C | T | 684316 |
| rs10163179 | 15 | 42879555 G | C | 713519 |
| rs181793901 | 15 | 42891461 C | T | 1034177 |
| rs562713483 | 15 | 42946939 A | G | 365285 |
| rs183754250 | 15 | 43005370 G | C | 1140689 |
| rs568687584 | 15 | 42986564 G | A | 82045 |
| rs532261184 | 15 | 42906837 T | A | 373521 |
| rs569243432 | 15 | 42937646 C | T | 1191454 |
| rs201351089 | 15 | 42943656 A | AT | 616372 |
| rs182385853 | 15 | 43009489 C | T | 1217468 |
| rs77171961 | 15 | 42882895 G | C | 656513 |
| rs183598354 | 15 | 42902427 T | G | 599665 |
| NA | 15 | 42882219 C | T | 74937 |
| rs575026169 | 15 | 42960629 A | G | 470540 |
| rs545585433 | 15 | 42935707 A | G | 243011 |
| rs9652418 | 15 | 42892728 T | C | 1022169 |
| rs138246348 | 15 | 42898800 C | T | 581748 |
| rs112267992 | 15 | 42895540 G | A | 577971 |
| rs79816516 | 15 | 42940199 A | G | 1144650 |
| rs189914718 | 15 | 42962288 A | G | 365285 |
| rs191913662 | 15 | 42998509 T | C | 290954 |
| rs780911612 | 15 | 42985057 C | T | 500462 |
| rs550493975 | 15 | 42995846 C | T | 647306 |
| rs547496925 | 15 | 42883035 G | T | 830404 |
| rs750639084 | 15 | 42955218 G | A | 483288 |
| rs550193589 | 15 | 42966620 GC | G | 1021913 |
| rs145410785 | 15 | 42886573 G | GT | 586904 |
| rs775555339 | 15 | 42899654 C | G | 515373 |
| rs570324067 | 15 | 42974474 G | A | 516136 |
| rs189897944 | 15 | 42893598 C | T | 650271 |
| rs115488672 | 15 | 43007595 A | C | 610199 |

|  |  |  |  |  |
| --- | --- | --- | --- | --- |
| rs749762665 | 15 | 42911969 T | C | 616956 |
| rs143677353 | 15 | 42971296 G | C | 318351 |
| rs536754439 | 15 | 42947912 A | C | 219280 |
| rs140027750 | 15 | 43001862 G | C | 628400 |
| rs147467293 | 15 | 42916903 C | T | 599568 |
| rs545441576 | 15 | 42930749 G | A | 365285 |
| rs143394146 | 15 | 42986676 A | G | 265704 |
| rs182535861 | 15 | 42908849 G | C | 584048 |
| rs115422270 | 15 | 42871599 G | C | 596435 |
| rs563472814 | 15 | 42883775 T | C | 16054 |
| rs140219968 | 15 | 42969456 C | T | 639752 |
| rs887227858 | 15 | 42924342 T | G | 456161 |
| rs551458553 | 15 | 42936471 C | A | 754632 |
| rs114643981 | 15 | 42977460 C | T | 617415 |
| rs370372581 | 15 | 42869497 A | G | 817831 |
| rs149502125 | 15 | 42886418 A | G | 467061 |
| rs528087643 | 15 | 42948574 C | T | 371142 |
| rs115755581 | 15 | 42872867 C | T | 639682 |
| rs184997894 | 15 | 42889955 G | A | 723791 |
| rs185447522 | 15 | 42957031 G | A | 257431 |
| rs112240126 | 15 | 42884761 G | A | 1117742 |
| NA | 15 | 42894125 A | C | 382277 |
| rs182793324 | 15 | 42959071 G | A | 365285 |
| rs567499517 | 15 | 42879317 G | A | 383632 |
| rs143444286 | 15 | 42984256 G | A | 229855 |
| rs546129700 | 15 | 42914280 A | T | 365285 |
| rs539820661 | 15 | 42893649 C | T | 586032 |
| rs186653054 | 15 | 42918662 C | T | 440655 |
| rs755153315 | 15 | 42929877 A | G | 405932 |
| rs566949602 | 15 | 42935789 A | G | 369062 |
| rs567259695 | 15 | 42978443 G | A | 70911 |
| NA | 15 | 42988491 A | G | 509311 |
| rs57258642 | 15 | 43003656 A | T | 834747 |
| rs547679651 | 15 | 42874132 T | A | 1162972 |
| rs532715301 | 15 | 42908030 A | G | 16054 |
| rs16973384 | 15 | 42904805 T | A | 747739 |
| rs529049962 | 15 | 43007788 G | A | 577971 |
| rs539573579 | 15 | 42883444 G | C | 1132440 |
| rs539890320 | 15 | 42962107 G | T | 975115 |
| rs544006711 | 15 | 43010245 G | A | 364588 |
| rs76471889 | 15 | 42909363 A | G | 659927 |
| rs750810228 | 15 | 42948041 A | G | 65118 |
| rs141248078 | 15 | 42879852 C | G | 230552 |
| rs78898799 | 15 | 42888226 G | A | 727791 |
| rs540721584 | 15 | 42975513 G | T | 606266 |
| rs142416794 | 15 | 42917406 G | A | 667346 |
| rs752564285 | 15 | 42920688 A | G | 512535 |
| rs116360099 | 15 | 43007017 G | A | 638096 |
| rs546865775 | 15 | 42922076 A | G | 24858 |
| rs763432324 | 15 | 43002932 G | A | 519487 |
| rs73406562 | 15 | 42877863 C | T | 1114172 |
| rs370868295 | 15 | 42898953 C | T | 581748 |

|  |  |  |  |  |
| --- | --- | --- | --- | --- |
| rs538275463 | 15 | 42921518 A | G | 365285 |
| rs138641056 | 15 | 42907396 G | C | 747739 |
| rs186242531 | 15 | 42880107 C | T | 120271 |
| rs147467293 | 15 | 42916903 C | A | 560837 |
| rs138113203 | 15 | 42874407 C | G | 577971 |
| rs191875909 | 15 | 42901586 T | A | 596435 |
| rs115827874 | 15 | 42874859 C | T | 577971 |
| rs545866797 | 15 | 42909991 G | A | 365285 |
| rs557571324 | 15 | 42894876 C | G | 553330 |
| rs181989685 | 15 | 42869480 C | T | 630096 |
| rs141314082 | 15 | 42900961 A | T | 596435 |
| rs770986137 | 15 | 42911151 T | C | 489349 |
| rs188247542 | 15 | 42999995 G | T | 818084 |
| rs758910906 | 15 | 42958938 G | A | 656955 |
| rs143795631 | 15 | 42901120 T | C | 596435 |
| rs117352983 | 15 | 42956718 T | C | 879738 |
| rs113194001 | 15 | 42981563 G | A | 453668 |
| rs530071996 | 15 | 42991882 T | C | 365285 |
| rs145017779 | 15 | 42898095 A | G | 596435 |
| rs191081200 | 15 | 42952913 C | T | 246437 |
| rs181164895 | 15 | 42936910 C | T | 606200 |
| rs78261202 | 15 | 43003113 G | T | 230552 |
| rs538043621 | 15 | 42911052 G | A | 654785 |
| rs74352881 | 15 | 42972364 A | C | 1231940 |
| NA | 15 | 42894048 T | A | 390459 |
| rs527955560 | 15 | 42960688 A | G | 230614 |
| rs116478053 | 15 | 42963301 C | G | 684873 |
| rs533842483 | 15 | 42979463 G | A | 357299 |
| rs530613898 | 15 | 42982113 C | T | 364588 |
| rs565851473 | 15 | 43005664 TACTC | T | 454028 |
| rs4924688 | 15 | 42910548 C | T | 691556 |
| rs765465286 | 15 | 42920641 A | G | 552624 |
| rs752536103 | 15 | 42954307 C | T | 500483 |
| rs192131974 | 15 | 42963378 T | C | 684872 |
| rs563762546 | 15 | 42968248 T | G | 840217 |
| NA | 15 | 42990765 C | T | 76062 |
| rs4923951 | 15 | 43004028 G | A | 547935 |
| rs374126429 | 15 | 42918675 A | C | 881584 |
| rs769391315 | 15 | 42894159 C | T | 225971 |
| rs148698759 | 15 | 42903408 C | T | 596435 |
| rs146992306 | 15 | 42973533 T | C | 376231 |
| rs559682263 | 15 | 42878911 T | TG | 793400 |
| rs568974261 | 15 | 42929329 C | T | 577971 |
| rs192367918 | 15 | 42943122 A | G | 290954 |
| rs185723917 | 15 | 42995861 G | A | 579712 |
| rs533415918 | 15 | 42896101 A | G | 364588 |
| rs565782439 | 15 | 42994289 A | G | 365285 |
| rs577527530 | 15 | 42868980 A | G | 470888 |
| rs184366727 | 15 | 42932036 T | C | 597267 |
| rs62019355 | 15 | 42938163 G | A | 1219499 |
| rs146581067 | 15 | 42947072 A | G | 1234884 |
| rs115887309 | 15 | 42954682 T | G | 739137 |

|  |  |  |  |  |
| --- | --- | --- | --- | --- |
| rs533752199 | 15 | 42992096 G | A | 365285 |
| rs12439524 | 15 | 42886117 C | T | 1141934 |
| NA | 15 | 42910898 G | A | 39170 |
| rs764077238 | 15 | 42883475 C | T | 622329 |
| rs140917325 | 15 | 42872335 G | A | 1156051 |
| rs188650242 | 15 | 42922912 A | G | 290954 |
| rs149365302 | 15 | 42956842 G | C | 360129 |
| rs535301205 | 15 | 42956781 T | C | 543691 |
| rs149025447 | 15 | 42923445 A | G | 577971 |
| rs572787507 | 15 | 42891799 T | C | 371148 |
| rs9652417 | 15 | 42892674 T | G | 732137 |
| rs558945520 | 15 | 42898456 TC | T | 378874 |
| rs571738986 | 15 | 42933157 AC | A | 290954 |
| rs535169145 | 15 | 43002549 C | T | 371074 |
| rs143933986 | 15 | 42888244 A | G | 1157340 |
| rs1053290276 | 15 | 42913357 G | A | 370024 |
| rs77245532 | 15 | 42975619 A | G | 937202 |
| rs529979641 | 15 | 43010416 T | G | 281436 |
| rs1050368245 | 15 | 42918465 T | C | 23043 |
| rs538541542 | 15 | 42990575 A | G | 625926 |
| rs139879027 | 15 | 42998863 G | A | 596435 |
| rs555246628 | 15 | 42880764 C | T | 609268 |
| rs567976615 | 15 | 42979462 G | A | 362320 |
| rs114991203 | 15 | 42990918 T | C | 577971 |
| rs13379865 | 15 | 42898498 C | T | 750639 |
| rs544613911 | 15 | 42907635 G | A | 281857 |
| rs375469449 | 15 | 42871338 G | C | 577971 |
| rs79927491 | 15 | 42880350 A | G | 628520 |
| rs777217494 | 15 | 42920742 A | G | 651509 |
| rs925991470 | 15 | 42965660 G | A | 600035 |
| rs146753941 | 15 | 43003871 C | T | 601266 |
| rs188728159 | 15 | 42943279 A | C | 365285 |
| rs201340789 | 15 | 42982237 G | C | 1146286 |
| rs531146879 | 15 | 42889936 T | C | 1157340 |
| rs552403828 | 15 | 42951233 C | A | 577971 |
| rs144638832 | 15 | 42912991 C | T | 697335 |
| rs192863692 | 15 | 42974806 G | C | 654476 |
| rs117506318 | 15 | 42872647 C | G | 1203848 |
| NA | 15 | 42916776 T | <INS_ME_SVA> | 235010 |
| rs548431781 | 15 | 42902807 C | T | 428971 |
| rs117527257 | 15 | 42904467 C | T | 700635 |
| rs185003255 | 15 | 42909008 G | A | 577971 |
| rs149407441 | 15 | 42937825 G | A | 1239794 |
| rs190421447 | 15 | 42937413 C | T | 290954 |
| rs559254882 | 15 | 42938251 C | A | 290954 |
| rs146651970 | 15 | 42965101 C | T | 658975 |
| rs573507159 | 15 | 42990952 C | G | 584026 |
| rs142777319 | 15 | 42902570 C | A | 27798 |
| rs16957037 | 15 | 42971188 C | T | 983844 |
| rs139430686 | 15 | 42870497 C | T | 1156050 |
| rs140942120 | 15 | 42885052 C | G | 1155599 |
| rs751529122 | 15 | 42923404 A | G | 657894 |

|  |  |  |  |  |
| --- | --- | --- | --- | --- |
| rs560061012 | 15 | 42936559 C | T | 188716 |
| rs566546449 | 15 | 42978632 C | T | 365285 |
| rs140844877 | 15 | 42916792 G | C | 1185057 |
| rs192855523 | 15 | 42935587 T | A | 597267 |
| rs184832341 | 15 | 42935608 G | A | 597267 |
| rs73408701 | 15 | 42968633 T | A | 852251 |
| rs761590293 | 15 | 42882398 C | T | 537933 |
| rs115788605 | 15 | 42947772 A | G | 734592 |
| rs143860655 | 15 | 43001506 C | T | 963423 |
| rs561463615 | 15 | 42881440 G | T | 365285 |
| rs74714963 | 15 | 42881700 C | T | 643291 |
| rs72711771 | 15 | 42888777 A | T | 1236611 |
| rs141601398 | 15 | 42927449 A | C | 618032 |
| rs113572612 | 15 | 42917539 T | G | 528942 |
| rs188759214 | 15 | 42932877 G | A | 553282 |
| rs188602770 | 15 | 42941109 T | A | 1204042 |
| rs546811853 | 15 | 42942308 G | A | 362320 |
| rs1877138 | 15 | 42960674 A | G | 745758 |
| rs146134970 | 15 | 42932113 G | C | 687366 |
| rs114995589 | 15 | 42961882 G | A | 687992 |
| rs559815648 | 15 | 42886131 T | G | 377764 |
| rs369695076 | 15 | 43002057 TTTCTC | T | 728707 |
| rs188799560 | 15 | 42985187 G | A | 578803 |
| rs188976332 | 15 | 43005416 C | G | 230552 |
| rs1006228942 | 15 | 42932049 A | G | 29487 |
| rs140632314 | 15 | 42941228 C | T | 1232196 |
| rs146135977 | 15 | 42889446 A | G | 1157341 |
| rs77371649 | 15 | 42894363 G | A | 659980 |
| rs16957002 | 15 | 42961681 G | A | 687992 |
| rs145526987 | 15 | 42997089 A | C | 252909 |
| rs113625023 | 15 | 42886168 T | C | 567394 |
| rs570397733 | 15 | 42925184 CA | C | 1049471 |
| rs7175580 | 15 | 42959671 G | A | 234899 |
| rs574401311 | 15 | 42991813 G | C | 585964 |
| rs776011349 | 15 | 43005955 A | G | 451417 |
| rs28537072 | 15 | 42958484 C | A | 1132277 |
| rs539933616 | 15 | 42925191 A | G | 1052360 |
| rs538709234 | 15 | 43005421 C | T | 439534 |
| rs144132229 | 15 | 43012462 G | A | 652871 |
| rs577790650 | 15 | 42876033 C | T | 606740 |
| rs544010780 | 15 | 42941389 G | A | 585653 |
| rs568962893 | 15 | 42978912 C | T | 374662 |
| rs776613042 | 15 | 42893979 C | T | 567394 |
| rs142117976 | 15 | 42895399 T | A | 1218556 |
| rs116205969 | 15 | 42912809 A | G | 697335 |
| rs577645292 | 15 | 43006346 C | T | 946709 |
| rs563005706 | 15 | 42875064 A | G | 364588 |
| rs561343338 | 15 | 42875196 T | C | 364588 |
| rs948178615 | 15 | 42895090 A | G | 29487 |
| rs144482277 | 15 | 42938217 G | C | 600211 |
| rs561258261 | 15 | 42936422 C | T | 369062 |
| rs532933640 | 15 | 42910158 A | G | 375149 |

|  |  |  |  |  |
| --- | --- | --- | --- | --- |
| rs16973395 | 15 | 42912191 T | C | 698911 |
| rs531652303 | 15 | 42915607 A | G | 602876 |
| rs116936632 | 15 | 42924234 G | T | 1210288 |
| rs558007975 | 15 | 42930036 C | A | 1191776 |
| rs145566791 | 15 | 42960083 C | T | 639752 |
| rs78906041 | 15 | 42986552 C | T | 716180 |
| rs144619496 | 15 | 42882183 G | A | 240083 |
| rs74670147 | 15 | 42894846 G | A | 735099 |
| rs188615523 | 15 | 42960991 G | T | 894155 |
| rs183757978 | 15 | 42976040 C | T | 371770 |
| NA | 15 | 42875223 T | C | 539571 |
| rs150633440 | 15 | 42916646 A | G | 361618 |
| rs139919565 | 15 | 42994529 G | A | 610199 |
| rs531877342 | 15 | 43007279 A | C | 1010830 |
| rs139887123 | 15 | 43006556 G | A | 473664 |
| rs529982887 | 15 | 42878999 A | G | 498050 |
| rs556503870 | 15 | 42903941 A | T | 625484 |
| rs192611335 | 15 | 42994690 G | A | 590250 |
| rs181607501 | 15 | 42902975 C | G | 784616 |
| rs907021416 | 15 | 42917347 C | T | 29487 |
| rs370739059 | 15 | 42991702 C | G | 610199 |
| rs397729129 | 15 | 42929277 TA | T | 719060 |
| rs113515483 | 15 | 42938045 T | G | 1144338 |
| rs552442784 | 15 | 42943859 C | T | 365285 |
| rs16957031 | 15 | 42969114 G | C | 937725 |
| rs769973104 | 15 | 42989600 T | C | 575686 |
| rs143163723 | 15 | 43007526 C | T | 1235523 |
| rs186882062 | 15 | 42936652 C | G | 600212 |
| rs78081991 | 15 | 42990078 T | C | 691249 |
| rs539580881 | 15 | 42875976 T | A | 365285 |
| rs74012611 | 15 | 42955459 T | C | 240842 |
| rs184795205 | 15 | 42974899 C | T | 1043198 |
| rs116406869 | 15 | 42934371 T | G | 596435 |
| rs112458684 | 15 | 42951912 C | T | 1225224 |
| rs528814808 | 15 | 42961066 G | GT | 360129 |
| rs2136902 | 15 | 42993387 A | G | 699839 |
| rs568646935 | 15 | 42894468 C | A | 498333 |
| rs894513132 | 15 | 42911619 C | T | 370024 |
| rs573673780 | 15 | 42979056 C | G | 384150 |
| rs530700596 | 15 | 43002359 G | A | 73112 |
| rs564485063 | 15 | 42868879 T | A | 472132 |
| rs177111135 | 15 | 42943857 G | A | 907662 |
| rs554997165 | 15 | 43012974 C | T | 369744 |
| rs117217576 | 15 | 42954954 T | C | 1168474 |
| rs115067542 | 15 | 42979733 G | A | 577971 |
| rs183110458 | 15 | 42893272 C | G | 1007709 |
| rs186462510 | 15 | 43005172 A | G | 370390 |
| rs201701873 | 15 | 42877759 A | G | 1160173 |
| rs567751521 | 15 | 42874434 A | C | 365285 |
| rs149780137 | 15 | 42947732 A | G | 1046954 |
| rs374659128 | 15 | 42983017 TTAA | T | 586032 |
| rs541324309 | 15 | 42907683 AT | A | 52673 |

|  |  |  |  |  |
| --- | --- | --- | --- | --- |
| rs570361618 | 15 | 42909402 C | T | 365285 |
| rs572122353 | 15 | 42981744 C | G | 488328 |
| rs189480172 | 15 | 42993596 T | A | 577971 |
| rs79147516 | 15 | 42884438 A | G | 643291 |
| rs920939645 | 15 | 42947341 G | A | 488890 |
| rs183802733 | 15 | 42955055 C | T | 724733 |
| rs141905011 | 15 | 42907400 G | A | 230552 |
| rs187486986 | 15 | 42924731 T | C | 375097 |
| rs573571036 | 15 | 42997345 G | C | 365285 |
| rs537145859 | 15 | 42918480 CTCCTCT | C | 680990 |
| rs115595275 | 15 | 42931033 A | T | 1044045 |
| rs538593137 | 15 | 42898914 C | T | 369062 |
| rs188636270 | 15 | 42884194 A | T | 141120 |
| rs540545255 | 15 | 42935552 G | A | 661188 |
| rs535848692 | 15 | 42936777 A | C | 468259 |
| rs535436686 | 15 | 42947292 C | T | 377567 |
| rs118052909 | 15 | 42953376 C | G | 501345 |
| rs185165874 | 15 | 42982303 G | C | 361266 |
| rs570255301 | 15 | 42940956 TC | T | 248165 |
| rs553626612 | 15 | 42973738 C | G | 365828 |
| rs549071019 | 15 | 42977706 A | G | 594812 |
| rs116670090 | 15 | 43005168 G | A | 610199 |
| rs537317862 | 15 | 42903870 G | A | 584685 |
| rs75974156 | 15 | 42876745 A | G | 773960 |
| rs187999553 | 15 | 42908886 G | A | 1152498 |
| rs149356155 | 15 | 43009603 T | C | 1048630 |
| rs572711131 | 15 | 42895174 G | T | 788125 |
| rs550161682 | 15 | 42947809 T | A | 622985 |
| rs940605313 | 15 | 42959877 A | C | 521763 |
| rs193137901 | 15 | 43006356 G | A | 944024 |
| rs16956932 | 15 | 42926883 A | G | 1011051 |
| rs573239574 | 15 | 42936686 G | A | 866320 |
| rs8028863 | 15 | 43012764 T | G | 835679 |
| rs147842892 | 15 | 42896551 C | T | 676615 |
| rs530331777 | 15 | 42955899 A | G | 813677 |
| rs1048283382 | 15 | 42872712 T | C | 24128 |
| rs1877137 | 15 | 42960383 G | T | 745758 |
| rs114646969 | 15 | 43000363 T | A | 610199 |
| rs140427579 | 15 | 42903954 A | T | 366861 |
| rs369242363 | 15 | 42978186 C | T | 708107 |
| rs540506511 | 15 | 42900849 G | A | 498651 |
| rs146377113 | 15 | 42975687 A | G | 611998 |
| rs116084720 | 15 | 42999985 A | G | 251077 |
| rs115541077 | 15 | 43005554 A | G | 230552 |
| rs200346352 | 15 | 42872718 T | TA | 378551 |
| rs35259107 | 15 | 42953805 GCTTT | G | 931448 |
| rs762907337 | 15 | 42895562 C | T | 599992 |
| rs9972600 | 15 | 42910955 T | C | 728583 |
| rs73406582 | 15 | 42912925 C | A | 841584 |
| rs184714358 | 15 | 42950446 G | A | 578803 |
| rs538040094 | 15 | 42921600 A | G | 360129 |
| rs114103856 | 15 | 42913632 T | C | 340983 |

|  |  |  |  |  |
| --- | --- | --- | --- | --- |
| rs187401921 | 15 | 42890524 C | T | 240083 |
| rs146143514 | 15 | 42895236 G | A | 365285 |
| rs140676957 | 15 | 42912825 T | G | 228357 |
| rs34928615 | 15 | 42994757 GTCTA | G | 595949 |
| rs548633040 | 15 | 42999100 G | A | 376498 |
| rs200261102 | 15 | 42889573 A | AT | 694893 |
| rs16957052 | 15 | 42979461 C | T | 1146216 |
| rs35172521 | 15 | 42886287 GAGAC | G | 590681 |
| rs115361836 | 15 | 42965711 C | A | 738961 |
| rs144867366 | 15 | 42973022 A | G | 540498 |
| rs559429992 | 15 | 43002693 C | G | 369063 |
| rs556518794 | 15 | 42871439 A | G | 498050 |
| rs369934831 | 15 | 42955652 A | G | 584677 |
| rs760673116 | 15 | 42973701 G | T | 29487 |
| rs538866117 | 15 | 42876265 C | T | 394819 |
| rs76906844 | 15 | 42905887 G | A | 1004254 |
| rs369310511 | 15 | 42976363 C | T | 590686 |
| rs544475476 | 15 | 42905554 G | T | 268700 |
| rs143824062 | 15 | 43005682 G | A | 373521 |
| rs192461105 | 15 | 42930927 G | A | 258107 |
| rs151006266 | 15 | 42969999 G | C | 510729 |
| rs187159912 | 15 | 42984840 G | A | 391283 |
| rs536218697 | 15 | 43003903 TTTTA | T | 781934 |
| rs545599183 | 15 | 42886286 A | C | 365724 |
| rs184838385 | 15 | 42906543 A | G | 240083 |
| rs542154361 | 15 | 42908599 C | T | 242004 |
| rs181968356 | 15 | 42976786 G | A | 653226 |
| rs768258586 | 15 | 42882365 T | C | 731573 |
| rs148320957 | 15 | 42894430 A | G | 1000847 |
| rs531975840 | 15 | 42880623 G | A | 369743 |
| rs16956991 | 15 | 42958767 G | C | 745757 |
| NA | 15 | 42926750 C | T | 29487 |
| rs141047217 | 15 | 42913701 G | A | 992133 |
| rs183600272 | 15 | 42998717 C | T | 618873 |
| rs751216502 | 15 | 42876578 C | T | 196435 |
| rs544840606 | 15 | 42879972 A | G | 425687 |
| rs114142198 | 15 | 42901178 C | T | 225966 |
| rs555690816 | 15 | 42938377 T | C | 921630 |
| rs116417375 | 15 | 42997962 A | G | 618873 |
| rs185150927 | 15 | 43006447 T | C | 610199 |
| rs34081955 | 15 | 42896068 AG | A | 357299 |
| rs79165890 | 15 | 42977116 T | C | 1149543 |
| rs567963038 | 15 | 43009945 C | T | 817214 |
| rs553559102 | 15 | 42899776 A | G | 588102 |
| rs189904276 | 15 | 42941288 C | T | 27798 |
| rs534900279 | 15 | 42869452 C | T | 943818 |
| rs973355371 | 15 | 42890932 C | G | 39525 |
| rs183072726 | 15 | 42938596 A | G | 231095 |
| rs527693525 | 15 | 42946090 T | C | 1154470 |
| rs556707023 | 15 | 42956960 GAATT | G | 577971 |
| rs112111204 | 15 | 42980085 C | T | 721466 |
| rs139168421 | 15 | 42970685 G | C | 227936 |

|  |  |  |  |  |
| --- | --- | --- | --- | --- |
| rs751662687 | 15 | 42897263 A | G | 542089 |
| rs189285257 | 15 | 42916517 A | G | 606200 |
| NA | 15 | 42957668 AAC | A | 29487 |
| rs550195031 | 15 | 42918826 G | A | 230614 |
| rs116814420 | 15 | 42994820 G | A | 230552 |
| rs555525085 | 15 | 42868746 C | G | 365285 |
| rs142686033 | 15 | 42899186 C | T | 581748 |
| rs77880860 | 15 | 42939905 T | G | 817734 |
| rs138430407 | 15 | 42878735 C | G | 658089 |
| rs570528702 | 15 | 42918366 C | T | 365048 |
| rs540889150 | 15 | 42946083 G | A | 369062 |
| rs142547271 | 15 | 42948936 G | A | 807405 |
| rs148764121 | 15 | 42999844 A | G | 1047874 |
| rs550483327 | 15 | 42911559 G | A | 229680 |
| rs189260934 | 15 | 42975397 C | T | 428159 |
| rs779412480 | 15 | 42882189 C | T | 675940 |
| rs775678373 | 15 | 42898704 A | C | 414165 |
| rs185468708 | 15 | 42899154 G | A | 1087429 |
| rs533154655 | 15 | 42941845 C | T | 925646 |
| rs55651630 | 15 | 42961994 A | G | 1240208 |
| rs61746531 | 15 | 42985398 C | T | 701158 |
| rs559067105 | 15 | 42880375 T | G | 608188 |
| rs529904754 | 15 | 42912690 T | A | 577971 |
| rs569708425 | 15 | 42946670 G | C | 379382 |
| rs116032750 | 15 | 43000294 T | G | 618873 |
| rs146410393 | 15 | 43000332 C | A | 618873 |
| rs539014277 | 15 | 43004880 A | G | 577971 |
| rs144084975 | 15 | 42870498 G | T | 654313 |
| rs575145889 | 15 | 42979732 C | T | 577971 |
| rs74889135 | 15 | 42890612 T | G | 544607 |
| rs543405550 | 15 | 42922960 G | A | 391810 |
| rs115794149 | 15 | 42980441 A | G | 663865 |
| rs776239194 | 15 | 42910488 AC | A | 426633 |
| rs75888272 | 15 | 42931637 C | T | 240083 |
| rs954895078 | 15 | 42929690 G | A | 535830 |
| NA | 15 | 42885209 A | G | 32609 |
| rs189549622 | 15 | 42951754 C | T | 918090 |
| rs114158464 | 15 | 42933076 A | G | 240083 |
| rs145869433 | 15 | 42983572 G | A | 982222 |
| rs115389917 | 15 | 42928867 A | G | 240083 |
| rs535352166 | 15 | 42990423 T | C | 751253 |
| rs542369195 | 15 | 42926249 C | T | 360811 |
| rs115223182 | 15 | 42935477 C | T | 696457 |
| rs35842593 | 15 | 42985999 C | A | 1220666 |
| rs143268428 | 15 | 42934332 G | A | 240083 |
| rs150271497 | 15 | 42936208 A | G | 240083 |
| rs565747190 | 15 | 42910420 T | C | 19880 |
| rs191375699 | 15 | 42930574 G | A | 369744 |
| rs766166696 | 15 | 42946519 C | T | 188592 |

| N_studies | POOLED_ALT_A | EFFECT_SIZE | SE | pvalue |
| --- | --- | --- | --- | --- |
| 165 | 0.0872 | -0.023775 | 0.0025895 | 4.26E-20 |
| 56 | 0.00164 | -0.259824 | 0.0307984 | 3.28E-17 |
| 42 | 0.00149 | -0.245416 | 0.0340888 | 6.05E-13 |
| 72 | 0.00111 | -0.199107 | 0.0289283 | 5.87E-12 |
| 161 | 0.22 | 0.0116246 | 0.00173425 | 2.04E-11 |
| 170 | 0.194 | 0.0114777 | 0.00176097 | 7.13E-11 |
| 170 | 0.221 | 0.0108313 | 0.00166518 | 7.79E-11 |
| 170 | 0.22 | 0.0107717 | 0.00166537 | 9.93E-11 |
| 170 | 0.221 | 0.0107536 | 0.00166465 | 1.05E-10 |
| 170 | 0.128 | -0.0136906 | 0.00212148 | 1.09E-10 |
| 170 | 0.22 | 0.0107395 | 0.001665 | 1.12E-10 |
| 170 | 0.145 | -0.0125561 | 0.00197215 | 1.93E-10 |
| 170 | 0.222 | 0.0105422 | 0.00165937 | 2.11E-10 |
| 170 | 0.145 | -0.0125137 | 0.00197115 | 2.18E-10 |
| 170 | 0.146 | -0.0124582 | 0.00196906 | 2.50E-10 |
| 170 | 0.145 | -0.0124201 | 0.00196457 | 2.58E-10 |
| 170 | 0.224 | 0.0104509 | 0.00165599 | 2.77E-10 |
| 170 | 0.211 | 0.0106081 | 0.00169123 | 3.55E-10 |
| 170 | 0.212 | 0.0104654 | 0.00168886 | 5.77E-10 |
| 170 | 0.142 | -0.0123619 | 0.00200169 | 6.59E-10 |
| 170 | 0.212 | 0.0104193 | 0.00168759 | 6.66E-10 |
| 170 | 0.144 | -0.0121048 | 0.0019638 | 7.09E-10 |
| 170 | 0.142 | -0.0123342 | 0.00200272 | 7.33E-10 |
| 170 | 0.225 | 0.0101563 | 0.00165799 | 9.03E-10 |
| 170 | 0.213 | 0.0102641 | 0.00168301 | 1.07E-09 |
| 170 | 0.224 | 0.0100992 | 0.00165687 | 1.09E-09 |
| 170 | 0.223 | 0.0101407 | 0.00166383 | 1.10E-09 |
| 170 | 0.224 | 0.0101029 | 0.00166363 | 1.26E-09 |
| 170 | 0.223 | 0.0100948 | 0.00166314 | 1.28E-09 |
| 170 | 0.223 | 0.0100556 | 0.00165982 | 1.38E-09 |
| 170 | 0.212 | 0.0102149 | 0.00169357 | 1.62E-09 |
| 170 | 0.144 | -0.0119941 | 0.00199607 | 1.87E-09 |
| 170 | 0.212 | 0.0100643 | 0.00168882 | 2.53E-09 |
| 170 | 0.211 | 0.0100567 | 0.00169731 | 3.12E-09 |
| 170 | 0.212 | 0.0100499 | 0.00169638 | 3.14E-09 |
| 166 | 0.21 | 0.0102071 | 0.0017256 | 3.32E-09 |
| 170 | 0.212 | 0.0100384 | 0.00169707 | 3.32E-09 |
| 170 | 0.211 | 0.0100328 | 0.00169819 | 3.46E-09 |
| 170 | 0.212 | 0.0100186 | 0.00169675 | 3.54E-09 |
| 170 | 0.212 | 0.0100036 | 0.00169626 | 3.69E-09 |
| 166 | 0.21 | 0.010112 | 0.00172472 | 4.55E-09 |
| 119 | 0.145 | -0.0125924 | 0.00215549 | 5.16E-09 |
| 166 | 0.212 | 0.00989853 | 0.00172027 | 8.71E-09 |
| 166 | 0.212 | 0.00988807 | 0.00171991 | 8.97E-09 |
| 167 | 0.212 | 0.00986199 | 0.00171974 | 9.78E-09 |
| 166 | 0.212 | 0.00982866 | 0.00171487 | 9.96E-09 |
| 166 | 0.212 | 0.00985291 | 0.00172037 | 1.02E-08 |
| 166 | 0.212 | 0.00983773 | 0.00172066 | 1.08E-08 |
| 170 | 0.22 | 0.00941526 | 0.00166654 | 1.61E-08 |
| 166 | 0.212 | 0.00972702 | 0.00172814 | 1.82E-08 |
| 166 | 0.218 | 0.0093886 | 0.00170268 | 3.51E-08 |

|  |  |  |  |  |
| --- | --- | --- | --- | --- |
| 111 | 0.222 | 0.0101353 | 0.00186232 | 5.26E-08 |
| 144 | 0.00436 | -0.0577434 | 0.0107551 | 7.92E-08 |
| 111 | 0.183 | -0.0113758 | 0.00212295 | 8.39E-08 |
| 111 | 0.229 | 0.00968685 | 0.00185547 | 1.78E-07 |
| 170 | 0.201 | 0.00897809 | 0.00172192 | 1.85E-07 |
| 170 | 0.2 | 0.00900438 | 0.00172699 | 1.85E-07 |
| 145 | 0.00441 | -0.054669 | 0.0105647 | 2.28E-07 |
| 170 | 0.201 | 0.00890744 | 0.00172258 | 2.33E-07 |
| 170 | 0.201 | 0.00887901 | 0.00172162 | 2.50E-07 |
| 170 | 0.201 | 0.00885532 | 0.00172279 | 2.75E-07 |
| 170 | 0.2 | 0.00878366 | 0.00172272 | 3.42E-07 |
| 170 | 0.2 | 0.00875828 | 0.00172781 | 4.00E-07 |
| 170 | 0.2 | 0.00868971 | 0.00172991 | 5.08E-07 |
| 170 | 0.2 | 0.00867414 | 0.00172829 | 5.20E-07 |
| 170 | 0.189 | 0.00885753 | 0.00176492 | 5.20E-07 |
| 170 | 0.189 | 0.00885531 | 0.00176489 | 5.24E-07 |
| 170 | 0.201 | 0.00863829 | 0.00172503 | 5.51E-07 |
| 170 | 0.189 | 0.00880007 | 0.00176313 | 6.00E-07 |
| 163 | 0.2 | 0.00867647 | 0.00174628 | 6.75E-07 |
| 170 | 0.189 | 0.00873847 | 0.00176288 | 7.16E-07 |
| 169 | 0.19 | 0.00878817 | 0.00177464 | 7.34E-07 |
| 170 | 0.2 | 0.00849891 | 0.00172143 | 7.93E-07 |
| 170 | 0.19 | 0.00867648 | 0.00175912 | 8.13E-07 |
| 170 | 0.19 | 0.00863824 | 0.00175825 | 8.97E-07 |
| 170 | 0.189 | 0.00859146 | 0.00176354 | 1.11E-06 |
| 170 | 0.189 | 0.00853821 | 0.00176855 | 1.38E-06 |
| 170 | 0.189 | 0.00849365 | 0.001769 | 1.58E-06 |
| 170 | 0.189 | 0.00848851 | 0.00176806 | 1.58E-06 |
| 170 | 0.189 | 0.00848996 | 0.00176852 | 1.58E-06 |
| 170 | 0.189 | 0.00848986 | 0.00176912 | 1.60E-06 |
| 170 | 0.19 | 0.00840134 | 0.00175787 | 1.76E-06 |
| 170 | 0.189 | 0.00843503 | 0.0017673 | 1.82E-06 |
| 170 | 0.189 | 0.00842192 | 0.00176712 | 1.88E-06 |
| 170 | 0.18 | 0.00859309 | 0.00180801 | 2.01E-06 |
| 170 | 0.189 | 0.00839701 | 0.00176957 | 2.08E-06 |
| 170 | 0.189 | 0.00839278 | 0.00176843 | 2.08E-06 |
| 170 | 0.18 | 0.00849886 | 0.0018046 | 2.48E-06 |
| 157 | 0.211 | 0.0091456 | 0.00194213 | 2.49E-06 |
| 170 | 0.18 | 0.00841383 | 0.00180894 | 3.30E-06 |
| 156 | 0.0303 | 0.0193278 | 0.00417742 | 3.71E-06 |
| 166 | 0.188 | 0.008126 | 0.00181037 | 7.17E-06 |
| 170 | 0.177 | 0.00814454 | 0.00183029 | 8.59E-06 |
| 170 | 0.181 | 0.00800273 | 0.00179908 | 8.66E-06 |
| 170 | 0.177 | 0.00813866 | 0.00184517 | 1.03E-05 |
| 113 | 0.186 | 0.00902491 | 0.00205916 | 1.17E-05 |
| 154 | 0.0258 | 0.0198137 | 0.00455107 | 1.34E-05 |
| 166 | 0.179 | 0.00806578 | 0.00185335 | 1.35E-05 |
| 170 | 0.0768 | 0.0113705 | 0.00263446 | 1.59E-05 |
| 162 | 0.186 | 0.00774033 | 0.00180008 | 1.71E-05 |
| 164 | 0.0267 | 0.0185634 | 0.00432834 | 1.80E-05 |
| 33 | 0.00032 | -0.324407 | 0.0770652 | 2.56E-05 |
| 109 | 0.206 | 0.00836771 | 0.00198768 | 2.56E-05 |

|  |  |  |  |  |
| --- | --- | --- | --- | --- |
| 111 | 0.173 | 0.00875006 | 0.00217519 | 5.75E-05 |
| 153 | 0.448 | -0.00595585 | 0.00148817 | 6.28E-05 |
| 164 | 0.0251 | 0.0177403 | 0.00446335 | 7.05E-05 |
| 166 | 0.247 | -0.00641665 | 0.00161485 | 7.08E-05 |
| 126 | 0.176 | 0.00785863 | 0.0019919 | 7.97E-05 |
| 127 | 0.0411 | 0.015397 | 0.00392435 | 8.73E-05 |
| 166 | 0.267 | -0.00616501 | 0.00158199 | 9.74E-05 |
| 170 | 0.0712 | 0.0102272 | 0.00273543 | 0.000185 |
| 167 | 0.267 | -0.00586438 | 0.00157309 | 0.000193 |
| 167 | 0.281 | -0.00573612 | 0.00154445 | 0.000204 |
| 166 | 0.282 | -0.00567605 | 0.00154894 | 0.000248 |
| 50 | 0.000643 | -0.142026 | 0.0392338 | 0.000295 |
| 161 | 0.0219 | 0.017573 | 0.00487267 | 0.00031 |
| 62 | 0.000659 | -0.142241 | 0.03997 | 0.000373 |
| 112 | 0.27 | -0.00604401 | 0.00173739 | 0.000504 |
| 164 | 0.0248 | 0.0157584 | 0.00454244 | 0.000522 |
| 124 | 0.279 | -0.00569539 | 0.00167824 | 0.00069 |
| 147 | 0.00971 | 0.024021 | 0.00728139 | 0.00097 |
| 3 | 0.000745 | -0.226716 | 0.0688255 | 0.000987 |
| 164 | 0.0436 | 0.0113535 | 0.00344911 | 0.000996 |
| 2 | 0.000102 | 0.438581 | 0.137459 | 0.00142 |
| 164 | 0.0429 | 0.0110465 | 0.00347623 | 0.00148 |
| 164 | 0.0228 | 0.0145113 | 0.0047356 | 0.00218 |
| 155 | 0.0166 | 0.0173004 | 0.00565672 | 0.00223 |
| 2 | 0.000636 | -0.304045 | 0.0995285 | 0.00225 |
| 112 | 0.272 | -0.0055959 | 0.00183229 | 0.00226 |
| 86 | 0.00115 | 0.0908896 | 0.0298086 | 0.0023 |
| 161 | 0.475 | -0.00437542 | 0.00144123 | 0.0024 |
| 164 | 0.0225 | 0.0142678 | 0.00477911 | 0.00283 |
| 4 | 0.000455 | 0.2277 | 0.0767342 | 0.003 |
| 161 | 0.0597 | -0.00888635 | 0.00300064 | 0.00306 |
| 136 | 0.00553 | 0.0286885 | 0.00973541 | 0.00321 |
| 136 | 0.00555 | 0.0285994 | 0.0097177 | 0.00325 |
| 97 | 0.00231 | -0.0481244 | 0.0164357 | 0.00341 |
| 15 | 0.000227 | 0.238844 | 0.0816373 | 0.00344 |
| 21 | 0.000134 | -0.276557 | 0.0950444 | 0.00362 |
| 136 | 0.00645 | 0.0274588 | 0.0094599 | 0.0037 |
| 95 | 0.00148 | -0.0680639 | 0.0235193 | 0.0038 |
| 44 | 0.0539 | 0.0150866 | 0.00532484 | 0.00461 |
| 23 | 0.000426 | 0.182525 | 0.0648538 | 0.00489 |
| 2 | 0.000229 | 0.302228 | 0.107789 | 0.00505 |
| 3 | 0.00576 | -0.0675959 | 0.0243659 | 0.00553 |
| 131 | 0.00451 | -0.0291561 | 0.010716 | 0.00651 |
| 2 | 0.000397 | -0.280087 | 0.10301 | 0.00655 |
| 170 | 0.284 | -0.00419014 | 0.00155514 | 0.00705 |
| 101 | 0.281 | 0.00738745 | 0.0027447 | 0.00711 |
| 17 | 0.000199 | 0.195385 | 0.0727412 | 0.00723 |
| 169 | 0.133 | 0.00553222 | 0.00207934 | 0.0078 |
| 169 | 0.133 | 0.00549699 | 0.00207993 | 0.00822 |
| 2 | 0.00174 | 0.0868088 | 0.032963 | 0.00845 |
| 2 | 0.000104 | 1.05473 | 0.401155 | 0.00856 |
| 108 | 0.176 | 0.00609174 | 0.00232176 | 0.0087 |

|  |  |  |  |  |
| --- | --- | --- | --- | --- |
| 169 | 0.133 | 0.00545807 | 0.00208252 | 0.00877 |
| 111 | 0.0247 | 0.0149111 | 0.0057228 | 0.00917 |
| 10 | 9.24E-05 | -0.332713 | 0.128686 | 0.00972 |
| 4 | 0.000112 | -0.474924 | 0.186071 | 0.0107 |
| 133 | 0.00423 | -0.0305939 | 0.0121012 | 0.0115 |
| 27 | 0.000405 | -0.143562 | 0.0570538 | 0.0119 |
| 2 | 0.000385 | -0.238848 | 0.0950248 | 0.012 |
| 2 | 0.000412 | -0.230051 | 0.0926103 | 0.013 |
| 12 | 0.00136 | -0.204099 | 0.0824371 | 0.0133 |
| 17 | 0.000627 | 0.121318 | 0.0492859 | 0.0138 |
| 22 | 0.000977 | 0.131723 | 0.0539552 | 0.0146 |
| 10 | 4.87E-05 | 0.426822 | 0.175602 | 0.0151 |
| 3 | 0.000536 | 0.164522 | 0.067766 | 0.0152 |
| 12 | 0.00061 | 0.257428 | 0.106841 | 0.016 |
| 8 | 0.00029 | 0.210741 | 0.087565 | 0.0161 |
| 156 | 0.0162 | 0.0135968 | 0.00568718 | 0.0168 |
| 7 | 0.00104 | 0.311077 | 0.130236 | 0.0169 |
| 10 | 2.00E-04 | -0.22903 | 0.0967849 | 0.018 |
| 19 | 0.000847 | -0.115814 | 0.048982 | 0.0181 |
| 84 | 0.00731 | -0.0358977 | 0.0152431 | 0.0185 |
| 82 | 0.00125 | 0.0506325 | 0.0215328 | 0.0187 |
| 39 | 0.000396 | -0.104541 | 0.044502 | 0.0188 |
| 108 | 0.231 | 0.0048316 | 0.00205808 | 0.0189 |
| 6 | 0.000387 | -0.123395 | 0.052595 | 0.019 |
| 143 | 0.0108 | 0.0163152 | 0.00696429 | 0.0191 |
| 52 | 0.000644 | 0.0904913 | 0.0387068 | 0.0194 |
| 111 | 0.00247 | 0.0375436 | 0.0160761 | 0.0195 |
| 3 | 0.00605 | -0.0547919 | 0.0234865 | 0.0197 |
| 139 | 0.00823 | 0.0199871 | 0.0085912 | 0.02 |
| 6 | 0.000955 | -0.131792 | 0.0571173 | 0.021 |
| 134 | 0.00444 | -0.028971 | 0.0125597 | 0.0211 |
| 116 | 0.00213 | -0.0344552 | 0.0149688 | 0.0213 |
| 2 | 0.000384 | 0.217787 | 0.0946816 | 0.0214 |
| 54 | 0.00119 | -0.0600923 | 0.026232 | 0.022 |
| 147 | 0.00519 | 0.022244 | 0.00971949 | 0.0221 |
| 2 | 0.00373 | 0.346482 | 0.151564 | 0.0223 |
| 2 | 0.00373 | 0.34624 | 0.151743 | 0.0225 |
| 2 | 3.20E-05 | 0.710086 | 0.311558 | 0.0227 |
| 4 | 0.000377 | 0.208819 | 0.0917996 | 0.0229 |
| 6 | 0.000409 | -0.153553 | 0.0675346 | 0.023 |
| 2 | 0.000909 | 0.175518 | 0.07788 | 0.0242 |
| 99 | 0.00553 | 0.0239963 | 0.0106483 | 0.0242 |
| 28 | 0.000915 | 0.100278 | 0.0445201 | 0.0243 |
| 150 | 0.00715 | -0.0192603 | 0.00856316 | 0.0245 |
| 77 | 0.00153 | -0.0498498 | 0.0222475 | 0.025 |
| 23 | 0.000274 | 0.136853 | 0.0611222 | 0.0252 |
| 5 | 0.00308 | 0.0667708 | 0.029833 | 0.0252 |
| 151 | 0.0131 | -0.015131 | 0.00679584 | 0.026 |
| 2 | 0.000129 | 0.281115 | 0.126852 | 0.0267 |
| 110 | 0.00247 | 0.0356947 | 0.0161466 | 0.0271 |
| 128 | 0.00419 | 0.0269802 | 0.0122273 | 0.0273 |
| 27 | 0.000369 | -0.131686 | 0.0597419 | 0.0275 |

|  |  |  |  |  |
| --- | --- | --- | --- | --- |
| 139 | 0.00843 | 0.0183195 | 0.00831414 | 0.0276 |
| 137 | 0.00714 | 0.0197586 | 0.00900738 | 0.0283 |
| 113 | 0.00224 | -0.0334176 | 0.0152572 | 0.0285 |
| 4 | 0.000403 | -0.202118 | 0.0925215 | 0.0289 |
| 13 | 0.000371 | 0.159122 | 0.0728641 | 0.029 |
| 4 | 0.000403 | -0.201911 | 0.0925688 | 0.0292 |
| 7 | 0.00314 | 0.152867 | 0.0701414 | 0.0293 |
| 3 | 0.000163 | -0.346022 | 0.158868 | 0.0294 |
| 16 | 0.000174 | 0.161736 | 0.0744412 | 0.0298 |
| 2 | 0.000421 | -0.197587 | 0.0911249 | 0.0301 |
| 17 | 0.000546 | -0.10638 | 0.0490585 | 0.0301 |
| 140 | 0.00931 | 0.0162946 | 0.00752873 | 0.0304 |
| 20 | 0.000328 | -0.157303 | 0.0728438 | 0.0308 |
| 13 | 0.000359 | 0.163711 | 0.0758549 | 0.0309 |
| 5 | 0.000272 | 0.262813 | 0.121842 | 0.031 |
| 160 | 0.0205 | -0.010562 | 0.00490861 | 0.0314 |
| 3 | 0.000495 | -0.149781 | 0.0701205 | 0.0327 |
| 9 | 0.000254 | 0.202723 | 0.0949787 | 0.0328 |
| 148 | 0.00942 | 0.0164097 | 0.00773868 | 0.034 |
| 4 | 6.08E-05 | -0.583137 | 0.275431 | 0.0342 |
| 3 | 0.000135 | 0.260071 | 0.122908 | 0.0343 |
| 2 | 0.184 | 0.00743061 | 0.00352828 | 0.0352 |
| 2 | 0.00416 | 0.0634614 | 0.0303595 | 0.0366 |
| 11 | 0.00038 | 0.303295 | 0.145374 | 0.037 |
| 23 | 0.00028 | -0.144943 | 0.0700467 | 0.0385 |
| 20 | 0.000226 | 0.134444 | 0.0652245 | 0.0393 |
| 99 | 0.00739 | -0.0207904 | 0.0100923 | 0.0394 |
| 40 | 0.000395 | 0.112837 | 0.0548681 | 0.0397 |
| 3 | 0.00809 | -0.0478586 | 0.023349 | 0.0404 |
| 4 | 0.00181 | 0.238746 | 0.116591 | 0.0406 |
| 6 | 0.000458 | -0.130856 | 0.0639555 | 0.0408 |
| 166 | 0.126 | 0.004327 | 0.00211613 | 0.0409 |
| 3 | 5.67E-05 | -0.675732 | 0.330593 | 0.041 |
| 3 | 0.000583 | -0.0999599 | 0.0489723 | 0.0412 |
| 12 | 0.00161 | 0.10443 | 0.0511408 | 0.0412 |
| 21 | 4.00E-04 | 0.124074 | 0.0608193 | 0.0413 |
| 166 | 0.126 | 0.00431517 | 0.00211629 | 0.0414 |
| 4 | 9.07E-05 | -0.407506 | 0.19995 | 0.0415 |
| 2 | 0.000134 | -0.328073 | 0.161012 | 0.0416 |
| 16 | 0.000165 | 0.197167 | 0.0969296 | 0.0419 |
| 4 | 8.88E-05 | -0.405249 | 0.199819 | 0.0426 |
| 9 | 0.000212 | 0.178337 | 0.0880165 | 0.0427 |
| 10 | 0.000343 | 0.129284 | 0.0638603 | 0.0429 |
| 4 | 0.000104 | -0.411931 | 0.203717 | 0.0432 |
| 4 | 0.000148 | 0.265492 | 0.131392 | 0.0433 |
| 89 | 0.00222 | 0.0422091 | 0.0208906 | 0.0433 |
| 141 | 0.0076 | 0.0176381 | 0.00873155 | 0.0434 |
| 11 | 0.000307 | -0.325951 | 0.161482 | 0.0435 |
| 9 | 0.000536 | 0.132921 | 0.0658329 | 0.0435 |
| 154 | 0.0158 | -0.0116143 | 0.00576071 | 0.0438 |
| 40 | 0.000448 | 0.0853324 | 0.0424966 | 0.0446 |
| 12 | 0.000535 | -0.104358 | 0.0519812 | 0.0447 |

|  |  |  |  |  |
| --- | --- | --- | --- | --- |
| 108 | 0.0785 | 0.00675492 | 0.00336643 | 0.0448 |
| 15 | 0.000166 | 0.162804 | 0.0811774 | 0.0449 |
| 160 | 0.0204 | -0.00988023 | 0.00494124 | 0.0455 |
| 4 | 0.000205 | 0.179681 | 0.0903255 | 0.0467 |
| 27 | 0.000276 | 0.138262 | 0.0699909 | 0.0482 |
| 6 | 0.000459 | -0.126639 | 0.0641622 | 0.0484 |
| 2 | 0.000149 | 0.773271 | 0.39191 | 0.0485 |
| 2 | 3.29E-05 | 0.588157 | 0.299622 | 0.0496 |
| 37 | 0.00675 | -0.0344793 | 0.0175651 | 0.0497 |
| 2 | 9.76E-05 | 1.07087 | 0.545717 | 0.0497 |
| 4 | 0.000329 | -0.201081 | 0.102702 | 0.0502 |
| 4 | 0.000106 | -0.336964 | 0.172362 | 0.0506 |
| 3 | 0.000119 | -0.273672 | 0.141451 | 0.053 |
| 70 | 0.000708 | 0.069927 | 0.0362064 | 0.0534 |
| 2 | 9.59E-05 | 0.282642 | 0.14742 | 0.0552 |
| 5 | 0.000175 | -0.254184 | 0.133125 | 0.0562 |
| 3 | 6.71E-05 | -0.378141 | 0.19863 | 0.0569 |
| 8 | 0.00011 | 0.226274 | 0.118858 | 0.0569 |
| 55 | 0.000639 | 0.069955 | 0.0368475 | 0.0576 |
| 60 | 0.00118 | 0.0564714 | 0.0299514 | 0.0594 |
| 2 | 7.91E-05 | 0.436318 | 0.232141 | 0.0602 |
| 77 | 0.00183 | -0.0342885 | 0.0182844 | 0.0608 |
| 20 | 0.000616 | 0.102859 | 0.0548744 | 0.0609 |
| 55 | 0.000397 | -0.0843981 | 0.0451425 | 0.0615 |
| 17 | 0.000942 | -0.121689 | 0.0651083 | 0.0616 |
| 2 | 7.91E-05 | 0.434843 | 0.232646 | 0.0616 |
| 5 | 0.000975 | -0.0911301 | 0.0488165 | 0.0619 |
| 39 | 0.000387 | 0.0970971 | 0.0525584 | 0.0647 |
| 3 | 0.000352 | -0.111408 | 0.0604546 | 0.0654 |
| 38 | 0.000386 | 0.103743 | 0.0563112 | 0.0654 |
| 2 | 0.00296 | -0.0739472 | 0.0401444 | 0.0655 |
| 39 | 0.00044 | 0.0909501 | 0.0496177 | 0.0668 |
| 4 | 0.000169 | 0.282902 | 0.155313 | 0.0685 |
| 109 | 0.166 | 0.00430205 | 0.00236424 | 0.0688 |
| 9 | 0.000299 | -0.169479 | 0.0934436 | 0.0697 |
| 12 | 0.000217 | -0.138104 | 0.0762296 | 0.07 |
| 82 | 0.00327 | -0.0282203 | 0.0156605 | 0.0715 |
| 2 | 0.00111 | 0.104821 | 0.0581937 | 0.0717 |
| 17 | 0.000373 | -0.107148 | 0.0595329 | 0.0719 |
| 65 | 0.00458 | 0.0234244 | 0.0130301 | 0.0722 |
| 11 | 9.12E-05 | 0.242386 | 0.135833 | 0.0744 |
| 31 | 0.000468 | 0.0793022 | 0.0448794 | 0.0772 |
| 85 | 0.00241 | 0.0328688 | 0.0186207 | 0.0775 |
| 2 | 5.40E-05 | -0.401712 | 0.228766 | 0.0791 |
| 136 | 0.00655 | 0.0164308 | 0.00936677 | 0.0794 |
| 12 | 0.000306 | -0.150606 | 0.085953 | 0.0797 |
| 103 | 0.00242 | 0.0259036 | 0.0147791 | 0.0797 |
| 23 | 0.000229 | 0.131326 | 0.0754751 | 0.0819 |
| 4 | 5.20E-05 | 0.306753 | 0.176845 | 0.0828 |
| 4 | 5.38E-05 | 0.55866 | 0.322517 | 0.0832 |
| 42 | 0.0013 | -0.0578374 | 0.0334784 | 0.0841 |
| 2 | 9.37E-05 | -0.254304 | 0.148216 | 0.0862 |

|  |  |  |  |  |
| --- | --- | --- | --- | --- |
| 125 | 0.00197 | 0.0282794 | 0.0164856 | 0.0863 |
| 144 | 0.00851 | 0.0132335 | 0.0077164 | 0.0863 |
| 22 | 0.000487 | 0.0809288 | 0.0472311 | 0.0866 |
| 22 | 0.000485 | 0.0806379 | 0.047351 | 0.0886 |
| 18 | 0.000151 | 0.164604 | 0.0969084 | 0.0894 |
| 30 | 0.000427 | -0.0962392 | 0.056699 | 0.0896 |
| 8 | 0.00239 | -0.140849 | 0.0831166 | 0.0902 |
| 166 | 0.117 | 0.0036815 | 0.00217595 | 0.0907 |
| 2 | 0.00144 | -0.0615379 | 0.0364424 | 0.0913 |
| 15 | 0.00033 | -0.127925 | 0.0758716 | 0.0918 |
| 36 | 0.000465 | -0.080033 | 0.0476149 | 0.0928 |
| 10 | 0.00471 | 0.0392972 | 0.0234123 | 0.0933 |
| 13 | 9.84E-05 | -0.184982 | 0.110315 | 0.0936 |
| 5 | 9.40E-05 | 0.267354 | 0.159437 | 0.0936 |
| 11 | 9.47E-05 | -0.185485 | 0.110717 | 0.0939 |
| 6 | 0.000757 | -0.106615 | 0.0637128 | 0.0943 |
| 4 | 0.000446 | 0.125932 | 0.0754913 | 0.0953 |
| 2 | 0.000208 | -0.221667 | 0.132926 | 0.0954 |
| 81 | 0.00146 | 0.0378422 | 0.0227086 | 0.0956 |
| 143 | 0.00468 | 0.0177845 | 0.0106971 | 0.0964 |
| 36 | 0.000505 | -0.0762214 | 0.0458577 | 0.0965 |
| 125 | 0.00523 | 0.0180558 | 0.0108764 | 0.0969 |
| 3 | 8.40E-05 | 0.256028 | 0.154361 | 0.0972 |
| 19 | 0.000213 | -0.153223 | 0.092906 | 0.0991 |
| 52 | 0.00135 | -0.0465271 | 0.0282229 | 0.0992 |
| 2 | 0.000558 | 0.132058 | 0.0801973 | 0.0996 |
| 155 | 0.00774 | 0.0136732 | 0.0083036 | 0.0996 |
| 3 | 0.000179 | 0.358562 | 0.218256 | 0.1 |
| 59 | 0.000983 | 0.0483537 | 0.0293992 | 0.1 |
| 148 | 0.00828 | 0.0125081 | 0.00761473 | 0.1 |
| 4 | 0.000105 | -0.210634 | 0.128451 | 0.101 |
| 22 | 0.000405 | 0.087325 | 0.0531724 | 0.101 |
| 13 | 0.00012 | -0.126646 | 0.0771735 | 0.101 |
| 8 | 0.000134 | -0.17176 | 0.105259 | 0.103 |
| 16 | 0.000406 | -0.101218 | 0.0622342 | 0.104 |
| 2 | 0.000463 | 0.135105 | 0.0832107 | 0.104 |
| 3 | 5.49E-05 | 0.323344 | 0.199646 | 0.105 |
| 3 | 0.000226 | -0.146182 | 0.0906514 | 0.107 |
| 11 | 0.000409 | -0.0926549 | 0.0574177 | 0.107 |
| 169 | 0.118 | 0.00347611 | 0.00216715 | 0.109 |
| 29 | 0.000214 | 0.118224 | 0.0737031 | 0.109 |
| 7 | 0.000563 | -0.166875 | 0.104493 | 0.11 |
| 24 | 0.000367 | 0.0895051 | 0.0561769 | 0.111 |
| 2 | 0.000122 | -0.233802 | 0.146676 | 0.111 |
| 60 | 0.00209 | 0.0353734 | 0.0222281 | 0.112 |
| 3 | 0.00612 | -0.11115 | 0.0702519 | 0.114 |
| 16 | 0.000466 | -0.0968743 | 0.0615396 | 0.115 |
| 3 | 0.00476 | -0.0393983 | 0.0249725 | 0.115 |
| 2 | 0.000315 | 0.158058 | 0.100648 | 0.116 |
| 2 | 0.000909 | 0.102318 | 0.0650259 | 0.116 |
| 3 | 5.77E-05 | 0.389699 | 0.248008 | 0.116 |
| 148 | 0.00944 | 0.0113211 | 0.0072116 | 0.116 |

|  |  |  |  |  |
| --- | --- | --- | --- | --- |
| 103 | 0.00176 | 0.0295916 | 0.0189294 | 0.118 |
| 11 | 0.000655 | -0.0637306 | 0.0409069 | 0.119 |
| 148 | 0.00943 | 0.0112184 | 0.00720582 | 0.12 |
| 8 | 0.000172 | -0.168352 | 0.108725 | 0.122 |
| 12 | 0.000663 | -0.0613126 | 0.0397954 | 0.123 |
| 120 | 0.00194 | 0.0245247 | 0.0158819 | 0.123 |
| 2 | 0.000303 | 0.15783 | 0.102547 | 0.124 |
| 13 | 0.000489 | 0.119726 | 0.0781314 | 0.125 |
| 9 | 7.12E-05 | 0.234042 | 0.153067 | 0.126 |
| 31 | 0.000697 | -0.0637532 | 0.0416761 | 0.126 |
| 157 | 0.0093 | 0.0109379 | 0.00714552 | 0.126 |
| 149 | 0.00835 | 0.0115763 | 0.0075871 | 0.127 |
| 5 | 0.000983 | -0.164586 | 0.107858 | 0.127 |
| 2 | 0.00116 | 0.0747775 | 0.0493064 | 0.129 |
| 14 | 0.000607 | 0.0885743 | 0.0585108 | 0.13 |
| 5 | 0.000218 | -0.175624 | 0.115973 | 0.13 |
| 2 | 0.00108 | -0.0990434 | 0.0655885 | 0.131 |
| 2 | 0.000195 | 0.262706 | 0.174035 | 0.131 |
| 156 | 0.00933 | 0.0107171 | 0.00710571 | 0.131 |
| 49 | 0.00113 | 0.0479389 | 0.0318329 | 0.132 |
| 2 | 0.00024 | 0.177899 | 0.118476 | 0.133 |
| 159 | 0.00975 | 0.0106514 | 0.00708963 | 0.133 |
| 167 | 0.118 | 0.00323949 | 0.00216146 | 0.134 |
| 169 | 0.117 | 0.00322754 | 0.00215511 | 0.134 |
| 169 | 0.118 | 0.00323384 | 0.00215855 | 0.134 |
| 25 | 0.00034 | 0.0892413 | 0.0596169 | 0.134 |
| 155 | 0.00926 | 0.0106991 | 0.00713861 | 0.134 |
| 5 | 0.000134 | 0.167792 | 0.112187 | 0.135 |
| 3 | 5.40E-05 | -0.302195 | 0.202761 | 0.136 |
| 5 | 0.000258 | -0.177293 | 0.118989 | 0.136 |
| 156 | 0.00933 | 0.0106765 | 0.00716759 | 0.136 |
| 8 | 1.00E-04 | 0.205015 | 0.137803 | 0.137 |
| 38 | 0.000336 | 0.0802358 | 0.0539463 | 0.137 |
| 4 | 0.000464 | 0.140605 | 0.0946005 | 0.137 |
| 54 | 0.00957 | 0.019308 | 0.0130037 | 0.138 |
| 2 | 0.000171 | -0.393084 | 0.265576 | 0.139 |
| 4 | 0.000201 | -0.184729 | 0.124912 | 0.139 |
| 35 | 0.000445 | 0.0766111 | 0.0518904 | 0.14 |
| 156 | 0.00932 | 0.0105756 | 0.0071594 | 0.14 |
| 156 | 0.00932 | 0.0105616 | 0.00715966 | 0.14 |
| 21 | 0.000155 | -0.126378 | 0.085839 | 0.141 |
| 159 | 0.00968 | 0.0104945 | 0.00713565 | 0.141 |
| 6 | 0.000114 | -0.158702 | 0.10792 | 0.141 |
| 156 | 0.00935 | 0.0105393 | 0.00716479 | 0.141 |
| 14 | 0.000113 | 0.131167 | 0.0893822 | 0.142 |
| 2 | 0.00019 | -0.32575 | 0.222585 | 0.143 |
| 52 | 0.000403 | -0.0614974 | 0.0419509 | 0.143 |
| 156 | 0.00933 | 0.0104922 | 0.00716032 | 0.143 |
| 156 | 0.00934 | 0.0104933 | 0.00716195 | 0.143 |
| 123 | 0.117 | 0.00345534 | 0.00236268 | 0.144 |
| 4 | 0.00152 | 0.180058 | 0.123221 | 0.144 |
| 156 | 0.00933 | 0.0104734 | 0.00715994 | 0.144 |

|  |  |  |  |  |
| --- | --- | --- | --- | --- |
| 156 | 0.00934 | 0.0104566 | 0.00716197 | 0.144 |
| 11 | 0.000275 | 0.100361 | 0.0688414 | 0.145 |
| 83 | 0.00156 | -0.0281288 | 0.0193496 | 0.146 |
| 8 | 0.00493 | -0.0293968 | 0.020278 | 0.147 |
| 155 | 0.00924 | 0.0103605 | 0.00714439 | 0.147 |
| 9 | 0.000156 | 0.140335 | 0.0968916 | 0.148 |
| 10 | 0.000414 | -0.0776459 | 0.053756 | 0.149 |
| 11 | 0.00112 | -0.0980176 | 0.0681038 | 0.15 |
| 2 | 0.000315 | 0.191953 | 0.133251 | 0.15 |
| 33 | 0.00027 | 0.0852155 | 0.0591872 | 0.15 |
| 101 | 0.00149 | 0.0297059 | 0.020683 | 0.151 |
| 116 | 0.00269 | 0.0214691 | 0.0149453 | 0.151 |
| 159 | 0.0097 | 0.0102032 | 0.00711255 | 0.151 |
| 53 | 0.00951 | -0.0191062 | 0.0133219 | 0.152 |
| 155 | 0.00922 | 0.0102525 | 0.00715029 | 0.152 |
| 6 | 0.000119 | 0.184946 | 0.129443 | 0.153 |
| 159 | 0.00968 | 0.0101156 | 0.00708255 | 0.153 |
| 44 | 0.000544 | -0.0597027 | 0.041849 | 0.154 |
| 156 | 0.00929 | 0.0102075 | 0.00715977 | 0.154 |
| 4 | 0.000249 | -0.185989 | 0.13076 | 0.155 |
| 156 | 0.00717 | -0.0119949 | 0.00842474 | 0.155 |
| 159 | 0.00973 | 0.0101411 | 0.00713552 | 0.155 |
| 155 | 0.00924 | 0.010162 | 0.00714398 | 0.155 |
| 2 | 7.09E-05 | 0.486344 | 0.34266 | 0.156 |
| 125 | 0.00351 | 0.0184017 | 0.0129748 | 0.156 |
| 132 | 0.00501 | 0.0149277 | 0.0105613 | 0.158 |
| 6 | 0.00173 | 0.0459182 | 0.0325775 | 0.159 |
| 152 | 0.00932 | 0.0101061 | 0.00717874 | 0.159 |
| 123 | 0.118 | 0.00330255 | 0.00235158 | 0.16 |
| 7 | 0.000361 | -0.180595 | 0.128875 | 0.161 |
| 10 | 0.000207 | -0.148215 | 0.105752 | 0.161 |
| 155 | 0.00934 | 0.00997175 | 0.00711471 | 0.161 |
| 12 | 8.92E-05 | 0.129147 | 0.0921761 | 0.161 |
| 156 | 0.0093 | 0.0100184 | 0.00715906 | 0.162 |
| 3 | 0.00206 | -0.063758 | 0.0456849 | 0.163 |
| 10 | 0.000685 | 0.206723 | 0.148054 | 0.163 |
| 146 | 0.00527 | -0.0135496 | 0.0097274 | 0.164 |
| 155 | 0.00924 | 0.00994447 | 0.00714392 | 0.164 |
| 3 | 0.000436 | -0.134645 | 0.0971343 | 0.166 |
| 102 | 0.249 | 0.00287556 | 0.00207458 | 0.166 |
| 158 | 0.00944 | 0.00986027 | 0.00711728 | 0.166 |
| 2 | 0.000156 | -0.228477 | 0.165403 | 0.167 |
| 153 | 0.0215 | -0.00698784 | 0.00507101 | 0.168 |
| 6 | 0.00117 | 0.0596243 | 0.0432559 | 0.168 |
| 165 | 0.131 | 0.00285472 | 0.00207389 | 0.169 |
| 18 | 0.00034 | 0.0914926 | 0.0664901 | 0.169 |
| 7 | 0.000184 | -0.145691 | 0.106063 | 0.17 |
| 4 | 7.20E-05 | -0.25886 | 0.188852 | 0.17 |
| 9 | 0.000243 | -0.096196 | 0.0701515 | 0.17 |
| 2 | 0.00026 | 0.149722 | 0.109289 | 0.171 |
| 10 | 0.000108 | -0.195696 | 0.14341 | 0.172 |
| 3 | 5.87E-05 | -0.203731 | 0.150111 | 0.175 |

|  |  |  |  |  |
| --- | --- | --- | --- | --- |
| 5 | 0.000204 | -0.131266 | 0.0966744 | 0.175 |
| 13 | 0.000154 | 0.113701 | 0.0838645 | 0.175 |
| 23 | 0.000194 | 0.108019 | 0.0795614 | 0.175 |
| 23 | 0.000312 | 0.0850849 | 0.0628919 | 0.176 |
| 2 | 0.000154 | -0.223977 | 0.165538 | 0.176 |
| 130 | 0.00453 | 0.0140204 | 0.0104051 | 0.178 |
| 13 | 0.000225 | -0.0980178 | 0.0730438 | 0.18 |
| 97 | 0.0108 | -0.0133684 | 0.00997934 | 0.18 |
| 3 | 0.000134 | -0.23833 | 0.178182 | 0.181 |
| 8 | 0.000117 | -0.154858 | 0.116123 | 0.182 |
| 161 | 0.0097 | 0.00951058 | 0.00713772 | 0.183 |
| 153 | 0.0421 | 0.00507701 | 0.00380872 | 0.183 |
| 12 | 0.000259 | -0.119464 | 0.0897291 | 0.183 |
| 6 | 0.00116 | 0.0581729 | 0.0438289 | 0.184 |
| 118 | 0.00494 | 0.0145162 | 0.0109476 | 0.185 |
| 6 | 6.99E-05 | 0.213684 | 0.161166 | 0.185 |
| 5 | 0.000292 | -0.121686 | 0.0920864 | 0.186 |
| 17 | 0.000252 | -0.113581 | 0.0858772 | 0.186 |
| 130 | 0.00519 | 0.0136203 | 0.0103145 | 0.187 |
| 4 | 0.000143 | 0.282325 | 0.214197 | 0.187 |
| 4 | 0.000337 | -0.151167 | 0.114466 | 0.187 |
| 31 | 0.000337 | 0.067115 | 0.0510679 | 0.189 |
| 52 | 0.000466 | -0.0595171 | 0.0452694 | 0.189 |
| 8 | 0.00501 | -0.0264174 | 0.0201231 | 0.189 |
| 11 | 0.000732 | 0.0748156 | 0.057192 | 0.191 |
| 19 | 0.000939 | -0.0564724 | 0.043257 | 0.192 |
| 8 | 0.00149 | 0.13856 | 0.106625 | 0.194 |
| 6 | 0.00225 | -0.045672 | 0.0351685 | 0.194 |
| 8 | 0.00151 | -0.134128 | 0.103373 | 0.194 |
| 133 | 0.00504 | 0.0124195 | 0.00955649 | 0.194 |
| 10 | 0.00092 | -0.057708 | 0.0445818 | 0.196 |
| 74 | 0.00433 | 0.019157 | 0.0148046 | 0.196 |
| 3 | 0.00016 | -0.219748 | 0.170428 | 0.197 |
| 9 | 0.000235 | -0.163642 | 0.127139 | 0.198 |
| 17 | 0.000275 | -0.0839291 | 0.0653973 | 0.199 |
| 4 | 0.000122 | -0.185706 | 0.144893 | 0.2 |
| 7 | 0.000117 | 0.18717 | 0.146384 | 0.201 |
| 3 | 0.000526 | 0.119501 | 0.0935527 | 0.201 |
| 2 | 0.000132 | -0.207866 | 0.163086 | 0.202 |
| 101 | 0.0226 | 0.00851682 | 0.00667951 | 0.202 |
| 28 | 0.000272 | 0.0780335 | 0.0611631 | 0.202 |
| 27 | 0.000267 | 0.0790289 | 0.0620123 | 0.203 |
| 73 | 0.000883 | -0.0321553 | 0.0252609 | 0.203 |
| 15 | 0.000303 | 0.0788111 | 0.0623628 | 0.206 |
| 2 | 0.00463 | -0.181133 | 0.143439 | 0.207 |
| 51 | 0.00132 | 0.0385586 | 0.0305632 | 0.207 |
| 14 | 0.000675 | 0.0556153 | 0.0441374 | 0.208 |
| 30 | 0.000723 | -0.0655889 | 0.0520494 | 0.208 |
| 101 | 0.00255 | 0.0212676 | 0.016976 | 0.21 |
| 9 | 0.000252 | -0.0950203 | 0.0758152 | 0.21 |
| 16 | 0.000224 | 0.0963921 | 0.077005 | 0.211 |
| 4 | 0.000937 | 0.0652651 | 0.0522107 | 0.211 |

|  |  |  |  |  |
| --- | --- | --- | --- | --- |
| 7 | 0.000923 | -0.178547 | 0.143175 | 0.212 |
| 2 | 7.49E-05 | 0.303085 | 0.24258 | 0.212 |
| 3 | 6.14E-05 | -0.232394 | 0.186447 | 0.213 |
| 48 | 0.000526 | 0.0474033 | 0.0382517 | 0.215 |
| 2 | 0.000115 | 0.234271 | 0.188818 | 0.215 |
| 9 | 0.000307 | 0.114347 | 0.0925109 | 0.216 |
| 4 | 8.56E-05 | 0.346567 | 0.280109 | 0.216 |
| 2 | 0.000178 | 0.196642 | 0.158977 | 0.216 |
| 4 | 0.00373 | -0.0466471 | 0.0378062 | 0.217 |
| 36 | 0.000341 | -0.0609572 | 0.0495245 | 0.218 |
| 8 | 0.000168 | -0.129803 | 0.105683 | 0.219 |
| 17 | 0.000474 | 0.0705819 | 0.0574447 | 0.219 |
| 7 | 0.000223 | -0.176337 | 0.143966 | 0.221 |
| 3 | 5.17E-05 | -0.230165 | 0.188364 | 0.222 |
| 3 | 0.000103 | -0.185566 | 0.152272 | 0.223 |
| 52 | 0.000516 | -0.0465801 | 0.0382999 | 0.224 |
| 157 | 0.00921 | 0.00874963 | 0.00719403 | 0.224 |
| 7 | 0.000155 | 0.165788 | 0.136287 | 0.224 |
| 3 | 0.00013 | 0.265917 | 0.220266 | 0.227 |
| 21 | 0.00102 | -0.0895556 | 0.0741597 | 0.227 |
| 3 | 0.000565 | -0.126778 | 0.10502 | 0.227 |
| 5 | 0.000178 | 0.153906 | 0.127685 | 0.228 |
| 26 | 0.000341 | 0.0692994 | 0.0576016 | 0.229 |
| 29 | 0.000646 | -0.0568146 | 0.0471817 | 0.229 |
| 101 | 0.00223 | -0.0218744 | 0.0182006 | 0.229 |
| 4 | 0.00026 | 0.123794 | 0.103066 | 0.23 |
| 4 | 0.000211 | 0.175696 | 0.146527 | 0.231 |
| 15 | 0.00062 | 0.110268 | 0.0920701 | 0.231 |
| 71 | 0.000876 | -0.0303627 | 0.0253312 | 0.231 |
| 6 | 0.00417 | 0.0269409 | 0.0225195 | 0.232 |
| 11 | 0.000169 | -0.119966 | 0.100448 | 0.232 |
| 5 | 0.000509 | -0.142351 | 0.119725 | 0.234 |
| 18 | 0.00125 | 0.0384198 | 0.0323037 | 0.234 |
| 2 | 0.000397 | -0.119589 | 0.100529 | 0.234 |
| 3 | 0.000103 | -0.179885 | 0.151431 | 0.235 |
| 5 | 0.000234 | 0.122348 | 0.103168 | 0.236 |
| 140 | 0.00432 | -0.0123424 | 0.0104545 | 0.238 |
| 52 | 0.0733 | -0.00548516 | 0.00465459 | 0.239 |
| 121 | 0.00314 | 0.0160063 | 0.0136012 | 0.239 |
| 5 | 3.96E-05 | -0.217192 | 0.184615 | 0.239 |
| 9 | 0.000173 | 0.0999795 | 0.0850957 | 0.24 |
| 5 | 0.000507 | 0.102255 | 0.0870638 | 0.24 |
| 7 | 0.00043 | -0.136773 | 0.116433 | 0.24 |
| 3 | 0.000127 | 0.188722 | 0.161024 | 0.241 |
| 35 | 0.000364 | -0.0953561 | 0.0814457 | 0.242 |
| 136 | 0.00413 | -0.0125807 | 0.0107519 | 0.242 |
| 6 | 0.000228 | -0.197552 | 0.168801 | 0.242 |
| 23 | 0.00063 | -0.0621979 | 0.0532342 | 0.243 |
| 3 | 0.000709 | 0.0606574 | 0.0520976 | 0.244 |
| 2 | 5.34E-05 | -0.240124 | 0.206621 | 0.245 |
| 140 | 0.00432 | -0.0121273 | 0.0104417 | 0.245 |
| 150 | 0.00576 | 0.0106364 | 0.00916996 | 0.246 |

|  |  |  |  |  |
| --- | --- | --- | --- | --- |
| 81 | 0.00263 | -0.0209556 | 0.0181224 | 0.248 |
| 80 | 0.00158 | 0.0247688 | 0.0214285 | 0.248 |
| 3 | 0.00267 | -0.110315 | 0.0954456 | 0.248 |
| 4 | 0.000219 | 0.111092 | 0.0963412 | 0.249 |
| 29 | 0.000626 | 0.0580898 | 0.0505239 | 0.25 |
| 5 | 0.000276 | 0.0962009 | 0.0836624 | 0.25 |
| 4 | 0.000218 | 0.110839 | 0.0963466 | 0.25 |
| 5 | 8.50E-05 | -0.146595 | 0.127662 | 0.251 |
| 3 | 7.26E-05 | -0.238704 | 0.207872 | 0.251 |
| 144 | 0.00956 | 0.00872402 | 0.00759214 | 0.251 |
| 102 | 0.00212 | -0.0229355 | 0.0200082 | 0.252 |
| 140 | 0.00443 | -0.0118018 | 0.0103073 | 0.252 |
| 8 | 0.000283 | 0.093926 | 0.0820414 | 0.252 |
| 9 | 4.24E-05 | 0.217621 | 0.189884 | 0.252 |
| 7 | 8.43E-05 | -0.148698 | 0.130184 | 0.253 |
| 2 | 0.0457 | -0.115425 | 0.101287 | 0.254 |
| 160 | 0.0157 | -0.00679644 | 0.00597716 | 0.256 |
| 27 | 0.000483 | 0.0899372 | 0.0793386 | 0.257 |
| 76 | 0.00146 | 0.0253576 | 0.0223937 | 0.257 |
| 51 | 0.000419 | 0.0462919 | 0.0408 | 0.257 |
| 151 | 0.0335 | 0.00496837 | 0.00438222 | 0.257 |
| 2 | 0.000643 | -0.0827816 | 0.073219 | 0.258 |
| 11 | 0.000433 | 0.100745 | 0.0890569 | 0.258 |
| 16 | 0.000767 | -0.0547061 | 0.0484104 | 0.258 |
| 4 | 4.10E-05 | 0.238451 | 0.211443 | 0.259 |
| 9 | 0.000512 | -0.0911753 | 0.0807747 | 0.259 |
| 3 | 0.000241 | -0.114924 | 0.10183 | 0.259 |
| 24 | 0.000265 | 0.0709523 | 0.0629696 | 0.26 |
| 12 | 0.000616 | -0.0956605 | 0.0850149 | 0.26 |
| 84 | 0.00191 | 0.0225998 | 0.0200938 | 0.261 |
| 3 | 0.00107 | 0.0673183 | 0.0600435 | 0.262 |
| 27 | 0.00015 | -0.0923135 | 0.0822924 | 0.262 |
| 18 | 0.000476 | -0.0576388 | 0.0513722 | 0.262 |
| 5 | 9.23E-05 | 0.220773 | 0.197086 | 0.263 |
| 139 | 0.00444 | -0.0115274 | 0.0102969 | 0.263 |
| 8 | 0.0028 | 0.0870792 | 0.0777642 | 0.263 |
| 150 | 0.0289 | 0.00525361 | 0.00469418 | 0.263 |
| 4 | 0.0027 | 0.103268 | 0.093138 | 0.268 |
| 161 | 0.0111 | 0.00744027 | 0.00672396 | 0.268 |
| 159 | 0.0103 | 0.00755888 | 0.00681834 | 0.268 |
| 3 | 5.87E-05 | -0.199133 | 0.17997 | 0.269 |
| 21 | 0.00083 | -0.0421197 | 0.0382471 | 0.271 |
| 2 | 0.00115 | -0.074999 | 0.0681885 | 0.271 |
| 29 | 0.000461 | -0.0587188 | 0.0534886 | 0.272 |
| 2 | 2.94E-05 | -0.431999 | 0.393744 | 0.273 |
| 78 | 0.00519 | -0.0129977 | 0.0118682 | 0.273 |
| 83 | 0.00193 | 0.0218646 | 0.019942 | 0.273 |
| 8 | 0.00436 | -0.0675361 | 0.061801 | 0.274 |
| 34 | 0.000309 | -0.0591416 | 0.0540923 | 0.274 |
| 4 | 0.000279 | 0.114398 | 0.104741 | 0.275 |
| 3 | 0.0031 | 0.0528781 | 0.0484846 | 0.275 |
| 2 | 0.000124 | 0.25551 | 0.234243 | 0.275 |

|  |  |  |  |  |
| --- | --- | --- | --- | --- |
| 5 | 0.000524 | -0.111634 | 0.102484 | 0.276 |
| 4 | 8.66E-05 | -0.144286 | 0.132444 | 0.276 |
| 24 | 0.000513 | 0.0546812 | 0.050195 | 0.276 |
| 28 | 0.00227 | 0.0394662 | 0.0362796 | 0.277 |
| 122 | 0.0196 | -0.00619446 | 0.00569686 | 0.277 |
| 123 | 0.0196 | -0.00632356 | 0.00581493 | 0.277 |
| 18 | 0.000126 | 0.105783 | 0.0973012 | 0.277 |
| 2 | 0.00171 | 0.0438522 | 0.0404475 | 0.278 |
| 7 | 0.000118 | -0.130705 | 0.120615 | 0.279 |
| 129 | 0.00387 | -0.0142543 | 0.0132298 | 0.281 |
| 87 | 0.00127 | -0.0253511 | 0.0236195 | 0.283 |
| 22 | 0.000282 | 0.0726076 | 0.0675975 | 0.283 |
| 8 | 0.000202 | 0.0947396 | 0.0885018 | 0.284 |
| 24 | 0.000256 | 0.0690775 | 0.0644935 | 0.284 |
| 2 | 7.60E-05 | -0.238062 | 0.222008 | 0.284 |
| 42 | 0.000365 | 0.0554281 | 0.0518154 | 0.285 |
| 12 | 0.000199 | -0.108077 | 0.101182 | 0.285 |
| 62 | 0.00102 | -0.0331902 | 0.0310614 | 0.285 |
| 7 | 0.000276 | 0.114441 | 0.107111 | 0.285 |
| 97 | 0.00276 | 0.0168372 | 0.0158063 | 0.287 |
| 6 | 0.000934 | -0.174237 | 0.163573 | 0.287 |
| 148 | 0.00508 | -0.0105076 | 0.00989252 | 0.288 |
| 12 | 0.000204 | -0.101367 | 0.0953747 | 0.288 |
| 7 | 0.00011 | -0.123329 | 0.116728 | 0.291 |
| 2 | 4.53E-05 | -0.27761 | 0.264446 | 0.294 |
| 147 | 0.00732 | 0.00891692 | 0.00850136 | 0.294 |
| 3 | 0.000432 | -0.0975757 | 0.0932569 | 0.295 |
| 119 | 0.00385 | 0.0135171 | 0.0129142 | 0.295 |
| 2 | 9.89E-05 | -0.223469 | 0.213679 | 0.296 |
| 67 | 0.000842 | -0.0309805 | 0.0296946 | 0.297 |
| 14 | 8.69E-05 | 0.130794 | 0.125337 | 0.297 |
| 8 | 0.000323 | 0.10284 | 0.098803 | 0.298 |
| 146 | 0.00497 | -0.0104704 | 0.0100504 | 0.298 |
| 102 | 0.00141 | -0.0205263 | 0.0197659 | 0.299 |
| 2 | 8.99E-05 | -0.21861 | 0.210871 | 0.3 |
| 106 | 0.00326 | -0.0143628 | 0.0138682 | 0.3 |
| 94 | 0.00119 | -0.022369 | 0.0215637 | 0.3 |
| 68 | 0.000919 | -0.0333266 | 0.03219 | 0.301 |
| 3 | 0.00161 | -0.146926 | 0.142242 | 0.302 |
| 5 | 0.000229 | -0.0898485 | 0.0871736 | 0.303 |
| 5 | 0.000245 | -0.145191 | 0.140838 | 0.303 |
| 3 | 0.0018 | -0.0552883 | 0.053644 | 0.303 |
| 6 | 0.000243 | 0.0972521 | 0.0945901 | 0.304 |
| 2 | 8.77E-05 | -0.215695 | 0.210197 | 0.305 |
| 2 | 7.79E-05 | 0.228267 | 0.222537 | 0.305 |
| 4 | 0.00168 | 0.0475436 | 0.0463656 | 0.305 |
| 134 | 0.00873 | 0.00821085 | 0.00800997 | 0.305 |
| 28 | 0.00022 | -0.0555483 | 0.0542073 | 0.305 |
| 5 | 0.000559 | 0.0594855 | 0.0580538 | 0.306 |
| 6 | 0.000434 | -0.118008 | 0.115636 | 0.307 |
| 32 | 0.000233 | -0.0579073 | 0.0566621 | 0.307 |
| 9 | 0.000423 | 0.0918565 | 0.0901541 | 0.308 |

|  |  |  |  |  |
| --- | --- | --- | --- | --- |
| 159 | 0.242 | -0.00172703 | 0.00169844 | 0.309 |
| 5 | 0.000241 | -0.143095 | 0.141336 | 0.311 |
| 2 | 5.43E-05 | -0.284299 | 0.280441 | 0.311 |
| 10 | 0.000287 | -0.165443 | 0.163367 | 0.311 |
| 11 | 0.00513 | 0.0477661 | 0.0471303 | 0.311 |
| 8 | 0.00123 | -0.0513182 | 0.0507754 | 0.312 |
| 8 | 0.0138 | -0.0354215 | 0.0351583 | 0.314 |
| 64 | 0.000941 | -0.0276671 | 0.0275563 | 0.315 |
| 150 | 0.00639 | -0.00893056 | 0.00888176 | 0.315 |
| 3 | 0.000105 | -0.146053 | 0.145423 | 0.315 |
| 4 | 0.000271 | 0.140026 | 0.14018 | 0.318 |
| 165 | 0.0698 | 0.00286218 | 0.00286832 | 0.318 |
| 27 | 0.000277 | -0.0621503 | 0.0623982 | 0.319 |
| 65 | 0.00135 | -0.0322784 | 0.0323753 | 0.319 |
| 3 | 9.20E-05 | 0.168203 | 0.169196 | 0.32 |
| 37 | 0.000501 | 0.0441385 | 0.0443948 | 0.32 |
| 95 | 0.0017 | -0.0178297 | 0.0179407 | 0.32 |
| 31 | 0.000568 | -0.0483475 | 0.0487593 | 0.321 |
| 2 | 0.000226 | 0.0946903 | 0.0955869 | 0.322 |
| 12 | 0.000777 | 0.0524769 | 0.0531056 | 0.323 |
| 19 | 0.000279 | -0.0587222 | 0.0595073 | 0.324 |
| 2 | 0.000227 | 0.0940747 | 0.0955199 | 0.325 |
| 154 | 0.00677 | -0.00891004 | 0.00907254 | 0.326 |
| 13 | 0.000245 | 0.100151 | 0.10225 | 0.327 |
| 8 | 0.00021 | 0.0953493 | 0.0974951 | 0.328 |
| 7 | 0.000747 | 0.0517606 | 0.0529573 | 0.328 |
| 5 | 0.0016 | -0.0293917 | 0.0300352 | 0.328 |
| 8 | 0.000746 | -0.136022 | 0.139346 | 0.329 |
| 4 | 0.000201 | -0.194017 | 0.198859 | 0.329 |
| 17 | 0.00123 | -0.0350139 | 0.035885 | 0.329 |
| 104 | 0.00808 | -0.00889947 | 0.00912001 | 0.329 |
| 26 | 0.000234 | -0.0515653 | 0.0528225 | 0.329 |
| 5 | 0.00133 | -0.0416239 | 0.0427594 | 0.33 |
| 65 | 0.0015 | 0.0253548 | 0.0260923 | 0.331 |
| 3 | 0.00067 | -0.139286 | 0.143322 | 0.331 |
| 31 | 0.00262 | 0.0287028 | 0.0295024 | 0.331 |
| 5 | 8.48E-05 | -0.133191 | 0.137774 | 0.334 |
| 103 | 0.00808 | -0.00881554 | 0.00912753 | 0.334 |
| 140 | 0.00789 | 0.00934564 | 0.00967808 | 0.334 |
| 97 | 0.00174 | -0.0171259 | 0.0177552 | 0.335 |
| 8 | 0.000746 | -0.134162 | 0.139399 | 0.336 |
| 160 | 0.0251 | 0.00448271 | 0.00465818 | 0.336 |
| 5 | 9.60E-05 | -0.12391 | 0.129169 | 0.337 |
| 160 | 0.0109 | 0.00650126 | 0.00676637 | 0.337 |
| 73 | 0.00105 | -0.025831 | 0.0269443 | 0.338 |
| 89 | 0.00135 | -0.021646 | 0.022581 | 0.338 |
| 102 | 0.00204 | -0.0184401 | 0.0192455 | 0.338 |
| 8 | 0.000155 | -0.103176 | 0.10774 | 0.338 |
| 15 | 0.000134 | -0.0691446 | 0.0721412 | 0.338 |
| 49 | 0.000582 | 0.0336475 | 0.0352612 | 0.34 |
| 16 | 0.000413 | 0.0749278 | 0.0786572 | 0.341 |
| 18 | 0.000173 | 0.0812708 | 0.0853526 | 0.341 |

|  |  |  |  |  |
| --- | --- | --- | --- | --- |
| 2 | 0.000133 | -0.188415 | 0.198426 | 0.342 |
| 2 | 0.000137 | -0.189296 | 0.199288 | 0.342 |
| 161 | 0.011 | 0.00641784 | 0.00675114 | 0.342 |
| 2 | 0.000106 | -0.183784 | 0.193915 | 0.343 |
| 135 | 0.00477 | -0.0100405 | 0.0105872 | 0.343 |
| 161 | 0.011 | 0.00638407 | 0.00674282 | 0.344 |
| 9 | 0.0121 | 0.0203039 | 0.0214365 | 0.344 |
| 117 | 0.00911 | 0.00742099 | 0.00784784 | 0.344 |
| 95 | 0.00174 | -0.0167495 | 0.017722 | 0.345 |
| 98 | 0.00174 | -0.0167194 | 0.0177304 | 0.346 |
| 95 | 0.00258 | 0.0154244 | 0.0164748 | 0.349 |
| 13 | 0.00157 | 0.027953 | 0.0298758 | 0.349 |
| 5 | 0.000728 | 0.160202 | 0.170898 | 0.349 |
| 2 | 0.00418 | 0.142605 | 0.152633 | 0.35 |
| 51 | 0.00143 | -0.0319691 | 0.0342306 | 0.35 |
| 6 | 8.29E-05 | -0.122934 | 0.132709 | 0.354 |
| 98 | 0.0164 | 0.0064247 | 0.00692842 | 0.354 |
| 3 | 6.69E-05 | 0.171307 | 0.184647 | 0.354 |
| 6 | 0.000554 | 0.11119 | 0.120088 | 0.354 |
| 4 | 0.00099 | -0.15015 | 0.162503 | 0.355 |
| 2 | 8.99E-05 | -0.212791 | 0.230711 | 0.356 |
| 42 | 0.000363 | 0.048054 | 0.0520208 | 0.356 |
| 5 | 0.00038 | 0.0762169 | 0.0824914 | 0.356 |
| 104 | 0.00153 | -0.0174523 | 0.0189579 | 0.357 |
| 25 | 0.00412 | 0.0152833 | 0.0165884 | 0.357 |
| 17 | 0.000283 | 0.0747956 | 0.0811988 | 0.357 |
| 3 | 0.000114 | 0.148619 | 0.161198 | 0.357 |
| 160 | 0.019 | -0.00493549 | 0.00536625 | 0.358 |
| 6 | 0.000792 | 0.0698267 | 0.0759784 | 0.358 |
| 6 | 0.000409 | -0.105519 | 0.115013 | 0.359 |
| 6 | 0.000137 | -0.124949 | 0.136467 | 0.36 |
| 6 | 0.000133 | -0.125088 | 0.136707 | 0.36 |
| 3 | 4.19E-05 | 0.151043 | 0.165396 | 0.361 |
| 8 | 0.000154 | 0.143418 | 0.157322 | 0.362 |
| 3 | 7.87E-05 | 0.152482 | 0.167427 | 0.362 |
| 57 | 0.00129 | -0.0273646 | 0.0301098 | 0.363 |
| 25 | 0.000499 | 0.0419076 | 0.0460722 | 0.363 |
| 94 | 0.00178 | -0.0182489 | 0.0201169 | 0.364 |
| 97 | 0.00174 | -0.0161438 | 0.0177698 | 0.364 |
| 8 | 0.000417 | 0.0699436 | 0.0770562 | 0.364 |
| 70 | 0.00255 | 0.0169354 | 0.0186941 | 0.365 |
| 3 | 8.69E-05 | -0.147077 | 0.16266 | 0.366 |
| 63 | 0.00133 | 0.0218961 | 0.0243525 | 0.369 |
| 107 | 0.00742 | -0.0084261 | 0.00939532 | 0.37 |
| 2 | 4.28E-05 | 0.202458 | 0.226139 | 0.371 |
| 2 | 6.87E-05 | 0.164029 | 0.183771 | 0.372 |
| 26 | 0.000261 | -0.0634871 | 0.0713292 | 0.373 |
| 16 | 0.000333 | 0.0742016 | 0.0832405 | 0.373 |
| 18 | 0.000426 | 0.0588402 | 0.0663977 | 0.376 |
| 14 | 0.000263 | 0.0903038 | 0.102321 | 0.377 |
| 16 | 0.000224 | -0.0793338 | 0.0897803 | 0.377 |
| 4 | 0.00013 | -0.0960366 | 0.108952 | 0.378 |

|  |  |  |  |  |
| --- | --- | --- | --- | --- |
| 92 | 0.000836 | -0.0241732 | 0.0274934 | 0.379 |
| 7 | 0.000118 | -0.113206 | 0.128606 | 0.379 |
| 35 | 0.00031 | 0.0492794 | 0.0560067 | 0.379 |
| 29 | 0.000163 | -0.0690867 | 0.0785158 | 0.379 |
| 7 | 0.000123 | -0.0797903 | 0.0906644 | 0.379 |
| 27 | 0.000246 | 0.0635516 | 0.0723754 | 0.38 |
| 43 | 0.0288 | -0.00473467 | 0.00539156 | 0.38 |
| 7 | 4.20E-05 | 0.187241 | 0.213917 | 0.381 |
| 8 | 0.00285 | -0.0661487 | 0.0755934 | 0.382 |
| 42 | 0.000829 | -0.0333523 | 0.0381363 | 0.382 |
| 53 | 0.306 | -0.00341646 | 0.00390852 | 0.382 |
| 5 | 0.000866 | 0.0419305 | 0.0479951 | 0.382 |
| 5 | 0.00052 | -0.0931681 | 0.106532 | 0.382 |
| 144 | 0.00557 | -0.00871299 | 0.0099812 | 0.383 |
| 17 | 0.000402 | 0.0461717 | 0.0529533 | 0.383 |
| 2 | 5.60E-05 | 0.183279 | 0.209997 | 0.383 |
| 145 | 0.00575 | -0.00857346 | 0.00984484 | 0.384 |
| 15 | 0.000177 | 0.0651998 | 0.0750623 | 0.385 |
| 5 | 0.00126 | -0.0463968 | 0.0533543 | 0.385 |
| 2 | 7.08E-05 | -0.167528 | 0.193157 | 0.386 |
| 26 | 0.000502 | 0.0398179 | 0.0459725 | 0.386 |
| 120 | 0.00289 | -0.01201 | 0.0138787 | 0.387 |
| 95 | 0.00173 | -0.0153507 | 0.0177581 | 0.387 |
| 4 | 0.00292 | 0.033883 | 0.0391413 | 0.387 |
| 11 | 6.36E-05 | -0.127452 | 0.147714 | 0.388 |
| 5 | 0.000243 | -0.0753127 | 0.0871867 | 0.388 |
| 97 | 0.00175 | -0.0153429 | 0.0177622 | 0.388 |
| 3 | 0.000176 | -0.111626 | 0.129234 | 0.388 |
| 4 | 0.000103 | 0.148387 | 0.17309 | 0.391 |
| 92 | 0.00116 | -0.0190934 | 0.0222352 | 0.391 |
| 88 | 0.00259 | -0.0159993 | 0.0187091 | 0.392 |
| 6 | 0.000113 | -0.111136 | 0.129963 | 0.392 |
| 2 | 0.000288 | -0.147619 | 0.17241 | 0.392 |
| 14 | 0.000164 | 0.0821054 | 0.09597 | 0.392 |
| 3 | 0.0011 | 0.0446934 | 0.052209 | 0.392 |
| 27 | 0.000722 | 0.0450406 | 0.0526838 | 0.393 |
| 2 | 0.000103 | -0.176594 | 0.20723 | 0.394 |
| 2 | 9.22E-05 | -0.173366 | 0.203724 | 0.395 |
| 2 | 9.44E-05 | -0.174405 | 0.20519 | 0.395 |
| 59 | 0.00114 | 0.0229818 | 0.0270087 | 0.395 |
| 2 | 0.000118 | -0.637101 | 0.749699 | 0.395 |
| 159 | 0.0411 | 0.00319383 | 0.00375733 | 0.395 |
| 2 | 0.000106 | -0.175963 | 0.207755 | 0.397 |
| 7 | 0.000791 | -0.124424 | 0.146997 | 0.397 |
| 2 | 8.99E-05 | -0.172186 | 0.203794 | 0.398 |
| 3 | 0.000128 | -0.130163 | 0.153865 | 0.398 |
| 3 | 0.000322 | -0.146655 | 0.173826 | 0.399 |
| 2 | 9.22E-05 | -0.171635 | 0.203844 | 0.4 |
| 137 | 0.00796 | 0.00710559 | 0.00845027 | 0.4 |
| 2 | 0.000115 | -0.176294 | 0.209651 | 0.4 |
| 3 | 8.38E-05 | -0.16111 | 0.191439 | 0.4 |
| 12 | 0.00156 | 0.0312695 | 0.0372447 | 0.401 |

|  |  |  |  |  |
| --- | --- | --- | --- | --- |
| 7 | 0.000791 | -0.123491 | 0.146998 | 0.401 |
| 13 | 0.000328 | -0.0578491 | 0.0689507 | 0.401 |
| 2 | 0.000115 | -0.175529 | 0.209783 | 0.403 |
| 62 | 0.000808 | 0.0221152 | 0.0264291 | 0.403 |
| 5 | 0.00025 | -0.101727 | 0.121909 | 0.404 |
| 2 | 8.09E-05 | -0.169487 | 0.203958 | 0.406 |
| 96 | 0.00172 | -0.0147908 | 0.0177923 | 0.406 |
| 2 | 0.00156 | -0.036082 | 0.0433962 | 0.406 |
| 65 | 0.0022 | -0.0172526 | 0.0208279 | 0.407 |
| 19 | 0.000119 | 0.0652691 | 0.078657 | 0.407 |
| 5 | 0.00122 | -0.120002 | 0.144926 | 0.408 |
| 2 | 4.35E-05 | -0.218176 | 0.264524 | 0.409 |
| 95 | 0.00174 | -0.014612 | 0.0176956 | 0.409 |
| 4 | 0.0069 | 0.014427 | 0.0174727 | 0.409 |
| 2 | 3.11E-05 | -0.214685 | 0.260654 | 0.41 |
| 75 | 0.00146 | 0.0184316 | 0.0224074 | 0.411 |
| 4 | 0.000154 | -0.110433 | 0.134665 | 0.412 |
| 94 | 0.0011 | -0.0180104 | 0.0219583 | 0.412 |
| 16 | 0.000399 | 0.0440738 | 0.0538106 | 0.413 |
| 2 | 0.000171 | -0.238599 | 0.292324 | 0.414 |
| 5 | 0.00115 | 0.113116 | 0.139245 | 0.417 |
| 78 | 0.0155 | -0.00699716 | 0.00864445 | 0.418 |
| 24 | 0.00102 | -0.0359596 | 0.044466 | 0.419 |
| 68 | 0.00216 | -0.0160724 | 0.0199002 | 0.419 |
| 99 | 0.00178 | -0.0142795 | 0.0176532 | 0.419 |
| 106 | 0.00246 | 0.0146988 | 0.0182818 | 0.421 |
| 27 | 0.000425 | -0.0335682 | 0.0417816 | 0.422 |
| 131 | 0.00552 | -0.00864192 | 0.0107642 | 0.422 |
| 49 | 0.00099 | -0.024152 | 0.0301349 | 0.423 |
| 9 | 0.00012 | -0.0894196 | 0.11158 | 0.423 |
| 100 | 0.00265 | -0.0133976 | 0.0167444 | 0.424 |
| 6 | 0.00167 | -0.0463408 | 0.05799 | 0.424 |
| 8 | 0.000412 | -0.0719553 | 0.0900766 | 0.424 |
| 2 | 0.00067 | 0.120258 | 0.150592 | 0.425 |
| 15 | 0.00193 | -0.0327452 | 0.0411441 | 0.426 |
| 2 | 7.18E-05 | -0.294465 | 0.370349 | 0.427 |
| 6 | 0.000478 | 0.0756369 | 0.095412 | 0.428 |
| 4 | 0.00119 | -0.0342846 | 0.0432226 | 0.428 |
| 39 | 0.000446 | -0.0330343 | 0.0418031 | 0.429 |
| 3 | 0.000297 | -0.0647043 | 0.081733 | 0.429 |
| 3 | 0.000138 | -0.13972 | 0.176839 | 0.429 |
| 22 | 0.000461 | -0.0386341 | 0.0488518 | 0.429 |
| 160 | 0.0213 | -0.00406517 | 0.00513688 | 0.429 |
| 8 | 9.21E-05 | -0.120715 | 0.153097 | 0.43 |
| 3 | 4.44E-05 | 0.115739 | 0.14681 | 0.43 |
| 22 | 0.000226 | -0.0521825 | 0.0662179 | 0.431 |
| 19 | 0.000279 | 0.0526593 | 0.0670753 | 0.432 |
| 103 | 0.00152 | -0.0149235 | 0.0190016 | 0.432 |
| 6 | 1.00E-04 | -0.100248 | 0.127896 | 0.433 |
| 12 | 0.000128 | 0.0705182 | 0.0899319 | 0.433 |
| 6 | 0.000359 | -0.0509265 | 0.065115 | 0.434 |
| 110 | 0.00161 | 0.0158326 | 0.0202737 | 0.435 |

|  |  |  |  |  |
| --- | --- | --- | --- | --- |
| 93 | 0.00113 | -0.016985 | 0.0217453 | 0.435 |
| 30 | 0.00157 | 0.0262204 | 0.0336461 | 0.436 |
| 3 | 0.000626 | -0.0524591 | 0.0672787 | 0.436 |
| 60 | 0.00109 | -0.0206779 | 0.0265229 | 0.436 |
| 3 | 0.000177 | -0.154511 | 0.199253 | 0.438 |
| 95 | 0.00114 | -0.0173706 | 0.0223919 | 0.438 |
| 5 | 0.00022 | 0.104137 | 0.134644 | 0.439 |
| 65 | 0.00107 | 0.02351 | 0.0303798 | 0.439 |
| 26 | 0.000356 | 0.0573773 | 0.0741769 | 0.439 |
| 4 | 0.00291 | -0.0403573 | 0.0522238 | 0.44 |
| 12 | 0.00047 | 0.0519925 | 0.0675381 | 0.441 |
| 157 | 0.0951 | 0.00197887 | 0.00256869 | 0.441 |
| 3 | 0.000151 | -0.104125 | 0.135632 | 0.443 |
| 78 | 0.00136 | -0.0182977 | 0.023838 | 0.443 |
| 3 | 0.000466 | 0.0642168 | 0.0839344 | 0.444 |
| 52 | 0.00162 | -0.0216973 | 0.0284181 | 0.445 |
| 169 | 0.0741 | -0.00209486 | 0.00274515 | 0.445 |
| 39 | 0.00213 | 0.0165686 | 0.0217589 | 0.446 |
| 4 | 0.000259 | -0.0631623 | 0.0827962 | 0.446 |
| 3 | 0.000355 | 0.135324 | 0.177919 | 0.447 |
| 94 | 0.00113 | -0.0164574 | 0.0216582 | 0.447 |
| 2 | 0.00146 | -0.0598615 | 0.0788631 | 0.448 |
| 60 | 0.00199 | 0.0159851 | 0.0210814 | 0.448 |
| 113 | 0.0108 | 0.00567069 | 0.00749159 | 0.449 |
| 2 | 8.65E-05 | -0.165083 | 0.217928 | 0.449 |
| 139 | 0.00565 | -0.00769149 | 0.0101748 | 0.45 |
| 3 | 0.000157 | -0.10252 | 0.136092 | 0.451 |
| 13 | 7.86E-05 | 0.100478 | 0.133329 | 0.451 |
| 161 | 0.012 | 0.00472764 | 0.00629161 | 0.452 |
| 2 | 8.12E-05 | 0.267485 | 0.357872 | 0.455 |
| 117 | 0.00338 | 0.00936953 | 0.0125519 | 0.455 |
| 17 | 0.000327 | -0.0649935 | 0.0872815 | 0.456 |
| 6 | 0.000125 | -0.0969557 | 0.130031 | 0.456 |
| 97 | 0.00178 | -0.0131835 | 0.0176954 | 0.456 |
| 31 | 0.000274 | -0.0436247 | 0.0587303 | 0.458 |
| 3 | 0.00114 | 0.0384814 | 0.0520215 | 0.459 |
| 18 | 0.00026 | -0.0533451 | 0.072101 | 0.459 |
| 15 | 0.000141 | -0.0515959 | 0.0698503 | 0.46 |
| 6 | 0.000216 | -0.0656116 | 0.0891533 | 0.462 |
| 2 | 5.03E-05 | -0.171796 | 0.233461 | 0.462 |
| 5 | 0.000117 | -0.0922596 | 0.125718 | 0.463 |
| 18 | 0.00024 | 0.0635437 | 0.0866654 | 0.463 |
| 52 | 0.00106 | -0.0208147 | 0.0283681 | 0.463 |
| 13 | 0.000137 | -0.0526541 | 0.0719533 | 0.464 |
| 94 | 0.00165 | -0.0170784 | 0.0233174 | 0.464 |
| 2 | 4.65E-05 | 0.207501 | 0.283883 | 0.465 |
| 3 | 0.00013 | -0.13704 | 0.187842 | 0.466 |
| 3 | 3.01E-05 | -0.163143 | 0.22382 | 0.466 |
| 44 | 0.000501 | -0.0353621 | 0.0486055 | 0.467 |
| 3 | 7.65E-05 | 0.129366 | 0.178222 | 0.468 |
| 148 | 0.00556 | -0.00712941 | 0.00984266 | 0.469 |
| 5 | 0.000148 | -0.0997207 | 0.138027 | 0.47 |

|  |  |  |  |  |
| --- | --- | --- | --- | --- |
| 5 | 0.000166 | 0.137908 | 0.191438 | 0.471 |
| 14 | 0.000183 | 0.0611372 | 0.0848901 | 0.471 |
| 27 | 0.000358 | 0.0500155 | 0.0693167 | 0.471 |
| 2 | 0.00171 | 0.0453631 | 0.0630101 | 0.472 |
| 35 | 0.000409 | -0.0363624 | 0.050545 | 0.472 |
| 4 | 0.00173 | 0.0237911 | 0.0330714 | 0.472 |
| 96 | 0.00175 | -0.0127683 | 0.017808 | 0.473 |
| 15 | 0.00014 | -0.0498645 | 0.0694848 | 0.473 |
| 170 | 0.0733 | -0.00199014 | 0.00277082 | 0.473 |
| 4 | 0.000209 | -0.0938986 | 0.131213 | 0.474 |
| 2 | 0.000122 | 0.098967 | 0.138289 | 0.474 |
| 2 | 0.000129 | -0.12533 | 0.175032 | 0.474 |
| 60 | 0.00117 | 0.0196792 | 0.0275088 | 0.474 |
| 5 | 0.000142 | -0.0990437 | 0.138774 | 0.475 |
| 12 | 0.000121 | -0.064879 | 0.0907743 | 0.475 |
| 2 | 5.71E-05 | 0.126042 | 0.176918 | 0.476 |
| 5 | 0.000148 | -0.0984883 | 0.138093 | 0.476 |
| 5 | 0.000168 | 0.0746513 | 0.104682 | 0.476 |
| 12 | 0.00012 | -0.0646584 | 0.0910007 | 0.477 |
| 117 | 0.0684 | -0.00237292 | 0.00334711 | 0.478 |
| 3 | 0.000545 | 0.0529258 | 0.0745545 | 0.478 |
| 13 | 0.000253 | -0.0697577 | 0.0982582 | 0.478 |
| 97 | 0.00143 | -0.0137557 | 0.0194024 | 0.478 |
| 9 | 0.000215 | 0.0829672 | 0.117246 | 0.479 |
| 8 | 0.000848 | 0.0981431 | 0.138487 | 0.479 |
| 17 | 0.000462 | 0.0382106 | 0.0539887 | 0.479 |
| 12 | 0.000163 | 0.0635871 | 0.0897664 | 0.479 |
| 15 | 0.000105 | 0.063749 | 0.0901364 | 0.479 |
| 4 | 0.000206 | -0.0918461 | 0.130183 | 0.48 |
| 6 | 8.62E-05 | -0.0935848 | 0.132632 | 0.48 |
| 19 | 0.000356 | 0.0384276 | 0.0545429 | 0.481 |
| 36 | 0.000437 | -0.0376575 | 0.0536927 | 0.483 |
| 34 | 0.000366 | 0.0379566 | 0.0541126 | 0.483 |
| 18 | 0.000201 | 0.0571464 | 0.0816475 | 0.484 |
| 19 | 0.000204 | 0.0570441 | 0.0815157 | 0.484 |
| 33 | 0.000674 | 0.0273825 | 0.0393679 | 0.487 |
| 63 | 0.00248 | -0.0126711 | 0.0182479 | 0.487 |
| 113 | 0.00166 | 0.0138508 | 0.0199711 | 0.488 |
| 6 | 0.00056 | -0.0877612 | 0.126689 | 0.488 |
| 2 | 0.000213 | 0.244351 | 0.352798 | 0.489 |
| 15 | 0.000139 | -0.0479209 | 0.0692096 | 0.489 |
| 30 | 0.000656 | -0.0293063 | 0.0424434 | 0.49 |
| 35 | 0.00052 | -0.0335357 | 0.0486023 | 0.49 |
| 5 | 0.000316 | -0.0937488 | 0.135872 | 0.49 |
| 11 | 0.00564 | 0.0166874 | 0.02418 | 0.49 |
| 5 | 9.85E-05 | -0.0947892 | 0.137701 | 0.491 |
| 2 | 0.000122 | -0.10454 | 0.151772 | 0.491 |
| 3 | 0.000101 | 0.113184 | 0.164836 | 0.492 |
| 4 | 0.00013 | -0.118082 | 0.171741 | 0.492 |
| 2 | 0.00064 | -0.0465226 | 0.0676547 | 0.492 |
| 5 | 0.000179 | -0.0923073 | 0.134707 | 0.493 |
| 152 | 0.0113 | -0.00480249 | 0.00700267 | 0.493 |

|  |  |  |  |  |
| --- | --- | --- | --- | --- |
| 2 | 0.000211 | -0.106898 | 0.15584 | 0.493 |
| 63 | 0.000648 | -0.021088 | 0.0307347 | 0.493 |
| 21 | 0.000225 | -0.0456966 | 0.0666287 | 0.493 |
| 42 | 0.000502 | -0.0276237 | 0.0403056 | 0.493 |
| 96 | 0.00132 | -0.0151803 | 0.0221721 | 0.494 |
| 88 | 0.00119 | -0.0155808 | 0.0228734 | 0.496 |
| 89 | 0.0013 | -0.0146097 | 0.0214996 | 0.497 |
| 4 | 5.95E-05 | -0.145298 | 0.213761 | 0.497 |
| 2 | 0.000277 | 0.0713713 | 0.104966 | 0.497 |
| 104 | 0.00396 | -0.00916365 | 0.0135112 | 0.498 |
| 108 | 0.00248 | 0.0103248 | 0.0152379 | 0.498 |
| 23 | 0.00033 | 0.0414305 | 0.061118 | 0.498 |
| 80 | 0.00253 | 0.0120987 | 0.0178654 | 0.498 |
| 5 | 0.000159 | -0.0893934 | 0.132393 | 0.5 |
| 5 | 0.000159 | -0.0896237 | 0.132958 | 0.5 |
| 5 | 4.28E-05 | -0.0988711 | 0.146449 | 0.5 |
| 3 | 0.000286 | 0.102559 | 0.152318 | 0.501 |
| 39 | 0.000708 | 0.0252553 | 0.0374969 | 0.501 |
| 8 | 0.00231 | 0.0565472 | 0.0841162 | 0.501 |
| 4 | 6.99E-05 | 0.125853 | 0.18761 | 0.502 |
| 3 | 0.000165 | 0.0741727 | 0.110842 | 0.503 |
| 9 | 0.000264 | 0.0678899 | 0.101344 | 0.503 |
| 5 | 0.000111 | 0.12796 | 0.191222 | 0.503 |
| 7 | 0.00127 | -0.0826325 | 0.12375 | 0.504 |
| 23 | 0.000467 | -0.0354093 | 0.0530073 | 0.504 |
| 53 | 0.000768 | 0.0227469 | 0.0340703 | 0.504 |
| 62 | 0.00126 | -0.0178672 | 0.0267582 | 0.504 |
| 134 | 0.00377 | 0.00784902 | 0.0117863 | 0.505 |
| 54 | 0.000991 | -0.0181551 | 0.0272634 | 0.505 |
| 22 | 0.000836 | 0.0493843 | 0.0740885 | 0.505 |
| 5 | 0.000153 | -0.0880969 | 0.132623 | 0.507 |
| 11 | 0.00086 | -0.0762199 | 0.114957 | 0.507 |
| 6 | 0.000737 | 0.0349782 | 0.0527104 | 0.507 |
| 27 | 0.000201 | -0.0515127 | 0.0777693 | 0.508 |
| 33 | 0.00045 | 0.0322695 | 0.0487531 | 0.508 |
| 3 | 0.000854 | -0.0512245 | 0.0778018 | 0.51 |
| 3 | 0.000104 | -0.0946309 | 0.14351 | 0.51 |
| 74 | 0.00362 | -0.010925 | 0.0165787 | 0.51 |
| 7 | 0.0011 | 0.0865404 | 0.131274 | 0.51 |
| 14 | 0.000124 | -0.0584943 | 0.0889646 | 0.511 |
| 29 | 0.00236 | -0.0245956 | 0.037433 | 0.511 |
| 26 | 0.000677 | -0.0241101 | 0.0368214 | 0.513 |
| 12 | 0.00012 | -0.0599114 | 0.0918521 | 0.514 |
| 3 | 0.000726 | 0.0881152 | 0.135068 | 0.514 |
| 12 | 0.00012 | -0.0596656 | 0.0918273 | 0.516 |
| 3 | 6.65E-05 | -0.209073 | 0.322541 | 0.517 |
| 25 | 0.00023 | -0.041404 | 0.0640627 | 0.518 |
| 64 | 0.00183 | 0.0118756 | 0.0183623 | 0.518 |
| 4 | 0.00113 | 0.0329399 | 0.0511168 | 0.519 |
| 114 | 0.00445 | -0.00753371 | 0.01169 | 0.519 |
| 2 | 2.94E-05 | -0.231337 | 0.359171 | 0.52 |
| 7 | 6.09E-05 | 0.110768 | 0.172315 | 0.52 |

|  |  |  |  |  |
| --- | --- | --- | --- | --- |
| 3 | 0.000864 | -0.0360217 | 0.0559336 | 0.52 |
| 2 | 0.000246 | 0.105916 | 0.165611 | 0.522 |
| 63 | 0.00121 | -0.021139 | 0.0329817 | 0.522 |
| 15 | 0.000377 | 0.070939 | 0.110775 | 0.522 |
| 55 | 0.000681 | 0.017969 | 0.028231 | 0.524 |
| 27 | 0.000455 | -0.0316985 | 0.0498591 | 0.525 |
| 3 | 0.000118 | 0.121736 | 0.191576 | 0.525 |
| 2 | 0.000246 | 0.105756 | 0.166645 | 0.526 |
| 11 | 0.000221 | -0.0648866 | 0.102239 | 0.526 |
| 4 | 0.000138 | -0.0741283 | 0.116844 | 0.526 |
| 3 | 0.000478 | 0.0572519 | 0.0903753 | 0.526 |
| 3 | 0.000478 | 0.057269 | 0.0903786 | 0.526 |
| 16 | 0.000493 | 0.0389005 | 0.0613647 | 0.526 |
| 114 | 0.00233 | -0.00940475 | 0.0148812 | 0.527 |
| 3 | 0.000478 | 0.0571419 | 0.0903857 | 0.527 |
| 8 | 0.000228 | 0.0515719 | 0.0816316 | 0.528 |
| 10 | 0.000157 | -0.0641556 | 0.101656 | 0.528 |
| 74 | 0.00108 | -0.0164405 | 0.0260612 | 0.528 |
| 6 | 0.000126 | -0.06632 | 0.105585 | 0.53 |
| 109 | 0.00293 | 0.01054 | 0.0167967 | 0.53 |
| 2 | 0.000125 | 0.201675 | 0.321942 | 0.531 |
| 3 | 0.00105 | -0.0624296 | 0.0996388 | 0.531 |
| 2 | 0.000246 | 0.104405 | 0.166897 | 0.532 |
| 28 | 0.000454 | -0.0313487 | 0.0501709 | 0.532 |
| 3 | 0.000158 | 0.0680905 | 0.109468 | 0.534 |
| 14 | 0.000124 | -0.0554628 | 0.0892741 | 0.534 |
| 11 | 0.000115 | -0.0597044 | 0.0963389 | 0.535 |
| 2 | 3.03E-05 | -0.587371 | 0.945855 | 0.535 |
| 3 | 9.33E-05 | 0.127841 | 0.207038 | 0.537 |
| 8 | 0.000334 | 0.053072 | 0.0863175 | 0.539 |
| 2 | 3.37E-05 | 0.820833 | 1.33705 | 0.539 |
| 9 | 0.000123 | -0.0487009 | 0.079552 | 0.54 |
| 164 | 0.0306 | 0.00247658 | 0.0040437 | 0.54 |
| 160 | 0.0282 | 0.00257586 | 0.00421594 | 0.541 |
| 15 | 0.000526 | 0.0284495 | 0.0466022 | 0.542 |
| 5 | 9.77E-05 | -0.0747775 | 0.122678 | 0.542 |
| 141 | 0.00851 | 0.00493474 | 0.00808761 | 0.542 |
| 2 | 0.000246 | 0.0606349 | 0.0996491 | 0.543 |
| 5 | 0.00112 | -0.090548 | 0.149091 | 0.544 |
| 5 | 6.00E-05 | -0.128176 | 0.21136 | 0.544 |
| 18 | 0.000585 | 0.049051 | 0.0807833 | 0.544 |
| 2 | 0.0122 | -0.0089741 | 0.0148607 | 0.546 |
| 50 | 0.000552 | 0.0235979 | 0.0390393 | 0.546 |
| 3 | 6.35E-05 | 0.139708 | 0.233175 | 0.549 |
| 18 | 0.00111 | -0.0299562 | 0.0499452 | 0.549 |
| 98 | 0.00248 | -0.0107323 | 0.0179229 | 0.549 |
| 7 | 4.43E-05 | 0.0993772 | 0.16673 | 0.551 |
| 4 | 0.00132 | -0.0270926 | 0.0457115 | 0.553 |
| 95 | 0.00133 | -0.0125638 | 0.0211554 | 0.553 |
| 147 | 0.0945 | 0.00156077 | 0.00262837 | 0.553 |
| 7 | 0.00018 | -0.0588337 | 0.0992477 | 0.553 |
| 43 | 0.000487 | 0.0242962 | 0.0413115 | 0.556 |

|  |  |  |  |  |
| --- | --- | --- | --- | --- |
| 34 | 0.000623 | -0.0235292 | 0.04006 | 0.557 |
| 10 | 9.85E-05 | -0.071065 | 0.120917 | 0.557 |
| 9 | 0.000387 | 0.0515219 | 0.0878441 | 0.558 |
| 4 | 0.00166 | -0.019501 | 0.0332506 | 0.558 |
| 84 | 0.000971 | -0.0141263 | 0.0241293 | 0.558 |
| 104 | 0.002 | 0.0107476 | 0.018343 | 0.558 |
| 37 | 0.000565 | -0.0247382 | 0.0423393 | 0.559 |
| 14 | 0.000185 | -0.0522763 | 0.0899141 | 0.561 |
| 19 | 0.000337 | 0.0327562 | 0.0563257 | 0.561 |
| 115 | 0.00339 | 0.00721767 | 0.0124111 | 0.561 |
| 5 | 0.00017 | -0.0904683 | 0.156509 | 0.563 |
| 7 | 0.000296 | -0.0579556 | 0.100437 | 0.564 |
| 3 | 0.000149 | -0.0809735 | 0.140851 | 0.565 |
| 141 | 0.0055 | -0.00558743 | 0.00970748 | 0.565 |
| 3 | 0.000236 | 0.0907063 | 0.157982 | 0.566 |
| 2 | 0.000119 | 0.0715437 | 0.125027 | 0.567 |
| 8 | 0.000381 | 0.0503038 | 0.0881403 | 0.568 |
| 40 | 0.000374 | -0.0283424 | 0.0496351 | 0.568 |
| 28 | 0.000456 | -0.0283825 | 0.0498497 | 0.569 |
| 114 | 0.00229 | -0.00856132 | 0.0150319 | 0.569 |
| 10 | 0.000179 | 0.0449717 | 0.079031 | 0.569 |
| 94 | 0.00984 | -0.00542418 | 0.00953854 | 0.57 |
| 15 | 0.000234 | 0.0480749 | 0.0845934 | 0.57 |
| 7 | 0.00024 | 0.0642891 | 0.113185 | 0.57 |
| 3 | 0.000101 | 0.124865 | 0.220315 | 0.571 |
| 5 | 0.000145 | 0.089311 | 0.157519 | 0.571 |
| 2 | 9.86E-05 | -0.105266 | 0.186254 | 0.572 |
| 4 | 0.00142 | -0.0295025 | 0.0522287 | 0.572 |
| 5 | 0.00201 | -0.0173056 | 0.0307408 | 0.573 |
| 23 | 0.000659 | -0.0251916 | 0.0449089 | 0.575 |
| 3 | 3.77E-05 | 0.128585 | 0.230752 | 0.577 |
| 20 | 0.000422 | -0.0382461 | 0.068726 | 0.578 |
| 146 | 0.00875 | -0.0047284 | 0.00851325 | 0.579 |
| 11 | 0.000175 | -0.0471515 | 0.0849021 | 0.579 |
| 4 | 0.000129 | 0.0896772 | 0.161508 | 0.579 |
| 8 | 0.000113 | -0.0677202 | 0.12224 | 0.58 |
| 8 | 5.05E-05 | 0.105658 | 0.191035 | 0.58 |
| 3 | 7.29E-05 | 0.0867371 | 0.157168 | 0.581 |
| 84 | 0.00117 | -0.0132652 | 0.0240061 | 0.581 |
| 10 | 0.000149 | -0.054686 | 0.0993341 | 0.582 |
| 56 | 0.000911 | -0.0174067 | 0.0316615 | 0.582 |
| 2 | 0.000111 | -0.0805598 | 0.146323 | 0.582 |
| 70 | 0.00305 | 0.00894998 | 0.016295 | 0.583 |
| 21 | 0.000359 | -0.0253016 | 0.046238 | 0.584 |
| 2 | 0.000187 | 0.189903 | 0.347817 | 0.585 |
| 15 | 0.000287 | -0.0453687 | 0.0830486 | 0.585 |
| 3 | 8.70E-05 | -0.0900804 | 0.165086 | 0.585 |
| 43 | 0.00128 | -0.0157825 | 0.0290487 | 0.587 |
| 103 | 0.00244 | 0.00906697 | 0.0167122 | 0.587 |
| 32 | 0.000519 | -0.0258511 | 0.0477513 | 0.588 |
| 2 | 2.94E-05 | -0.13255 | 0.245702 | 0.59 |
| 72 | 0.00101 | 0.0127087 | 0.023567 | 0.59 |

|  |  |  |  |  |
| --- | --- | --- | --- | --- |
| 33 | 0.00148 | 0.0197107 | 0.0366317 | 0.591 |
| 90 | 0.00159 | -0.0100332 | 0.0186513 | 0.591 |
| 3 | 0.0042 | 0.0141232 | 0.0263619 | 0.592 |
| 22 | 0.00024 | 0.0406229 | 0.076258 | 0.594 |
| 95 | 0.00122 | -0.0115494 | 0.0216717 | 0.594 |
| 37 | 0.000379 | -0.0259027 | 0.0486561 | 0.594 |
| 115 | 0.00306 | 0.00842518 | 0.0157855 | 0.594 |
| 3 | 0.000463 | 0.048995 | 0.0921591 | 0.595 |
| 53 | 0.000913 | 0.0194736 | 0.0365917 | 0.595 |
| 5 | 0.00021 | 0.0725219 | 0.136952 | 0.596 |
| 6 | 0.0011 | -0.0705081 | 0.133362 | 0.597 |
| 19 | 0.00108 | -0.026334 | 0.0498079 | 0.597 |
| 11 | 0.000171 | -0.0447618 | 0.0848508 | 0.598 |
| 143 | 0.0052 | -0.00528354 | 0.010008 | 0.598 |
| 51 | 0.000512 | -0.0186172 | 0.0352917 | 0.598 |
| 5 | 0.000116 | 0.0625524 | 0.11866 | 0.598 |
| 2 | 7.01E-05 | 0.102916 | 0.195605 | 0.599 |
| 60 | 0.00233 | -0.00888507 | 0.0168758 | 0.599 |
| 45 | 0.000698 | -0.0194866 | 0.0370151 | 0.599 |
| 17 | 0.000653 | -0.0216197 | 0.0411468 | 0.599 |
| 36 | 0.000819 | 0.021246 | 0.0403577 | 0.599 |
| 4 | 0.000976 | 0.0393307 | 0.0747058 | 0.599 |
| 26 | 0.000393 | 0.0308843 | 0.0587229 | 0.599 |
| 83 | 0.00112 | -0.012515 | 0.0238719 | 0.6 |
| 12 | 0.000136 | 0.0920389 | 0.175538 | 0.6 |
| 19 | 0.000191 | -0.0471655 | 0.0900528 | 0.6 |
| 3 | 3.11E-05 | -0.0979518 | 0.187123 | 0.601 |
| 98 | 0.00162 | -0.00949554 | 0.0181815 | 0.601 |
| 3 | 3.20E-05 | -0.0978091 | 0.187566 | 0.602 |
| 90 | 0.00118 | -0.0111051 | 0.0212877 | 0.602 |
| 23 | 0.000375 | 0.0352966 | 0.0678419 | 0.603 |
| 2 | 0.000124 | -0.0738884 | 0.142521 | 0.604 |
| 24 | 0.000771 | 0.0260428 | 0.0502058 | 0.604 |
| 27 | 0.000458 | -0.0257806 | 0.0498409 | 0.605 |
| 85 | 0.00123 | -0.0125543 | 0.0242886 | 0.605 |
| 52 | 0.00101 | -0.0140665 | 0.0271899 | 0.605 |
| 88 | 0.00157 | -0.00969213 | 0.0187908 | 0.606 |
| 6 | 3.60E-05 | 0.0986448 | 0.191131 | 0.606 |
| 25 | 0.000599 | 0.025968 | 0.0504496 | 0.607 |
| 8 | 0.00084 | 0.0547613 | 0.106411 | 0.607 |
| 17 | 0.000138 | 0.0567648 | 0.110776 | 0.608 |
| 3 | 0.00391 | 0.0641625 | 0.125146 | 0.608 |
| 87 | 0.00151 | -0.00995785 | 0.0194211 | 0.608 |
| 2 | 7.16E-05 | -0.19554 | 0.381735 | 0.608 |
| 3 | 9.58E-05 | -0.0837321 | 0.16378 | 0.609 |
| 17 | 0.000136 | 0.0565977 | 0.110869 | 0.61 |
| 8 | 0.00161 | -0.0510139 | 0.0999931 | 0.61 |
| 3 | 0.000116 | 0.0740213 | 0.145565 | 0.611 |
| 9 | 0.000107 | 0.0667784 | 0.131404 | 0.611 |
| 2 | 0.000133 | -0.0865299 | 0.170383 | 0.612 |
| 14 | 0.000122 | -0.044685 | 0.088452 | 0.613 |
| 7 | 0.000102 | 0.0730123 | 0.144753 | 0.614 |

|  |  |  |  |  |
| --- | --- | --- | --- | --- |
| 3 | 0.000399 | -0.0547225 | 0.109178 | 0.616 |
| 7 | 0.000102 | 0.0612384 | 0.122581 | 0.617 |
| 2 | 0.000281 | -0.0565501 | 0.113247 | 0.618 |
| 100 | 0.00175 | -0.00885744 | 0.0177491 | 0.618 |
| 2 | 8.91E-05 | 0.0691654 | 0.138625 | 0.618 |
| 19 | 0.000172 | 0.0418199 | 0.0842166 | 0.619 |
| 20 | 0.000428 | -0.030113 | 0.0606279 | 0.619 |
| 8 | 0.000353 | -0.0306871 | 0.0617248 | 0.619 |
| 2 | 5.39E-05 | 0.101158 | 0.204078 | 0.62 |
| 4 | 0.000622 | 0.037172 | 0.0749912 | 0.62 |
| 13 | 9.26E-05 | 0.0608303 | 0.122686 | 0.62 |
| 2 | 6.14E-05 | 0.210608 | 0.42633 | 0.621 |
| 59 | 0.00201 | -0.0108023 | 0.0219173 | 0.622 |
| 8 | 0.000433 | -0.0462557 | 0.0938849 | 0.622 |
| 15 | 0.000138 | -0.0506728 | 0.10289 | 0.622 |
| 12 | 0.00014 | -0.048064 | 0.0976638 | 0.623 |
| 9 | 0.000124 | -0.0523674 | 0.106506 | 0.623 |
| 6 | 9.74E-05 | 0.0735543 | 0.15017 | 0.624 |
| 22 | 0.000478 | 0.0386329 | 0.0787506 | 0.624 |
| 3 | 0.000192 | 0.0610649 | 0.125626 | 0.627 |
| 26 | 0.000258 | -0.0316957 | 0.0653977 | 0.628 |
| 2 | 0.00346 | 0.01358 | 0.0280283 | 0.628 |
| 155 | 0.0246 | 0.00235873 | 0.00487044 | 0.628 |
| 53 | 0.00102 | -0.0161875 | 0.0335017 | 0.629 |
| 7 | 0.00153 | -0.0526204 | 0.109253 | 0.63 |
| 5 | 0.000108 | -0.0644569 | 0.134366 | 0.631 |
| 25 | 0.000595 | -0.029419 | 0.0615556 | 0.633 |
| 4 | 3.95E-05 | 0.104154 | 0.217904 | 0.633 |
| 2 | 0.00545 | 0.0661721 | 0.139282 | 0.635 |
| 10 | 0.000249 | -0.0385853 | 0.0815505 | 0.636 |
| 20 | 0.000314 | 0.0322721 | 0.0683408 | 0.637 |
| 26 | 0.000175 | 0.037014 | 0.0787348 | 0.638 |
| 10 | 0.000877 | -0.0261033 | 0.0554856 | 0.638 |
| 83 | 0.00103 | -0.0108361 | 0.023013 | 0.638 |
| 120 | 0.00281 | -0.00644256 | 0.0137373 | 0.639 |
| 34 | 0.000521 | 0.0211616 | 0.0451112 | 0.639 |
| 119 | 0.00381 | 0.00587599 | 0.0125288 | 0.639 |
| 9 | 0.000304 | 0.0398052 | 0.085124 | 0.64 |
| 88 | 0.00154 | -0.00901769 | 0.0192884 | 0.64 |
| 65 | 0.000977 | -0.0160268 | 0.03423 | 0.64 |
| 14 | 0.000318 | 0.0316498 | 0.0679413 | 0.641 |
| 10 | 0.00017 | 0.0820985 | 0.177335 | 0.643 |
| 27 | 0.00247 | 0.0116284 | 0.0252028 | 0.645 |
| 16 | 0.000134 | 0.0340402 | 0.073797 | 0.645 |
| 35 | 0.000423 | 0.0187951 | 0.0408652 | 0.646 |
| 2 | 2.85E-05 | -0.0979761 | 0.213721 | 0.647 |
| 16 | 0.000233 | 0.0332134 | 0.0731298 | 0.65 |
| 155 | 0.0247 | 0.002202 | 0.00485625 | 0.65 |
| 12 | 0.000826 | 0.0479078 | 0.105431 | 0.65 |
| 4 | 0.000494 | -0.0436231 | 0.0965303 | 0.651 |
| 28 | 0.00245 | 0.0114543 | 0.0253247 | 0.651 |
| 4 | 8.87E-05 | -0.0633067 | 0.140571 | 0.652 |

|  |  |  |  |  |
| --- | --- | --- | --- | --- |
| 7 | 0.00153 | -0.0493019 | 0.109266 | 0.652 |
| 12 | 0.000389 | -0.0317919 | 0.0705049 | 0.652 |
| 16 | 0.000171 | -0.0325056 | 0.0723755 | 0.653 |
| 2 | 0.000291 | 0.0925165 | 0.206084 | 0.653 |
| 79 | 0.000929 | -0.0114034 | 0.0254349 | 0.654 |
| 145 | 0.00756 | -0.00390767 | 0.0087104 | 0.654 |
| 100 | 0.00143 | -0.00863717 | 0.0192906 | 0.654 |
| 85 | 0.00113 | -0.0100049 | 0.0224046 | 0.655 |
| 26 | 0.00238 | 0.0145637 | 0.0326817 | 0.656 |
| 24 | 0.00034 | 0.0263928 | 0.0594384 | 0.657 |
| 3 | 0.00039 | 0.0392859 | 0.088727 | 0.658 |
| 8 | 0.00171 | -0.0216424 | 0.0490557 | 0.659 |
| 3 | 7.54E-05 | -0.076759 | 0.17382 | 0.659 |
| 3 | 4.94E-05 | -0.0873044 | 0.198517 | 0.66 |
| 10 | 0.00037 | 0.0445935 | 0.1013 | 0.66 |
| 34 | 0.000243 | 0.0271575 | 0.0618251 | 0.66 |
| 2 | 0.00052 | 0.0415225 | 0.094373 | 0.66 |
| 6 | 0.000383 | -0.0370141 | 0.0844524 | 0.661 |
| 9 | 0.000372 | 0.0379693 | 0.086852 | 0.662 |
| 2 | 4.56E-05 | 0.0847033 | 0.19356 | 0.662 |
| 2 | 7.29E-05 | 0.137728 | 0.314901 | 0.662 |
| 3 | 0.000312 | -0.0884278 | 0.202733 | 0.663 |
| 6 | 0.00267 | 0.031485 | 0.0723099 | 0.663 |
| 7 | 0.000823 | 0.0467527 | 0.107445 | 0.663 |
| 4 | 3.24E-05 | -0.0791596 | 0.182143 | 0.664 |
| 113 | 0.00194 | -0.0070128 | 0.0162193 | 0.665 |
| 13 | 0.000168 | 0.0466228 | 0.10798 | 0.666 |
| 25 | 0.000384 | -0.0233922 | 0.0541998 | 0.666 |
| 27 | 0.000884 | -0.0186495 | 0.0436548 | 0.669 |
| 2 | 0.000274 | 0.0441433 | 0.103138 | 0.669 |
| 38 | 0.00216 | 0.00876476 | 0.0205462 | 0.67 |
| 97 | 0.00123 | -0.00883509 | 0.0207062 | 0.67 |
| 115 | 0.002 | 0.00722475 | 0.0169553 | 0.67 |
| 4 | 0.00169 | -0.0141676 | 0.0334877 | 0.672 |
| 4 | 0.000573 | -0.0594738 | 0.140602 | 0.672 |
| 7 | 0.000421 | -0.0434138 | 0.103061 | 0.674 |
| 8 | 0.000462 | -0.0391211 | 0.0931239 | 0.674 |
| 9 | 0.000238 | -0.0459725 | 0.109494 | 0.675 |
| 45 | 0.00184 | -0.0112119 | 0.0270753 | 0.679 |
| 15 | 0.000285 | 0.0289601 | 0.0701535 | 0.68 |
| 7 | 0.000112 | 0.0461975 | 0.112078 | 0.68 |
| 5 | 0.000157 | -0.0760193 | 0.184316 | 0.68 |
| 36 | 0.000878 | -0.0163855 | 0.0397159 | 0.68 |
| 4 | 4.41E-05 | 0.0604179 | 0.146477 | 0.68 |
| 3 | 0.000145 | 0.0510067 | 0.123924 | 0.681 |
| 11 | 0.000436 | 0.0353069 | 0.0858029 | 0.681 |
| 29 | 0.000586 | 0.0180618 | 0.0438932 | 0.681 |
| 88 | 0.00118 | -0.00881714 | 0.0214977 | 0.682 |
| 89 | 0.0013 | -0.00947557 | 0.0232383 | 0.683 |
| 11 | 0.000413 | -0.0265653 | 0.0650263 | 0.683 |
| 3 | 0.000776 | -0.0237503 | 0.0581821 | 0.683 |
| 75 | 0.00556 | -0.00607236 | 0.0149085 | 0.684 |

|  |  |  |  |  |
| --- | --- | --- | --- | --- |
| 8 | 0.000214 | 0.0448255 | 0.110505 | 0.685 |
| 12 | 0.00293 | 0.0129505 | 0.0319848 | 0.686 |
| 12 | 0.000197 | -0.0316938 | 0.0783378 | 0.686 |
| 39 | 0.00216 | 0.00827496 | 0.020551 | 0.687 |
| 31 | 0.000251 | -0.0266578 | 0.0661308 | 0.687 |
| 3 | 3.47E-05 | 0.0862888 | 0.215027 | 0.688 |
| 8 | 0.000532 | 0.0716015 | 0.178294 | 0.688 |
| 112 | 0.00282 | 0.00551618 | 0.013716 | 0.688 |
| 24 | 0.000572 | -0.0245779 | 0.0614282 | 0.689 |
| 23 | 0.00038 | -0.0229864 | 0.0574695 | 0.689 |
| 9 | 0.000454 | -0.0362907 | 0.0911357 | 0.69 |
| 26 | 0.00036 | -0.0289445 | 0.0725363 | 0.69 |
| 8 | 0.00337 | 0.0277975 | 0.0700477 | 0.691 |
| 2 | 5.14E-05 | -0.111136 | 0.279628 | 0.691 |
| 34 | 0.00119 | -0.0136848 | 0.0345965 | 0.692 |
| 41 | 0.00054 | 0.0180261 | 0.0454383 | 0.692 |
| 5 | 5.01E-05 | -0.106345 | 0.269523 | 0.693 |
| 5 | 5.87E-05 | 0.0547684 | 0.139224 | 0.694 |
| 2 | 0.000443 | -0.0276139 | 0.0708683 | 0.697 |
| 30 | 0.00225 | 0.00900123 | 0.023127 | 0.697 |
| 2 | 9.69E-05 | 0.0835081 | 0.215559 | 0.698 |
| 2 | 3.72E-05 | 0.198973 | 0.512404 | 0.698 |
| 6 | 0.000207 | -0.0305414 | 0.078623 | 0.698 |
| 11 | 7.08E-05 | -0.0530317 | 0.137318 | 0.699 |
| 64 | 0.000407 | 0.0145071 | 0.0375343 | 0.699 |
| 20 | 0.00115 | -0.016032 | 0.0414353 | 0.699 |
| 71 | 0.00175 | 0.00959146 | 0.0248891 | 0.7 |
| 4 | 0.000126 | 0.0696795 | 0.181023 | 0.7 |
| 6 | 0.000644 | -0.0267543 | 0.0695723 | 0.701 |
| 8 | 8.10E-05 | 0.063869 | 0.167975 | 0.704 |
| 12 | 0.000341 | -0.0207682 | 0.0548625 | 0.705 |
| 2 | 0.000143 | -0.048869 | 0.129691 | 0.706 |
| 4 | 0.000112 | 0.0538295 | 0.142752 | 0.706 |
| 23 | 0.00135 | -0.0183036 | 0.0486901 | 0.707 |
| 5 | 0.00182 | 0.040106 | 0.106859 | 0.707 |
| 70 | 0.00117 | -0.0102091 | 0.0272939 | 0.708 |
| 19 | 0.000242 | 0.0260935 | 0.069972 | 0.709 |
| 3 | 8.26E-05 | 0.0591589 | 0.159195 | 0.71 |
| 111 | 0.00444 | 0.00465999 | 0.0125839 | 0.711 |
| 5 | 0.00143 | -0.018198 | 0.0493208 | 0.712 |
| 2 | 0.00422 | -0.0334731 | 0.0905781 | 0.712 |
| 114 | 0.00234 | -0.00547897 | 0.014856 | 0.712 |
| 4 | 0.000207 | -0.0350353 | 0.0952829 | 0.713 |
| 53 | 0.000553 | -0.0139437 | 0.0380838 | 0.714 |
| 4 | 0.000113 | 0.0519432 | 0.142039 | 0.715 |
| 4 | 0.000113 | 0.0518688 | 0.141993 | 0.715 |
| 48 | 0.000703 | 0.0132403 | 0.036368 | 0.716 |
| 3 | 2.68E-05 | -0.0713202 | 0.197102 | 0.717 |
| 7 | 0.00112 | -0.0495668 | 0.137491 | 0.718 |
| 7 | 0.000326 | -0.0298313 | 0.08301 | 0.719 |
| 2 | 2.77E-05 | -0.0766517 | 0.213368 | 0.719 |
| 3 | 5.68E-05 | -0.0676909 | 0.18798 | 0.719 |

|  |  |  |  |  |
| --- | --- | --- | --- | --- |
| 99 | 0.00123 | -0.00743634 | 0.0206618 | 0.719 |
| 20 | 0.00149 | -0.0187437 | 0.0522882 | 0.72 |
| 5 | 0.000464 | -0.0389922 | 0.109115 | 0.721 |
| 23 | 0.0019 | 0.0138476 | 0.0387789 | 0.721 |
| 52 | 0.000785 | 0.0134992 | 0.0378527 | 0.721 |
| 9 | 6.03E-05 | -0.0601738 | 0.169319 | 0.722 |
| 5 | 0.00141 | -0.0174064 | 0.0488343 | 0.722 |
| 15 | 0.000385 | 0.024429 | 0.0690383 | 0.723 |
| 2 | 8.05E-05 | 0.0925012 | 0.260574 | 0.723 |
| 2 | 6.36E-05 | 0.0684742 | 0.193475 | 0.723 |
| 136 | 0.00823 | 0.00293422 | 0.00827688 | 0.723 |
| 2 | 6.36E-05 | 0.0685084 | 0.193216 | 0.723 |
| 2 | 3.03E-05 | 0.184214 | 0.518882 | 0.723 |
| 4 | 0.00128 | -0.043146 | 0.12191 | 0.723 |
| 22 | 0.000158 | 0.0377788 | 0.106889 | 0.724 |
| 8 | 0.000695 | 0.0449563 | 0.127503 | 0.724 |
| 2 | 5.97E-05 | -0.0890359 | 0.251821 | 0.724 |
| 7 | 2.83E-05 | 0.0678508 | 0.192122 | 0.724 |
| 9 | 0.00395 | 0.0101488 | 0.0288832 | 0.725 |
| 2 | 0.000908 | 0.020908 | 0.059659 | 0.726 |
| 18 | 0.00049 | -0.0217691 | 0.0622018 | 0.726 |
| 111 | 0.00405 | -0.00417182 | 0.0119341 | 0.727 |
| 18 | 0.000175 | 0.0250795 | 0.0717043 | 0.727 |
| 16 | 0.000193 | -0.0252777 | 0.072568 | 0.728 |
| 2 | 6.70E-05 | 0.0690386 | 0.198663 | 0.728 |
| 18 | 0.000531 | 0.0206144 | 0.0592982 | 0.728 |
| 7 | 0.000613 | -0.0219134 | 0.0630826 | 0.728 |
| 3 | 0.000194 | -0.0684431 | 0.197472 | 0.729 |
| 10 | 0.000364 | -0.0252426 | 0.0732746 | 0.73 |
| 23 | 0.000732 | -0.0125988 | 0.0365219 | 0.73 |
| 61 | 0.000666 | -0.0107094 | 0.0310264 | 0.73 |
| 2 | 6.01E-05 | 0.0682321 | 0.198538 | 0.731 |
| 2 | 6.01E-05 | 0.0678697 | 0.198153 | 0.732 |
| 112 | 0.00269 | 0.00486952 | 0.0142391 | 0.732 |
| 22 | 0.000479 | 0.0177876 | 0.0518835 | 0.732 |
| 35 | 0.000351 | 0.0171743 | 0.0503639 | 0.733 |
| 2 | 6.36E-05 | 0.0653833 | 0.192708 | 0.734 |
| 4 | 0.000214 | 0.0406525 | 0.119408 | 0.734 |
| 53 | 0.00163 | 0.00677129 | 0.0200713 | 0.736 |
| 2 | 0.000909 | 0.0201254 | 0.0596151 | 0.736 |
| 4 | 0.000362 | 0.0330391 | 0.098146 | 0.736 |
| 47 | 0.000956 | 0.0110931 | 0.032991 | 0.737 |
| 3 | 3.02E-05 | -0.0906263 | 0.272012 | 0.739 |
| 29 | 0.00149 | 0.012895 | 0.03877 | 0.739 |
| 4 | 0.000549 | 0.0206971 | 0.0620089 | 0.739 |
| 19 | 0.000413 | -0.02005 | 0.0604134 | 0.74 |
| 95 | 0.00248 | -0.00580758 | 0.0174924 | 0.74 |
| 30 | 0.000276 | 0.0235271 | 0.0707704 | 0.74 |
| 39 | 0.000434 | -0.0151905 | 0.0460484 | 0.741 |
| 9 | 0.000173 | -0.0241729 | 0.0732746 | 0.741 |
| 38 | 0.00389 | -0.00517943 | 0.0157027 | 0.742 |
| 10 | 0.000302 | 0.0198841 | 0.0604198 | 0.742 |

|  |  |  |  |  |
| --- | --- | --- | --- | --- |
| 89 | 0.00215 | 0.00617564 | 0.0187866 | 0.742 |
| 11 | 0.000385 | -0.0281477 | 0.0854081 | 0.742 |
| 5 | 0.00141 | -0.0160803 | 0.0489865 | 0.743 |
| 33 | 0.000481 | -0.0151985 | 0.046968 | 0.746 |
| 46 | 0.000897 | 0.0106679 | 0.0330509 | 0.747 |
| 115 | 0.00361 | -0.0046291 | 0.0144483 | 0.749 |
| 3 | 9.53E-05 | 0.0630297 | 0.196591 | 0.749 |
| 2 | 6.36E-05 | 0.0607257 | 0.19059 | 0.75 |
| 2 | 6.36E-05 | 0.0606644 | 0.190588 | 0.75 |
| 13 | 0.000468 | 0.023861 | 0.0750734 | 0.751 |
| 8 | 0.000444 | -0.0297622 | 0.0936545 | 0.751 |
| 3 | 7.17E-05 | 0.0837952 | 0.263663 | 0.751 |
| 2 | 0.0011 | 0.0201833 | 0.0635495 | 0.751 |
| 17 | 0.000676 | 0.0142536 | 0.0450816 | 0.752 |
| 4 | 0.00011 | 0.0450678 | 0.142522 | 0.752 |
| 25 | 0.000279 | 0.025532 | 0.0812222 | 0.753 |
| 119 | 0.00524 | 0.00314304 | 0.00997585 | 0.753 |
| 22 | 0.00407 | 0.00653491 | 0.0208476 | 0.754 |
| 3 | 0.000207 | -0.0365748 | 0.117314 | 0.755 |
| 7 | 0.000201 | 0.0382458 | 0.122741 | 0.755 |
| 13 | 0.00101 | 0.0233739 | 0.0749405 | 0.755 |
| 20 | 0.00017 | 0.0220905 | 0.070837 | 0.755 |
| 19 | 0.000431 | -0.018894 | 0.0610418 | 0.757 |
| 50 | 0.000487 | -0.0114979 | 0.037388 | 0.758 |
| 2 | 7.67E-05 | 0.0645869 | 0.209462 | 0.758 |
| 95 | 0.00242 | -0.00464098 | 0.0150328 | 0.758 |
| 3 | 0.000896 | -0.0265126 | 0.086294 | 0.759 |
| 4 | 0.000451 | 0.0240284 | 0.078619 | 0.76 |
| 113 | 0.0027 | 0.00434546 | 0.0142542 | 0.76 |
| 10 | 0.000202 | -0.0295981 | 0.0970591 | 0.76 |
| 139 | 0.00365 | 0.00348615 | 0.0114317 | 0.76 |
| 8 | 0.000324 | 0.0263005 | 0.0864953 | 0.761 |
| 37 | 0.000309 | -0.0175594 | 0.0576859 | 0.761 |
| 2 | 0.000247 | 0.0619526 | 0.204501 | 0.762 |
| 10 | 0.000234 | -0.025321 | 0.0836137 | 0.762 |
| 5 | 2.00E-04 | -0.0429115 | 0.142714 | 0.764 |
| 100 | 0.00197 | 0.00582864 | 0.0194852 | 0.765 |
| 3 | 0.000193 | 0.0423938 | 0.141885 | 0.765 |
| 2 | 5.54E-05 | -0.0536412 | 0.180431 | 0.766 |
| 91 | 0.00118 | -0.00675775 | 0.0226647 | 0.766 |
| 2 | 0.000661 | -0.0225976 | 0.0758984 | 0.766 |
| 2 | 6.36E-05 | 0.0565423 | 0.190152 | 0.766 |
| 21 | 0.000367 | 0.0191679 | 0.064645 | 0.767 |
| 3 | 3.55E-05 | -0.0516552 | 0.174478 | 0.767 |
| 112 | 0.00321 | 0.00433304 | 0.014706 | 0.768 |
| 2 | 0.000227 | -0.0297672 | 0.101764 | 0.77 |
| 90 | 0.00362 | 0.00410217 | 0.0140908 | 0.771 |
| 3 | 5.45E-05 | -0.0710976 | 0.245398 | 0.772 |
| 4 | 0.000203 | 0.0307687 | 0.106192 | 0.772 |
| 58 | 0.00134 | 0.00789135 | 0.0272832 | 0.772 |
| 10 | 0.000242 | -0.0219099 | 0.0758455 | 0.773 |
| 5 | 0.00013 | 0.0345799 | 0.119761 | 0.773 |

|  |  |  |  |  |
| --- | --- | --- | --- | --- |
| 40 | 0.000471 | 0.0149651 | 0.0520131 | 0.774 |
| 4 | 0.00011 | 0.0411432 | 0.14345 | 0.774 |
| 35 | 0.000308 | -0.0230008 | 0.0805816 | 0.775 |
| 6 | 0.000119 | 0.0336801 | 0.117769 | 0.775 |
| 6 | 0.000431 | -0.0297904 | 0.104506 | 0.776 |
| 2 | 0.000371 | 0.0278495 | 0.097684 | 0.776 |
| 7 | 0.000199 | -0.0287571 | 0.101009 | 0.776 |
| 11 | 0.00032 | -0.0217081 | 0.0765012 | 0.777 |
| 3 | 5.20E-05 | -0.0790236 | 0.280802 | 0.778 |
| 5 | 0.00168 | 0.0423023 | 0.149995 | 0.778 |
| 7 | 5.71E-05 | -0.0424966 | 0.150638 | 0.778 |
| 11 | 0.000105 | 0.0330313 | 0.119935 | 0.783 |
| 65 | 0.00131 | 0.00710183 | 0.025809 | 0.783 |
| 6 | 8.10E-05 | -0.0370486 | 0.134395 | 0.783 |
| 57 | 0.00196 | -0.00612105 | 0.0225566 | 0.786 |
| 13 | 0.000154 | 0.0263408 | 0.0972341 | 0.786 |
| 4 | 0.00125 | -0.0145194 | 0.053432 | 0.786 |
| 6 | 8.10E-05 | 0.0327712 | 0.121274 | 0.787 |
| 12 | 0.000166 | 0.0218975 | 0.0809699 | 0.787 |
| 4 | 8.16E-05 | -0.0542043 | 0.200669 | 0.787 |
| 85 | 0.001 | -0.00657551 | 0.0244036 | 0.788 |
| 4 | 0.000252 | 0.0229414 | 0.0852507 | 0.788 |
| 2 | 0.0016 | -0.0154158 | 0.0572914 | 0.788 |
| 3 | 1.00E-04 | -0.0369558 | 0.137937 | 0.789 |
| 2 | 0.000159 | 0.125008 | 0.46648 | 0.789 |
| 2 | 0.000528 | 0.0191857 | 0.0721035 | 0.79 |
| 3 | 4.18E-05 | -0.0798022 | 0.301093 | 0.791 |
| 4 | 0.00191 | -0.0108573 | 0.0410016 | 0.791 |
| 7 | 0.000138 | -0.0300207 | 0.113206 | 0.791 |
| 3 | 5.28E-05 | -0.062762 | 0.237344 | 0.791 |
| 2 | 0.000543 | -0.0392454 | 0.148615 | 0.792 |
| 7 | 0.000609 | -0.0165804 | 0.0629341 | 0.792 |
| 31 | 0.000387 | 0.0127086 | 0.0482937 | 0.792 |
| 109 | 0.00216 | -0.0044175 | 0.0168012 | 0.793 |
| 5 | 0.00168 | 0.0394478 | 0.150056 | 0.793 |
| 24 | 0.000598 | -0.0161646 | 0.0618875 | 0.794 |
| 2 | 3.37E-05 | -0.0726197 | 0.278235 | 0.794 |
| 71 | 0.000943 | 0.0067944 | 0.0260911 | 0.795 |
| 51 | 0.000611 | -0.00813673 | 0.0313364 | 0.795 |
| 3 | 0.00021 | 0.0318175 | 0.12241 | 0.795 |
| 9 | 0.000374 | 0.0260014 | 0.100399 | 0.796 |
| 7 | 0.000246 | 0.0454142 | 0.175363 | 0.796 |
| 2 | 0.000132 | 0.0866969 | 0.337847 | 0.797 |
| 20 | 0.000499 | -0.0158575 | 0.0615028 | 0.797 |
| 7 | 0.000232 | 0.0293118 | 0.114495 | 0.798 |
| 5 | 0.000181 | -0.0311145 | 0.12242 | 0.799 |
| 21 | 0.00022 | 0.0200814 | 0.0788364 | 0.799 |
| 7 | 6.66E-05 | 0.0315769 | 0.123814 | 0.799 |
| 6 | 0.000624 | 0.0515255 | 0.203071 | 0.8 |
| 6 | 0.000649 | 0.0146285 | 0.057651 | 0.8 |
| 85 | 0.00097 | -0.00616706 | 0.0244398 | 0.801 |
| 3 | 9.80E-05 | 0.0371907 | 0.147443 | 0.801 |

|  |  |  |  |  |
| --- | --- | --- | --- | --- |
| 2 | 0.00037 | 0.0247267 | 0.0981496 | 0.801 |
| 24 | 0.000606 | -0.0156194 | 0.0623659 | 0.802 |
| 12 | 0.000362 | -0.0262675 | 0.105118 | 0.803 |
| 11 | 0.000136 | -0.034664 | 0.139107 | 0.803 |
| 2 | 5.02E-05 | -0.0447431 | 0.182082 | 0.806 |
| 3 | 6.04E-05 | -0.0687868 | 0.279683 | 0.806 |
| 2 | 5.02E-05 | -0.0445835 | 0.182172 | 0.807 |
| 2 | 0.00363 | 0.00661317 | 0.0270912 | 0.807 |
| 20 | 0.000176 | -0.0239253 | 0.0987264 | 0.809 |
| 6 | 0.000205 | 0.0256449 | 0.106921 | 0.81 |
| 3 | 6.04E-05 | -0.0670532 | 0.279609 | 0.81 |
| 16 | 0.000285 | -0.0183769 | 0.0762753 | 0.81 |
| 116 | 0.00258 | -0.00409702 | 0.0170494 | 0.81 |
| 14 | 0.00048 | 0.0119419 | 0.0499575 | 0.811 |
| 3 | 6.04E-05 | -0.0663744 | 0.279627 | 0.812 |
| 70 | 0.00107 | 0.00678995 | 0.0285503 | 0.812 |
| 10 | 0.000154 | 0.0310427 | 0.130298 | 0.812 |
| 2 | 0.00037 | 0.0236635 | 0.0993054 | 0.812 |
| 3 | 6.04E-05 | -0.0661527 | 0.279342 | 0.813 |
| 3 | 7.91E-05 | -0.0502944 | 0.213528 | 0.814 |
| 5 | 7.75E-05 | 0.0372094 | 0.159386 | 0.815 |
| 2 | 0.000338 | 0.0427817 | 0.183128 | 0.815 |
| 35 | 0.000588 | -0.00932873 | 0.0401849 | 0.816 |
| 161 | 0.0192 | -0.00120163 | 0.00517957 | 0.817 |
| 4 | 0.000115 | 0.0298146 | 0.129364 | 0.818 |
| 3 | 8.67E-05 | 0.0628393 | 0.272619 | 0.818 |
| 7 | 0.000473 | -0.0216424 | 0.0941312 | 0.818 |
| 2 | 0.00643 | -0.00556433 | 0.0242089 | 0.818 |
| 3 | 0.000247 | 0.0335745 | 0.14587 | 0.818 |
| 10 | 0.00182 | 0.00630834 | 0.0274638 | 0.818 |
| 22 | 0.00103 | -0.0122914 | 0.0537616 | 0.819 |
| 3 | 0.000561 | 0.0142199 | 0.062132 | 0.819 |
| 16 | 0.000375 | 0.0155386 | 0.067784 | 0.819 |
| 7 | 0.000472 | -0.0215426 | 0.0941245 | 0.819 |
| 70 | 0.00361 | 0.00375662 | 0.0165128 | 0.82 |
| 2 | 0.002 | 0.0163841 | 0.0718796 | 0.82 |
| 36 | 0.000329 | 0.012849 | 0.056385 | 0.82 |
| 66 | 0.000868 | -0.00714282 | 0.0315011 | 0.821 |
| 5 | 0.00123 | -0.0124278 | 0.0555377 | 0.823 |
| 3 | 6.12E-05 | -0.0627283 | 0.279769 | 0.823 |
| 5 | 0.00123 | 0.0097043 | 0.0435786 | 0.824 |
| 29 | 0.000524 | -0.0085262 | 0.0385187 | 0.825 |
| 2 | 3.46E-05 | -0.0588867 | 0.265618 | 0.825 |
| 2 | 8.25E-05 | 0.0411417 | 0.18595 | 0.825 |
| 3 | 3.88E-05 | 0.0531348 | 0.240063 | 0.825 |
| 2 | 0.000491 | 0.0185404 | 0.0842456 | 0.826 |
| 2 | 0.000271 | -0.0207717 | 0.0949838 | 0.827 |
| 38 | 0.0014 | 0.00797146 | 0.0369575 | 0.829 |
| 4 | 6.70E-05 | 0.0382665 | 0.176658 | 0.829 |
| 145 | 0.00895 | 0.00172147 | 0.00797022 | 0.829 |
| 159 | 0.0115 | 0.00145728 | 0.0068081 | 0.831 |
| 28 | 0.000411 | -0.0096615 | 0.0452856 | 0.831 |

|  |  |  |  |  |
| --- | --- | --- | --- | --- |
| 2 | 0.000286 | -0.0202164 | 0.0946676 | 0.831 |
| 99 | 0.00179 | -0.00428471 | 0.020144 | 0.832 |
| 5 | 0.000498 | 0.0360193 | 0.169578 | 0.832 |
| 14 | 0.000697 | 0.00745494 | 0.035387 | 0.833 |
| 114 | 0.00242 | -0.00300878 | 0.0143771 | 0.834 |
| 2 | 7.91E-05 | 0.0393091 | 0.187398 | 0.834 |
| 2 | 0.000522 | 0.0144926 | 0.0696249 | 0.835 |
| 8 | 0.000239 | 0.020014 | 0.0966516 | 0.836 |
| 2 | 4.33E-05 | -0.0737278 | 0.359427 | 0.837 |
| 4 | 0.0104 | -0.00302834 | 0.014853 | 0.838 |
| 21 | 0.000544 | -0.0126387 | 0.0619493 | 0.838 |
| 5 | 0.00172 | 0.00734125 | 0.0360183 | 0.838 |
| 2 | 7.91E-05 | 0.0381322 | 0.186691 | 0.838 |
| 5 | 0.000317 | 0.0244907 | 0.119526 | 0.838 |
| 115 | 0.00242 | -0.00293321 | 0.0144344 | 0.839 |
| 3 | 0.000114 | -0.0271207 | 0.133263 | 0.839 |
| 41 | 4.00E-04 | 0.00802107 | 0.0393868 | 0.839 |
| 4 | 0.000142 | -0.0471263 | 0.231815 | 0.839 |
| 2 | 0.00108 | 0.0313037 | 0.155057 | 0.84 |
| 7 | 0.00079 | 0.00925066 | 0.0457207 | 0.84 |
| 3 | 7.13E-05 | -0.0312757 | 0.154965 | 0.84 |
| 18 | 0.000222 | -0.0156251 | 0.0779575 | 0.841 |
| 3 | 0.00181 | 0.00904271 | 0.0450674 | 0.841 |
| 2 | 0.000189 | -0.0492711 | 0.245046 | 0.841 |
| 25 | 0.000576 | -0.0122415 | 0.0613054 | 0.842 |
| 13 | 0.00116 | 0.0106976 | 0.053608 | 0.842 |
| 2 | 3.46E-05 | -0.100398 | 0.506694 | 0.843 |
| 5 | 0.000107 | -0.0464032 | 0.233885 | 0.843 |
| 53 | 0.00101 | 0.00629123 | 0.0320009 | 0.844 |
| 31 | 0.000282 | 0.0123454 | 0.062723 | 0.844 |
| 4 | 8.40E-05 | -0.0287496 | 0.145742 | 0.844 |
| 2 | 0.000758 | -0.0140638 | 0.0719492 | 0.845 |
| 95 | 0.0026 | -0.00273777 | 0.0141387 | 0.846 |
| 115 | 0.00244 | -0.00278237 | 0.0144167 | 0.847 |
| 2 | 2.94E-05 | 0.280156 | 1.45077 | 0.847 |
| 13 | 0.000189 | 0.0195214 | 0.101542 | 0.848 |
| 31 | 0.000681 | 0.00746521 | 0.038913 | 0.848 |
| 129 | 0.00449 | -0.00211155 | 0.0110851 | 0.849 |
| 3 | 0.000768 | -0.0292275 | 0.153967 | 0.849 |
| 7 | 4.40E-05 | -0.0387249 | 0.204594 | 0.85 |
| 8 | 0.000172 | 0.0195261 | 0.103116 | 0.85 |
| 2 | 3.11E-05 | 0.0445882 | 0.235145 | 0.85 |
| 164 | 0.0264 | -0.000826121 | 0.00436911 | 0.85 |
| 2 | 7.73E-05 | 0.034996 | 0.186511 | 0.851 |
| 2 | 7.56E-05 | 0.0349798 | 0.186441 | 0.851 |
| 8 | 6.83E-05 | -0.0275849 | 0.147028 | 0.851 |
| 3 | 0.00014 | -0.0188571 | 0.100204 | 0.851 |
| 7 | 0.00103 | 0.0244046 | 0.130814 | 0.852 |
| 89 | 0.00127 | -0.00405581 | 0.0217647 | 0.852 |
| 114 | 0.00243 | -0.00266285 | 0.0143474 | 0.853 |
| 114 | 0.00241 | -0.00267916 | 0.0144568 | 0.853 |
| 11 | 0.000417 | -0.0103773 | 0.0560911 | 0.853 |

|  |  |  |  |  |
| --- | --- | --- | --- | --- |
| 22 | 0.000223 | 0.0189421 | 0.102013 | 0.853 |
| 2 | 0.000435 | -0.0174312 | 0.0941891 | 0.853 |
| 127 | 0.00538 | 0.00184756 | 0.0100071 | 0.854 |
| 4 | 6.36E-05 | 0.0321011 | 0.17488 | 0.854 |
| 4 | 6.53E-05 | 0.0319551 | 0.17486 | 0.855 |
| 36 | 0.000558 | 0.00812346 | 0.0445445 | 0.855 |
| 9 | 0.000147 | 0.0192474 | 0.105942 | 0.856 |
| 24 | 0.000342 | 0.0107569 | 0.0591335 | 0.856 |
| 74 | 0.00211 | 0.00342935 | 0.0190083 | 0.857 |
| 2 | 4.79E-05 | -0.04172 | 0.233588 | 0.858 |
| 7 | 0.000261 | -0.0173467 | 0.0971513 | 0.858 |
| 163 | 0.0389 | 0.00066787 | 0.00373685 | 0.858 |
| 7 | 0.00047 | -0.0171585 | 0.0962676 | 0.859 |
| 29 | 0.000234 | 0.0124549 | 0.0706755 | 0.86 |
| 29 | 0.000443 | 0.00875761 | 0.0496335 | 0.86 |
| 118 | 0.00282 | 0.00255464 | 0.0144758 | 0.86 |
| 3 | 0.00167 | 0.00816782 | 0.0464048 | 0.86 |
| 24 | 0.000286 | -0.0104045 | 0.0590827 | 0.86 |
| 8 | 0.000401 | -0.0168799 | 0.0967284 | 0.861 |
| 13 | 0.000153 | 0.0135856 | 0.077746 | 0.861 |
| 4 | 0.000473 | -0.0140563 | 0.0808507 | 0.862 |
| 20 | 0.000383 | -0.00829508 | 0.0476852 | 0.862 |
| 3 | 2.68E-05 | -0.0329322 | 0.190382 | 0.863 |
| 2 | 0.000119 | 0.0450042 | 0.261671 | 0.863 |
| 8 | 0.00117 | -0.0203257 | 0.118406 | 0.864 |
| 150 | 0.00612 | -0.00156853 | 0.00913004 | 0.864 |
| 115 | 0.00244 | -0.00243654 | 0.0144105 | 0.866 |
| 60 | 0.00104 | -0.00621346 | 0.0367062 | 0.866 |
| 13 | 0.000154 | 0.0131214 | 0.0777565 | 0.866 |
| 5 | 0.000411 | -0.0130005 | 0.0769696 | 0.866 |
| 38 | 0.000335 | 0.00942797 | 0.0562986 | 0.867 |
| 93 | 0.0043 | -0.0021504 | 0.0128024 | 0.867 |
| 2 | 0.000113 | 0.0519595 | 0.309502 | 0.867 |
| 4 | 9.39E-05 | 0.0193352 | 0.11757 | 0.869 |
| 4 | 0.000105 | 0.0211422 | 0.128673 | 0.869 |
| 100 | 0.00141 | -0.00323065 | 0.0197577 | 0.87 |
| 96 | 0.0046 | -0.00200422 | 0.0123455 | 0.871 |
| 11 | 0.000218 | -0.0194041 | 0.11911 | 0.871 |
| 12 | 0.000108 | 0.0149642 | 0.0926422 | 0.872 |
| 40 | 0.000491 | 0.00850789 | 0.0533614 | 0.873 |
| 3 | 7.09E-05 | 0.0526817 | 0.328495 | 0.873 |
| 4 | 0.000125 | 0.023621 | 0.148555 | 0.874 |
| 38 | 0.00034 | 0.00857644 | 0.0555343 | 0.877 |
| 152 | 0.0175 | 0.000906771 | 0.00583673 | 0.877 |
| 13 | 0.000189 | 0.0156772 | 0.101607 | 0.877 |
| 56 | 0.000462 | 0.0057777 | 0.0375791 | 0.878 |
| 2 | 0.000577 | 0.0119047 | 0.0778904 | 0.879 |
| 2 | 0.000577 | 0.0118728 | 0.0779205 | 0.879 |
| 8 | 0.00137 | 0.0190335 | 0.124631 | 0.879 |
| 3 | 7.33E-05 | -0.0258431 | 0.169526 | 0.879 |
| 3 | 0.000381 | 0.014644 | 0.0968309 | 0.88 |
| 5 | 0.00042 | -0.0142708 | 0.0956233 | 0.881 |

|  |  |  |  |  |
| --- | --- | --- | --- | --- |
| 14 | 0.000191 | 0.0150702 | 0.100412 | 0.881 |
| 53 | 0.00105 | -0.00489753 | 0.0327291 | 0.881 |
| 156 | 0.0318 | 0.000611811 | 0.00408477 | 0.881 |
| 113 | 0.00273 | 0.00212118 | 0.0141663 | 0.881 |
| 7 | 8.05E-05 | -0.0229186 | 0.152472 | 0.881 |
| 22 | 0.000171 | 0.0101457 | 0.0677555 | 0.881 |
| 2 | 0.000106 | -0.0474187 | 0.319327 | 0.882 |
| 22 | 0.000562 | -0.00916362 | 0.0618893 | 0.882 |
| 58 | 0.0022 | -0.00246615 | 0.0166658 | 0.882 |
| 4 | 0.00259 | -0.00622871 | 0.0418777 | 0.882 |
| 26 | 0.000945 | 0.00486815 | 0.033082 | 0.883 |
| 4 | 0.000795 | -0.0119869 | 0.0815825 | 0.883 |
| 5 | 7.29E-05 | 0.0212164 | 0.144131 | 0.883 |
| 48 | 0.000899 | 0.00422507 | 0.0289838 | 0.884 |
| 19 | 0.000214 | 0.0131925 | 0.09104 | 0.885 |
| 11 | 0.000143 | 0.0160257 | 0.111878 | 0.886 |
| 29 | 0.000409 | -0.00802717 | 0.0557454 | 0.886 |
| 5 | 3.90E-05 | 0.024296 | 0.169579 | 0.886 |
| 21 | 0.00027 | -0.0103065 | 0.0723623 | 0.887 |
| 8 | 0.00115 | -0.0168552 | 0.118485 | 0.887 |
| 5 | 6.96E-05 | 0.0205785 | 0.144187 | 0.887 |
| 55 | 0.0344 | -0.000963057 | 0.00684453 | 0.888 |
| 90 | 0.00121 | -0.00319859 | 0.0226509 | 0.888 |
| 2 | 0.000957 | 0.010809 | 0.0769948 | 0.888 |
| 82 | 0.0012 | -0.00318319 | 0.0225635 | 0.888 |
| 25 | 0.000168 | 0.0113395 | 0.0811514 | 0.889 |
| 160 | 0.0164 | -0.000776228 | 0.00558655 | 0.889 |
| 3 | 6.83E-05 | -0.0244871 | 0.177075 | 0.89 |
| 14 | 0.00019 | -0.00924076 | 0.0669148 | 0.89 |
| 2 | 0.000164 | -0.0217961 | 0.15866 | 0.891 |
| 2 | 7.47E-05 | -0.0394661 | 0.290949 | 0.892 |
| 51 | 0.000584 | -0.00417441 | 0.0307886 | 0.892 |
| 3 | 7.63E-05 | -0.0231304 | 0.171366 | 0.893 |
| 141 | 0.00565 | -0.00127513 | 0.0096111 | 0.894 |
| 2 | 5.97E-05 | 0.0522132 | 0.39217 | 0.894 |
| 18 | 0.000681 | 0.01157 | 0.0864692 | 0.894 |
| 21 | 0.000461 | 0.0105735 | 0.080002 | 0.895 |
| 3 | 0.000112 | -0.0175916 | 0.133799 | 0.895 |
| 8 | 7.94E-05 | 0.0168137 | 0.127389 | 0.895 |
| 2 | 0.00127 | -0.0103718 | 0.0798263 | 0.897 |
| 11 | 0.000127 | -0.0127886 | 0.10025 | 0.898 |
| 51 | 5.00E-04 | -0.00455476 | 0.0356823 | 0.898 |
| 3 | 0.000341 | 0.0119228 | 0.0940496 | 0.899 |
| 113 | 0.00198 | 0.0021317 | 0.017057 | 0.901 |
| 2 | 2.68E-05 | -0.0235817 | 0.188988 | 0.901 |
| 85 | 0.00223 | 0.00209139 | 0.0169656 | 0.902 |
| 4 | 0.000563 | -0.00970825 | 0.0790309 | 0.902 |
| 95 | 0.00188 | -0.00205825 | 0.016939 | 0.903 |
| 2 | 0.000166 | -0.0189455 | 0.157297 | 0.904 |
| 52 | 0.000557 | 0.00382514 | 0.0319776 | 0.905 |
| 3 | 4.44E-05 | -0.018748 | 0.157118 | 0.905 |
| 15 | 0.00148 | 0.00926303 | 0.0784991 | 0.906 |

|  |  |  |  |  |
| --- | --- | --- | --- | --- |
| 2 | 0.000364 | 0.011582 | 0.0992834 | 0.907 |
| 17 | 0.000203 | -0.00897798 | 0.0769716 | 0.907 |
| 2 | 6.57E-05 | 0.0288553 | 0.25017 | 0.908 |
| 7 | 0.000274 | -0.0110441 | 0.0974616 | 0.91 |
| 3 | 0.000141 | 0.0135879 | 0.120703 | 0.91 |
| 19 | 0.000241 | -0.00730444 | 0.0646329 | 0.91 |
| 2 | 0.000139 | -0.0276237 | 0.247104 | 0.911 |
| 5 | 0.000256 | 0.0118299 | 0.106444 | 0.912 |
| 2 | 0.000448 | -0.010388 | 0.0938773 | 0.912 |
| 25 | 0.000687 | 0.00500495 | 0.0456158 | 0.913 |
| 49 | 0.000546 | -0.00350578 | 0.0325334 | 0.914 |
| 3 | 0.000327 | -0.0112251 | 0.105457 | 0.915 |
| 18 | 0.000616 | -0.00805705 | 0.0763979 | 0.916 |
| 39 | 0.000318 | -0.00605504 | 0.0580134 | 0.917 |
| 11 | 0.000235 | 0.00878673 | 0.0840556 | 0.917 |
| 6 | 0.000585 | 0.0086614 | 0.0829177 | 0.917 |
| 16 | 0.000441 | -0.00781976 | 0.0760917 | 0.918 |
| 2 | 0.000213 | 0.0133129 | 0.129016 | 0.918 |
| 8 | 0.000651 | 0.00939892 | 0.0924112 | 0.919 |
| 3 | 7.24E-05 | -0.0214091 | 0.211827 | 0.919 |
| 7 | 0.00118 | -0.00441426 | 0.0434026 | 0.919 |
| 5 | 0.000103 | 0.0143484 | 0.141447 | 0.919 |
| 24 | 0.000315 | 0.00595663 | 0.0595049 | 0.92 |
| 38 | 0.000444 | 0.00407864 | 0.0408832 | 0.921 |
| 110 | 0.00218 | 0.00166066 | 0.0167552 | 0.921 |
| 53 | 0.000594 | -0.00300002 | 0.0303417 | 0.921 |
| 58 | 0.000604 | -0.00394308 | 0.0404204 | 0.922 |
| 30 | 0.000625 | -0.00449497 | 0.0457821 | 0.922 |
| 11 | 0.000167 | -0.00893898 | 0.0913776 | 0.922 |
| 61 | 0.0011 | -0.00267853 | 0.0274522 | 0.922 |
| 45 | 0.000922 | -0.00354263 | 0.0373612 | 0.924 |
| 51 | 0.0012 | 0.00225592 | 0.0235255 | 0.924 |
| 31 | 0.000402 | -0.00425819 | 0.0451189 | 0.925 |
| 27 | 0.0017 | 0.00274989 | 0.029808 | 0.926 |
| 49 | 0.000731 | 0.00293881 | 0.0326158 | 0.928 |
| 6 | 0.000767 | 0.0143725 | 0.162175 | 0.929 |
| 24 | 0.000289 | -0.00526171 | 0.0589276 | 0.929 |
| 5 | 8.11E-05 | 0.0127747 | 0.142842 | 0.929 |
| 3 | 5.04E-05 | 0.0199867 | 0.228389 | 0.93 |
| 29 | 0.000444 | 0.00504475 | 0.0578057 | 0.93 |
| 14 | 0.000226 | 0.00876834 | 0.10088 | 0.931 |
| 12 | 0.000641 | 0.00457195 | 0.05359 | 0.932 |
| 3 | 7.57E-05 | -0.019849 | 0.232518 | 0.932 |
| 2 | 8.89E-05 | 0.0154684 | 0.181853 | 0.932 |
| 5 | 0.000329 | 0.0101974 | 0.121589 | 0.933 |
| 55 | 0.00644 | -0.00104574 | 0.0126515 | 0.934 |
| 6 | 7.75E-05 | -0.0113505 | 0.139041 | 0.935 |
| 18 | 0.000734 | -0.00498272 | 0.0621077 | 0.936 |
| 43 | 0.00143 | 0.00285799 | 0.0365893 | 0.938 |
| 3 | 3.02E-05 | -0.015248 | 0.196056 | 0.938 |
| 2 | 4.72E-05 | -0.0158835 | 0.209326 | 0.94 |
| 12 | 0.000378 | 0.00756357 | 0.102477 | 0.941 |

|  |  |  |  |  |
| --- | --- | --- | --- | --- |
| 2 | 0.000137 | -0.0200719 | 0.281122 | 0.943 |
| 2 | 0.000153 | -0.0116364 | 0.163092 | 0.943 |
| 2 | 0.00042 | -0.00857709 | 0.119932 | 0.943 |
| 62 | 0.00875 | 0.000882942 | 0.0125429 | 0.944 |
| 6 | 0.00112 | -0.00295747 | 0.0422997 | 0.944 |
| 51 | 0.00459 | -0.00121178 | 0.0175488 | 0.945 |
| 96 | 0.0012 | -0.00146447 | 0.0214373 | 0.946 |
| 4 | 0.000379 | 0.00604242 | 0.0916006 | 0.947 |
| 23 | 0.000256 | -0.0039637 | 0.0591174 | 0.947 |
| 37 | 0.00133 | 0.00248082 | 0.0371786 | 0.947 |
| 3 | 0.00387 | -0.0017044 | 0.0256433 | 0.947 |
| 11 | 0.000143 | 0.00725755 | 0.111863 | 0.948 |
| 20 | 0.000408 | -0.00371082 | 0.056816 | 0.948 |
| 8 | 0.00125 | -0.0067903 | 0.103385 | 0.948 |
| 6 | 0.000124 | 0.00811198 | 0.125644 | 0.949 |
| 94 | 0.00208 | -0.00142828 | 0.0224805 | 0.949 |
| 22 | 0.000306 | -0.00411331 | 0.0655303 | 0.95 |
| 26 | 0.000279 | -0.00528245 | 0.0862679 | 0.951 |
| 3 | 0.000138 | 0.0102707 | 0.168866 | 0.952 |
| 6 | 0.000876 | 0.00410981 | 0.0703441 | 0.953 |
| 28 | 0.000569 | -0.00280104 | 0.0475885 | 0.953 |
| 8 | 0.00142 | -0.00271895 | 0.046194 | 0.953 |
| 33 | 0.00124 | -0.00192461 | 0.0328163 | 0.953 |
| 3 | 5.47E-05 | 0.0109173 | 0.198339 | 0.956 |
| 2 | 0.000148 | -0.0149842 | 0.282454 | 0.958 |
| 8 | 0.000568 | -0.00497211 | 0.0943893 | 0.958 |
| 36 | 0.000511 | 0.00249052 | 0.0470298 | 0.958 |
| 46 | 0.000961 | -0.00161634 | 0.0315785 | 0.959 |
| 85 | 0.00122 | -0.0012469 | 0.024312 | 0.959 |
| 3 | 0.000465 | 0.00468126 | 0.094073 | 0.96 |
| 24 | 0.000298 | -0.0029539 | 0.0593091 | 0.96 |
| 8 | 0.00122 | -0.00567056 | 0.117031 | 0.961 |
| 62 | 0.000813 | 0.00151125 | 0.0318315 | 0.962 |
| 6 | 7.68E-05 | -0.00687211 | 0.146468 | 0.963 |
| 20 | 0.000387 | 0.00449001 | 0.0996374 | 0.964 |
| 2 | 0.000173 | -0.0053054 | 0.116699 | 0.964 |
| 2 | 6.64E-05 | -0.0158659 | 0.353169 | 0.964 |
| 69 | 0.00121 | -0.00138849 | 0.0305613 | 0.964 |
| 6 | 6.46E-05 | -0.00562036 | 0.125532 | 0.964 |
| 5 | 8.85E-05 | 0.00636183 | 0.141972 | 0.964 |
| 2 | 0.015 | 0.000577913 | 0.0130109 | 0.965 |
| 97 | 0.00258 | -0.000613965 | 0.0141876 | 0.965 |
| 27 | 0.000289 | 0.00230615 | 0.052737 | 0.965 |
| 4 | 0.000133 | -0.00570181 | 0.139267 | 0.967 |
| 7 | 0.00115 | 0.0062642 | 0.149903 | 0.967 |
| 55 | 0.0017 | 0.000801065 | 0.0198664 | 0.968 |
| 9 | 0.00161 | -0.00340853 | 0.0859458 | 0.968 |
| 3 | 0.00013 | 0.0121191 | 0.301978 | 0.968 |
| 114 | 0.00502 | 0.00040941 | 0.0106652 | 0.969 |
| 2 | 5.62E-05 | -0.0130562 | 0.338233 | 0.969 |
| 16 | 0.000205 | 0.00225909 | 0.0607655 | 0.97 |
| 3 | 8.56E-05 | -0.00557404 | 0.165364 | 0.973 |

|  |  |  |  |  |
| --- | --- | --- | --- | --- |
| 16 | 0.000409 | 0.00167343 | 0.0517513 | 0.974 |
| 5 | 0.000167 | -0.00409554 | 0.125706 | 0.974 |
| 8 | 0.00125 | -0.00340754 | 0.103342 | 0.974 |
| 4 | 8.67E-05 | 0.00659569 | 0.212436 | 0.975 |
| 2 | 8.24E-05 | -0.00574479 | 0.182713 | 0.975 |
| 2 | 0.000168 | -0.0044224 | 0.147925 | 0.976 |
| 3 | 0.000201 | 0.00351911 | 0.115191 | 0.976 |
| 50 | 0.00411 | -0.00073632 | 0.0279979 | 0.979 |
| 9 | 0.000302 | -0.00202896 | 0.0849177 | 0.981 |
| 3 | 0.000144 | 0.00336579 | 0.140472 | 0.981 |
| 2 | 0.000648 | 0.0016875 | 0.071312 | 0.981 |
| 89 | 0.00116 | 0.000538044 | 0.025581 | 0.983 |
| 53 | 0.000594 | 0.000658918 | 0.0303675 | 0.983 |
| 2 | 8.05E-05 | -0.00704545 | 0.340868 | 0.984 |
| 8 | 0.000265 | -0.00177138 | 0.0883086 | 0.984 |
| 19 | 0.000941 | 0.000712036 | 0.0385415 | 0.985 |
| 6 | 4.95E-05 | 0.00349726 | 0.184911 | 0.985 |
| 65 | 0.000813 | 0.000504387 | 0.0264684 | 0.985 |
| 59 | 0.000572 | -0.000705769 | 0.0369339 | 0.985 |
| 169 | 0.191 | -3.32E-05 | 0.00181938 | 0.985 |
| 19 | 0.000142 | 0.00125464 | 0.074056 | 0.986 |
| 4 | 6.58E-05 | -0.00345307 | 0.212704 | 0.987 |
| 2 | 0.000108 | -0.0032827 | 0.196844 | 0.987 |
| 5 | 0.00247 | -0.000723555 | 0.0434134 | 0.987 |
| 6 | 6.54E-05 | -0.00200703 | 0.125708 | 0.987 |
| 6 | 6.54E-05 | -0.00211418 | 0.125714 | 0.987 |
| 2 | 4.33E-05 | 0.00463929 | 0.288622 | 0.987 |
| 8 | 0.000298 | 0.00132534 | 0.0858082 | 0.988 |
| 2 | 4.24E-05 | -0.00442072 | 0.288715 | 0.988 |
| 18 | 0.000745 | 0.000486754 | 0.035924 | 0.989 |
| 7 | 0.000107 | -0.00172389 | 0.122947 | 0.989 |
| 13 | 0.000101 | 0.00124186 | 0.0890232 | 0.989 |
| 3 | 0.00334 | 0.000260729 | 0.0259186 | 0.992 |
| 2 | 6.66E-05 | -0.00273229 | 0.280274 | 0.992 |
| 26 | 0.000398 | 0.000451821 | 0.0507322 | 0.993 |
| 3 | 0.000797 | -0.001294 | 0.174572 | 0.994 |
| 60 | 0.000763 | -0.000228573 | 0.0283617 | 0.994 |
| 2 | 6.66E-05 | 0.00184243 | 0.279369 | 0.995 |
| 78 | 0.00145 | -0.000151605 | 0.0223794 | 0.995 |
| 2 | 6.87E-05 | -0.00132956 | 0.28208 | 0.996 |
| 32 | 0.000344 | 0.000303913 | 0.059735 | 0.996 |
| 2 | 6.79E-05 | -0.00117762 | 0.2845 | 0.997 |
| 20 | 0.00023 | 0.000243674 | 0.0698102 | 0.997 |
| 146 | 0.00891 | -2.75E-05 | 0.00801946 | 0.997 |
| 2 | 6.66E-05 | 0.000668524 | 0.278543 | 0.998 |
| 2 | 6.46E-05 | -0.00055671 | 0.2773 | 0.998 |
| 4 | 0.00169 | -9.85E-05 | 0.123226 | 0.999 |
| 3 | 0.00147 | -6.79E-05 | 0.0419145 | 0.999 |
| 12 | 0.00018 | -0.000175776 | 0.126597 | 0.999 |

| rsID | CHROM | POS_b37 | REF | ALT | N | N_studies | POOLED_ALT_AF |
| --- | --- | --- | --- | --- | --- | --- | --- |
| rs62019326 | 15 | 42917387 | G | A | 1245174 | 171 | 0.0875 |
| rs532800789 | 15 | 42966538 | A | G | 841791 | 57 | 0.00167 |
| rs187892402 | 15 | 42994453 | G | A | 759354 | 75 | 0.00113 |
| rs12914539 | 15 | 42910624 | C | A | 1253061 | 176 | 0.128 |
| rs62019322 | 15 | 42897374 | T | C | 1253234 | 176 | 0.145 |
| rs62019320 | 15 | 42895389 | C | T | 1253234 | 176 | 0.144 |
| rs17767270 | 15 | 42898612 | C | T | 1253234 | 176 | 0.146 |
| rs17767264 | 15 | 42886656 | C | T | 1253237 | 176 | 0.144 |
| rs79044457 | 15 | 42895208 | G | T | 1253211 | 176 | 0.142 |
| rs538041733 | 15 | 42938029 | C | G | 802705 | 43 | 0.0015 |
| rs78366241 | 15 | 42895209 | C | T | 1253210 | 176 | 0.142 |
| rs11070375 | 15 | 42870587 | A | G | 1253144 | 176 | 0.144 |
| rs12910538 | 15 | 42910357 | G | C | 1253045 | 176 | 0.144 |
| rs202077402 | 15 | 42982090 | A | G | 1237449 | 148 | 0.00442 |
| rs550777038 | 15 | 42969019 | C | T | 1229932 | 147 | 0.00435 |
| rs56172155 | 15 | 42882182 | C | T | 1092509 | 127 | 0.145 |
| rs1995939 | 15 | 43012517 | A | G | 1253211 | 176 | 0.22 |
| rs3199486 | 15 | 43012195 | T | C | 1253212 | 176 | 0.22 |
| rs2280048 | 15 | 43011297 | C | A | 1253185 | 176 | 0.22 |
| rs2412740 | 15 | 43009448 | T | C | 1247691 | 175 | 0.22 |
| rs12912064 | 15 | 43004170 | C | T | 1247699 | 175 | 0.222 |
| rs12914570 | 15 | 42970899 | A | G | 1247576 | 175 | 0.194 |
| rs7169202 | 15 | 42973053 | G | C | 1247665 | 175 | 0.224 |
| rs12102055 | 15 | 42997546 | T | A | 1247700 | 175 | 0.224 |
| rs112840248 | 15 | 42950885 | C | T | 1253119 | 176 | 0.223 |
| rs28562951 | 15 | 42998365 | T | C | 1247700 | 175 | 0.211 |
| rs4924690 | 15 | 42950219 | C | G | 1253119 | 176 | 0.223 |
| rs28479391 | 15 | 42963400 | A | G | 1247636 | 175 | 0.224 |
| rs9269 | 15 | 43012274 | A | G | 1253212 | 176 | 0.212 |
| rs8043472 | 15 | 42948862 | A | G | 1247615 | 175 | 0.224 |
| rs2136900 | 15 | 42993105 | A | G | 1247698 | 175 | 0.212 |
| rs7163030 | 15 | 42971620 | T | C | 1247657 | 175 | 0.223 |
| rs28744617 | 15 | 42981022 | A | G | 1253098 | 176 | 0.22 |
| rs16957020 | 15 | 42964426 | T | C | 1247649 | 175 | 0.212 |
| rs6493059 | 15 | 42977526 | A | C | 1247654 | 175 | 0.212 |
| rs67391479 | 15 | 43004351 | G | A | 1096338 | 166 | 0.22 |
| rs28823029 | 15 | 42952109 | G | C | 1247647 | 175 | 0.211 |
| rs10400824 | 15 | 42951277 | C | T | 1247651 | 175 | 0.212 |
| rs28811114 | 15 | 42952112 | C | T | 1247651 | 175 | 0.211 |
| rs9920163 | 15 | 42942381 | G | T | 1247642 | 175 | 0.211 |
| rs4924691 | 15 | 42950237 | T | C | 1253136 | 176 | 0.212 |
| rs28798437 | 15 | 42952005 | A | C | 1247652 | 175 | 0.212 |
| rs9919994 | 15 | 43011537 | G | T | 1247691 | 175 | 0.213 |
| rs60284149 | 15 | 42870823 | GT | G | 1063443 | 118 | 0.184 |
| rs623271 | 15 | 42928208 | G | A | 1240619 | 173 | 0.212 |
| rs1197551 | 15 | 42931717 | T | A | 1238999 | 172 | 0.209 |
| rs1705356 | 15 | 42922656 | C | T | 1238962 | 172 | 0.212 |
| rs1197549 | 15 | 42929154 | C | T | 1238992 | 172 | 0.212 |
| rs1197548 | 15 | 42928764 | G | A | 1238988 | 172 | 0.212 |
| rs1212814 | 15 | 42933603 | C | G | 1238998 | 172 | 0.212 |
| rs3106286 | 15 | 42927717 | T | C | 1238983 | 172 | 0.212 |
| rs3099758 | 15 | 42927984 | C | T | 1238989 | 172 | 0.212 |
| rs10162939 | 15 | 42930777 | A | T | 1239001 | 172 | 0.21 |
| rs9944248 | 15 | 42965191 | A | G | 1247672 | 175 | 0.2 |
| rs2136901 | 15 | 42993266 | G | A | 1247716 | 175 | 0.201 |
| rs35616173 | 15 | 43001799 | T | C | 1247720 | 175 | 0.201 |
| rs7166373 | 15 | 42954888 | C | T | 1247672 | 175 | 0.199 |
| rs4923950 | 15 | 42959743 | A | G | 1247675 | 175 | 0.2 |
| rs12905592 | 15 | 43002536 | G | A | 1247722 | 175 | 0.201 |
| rs34191150 | 15 | 43005718 | A | G | 1247717 | 175 | 0.201 |
| rs3106285 | 15 | 42927668 | T | C | 1238984 | 172 | 0.217 |
| rs7182479 | 15 | 42999565 | G | A | 1247718 | 175 | 0.199 |
| rs6493061 | 15 | 42988412 | C | A | 1247711 | 175 | 0.2 |
| rs3742993 | 15 | 42983923 | A | G | 1253191 | 176 | 0.188 |
| rs35738819 | 15 | 42966288 | G | A | 1247670 | 175 | 0.201 |
| rs2136899 | 15 | 42990386 | A | G | 1247714 | 175 | 0.189 |
| rs8030587 | 15 | 42981806 | G | A | 1253182 | 176 | 0.2 |
| rs12905187 | 15 | 43002534 | A | T | 1247720 | 175 | 0.189 |
| rs57298545 | 15 | 42956620 | A | G | 1247689 | 175 | 0.189 |
| rs4924694 | 15 | 42991066 | T | A | 1247713 | 175 | 0.189 |
| rs7176860 | 15 | 42998858 | G | A | 1247719 | 175 | 0.188 |
| rs938047 | 15 | 42955967 | A | G | 1247687 | 175 | 0.189 |
| rs6493060 | 15 | 42977676 | G | A | 1225881 | 169 | 0.2 |
| rs34603230 | 15 | 42958216 | A | G | 1247679 | 175 | 0.189 |
| rs4545789 | 15 | 42958928 | A | C | 1247691 | 175 | 0.189 |
| rs58938096 | 15 | 43005804 | G | T | 1247716 | 175 | 0.19 |
| rs4924692 | 15 | 42950297 | G | A | 1253174 | 176 | 0.188 |
| rs8039711 | 15 | 42957180 | G | A | 1247692 | 175 | 0.188 |
| rs8024902 | 15 | 42957532 | T | G | 1247691 | 175 | 0.188 |
| rs8039765 | 15 | 42957373 | C | A | 1247691 | 175 | 0.189 |
| rs7183487 | 15 | 42999553 | T | A | 1247716 | 175 | 0.19 |
| rs2175370 | 15 | 42960322 | G | A | 1247694 | 175 | 0.188 |
| rs12903762 | 15 | 43006385 | C | A | 1226163 | 174 | 0.19 |
| rs8031218 | 15 | 42982340 | C | T | 1253187 | 176 | 0.19 |
| rs1058846 | 15 | 43012024 | C | T | 1253233 | 176 | 0.18 |
| rs12902830 | 15 | 42997382 | C | T | 1253215 | 176 | 0.179 |
| rs672054 | 15 | 42918017 | C | T | 1238974 | 172 | 0.188 |
| rs71474496 | 15 | 42918141 | A | G | 1253062 | 176 | 0.176 |
| rs61122145 | 15 | 42991887 | G | A | 1253211 | 176 | 0.179 |
| rs11638835 | 15 | 42918341 | G | A | 1238986 | 172 | 0.178 |
| rs12911585 | 15 | 42934545 | T | C | 1253111 | 176 | 0.177 |
| rs1197550 | 15 | 42930891 | T | A | 999640 | 161 | 0.21 |
| rs2241998 | 15 | 42869028 | G | A | 1239007 | 172 | 0.247 |
| rs34070722 | 15 | 42938962 | G | A | 1247592 | 175 | 0.181 |
| rs398039432 | 15 | 43000191 | G | GA | 1071779 | 118 | 0.222 |
| rs12594855 | 15 | 42939904 | G | A | 1218080 | 167 | 0.186 |
| rs111810960 | 15 | 43007276 | CA | C | 1058031 | 116 | 0.206 |
| rs34117505 | 15 | 42925825 | C | T | 1182683 | 159 | 0.449 |
| rs35392023 | 15 | 43001660 | G | A | 1118207 | 134 | 0.176 |
| rs34868427 | 15 | 42995565 | GT | G | 1073372 | 119 | 0.229 |
| rs1667493 | 15 | 42889170 | T | C | 1231863 | 172 | 0.267 |
| rs530032660 | 15 | 42952212 | G | C | 787620 | 65 | 0.000663 |
| rs398043203 | 15 | 42933807 | CT | C | 1076692 | 120 | 0.186 |
| rs6493053 | 15 | 42886074 | A | G | 1240795 | 173 | 0.267 |
| rs61683241 | 15 | 43007718 | A | AT | 1063980 | 118 | 0.173 |
| rs35915660 | 15 | 42873866 | TCAG | T | 1075629 | 119 | 0.27 |
| rs12594951 | 15 | 42934631 | A | C | 1212273 | 167 | 0.475 |
| rs111730580 | 15 | 42907211 | AT | A | 648551 | 109 | 0.28 |
| rs12903416 | 15 | 42881039 | T | C | 1110560 | 132 | 0.279 |
| rs71474497 | 15 | 42951427 | T | C | 1227067 | 159 | 0.0424 |
| rs150746990 | 15 | 42932515 | A | T | 937314 | 91 | 0.00146 |
| rs9783683 | 15 | 42875276 | A | C | 1239121 | 172 | 0.282 |
| rs7168835 | 15 | 42876014 | C | T | 1240795 | 173 | 0.281 |
| rs191874576 | 15 | 42881098 | C | T | 416095 | 3 | 0.000127 |
| rs11399920 | 15 | 42917845 | AT | A | 1043714 | 115 | 0.0779 |
| rs180854628 | 15 | 42988525 | C | A | 397968 | 2 | 0.000629 |
| rs12903726 | 15 | 42954739 | A | C | 1249619 | 169 | 0.0217 |
| rs11632169 | 15 | 42903115 | A | G | 1238730 | 174 | 0.133 |
| rs11070376 | 15 | 42879757 | T | G | 1239142 | 172 | 0.126 |
| rs1197546 | 15 | 42902246 | G | A | 1238732 | 174 | 0.132 |
| rs1197547 | 15 | 42905340 | T | G | 1253134 | 176 | 0.283 |
| rs539464370 | 15 | 43008093 | C | T | 398221 | 3 | 0.000159 |
| rs12904010 | 15 | 42881210 | A | G | 1239138 | 172 | 0.126 |
| rs4447398 | 15 | 42904904 | C | A | 1238728 | 174 | 0.133 |
| rs544743742 | 15 | 42938738 | A | T | 402794 | 3 | 0.000161 |
| rs61750788 | 15 | 42953382 | T | G | 1251701 | 171 | 0.0247 |
| rs67470909 | 15 | 43010435 | T | C | 1253139 | 176 | 0.0769 |
| rs143379946 | 15 | 42974907 | C | T | 1242142 | 152 | 0.00976 |
| rs12904592 | 15 | 42890198 | G | A | 1060102 | 120 | 0.272 |
| rs62019357 | 15 | 42948374 | C | T | 1227999 | 161 | 0.096 |
| rs117066630 | 15 | 42953718 | T | C | 1251684 | 171 | 0.0227 |
| rs876992 | 15 | 42878456 | C | T | 1230227 | 171 | 0.131 |
| rs532715301 | 15 | 42908030 | A | G | 16199 | 5 | 0.00173 |
| rs767904319 | 15 | 42941881 | C | G | 627001 | 34 | 0.000478 |
| rs117954128 | 15 | 42974055 | G | A | 1251688 | 171 | 0.0224 |
| rs563472814 | 15 | 42883775 | T | C | 16199 | 5 | 0.00173 |
| rs557984924 | 15 | 42869457 | T | C | 397969 | 2 | 0.000908 |
| rs78562640 | 15 | 43002759 | A | G | 1216971 | 156 | 0.0291 |
| rs77298618 | 15 | 42876902 | T | G | 1250336 | 168 | 0.0206 |
| NA | 15 | 42894125 | A | C | 395738 | 3 | 0.000236 |
| NA | 15 | 42873849 | G | C | 406215 | 4 | 0.000242 |
| rs77003671 | 15 | 42896234 | T | C | 1250335 | 168 | 0.0205 |
| rs28498538 | 15 | 43011353 | G | C | 1251793 | 172 | 0.0267 |
| rs559353595 | 15 | 43009409 | G | A | 404155 | 4 | 0.000746 |
| rs78325423 | 15 | 43012549 | G | C | 1220348 | 157 | 0.0338 |
| rs12901400 | 15 | 42942262 | C | T | 1251792 | 172 | 0.0434 |
| rs35006192 | 15 | 42935698 | G | A | 1253090 | 176 | 0.0712 |
| rs145113087 | 15 | 43005947 | G | C | 809018 | 86 | 0.00111 |
| rs532786415 | 15 | 42902395 | G | T | 555698 | 32 | 0.000503 |
| rs563855481 | 15 | 42889884 | C | T | 433211 | 6 | 0.00012 |
| rs34928615 | 15 | 42994757 | GTCTA | G | 594940 | 67 | 0.00866 |
| rs548604256 | 15 | 42948022 | A | G | 306281 | 19 | 0.000968 |
| rs56286330 | 15 | 42950486 | G | GGT | 1029363 | 109 | 0.248 |
| rs67741694 | 15 | 42940334 | A | T | 1251774 | 172 | 0.0428 |
| rs548861293 | 15 | 42918759 | C | T | 399076 | 2 | 0.000108 |
| rs28431056 | 15 | 43010138 | C | T | 1202895 | 142 | 0.0055 |
| rs1018744820 | 15 | 42918643 | T | C | 564380 | 30 | 0.000331 |
| rs773669948 | 15 | 42883556 | C | T | 859232 | 65 | 0.00148 |
| rs565298 | 15 | 42893192 | C | A | 1238808 | 174 | 0.118 |
| rs73410940 | 15 | 43010574 | G | A | 1202895 | 142 | 0.00552 |
| rs12591415 | 15 | 43009399 | C | T | 1200528 | 141 | 0.00643 |
| rs147646076 | 15 | 42929139 | G | C | 1235442 | 151 | 0.00949 |
| rs6493054 | 15 | 42886504 | A | G | 1244304 | 175 | 0.117 |
| rs369934831 | 15 | 42955652 | A | G | 577454 | 18 | 0.000417 |
| rs113551877 | 15 | 42918582 | C | T | 922640 | 107 | 0.0229 |
| rs28664781 | 15 | 42953809 | T | G | 1222308 | 164 | 0.0303 |
| rs558776330 | 15 | 42929884 | T | C | 397969 | 2 | 0.000124 |
| rs544593985 | 15 | 42909007 | C | T | 1072352 | 73 | 0.00302 |
| rs28650274 | 15 | 42944829 | C | T | 1251706 | 171 | 0.0251 |
| rs1705360 | 15 | 42889047 | A | G | 1238807 | 174 | 0.117 |
| rs577245802 | 15 | 42901367 | AAG | A | 401375 | 4 | 0.000105 |
| rs557974103 | 15 | 42955918 | T | G | 397969 | 2 | 0.000402 |
| rs11637248 | 15 | 42882016 | G | C | 1234541 | 172 | 0.117 |
| rs533443278 | 15 | 42994547 | G | A | 637836 | 5 | 0.000175 |
| rs1814518 | 15 | 42873672 | G | A | 1239158 | 172 | 0.117 |
| rs766798280 | 15 | 42949754 | A | G | 402209 | 4 | 0.000129 |
| rs186242531 | 15 | 42880107 | C | T | 92401 | 11 | 0.000455 |
| rs1044735653 | 15 | 43005905 | C | T | 591790 | 18 | 0.000415 |
| rs28882725 | 15 | 42924027 | A | T | 1219610 | 161 | 0.0259 |
| rs114824872 | 15 | 42920562 | G | A | 861738 | 49 | 0.000641 |
| rs538043621 | 15 | 42911052 | G | A | 658685 | 35 | 0.000584 |
| rs78261202 | 15 | 43003113 | G | C | 751608 | 58 | 0.00954 |
| rs55773330 | 15 | 42956987 | AAC | A | 1052427 | 115 | 0.232 |
| rs138641635 | 15 | 42998558 | A | C | 1241681 | 153 | 0.00518 |
| rs202031252 | 15 | 42935804 | TCGGCTGAGC | T | 29486 | 8 | 0.00436 |
| rs185165874 | 15 | 42982303 | G | C | 393950 | 2 | 0.000208 |
| rs761481866 | 15 | 42902640 | G | T | 651490 | 36 | 0.000438 |
| rs931528965 | 15 | 42918444 | T | C | 513086 | 13 | 0.000374 |
| rs147330525 | 15 | 42999707 | G | A | 774296 | 13 | 0.000535 |
| rs200378215 | 15 | 42955816 | A | G | 1245393 | 157 | 0.0058 |
| rs564756186 | 15 | 42959022 | A | AT | 627296 | 3 | 0.000313 |
| rs568429587 | 15 | 42934240 | AT | A | 896229 | 79 | 0.00437 |
| NA | 15 | 42954809 | C | A | 530411 | 13 | 0.000377 |
| rs145395424 | 15 | 42904938 | T | C | 1217620 | 134 | 0.0052 |
| rs573466110 | 15 | 42933777 | A | G | 401746 | 3 | 0.000536 |
| rs766573983 | 15 | 42941572 | A | G | 579035 | 36 | 0.000294 |
| rs373035305 | 15 | 42867925 | G | A | 443872 | 7 | 0.000207 |
| NA | 15 | 42948935 | C | T | 408980 | 4 | 0.00011 |
| rs771779746 | 15 | 42982372 | C | G | 640063 | 10 | 0.000344 |
| rs479519 | 15 | 42892809 | C | T | 397287 | 4 | 0.000434 |
| rs763038765 | 15 | 42908885 | C | T | 29486 | 8 | 0.00161 |
| rs74009101 | 15 | 42895210 | T | C | 691189 | 10 | 0.000225 |
| rs7175580 | 15 | 42959671 | G | A | 238830 | 2 | 0.000113 |
| rs370116985 | 15 | 42918624 | T | C | 49381 | 12 | 0.000334 |
| rs554337611 | 15 | 42881817 | T | C | 571652 | 13 | 0.000168 |
| rs148862329 | 15 | 42953372 | A | G | 1190456 | 125 | 0.00524 |
| rs2136903 | 15 | 42941469 | A | G | 1121906 | 135 | 0.0412 |
| rs748629198 | 15 | 42910271 | G | A | 479614 | 6 | 9.90E-05 |
| rs539697836 | 15 | 42889573 | A | T | 311658 | 15 | 0.000319 |
| rs553736195 | 15 | 43011409 | A | C | 393138 | 2 | 5.09E-05 |
| rs547182963 | 15 | 43011781 | C | T | 401013 | 4 | 8.85E-05 |
| rs563268103 | 15 | 42925722 | A | G | 393138 | 2 | 0.000248 |
| rs199940227 | 15 | 42984727 | T | G | 364275 | 22 | 0.000528 |
| rs574547539 | 15 | 42920203 | T | G | 729553 | 58 | 0.000663 |
| rs142114926 | 15 | 42991816 | T | C | 1229889 | 155 | 0.0133 |
| rs967418990 | 15 | 42893416 | G | T | 507038 | 7 | 8.10E-05 |
| rs536290738 | 15 | 42947564 | A | G | 442707 | 7 | 0.000282 |
| rs147055430 | 15 | 43005504 | T | A | 393955 | 2 | 0.000114 |
| rs565991513 | 15 | 42909088 | CA | C | 961541 | 115 | 0.0248 |
| rs1042575965 | 15 | 42956237 | G | C | 571730 | 18 | 0.000365 |
| rs537604330 | 15 | 43003401 | G | A | 389982 | 2 | 0.00416 |
| rs544454539 | 15 | 42964412 | C | T | 640705 | 6 | 0.000121 |
| rs537145859 | 15 | 42918480 | CTCCTCT | C | 718679 | 26 | 0.000673 |
| rs565180029 | 15 | 43004581 | C | T | 408747 | 5 | 0.000927 |
| rs113206636 | 15 | 43010551 | G | GGGGA | 1072590 | 106 | 0.00551 |
| rs559750005 | 15 | 42893677 | G | A | 612752 | 24 | 0.000294 |
| rs574256734 | 15 | 42898994 | A | G | 414158 | 6 | 0.00227 |
| rs140844877 | 15 | 42916792 | G | C | 1198559 | 133 | 0.00537 |
| rs542874709 | 15 | 42949334 | C | T | 574769 | 21 | 0.000242 |
| rs557162342 | 15 | 42888744 | G | A | 1043834 | 108 | 0.0015 |
| rs146581067 | 15 | 42947072 | A | G | 1248413 | 167 | 0.0115 |
| rs531975840 | 15 | 42880623 | G | A | 404155 | 4 | 0.000468 |
| rs755384675 | 15 | 42967612 | A | G | 407939 | 3 | 0.000127 |
| rs866926503 | 15 | 43005024 | C | T | 713447 | 51 | 0.000481 |
| rs779412480 | 15 | 42882189 | C | T | 678351 | 18 | 0.00091 |
| rs115196427 | 15 | 43008297 | A | G | 238830 | 3 | 6.28E-05 |
| rs551482565 | 15 | 42905800 | C | T | 697272 | 13 | 0.000151 |
| rs191282954 | 15 | 42915084 | G | T | 404155 | 4 | 0.000379 |
| rs672150 | 15 | 42918082 | C | T | 669373 | 16 | 0.000801 |
| rs149018034 | 15 | 42948125 | C | T | 764711 | 24 | 0.000332 |
| rs180869067 | 15 | 42877641 | G | T | 484782 | 4 | 0.000105 |
| rs369310511 | 15 | 42976363 | C | T | 586369 | 20 | 0.00031 |
| rs545253441 | 15 | 42970464 | C | T | 66244 | 12 | 0.000611 |
| rs62019355 | 15 | 42938163 | G | A | 1227899 | 151 | 0.00896 |
| rs115788605 | 15 | 42947772 | A | G | 764712 | 24 | 0.000335 |
| rs766696626 | 15 | 42955169 | A | G | 783447 | 38 | 0.000708 |
| rs187403913 | 15 | 42909578 | G | A | 498310 | 16 | 0.000118 |
| rs79313070 | 15 | 43002191 | G | T | 1177281 | 132 | 0.00423 |
| rs138423727 | 15 | 43005653 | T | C | 1244303 | 167 | 0.0412 |
| rs144343572 | 15 | 42871691 | C | T | 767761 | 27 | 0.00044 |
| rs114984341 | 15 | 42879064 | C | T | 771538 | 28 | 0.000439 |
| rs117143870 | 15 | 42938044 | C | T | 1239394 | 148 | 0.00468 |
| rs79994978 | 15 | 42902639 | T | C | 776368 | 29 | 0.000443 |
| rs548883046 | 15 | 42921116 | T | G | 810882 | 104 | 0.00206 |
| rs187407014 | 15 | 42931263 | A | G | 895564 | 66 | 0.000909 |
| rs778633745 | 15 | 42874075 | A | G | 427291 | 6 | 0.000173 |
| rs78263141 | 15 | 42880383 | G | T | 771538 | 28 | 0.000439 |
| rs142547271 | 15 | 42948936 | G | A | 816587 | 94 | 0.00119 |
| rs182793324 | 15 | 42959071 | G | A | 397969 | 2 | 0.00162 |
| rs555489047 | 15 | 42886831 | G | GT | 401456 | 5 | 0.00125 |
| rs184321765 | 15 | 42881661 | A | G | 393388 | 4 | 4.19E-05 |
| rs544510397 | 15 | 42938358 | T | TCAA | 69843 | 2 | 0.00145 |
| rs539637890 | 15 | 42970185 | C | G | 931794 | 103 | 0.00244 |
| rs544111850 | 15 | 42920659 | T | C | 558817 | 14 | 0.000167 |
| rs79158309 | 15 | 42877590 | A | G | 767761 | 27 | 0.000442 |
| rs117807571 | 15 | 42932609 | C | T | 414158 | 6 | 0.00141 |
| rs570354852 | 15 | 42884271 | G | GTGATC | 717296 | 82 | 0.00539 |
| rs116565691 | 15 | 42888132 | G | A | 632244 | 5 | 8.94E-05 |
| rs78196628 | 15 | 42921854 | C | T | 1210609 | 132 | 0.00394 |
| rs111612014 | 15 | 42874007 | A | G | 684177 | 3 | 0.000371 |
| rs186290703 | 15 | 42932876 | C | T | 800349 | 109 | 0.00249 |
| rs74345837 | 15 | 43005264 | T | C | 1244425 | 167 | 0.0599 |
| rs558891400 | 15 | 42956104 | C | T | 665639 | 39 | 0.000502 |
| rs151006266 | 15 | 42969999 | G | C | 515113 | 29 | 0.000592 |
| rs35842593 | 15 | 42985999 | C | A | 1228818 | 150 | 0.00893 |
| rs565169604 | 15 | 42936375 | C | T | 390860 | 2 | 0.000432 |
| rs184949186 | 15 | 42954225 | T | A | 19463 | 3 | 0.00162 |
| rs747499658 | 15 | 42935069 | T | C | 689623 | 17 | 0.000396 |
| rs138819618 | 15 | 42891380 | C | T | 1231047 | 153 | 0.00947 |
| rs535717612 | 15 | 42909006 | A | G | 1000604 | 94 | 0.00214 |
| rs749762665 | 15 | 42911969 | T | C | 616167 | 36 | 0.000471 |
| rs577186934 | 15 | 42957491 | T | C | 493290 | 15 | 0.000504 |
| rs769089032 | 15 | 42910077 | C | T | 475343 | 3 | 0.000557 |
| rs370894209 | 15 | 42918444 | TTCTTCC | T | 728346 | 55 | 0.00957 |
| rs191659987 | 15 | 42874061 | A | G | 599625 | 39 | 0.00371 |
| rs544793721 | 15 | 42979703 | A | G | 412075 | 5 | 0.00142 |
| rs1050706910 | 15 | 42899088 | A | G | 434786 | 5 | 0.000107 |

|  |  |  |  |  |  |  |  |
| --- | --- | --- | --- | --- | --- | --- | --- |
| rs576555473 | 15 | 42919375 | T | C | 595466 | 23 | 0.000259 |
| rs764967488 | 15 | 42949280 | C | T | 453625 | 8 | 9.59E-05 |
| rs537835505 | 15 | 42915072 | G | A | 398916 | 3 | 0.00577 |
| rs149873427 | 15 | 42893027 | C | T | 719261 | 30 | 0.000918 |
| rs566475856 | 15 | 42902894 | G | A | 952771 | 48 | 0.000577 |
| rs11314415 | 15 | 42916751 | CT | C | 817978 | 55 | 0.0738 |
| rs751662687 | 15 | 42897263 | A | G | 566128 | 16 | 0.000401 |
| rs16957031 | 15 | 42969114 | G | A | 471928 | 3 | 0.000125 |
| rs186462510 | 15 | 43005172 | A | G | 403068 | 4 | 0.000572 |
| rs746547270 | 15 | 42897884 | G | A | 607245 | 7 | 0.000394 |
| rs12442544 | 15 | 42995418 | C | G | 688809 | 71 | 0.000736 |
| NA | 15 | 42882955 | R | I | 1339 | 2 | 0.0564 |
| rs117112275 | 15 | 42887565 | T | C | 1243846 | 156 | 0.0072 |
| rs148181870 | 15 | 42911523 | A | G | 1218890 | 129 | 0.00199 |
| rs779112444 | 15 | 42992926 | C | A | 457643 | 9 | 0.000116 |
| rs140235780 | 15 | 42995538 | C | T | 459940 | 7 | 0.000102 |
| rs762148444 | 15 | 42929474 | G | A | 629425 | 29 | 0.000246 |
| NA | 15 | 42957127 | G | T | 94449 | 7 | 0.00036 |
| rs148176549 | 15 | 43009749 | G | A | 1251654 | 171 | 0.0307 |
| rs766166696 | 15 | 42946519 | C | T | 161496 | 10 | 0.000167 |
| rs563833340 | 15 | 42977303 | G | A | 465888 | 24 | 0.000963 |
| rs115582718 | 15 | 42996987 | A | G | 832668 | 23 | 0.000384 |
| rs115969538 | 15 | 42896702 | T | C | 619457 | 3 | 7.10E-05 |
| rs146015002 | 15 | 42929887 | A | G | 631733 | 4 | 0.000306 |
| rs139768798 | 15 | 42879951 | C | T | 393759 | 3 | 0.000857 |
| rs563962097 | 15 | 42924629 | A | G | 902907 | 83 | 0.00161 |
| rs115361836 | 15 | 42965711 | C | A | 753890 | 23 | 0.000255 |
| rs537317862 | 15 | 42903870 | G | A | 587344 | 22 | 0.000318 |
| rs147467293 | 15 | 42916903 | C | A | 568341 | 10 | 0.000141 |
| rs548121692 | 15 | 42915911 | G | A | 402071 | 3 | 0.000502 |
| rs398027029 | 15 | 42870304 | A | AT | 1051768 | 116 | 0.165 |
| rs1027710403 | 15 | 42925755 | A | C | 420291 | 4 | 0.000271 |
| rs184215065 | 15 | 43008342 | G | C | 394085 | 2 | 0.00297 |
| rs542412856 | 15 | 42868135 | G | A | 614586 | 2 | 3.74E-05 |
| rs567027783 | 15 | 42900648 | T | A | 404155 | 4 | 0.00062 |
| rs535171426 | 15 | 42946687 | C | A | 397969 | 2 | 0.000211 |
| rs562182558 | 15 | 42880682 | G | T | 630688 | 4 | 0.000292 |
| rs574691968 | 15 | 42935706 | T | TA | 650709 | 13 | 0.00159 |
| rs187495317 | 15 | 42963968 | G | A | 987773 | 145 | 0.00789 |
| rs200568949 | 15 | 42978892 | A | G | 951983 | 62 | 0.000674 |
| rs567681136 | 15 | 42913727 | T | C | 22048 | 5 | 0.00152 |
| rs779017439 | 15 | 42970028 | G | A | 426496 | 5 | 0.000205 |
| rs751263954 | 15 | 43006721 | G | A | 446220 | 7 | 4.15E-05 |
| rs530299717 | 15 | 42922023 | A | C | 397968 | 2 | 0.00108 |
| rs565782439 | 15 | 42994289 | A | G | 397969 | 2 | 0.000271 |
| rs143056573 | 15 | 42874004 | A | G | 614586 | 2 | 4.88E-05 |
| rs182572446 | 15 | 42878337 | G | C | 629450 | 31 | 0.000635 |
| rs1197545 | 15 | 42893485 | A | G | 1101626 | 131 | 0.117 |

|  |  |  |  |  |  |  |  |
| --- | --- | --- | --- | --- | --- | --- | --- |
| rs533752199 | 15 | 42992096 | G | A | 397969 | 2 | 0.000285 |
| rs58903190 | 15 | 42946958 | C | T | 858074 | 29 | 0.000224 |
| rs1009186043 | 15 | 42989671 | A | G | 468001 | 5 | 8.23E-05 |
| rs181335046 | 15 | 43001629 | A | G | 1212450 | 132 | 0.00566 |
| rs187388117 | 15 | 43011625 | C | T | 907058 | 94 | 0.00226 |
| rs531993169 | 15 | 42916749 | G | T | 422277 | 8 | 0.00412 |
| rs552109201 | 15 | 42885029 | T | C | 392813 | 2 | 0.000892 |
| rs772067408 | 15 | 42982047 | G | T | 94449 | 7 | 0.000429 |
| rs1211642 | 15 | 42890450 | C | T | 1101626 | 131 | 0.118 |
| rs539858496 | 15 | 42904674 | C | T | 609017 | 39 | 0.000401 |
| rs183773847 | 15 | 42915614 | A | C | 922419 | 79 | 0.00374 |
| rs557804844 | 15 | 42883641 | G | A | 392813 | 2 | 0.000894 |
| rs553534463 | 15 | 42993913 | C | T | 644892 | 33 | 0.00046 |
| rs147950526 | 15 | 42913198 | C | T | 705294 | 8 | 0.000196 |
| rs570528702 | 15 | 42918366 | C | T | 397732 | 3 | 0.000138 |
| rs563158318 | 15 | 42932912 | G | A | 394897 | 3 | 7.85E-05 |
| rs561457101 | 15 | 43011558 | A | G | 459133 | 18 | 0.00051 |
| rs16957002 | 15 | 42961681 | G | A | 720332 | 13 | 0.00015 |
| rs530185347 | 15 | 42923671 | G | T | 607574 | 25 | 0.000269 |
| rs566759400 | 15 | 42942150 | C | T | 647572 | 35 | 0.000417 |
| rs771776965 | 15 | 42908221 | C | T | 84680 | 4 | 0.000218 |
| rs150157757 | 15 | 42919458 | T | C | 721893 | 102 | 0.00213 |
| rs529979641 | 15 | 43010416 | T | G | 264063 | 3 | 0.000148 |
| rs775555339 | 15 | 42899654 | C | G | 519853 | 3 | 0.000198 |
| rs141270837 | 15 | 42998672 | A | G | 1202886 | 146 | 0.00761 |
| rs192615557 | 15 | 42954183 | G | T | 394897 | 3 | 7.85E-05 |
| rs191154656 | 15 | 42874605 | A | T | 397732 | 3 | 9.81E-05 |
| rs188267549 | 15 | 42939749 | C | G | 1237103 | 145 | 0.00551 |
| rs114995589 | 15 | 42961882 | G | A | 720332 | 13 | 0.000149 |
| rs74244935 | 15 | 42871706 | A | G | 469895 | 18 | 0.000147 |
| rs1877138 | 15 | 42960674 | A | G | 760690 | 24 | 0.000285 |
| rs189297009 | 15 | 42955222 | A | G | 715798 | 56 | 0.000548 |
| rs551937058 | 15 | 42893450 | C | T | 397732 | 3 | 9.81E-05 |
| rs573580149 | 15 | 42911336 | A | C | 398999 | 4 | 0.00118 |
| rs397722507 | 15 | 42953149 | GT | G | 389982 | 2 | 0.184 |
| NA | 15 | 42953936 | T | A | 395960 | 3 | 7.96E-05 |
| rs114142198 | 15 | 42901178 | C | T | 229897 | 2 | 6.74E-05 |
| rs564907467 | 15 | 42952039 | G | A | 606896 | 27 | 0.000274 |
| rs557922265 | 15 | 42988143 | A | G | 625012 | 5 | 0.00164 |
| rs547275377 | 15 | 42872870 | C | T | 489230 | 35 | 0.00143 |
| rs528091580 | 15 | 42925476 | AG | A | 458628 | 2 | 0.00171 |
| rs577549839 | 15 | 42940429 | C | T | 105500 | 15 | 0.000398 |
| rs145552657 | 15 | 42996251 | A | G | 1250292 | 169 | 0.0252 |
| rs16956991 | 15 | 42958767 | G | C | 760690 | 24 | 0.000297 |
| rs3742994 | 15 | 42983742 | C | A | 424684 | 8 | 0.000102 |
| NA | 15 | 42949475 | T | C | 26987 | 2 | 0.00106 |
| rs536362397 | 15 | 42984299 | T | G | 402692 | 4 | 0.00692 |
| rs539014277 | 15 | 43004880 | A | G | 614586 | 2 | 4.39E-05 |

|  |  |  |  |  |  |  |  |
| --- | --- | --- | --- | --- | --- | --- | --- |
| rs145276351 | 15 | 42914910 | A | G | 614586 | 2 | 5.53E-05 |
| rs138442121 | 15 | 42948357 | C | T | 1236482 | 157 | 0.0248 |
| rs1001761276 | 15 | 42919787 | C | T | 622500 | 35 | 0.000235 |
| rs114435521 | 15 | 42938902 | G | A | 708242 | 13 | 0.000337 |
| rs4923951 | 15 | 43004028 | G | A | 556335 | 36 | 0.000328 |
| rs114829780 | 15 | 42940955 | T | C | 753155 | 20 | 0.000238 |
| rs147888154 | 15 | 42877151 | G | A | 677773 | 15 | 0.000662 |
| rs541664623 | 15 | 42961778 | C | T | 391120 | 2 | 0.00114 |
| rs570324067 | 15 | 42974474 | G | A | 509747 | 59 | 0.00136 |
| rs542582189 | 15 | 42979085 | G | A | 46909 | 8 | 0.00033 |
| rs147473758 | 15 | 42883617 | C | T | 1179051 | 118 | 0.00223 |
| NA | 15 | 42911211 | A | C | 26361 | 5 | 0.00451 |
| rs577253212 | 15 | 42922291 | T | C | 412309 | 10 | 0.00469 |
| rs546146720 | 15 | 42945342 | T | A | 595467 | 23 | 0.000266 |
| NA | 15 | 42896073 | C | G | 29486 | 8 | 0.00285 |
| rs397729129 | 15 | 42929277 | TA | T | 754500 | 58 | 0.0345 |
| rs544010780 | 15 | 42941389 | G | A | 617992 | 4 | 7.04E-05 |
| rs117100467 | 15 | 43008831 | G | A | 1235377 | 146 | 0.0081 |
| rs140632314 | 15 | 42941228 | C | T | 1245579 | 156 | 0.00612 |
| rs568687731 | 15 | 42992682 | A | T | 392813 | 2 | 0.00041 |
| rs76175552 | 15 | 42887079 | A | T | 1237194 | 147 | 0.00831 |
| rs577527530 | 15 | 42868980 | A | G | 504483 | 40 | 0.00137 |
| rs575145889 | 15 | 42979732 | C | T | 614586 | 2 | 4.31E-05 |
| rs548633040 | 15 | 42999100 | G | A | 409690 | 6 | 0.00112 |
| rs145521822 | 15 | 42914681 | T | C | 438562 | 12 | 0.00158 |
| rs751457103 | 15 | 42931670 | A | G | 728392 | 57 | 0.000389 |
| rs74890375 | 15 | 42958672 | A | G | 242607 | 4 | 0.000221 |
| rs865821954 | 15 | 42905124 | G | T | 489667 | 21 | 0.000318 |
| rs114690791 | 15 | 42937403 | A | G | 782206 | 20 | 0.000271 |
| rs529049962 | 15 | 43007788 | G | A | 614586 | 2 | 3.42E-05 |
| rs532387773 | 15 | 42899343 | A | G | 507757 | 19 | 0.000399 |
| rs541928952 | 15 | 42902878 | G | A | 614586 | 2 | 7.89E-05 |
| rs569316332 | 15 | 42901679 | C | G | 739475 | 68 | 0.00108 |
| rs963090177 | 15 | 42915175 | C | T | 394638 | 2 | 0.000115 |
| rs146377113 | 15 | 42975687 | A | G | 647494 | 12 | 0.000619 |
| rs116936632 | 15 | 42924234 | G | T | 1222216 | 164 | 0.0318 |
| rs116820442 | 15 | 42921051 | C | T | 756948 | 11 | 0.000785 |
| rs575362514 | 15 | 42989281 | A | G | 462484 | 10 | 0.000165 |
| rs185252821 | 15 | 42868239 | G | C | 412420 | 5 | 0.000108 |
| rs193137901 | 15 | 43006356 | G | A | 955182 | 62 | 0.00111 |
| rs55959547 | 15 | 42976518 | A | G | 1236888 | 160 | 0.0168 |
| rs189193514 | 15 | 42928407 | T | C | 796656 | 66 | 0.00453 |
| rs1877137 | 15 | 42960383 | G | T | 760690 | 24 | 0.000288 |
| rs564597643 | 15 | 42930718 | CA | C | 618602 | 40 | 0.00672 |
| NA | 15 | 42931548 | C | T | 6702 | 2 | 0.00545 |
| rs551035769 | 15 | 42937527 | G | A | 592314 | 20 | 0.000157 |
| rs189800151 | 15 | 42960589 | T | C | 1206698 | 138 | 0.00863 |
| rs567976615 | 15 | 42979462 | G | A | 395005 | 3 | 0.00181 |

|  |  |  |  |  |  |  |  |
| --- | --- | --- | --- | --- | --- | --- | --- |
| rs150250448 | 15 | 42878578 | G | T | 773223 | 12 | 0.000118 |
| rs188636270 | 15 | 42884194 | A | T | 140892 | 18 | 0.000614 |
| rs16957052 | 15 | 42979461 | C | T | 1153231 | 92 | 0.0012 |
| rs201923100 | 15 | 42944469 | TA | T | 1066849 | 102 | 0.0164 |
| rs12594724 | 15 | 42999650 | A | G | 1218469 | 154 | 0.00943 |
| rs139898805 | 15 | 42951125 | G | A | 614586 | 2 | 3.99E-05 |
| rs147801000 | 15 | 42963833 | AC | A | 614586 | 2 | 0.000193 |
| rs568974261 | 15 | 42929329 | C | T | 614586 | 2 | 3.42E-05 |
| rs546811853 | 15 | 42942308 | G | A | 395005 | 3 | 0.00168 |
| rs569220966 | 15 | 42943178 | A | G | 403830 | 4 | 0.000457 |
| rs144624716 | 15 | 42959424 | C | T | 1236488 | 157 | 0.0247 |
| NA | 15 | 42988850 | G | A | 20834 | 3 | 0.0061 |
| rs4334272 | 15 | 42994169 | T | C | 1218469 | 154 | 0.00942 |
| rs757758595 | 15 | 42959406 | G | A | 414826 | 4 | 0.000189 |
| rs768060909 | 15 | 42962501 | A | G | 539813 | 10 | 0.000287 |
| NA | 15 | 42923365 | A | C | 581462 | 25 | 0.000377 |
| rs142703729 | 15 | 42915717 | A | G | 1232445 | 145 | 0.00843 |
| rs544840606 | 15 | 42879972 | A | G | 458627 | 2 | 0.000176 |
| rs750810228 | 15 | 42948041 | A | G | 67112 | 8 | 0.000253 |
| rs539519966 | 15 | 42918498 | C | T | 403031 | 4 | 0.000756 |
| rs780649253 | 15 | 42933552 | A | G | 443540 | 6 | 0.000347 |
| rs138657899 | 15 | 42919534 | T | C | 658895 | 42 | 7.00E-04 |
| rs188271346 | 15 | 42910848 | G | A | 634840 | 34 | 0.000573 |
| rs182258163 | 15 | 42936560 | G | A | 1236829 | 147 | 0.00446 |
| rs112458684 | 15 | 42951912 | C | T | 1237434 | 146 | 0.00566 |
| rs4923949 | 15 | 42892748 | A | G | 1047045 | 106 | 0.00731 |
| rs148793160 | 15 | 42882955 | C | CGA | 1107820 | 124 | 0.0681 |
| rs141192915 | 15 | 42897985 | C | G | 397969 | 2 | 0.000563 |
| rs143444286 | 15 | 42984256 | G | A | 233429 | 2 | 0.000159 |
| rs186674983 | 15 | 42911925 | G | A | 1235353 | 144 | 0.00415 |
| rs117478364 | 15 | 42959003 | A | T | 1217047 | 152 | 0.00852 |
| rs145017779 | 15 | 42898095 | A | G | 614586 | 2 | 5.78E-05 |
| rs72711791 | 15 | 43007646 | A | G | 1253180 | 176 | 0.0731 |
| rs141314082 | 15 | 42900961 | A | T | 614586 | 2 | 5.86E-05 |
| rs145590620 | 15 | 42933274 | T | C | 1236831 | 147 | 0.00435 |
| rs115422270 | 15 | 42871599 | G | C | 614586 | 2 | 4.96E-05 |
| rs117506318 | 15 | 42872647 | C | G | 1215207 | 132 | 0.0045 |
| rs143795631 | 15 | 42901120 | T | C | 614586 | 2 | 5.86E-05 |
| rs187874698 | 15 | 42934546 | C | T | 614586 | 2 | 8.87E-05 |
| rs569388978 | 15 | 42953642 | C | T | 398915 | 3 | 0.00605 |
| NA | 15 | 42990765 | C | T | 76317 | 2 | 0.00199 |
| rs777344032 | 15 | 42872371 | G | A | 6702 | 2 | 0.00463 |
| rs138864471 | 15 | 42936564 | A | G | 1236137 | 146 | 0.00446 |
| rs149176049 | 15 | 42982891 | A | C | 1216357 | 126 | 0.00198 |
| rs1038998860 | 15 | 42872719 | A | T | 457987 | 10 | 0.000218 |
| rs144867366 | 15 | 42973022 | A | G | 547103 | 37 | 0.00133 |
| rs150402119 | 15 | 42894680 | G | C | 974390 | 111 | 0.00248 |
| rs565472380 | 15 | 42979114 | T | C | 411611 | 5 | 0.00311 |
| rs191875909 | 15 | 42901586 | T | A | 614586 | 2 | 5.86E-05 |
| rs541693905 | 15 | 42920232 | A | T | 620639 | 45 | 0.000516 |
| rs117731492 | 15 | 42934853 | A | G | 1236828 | 147 | 0.00434 |
| rs146027773 | 15 | 43009197 | C | T | 887981 | 45 | 0.000896 |
| rs184838385 | 15 | 42906543 | A | G | 614586 | 2 | 5.94E-05 |
| rs534849153 | 15 | 42914548 | C | CCTCT | 849260 | 75 | 0.00172 |
| rs147213920 | 15 | 42984783 | T | C | 578499 | 13 | 0.000328 |
| rs188728159 | 15 | 42943279 | A | C | 397969 | 2 | 0.000751 |
| rs760584528 | 15 | 42966962 | G | T | 937777 | 88 | 0.0012 |
| rs774795729 | 15 | 42945865 | A | G | 707884 | 22 | 0.00067 |
| NA | 15 | 43007718 | R | I | 1339 | 2 | 0.152 |
| rs148698759 | 15 | 42903408 | C | T | 614586 | 2 | 5.94E-05 |
| NA | 15 | 42908082 | G | C | 6702 | 2 | 0.00418 |
| rs74475031 | 15 | 42979146 | A | G | 1187160 | 141 | 0.00714 |
| rs765294767 | 15 | 42876417 | T | G | 515721 | 12 | 0.000486 |
| rs539573579 | 15 | 42883444 | G | C | 1141947 | 71 | 0.000947 |
| rs188303137 | 15 | 42888701 | G | A | 941816 | 107 | 0.00393 |
| rs187216270 | 15 | 42904022 | A | T | 633707 | 39 | 0.000383 |
| rs72711781 | 15 | 42952175 | A | G | 1251652 | 175 | 0.0739 |
| rs551993514 | 15 | 42992228 | C | T | 537976 | 31 | 0.000576 |
| rs150895411 | 15 | 42889015 | T | G | 1202287 | 126 | 0.0049 |
| rs574731559 | 15 | 42941284 | A | G | 393383 | 2 | 7.63E-05 |
| rs916413970 | 15 | 42946588 | A | G | 442218 | 10 | 4.41E-05 |
| rs140243268 | 15 | 43000117 | A | G | 1245090 | 156 | 0.00827 |
| rs534886977 | 15 | 42910451 | C | T | 397969 | 2 | 0.000469 |
| rs572358923 | 15 | 43000193 | A | G | 412821 | 6 | 0.000785 |
| rs181167607 | 15 | 42906448 | G | T | 1230572 | 148 | 0.0109 |
| rs150633440 | 15 | 42916646 | A | G | 394306 | 4 | 0.000795 |
| rs74012616 | 15 | 42970814 | A | G | 652845 | 8 | 0.000121 |
| rs141601398 | 15 | 42927449 | A | C | 652423 | 7 | 0.000454 |
| rs765034095 | 15 | 42997643 | T | G | 405752 | 2 | 6.16E-05 |
| rs751302053 | 15 | 42871640 | G | A | 860337 | 64 | 0.00135 |
| rs189832961 | 15 | 42922492 | C | T | 401160 | 3 | 0.000147 |
| rs529982887 | 15 | 42878999 | A | G | 511859 | 10 | 0.000144 |
| rs561258261 | 15 | 42936422 | C | T | 401746 | 3 | 0.00039 |
| rs1052365338 | 15 | 42897631 | G | T | 29486 | 8 | 0.00149 |
| rs566366165 | 15 | 42929197 | G | A | 805090 | 82 | 0.00146 |
| rs563666899 | 15 | 42944936 | C | T | 949767 | 98 | 0.00136 |
| rs143155792 | 15 | 42921007 | C | T | 234483 | 2 | 0.000171 |
| rs556518794 | 15 | 42871439 | A | G | 511859 | 10 | 0.000144 |
| NA | 15 | 42991741 | G | A | 94449 | 7 | 0.000556 |
| rs544507849 | 15 | 42999666 | CT | C | 1036326 | 115 | 0.176 |
| rs547797147 | 15 | 42947514 | C | T | 712963 | 29 | 0.00234 |
| rs569045927 | 15 | 42959642 | A | G | 714215 | 30 | 0.00233 |
| rs146134970 | 15 | 42932113 | G | C | 722014 | 8 | 0.000389 |
| rs574542771 | 15 | 42922405 | G | A | 791527 | 54 | 0.00164 |
| rs181996302 | 15 | 42887884 | T | C | 767168 | 15 | 0.000184 |
| rs185512143 | 15 | 42943200 | C | T | 418917 | 6 | 9.43E-05 |
| rs183583949 | 15 | 42959536 | C | T | 727290 | 62 | 0.00106 |
| rs139864533 | 15 | 42889447 | C | T | 761784 | 67 | 0.000869 |
| rs117015041 | 15 | 42912674 | T | A | 685078 | 13 | 0.000475 |
| rs574639690 | 15 | 42923049 | T | G | 565791 | 30 | 0.00163 |
| rs35259107 | 15 | 42953805 | GCTTT | G | 956020 | 58 | 0.00638 |
| rs182241507 | 15 | 42967326 | C | G | 403830 | 4 | 0.000337 |
| rs12324135 | 15 | 42942696 | C | T | 1250553 | 167 | 0.011 |
| rs192131974 | 15 | 42963378 | T | C | 719520 | 7 | 0.000459 |
| rs1054031892 | 15 | 42910298 | A | G | 412810 | 4 | 0.000199 |
| rs148448542 | 15 | 42911342 | A | G | 1163182 | 103 | 0.00269 |
| rs140183846 | 15 | 42916970 | C | A | 652424 | 7 | 0.000408 |
| rs755153315 | 15 | 42929877 | A | G | 441236 | 7 | 0.000142 |
| rs775814927 | 15 | 42953874 | C | T | 96964 | 13 | 0.00017 |
| rs116478053 | 15 | 42963301 | C | G | 719521 | 7 | 0.000459 |
| rs567987273 | 15 | 42973305 | T | C | 416582 | 3 | 5.76E-05 |
| rs541890756 | 15 | 42931425 | G | A | 581694 | 19 | 0.000425 |
| rs188976332 | 15 | 43005416 | C | G | 234483 | 2 | 0.000117 |
| rs564138574 | 15 | 43007719 | T | A | 416093 | 9 | 0.00398 |
| rs542175917 | 15 | 42916758 | T | C | 833198 | 54 | 0.00132 |
| rs75021400 | 15 | 42981377 | C | T | 891650 | 67 | 0.000406 |
| rs12440556 | 15 | 42874506 | C | G | 890446 | 41 | 0.000537 |
| rs770986137 | 15 | 42911151 | T | C | 490505 | 14 | 0.000292 |
| rs115363065 | 15 | 42977266 | G | C | 727314 | 15 | 0.000132 |
| rs371614413 | 15 | 43011821 | A | G | 846709 | 69 | 0.000958 |
| rs554997165 | 15 | 43012974 | C | T | 402071 | 3 | 0.000336 |
| rs568003335 | 15 | 42894191 | C | G | 968209 | 109 | 0.00247 |
| rs56801471 | 15 | 42896146 | A | G | 1199249 | 139 | 0.00423 |
| rs78994161 | 15 | 42975022 | C | A | 1186212 | 140 | 0.00654 |
| rs76787264 | 15 | 42983155 | G | A | 743325 | 13 | 0.000202 |
| rs563451455 | 15 | 42968307 | A | G | 773178 | 29 | 0.000471 |
| rs777539145 | 15 | 43010965 | A | G | 602498 | 6 | 0.000336 |
| NA | 15 | 42966323 | A | G | 394638 | 2 | 6.33E-05 |
| rs754784581 | 15 | 42982890 | G | GAGC | 90248 | 6 | 0.00167 |
| rs567894069 | 15 | 43011180 | T | A | 407817 | 5 | 0.000277 |
| NA | 15 | 42916776 | T | IS_ME_S\ | 238585 | 3 | 0.000773 |
| rs150519590 | 15 | 42907387 | T | C | 960029 | 111 | 0.00162 |
| rs72711776 | 15 | 42916041 | T | G | 1231955 | 159 | 0.0217 |
| rs572736554 | 15 | 42918591 | T | C | 826581 | 59 | 0.00125 |
| rs536329199 | 15 | 42991579 | T | C | 623415 | 4 | 3.77E-05 |
| rs552708360 | 15 | 42953454 | A | C | 541238 | 5 | 0.000716 |
| rs117842035 | 15 | 42978426 | T | A | 831015 | 26 | 0.000224 |
| rs573215252 | 15 | 42985549 | A | G | 36601 | 12 | 0.00082 |
| rs545599183 | 15 | 42886286 | A | C | 398051 | 3 | 5.15E-05 |
| rs139197926 | 15 | 42905546 | C | T | 488522 | 13 | 0.000292 |
| rs759558730 | 15 | 42978435 | A | C | 415962 | 3 | 0.000142 |
| rs80319409 | 15 | 42921919 | T | C | 1224605 | 140 | 0.00483 |
| rs146394639 | 15 | 42927581 | G | A | 1240908 | 153 | 0.00529 |
| rs28699522 | 15 | 42946935 | G | C | 1250292 | 166 | 0.0109 |
| rs144738245 | 15 | 42957109 | A | G | 826173 | 103 | 0.00224 |
| rs925195333 | 15 | 42901924 | C | T | 409125 | 6 | 4.03E-05 |
| rs193223759 | 15 | 42902603 | C | T | 959163 | 114 | 0.00168 |
| rs187036267 | 15 | 42947859 | C | T | 406794 | 5 | 0.00201 |
| rs114793100 | 15 | 42932355 | A | T | 1243078 | 153 | 0.00501 |
| rs535583406 | 15 | 42893630 | C | T | 698814 | 8 | 0.000114 |
| rs556707023 | 15 | 42956960 | GAATT | G | 614586 | 2 | 5.61E-05 |
| rs28680600 | 15 | 42979105 | A | G | 1247954 | 162 | 0.00934 |
| rs544930217 | 15 | 42991793 | A | G | 853444 | 57 | 0.00116 |
| rs73410937 | 15 | 43006957 | T | G | 1247956 | 162 | 0.00933 |
| rs114129956 | 15 | 43008736 | C | T | 631603 | 4 | 3.96E-05 |
| rs538352673 | 15 | 42980127 | A | G | 393138 | 2 | 0.000226 |
| rs73410931 | 15 | 43003939 | G | C | 1247955 | 162 | 0.00931 |
| rs555246628 | 15 | 42880764 | C | T | 608282 | 16 | 0.000221 |
| rs113194001 | 15 | 42981563 | G | A | 448900 | 7 | 0.000157 |
| rs143587129 | 15 | 42982912 | A | G | 614586 | 2 | 6.18E-05 |
| rs528267195 | 15 | 42990112 | C | T | 811007 | 48 | 0.0011 |
| rs75988797 | 15 | 43001332 | G | A | 1247956 | 162 | 0.00933 |
| rs12594837 | 15 | 42977290 | C | T | 1245089 | 157 | 0.00837 |
| rs570784384 | 15 | 42876851 | A | C | 402071 | 3 | 0.00107 |
| rs12592374 | 15 | 42991299 | A | G | 1248802 | 165 | 0.0103 |
| rs551210946 | 15 | 42991368 | G | C | 538186 | 13 | 0.000339 |
| rs73410927 | 15 | 43000977 | A | G | 1247955 | 162 | 0.00933 |
| rs112280084 | 15 | 42945127 | C | T | 618363 | 3 | 0.000139 |
| rs12594725 | 15 | 42999641 | C | G | 1247955 | 162 | 0.00932 |
| rs74477058 | 15 | 43001334 | C | G | 1243806 | 157 | 0.00931 |
| rs559872238 | 15 | 42900330 | G | C | 614586 | 2 | 3.42E-05 |
| rs887227858 | 15 | 42924342 | T | G | 460680 | 9 | 0.000102 |
| rs77096807 | 15 | 42982124 | A | G | 830376 | 25 | 0.000237 |
| rs60723187 | 15 | 42997347 | C | T | 1247956 | 162 | 0.00931 |
| rs563630307 | 15 | 42924465 | G | T | 490111 | 19 | 0.000276 |
| rs577736918 | 15 | 42995463 | T | C | 403470 | 4 | 8.67E-05 |
| rs12594696 | 15 | 42999540 | A | G | 1247954 | 162 | 0.00932 |
| rs1048283382 | 15 | 42872712 | T | C | 24127 | 6 | 0.000767 |
| rs778050361 | 15 | 42885190 | T | G | 417972 | 4 | 0.000195 |
| NA | 15 | 42903471 | G | A | 395738 | 3 | 0.000125 |
| rs751297558 | 15 | 42992954 | A | G | 458627 | 2 | 0.000519 |
| rs61732534 | 15 | 42985873 | C | T | 1202360 | 116 | 0.00282 |
| rs114991203 | 15 | 42990918 | T | C | 614586 | 2 | 0.000183 |
| rs73410923 | 15 | 42997586 | C | T | 1247956 | 162 | 0.00931 |
| rs368247671 | 15 | 42870824 | T | G | 782022 | 16 | 0.000203 |
| rs146007972 | 15 | 42886739 | A | G | 666479 | 10 | 0.000147 |
| rs545539486 | 15 | 42893768 | C | T | 742998 | 37 | 0.00122 |
| rs151101516 | 15 | 42913469 | C | T | 465326 | 18 | 0.000269 |
| rs113572612 | 15 | 42917539 | T | G | 539552 | 28 | 0.000213 |
| rs186343429 | 15 | 42891962 | G | A | 816050 | 11 | 0.000517 |
| rs557439861 | 15 | 42926132 | G | A | 537737 | 11 | 6.23E-05 |
| rs16957055 | 15 | 42980390 | C | T | 1247261 | 161 | 0.00935 |

|  |  |  |  |  |  |  |  |
| --- | --- | --- | --- | --- | --- | --- | --- |
| rs189150192 | 15 | 42944420 | C | T | 757144 | 49 | 0.00182 |
| NA | 15 | 42960106 | A | T | 42485 | 10 | 0.000847 |
| rs79165890 | 15 | 42977116 | T | C | 1161261 | 101 | 0.00261 |
| rs8036733 | 15 | 42998776 | G | T | 1249623 | 164 | 0.00942 |
| rs751529122 | 15 | 42923404 | A | G | 635059 | 7 | 0.000439 |
| rs191917476 | 15 | 42988641 | G | A | 467689 | 15 | 0.000247 |
| rs565747190 | 15 | 42910420 | T | C | 20031 | 4 | 0.0017 |
| rs8028699 | 15 | 42993469 | T | C | 1247955 | 162 | 0.00928 |
| rs114621323 | 15 | 42924567 | C | T | 722015 | 8 | 0.000432 |
| rs565034967 | 15 | 42963671 | C | T | 535228 | 8 | 0.000263 |
| rs16957063 | 15 | 42984288 | A | G | 1247264 | 161 | 0.00925 |
| rs148183291 | 15 | 42991986 | C | T | 614586 | 2 | 4.23E-05 |
| rs970534725 | 15 | 43012893 | T | G | 419245 | 7 | 0.000383 |
| rs201340789 | 15 | 42982237 | G | C | 1158002 | 100 | 0.00263 |
| NA | 15 | 42917845 | I | R | 1339 | 2 | 0.05 |
| rs113567591 | 15 | 42921313 | G | C | 1238230 | 153 | 0.00561 |
| rs62019359 | 15 | 42976706 | C | T | 1243433 | 168 | 0.0283 |
| rs3742992 | 15 | 42983943 | C | G | 1247264 | 161 | 0.00925 |
| rs573499753 | 15 | 42931951 | A | G | 25608 | 6 | 0.00105 |
| NA | 15 | 42991942 | T | C | 26547 | 7 | 0.00153 |
| rs8028365 | 15 | 42993930 | C | T | 1247956 | 162 | 0.00929 |
| rs6493055 | 15 | 42912620 | A | G | 1233279 | 150 | 0.00565 |
| rs576914901 | 15 | 42919272 | C | T | 462083 | 12 | 0.000171 |
| rs187191059 | 15 | 42941341 | C | T | 458627 | 2 | 0.000122 |
| rs758910906 | 15 | 42958938 | G | A | 674126 | 13 | 0.000478 |
| rs73408611 | 15 | 42945580 | C | T | 1250554 | 167 | 0.011 |
| rs149080202 | 15 | 42966135 | G | A | 234500 | 3 | 0.000354 |
| rs56839966 | 15 | 42945683 | G | T | 1250555 | 167 | 0.011 |
| rs16957061 | 15 | 42984182 | A | G | 1247262 | 161 | 0.00927 |
| rs939140502 | 15 | 42869541 | T | C | 30258 | 4 | 0.00273 |
| rs9652418 | 15 | 42892728 | T | C | 1031911 | 101 | 0.00199 |
| rs546585224 | 15 | 42979250 | C | T | 411270 | 2 | 0.000135 |
| NA | 15 | 42989408 | C | T | 26547 | 7 | 0.00153 |
| rs567499517 | 15 | 42879317 | G | A | 418410 | 3 | 9.80E-05 |
| rs554716041 | 15 | 42922198 | C | T | 346510 | 58 | 0.307 |
| rs181164895 | 15 | 42936910 | C | T | 641543 | 5 | 7.56E-05 |
| rs550161682 | 15 | 42947809 | T | A | 623293 | 30 | 0.000654 |
| rs138121440 | 15 | 42974713 | C | T | 1217173 | 141 | 0.00507 |
| rs58237318 | 15 | 42984858 | G | T | 1247263 | 161 | 0.00923 |
| rs143163723 | 15 | 43007526 | C | T | 1249051 | 168 | 0.0164 |
| rs764077238 | 15 | 42883475 | C | T | 645910 | 14 | 0.00069 |
| rs556503870 | 15 | 42903941 | A | T | 622446 | 27 | 0.000432 |
| rs1024511261 | 15 | 42942332 | A | T | 439347 | 9 | 0.00052 |
| rs528882270 | 15 | 42932699 | C | G | 561886 | 31 | 0.000437 |
| rs562465519 | 15 | 42894211 | G | A | 404353 | 3 | 0.000141 |
| rs578228153 | 15 | 42911780 | A | G | 706690 | 38 | 0.000843 |
| rs756488144 | 15 | 42911941 | G | A | 547845 | 25 | 0.000923 |
| rs776350968 | 15 | 43011503 | G | A | 20003 | 3 | 0.0016 |

|  |  |  |  |  |  |  |  |
| --- | --- | --- | --- | --- | --- | --- | --- |
| rs113915421 | 15 | 42991684 | C | T | 1249471 | 163 | 0.00929 |
| rs780449628 | 15 | 42944948 | A | C | 633123 | 10 | 0.000349 |
| rs545079807 | 15 | 42964221 | A | G | 403830 | 4 | 0.000196 |
| rs564485063 | 15 | 42868879 | T | A | 499120 | 11 | 0.000123 |
| rs149462255 | 15 | 42903026 | T | G | 236544 | 2 | 6.98E-05 |
| rs536362302 | 15 | 42977617 | T | C | 900272 | 58 | 0.00133 |
| rs12594858 | 15 | 42939916 | G | A | 1245283 | 156 | 0.00511 |
| rs560029646 | 15 | 43003736 | T | C | 636814 | 32 | 0.000262 |
| rs77080857 | 15 | 42928937 | T | C | 773768 | 16 | 0.000178 |
| rs184832341 | 15 | 42935608 | G | A | 632610 | 4 | 6.32E-05 |
| rs574188738 | 15 | 42900199 | C | G | 28623 | 10 | 0.000681 |
| rs192855523 | 15 | 42935587 | T | A | 632610 | 4 | 6.24E-05 |
| rs74626491 | 15 | 42970123 | C | A | 832835 | 28 | 0.000726 |
| NA | 15 | 43002106 | T | C | 29681 | 4 | 0.00153 |
| rs114356937 | 15 | 42974042 | T | C | 647987 | 5 | 0.000269 |
| rs3742995 | 15 | 42982819 | A | G | 1247263 | 161 | 0.00925 |
| rs78617381 | 15 | 42912689 | A | T | 413986 | 2 | 5.80E-05 |
| rs572370881 | 15 | 42902097 | C | G | 471928 | 3 | 9.22E-05 |
| rs140017728 | 15 | 42921751 | A | G | 582337 | 37 | 0.00214 |
| rs117217576 | 15 | 42954954 | T | C | 1180525 | 118 | 0.00199 |
| rs116507617 | 15 | 42915653 | A | G | 624350 | 4 | 0.000146 |
| rs572122353 | 15 | 42981744 | C | G | 520937 | 17 | 0.000195 |
| rs79816516 | 15 | 42940199 | A | G | 1140412 | 83 | 0.00116 |
| rs114982552 | 15 | 42913679 | G | A | 624350 | 4 | 0.000145 |
| rs114863961 | 15 | 42967544 | G | A | 402544 | 4 | 4.60E-05 |
| rs375469449 | 15 | 42871338 | G | C | 614586 | 2 | 3.25E-05 |
| rs147095009 | 15 | 42910982 | A | T | 436586 | 7 | 5.84E-05 |
| rs547725013 | 15 | 42953580 | G | A | 438277 | 9 | 0.000245 |
| rs61750914 | 15 | 42984825 | C | T | 728876 | 7 | 0.000126 |
| rs9972600 | 15 | 42910955 | T | C | 739198 | 17 | 0.000739 |
| rs184366727 | 15 | 42932036 | T | C | 632610 | 4 | 6.56E-05 |
| rs189285257 | 15 | 42916517 | A | G | 626434 | 5 | 0.000165 |
| rs370372581 | 15 | 42869497 | A | G | 856208 | 61 | 0.00193 |
| rs558945520 | 15 | 42898456 | TC | T | 419080 | 5 | 0.00169 |
| rs888907858 | 15 | 42900841 | A | C | 529486 | 4 | 7.08E-05 |
| rs752564285 | 15 | 42920688 | A | G | 516773 | 20 | 0.000219 |
| rs778806352 | 15 | 42938861 | G | C | 691610 | 17 | 0.00152 |
| rs114547623 | 15 | 42940503 | G | A | 728190 | 9 | 0.000442 |
| rs530613898 | 15 | 42982113 | C | T | 396915 | 3 | 0.000248 |
| rs538709234 | 15 | 43005421 | C | T | 462563 | 10 | 0.000226 |
| rs630620 | 15 | 42909082 | C | T | 916216 | 102 | 0.0106 |
| rs537457295 | 15 | 42942607 | C | T | 397905 | 4 | 0.000101 |
| rs542369195 | 15 | 42926249 | C | T | 393138 | 2 | 7.25E-05 |
| rs146143514 | 15 | 42895236 | G | A | 397969 | 2 | 0.000153 |
| rs184696803 | 15 | 42910131 | T | A | 392813 | 2 | 7.00E-05 |
| rs544930217 | 15 | 42991793 | A | T | 490218 | 17 | 0.000278 |
| rs559499569 | 15 | 42869508 | T | C | 419220 | 6 | 0.000113 |
| rs920583154 | 15 | 42906931 | G | A | 26316 | 7 | 0.00127 |
| rs543084498 | 15 | 42990117 | T | A | 495644 | 21 | 0.000606 |
| rs545694138 | 15 | 42998053 | C | T | 799327 | 38 | 0.000506 |
| rs188508115 | 15 | 43003063 | A | C | 687821 | 20 | 0.000844 |
| rs370295021 | 15 | 42867875 | C | G | 654013 | 4 | 0.000118 |
| rs115549874 | 15 | 42958108 | G | A | 725696 | 8 | 0.000449 |
| NA | 15 | 42956315 | T | C | 128382 | 12 | 0.000608 |
| rs12323990 | 15 | 42971515 | C | T | 1248462 | 163 | 0.00921 |
| rs112165067 | 15 | 42906924 | G | A | 1194334 | 139 | 0.00446 |
| rs1052463329 | 15 | 42930144 | A | G | 393383 | 2 | 4.96E-05 |
| rs773809476 | 15 | 42938556 | G | A | 844741 | 55 | 0.00109 |
| rs138147686 | 15 | 42911919 | G | A | 618363 | 3 | 5.50E-05 |
| rs532933640 | 15 | 42910158 | A | G | 407480 | 5 | 0.000422 |
| rs140413699 | 15 | 42924271 | C | G | 1151522 | 115 | 0.00411 |
| rs576086194 | 15 | 42910995 | A | AGCATG | 623519 | 3 | 0.000172 |
| rs181607501 | 15 | 42902975 | C | G | 783318 | 18 | 0.000274 |
| rs372595609 | 15 | 42918728 | C | T | 465662 | 25 | 0.000384 |
| rs554263361 | 15 | 42991801 | C | T | 678886 | 22 | 0.00111 |
| rs140540731 | 15 | 42943177 | C | T | 725695 | 8 | 0.00042 |
| rs540889150 | 15 | 42946083 | G | A | 401746 | 2 | 0.00065 |
| rs777092034 | 15 | 42982568 | C | T | 607336 | 15 | 0.00023 |
| rs116088885 | 15 | 42879631 | G | A | 631779 | 3 | 9.73E-05 |
| rs571444516 | 15 | 42879849 | C | G | 608935 | 40 | 0.00041 |
| rs367880689 | 15 | 42880211 | TTG | T | 615417 | 3 | 7.31E-05 |
| rs12911561 | 15 | 42868357 | C | T | 805795 | 32 | 0.000331 |
| rs146948676 | 15 | 42906046 | C | A | 668797 | 6 | 0.000124 |
| rs113515483 | 15 | 42938045 | T | G | 1139999 | 83 | 0.00119 |
| rs567867043 | 15 | 42964780 | C | T | 405477 | 5 | 0.000991 |
| rs573571482 | 15 | 42932937 | T | G | 403177 | 4 | 0.00372 |
| rs776432879 | 15 | 42905068 | G | C | 389982 | 2 | 0.000131 |
| rs535436686 | 15 | 42947292 | C | T | 410396 | 6 | 0.000588 |
| rs550441855 | 15 | 42956924 | A | G | 715413 | 52 | 0.000665 |
| rs535962355 | 15 | 42960676 | T | G | 834209 | 102 | 0.0026 |
| rs530071996 | 15 | 42991882 | T | C | 397969 | 2 | 0.000381 |
| rs543405550 | 15 | 42922960 | G | A | 407521 | 6 | 0.00011 |
| rs539505032 | 15 | 42937661 | G | T | 531303 | 27 | 0.000268 |
| rs550493975 | 15 | 42995846 | C | T | 684177 | 3 | 3.51E-05 |
| rs144945764 | 15 | 42999341 | T | TA | 1112941 | 123 | 0.00914 |
| rs563498229 | 15 | 42883265 | G | A | 390860 | 2 | 0.000168 |
| rs529629900 | 15 | 42887224 | G | A | 397969 | 2 | 0.00029 |
| rs761344257 | 15 | 42935771 | C | T | 505188 | 16 | 0.000428 |
| rs190287691 | 15 | 42989886 | C | T | 949665 | 112 | 0.00293 |
| rs151012902 | 15 | 42899817 | G | A | 1247618 | 168 | 0.0158 |
| rs561303050 | 15 | 42932406 | G | A | 524767 | 24 | 0.000275 |
| rs150050265 | 15 | 42917181 | T | A | 1198149 | 119 | 0.00383 |
| rs779133096 | 15 | 43013154 | C | T | 604876 | 12 | 0.000379 |
| rs185405546 | 15 | 42885977 | T | C | 666686 | 65 | 0.00259 |
| rs6493056 | 15 | 42913470 | G | A | 1237608 | 159 | 0.00687 |
| rs142508976 | 15 | 42952718 | G | A | 578561 | 36 | 0.00215 |
| rs539879268 | 15 | 42961387 | A | G | 404971 | 4 | 0.000225 |
| rs566075663 | 15 | 42963258 | A | C | 404971 | 4 | 0.000225 |
| rs561020009 | 15 | 42941197 | G | C | 407932 | 5 | 0.00169 |
| rs556362354 | 15 | 42918499 | C | T | 403031 | 4 | 0.000953 |
| rs748457096 | 15 | 42957287 | G | C | 145306 | 2 | 0.00321 |
| rs751191269 | 15 | 43011782 | G | C | 162492 | 4 | 0.000554 |
| rs140219968 | 15 | 42969456 | C | T | 667708 | 7 | 5.54E-05 |
| rs762517984 | 15 | 42945917 | C | A | 452751 | 8 | 0.000248 |
| rs558809042 | 15 | 42922202 | T | C | 725220 | 15 | 0.000623 |
| rs543405408 | 15 | 42886259 | G | A | 706153 | 90 | 0.00263 |
| rs866924731 | 15 | 42869204 | T | G | 29486 | 8 | 0.00337 |
| rs572679906 | 15 | 42877298 | G | T | 556765 | 17 | 0.000173 |
| rs146582932 | 15 | 42897448 | G | T | 1237920 | 162 | 0.0163 |
| rs777217494 | 15 | 42920742 | A | G | 656194 | 54 | 0.00105 |
| rs543953990 | 15 | 42965920 | G | A | 1237138 | 145 | 0.00569 |
| rs189176833 | 15 | 42906239 | G | A | 818922 | 24 | 0.000368 |
| rs139879027 | 15 | 42998863 | G | A | 614586 | 2 | 6.83E-05 |
| rs140427579 | 15 | 42903954 | A | T | 400053 | 3 | 5.50E-05 |
| rs577645292 | 15 | 43006346 | C | T | 948903 | 52 | 0.000469 |
| rs1056686032 | 15 | 42907056 | G | A | 29486 | 8 | 0.00151 |
| rs147467293 | 15 | 42916903 | C | T | 635230 | 6 | 0.000415 |
| rs1006228942 | 15 | 42932049 | A | G | 29486 | 8 | 0.00117 |
| rs574640812 | 15 | 42972264 | A | G | 30093 | 8 | 0.000532 |
| rs2136902 | 15 | 42993387 | A | G | 711484 | 18 | 0.000687 |
| rs181739281 | 15 | 42874183 | A | C | 622656 | 3 | 6.34E-05 |
| rs57568065 | 15 | 42945796 | CT | C | 1102068 | 117 | 0.0108 |
| rs8023743 | 15 | 42957542 | C | G | 1248782 | 165 | 0.00976 |
| rs539954662 | 15 | 42992689 | T | C | 712884 | 66 | 0.00101 |
| rs542480757 | 15 | 42884760 | C | T | 397969 | 2 | 4.65E-05 |
| rs538996660 | 15 | 42929482 | CT | C | 406042 | 3 | 0.000137 |
| rs115569250 | 15 | 42994390 | C | T | 704430 | 17 | 0.000161 |
| rs752837000 | 15 | 43002061 | T | C | 481532 | 9 | 0.000242 |
| rs954895078 | 15 | 42929690 | G | A | 551679 | 25 | 0.000388 |
| rs138677372 | 15 | 42931457 | C | T | 1215712 | 137 | 0.00456 |
| rs573239574 | 15 | 42936686 | G | A | 877257 | 50 | 0.00118 |
| rs578043321 | 15 | 42942579 | A | C | 429730 | 13 | 0.00293 |
| rs571684887 | 15 | 42956687 | G | A | 414343 | 8 | 0.00493 |
| rs78906041 | 15 | 42986552 | C | T | 710667 | 16 | 0.000164 |
| rs146651970 | 15 | 42965101 | C | T | 667708 | 7 | 6.66E-05 |
| rs571668597 | 15 | 43003370 | C | T | 425052 | 9 | 0.00124 |
| rs74889135 | 15 | 42890612 | T | G | 576364 | 18 | 0.000729 |
| rs9652442 | 15 | 42891654 | T | C | 668797 | 6 | 1.00E-04 |
| rs555887680 | 15 | 42954944 | T | C | 529368 | 10 | 0.000417 |
| rs189914718 | 15 | 42962288 | A | G | 397968 | 2 | 0.000655 |
| rs778466650 | 15 | 42903801 | A | ATTGTCC | 76317 | 2 | 0.0017 |
| rs60753136 | 15 | 42915693 | G | A | 1233145 | 150 | 0.00583 |
| rs112377990 | 15 | 42928463 | C | T | 1139999 | 83 | 0.00128 |
| rs115889964 | 15 | 42964282 | T | C | 722986 | 7 | 0.000259 |
| rs907021416 | 15 | 42917347 | C | T | 29486 | 8 | 0.00115 |
| rs534149147 | 15 | 42973156 | A | G | 492031 | 6 | 0.00036 |
| rs561463615 | 15 | 42881440 | G | T | 400053 | 3 | 5.25E-05 |
| rs538247556 | 15 | 42943741 | C | T | 905187 | 87 | 0.0019 |
| rs187159912 | 15 | 42984840 | G | A | 419860 | 8 | 0.00144 |
| rs541093919 | 15 | 42897691 | T | G | 441424 | 10 | 0.00025 |
| rs186941746 | 15 | 42933630 | T | C | 614437 | 32 | 0.000679 |
| rs1006058718 | 15 | 43012832 | G | A | 560180 | 18 | 0.000162 |
| rs936199949 | 15 | 42893312 | C | T | 451340 | 14 | 0.000557 |
| rs183399424 | 15 | 42941468 | C | T | 445183 | 10 | 7.30E-05 |
| rs563762546 | 15 | 42968248 | T | G | 878270 | 71 | 0.00362 |
| rs188759214 | 15 | 42932877 | G | A | 561374 | 27 | 0.00044 |
| rs6493058 | 15 | 42962576 | T | C | 1123798 | 80 | 0.000959 |
| rs114643981 | 15 | 42977460 | C | T | 630445 | 5 | 7.93E-05 |
| rs77371649 | 15 | 42894363 | G | A | 671230 | 62 | 0.00107 |
| rs910876412 | 15 | 42934648 | A | G | 21411 | 6 | 0.000934 |
| rs533049646 | 15 | 42924634 | A | G | 409073 | 5 | 0.000281 |
| rs183344754 | 15 | 42985129 | A | G | 44610 | 10 | 0.00528 |
| rs144580936 | 15 | 42870421 | G | A | 561783 | 37 | 0.000447 |
| rs549395191 | 15 | 42911679 | G | A | 523064 | 9 | 0.000667 |
| rs567595733 | 15 | 42936635 | G | A | 724051 | 23 | 0.000452 |
| rs549801198 | 15 | 42897008 | ATTTT | A | 477794 | 28 | 0.00057 |
| NA | 15 | 42945796 | R | D | 1339 | 2 | 0.0134 |
| rs75079841 | 15 | 42923403 | T | C | 641543 | 5 | 0.000138 |
| rs1037588084 | 15 | 42875428 | G | A | 148837 | 33 | 0.000413 |
| rs188799560 | 15 | 42985187 | G | A | 615417 | 3 | 2.60E-05 |
| rs150750050 | 15 | 43011145 | C | A | 1184639 | 121 | 0.00201 |
| rs577962095 | 15 | 42908997 | G | A | 407931 | 5 | 0.00171 |
| rs142645685 | 15 | 42899443 | C | G | 668797 | 6 | 0.000114 |
| rs541403134 | 15 | 42933835 | A | C | 614586 | 2 | 9.84E-05 |
| rs111805819 | 15 | 42947077 | A | G | 748474 | 20 | 0.000303 |
| rs180793543 | 15 | 42997839 | G | A | 615417 | 3 | 2.60E-05 |
| rs781370951 | 15 | 43001949 | C | T | 423400 | 4 | 0.000112 |
| rs529269398 | 15 | 42972145 | A | G | 560517 | 17 | 0.000231 |
| rs7178483 | 15 | 42956626 | A | G | 1248782 | 165 | 0.0097 |
| rs528814808 | 15 | 42961066 | G | GT | 392813 | 2 | 5.98E-05 |
| rs139887123 | 15 | 43006556 | G | A | 500291 | 19 | 0.000191 |
| rs565324621 | 15 | 43011460 | C | T | 393383 | 2 | 0.000278 |
| rs570397733 | 15 | 42925184 | CA | C | 1079573 | 96 | 0.00436 |
| rs556801807 | 15 | 42932670 | A | G | 845637 | 49 | 0.000533 |
| rs556120932 | 15 | 42932790 | C | T | 400053 | 3 | 0.000556 |
| rs16957043 | 15 | 42975053 | G | A | 1249747 | 167 | 0.012 |
| rs561702892 | 15 | 42992719 | A | G | 517711 | 11 | 0.000203 |
| rs547679651 | 15 | 42874132 | T | A | 1172726 | 112 | 0.0022 |
| rs145566791 | 15 | 42960083 | C | T | 667708 | 7 | 7.79E-05 |
| rs550730817 | 15 | 42964752 | G | A | 18896 | 4 | 0.00185 |
| rs148013951 | 15 | 42906336 | C | T | 1159427 | 102 | 0.0025 |
| rs986243537 | 15 | 42967543 | C | T | 44803 | 5 | 0.000714 |
| rs564244442 | 15 | 43008093 | C | CTGG | 390358 | 2 | 0.000386 |
| rs140074516 | 15 | 42903809 | T | C | 397969 | 2 | 0.000123 |
| rs544718627 | 15 | 42911991 | G | T | 25486 | 7 | 0.00116 |
| rs189897944 | 15 | 42893598 | C | T | 683823 | 12 | 0.000214 |
| NA | 15 | 43007276 | R | D | 1339 | 2 | 0.203 |
| rs181391693 | 15 | 42923116 | A | G | 471656 | 15 | 0.000332 |
| rs569918428 | 15 | 42982934 | G | A | 394638 | 2 | 4.31E-05 |
| rs530272524 | 15 | 42873680 | A | G | 279687 | 13 | 0.00114 |
| rs555231317 | 15 | 42952781 | A | C | 144056 | 21 | 0.00101 |
| rs551791019 | 15 | 42941985 | C | G | 702303 | 55 | 0.000918 |
| rs185998838 | 15 | 42971310 | T | G | 295141 | 2 | 6.78E-05 |
| rs544175764 | 15 | 42971579 | G | T | 295141 | 2 | 6.78E-05 |
| rs191913662 | 15 | 42998509 | T | C | 295141 | 2 | 6.78E-05 |
| rs572371078 | 15 | 43007480 | T | C | 414343 | 8 | 0.00501 |
| rs776613042 | 15 | 42893979 | C | T | 576040 | 36 | 0.000313 |
| rs111315571 | 15 | 42915042 | A | G | 614155 | 13 | 0.000127 |
| rs569336802 | 15 | 42964263 | T | A | 549870 | 5 | 0.000707 |
| rs549260871 | 15 | 42973275 | G | A | 843998 | 118 | 0.00308 |
| rs138641056 | 15 | 42907396 | G | C | 768125 | 23 | 0.000602 |
| rs112369865 | 15 | 42910650 | G | C | 614155 | 13 | 0.000127 |
| rs566949602 | 15 | 42935789 | A | G | 401746 | 3 | 5.73E-05 |
| rs780911612 | 15 | 42985057 | C | T | 522009 | 21 | 0.000358 |
| rs9944270 | 15 | 42962209 | A | T | 1248780 | 165 | 0.00969 |
| rs772774346 | 15 | 42980278 | A | C | 546183 | 14 | 0.000401 |
| rs75442140 | 15 | 42868643 | T | A | 236022 | 3 | 0.000269 |
| rs563800652 | 15 | 42995026 | T | C | 539586 | 22 | 0.000401 |
| rs142549145 | 15 | 42996712 | C | T | 623415 | 4 | 3.05E-05 |
| rs113206433 | 15 | 42893203 | A | G | 1167906 | 124 | 0.00289 |
| rs142416794 | 15 | 42917406 | G | A | 686261 | 4 | 0.000178 |
| rs549202838 | 15 | 42918182 | C | T | 601479 | 26 | 0.000603 |
| rs568962893 | 15 | 42978912 | C | T | 406989 | 4 | 0.000123 |
| rs149230374 | 15 | 42986336 | A | G | 413193 | 5 | 0.000238 |
| rs555525085 | 15 | 42868746 | C | G | 397969 | 2 | 0.000165 |
| NA | 15 | 42939984 | T | G | 112399 | 24 | 0.0014 |
| rs535068320 | 15 | 42949070 | G | T | 617306 | 3 | 0.000408 |
| rs117366041 | 15 | 42914185 | A | G | 571177 | 26 | 0.00105 |
| rs10163044 | 15 | 42903456 | G | C | 669628 | 7 | 0.000117 |
| rs114876104 | 15 | 42921192 | A | G | 643627 | 6 | 0.000133 |
| rs12439524 | 15 | 42886117 | C | T | 1147576 | 98 | 0.00178 |
| rs530835353 | 15 | 42919320 | T | C | 389982 | 2 | 0.000309 |
| rs753886624 | 15 | 42938027 | A | G | 426309 | 5 | 0.000249 |
| rs182392365 | 15 | 42989697 | G | A | 406985 | 4 | 0.000542 |
| NA | 15 | 43007040 | G | A | 6702 | 2 | 0.00373 |
| rs868042277 | 15 | 43008884 | A | G | 6702 | 2 | 0.00373 |
| rs747908929 | 15 | 43012424 | C | T | 123734 | 2 | 0.000626 |
| rs143568497 | 15 | 42920277 | C | T | 1203034 | 116 | 0.00271 |
| rs539402884 | 15 | 43001119 | T | A | 19702 | 4 | 0.00107 |
| NA | 15 | 42873866 | I | R | 1339 | 2 | 0.249 |

|  |  |  |  |  |  |  |  |
| --- | --- | --- | --- | --- | --- | --- | --- |
| rs754291307 | 15 | 42894385 | A | T | 443132 | 9 | 6.66E-05 |
| rs769686927 | 15 | 42926127 | C | T | 499441 | 15 | 0.000261 |
| rs577438946 | 15 | 42939829 | T | C | 404155 | 4 | 9.77E-05 |
| rs535464309 | 15 | 42946008 | T | TA | 390997 | 2 | 5.37E-05 |
| rs752599758 | 15 | 42964444 | A | T | 480340 | 8 | 0.000195 |
| rs553674710 | 15 | 43004059 | T | C | 622670 | 3 | 7.31E-05 |
| NA | 15 | 42908353 | T | C | 89846 | 16 | 0.00044 |
| rs188271981 | 15 | 42919241 | A | G | 915405 | 61 | 0.0023 |
| rs73408619 | 15 | 42949817 | G | C | 1248780 | 165 | 0.00968 |
| rs148437368 | 15 | 42938858 | T | G | 409066 | 5 | 0.000849 |
| rs184714358 | 15 | 42950446 | G | A | 615417 | 3 | 3.01E-05 |
| rs199897680 | 15 | 42975162 | T | C | 454335 | 21 | 0.000479 |
| rs185791709 | 15 | 42888179 | C | T | 295141 | 2 | 7.12E-05 |
| rs114191253 | 15 | 42924026 | T | A | 1192419 | 129 | 0.00355 |
| rs111336295 | 15 | 42984179 | C | G | 230866 | 4 | 0.000227 |
| rs181125969 | 15 | 42977568 | G | A | 409856 | 3 | 0.00112 |
| rs567751521 | 15 | 42874434 | A | C | 397969 | 2 | 0.000162 |
| rs539580881 | 15 | 42875976 | T | A | 397969 | 2 | 0.000161 |
| rs560061012 | 15 | 42936559 | C | T | 155730 | 20 | 0.000218 |
| rs552741284 | 15 | 43010039 | T | A | 405848 | 4 | 0.000439 |
| rs113625023 | 15 | 42886168 | T | C | 576040 | 36 | 0.000308 |
| rs781419142 | 15 | 42923300 | C | T | 566711 | 31 | 0.000664 |
| rs187519157 | 15 | 42904780 | T | C | 295141 | 2 | 6.27E-05 |
| rs185003255 | 15 | 42909008 | G | A | 614586 | 2 | 2.93E-05 |
| rs770886983 | 15 | 42894934 | A | G | 677353 | 39 | 0.000853 |
| rs145239633 | 15 | 42982455 | A | G | 398055 | 2 | 0.000155 |
| rs546043408 | 15 | 42881894 | T | G | 397905 | 4 | 0.000274 |
| rs199708541 | 15 | 42896064 | TC | T | 1115513 | 129 | 0.0196 |
| NA | 15 | 42896064 | R | D | 1339 | 2 | 0.0183 |
| rs973227743 | 15 | 42943691 | C | T | 462276 | 10 | 0.00038 |
| rs182501865 | 15 | 42948152 | G | A | 585873 | 22 | 0.000515 |
| rs188846832 | 15 | 42988440 | G | T | 393953 | 2 | 0.000156 |
| rs569639072 | 15 | 42895430 | C | G | 295141 | 2 | 6.27E-05 |
| rs564661828 | 15 | 42928648 | G | A | 743220 | 96 | 0.00168 |
| rs8043143 | 15 | 42948674 | A | G | 1248779 | 165 | 0.00973 |
| rs568463978 | 15 | 42932447 | C | T | 295141 | 2 | 6.61E-05 |
| NA | 15 | 42902586 | A | G | 409039 | 3 | 0.00021 |
| rs546152162 | 15 | 42929192 | A | T | 540530 | 29 | 0.000662 |
| rs550103536 | 15 | 42939466 | T | C | 815623 | 93 | 0.00179 |
| rs72711786 | 15 | 42994336 | C | T | 1184121 | 124 | 0.0039 |
| rs148235732 | 15 | 42916043 | G | A | 695194 | 4 | 0.000114 |
| rs566924050 | 15 | 42991956 | G | A | 622659 | 3 | 0.000182 |
| rs752932916 | 15 | 42895952 | T | C | 1214 | 2 | 0.0457 |
| rs571444533 | 15 | 42918107 | T | C | 825210 | 46 | 0.000962 |
| rs536218697 | 15 | 43003903 | TTTTA | T | 820882 | 36 | 0.00125 |
| rs553908774 | 15 | 43005841 | C | T | 394134 | 2 | 0.00112 |
| rs553920574 | 15 | 42924012 | G | C | 295141 | 2 | 6.61E-05 |
| rs564355348 | 15 | 42933848 | A | G | 618363 | 3 | 9.70E-05 |

|  |  |  |  |  |  |  |  |
| --- | --- | --- | --- | --- | --- | --- | --- |
| rs149407441 | 15 | 42937825 | G | A | 1251621 | 171 | 0.0263 |
| rs12438056 | 15 | 42959540 | G | A | 1201882 | 153 | 0.0952 |
| rs770269074 | 15 | 42898276 | A | G | 584219 | 4 | 0.000531 |
| rs117527257 | 15 | 42904467 | C | T | 730793 | 8 | 0.000168 |
| rs564833722 | 15 | 42919265 | G | A | 295141 | 2 | 6.61E-05 |
| rs539933616 | 15 | 42925191 | A | G | 1084606 | 102 | 0.00464 |
| rs542927420 | 15 | 42971913 | A | G | 733255 | 60 | 0.001 |
| rs150492102 | 15 | 42999937 | TTG | T | 660718 | 11 | 0.000126 |
| rs559067105 | 15 | 42880375 | T | G | 625580 | 3 | 6.63E-05 |
| rs192863692 | 15 | 42974806 | G | C | 651850 | 28 | 0.000703 |
| rs144291102 | 15 | 42912619 | C | T | 234483 | 2 | 0.000171 |
| rs547585065 | 15 | 42920939 | T | C | 847686 | 62 | 0.00205 |
| rs1029083837 | 15 | 43006299 | C | T | 399953 | 3 | 0.00012 |
| rs572787507 | 15 | 42891799 | T | C | 403831 | 4 | 0.0104 |
| rs530440643 | 15 | 42899141 | A | C | 409620 | 6 | 0.00172 |
| rs79533437 | 15 | 42948926 | A | G | 1141415 | 86 | 0.00115 |
| rs550166365 | 15 | 42869382 | C | G | 419707 | 3 | 0.000279 |
| rs187131404 | 15 | 42911059 | C | T | 1002668 | 77 | 0.00139 |
| rs7179256 | 15 | 42962258 | T | A | 1125909 | 80 | 0.00111 |
| rs76594962 | 15 | 42979456 | T | G | 632348 | 5 | 4.27E-05 |
| rs150536629 | 15 | 42895591 | G | A | 1162857 | 114 | 0.00231 |
| rs28736688 | 15 | 42945022 | G | A | 1250559 | 167 | 0.0097 |
| rs561421938 | 15 | 42937189 | T | A | 617306 | 3 | 0.000382 |
| rs770699990 | 15 | 42901032 | G | A | 816238 | 87 | 0.00128 |
| rs562709108 | 15 | 42909133 | A | G | 397969 | 2 | 0.00345 |
| rs569243432 | 15 | 42937646 | C | T | 1199423 | 116 | 0.00272 |
| rs538838662 | 15 | 42900649 | C | G | 403151 | 5 | 0.00133 |
| rs561000481 | 15 | 42919070 | C | CGTCT | 614586 | 2 | 8.05E-05 |
| rs534940697 | 15 | 42966921 | G | GTA | 394653 | 4 | 5.19E-05 |
| rs16956958 | 15 | 42944616 | G | A | 234483 | 2 | 0.000495 |
| NA | 15 | 42950486 | D | R | 1339 | 2 | 0.226 |
| rs553415415 | 15 | 42998513 | G | A | 404790 | 5 | 7.16E-05 |
| rs561497228 | 15 | 42979405 | A | T | 856035 | 64 | 0.00128 |
| rs948178615 | 15 | 42895090 | A | G | 29486 | 8 | 0.00137 |
| rs899259002 | 15 | 42900771 | A | G | 614155 | 13 | 0.000125 |
| rs76906844 | 15 | 42905887 | G | A | 1011436 | 92 | 0.00209 |
| rs146753941 | 15 | 43003871 | C | T | 619417 | 3 | 8.07E-05 |
| rs543709938 | 15 | 42941372 | C | T | 905445 | 73 | 0.00116 |
| rs569786942 | 15 | 42875501 | T | C | 704573 | 21 | 0.00028 |
| rs559682263 | 15 | 42878911 | T | TG | 814465 | 26 | 0.000523 |
| rs16973384 | 15 | 42904805 | T | A | 766285 | 23 | 0.000597 |
| rs537314110 | 15 | 42954801 | C | T | 539995 | 12 | 9.54E-05 |
| rs563742211 | 15 | 42992049 | C | T | 780574 | 87 | 0.0024 |
| rs551458553 | 15 | 42936471 | C | A | 765572 | 68 | 0.00133 |
| rs139264905 | 15 | 42908568 | G | A | 1246330 | 166 | 0.019 |
| rs540950607 | 15 | 42952786 | A | C | 244187 | 3 | 8.40E-05 |
| rs141349705 | 15 | 42970404 | C | T | 653536 | 5 | 0.000207 |
| rs547010092 | 15 | 42893819 | G | A | 852209 | 34 | 0.000343 |

|  |  |  |  |  |  |  |  |
| --- | --- | --- | --- | --- | --- | --- | --- |
| rs762907337 | 15 | 42895562 | C | T | 624874 | 6 | 7.68E-05 |
| rs192196303 | 15 | 42909620 | G | A | 1013330 | 85 | 0.00125 |
| NA | 15 | 42913750 | A | ACAGAG | 29486 | 8 | 0.00239 |
| rs189904276 | 15 | 42941288 | C | T | 27797 | 7 | 0.00115 |
| rs149354576 | 15 | 42931464 | C | A | 1238451 | 150 | 0.00759 |
| rs183125787 | 15 | 42934233 | T | A | 670279 | 7 | 0.000159 |
| rs184939172 | 15 | 42913788 | A | T | 655066 | 6 | 0.000162 |
| rs756077457 | 15 | 42920689 | A | G | 216499 | 30 | 0.000494 |
| rs57481952 | 15 | 42992989 | A | G | 758589 | 23 | 0.000767 |
| rs181793901 | 15 | 42891461 | C | T | 1032131 | 47 | 0.000481 |
| NA | 15 | 42894048 | T | A | 426762 | 4 | 0.000108 |
| rs1030428778 | 15 | 42929667 | C | A | 26547 | 7 | 0.000923 |
| rs192461105 | 15 | 42930927 | G | A | 260476 | 7 | 0.000885 |
| rs115134206 | 15 | 42943048 | A | G | 254123 | 5 | 0.000226 |
| rs78594152 | 15 | 42901445 | G | A | 716956 | 11 | 0.000429 |
| rs563034338 | 15 | 42924040 | T | A | 659787 | 33 | 0.000542 |
| rs114786341 | 15 | 42953446 | G | A | 244187 | 3 | 8.40E-05 |
| rs141905011 | 15 | 42907400 | G | A | 234483 | 2 | 0.000143 |
| rs576049319 | 15 | 42894935 | T | C | 911072 | 49 | 0.00052 |
| rs558670205 | 15 | 42948234 | C | T | 901406 | 85 | 0.00189 |
| rs555144614 | 15 | 42980924 | A | G | 399291 | 3 | 0.000773 |
| rs145410785 | 15 | 42886573 | G | GT | 623519 | 3 | 5.37E-05 |
| rs752536103 | 15 | 42954307 | C | T | 505466 | 14 | 0.000355 |
| rs60111534 | 15 | 42868169 | CCG | C | 1087139 | 113 | 0.00745 |
| rs577581215 | 15 | 42885609 | T | C | 539458 | 23 | 0.000396 |
| rs550361351 | 15 | 42942246 | C | G | 582789 | 20 | 0.000513 |
| rs555851843 | 15 | 42908243 | T | C | 885478 | 85 | 0.00266 |
| rs542299362 | 15 | 42953963 | A | C | 31574 | 7 | 0.00314 |
| rs543049266 | 15 | 42919059 | T | G | 399291 | 3 | 5.76E-05 |
| rs183110458 | 15 | 42893272 | C | G | 1036847 | 88 | 0.00222 |
| rs192367918 | 15 | 42943122 | A | G | 295141 | 2 | 8.64E-05 |
| rs924868347 | 15 | 42871846 | A | G | 439262 | 7 | 4.10E-05 |
| rs564939782 | 15 | 42905529 | A | G | 401746 | 3 | 0.000126 |
| rs116067069 | 15 | 42888213 | A | G | 684070 | 7 | 0.000237 |
| rs142940493 | 15 | 42931797 | C | T | 614586 | 2 | 2.85E-05 |
| rs9652417 | 15 | 42892674 | T | G | 756803 | 20 | 0.000539 |
| NA | 15 | 42907211 | I | R | 1339 | 2 | 0.251 |
| rs114642146 | 15 | 42922785 | T | G | 643627 | 6 | 0.00013 |
| rs112111204 | 15 | 42980085 | C | T | 733594 | 17 | 0.000198 |
| rs750361625 | 15 | 42935674 | G | C | 257646 | 19 | 0.00108 |
| rs544613911 | 15 | 42907635 | G | A | 287737 | 15 | 0.00117 |
| rs183904150 | 15 | 43011179 | G | A | 1242012 | 161 | 0.00776 |
| rs191328807 | 15 | 42897371 | A | G | 402070 | 3 | 0.000713 |
| rs775556799 | 15 | 42902599 | T | G | 399762 | 3 | 9.51E-05 |
| rs763235317 | 15 | 42966791 | T | C | 637697 | 32 | 0.000501 |
| rs763967566 | 15 | 42968068 | A | T | 499495 | 16 | 0.00019 |
| rs529322354 | 15 | 42968801 | A | T | 881824 | 80 | 0.00145 |
| rs567833296 | 15 | 42910039 | C | T | 625364 | 5 | 0.000248 |
| rs189549622 | 15 | 42951754 | C | T | 928287 | 60 | 0.000763 |
| rs545235526 | 15 | 42985103 | G | A | 626436 | 4 | 4.31E-05 |
| rs558007975 | 15 | 42930036 | C | A | 1201599 | 117 | 0.00274 |
| rs73406601 | 15 | 42938683 | T | C | 986174 | 85 | 0.00112 |
| rs79710890 | 15 | 42953617 | G | T | 1147122 | 89 | 0.0012 |
| rs185150927 | 15 | 43006447 | T | C | 628350 | 4 | 8.51E-05 |
| rs149502125 | 15 | 42886418 | A | G | 460154 | 11 | 0.000158 |
| rs566288410 | 15 | 42892849 | G | A | 397969 | 2 | 0.00116 |
| rs560294077 | 15 | 42933084 | CAT | C | 440025 | 12 | 0.000191 |
| rs77336982 | 15 | 42999952 | C | T | 1143217 | 85 | 0.00116 |
| rs745796005 | 15 | 43001017 | A | G | 455177 | 12 | 8.02E-05 |
| NA | 15 | 42870304 | D | R | 1339 | 2 | 0.127 |
| rs533124511 | 15 | 42883247 | G | C | 392813 | 2 | 0.000134 |
| rs145036897 | 15 | 42896996 | C | T | 1210046 | 139 | 0.00805 |
| rs755438104 | 15 | 42936547 | C | T | 503503 | 6 | 0.000117 |
| NA | 15 | 42930850 | A | T | 27797 | 7 | 0.00112 |
| rs190421447 | 15 | 42937413 | C | T | 295141 | 2 | 7.96E-05 |
| rs188615523 | 15 | 42960991 | G | T | 906743 | 58 | 0.00217 |
| rs186959564 | 15 | 42870931 | T | C | 1103419 | 153 | 0.00877 |
| rs140413699 | 15 | 42924271 | C | T | 441627 | 8 | 0.000118 |
| rs559254882 | 15 | 42938251 | C | A | 295141 | 2 | 7.96E-05 |
| rs756126576 | 15 | 42869616 | G | A | 590843 | 20 | 0.00026 |
| rs541758310 | 15 | 42918606 | T | C | 781440 | 28 | 0.000643 |
| rs182260499 | 15 | 42956967 | T | C | 717206 | 30 | 0.00071 |
| rs191115724 | 15 | 42884735 | C | T | 632175 | 5 | 0.000206 |
| rs141336135 | 15 | 42891358 | G | C | 1142251 | 95 | 0.000858 |
| rs548431781 | 15 | 42902807 | C | T | 435269 | 5 | 4.10E-05 |
| rs150908758 | 15 | 43006166 | G | A | 684177 | 3 | 3.00E-05 |
| rs148022476 | 15 | 42882970 | G | A | 623519 | 3 | 4.73E-05 |
| rs116719583 | 15 | 42937465 | T | G | 665333 | 8 | 9.99E-05 |
| rs547575932 | 15 | 42896069 | G | A | 1116812 | 131 | 0.0196 |
| rs78373841 | 15 | 42989002 | A | G | 1159163 | 99 | 0.00122 |
| rs188599187 | 15 | 42991957 | T | G | 417811 | 7 | 0.000297 |
| NA | 15 | 42926750 | C | T | 29486 | 8 | 0.00122 |
| rs59881302 | 15 | 42930419 | G | A | 987679 | 85 | 0.00117 |
| rs185487313 | 15 | 42905168 | C | A | 804973 | 20 | 0.000347 |
| rs77757620 | 15 | 42953597 | C | T | 1149384 | 90 | 0.00113 |
| rs571738986 | 15 | 42933157 | AC | A | 295141 | 2 | 8.30E-05 |
| rs150051531 | 15 | 42952133 | T | C | 666478 | 11 | 0.000412 |
| rs138770997 | 15 | 42879489 | C | G | 242743 | 2 | 0.000247 |
| rs188602770 | 15 | 42941109 | T | A | 1212447 | 122 | 0.00285 |
| rs535004528 | 15 | 42977084 | C | T | 405847 | 4 | 0.00173 |
| NA | 15 | 42995565 | R | D | 1339 | 2 | 0.217 |
| rs527386875 | 15 | 42923528 | C | T | 693334 | 9 | 0.000177 |
| rs143440000 | 15 | 42879189 | G | A | 242743 | 2 | 0.000249 |
| rs534058064 | 15 | 42881762 | G | T | 625603 | 4 | 0.00047 |
| rs72711771 | 15 | 42888777 | A | T | 1250233 | 171 | 0.039 |
| rs79130229 | 15 | 42896228 | G | C | 670560 | 10 | 0.000173 |
| NA | 15 | 42909088 | R | D | 1339 | 2 | 0.0205 |
| rs78898799 | 15 | 42888226 | G | A | 752456 | 19 | 0.000494 |
| rs538040094 | 15 | 42921600 | A | G | 392813 | 2 | 4.58E-05 |
| rs188650242 | 15 | 42922912 | A | G | 295141 | 2 | 8.30E-05 |
| rs557130886 | 15 | 42987901 | G | T | 751916 | 44 | 0.00131 |
| rs78859522 | 15 | 42952782 | C | A | 640108 | 11 | 0.00563 |
| rs114646969 | 15 | 43000363 | T | A | 628350 | 4 | 7.96E-05 |
| rs140192774 | 15 | 42881857 | C | T | 710499 | 8 | 0.000113 |
| rs142117976 | 15 | 42895399 | T | A | 1226954 | 157 | 0.0176 |
| rs115223182 | 15 | 42935477 | C | T | 727615 | 19 | 0.000228 |
| rs576866690 | 15 | 42945758 | G | A | 402071 | 3 | 0.00063 |
| rs540545255 | 15 | 42935552 | G | A | 654348 | 35 | 0.000316 |
| rs149348057 | 15 | 42998848 | A | G | 1134708 | 80 | 0.00102 |
| rs553559102 | 15 | 42899776 | A | G | 618363 | 3 | 0.000131 |
| rs538866117 | 15 | 42876265 | C | T | 404355 | 5 | 0.000122 |
| rs537715420 | 15 | 42911798 | G | A | 497691 | 7 | 0.000207 |
| rs539460492 | 15 | 42936407 | C | G | 614586 | 2 | 2.68E-05 |
| rs180867650 | 15 | 43008642 | C | A | 1218914 | 143 | 0.00943 |
| rs117352983 | 15 | 42956718 | T | C | 915231 | 73 | 0.00104 |
| rs73406557 | 15 | 42868515 | C | G | 1123229 | 86 | 0.00128 |
| rs186882062 | 15 | 42936652 | C | G | 618363 | 2 | 6.55E-05 |
| rs544284211 | 15 | 42957625 | T | A | 416872 | 4 | 0.000202 |
| rs941905115 | 15 | 42899346 | A | G | 29486 | 8 | 0.000848 |
| rs182390722 | 15 | 42930168 | A | G | 1021742 | 88 | 0.0015 |
| rs116577253 | 15 | 42873997 | A | C | 242743 | 2 | 0.000249 |
| rs80347429 | 15 | 42875246 | A | G | 741320 | 19 | 0.000409 |
| rs541589109 | 15 | 42900851 | A | G | 1164322 | 116 | 0.00232 |
| rs140656467 | 15 | 43012692 | G | A | 715982 | 79 | 0.00255 |
| rs149365302 | 15 | 42956842 | G | C | 392813 | 2 | 0.000524 |
| rs145400823 | 15 | 42986274 | C | T | 407170 | 5 | 0.00027 |
| rs143499451 | 15 | 42953711 | T | C | 618363 | 3 | 8.33E-05 |
| rs546146429 | 15 | 42961977 | G | A | 859163 | 57 | 0.00094 |
| rs761997441 | 15 | 42900978 | T | C | 766637 | 47 | 0.00071 |
| rs7167971 | 15 | 42912455 | A | G | 614586 | 2 | 8.14E-05 |
| rs538541542 | 15 | 42990575 | A | G | 663683 | 9 | 0.000795 |
| NA | 15 | 42910898 | G | A | 37017 | 4 | 0.000513 |
| rs79749780 | 15 | 42963806 | T | A | 234483 | 2 | 0.000463 |
| rs149616782 | 15 | 42898801 | G | A | 238260 | 3 | 0.000313 |
| rs529923027 | 15 | 42957022 | A | C | 873928 | 85 | 0.0157 |
| NA | 15 | 42875223 | T | C | 555710 | 25 | 0.000927 |
| rs141935808 | 15 | 42905231 | AG | A | 635348 | 6 | 0.000464 |
| rs560437326 | 15 | 42872620 | T | G | 1001108 | 40 | 0.000432 |
| NA | 15 | 42885209 | A | G | 31269 | 2 | 0.000816 |
| rs113245184 | 15 | 42933898 | G | A | 669357 | 36 | 0.000337 |
| rs188247542 | 15 | 42999995 | G | T | 822924 | 112 | 0.0026 |
| rs183754250 | 15 | 43005370 | G | C | 1154050 | 98 | 0.00244 |
| rs539428789 | 15 | 42946624 | G | T | 234483 | 2 | 9.60E-05 |
| rs1036190902 | 15 | 42959670 | C | T | 394638 | 2 | 4.18E-05 |

|  |  |  |  |  |  |  |  |
| --- | --- | --- | --- | --- | --- | --- | --- |
| rs572610576 | 15 | 42969282 | C | G | 234483 | 2 | 0.000124 |
| rs542649755 | 15 | 42995728 | AT | A | 744562 | 49 | 0.0541 |
| rs143824062 | 15 | 43005682 | G | A | 405848 | 3 | 0.000139 |
| rs548248907 | 15 | 42974169 | C | A | 214299 | 26 | 0.0024 |
| rs533842483 | 15 | 42979463 | G | A | 389982 | 2 | 0.00643 |
| rs9806648 | 15 | 43007688 | G | A | 699559 | 16 | 0.000168 |
| rs200569969 | 15 | 42906593 | TAGG | T | 630273 | 4 | 0.000118 |
| rs57133852 | 15 | 42923960 | GA | G | 1020647 | 82 | 0.00524 |
| rs116267701 | 15 | 42876795 | C | T | 240875 | 5 | 0.000241 |
| rs530364055 | 15 | 42880494 | A | G | 397969 | 2 | 0.000217 |
| rs75939632 | 15 | 42907791 | A | T | 738321 | 9 | 0.000514 |
| rs545866797 | 15 | 42909991 | G | A | 397968 | 2 | 0.00362 |
| rs202017657 | 15 | 42981101 | C | G | 1181560 | 107 | 0.00241 |
| rs10163179 | 15 | 42879555 | G | C | 741321 | 19 | 0.000426 |
| rs74670147 | 15 | 42894846 | G | A | 759767 | 21 | 0.000556 |
| rs113910756 | 15 | 42965461 | G | A | 614586 | 2 | 3.50E-05 |
| rs568687584 | 15 | 42986564 | G | A | 82301 | 3 | 0.000899 |
| rs143689566 | 15 | 42897434 | A | G | 1161875 | 113 | 0.00229 |
| rs138246348 | 15 | 42898800 | C | T | 620447 | 4 | 0.000186 |
| rs573060310 | 15 | 42943327 | C | T | 391120 | 2 | 0.000485 |
| NA | 15 | 42993022 | T | G | 20911 | 5 | 0.00115 |
| rs144334043 | 15 | 42873927 | G | GT | 615417 | 3 | 5.69E-05 |
| rs370739059 | 15 | 42991702 | C | G | 628350 | 4 | 6.68E-05 |
| rs543662054 | 15 | 42890537 | A | G | 397969 | 2 | 0.000217 |
| NA | 15 | 42956987 | R | D | 1339 | 2 | 0.205 |
| rs942809430 | 15 | 42996893 | G | C | 564026 | 18 | 0.000477 |
| rs139919565 | 15 | 42994529 | G | A | 628350 | 4 | 7.00E-05 |
| rs76712048 | 15 | 42925075 | G | A | 1247492 | 168 | 0.0213 |
| rs17767439 | 15 | 42927182 | A | G | 1244916 | 160 | 0.0114 |
| rs766671995 | 15 | 42966902 | C | T | 431690 | 4 | 0.00013 |
| rs572711131 | 15 | 42895174 | G | T | 784023 | 57 | 0.00063 |
| rs112267992 | 15 | 42895540 | G | C | 571540 | 10 | 0.000117 |
| NA | 15 | 42929277 | I | R | 1339 | 2 | 0.0164 |
| rs576093418 | 15 | 42931389 | G | A | 693334 | 9 | 0.000181 |
| rs8039881 | 15 | 42935849 | A | G | 1240999 | 162 | 0.00721 |
| rs1056030401 | 15 | 42917567 | C | T | 586642 | 21 | 0.000214 |
| rs755722649 | 15 | 42874521 | G | C | 592145 | 21 | 0.000254 |
| rs189531502 | 15 | 42951301 | A | C | 1174088 | 117 | 0.00445 |
| NA | 15 | 42957668 | AAC | A | 29486 | 8 | 0.00125 |
| rs938046 | 15 | 43010364 | C | T | 1034862 | 99 | 0.00246 |
| rs143222805 | 15 | 42889798 | G | C | 1248371 | 171 | 0.0699 |
| rs570478377 | 15 | 42936300 | A | G | 614586 | 2 | 2.77E-05 |
| rs189020874 | 15 | 42954642 | C | T | 426597 | 7 | 0.000206 |
| rs549319649 | 15 | 42967398 | C | G | 991744 | 78 | 0.00186 |
| rs115595492 | 15 | 42888436 | C | G | 664960 | 5 | 9.60E-05 |
| rs750624051 | 15 | 42954133 | C | A | 550294 | 8 | 0.000107 |
| rs572690483 | 15 | 42880406 | A | G | 644476 | 59 | 0.00115 |
| rs1005109579 | 15 | 42941159 | T | G | 426641 | 5 | 0.000402 |

|  |  |  |  |  |  |  |  |
| --- | --- | --- | --- | --- | --- | --- | --- |
| rs188648617 | 15 | 42969278 | G | A | 345155 | 11 | 0.000426 |
| rs114831533 | 15 | 42885803 | C | A | 240875 | 5 | 0.000243 |
| rs547550289 | 15 | 42982862 | A | G | 393138 | 2 | 0.000644 |
| rs76473483 | 15 | 42975557 | G | C | 1141856 | 85 | 0.00116 |
| rs187974547 | 15 | 42979226 | C | G | 1015474 | 62 | 0.000818 |
| rs184219872 | 15 | 42988314 | G | A | 626239 | 4 | 0.000402 |
| rs143860655 | 15 | 43001506 | C | T | 972237 | 76 | 0.00214 |
| NA | 15 | 42955996 | A | T | 40976 | 6 | 0.00116 |
| rs778100847 | 15 | 42986370 | C | T | 476027 | 14 | 0.000297 |
| NA | 15 | 42999341 | R | I | 1339 | 2 | 0.0127 |
| rs185424711 | 15 | 43009737 | C | T | 353425 | 8 | 0.000417 |
| rs147842892 | 15 | 42896551 | C | T | 710699 | 28 | 0.00171 |
| rs79365604 | 15 | 42976440 | G | A | 1228685 | 154 | 0.00735 |
| rs61746531 | 15 | 42985398 | C | T | 734542 | 18 | 0.000133 |
| rs550580422 | 15 | 42885935 | C | T | 401746 | 3 | 0.000107 |
| rs148150965 | 15 | 42924487 | CT | C | 947886 | 43 | 0.00126 |
| NA | 15 | 42987552 | G | A | 28936 | 3 | 0.00145 |
| rs77736048 | 15 | 42909596 | T | G | 1061799 | 110 | 0.00249 |
| rs560512058 | 15 | 42992562 | C | G | 638126 | 5 | 6.35E-05 |
| rs181359744 | 15 | 42941161 | C | G | 1019711 | 90 | 0.00152 |
| rs572428417 | 15 | 42948976 | A | C | 403068 | 4 | 0.000497 |
| rs61192504 | 15 | 42989060 | C | A | 1159231 | 98 | 0.00122 |
| rs535352166 | 15 | 42990423 | T | C | 766413 | 32 | 0.000345 |
| rs184122994 | 15 | 42990730 | T | G | 626239 | 4 | 0.000401 |
| rs10163179 | 15 | 42879555 | G | A | 825451 | 66 | 0.00224 |
| rs535169145 | 15 | 43002549 | C | T | 403909 | 5 | 0.000313 |
| NA | 15 | 42963392 | G | A | 22669 | 5 | 0.000728 |
| rs188552931 | 15 | 42901005 | G | A | 234483 | 2 | 0.000211 |
| rs185723917 | 15 | 42995861 | G | A | 616436 | 3 | 3.81E-05 |
| rs182223038 | 15 | 42941333 | T | C | 402071 | 3 | 9.70E-05 |
| rs578201778 | 15 | 42970456 | T | TTG | 744703 | 18 | 0.000959 |
| rs536364283 | 15 | 42977319 | T | C | 699969 | 19 | 0.000341 |
| rs185468708 | 15 | 42899154 | G | A | 1097285 | 66 | 0.000824 |
| rs550555857 | 15 | 42987196 | T | TA | 981992 | 65 | 0.00201 |
| rs73406561 | 15 | 42873793 | C | A | 1109687 | 78 | 0.000926 |
| NA | 15 | 42981031 | C | G | 426156 | 3 | 0.000238 |
| NA | 15 | 42874103 | CTT | C | 29486 | 8 | 0.000746 |
| rs115660182 | 15 | 42889346 | T | G | 664960 | 5 | 9.30E-05 |
| NA | 15 | 42915711 | G | T | 19722 | 5 | 0.00122 |
| rs973355371 | 15 | 42890932 | C | G | 39526 | 9 | 0.00161 |
| NA | 15 | 42897419 | G | A | 29486 | 8 | 0.000746 |
| rs16957011 | 15 | 42961901 | T | C | 694715 | 12 | 0.000158 |
| rs145525359 | 15 | 42911771 | G | A | 1220343 | 129 | 0.00518 |
| rs142591139 | 15 | 42988901 | G | A | 1236752 | 160 | 0.0159 |
| rs28461422 | 15 | 42883883 | G | A | 272147 | 4 | 0.000215 |
| rs540506511 | 15 | 42900849 | G | A | 489710 | 13 | 0.000233 |
| rs778682289 | 15 | 42927059 | C | T | 535283 | 10 | 8.87E-05 |
| rs767425815 | 15 | 42935673 | C | A | 249592 | 18 | 0.00111 |

|  |  |  |  |  |  |  |  |
| --- | --- | --- | --- | --- | --- | --- | --- |
| rs144424767 | 15 | 42884767 | C | T | 663676 | 6 | 0.000101 |
| rs192212691 | 15 | 42929560 | G | A | 455098 | 19 | 0.000761 |
| rs549071019 | 15 | 42977706 | A | G | 631343 | 7 | 0.00114 |
| NA | 15 | 42879024 | A | C | 394638 | 2 | 5.07E-05 |
| rs565841251 | 15 | 42910911 | C | T | 485692 | 30 | 0.000264 |
| rs562642851 | 15 | 42944671 | C | A | 404954 | 4 | 0.00169 |
| rs759661465 | 15 | 42985854 | G | T | 389982 | 2 | 0.00155 |
| rs753974961 | 15 | 42992817 | T | C | 679010 | 22 | 0.000334 |
| rs559676742 | 15 | 42994563 | G | A | 773768 | 15 | 0.000167 |
| NA | 15 | 43000191 | R | I | 1339 | 2 | 0.213 |
| rs150824141 | 15 | 42885513 | AT | A | 928066 | 45 | 0.0289 |
| rs532261184 | 15 | 42906837 | T | A | 405847 | 4 | 0.000448 |
| rs146992306 | 15 | 42973533 | T | C | 409066 | 5 | 0.00122 |
| rs760673116 | 15 | 42973701 | G | T | 29486 | 8 | 0.00125 |
| rs570361618 | 15 | 42909402 | C | T | 397969 | 2 | 0.000368 |
| rs374126429 | 15 | 42918675 | A | C | 887477 | 65 | 0.000873 |
| rs148223590 | 15 | 42874204 | G | A | 614586 | 2 | 6.83E-05 |
| rs186653054 | 15 | 42918662 | C | T | 473746 | 5 | 0.00192 |
| rs112210775 | 15 | 42939130 | G | A | 230866 | 4 | 0.000271 |
| rs188499850 | 15 | 42974646 | G | A | 574507 | 26 | 0.000359 |
| rs183600272 | 15 | 42998717 | C | T | 649064 | 6 | 7.47E-05 |
| rs201049061 | 15 | 42907744 | CAT | C | 657903 | 10 | 0.000917 |
| rs150454654 | 15 | 42898683 | G | A | 242743 | 2 | 0.000185 |
| rs141161691 | 15 | 42992007 | C | T | 612657 | 27 | 0.000157 |
| rs116670090 | 15 | 43005168 | G | A | 628350 | 4 | 9.95E-05 |
| rs185447522 | 15 | 42957031 | G | A | 625580 | 3 | 2.56E-05 |
| rs541989351 | 15 | 42993931 | G | A | 639351 | 56 | 0.000816 |
| rs1010566444 | 15 | 42933566 | T | G | 608549 | 12 | 0.000126 |
| rs12324838 | 15 | 42940790 | C | T | 1243439 | 157 | 0.0064 |
| rs555690816 | 15 | 42938377 | T | C | 931615 | 70 | 0.00122 |
| rs527471622 | 15 | 42938913 | T | C | 668170 | 39 | 0.000378 |
| rs565647014 | 15 | 42947538 | A | G | 402071 | 3 | 5.47E-05 |
| rs543357875 | 15 | 42975078 | C | T | 773517 | 78 | 0.00154 |
| NA | 15 | 42870823 | R | D | 1339 | 2 | 0.162 |
| rs751216502 | 15 | 42876578 | C | T | 157873 | 17 | 0.000469 |
| rs560294808 | 15 | 42943755 | T | C | 608549 | 12 | 0.000126 |
| rs545441576 | 15 | 42930749 | G | A | 397969 | 2 | 0.000376 |
| rs552442784 | 15 | 42943859 | C | T | 397968 | 2 | 0.000966 |
| rs150729042 | 15 | 42950747 | C | G | 697435 | 13 | 0.000158 |
| rs546974729 | 15 | 42963858 | G | A | 903305 | 98 | 0.00276 |
| rs111656044 | 15 | 42888606 | A | G | 611482 | 12 | 0.00012 |
| rs538472001 | 15 | 42918459 | CCTCTTCCTC | T | 928021 | 88 | 0.00327 |
| rs139534250 | 15 | 42928165 | C | A | 1238928 | 145 | 0.0052 |
| rs183072726 | 15 | 42938596 | A | G | 235030 | 3 | 0.00013 |
| rs112297283 | 15 | 42886367 | G | T | 837032 | 13 | 0.000137 |
| rs575749928 | 15 | 42893616 | C | T | 611482 | 12 | 0.00012 |
| rs559047960 | 15 | 43005859 | GTCTTT | G | 618363 | 3 | 0.000228 |
| rs554039449 | 15 | 42880312 | G | T | 402618 | 4 | 0.00132 |
| rs148320957 | 15 | 42894430 | A | G | 1018041 | 88 | 0.00121 |
| rs535848692 | 15 | 42936777 | A | C | 474144 | 10 | 0.000246 |
| rs531519065 | 15 | 42909557 | A | G | 909246 | 98 | 0.00251 |
| rs10162744 | 15 | 42879383 | A | C | 661183 | 5 | 8.60E-05 |
| rs541657557 | 15 | 42888732 | A | G | 741919 | 28 | 0.000338 |
| rs142987541 | 15 | 43006485 | C | T | 832996 | 45 | 0.000398 |
| rs114743946 | 15 | 42896913 | T | C | 771688 | 24 | 0.000566 |
| rs576444632 | 15 | 42923161 | C | T | 503590 | 8 | 0.000409 |
| rs541248519 | 15 | 42917177 | G | A | 594837 | 20 | 0.000444 |
| rs190820260 | 15 | 42972022 | A | T | 1020966 | 91 | 0.00158 |
| NA | 15 | 42983932 | G | A | 27797 | 7 | 0.000791 |
| rs549644430 | 15 | 43005010 | G | A | 1014598 | 85 | 0.00156 |
| rs16973396 | 15 | 42915785 | A | T | 768194 | 25 | 0.000699 |
| rs141012846 | 15 | 42931366 | A | G | 417047 | 4 | 0.000126 |
| rs562713483 | 15 | 42946939 | A | G | 397969 | 2 | 7.92E-05 |
| rs538275463 | 15 | 42921518 | A | G | 397969 | 2 | 0.000374 |
| rs202110998 | 15 | 42984712 | G | A | 739366 | 28 | 0.00221 |
| rs574436638 | 15 | 42996347 | G | A | 534516 | 22 | 0.000193 |
| rs562269594 | 15 | 42933234 | T | A | 568871 | 15 | 0.000677 |
| rs767836352 | 15 | 42894878 | A | G | 684790 | 20 | 0.000827 |
| rs202074007 | 15 | 42967130 | G | T | 1215290 | 118 | 0.00214 |
| NA | 15 | 42982744 | G | C | 27797 | 7 | 0.000791 |
| rs541065612 | 15 | 42924612 | T | C | 402071 | 3 | 0.000527 |
| rs7162040 | 15 | 42999994 | T | C | 1168991 | 101 | 0.00142 |
| rs182845723 | 15 | 42924574 | T | C | 1172156 | 107 | 0.00177 |
| rs187967591 | 15 | 42948375 | G | A | 392812 | 2 | 4.46E-05 |
| rs201643568 | 15 | 42984156 | C | G | 1204652 | 121 | 0.00337 |
| rs778329983 | 15 | 43009275 | C | T | 408773 | 4 | 9.91E-05 |
| rs533415918 | 15 | 42896101 | A | G | 396915 | 2 | 0.000498 |
| rs545016214 | 15 | 42940515 | C | T | 1197878 | 118 | 0.00337 |
| rs181506969 | 15 | 42971266 | G | A | 1018719 | 89 | 0.00156 |
| rs374659128 | 15 | 42983017 | TTAA | T | 622659 | 3 | 4.42E-05 |
| rs189692396 | 15 | 42879140 | T | C | 628358 | 4 | 8.67E-05 |
| rs115044157 | 15 | 42891021 | C | T | 769966 | 24 | 0.000192 |
| rs189296046 | 15 | 42906614 | C | A | 400053 | 3 | 0.000257 |
| rs531332837 | 15 | 42907087 | C | T | 680328 | 14 | 0.000383 |
| rs149026782 | 15 | 42959209 | C | T | 615331 | 3 | 3.01E-05 |
| rs370811683 | 15 | 42964861 | A | AG | 641951 | 10 | 0.000725 |
| rs560093737 | 15 | 42979758 | T | G | 403830 | 4 | 0.000328 |
| rs144819529 | 15 | 42958167 | A | G | 615331 | 3 | 3.01E-05 |
| rs78081991 | 15 | 42990078 | T | C | 697839 | 12 | 0.000181 |
| rs555394489 | 15 | 42895336 | C | A | 59567 | 11 | 0.000369 |
| rs567963038 | 15 | 43009945 | C | T | 825410 | 25 | 3.00E-04 |
| rs116809323 | 15 | 42868836 | G | C | 616670 | 3 | 0.000144 |
| rs569234592 | 15 | 42907158 | G | T | 397969 | 2 | 9.55E-05 |
| rs117170089 | 15 | 42997618 | G | T | 979168 | 64 | 0.00135 |
| rs113162826 | 15 | 42897162 | C | T | 647564 | 39 | 0.000344 |
| rs141930037 | 15 | 42901502 | C | G | 614586 | 2 | 2.85E-05 |
| rs531652303 | 15 | 42915607 | A | G | 611581 | 55 | 0.00106 |
| rs550992626 | 15 | 42900445 | A | G | 619157 | 31 | 0.000199 |
| rs748384434 | 15 | 42989599 | T | G | 602741 | 27 | 0.000184 |
| rs764650416 | 15 | 42993496 | G | T | 509303 | 17 | 0.000343 |
| rs577790650 | 15 | 42876033 | C | T | 610833 | 41 | 0.000513 |
| rs188267149 | 15 | 42891458 | C | G | 177388 | 28 | 0.00221 |
| rs575816891 | 15 | 42961747 | C | G | 878370 | 77 | 0.00145 |
| rs201393731 | 15 | 42979099 | C | T | 736709 | 35 | 0.000443 |
| rs181986747 | 15 | 42995783 | C | T | 652229 | 7 | 0.000123 |
| NA | 15 | 42885513 | R | D | 1339 | 2 | 0.0232 |
| rs75548024 | 15 | 42887849 | A | G | 439557 | 4 | 5.60E-05 |
| rs115420503 | 15 | 42909687 | A | T | 1072782 | 119 | 0.00269 |
| rs144400192 | 15 | 42933225 | A | G | 237411 | 3 | 0.000457 |
| rs9652443 | 15 | 42892631 | T | C | 667875 | 7 | 8.46E-05 |
| rs143562438 | 15 | 42893927 | A | G | 234483 | 2 | 0.000186 |
| rs142562373 | 15 | 42965599 | C | CA | 990499 | 55 | 0.000995 |
| rs376229251 | 15 | 42978141 | A | C | 295141 | 2 | 7.28E-05 |
| rs112745231 | 15 | 42982708 | A | G | 650739 | 33 | 0.000303 |
| rs532451462 | 15 | 42901557 | C | T | 405502 | 6 | 4.07E-05 |
| rs77836884 | 15 | 42986787 | C | T | 798186 | 17 | 0.000119 |
| rs373520291 | 15 | 42991781 | T | C | 644603 | 4 | 0.000247 |
| rs555203230 | 15 | 42994269 | C | A | 266647 | 7 | 0.000306 |
| rs555233136 | 15 | 42960377 | T | C | 791285 | 27 | 0.000711 |
| rs181989685 | 15 | 42869480 | C | T | 659512 | 6 | 2.00E-04 |
| rs190857140 | 15 | 42870763 | G | A | 1185646 | 106 | 0.00156 |
| rs183757978 | 15 | 42976040 | C | T | 404962 | 4 | 0.00261 |
| rs776011349 | 15 | 43005955 | A | G | 463564 | 3 | 0.000113 |
| rs537743332 | 15 | 42886644 | C | T | 398915 | 3 | 0.00473 |
| rs764365243 | 15 | 42918606 | TTCC | T | 389982 | 2 | 0.0123 |
| rs528276071 | 15 | 42979359 | CAGCACA | C | 171213 | 30 | 0.00233 |
| rs146774151 | 15 | 42982996 | G | C | 617549 | 3 | 8.02E-05 |
| rs78261202 | 15 | 43003113 | G | T | 234483 | 2 | 0.000343 |
| rs115541077 | 15 | 43005554 | A | G | 234483 | 2 | 8.96E-05 |
| rs151260934 | 15 | 42951968 | T | G | 614586 | 2 | 3.58E-05 |
| rs570832687 | 15 | 42976712 | G | C | 29486 | 8 | 0.00231 |
| rs369454740 | 15 | 42983848 | T | C | 72007 | 3 | 0.00106 |
| rs116602451 | 15 | 42932722 | T | C | 614586 | 2 | 7.32E-05 |
| NA | 15 | 42933807 | R | D | 1339 | 2 | 0.169 |
| rs112240126 | 15 | 42884761 | G | A | 1119860 | 83 | 0.000998 |
| rs187486986 | 15 | 42924731 | T | C | 408289 | 6 | 0.000249 |
| rs567259695 | 15 | 42978443 | G | A | 71675 | 2 | 0.000544 |
| rs115488672 | 15 | 43007595 | A | C | 645543 | 5 | 0.000124 |
| rs756943396 | 15 | 42947980 | T | A | 498514 | 12 | 0.000311 |
| rs940605313 | 15 | 42959877 | A | C | 539337 | 12 | 0.000172 |
| rs769391315 | 15 | 42894159 | C | T | 199528 | 3 | 0.00135 |
| rs116406869 | 15 | 42934371 | T | G | 614586 | 2 | 7.32E-05 |
| rs201119685 | 15 | 42994073 | T | TC | 618688 | 3 | 0.000116 |
| rs143643681 | 15 | 42903812 | C | T | 618688 | 3 | 0.000126 |

|  |  |  |  |  |  |  |  |
| --- | --- | --- | --- | --- | --- | --- | --- |
| rs562232008 | 15 | 42936049 | A | G | 834242 | 58 | 0.00116 |
| rs144482277 | 15 | 42938217 | G | C | 618363 | 2 | 7.12E-05 |
| rs200058353 | 15 | 42983771 | C | G | 830577 | 25 | 0.000424 |
| rs76465418 | 15 | 42887258 | C | A | 740624 | 17 | 0.000485 |
| rs74714963 | 15 | 42881700 | C | T | 672199 | 7 | 0.000255 |
| rs182535861 | 15 | 42908849 | G | C | 584484 | 11 | 0.000346 |
| rs201349756 | 15 | 42978424 | C | T | 454528 | 14 | 0.000587 |
| rs78904754 | 15 | 42994141 | C | T | 614586 | 2 | 3.09E-05 |
| rs151166224 | 15 | 42999982 | A | G | 1134777 | 96 | 0.00161 |
| rs79927491 | 15 | 42880350 | A | G | 623519 | 3 | 0.000108 |
| rs572298610 | 15 | 42869451 | A | G | 585584 | 40 | 0.00203 |
| rs370550265 | 15 | 42891689 | C | T | 392813 | 2 | 0.000126 |
| rs546129700 | 15 | 42914280 | A | T | 397969 | 2 | 0.000519 |
| rs546865775 | 15 | 42922076 | A | G | 24858 | 6 | 0.000624 |
| rs753994390 | 15 | 42964301 | C | G | 456991 | 10 | 0.000241 |
| rs565525729 | 15 | 42983143 | C | A | 398919 | 4 | 0.00295 |
| rs74715782 | 15 | 42903428 | G | A | 695637 | 11 | 0.000427 |
| rs191655165 | 15 | 42904217 | A | G | 618363 | 3 | 0.000202 |
| rs189916861 | 15 | 42934339 | G | A | 903556 | 44 | 0.000525 |
| rs150313767 | 15 | 42958792 | G | A | 237411 | 3 | 0.000472 |
| rs573507159 | 15 | 42990952 | C | G | 614586 | 2 | 0.000148 |
| rs576898675 | 15 | 42900931 | A | G | 392813 | 2 | 4.84E-05 |
| rs138717526 | 15 | 42902159 | T | C | 774212 | 25 | 0.000589 |
| rs1045397885 | 15 | 42916252 | C | A | 398410 | 4 | 0.000131 |
| rs568016106 | 15 | 42959617 | G | A | 237411 | 3 | 0.000472 |
| rs539091007 | 15 | 42893886 | G | A | 614586 | 2 | 2.93E-05 |
| rs566546449 | 15 | 42978632 | C | T | 397969 | 2 | 0.00044 |
| rs576523100 | 15 | 42992120 | G | T | 392813 | 2 | 0.000115 |
| rs75974156 | 15 | 42876745 | A | G | 783464 | 35 | 0.000431 |
| NA | 15 | 42882219 | C | T | 75193 | 2 | 0.000246 |
| rs192447262 | 15 | 42889845 | C | T | 614586 | 2 | 2.93E-05 |
| rs777242940 | 15 | 42944170 | CTG | C | 458627 | 2 | 0.00142 |
| rs185205489 | 15 | 42961844 | C | T | 237411 | 3 | 0.00047 |
| rs79774962 | 15 | 43005092 | C | T | 1160013 | 96 | 0.00141 |
| rs143805766 | 15 | 43009642 | G | A | 834188 | 21 | 0.000426 |
| rs201351089 | 15 | 42943656 | A | AT | 642203 | 8 | 0.000198 |
| rs115794149 | 15 | 42980441 | A | G | 687254 | 11 | 9.60E-05 |
| rs533630312 | 15 | 42903619 | C | T | 412832 | 6 | 0.000184 |
| rs183794720 | 15 | 42935957 | T | C | 614586 | 2 | 3.58E-05 |
| rs185622584 | 15 | 42940650 | C | T | 398915 | 3 | 0.00422 |
| rs552552047 | 15 | 42874020 | C | T | 798433 | 19 | 0.00036 |
| rs186670557 | 15 | 42893536 | A | G | 665791 | 6 | 8.34E-05 |
| rs192352355 | 15 | 42959921 | C | T | 1045706 | 50 | 0.000404 |
| rs2899060 | 15 | 42899437 | G | T | 246520 | 3 | 0.000357 |
| rs185441627 | 15 | 42937866 | T | G | 635105 | 8 | 0.000149 |
| rs569708425 | 15 | 42946670 | G | C | 411709 | 5 | 0.00248 |
| rs141542733 | 15 | 42911557 | C | T | 396915 | 3 | 0.00043 |
| rs553969178 | 15 | 42915402 | A | C | 665927 | 16 | 0.00126 |

|  |  |  |  |  |  |  |  |
| --- | --- | --- | --- | --- | --- | --- | --- |
| rs373300237 | 15 | 42918621 | CTCT | C | 1064481 | 100 | 0.00989 |
| rs535301205 | 15 | 42956781 | T | C | 568465 | 7 | 0.00024 |
| rs550267054 | 15 | 42933960 | G | A | 542813 | 16 | 0.000232 |
| rs763159117 | 15 | 42949738 | A | C | 435549 | 9 | 6.20E-05 |
| rs149335396 | 15 | 42882419 | T | G | 1164777 | 123 | 0.00281 |
| rs61707019 | 15 | 42918962 | G | A | 983709 | 84 | 0.00124 |
| rs117522776 | 15 | 42963841 | C | T | 622448 | 25 | 0.000145 |
| rs763432324 | 15 | 43002932 | G | A | 528084 | 5 | 0.000619 |
| rs567147466 | 15 | 43004874 | A | G | 450700 | 9 | 4.10E-05 |
| rs144132229 | 15 | 43012462 | G | A | 669863 | 9 | 0.000104 |
| rs559815648 | 15 | 42886131 | T | G | 410452 | 4 | 0.00047 |
| rs558081449 | 15 | 43005954 | G | C | 992445 | 76 | 0.00108 |
| rs73406562 | 15 | 42877863 | C | T | 1117473 | 83 | 0.000963 |
| rs35172521 | 15 | 42886287 | GAGAC | G | 627296 | 4 | 0.000367 |
| NA | 15 | 42973934 | G | T | 511832 | 6 | 0.000637 |
| rs115595275 | 15 | 42931033 | A | T | 1044601 | 45 | 0.000547 |
| rs116814420 | 15 | 42994820 | G | A | 234483 | 2 | 8.32E-05 |
| rs75617784 | 15 | 42914904 | G | A | 615417 | 3 | 5.69E-05 |
| rs898531013 | 15 | 42956882 | T | A | 563214 | 14 | 0.000175 |
| rs141297638 | 15 | 42893808 | G | A | 514237 | 89 | 0.00728 |
| rs111312181 | 15 | 42898767 | C | T | 647564 | 39 | 0.000346 |
| rs13379865 | 15 | 42898498 | C | T | 776802 | 25 | 0.00057 |
| rs144638832 | 15 | 42912991 | C | T | 717457 | 11 | 0.000187 |
| rs147795392 | 15 | 42953329 | A | G | 945451 | 59 | 0.00113 |
| rs145900049 | 15 | 42978144 | A | T | 614586 | 2 | 2.93E-05 |
| rs202070432 | 15 | 42983192 | A | G | 955189 | 65 | 0.00187 |
| rs541324309 | 15 | 42907683 | AT | A | 53568 | 14 | 0.00147 |
| rs1020271692 | 15 | 42972418 | T | C | 532927 | 11 | 0.000175 |
| rs748249971 | 15 | 42888005 | T | A | 610821 | 21 | 0.000245 |
| rs763317493 | 15 | 42905383 | ACTC | A | 458627 | 2 | 0.00045 |
| rs16973395 | 15 | 42912191 | T | C | 719541 | 12 | 0.000189 |
| rs143677353 | 15 | 42971296 | G | C | 710303 | 5 | 5.28E-05 |
| rs777870405 | 15 | 42993947 | C | A | 590795 | 38 | 0.000455 |
| rs141247863 | 15 | 42916512 | CAG | C | 397920 | 5 | 0.000293 |
| rs533481642 | 15 | 42947115 | A | G | 24165 | 6 | 0.00182 |
| rs559982616 | 15 | 42936475 | A | G | 389583 | 2 | 5.52E-05 |
| rs149780137 | 15 | 42947732 | A | G | 1047381 | 48 | 0.000562 |
| rs77245532 | 15 | 42975619 | A | G | 933106 | 39 | 0.000403 |
| rs151016272 | 15 | 43008947 | A | G | 403830 | 4 | 0.00141 |
| rs545978244 | 15 | 42868026 | C | A | 407603 | 5 | 0.00142 |
| rs749066345 | 15 | 42880005 | C | T | 474548 | 14 | 0.000291 |
| rs151217518 | 15 | 42923871 | G | A | 614586 | 2 | 2.93E-05 |
| rs551667710 | 15 | 42960145 | A | C | 569499 | 20 | 0.00048 |
| rs1053290276 | 15 | 42913357 | G | A | 402708 | 3 | 0.000107 |
| rs55651630 | 15 | 42961994 | A | G | 1248900 | 175 | 0.191 |
| rs57258642 | 15 | 43003656 | A | T | 835210 | 25 | 0.000372 |
| NA | 15 | 42918201 | C | T | 40976 | 7 | 0.00118 |
| rs573571036 | 15 | 42997345 | G | C | 397969 | 2 | 0.000452 |
| rs530762141 | 15 | 43009056 | G | A | 424065 | 6 | 0.000159 |
| NA | 15 | 42879347 | CCCAG | TCCAG | 226295 | 2 | 8.84E-05 |
| rs146135977 | 15 | 42889446 | A | G | 1164321 | 116 | 0.00241 |
| rs183088570 | 15 | 42908063 | T | C | 176662 | 22 | 0.00208 |
| rs140917325 | 15 | 42872335 | G | A | 1164322 | 116 | 0.0024 |
| rs537910025 | 15 | 42875968 | T | C | 418214 | 7 | 0.000219 |
| rs114641497 | 15 | 42972198 | C | T | 710303 | 5 | 5.28E-05 |
| rs112765931 | 15 | 42931212 | GAAAC | G | 1001425 | 71 | 0.00213 |
| rs925991470 | 15 | 42965660 | G | A | 603848 | 30 | 0.000282 |
| rs139168421 | 15 | 42970685 | G | C | 232059 | 3 | 8.83E-05 |
| NA | 15 | 42995728 | R | D | 1339 | 2 | 0.0362 |
| rs770679768 | 15 | 43005828 | T | G | 521526 | 8 | 0.000244 |
| rs529032170 | 15 | 42960888 | T | G | 246870 | 2 | 0.000126 |
| rs142686033 | 15 | 42899186 | C | T | 618363 | 3 | 0.000208 |
| rs528635271 | 15 | 42935706 | T | A | 476666 | 12 | 0.000263 |
| rs79331266 | 15 | 42953161 | T | G | 230866 | 4 | 0.000574 |
| rs190350231 | 15 | 42892087 | G | A | 845024 | 119 | 0.00365 |
| rs574063567 | 15 | 42893636 | G | A | 276755 | 5 | 0.000186 |
| rs531146879 | 15 | 42889936 | T | C | 1164323 | 116 | 0.00241 |
| rs559447218 | 15 | 42978245 | A | T | 402071 | 3 | 0.000402 |
| rs8039346 | 15 | 42987070 | G | C | 1151693 | 95 | 0.00113 |
| rs189545083 | 15 | 42992904 | A | G | 1089079 | 107 | 0.00198 |
| rs531067051 | 15 | 43011162 | C | T | 470549 | 12 | 7.65E-05 |
| rs142780671 | 15 | 42910004 | T | C | 235314 | 3 | 0.000193 |
| rs571609256 | 15 | 42996263 | G | A | 1076159 | 62 | 0.000645 |
| rs7164196 | 15 | 42869986 | G | T | 709495 | 12 | 0.000216 |
| rs184997894 | 15 | 42889955 | G | A | 757448 | 11 | 0.000158 |
| rs534725679 | 15 | 42985477 | A | G | 402554 | 4 | 5.96E-05 |
| rs142916862 | 15 | 42913461 | A | G | 567990 | 17 | 0.000436 |
| rs76108347 | 15 | 42955118 | A | C | 734882 | 18 | 0.000201 |
| rs143933986 | 15 | 42888244 | A | G | 1164322 | 116 | 0.0024 |
| rs188518430 | 15 | 42895715 | C | A | 618363 | 3 | 0.000218 |
| rs115887309 | 15 | 42954682 | T | G | 774082 | 30 | 0.000389 |
| rs143394146 | 15 | 42986676 | A | G | 263193 | 6 | 0.000199 |
| rs149356155 | 15 | 43009603 | T | C | 1050533 | 51 | 6.00E-04 |
| rs943592293 | 15 | 42950524 | C | G | 29486 | 8 | 0.0028 |
| rs746489852 | 15 | 43002363 | A | G | 502933 | 8 | 0.000424 |
| rs8028863 | 15 | 43012764 | T | G | 837583 | 26 | 0.000392 |
| rs75891360 | 15 | 42891957 | A | C | 687584 | 9 | 0.00038 |
| rs776239194 | 15 | 42910488 | AC | A | 459573 | 3 | 0.00337 |
| rs150192451 | 15 | 42931427 | C | G | 573302 | 17 | 0.000629 |
| rs562486786 | 15 | 42958834 | A | G | 514697 | 3 | 0.000154 |
| rs183640181 | 15 | 42963391 | C | T | 630896 | 4 | 0.000211 |
| rs138430407 | 15 | 42878735 | C | G | 687584 | 9 | 0.000296 |
| rs140942120 | 15 | 42885052 | C | G | 1164322 | 116 | 0.00239 |
| rs73406582 | 15 | 42912925 | C | A | 872182 | 42 | 0.0014 |
| NA | 15 | 42916751 | R | D | 1339 | 2 | 0.0321 |
| rs574401311 | 15 | 42991813 | G | C | 622770 | 4 | 9.55E-05 |

|  |  |  |  |  |  |  |  |
| --- | --- | --- | --- | --- | --- | --- | --- |
| rs141047217 | 15 | 42913701 | G | A | 992066 | 60 | 0.000824 |
| rs527693525 | 15 | 42946090 | T | C | 1167916 | 119 | 0.00506 |
| rs530331777 | 15 | 42955899 | A | G | 810981 | 48 | 0.000756 |
| rs115058341 | 15 | 42901551 | A | T | 695636 | 10 | 0.000418 |
| rs913429324 | 15 | 42999523 | G | A | 27797 | 7 | 0.00104 |
| rs1050368245 | 15 | 42918465 | T | C | 20742 | 2 | 0.00118 |
| rs144697433 | 15 | 42891968 | C | G | 1160999 | 112 | 0.00194 |
| NA | 15 | 42988491 | A | G | 519885 | 6 | 0.000632 |
| rs8036993 | 15 | 43010887 | G | A | 668605 | 8 | 0.000165 |
| rs868202680 | 15 | 42868358 | C | T | 446040 | 7 | 0.000201 |
| rs141248078 | 15 | 42879852 | C | G | 234483 | 2 | 0.000132 |
| rs185865927 | 15 | 42884511 | C | T | 616739 | 3 | 5.19E-05 |
| rs535127983 | 15 | 42889684 | G | A | 57422 | 13 | 0.00131 |
| rs540721584 | 15 | 42975513 | G | T | 630115 | 5 | 0.000232 |
| rs140746565 | 15 | 42978102 | C | T | 710303 | 5 | 5.49E-05 |
| rs28537072 | 15 | 42958484 | C | A | 1136919 | 103 | 0.00142 |
| rs140924205 | 15 | 42977810 | T | G | 710303 | 5 | 5.49E-05 |
| rs569984356 | 15 | 42869414 | C | T | 566560 | 27 | 0.000264 |
| rs778181709 | 15 | 42999548 | A | G | 515603 | 18 | 0.000215 |
| rs188252842 | 15 | 42872751 | C | T | 766512 | 15 | 0.000172 |
| rs79147516 | 15 | 42884438 | A | G | 672199 | 7 | 0.000269 |
| rs569250173 | 15 | 42904514 | CAGTG | C | 758546 | 26 | 0.00417 |
| rs895643189 | 15 | 42889532 | A | G | 19722 | 5 | 0.00112 |
| rs539820661 | 15 | 42893649 | C | T | 622659 | 3 | 4.18E-05 |
| rs550483327 | 15 | 42911559 | G | A | 254943 | 3 | 8.20E-05 |
| rs10162619 | 15 | 42879686 | T | G | 272978 | 5 | 0.000169 |
| rs894513132 | 15 | 42911619 | C | T | 402708 | 3 | 0.000106 |
| rs116205969 | 15 | 42912809 | A | G | 717457 | 11 | 0.000187 |
| rs118052909 | 15 | 42953376 | C | G | 515726 | 16 | 0.000457 |
| rs761882822 | 15 | 42954006 | T | G | 619382 | 31 | 0.000383 |
| rs28378282 | 15 | 42884072 | A | T | 272978 | 5 | 0.000167 |
| rs116417375 | 15 | 42997962 | A | G | 649064 | 6 | 6.24E-05 |
| rs551970799 | 15 | 42929307 | C | T | 438778 | 7 | 4.40E-05 |
| rs147737228 | 15 | 42881116 | G | A | 272978 | 5 | 0.000163 |
| rs181968356 | 15 | 42976786 | G | A | 666447 | 38 | 0.000507 |
| rs544006711 | 15 | 43010245 | G | A | 396915 | 3 | 0.000212 |
| rs150784222 | 15 | 42985224 | A | G | 400053 | 3 | 3.00E-04 |
| rs183598354 | 15 | 42902427 | T | G | 629491 | 40 | 0.00032 |
| rs556127396 | 15 | 42998583 | A | G | 397969 | 2 | 0.000176 |
| rs571385368 | 15 | 42872500 | T | TC | 782347 | 22 | 0.000362 |
| rs186001502 | 15 | 42903616 | T | G | 927535 | 88 | 0.00139 |
| rs769973104 | 15 | 42989600 | T | C | 600069 | 26 | 0.000178 |
| rs116084720 | 15 | 42999985 | A | G | 236544 | 2 | 7.61E-05 |
| rs190352047 | 15 | 42878577 | C | G | 408039 | 5 | 0.000652 |
| rs1033036112 | 15 | 42933713 | C | T | 234445 | 30 | 0.00155 |
| rs187488302 | 15 | 42941594 | C | T | 916880 | 52 | 0.000519 |
| rs887317340 | 15 | 42958646 | G | A | 633629 | 30 | 0.000267 |
| rs116737895 | 15 | 42986042 | C | T | 551939 | 21 | 0.000461 |

|  |  |  |  |  |  |  |  |
| --- | --- | --- | --- | --- | --- | --- | --- |
| rs535868330 | 15 | 42916538 | TATAAC | T | 515320 | 25 | 0.00411 |
| rs376374890 | 15 | 43009962 | C | T | 577616 | 18 | 0.000345 |
| rs139430686 | 15 | 42870497 | C | T | 1164320 | 116 | 0.00241 |
| rs761953576 | 15 | 42886839 | T | C | 511074 | 7 | 0.00024 |
| rs201252548 | 15 | 43011726 | G | A | 637397 | 7 | 2.67E-05 |
| rs575026169 | 15 | 42960629 | A | G | 479810 | 8 | 0.000234 |
| rs7163324 | 15 | 42988175 | C | T | 1149860 | 93 | 0.00113 |
| rs540055435 | 15 | 42873773 | AGC | A | 226381 | 2 | 8.61E-05 |
| rs145869433 | 15 | 42983572 | G | A | 986025 | 77 | 0.00149 |
| rs77876077 | 15 | 42869977 | G | C | 801207 | 16 | 0.000398 |
| rs372097263 | 15 | 42879057 | G | T | 226381 | 2 | 8.83E-05 |
| rs114539185 | 15 | 42914152 | T | G | 634451 | 4 | 0.000163 |
| rs369242363 | 15 | 42978186 | C | T | 698727 | 26 | 0.000471 |
| rs144271727 | 15 | 42876084 | T | G | 429792 | 4 | 4.50E-05 |
| rs569941620 | 15 | 42937337 | C | T | 496663 | 4 | 0.00013 |
| rs146410393 | 15 | 43000332 | C | A | 649064 | 6 | 6.39E-05 |
| rs141553512 | 15 | 43006043 | T | G | 355240 | 15 | 0.000528 |
| rs1017461242 | 15 | 42950058 | C | A | 551939 | 21 | 0.000463 |
| rs552403828 | 15 | 42951233 | C | A | 614586 | 2 | 2.93E-05 |
| rs557261358 | 15 | 42988192 | A | G | 411708 | 6 | 0.00116 |
| rs116032750 | 15 | 43000294 | T | G | 649064 | 6 | 6.32E-05 |
| rs369695076 | 15 | 43002057 | TTTCTC | T | 758974 | 18 | 0.000387 |
| rs570880299 | 15 | 42883424 | G | A | 515369 | 8 | 0.000296 |
| rs7174847 | 15 | 42913750 | A | T | 695079 | 9 | 0.000212 |
| rs60051108 | 15 | 42920507 | C | T | 952202 | 59 | 0.00121 |
| rs191342069 | 15 | 42968129 | C | T | 680378 | 42 | 0.000536 |
| rs150686043 | 15 | 42921661 | A | G | 400053 | 3 | 0.000312 |
| rs549994482 | 15 | 42924359 | T | G | 233604 | 2 | 8.56E-05 |
| rs115827874 | 15 | 42874859 | C | T | 614586 | 2 | 4.72E-05 |
| rs183802733 | 15 | 42955055 | C | T | 742502 | 19 | 0.00025 |
| rs765405244 | 15 | 42993620 | G | A | 458627 | 2 | 0.000268 |
| rs895701914 | 15 | 42924428 | A | G | 535579 | 10 | 9.99E-05 |
| rs184795205 | 15 | 42974899 | C | T | 1046875 | 48 | 0.000588 |
| rs535923283 | 15 | 42869430 | C | T | 972441 | 57 | 0.00158 |
| rs183580304 | 15 | 42891278 | T | C | 687584 | 9 | 0.000372 |
| rs547496925 | 15 | 42883035 | G | T | 839755 | 116 | 0.00322 |
| NA | 15 | 42884547 | A | G | 394638 | 2 | 0.000109 |
| rs200261102 | 15 | 42889573 | A | AT | 730069 | 53 | 0.00462 |
| rs77880860 | 15 | 42939905 | T | G | 830890 | 50 | 0.00412 |
| rs575135631 | 15 | 42980297 | A | G | 390358 | 2 | 4.74E-05 |
| rs74009211 | 15 | 43008971 | C | A | 722327 | 17 | 0.000588 |
| rs138113203 | 15 | 42874407 | C | G | 614586 | 2 | 4.72E-05 |
| rs572054086 | 15 | 42890464 | C | T | 226381 | 2 | 9.72E-05 |
| rs138257631 | 15 | 42921324 | C | A | 1212301 | 138 | 0.00833 |
| rs184720517 | 15 | 42991842 | T | C | 821929 | 23 | 0.000639 |
| rs140027750 | 15 | 43001862 | G | C | 666863 | 6 | 0.000112 |
| rs189480172 | 15 | 42993596 | T | A | 614586 | 2 | 6.75E-05 |
| rs986169146 | 15 | 42950972 | G | A | 472066 | 15 | 0.000139 |
| rs34081955 | 15 | 42896068 | AG | A | 389982 | 2 | 0.015 |
| NA | 15 | 42944185 | G | A | 85076 | 3 | 0.0022 |
| rs112267992 | 15 | 42895540 | G | A | 614586 | 2 | 5.21E-05 |
| rs149877181 | 15 | 42972932 | G | A | 1154398 | 109 | 0.0033 |
| rs144084975 | 15 | 42870498 | G | T | 683807 | 8 | 0.000291 |
| rs115105685 | 15 | 42874825 | T | A | 264894 | 4 | 0.000162 |
| rs193064675 | 15 | 42882083 | G | A | 927971 | 45 | 0.000494 |
| rs16956932 | 15 | 42926883 | A | G | 1012860 | 43 | 0.000939 |
| NA | 15 | 42977820 | C | A | 29486 | 8 | 0.000695 |
| rs562680697 | 15 | 43001428 | T | TAC | 83954 | 4 | 0.000578 |
| rs138157695 | 15 | 42993425 | G | A | 527851 | 27 | 0.000362 |
| rs148764121 | 15 | 42999844 | A | G | 1049514 | 50 | 0.000597 |
| rs774605196 | 15 | 42987180 | CA | C | 99077 | 9 | 0.012 |
| rs569851918 | 15 | 42988920 | GGAGA | G | 392813 | 2 | 0.00111 |
| rs535667563 | 15 | 42918555 | C | T | 808488 | 38 | 0.000505 |
| rs9972577 | 15 | 42910951 | C | T | 686714 | 10 | 0.000364 |
| rs540343022 | 15 | 42915699 | T | C | 396915 | 3 | 0.00047 |
| rs148382999 | 15 | 42942895 | A | G | 714079 | 21 | 0.00145 |
| rs563033349 | 15 | 42959907 | T | C | 397968 | 2 | 0.000921 |
| rs77922743 | 15 | 42993355 | C | G | 1152792 | 95 | 0.00119 |
| rs60484860 | 15 | 43008972 | G | A | 916051 | 50 | 0.00144 |
| rs551060551 | 15 | 43004638 | T | C | 684177 | 3 | 7.89E-05 |
| rs568646935 | 15 | 42894468 | C | A | 493437 | 18 | 0.000496 |
| rs201346447 | 15 | 42967124 | A | G | 1096816 | 71 | 0.00102 |
| rs200346352 | 15 | 42872718 | T | TA | 405032 | 4 | 0.000332 |
| rs539366399 | 15 | 42973867 | C | T | 393138 | 2 | 0.000212 |
| rs116318862 | 15 | 42989917 | A | G | 1031918 | 99 | 0.00173 |
| NA | 15 | 42887096 | A | G | 20003 | 2 | 0.0042 |
| rs370868295 | 15 | 42898953 | C | T | 618363 | 3 | 9.30E-05 |
| rs190028056 | 15 | 42941943 | G | A | 228044 | 2 | 0.000118 |
| rs148023580 | 15 | 42960312 | G | A | 1145692 | 100 | 0.00132 |
| rs566582680 | 15 | 42886495 | G | A | 614586 | 2 | 6.35E-05 |
| rs114128771 | 15 | 42934908 | C | T | 723295 | 12 | 0.00014 |
| rs144363278 | 15 | 42968140 | G | A | 317626 | 10 | 0.000168 |
| NA | 15 | 42975732 | T | C | 72843 | 3 | 0.003 |
| rs59900882 | 15 | 43002685 | C | G | 1169211 | 102 | 0.0014 |
| rs542154361 | 15 | 42908599 | C | T | 237019 | 5 | 0.000576 |
| rs765293647 | 15 | 42921511 | C | A | 526605 | 13 | 0.000486 |
| rs556659632 | 15 | 42933766 | C | T | 396915 | 3 | 0.000215 |
| rs367902230 | 15 | 42935270 | TATAAA | T | 234483 | 2 | 7.46E-05 |
| rs541204716 | 15 | 42889085 | C | T | 397969 | 2 | 0.000648 |
| rs4924688 | 15 | 42910548 | C | T | 726226 | 25 | 0.000998 |
| rs570255301 | 15 | 42940956 | TC | T | 243657 | 6 | 0.000667 |
| rs116360099 | 15 | 43007017 | G | A | 649064 | 6 | 6.47E-05 |
| rs77171961 | 15 | 42882895 | G | C | 687584 | 9 | 0.000319 |
| rs570858258 | 15 | 43008007 | A | C | 242715 | 4 | 0.000111 |
| rs182385853 | 15 | 43009489 | C | T | 1230916 | 146 | 0.00365 |
| rs889746332 | 15 | 42875974 | C | T | 437910 | 7 | 0.00015 |

|  |  |  |  |  |  |  |  |
| --- | --- | --- | --- | --- | --- | --- | --- |
| rs557571324 | 15 | 42894876 | C | G | 566103 | 20 | 0.000178 |
| rs759560583 | 15 | 42914523 | T | C | 229719 | 7 | 0.00185 |
| rs552844514 | 15 | 42950749 | T | G | 70390 | 3 | 0.000668 |
| rs137999320 | 15 | 42868678 | T | G | 226381 | 2 | 9.06E-05 |
| rs556780600 | 15 | 42925880 | C | T | 581690 | 18 | 0.000133 |
| rs539280750 | 15 | 42929787 | T | C | 571739 | 17 | 0.00048 |
| rs116577053 | 15 | 42941664 | C | G | 240085 | 2 | 9.79E-05 |
| rs115532536 | 15 | 42879312 | C | T | 687584 | 9 | 0.000297 |
| rs756135004 | 15 | 42904832 | G | C | 456238 | 6 | 0.000212 |
| rs562322333 | 15 | 42992185 | G | A | 401746 | 3 | 0.000866 |
| rs76471889 | 15 | 42909363 | A | G | 686713 | 10 | 0.000368 |
| NA | 15 | 42965470 | G | A | 203604 | 30 | 0.00286 |
| rs552485294 | 15 | 42982602 | G | A | 822483 | 21 | 0.000227 |
| rs186697829 | 15 | 42878575 | C | A | 1028094 | 99 | 0.00234 |
| rs543442379 | 15 | 42920839 | T | G | 428643 | 4 | 5.70E-05 |
| rs528087643 | 15 | 42948574 | C | T | 403825 | 4 | 0.00124 |
| rs188025826 | 15 | 42870741 | A | G | 226381 | 2 | 8.83E-05 |
| rs571391144 | 15 | 42904356 | T | C | 397969 | 2 | 5.03E-05 |
| rs187999553 | 15 | 42908886 | G | A | 1167021 | 116 | 0.0022 |
| rs61753412 | 15 | 42986471 | A | T | 1104190 | 71 | 0.00088 |
| rs555742898 | 15 | 42994775 | T | C | 405848 | 4 | 0.000987 |
| NA | 15 | 42999666 | R | D | 1339 | 2 | 0.165 |
| rs114103856 | 15 | 42913632 | T | C | 328420 | 10 | 0.000399 |
| rs535188924 | 15 | 42991899 | TA | T | 264340 | 11 | 0.000895 |
| rs189324154 | 15 | 42868078 | G | A | 226381 | 2 | 7.95E-05 |
| rs543217741 | 15 | 42940658 | G | C | 229897 | 2 | 0.000285 |
| rs550193589 | 15 | 42966620 | GC | G | 1055890 | 95 | 0.00367 |
| rs769158133 | 15 | 42992247 | T | C | 551616 | 16 | 0.000255 |
| rs142543234 | 15 | 42873991 | A | G | 226381 | 2 | 9.06E-05 |
| rs201416428 | 15 | 42890742 | A | AT | 622200 | 3 | 9.16E-05 |
| rs7342639 | 15 | 42925904 | C | T | 690390 | 13 | 0.000261 |
| rs1013279760 | 15 | 42926283 | G | A | 394638 | 2 | 7.86E-05 |
| rs577240569 | 15 | 42959807 | G | A | 573216 | 36 | 0.000827 |
| rs546611281 | 15 | 42985711 | T | C | 802768 | 18 | 0.000224 |
| rs768035777 | 15 | 42919268 | G | A | 439612 | 8 | 0.000318 |
| rs545585433 | 15 | 42935707 | A | G | 247535 | 5 | 0.000204 |
| rs151310546 | 15 | 42881602 | C | T | 226381 | 2 | 9.50E-05 |
| rs573673780 | 15 | 42979056 | C | G | 416833 | 8 | 7.44E-05 |
| rs533919230 | 15 | 42922349 | G | A | 643488 | 5 | 5.91E-05 |
| rs530454876 | 15 | 42952699 | G | A | 528090 | 24 | 0.000373 |
| rs189260934 | 15 | 42975397 | C | T | 435357 | 6 | 0.000269 |
| rs771882568 | 15 | 42990748 | T | C | 733009 | 26 | 0.000696 |
| rs187732802 | 15 | 42905119 | T | G | 226381 | 2 | 0.000135 |
| rs920939645 | 15 | 42947341 | G | A | 516522 | 3 | 0.000133 |
| rs545657563 | 15 | 42997366 | C | T | 802768 | 18 | 0.000223 |
| rs139919621 | 15 | 42946665 | G | T | 729473 | 17 | 0.000198 |
| rs201701873 | 15 | 42877759 | A | G | 1175715 | 99 | 0.00188 |
| rs557425991 | 15 | 42950657 | A | G | 404615 | 5 | 0.000143 |

|  |  |  |  |  |  |  |  |
| --- | --- | --- | --- | --- | --- | --- | --- |
| rs114643417 | 15 | 42892578 | G | A | 226381 | 2 | 0.000104 |
| rs114533051 | 15 | 42980487 | T | A | 1025500 | 95 | 0.00169 |
| rs545086800 | 15 | 42972524 | G | A | 687329 | 18 | 0.00121 |
| rs559429992 | 15 | 43002693 | C | G | 405847 | 4 | 0.00385 |
| rs118027222 | 15 | 42908213 | G | A | 1172139 | 114 | 0.00448 |
| NA | 15 | 42901140 | C | T | 6702 | 2 | 0.005 |
| rs138873449 | 15 | 42891145 | C | T | 226381 | 2 | 0.000104 |
| rs549133434 | 15 | 42960894 | G | A | 613214 | 9 | 0.000161 |
| rs139999040 | 15 | 42883642 | C | T | 687584 | 9 | 0.000328 |
| rs61218673 | 15 | 42981978 | G | A | 667397 | 9 | 8.54E-05 |
| rs141596381 | 15 | 42898603 | T | C | 226381 | 2 | 0.000133 |
| rs185606555 | 15 | 42902671 | A | G | 226381 | 2 | 0.000115 |
| rs534891696 | 15 | 42868588 | C | T | 398915 | 3 | 0.00206 |
| rs768258586 | 15 | 42882365 | T | C | 726104 | 43 | 0.000951 |
| rs374381156 | 15 | 42946538 | TTAGAATCAC | A | 634947 | 6 | 0.000113 |
| rs116233593 | 15 | 42986987 | G | A | 1031173 | 98 | 0.00172 |
| rs561343338 | 15 | 42875196 | T | C | 396915 | 2 | 0.000586 |
| rs566344796 | 15 | 42878328 | G | A | 537292 | 29 | 0.000253 |
| rs568813363 | 15 | 42952486 | G | A | 406793 | 5 | 0.000513 |
| rs765261127 | 15 | 42954125 | A | C | 774726 | 50 | 0.000778 |
| rs771981477 | 15 | 42893470 | G | A | 29486 | 8 | 0.0138 |
| rs139196909 | 15 | 42922603 | T | C | 1146034 | 102 | 0.00175 |
| rs539890320 | 15 | 42962107 | G | T | 975315 | 47 | 0.000616 |
| rs144114925 | 15 | 42903341 | A | G | 226381 | 2 | 0.000115 |
| NA | 15 | 43006385 | C | T | 493971 | 6 | 0.000108 |
| rs563005706 | 15 | 42875064 | A | G | 396915 | 2 | 0.000586 |
| rs143110514 | 15 | 42892381 | C | T | 1228293 | 140 | 0.0038 |
| rs550231775 | 15 | 42925881 | G | A | 411709 | 6 | 0.00116 |
| rs115491632 | 15 | 42982583 | G | T | 712364 | 5 | 5.40E-05 |
| rs533154655 | 15 | 42941845 | C | T | 954443 | 60 | 0.000559 |
| rs113697440 | 15 | 42970776 | C | T | 1031875 | 101 | 0.00178 |
| rs2412739 | 15 | 42972955 | T | C | 716200 | 20 | 0.000847 |
| rs182571354 | 15 | 42870794 | A | G | 226381 | 2 | 0.000106 |
| rs184935062 | 15 | 42887479 | T | G | 463675 | 18 | 0.000149 |
| rs553132117 | 15 | 42956733 | C | CA | 803316 | 16 | 0.000309 |
| rs115067542 | 15 | 42979733 | G | A | 614586 | 2 | 2.60E-05 |
| rs114784267 | 15 | 42996974 | G | C | 268934 | 4 | 0.000115 |
| rs185294220 | 15 | 43001221 | G | C | 1226444 | 135 | 0.00451 |
| rs148405956 | 15 | 43008230 | T | C | 837426 | 12 | 0.000122 |
| rs111607613 | 15 | 43012888 | G | A | 226764 | 3 | 0.000209 |
| rs564955678 | 15 | 42965237 | G | A | 394084 | 3 | 0.00806 |
| rs140771911 | 15 | 42882036 | C | T | 803121 | 16 | 0.000457 |
| rs138565386 | 15 | 42970310 | T | C | 1070608 | 56 | 0.000684 |
| rs147572892 | 15 | 42868482 | CG | C | 629950 | 5 | 0.000221 |
| rs114844882 | 15 | 42909326 | T | C | 657707 | 5 | 9.88E-05 |
| rs142538022 | 15 | 42911994 | A | G | 614586 | 2 | 6.92E-05 |
| rs57460884 | 15 | 43008756 | C | T | 1169347 | 103 | 0.0015 |
| rs544475476 | 15 | 42905554 | G | T | 240298 | 26 | 0.000277 |

|  |  |  |  |  |  |  |  |
| --- | --- | --- | --- | --- | --- | --- | --- |
| rs143692879 | 15 | 42977628 | T | C | 1199282 | 134 | 0.00501 |
| rs183414402 | 15 | 42978417 | T | C | 883155 | 33 | 0.000425 |
| rs757526455 | 15 | 42919492 | T | G | 397223 | 3 | 0.000167 |
| rs16957031 | 15 | 42969114 | G | C | 948736 | 85 | 0.0012 |
| rs142075513 | 15 | 42923651 | G | A | 667679 | 7 | 0.000329 |
| rs376018737 | 15 | 42983973 | C | T | 483509 | 6 | 0.000367 |
| rs561210836 | 15 | 42871263 | G | A | 398915 | 3 | 0.00114 |
| rs116232111 | 15 | 42994170 | G | A | 1030898 | 99 | 0.00177 |
| rs531877342 | 15 | 43007279 | A | C | 1019083 | 46 | 0.000868 |
| rs12910030 | 15 | 42868286 | A | G | 1200721 | 165 | 0.242 |
| rs779195925 | 15 | 42892283 | A | G | 666310 | 14 | 0.000646 |
| rs538593137 | 15 | 42898914 | C | T | 401746 | 3 | 0.000331 |
| NA | 15 | 42981870 | T | C | 407939 | 2 | 0.000131 |
| rs114426631 | 15 | 42871821 | C | T | 264063 | 3 | 0.000161 |
| rs142520634 | 15 | 42890090 | G | A | 687584 | 9 | 0.000363 |
| rs376095487 | 15 | 42953467 | C | T | 234483 | 2 | 0.000104 |
| rs573448038 | 15 | 42893064 | T | C | 614585 | 2 | 0.000286 |
| rs765465286 | 15 | 42920641 | A | G | 593812 | 4 | 0.000505 |
| rs529710891 | 15 | 42935792 | G | T | 873525 | 37 | 0.000392 |
| rs115755581 | 15 | 42872867 | C | T | 653099 | 4 | 7.50E-05 |
| rs143081363 | 15 | 42991837 | C | T | 1034816 | 99 | 0.00173 |
| rs760057838 | 15 | 42869202 | G | A | 395738 | 3 | 0.000147 |
| rs149025447 | 15 | 42923445 | A | G | 614586 | 2 | 4.07E-05 |
| rs187890018 | 15 | 42973471 | G | C | 842183 | 14 | 0.000144 |
| rs745743014 | 15 | 42930019 | C | T | 490783 | 13 | 0.000218 |
| rs76506589 | 15 | 42990064 | A | G | 943028 | 54 | 0.00104 |
| rs565914422 | 15 | 42868570 | G | A | 403816 | 4 | 0.00114 |
| rs534900279 | 15 | 42869452 | C | T | 976774 | 57 | 0.00166 |
| NA | 15 | 42918707 | T | C | 419245 | 7 | 0.000428 |
| rs11273757 | 15 | 42964854 | 3AGGGGGCATC | G | 1001244 | 57 | 0.000974 |
| rs549243536 | 15 | 42968341 | T | C | 614586 | 2 | 2.60E-05 |
| rs8039217 | 15 | 42987137 | C | A | 1154881 | 95 | 0.0011 |
| rs551025057 | 15 | 42902777 | A | G | 448297 | 10 | 8.36E-05 |
| rs145526987 | 15 | 42997089 | A | C | 250480 | 4 | 0.000397 |
| rs191375699 | 15 | 42930574 | G | A | 397970 | 2 | 0.00149 |
| rs546203561 | 15 | 42923268 | C | A | 617306 | 3 | 0.00042 |
| rs60472854 | 15 | 42969243 | G | A | 623519 | 2 | 3.29E-05 |
| rs116760018 | 15 | 42887668 | T | G | 631603 | 4 | 0.000165 |
| rs529590315 | 15 | 42941017 | T | C | 458627 | 2 | 4.14E-05 |
| rs536754439 | 15 | 42947912 | A | C | 194735 | 37 | 0.000344 |
| rs750639084 | 15 | 42955218 | G | A | 510920 | 2 | 0.000207 |
| rs529988191 | 15 | 42956480 | C | G | 842183 | 14 | 0.000144 |
| rs527955560 | 15 | 42960688 | A | G | 234545 | 3 | 8.53E-05 |
| rs13380302 | 15 | 42900391 | G | C | 657707 | 5 | 8.51E-05 |
| rs1007016522 | 15 | 42910442 | T | A | 419245 | 7 | 0.000428 |
| rs577168290 | 15 | 42953432 | G | A | 403068 | 3 | 9.43E-05 |
| rs187995678 | 15 | 42990326 | C | T | 527947 | 20 | 0.000267 |
| rs532830494 | 15 | 42968254 | T | G | 871302 | 64 | 0.00248 |

|  |  |  |  |  |  |  |  |
| --- | --- | --- | --- | --- | --- | --- | --- |
| rs114416020 | 15 | 42981421 | C | T | 1026244 | 96 | 0.00172 |
| rs143465521 | 15 | 42997507 | G | A | 405848 | 4 | 0.000471 |
| rs17711135 | 15 | 42943857 | G | A | 918704 | 53 | 0.000506 |
| rs115593137 | 15 | 42970388 | G | A | 1029237 | 98 | 0.00173 |
| rs758857318 | 15 | 42989972 | T | A | 546003 | 9 | 0.000656 |
| rs61997204 | 15 | 42986065 | G | A | 701370 | 4 | 5.56E-05 |
| rs761590293 | 15 | 42882398 | C | T | 547274 | 9 | 0.000153 |
| rs190551061 | 15 | 42907287 | A | T | 268965 | 4 | 0.000584 |
| rs781346857 | 15 | 42916524 | A | G | 633256 | 30 | 0.000541 |
| rs116745790 | 15 | 42980023 | G | A | 1025499 | 95 | 0.00173 |
| rs557266111 | 15 | 42996335 | C | G | 903291 | 40 | 0.000459 |
| rs530700596 | 15 | 43002359 | G | A | 73368 | 2 | 0.00127 |
| rs577807047 | 15 | 42889101 | C | T | 448453 | 14 | 0.000165 |
| rs192217223 | 15 | 42899883 | C | A | 957536 | 50 | 0.000952 |
| rs974904581 | 15 | 42918564 | T | C | 625429 | 32 | 0.000308 |
| rs117775549 | 15 | 42973997 | C | T | 1208052 | 124 | 0.00312 |
| rs113103629 | 15 | 42987592 | C | T | 1112210 | 74 | 0.000885 |
| rs546340248 | 15 | 42935820 | G | C | 1087706 | 109 | 0.00807 |
| rs73408701 | 15 | 42968633 | T | A | 857320 | 32 | 0.000545 |
| rs553626612 | 15 | 42973738 | C | G | 398516 | 3 | 7.65E-05 |
| rs9806554 | 15 | 42876753 | T | A | 264063 | 3 | 0.000167 |
| rs768291062 | 15 | 42973213 | A | G | 644361 | 32 | 0.000379 |
| rs538590478 | 15 | 43004069 | G | A | 614586 | 2 | 3.42E-05 |
| rs565851473 | 15 | 43005664 | TACTC | T | 486971 | 10 | 0.00184 |
| rs551812998 | 15 | 42887789 | T | C | 239954 | 5 | 0.000156 |
| rs754491580 | 15 | 42898085 | A | T | 822304 | 57 | 0.00204 |
| rs79114507 | 15 | 42930636 | G | A | 723635 | 17 | 0.000238 |
| rs117486394 | 15 | 42984892 | A | C | 719070 | 14 | 0.000105 |
| rs764114296 | 15 | 42964236 | A | G | 720014 | 60 | 0.00121 |
| rs74352881 | 15 | 42972364 | A | C | 1246744 | 168 | 0.0193 |
| rs566567441 | 15 | 42965971 | G | CCATAATC | 776197 | 9 | 0.000126 |
| rs114736360 | 15 | 42969940 | C | T | 1028454 | 97 | 0.00173 |
| rs577202617 | 15 | 43000362 | A | T | 439232 | 5 | 0.000182 |
| rs528684067 | 15 | 42896224 | G | C | 1093867 | 72 | 0.00107 |
| rs564632546 | 15 | 42896159 | C | A | 541492 | 17 | 0.00014 |
| rs550195031 | 15 | 42918826 | G | A | 234545 | 4 | 8.74E-05 |
| rs140676957 | 15 | 42912825 | T | G | 232290 | 2 | 0.000418 |
| rs546990102 | 15 | 42894164 | C | G | 541492 | 17 | 0.000139 |
| rs116333487 | 15 | 42977839 | G | C | 1025499 | 95 | 0.00171 |
| rs192611335 | 15 | 42994690 | G | A | 626864 | 5 | 3.83E-05 |
| rs529904754 | 15 | 42912690 | T | A | 618618 | 3 | 0.000108 |
| rs750109325 | 15 | 42934843 | C | T | 389982 | 2 | 0.00174 |
| rs16957037 | 15 | 42971188 | C | T | 994888 | 91 | 0.00127 |
| rs754988459 | 15 | 42871305 | C | T | 191475 | 2 | 0.000402 |
| rs145054253 | 15 | 42932805 | G | A | 233093 | 3 | 0.000189 |
| rs183222293 | 15 | 42973350 | G | A | 842183 | 14 | 0.000144 |
| rs139275104 | 15 | 42988857 | C | T | 840099 | 13 | 0.000136 |
| rs142777319 | 15 | 42902570 | C | A | 27797 | 7 | 0.00103 |

|  |  |  |  |  |  |  |  |
| --- | --- | --- | --- | --- | --- | --- | --- |
| rs116259624 | 15 | 42975260 | C | T | 1032436 | 97 | 0.0017 |
| rs527501558 | 15 | 42935817 | CTA | C | 1086368 | 107 | 0.00807 |
| NA | 15 | 42944469 | R | D | 1339 | 2 | 0.0157 |
| rs532648900 | 15 | 42992572 | G | A | 66330 | 13 | 0.00133 |

| EFFECT_SIZE | SE | pvalue |
| --- | --- | --- |
| 0.0289711 | 0.00257489 | 2.28E-29 |
| 0.297366 | 0.0307943 | 4.61E-22 |
| 0.267029 | 0.029026 | 3.59E-20 |
| 0.0171269 | 0.00210829 | 4.53E-16 |
| 0.0154596 | 0.00195907 | 2.99E-15 |
| 0.0154077 | 0.00195802 | 3.57E-15 |
| 0.0153548 | 0.00195592 | 4.15E-15 |
| 0.0152443 | 0.00195122 | 5.60E-15 |
| 0.0154782 | 0.00198886 | 7.11E-15 |
| 0.264523 | 0.0339968 | 7.21E-15 |
| 0.0154681 | 0.00198946 | 7.54E-15 |
| 0.0151229 | 0.00195033 | 8.90E-15 |
| 0.0152232 | 0.00198336 | 1.65E-14 |
| 0.0753371 | 0.0104864 | 6.76E-13 |
| 0.0766565 | 0.0106881 | 7.38E-13 |
| 0.0142429 | 0.00211351 | 1.60E-11 |
| -0.0105118 | 0.00165043 | 1.90E-10 |
| -0.0105033 | 0.00165011 | 1.95E-10 |
| -0.0104723 | 0.00165064 | 2.23E-10 |
| -0.0103961 | 0.00165456 | 3.31E-10 |
| -0.0100812 | 0.00164887 | 9.72E-10 |
| -0.0106769 | 0.00175013 | 1.06E-09 |
| -0.0100431 | 0.00164652 | 1.06E-09 |
| -0.00999516 | 0.00164543 | 1.24E-09 |
| -0.0100069 | 0.00164946 | 1.31E-09 |
| -0.0101574 | 0.00168032 | 1.49E-09 |
| -0.00995526 | 0.00164901 | 1.57E-09 |
| -0.00992539 | 0.00164765 | 1.70E-09 |
| -0.0100578 | 0.00167269 | 1.82E-09 |
| -0.00992408 | 0.00165344 | 1.95E-09 |
| -0.0100722 | 0.00167819 | 1.95E-09 |
| -0.00989048 | 0.00164938 | 2.02E-09 |
| -0.00987085 | 0.00165186 | 2.29E-09 |
| -0.0100173 | 0.00168299 | 2.65E-09 |
| -0.00997979 | 0.00167818 | 2.73E-09 |
| -0.0102744 | 0.00172793 | 2.75E-09 |
| -0.00997178 | 0.00168669 | 3.38E-09 |
| -0.00995922 | 0.00168571 | 3.46E-09 |
| -0.00996156 | 0.00168644 | 3.49E-09 |
| -0.00994014 | 0.00168751 | 3.85E-09 |
| -0.00990296 | 0.00168154 | 3.88E-09 |
| -0.00991721 | 0.00168616 | 4.06E-09 |
| -0.00983051 | 0.00167208 | 4.12E-09 |
| 0.0120649 | 0.00207545 | 6.13E-09 |
| -0.0098498 | 0.00170508 | 7.62E-09 |
| -0.00980586 | 0.00171075 | 9.93E-09 |
| -0.00980097 | 0.00171338 | 1.06E-08 |

|  |  |  |
| --- | --- | --- |
| -0.00971078 | 0.00170459 | 1.22E-08 |
| -0.00970886 | 0.00170531 | 1.25E-08 |
| -0.00964931 | 0.00169965 | 1.37E-08 |
| -0.00968411 | 0.0017059 | 1.37E-08 |
| -0.00967848 | 0.0017055 | 1.39E-08 |
| -0.00969754 | 0.00170998 | 1.42E-08 |
| -0.00954277 | 0.00171752 | 2.76E-08 |
| -0.00950195 | 0.00171138 | 2.82E-08 |
| -0.00945941 | 0.00171191 | 3.28E-08 |
| -0.00949712 | 0.00171965 | 3.34E-08 |
| -0.00948539 | 0.00171817 | 3.38E-08 |
| -0.0094455 | 0.0017123 | 3.46E-08 |
| -0.00941621 | 0.00171128 | 3.75E-08 |
| -0.0092834 | 0.00168818 | 3.82E-08 |
| -0.00943018 | 0.0017164 | 3.93E-08 |
| -0.0093888 | 0.00171251 | 4.19E-08 |
| -0.00956193 | 0.00174865 | 4.55E-08 |
| -0.00934631 | 0.00171483 | 5.03E-08 |
| -0.00952596 | 0.00175226 | 5.44E-08 |
| -0.00926612 | 0.00170708 | 5.70E-08 |
| -0.00949663 | 0.00175408 | 6.16E-08 |
| -0.0094839 | 0.00175678 | 6.72E-08 |
| -0.00942946 | 0.00175251 | 7.43E-08 |
| -0.00943761 | 0.00175409 | 7.43E-08 |
| -0.00944763 | 0.00175656 | 7.51E-08 |
| -0.00930226 | 0.00173111 | 7.72E-08 |
| -0.00942927 | 0.00175794 | 8.15E-08 |
| -0.00941732 | 0.00175749 | 8.40E-08 |
| -0.00935923 | 0.00174749 | 8.52E-08 |
| -0.0093971 | 0.00175474 | 8.54E-08 |
| -0.00940029 | 0.00175856 | 9.02E-08 |
| -0.0093992 | 0.00175844 | 9.03E-08 |
| -0.00939141 | 0.00175781 | 9.16E-08 |
| -0.00931462 | 0.0017483 | 9.94E-08 |
| -0.00933566 | 0.00175795 | 1.09E-07 |
| -0.00936706 | 0.00176608 | 1.13E-07 |
| -0.0091439 | 0.00174293 | 1.55E-07 |
| -0.00938118 | 0.00179009 | 1.60E-07 |
| -0.00934334 | 0.00179328 | 1.89E-07 |
| -0.00924418 | 0.00179516 | 2.61E-07 |
| -0.00942469 | 0.0018307 | 2.63E-07 |
| -0.00922506 | 0.00179441 | 2.73E-07 |
| -0.00936067 | 0.00183786 | 3.52E-07 |
| -0.00918687 | 0.00181588 | 4.21E-07 |
| -0.00971145 | 0.00192689 | 4.66E-07 |
| 0.0078962 | 0.00160244 | 8.32E-07 |
| -0.00881037 | 0.00178931 | 8.48E-07 |
| -0.00866782 | 0.00182127 | 1.94E-06 |

|  |  |  |
| --- | --- | --- |
| -0.00845776 | 0.00178924 | 2.28E-06 |
| -0.0090262 | 0.00194312 | 3.40E-06 |
| 0.00677477 | 0.00147137 | 4.14E-06 |
| -0.00853168 | 0.00195086 | 1.22E-05 |
| -0.00789223 | 0.00181285 | 1.34E-05 |
| 0.00682142 | 0.00156966 | 1.39E-05 |
| 0.17425 | 0.0402064 | 1.46E-05 |
| -0.00870221 | 0.00201226 | 1.53E-05 |
| 0.00657117 | 0.00156062 | 2.55E-05 |
| -0.00883056 | 0.00212546 | 3.26E-05 |
| 0.00703088 | 0.00170037 | 3.55E-05 |
| 0.00588824 | 0.00142572 | 3.63E-05 |
| -0.0105104 | 0.00269057 | 9.37E-05 |
| 0.00625464 | 0.00164525 | 0.000144 |
| -0.0143412 | 0.00378971 | 0.000154 |
| 0.0889942 | 0.0236961 | 0.000173 |
| 0.00575756 | 0.00153655 | 0.000179 |
| 0.00567369 | 0.00153212 | 0.000213 |
| 0.573668 | 0.155401 | 0.000223 |
| -0.0115066 | 0.00327817 | 0.000448 |
| 0.338397 | 0.0967093 | 0.000467 |
| -0.016926 | 0.00484694 | 0.000479 |
| -0.00712783 | 0.00206948 | 0.000573 |
| -0.00719127 | 0.00209855 | 0.000611 |
| -0.0070889 | 0.00207002 | 0.000616 |
| 0.00528601 | 0.00154336 | 0.000615 |
| 0.56385 | 0.165753 | 0.00067 |
| -0.00713774 | 0.00209887 | 0.000672 |
| -0.00696886 | 0.0020725 | 0.000772 |
| 0.510293 | 0.154928 | 0.000989 |
| -0.0147894 | 0.00451805 | 0.00106 |
| -0.00845474 | 0.00261572 | 0.00123 |
| -0.0234025 | 0.00724476 | 0.00124 |
| 0.0056747 | 0.00179312 | 0.00155 |
| -0.0079624 | 0.00255445 | 0.00183 |
| -0.014367 | 0.00471201 | 0.0023 |
| -0.00608788 | 0.00205704 | 0.00308 |
| -0.433598 | 0.146716 | 0.00312 |
| 0.140536 | 0.0477269 | 0.00323 |
| -0.0140005 | 0.00475924 | 0.00326 |
| -0.431173 | 0.146638 | 0.00328 |
| -0.217707 | 0.0745415 | 0.00349 |
| -0.0133701 | 0.00466291 | 0.00414 |
| 0.0139 | 0.00485597 | 0.0042 |
| 0.251893 | 0.0880024 | 0.00421 |
| 0.232747 | 0.0818028 | 0.00444 |
| 0.0139 | 0.00489071 | 0.00448 |
| -0.0121802 | 0.00429052 | 0.00453 |

|  |  |  |
| --- | --- | --- |
| 0.184419 | 0.0655763 | 0.00492 |
| -0.0122066 | 0.00435025 | 0.00502 |
| -0.00951097 | 0.00342355 | 0.00547 |
| -0.00752264 | 0.00271746 | 0.00564 |
| -0.086818 | 0.031465 | 0.00579 |
| 0.13232 | 0.0480563 | 0.0059 |
| -0.398126 | 0.145958 | 0.00638 |
| -0.0341139 | 0.0125601 | 0.00661 |
| 0.16668 | 0.0622226 | 0.00739 |
| -0.00539283 | 0.00201792 | 0.00753 |
| -0.00919781 | 0.0034507 | 0.00769 |
| -0.340153 | 0.127795 | 0.00777 |
| -0.0254672 | 0.00962814 | 0.00817 |
| 0.20056 | 0.0764606 | 0.00871 |
| -0.066518 | 0.0255494 | 0.00923 |
| -0.00558232 | 0.00215572 | 0.00961 |
| -0.0248015 | 0.00961259 | 0.00988 |
| -0.0239585 | 0.00936027 | 0.0105 |
| -0.0192952 | 0.00754457 | 0.0105 |
| -0.00538993 | 0.00213807 | 0.0117 |
| -0.146891 | 0.0582679 | 0.0117 |
| -0.01642 | 0.0065257 | 0.0119 |
| -0.0103419 | 0.00414162 | 0.0125 |
| 0.35403 | 0.141954 | 0.0126 |
| -0.0402327 | 0.0161558 | 0.0128 |
| -0.010997 | 0.00442472 | 0.0129 |
| -0.00532038 | 0.00214693 | 0.0132 |
| 0.406475 | 0.164192 | 0.0133 |
| 0.243452 | 0.0983532 | 0.0133 |
| -0.00530822 | 0.00215011 | 0.0136 |
| 0.317735 | 0.12892 | 0.0137 |
| -0.00531608 | 0.00215858 | 0.0138 |
| 0.408303 | 0.166002 | 0.0139 |
| 0.264052 | 0.107397 | 0.0139 |
| -0.148657 | 0.0608386 | 0.0145 |
| -0.0110244 | 0.00451421 | 0.0146 |
| 0.0958865 | 0.0392878 | 0.0147 |
| 0.0998881 | 0.0411028 | 0.0151 |
| -0.0307839 | 0.0127152 | 0.0155 |
| -0.00478598 | 0.00200775 | 0.0171 |
| -0.0229539 | 0.0096888 | 0.0178 |
| 0.14576 | 0.0617836 | 0.0183 |
| 0.296991 | 0.126655 | 0.019 |
| -0.125182 | 0.0540799 | 0.0206 |
| -0.173015 | 0.0751238 | 0.0213 |
| 0.11985 | 0.0522982 | 0.0219 |
| -0.0207769 | 0.00907995 | 0.0221 |
| 0.396608 | 0.173452 | 0.0222 |

|  |  |  |
| --- | --- | --- |
| -0.0326895 | 0.0144029 | 0.0232 |
| -0.163267 | 0.0720241 | 0.0234 |
| -0.0229741 | 0.010241 | 0.0249 |
| -0.146153 | 0.0653927 | 0.0254 |
| -0.125126 | 0.056264 | 0.0262 |
| -0.292397 | 0.131606 | 0.0263 |
| 0.398937 | 0.181259 | 0.0277 |
| 0.124603 | 0.0566894 | 0.0279 |
| -0.293074 | 0.133478 | 0.0281 |
| 0.218318 | 0.0999588 | 0.029 |
| 0.267616 | 0.123213 | 0.0299 |
| 0.66772 | 0.308007 | 0.0302 |
| 0.342901 | 0.159665 | 0.0317 |
| -0.165678 | 0.0773388 | 0.0322 |
| -0.0211248 | 0.00989469 | 0.0328 |
| -0.00814583 | 0.00384487 | 0.0341 |
| -0.305647 | 0.144423 | 0.0343 |
| 0.188969 | 0.0893092 | 0.0344 |
| 0.480606 | 0.227569 | 0.0347 |
| 0.334418 | 0.159158 | 0.0356 |
| -0.228944 | 0.109369 | 0.0363 |
| -0.105879 | 0.0507571 | 0.037 |
| -0.0745321 | 0.0360928 | 0.0389 |
| 0.0138852 | 0.00676318 | 0.0401 |
| 0.343505 | 0.168165 | 0.0411 |
| -0.169996 | 0.0830315 | 0.0406 |
| -0.369911 | 0.181262 | 0.0413 |
| -0.0113244 | 0.00560352 | 0.0433 |
| 0.122543 | 0.0605321 | 0.0429 |
| -0.0584819 | 0.0289486 | 0.0434 |
| 0.211772 | 0.10506 | 0.0438 |
| 0.0901762 | 0.0449164 | 0.0447 |
| -0.100468 | 0.050148 | 0.0451 |
| -0.0207531 | 0.0103962 | 0.0459 |
| -0.13908 | 0.0697061 | 0.046 |
| 0.0668328 | 0.0335464 | 0.0463 |
| -0.0198046 | 0.00994492 | 0.0464 |
| -0.146864 | 0.0737211 | 0.0464 |
| -0.040477 | 0.0203826 | 0.047 |
| -0.0133386 | 0.00673872 | 0.0478 |
| 0.175907 | 0.0891422 | 0.0485 |
| -0.33145 | 0.169095 | 0.05 |
| 0.0884219 | 0.0450816 | 0.0498 |
| 0.0738177 | 0.0377488 | 0.0505 |
| 0.516614 | 0.264286 | 0.0506 |
| -0.165513 | 0.0848184 | 0.051 |
| -0.172398 | 0.0884889 | 0.0514 |
| 0.0809895 | 0.0415528 | 0.0513 |

|  |  |  |
| --- | --- | --- |
| -0.114502 | 0.0588864 | 0.0518 |
| 0.238321 | 0.12335 | 0.0534 |
| -0.129613 | 0.0671902 | 0.0537 |
| -0.20606 | 0.107238 | 0.0547 |
| -0.0152118 | 0.00794296 | 0.0555 |
| -0.112199 | 0.0585731 | 0.0554 |
| -0.070965 | 0.037133 | 0.056 |
| -0.191218 | 0.100507 | 0.0571 |
| -0.0231112 | 0.0121652 | 0.0575 |
| -0.00704839 | 0.00372609 | 0.0585 |
| 0.0947535 | 0.050711 | 0.0617 |
| 0.0950712 | 0.0508244 | 0.0614 |
| -0.019906 | 0.0106428 | 0.0614 |
| 0.10238 | 0.0548842 | 0.0621 |
| 0.0358849 | 0.019272 | 0.0626 |
| 0.0517051 | 0.0277172 | 0.0621 |
| 0.207866 | 0.111741 | 0.0629 |
| 0.0944037 | 0.0510111 | 0.0642 |
| 0.04699 | 0.0253956 | 0.0643 |
| 0.0997382 | 0.0540582 | 0.065 |
| 0.0943271 | 0.0513474 | 0.0662 |
| -0.381573 | 0.208522 | 0.0673 |
| 0.149862 | 0.0818837 | 0.0672 |
| 0.0307968 | 0.0168039 | 0.0668 |
| -0.17543 | 0.0959973 | 0.0676 |
| 0.0923232 | 0.0508488 | 0.0694 |
| 0.0845942 | 0.0468059 | 0.0707 |
| -0.0259283 | 0.0143656 | 0.0711 |
| -0.349273 | 0.194505 | 0.0725 |
| 0.023623 | 0.0131991 | 0.0735 |
| 0.101893 | 0.0570899 | 0.0743 |
| -0.0328405 | 0.0183626 | 0.0737 |
| 0.00532644 | 0.00297858 | 0.0737 |
| -0.0795097 | 0.0445982 | 0.0746 |
| 0.0828927 | 0.046508 | 0.0747 |
| -0.0142557 | 0.00799064 | 0.0744 |
| 0.168515 | 0.0948343 | 0.0756 |
| -0.220656 | 0.123994 | 0.0751 |
| -0.092383 | 0.0520461 | 0.0759 |
| -0.0135925 | 0.0076825 | 0.0768 |
| -0.0336806 | 0.019037 | 0.0769 |
| 0.0967345 | 0.0548116 | 0.0776 |
| -0.0982409 | 0.0562644 | 0.0808 |
| 0.0877094 | 0.0503922 | 0.0818 |
| 0.0224164 | 0.0130055 | 0.0848 |
| -0.0269539 | 0.0157068 | 0.0862 |
| 0.0805694 | 0.0470168 | 0.0866 |
| 0.257656 | 0.150638 | 0.0872 |

|  |  |  |
| --- | --- | --- |
| -0.110412 | 0.0646827 | 0.0878 |
| 0.223108 | 0.130824 | 0.0881 |
| 0.0396888 | 0.0232799 | 0.0882 |
| -0.0735909 | 0.0433916 | 0.0899 |
| -0.0600184 | 0.0356927 | 0.0927 |
| 0.00761541 | 0.00453538 | 0.0931 |
| 0.0875488 | 0.0526519 | 0.0964 |
| 0.249861 | 0.150819 | 0.0976 |
| -0.124181 | 0.0748846 | 0.0973 |
| 0.0884237 | 0.0535889 | 0.0989 |
| -0.0585252 | 0.0356331 | 0.1 |
| 0.13902 | 0.084812 | 0.101 |
| 0.0139045 | 0.00852132 | 0.103 |
| -0.0266988 | 0.0163656 | 0.103 |
| 0.183631 | 0.112546 | 0.103 |
| -0.216745 | 0.133229 | 0.104 |
| 0.110401 | 0.0686783 | 0.108 |
| 0.208997 | 0.130321 | 0.109 |
| -0.00644293 | 0.00402646 | 0.11 |
| -0.25153 | 0.157561 | 0.11 |
| -0.083647 | 0.0529336 | 0.114 |
| -0.0924273 | 0.0588503 | 0.116 |
| -0.246325 | 0.157151 | 0.117 |
| 0.212758 | 0.135975 | 0.118 |
| 0.115817 | 0.0741855 | 0.118 |
| -0.0331632 | 0.0212502 | 0.119 |
| -0.0912321 | 0.0586521 | 0.12 |
| -0.0934436 | 0.0602026 | 0.121 |
| 0.21328 | 0.137369 | 0.121 |
| 0.103083 | 0.066577 | 0.122 |
| -0.00356516 | 0.00230584 | 0.122 |
| -0.151567 | 0.0980676 | 0.122 |
| 0.0593211 | 0.0383386 | 0.122 |
| 0.448194 | 0.291121 | 0.124 |
| -0.110883 | 0.0719973 | 0.124 |
| 0.192308 | 0.125053 | 0.124 |
| 0.121051 | 0.07906 | 0.126 |
| -0.0756063 | 0.0495441 | 0.127 |
| -0.014705 | 0.0096372 | 0.127 |
| 0.0472673 | 0.0309819 | 0.127 |
| -0.185327 | 0.121774 | 0.128 |
| -0.132514 | 0.0869377 | 0.127 |
| -0.256352 | 0.168261 | 0.128 |
| 0.0940756 | 0.0622381 | 0.131 |
| 0.137187 | 0.0908714 | 0.131 |
| 0.44951 | 0.298648 | 0.132 |
| 0.0607501 | 0.0403564 | 0.132 |
| -0.00349175 | 0.00231599 | 0.132 |

|  |  |  |
| --- | --- | --- |
| 0.136562 | 0.0905807 | 0.132 |
| -0.0870081 | 0.0579741 | 0.133 |
| -0.20207 | 0.134773 | 0.134 |
| 0.0159643 | 0.0106694 | 0.135 |
| -0.0312296 | 0.0209112 | 0.135 |
| -0.0320015 | 0.0214801 | 0.136 |
| -0.0858302 | 0.0576925 | 0.137 |
| 0.174789 | 0.117881 | 0.138 |
| -0.00341099 | 0.00230475 | 0.139 |
| -0.0829683 | 0.0560035 | 0.138 |
| 0.0237168 | 0.0160196 | 0.139 |
| -0.0850234 | 0.0576493 | 0.14 |
| 0.0717216 | 0.0486186 | 0.14 |
| 0.146182 | 0.0995594 | 0.142 |
| 0.204174 | 0.138979 | 0.142 |
| -0.314834 | 0.214299 | 0.142 |
| 0.0721223 | 0.0490866 | 0.142 |
| -0.112593 | 0.077195 | 0.145 |
| -0.0910528 | 0.0624734 | 0.145 |
| 0.0729462 | 0.0500567 | 0.145 |
| -0.272354 | 0.187307 | 0.146 |
| 0.029104 | 0.0200371 | 0.146 |
| 0.367023 | 0.252539 | 0.146 |
| -0.145559 | 0.100403 | 0.147 |
| -0.0125612 | 0.00867374 | 0.148 |
| -0.309485 | 0.214908 | 0.15 |
| 0.215237 | 0.149767 | 0.151 |
| 0.0138841 | 0.00967361 | 0.151 |
| -0.11074 | 0.0771842 | 0.151 |
| -0.13712 | 0.0958424 | 0.153 |
| -0.0839241 | 0.0586213 | 0.152 |
| 0.0541599 | 0.037876 | 0.153 |
| 0.21473 | 0.150587 | 0.154 |
| 0.059235 | 0.0415189 | 0.154 |
| -0.00480096 | 0.00336658 | 0.154 |
| -0.216452 | 0.151836 | 0.154 |
| 0.499546 | 0.351028 | 0.155 |
| -0.0872825 | 0.0613949 | 0.155 |
| 0.0406013 | 0.0286249 | 0.156 |
| -0.0516329 | 0.0364818 | 0.157 |
| -0.0460706 | 0.0325495 | 0.157 |
| -0.156565 | 0.110418 | 0.156 |
| 0.0065412 | 0.00461479 | 0.156 |
| -0.0831963 | 0.0588218 | 0.157 |
| -0.169138 | 0.119713 | 0.158 |
| 0.209744 | 0.14854 | 0.158 |
| -0.0235641 | 0.0166942 | 0.158 |
| 0.378319 | 0.267974 | 0.158 |

|  |  |  |
| --- | --- | --- |
| -0.245057 | 0.174217 | 0.16 |
| 0.00677392 | 0.00481075 | 0.159 |
| -0.0861186 | 0.0612679 | 0.16 |
| -0.125494 | 0.0893861 | 0.16 |
| -0.0805509 | 0.0573579 | 0.16 |
| -0.0963841 | 0.0687738 | 0.161 |
| 0.0630828 | 0.0451039 | 0.162 |
| 0.0920484 | 0.0658464 | 0.162 |
| -0.0381872 | 0.0272714 | 0.161 |
| 0.245861 | 0.175824 | 0.162 |
| 0.0212679 | 0.0152529 | 0.163 |
| -0.100296 | 0.0719881 | 0.164 |
| -0.0313514 | 0.022525 | 0.164 |
| -0.088008 | 0.0631345 | 0.163 |
| 0.10523 | 0.0757101 | 0.165 |
| 0.00927182 | 0.0066748 | 0.165 |
| -0.407057 | 0.294766 | 0.167 |
| -0.011798 | 0.00854621 | 0.167 |
| 0.0125214 | 0.00909648 | 0.169 |
| -0.127611 | 0.0926643 | 0.168 |
| -0.0113252 | 0.00826788 | 0.171 |
| -0.0502526 | 0.0367882 | 0.172 |
| 0.366314 | 0.267988 | 0.172 |
| -0.0548434 | 0.0406257 | 0.177 |
| -0.0480884 | 0.0356663 | 0.178 |
| 0.0601385 | 0.044739 | 0.179 |
| 0.276448 | 0.205797 | 0.179 |
| -0.0875609 | 0.0654629 | 0.181 |
| -0.0797574 | 0.0598098 | 0.182 |
| 0.356598 | 0.26795 | 0.183 |
| -0.0890828 | 0.0671152 | 0.184 |
| -0.284044 | 0.213892 | 0.184 |
| -0.0408486 | 0.0308427 | 0.185 |
| -0.215008 | 0.162488 | 0.186 |
| -0.0700556 | 0.0530639 | 0.187 |
| -0.00535634 | 0.00406556 | 0.188 |
| -0.0694916 | 0.0528398 | 0.188 |
| 0.1347 | 0.102526 | 0.189 |
| 0.155406 | 0.118475 | 0.19 |
| 0.0358778 | 0.0273606 | 0.19 |
| -0.00735233 | 0.00562077 | 0.191 |
| -0.017012 | 0.0130207 | 0.191 |
| -0.0762175 | 0.0584428 | 0.192 |
| 0.0219596 | 0.0168961 | 0.194 |
| -0.178091 | 0.136885 | 0.193 |
| 0.110577 | 0.0852962 | 0.195 |
| -0.0103024 | 0.00795059 | 0.195 |
| -0.0556339 | 0.0429088 | 0.195 |

|  |  |  |
| --- | --- | --- |
| -0.11375 | 0.0880195 | 0.196 |
| -0.100613 | 0.0777625 | 0.196 |
| -0.0277828 | 0.0214827 | 0.196 |
| -0.00874072 | 0.00678109 | 0.197 |
| -0.0092152 | 0.00715092 | 0.198 |
| -0.219086 | 0.17012 | 0.198 |
| 0.211495 | 0.165102 | 0.2 |
| 0.326019 | 0.256342 | 0.203 |
| -0.0562245 | 0.0441826 | 0.203 |
| -0.0927326 | 0.0729277 | 0.204 |
| 0.00613105 | 0.00482604 | 0.204 |
| -0.088341 | 0.0694752 | 0.204 |
| -0.00905974 | 0.00714525 | 0.205 |
| -0.16031 | 0.126593 | 0.205 |
| -0.0862975 | 0.0680962 | 0.205 |
| 0.0687681 | 0.0546406 | 0.208 |
| -0.0100706 | 0.00802897 | 0.21 |
| -0.138878 | 0.111095 | 0.211 |
| -0.211154 | 0.169382 | 0.213 |
| -0.0777064 | 0.0624815 | 0.214 |
| 0.0995629 | 0.0801598 | 0.214 |
| 0.0465457 | 0.0375681 | 0.215 |
| 0.0526928 | 0.042637 | 0.217 |
| 0.0126602 | 0.0102422 | 0.216 |
| 0.0117927 | 0.00957638 | 0.218 |
| 0.0122749 | 0.00998205 | 0.219 |
| 0.00400948 | 0.00326884 | 0.22 |
| -0.0935634 | 0.076211 | 0.22 |
| 0.574346 | 0.467473 | 0.219 |
| 0.0130804 | 0.0106889 | 0.221 |
| -0.00936475 | 0.00764468 | 0.221 |
| -0.373428 | 0.305384 | 0.221 |
| 0.00337517 | 0.00276 | 0.221 |
| -0.373159 | 0.305748 | 0.222 |
| 0.0126822 | 0.0103857 | 0.222 |
| -0.374169 | 0.307602 | 0.224 |
| -0.013454 | 0.0110474 | 0.223 |
| -0.371946 | 0.305791 | 0.224 |
| 0.162042 | 0.133507 | 0.225 |
| 0.0273101 | 0.0224869 | 0.225 |
| 0.0908037 | 0.0747021 | 0.224 |
| 0.171857 | 0.141626 | 0.225 |
| 0.0124008 | 0.0102309 | 0.225 |
| -0.0189809 | 0.0156764 | 0.226 |
| -0.125348 | 0.103845 | 0.227 |
| -0.0468952 | 0.0388472 | 0.227 |
| -0.0193616 | 0.0160549 | 0.228 |
| -0.0342506 | 0.0284113 | 0.228 |

|  |  |  |
| --- | --- | --- |
| -0.366625 | 0.305864 | 0.231 |
| 0.0585295 | 0.0487524 | 0.23 |
| 0.0124479 | 0.0103752 | 0.23 |
| 0.0397015 | 0.0330567 | 0.23 |
| -0.36759 | 0.306708 | 0.231 |
| -0.0296513 | 0.024834 | 0.232 |
| -0.0801329 | 0.0671149 | 0.232 |
| -0.0826225 | 0.0692471 | 0.233 |
| 0.0274136 | 0.0229748 | 0.233 |
| -0.0537444 | 0.0451232 | 0.234 |
| 0.0660992 | 0.0555976 | 0.234 |
| -0.36301 | 0.305952 | 0.235 |
| -0.180214 | 0.151854 | 0.235 |
| -0.0106341 | 0.00896547 | 0.236 |
| 0.0793337 | 0.0671148 | 0.237 |
| 0.0304741 | 0.0257767 | 0.237 |
| -0.0160313 | 0.0135687 | 0.237 |
| -0.0625839 | 0.0530479 | 0.238 |
| 0.00322791 | 0.00273264 | 0.238 |
| 0.0574254 | 0.048653 | 0.238 |
| -0.0128407 | 0.010922 | 0.24 |
| -0.265501 | 0.226075 | 0.24 |
| -0.213115 | 0.181224 | 0.24 |
| -0.00885576 | 0.00754665 | 0.241 |
| -0.0925058 | 0.0790875 | 0.242 |
| -0.0865002 | 0.0740309 | 0.243 |
| -0.00808119 | 0.00693602 | 0.244 |
| 0.0911366 | 0.0781513 | 0.244 |
| 0.124281 | 0.106595 | 0.244 |
| -0.110059 | 0.0947379 | 0.245 |
| -0.298404 | 0.257009 | 0.246 |
| -0.0279687 | 0.0242388 | 0.249 |
| 0.160454 | 0.139038 | 0.248 |
| 0.130551 | 0.11347 | 0.25 |
| 0.104593 | 0.0909496 | 0.25 |
| -0.123241 | 0.10732 | 0.251 |
| -0.0263748 | 0.0229829 | 0.251 |
| 0.0241526 | 0.0210303 | 0.251 |
| 0.30394 | 0.265202 | 0.252 |
| 0.130027 | 0.113568 | 0.252 |
| 0.121833 | 0.106395 | 0.252 |
| -0.00259035 | 0.00226526 | 0.253 |
| -0.0275338 | 0.0241413 | 0.254 |
| -0.0276798 | 0.0242622 | 0.254 |
| -0.10811 | 0.0952382 | 0.256 |
| 0.0316728 | 0.0280985 | 0.26 |
| 0.0824216 | 0.0731948 | 0.26 |
| 0.184387 | 0.163934 | 0.261 |

|  |  |  |
| --- | --- | --- |
| 0.0347304 | 0.0308963 | 0.261 |
| 0.0331269 | 0.0295153 | 0.262 |
| -0.0873086 | 0.0777279 | 0.261 |
| -0.037628 | 0.0334955 | 0.261 |
| -0.0140294 | 0.0125008 | 0.262 |
| 0.123246 | 0.109882 | 0.262 |
| -0.00749004 | 0.00667914 | 0.262 |
| -0.103667 | 0.0927331 | 0.264 |
| 0.12698 | 0.113892 | 0.265 |
| -0.0175692 | 0.01576 | 0.265 |
| -0.112904 | 0.101254 | 0.265 |
| 0.120058 | 0.1075 | 0.264 |
| 0.198243 | 0.177568 | 0.264 |
| -0.103443 | 0.0927395 | 0.265 |
| 0.257087 | 0.231046 | 0.266 |
| 0.0583787 | 0.0525362 | 0.266 |
| 0.292272 | 0.263115 | 0.267 |
| -0.0306469 | 0.027575 | 0.266 |
| 0.0309241 | 0.0278687 | 0.267 |
| 0.0414498 | 0.037475 | 0.269 |
| 0.0467973 | 0.0424247 | 0.27 |
| -0.0847382 | 0.0768605 | 0.27 |
| -0.0814485 | 0.0738674 | 0.27 |
| 0.0349036 | 0.0316607 | 0.27 |
| -0.0999546 | 0.0907343 | 0.271 |
| -0.0177627 | 0.016142 | 0.271 |
| 0.0132441 | 0.0120809 | 0.273 |
| -0.0102449 | 0.00932878 | 0.272 |
| -0.0830877 | 0.0757483 | 0.273 |
| -0.0492118 | 0.044987 | 0.274 |
| -0.0698426 | 0.0639733 | 0.275 |
| 0.212284 | 0.194623 | 0.275 |
| 0.0655235 | 0.0600364 | 0.275 |
| -0.103903 | 0.0954782 | 0.276 |
| 0.165293 | 0.152098 | 0.277 |
| -0.0220204 | 0.0203388 | 0.279 |
| 0.005417 | 0.00500433 | 0.279 |
| 0.0324477 | 0.0299524 | 0.279 |
| 0.204706 | 0.189785 | 0.281 |
| -0.054437 | 0.0505426 | 0.281 |
| 0.05779 | 0.0537089 | 0.282 |
| -0.12227 | 0.113631 | 0.282 |
| -0.216442 | 0.20165 | 0.283 |
| 0.090335 | 0.0841773 | 0.283 |
| -0.147244 | 0.137389 | 0.284 |
| 0.0112778 | 0.0105453 | 0.285 |
| 0.0103505 | 0.00966809 | 0.284 |
| -0.00716578 | 0.00671746 | 0.286 |

|  |  |  |
| --- | --- | --- |
| 0.0196922 | 0.0185189 | 0.288 |
| 0.209074 | 0.196997 | 0.289 |
| -0.0212023 | 0.0199997 | 0.289 |
| -0.0312618 | 0.0294414 | 0.288 |
| 0.0105741 | 0.00998219 | 0.289 |
| -0.123351 | 0.116746 | 0.291 |
| -0.324424 | 0.307227 | 0.291 |
| -0.00744862 | 0.00704487 | 0.29 |
| 0.0275376 | 0.0260483 | 0.29 |
| -0.007507 | 0.00710539 | 0.291 |
| -0.222874 | 0.211564 | 0.292 |
| -0.109256 | 0.104145 | 0.294 |
| -0.00745702 | 0.00710843 | 0.294 |
| 0.0852482 | 0.0813851 | 0.295 |
| 0.146113 | 0.139611 | 0.295 |
| 0.240362 | 0.231278 | 0.299 |
| 0.0321081 | 0.0309106 | 0.299 |
| -0.00737345 | 0.00710261 | 0.299 |
| -0.00778696 | 0.00751753 | 0.3 |
| -0.0597258 | 0.0576967 | 0.301 |
| -0.00699743 | 0.00676316 | 0.301 |
| 0.0706685 | 0.0683539 | 0.301 |
| -0.00733883 | 0.00710264 | 0.301 |
| 0.221441 | 0.21441 | 0.302 |
| -0.00732534 | 0.00710093 | 0.302 |
| -0.00735251 | 0.00711902 | 0.302 |
| -0.290559 | 0.282448 | 0.304 |
| -0.128092 | 0.124427 | 0.303 |
| 0.0539328 | 0.0523662 | 0.303 |
| -0.00730986 | 0.00710017 | 0.303 |
| -0.0705408 | 0.0686933 | 0.304 |
| -0.295832 | 0.287693 | 0.304 |
| -0.00729093 | 0.00710065 | 0.305 |
| 0.165367 | 0.161185 | 0.305 |
| 0.100975 | 0.0984899 | 0.305 |
| -0.124761 | 0.121694 | 0.305 |
| -0.0939055 | 0.0915025 | 0.305 |
| -0.0138633 | 0.0135599 | 0.307 |
| 0.240251 | 0.234868 | 0.306 |
| -0.00725924 | 0.00710031 | 0.307 |
| 0.0728435 | 0.0717133 | 0.31 |
| 0.0991902 | 0.0975202 | 0.309 |
| -0.0341887 | 0.0336302 | 0.309 |
| 0.0613819 | 0.0603638 | 0.309 |
| 0.0745149 | 0.0733216 | 0.309 |
| -0.045598 | 0.0452573 | 0.314 |
| 0.148473 | 0.147213 | 0.313 |
| -0.00710674 | 0.00705274 | 0.314 |

|  |  |  |
| --- | --- | --- |
| 0.0272395 | 0.0271128 | 0.315 |
| 0.118623 | 0.117921 | 0.314 |
| -0.0141082 | 0.0140314 | 0.315 |
| -0.00710728 | 0.00706683 | 0.315 |
| -0.0549002 | 0.0547059 | 0.316 |
| 0.0797093 | 0.0793768 | 0.315 |
| 0.122459 | 0.122376 | 0.317 |
| -0.00711683 | 0.00710049 | 0.316 |
| -0.0919242 | 0.0919145 | 0.317 |
| 0.0819441 | 0.0820204 | 0.318 |
| -0.00707424 | 0.0070851 | 0.318 |
| -0.148319 | 0.148377 | 0.317 |
| 0.0662718 | 0.0664709 | 0.319 |
| -0.0139038 | 0.0139788 | 0.32 |
| 0.0887679 | 0.0893378 | 0.32 |
| 0.00973416 | 0.00981241 | 0.321 |
| -0.00415186 | 0.00418063 | 0.321 |
| -0.007036 | 0.00708519 | 0.321 |
| -0.135002 | 0.136228 | 0.322 |
| -0.109085 | 0.110065 | 0.322 |
| -0.00703366 | 0.00710011 | 0.322 |
| 0.00982487 | 0.00993921 | 0.323 |
| -0.107961 | 0.109185 | 0.323 |
| 0.140443 | 0.142026 | 0.323 |
| -0.0484998 | 0.0490411 | 0.323 |
| -0.00661447 | 0.00670266 | 0.324 |
| -0.0976759 | 0.0990403 | 0.324 |
| -0.00659498 | 0.00669397 | 0.325 |
| -0.0069719 | 0.00707943 | 0.325 |
| 0.0906014 | 0.092158 | 0.326 |
| -0.0192734 | 0.0196168 | 0.326 |
| -0.116687 | 0.118724 | 0.326 |
| -0.10814 | 0.110053 | 0.326 |
| 0.132127 | 0.134831 | 0.327 |
| 0.00379694 | 0.00387536 | 0.327 |
| -0.150998 | 0.154074 | 0.327 |
| -0.0449228 | 0.0458278 | 0.327 |
| -0.00930767 | 0.0095005 | 0.327 |
| -0.00694355 | 0.00709129 | 0.327 |
| -0.00542396 | 0.00555125 | 0.329 |
| -0.0350034 | 0.0359032 | 0.33 |
| 0.0539162 | 0.0552481 | 0.329 |
| -0.0626234 | 0.0642462 | 0.33 |
| 0.0544676 | 0.0559317 | 0.33 |
| 0.161698 | 0.166566 | 0.332 |
| 0.0363057 | 0.0374456 | 0.332 |
| -0.0418298 | 0.0431269 | 0.332 |
| 0.136176 | 0.140559 | 0.333 |

|  |  |  |
| --- | --- | --- |
| -0.00685857 | 0.00709315 | 0.334 |
| -0.0591845 | 0.0613422 | 0.335 |
| -0.113057 | 0.117333 | 0.335 |
| 0.096168 | 0.100301 | 0.338 |
| -0.327203 | 0.340947 | 0.337 |
| 0.0287568 | 0.0299988 | 0.338 |
| 0.0093942 | 0.00982238 | 0.339 |
| 0.0576473 | 0.0602752 | 0.339 |
| 0.0694569 | 0.0728417 | 0.34 |
| -0.162103 | 0.170097 | 0.341 |
| -0.151336 | 0.159245 | 0.342 |
| -0.161795 | 0.170116 | 0.342 |
| 0.0495397 | 0.0521652 | 0.342 |
| -0.107076 | 0.112571 | 0.342 |
| -0.101609 | 0.107139 | 0.343 |
| -0.0067177 | 0.00708464 | 0.343 |
| -0.18112 | 0.191527 | 0.344 |
| 0.125976 | 0.133465 | 0.345 |
| -0.0194625 | 0.0206336 | 0.346 |
| 0.0159681 | 0.0169205 | 0.345 |
| -0.131588 | 0.139731 | 0.346 |
| -0.0722574 | 0.0769412 | 0.348 |
| -0.0213814 | 0.0228584 | 0.35 |
| -0.130434 | 0.139665 | 0.35 |
| 0.297186 | 0.318447 | 0.351 |
| 0.471173 | 0.505767 | 0.352 |
| -0.159011 | 0.170616 | 0.351 |
| 0.0986725 | 0.105948 | 0.352 |
| 0.102514 | 0.110026 | 0.351 |
| -0.0571758 | 0.0616522 | 0.354 |
| -0.159285 | 0.171737 | 0.354 |
| -0.115394 | 0.124581 | 0.354 |
| -0.0206723 | 0.022375 | 0.356 |
| -0.0318796 | 0.0345479 | 0.356 |
| 0.168353 | 0.182221 | 0.356 |
| 0.0730685 | 0.0791245 | 0.356 |
| -0.0268327 | 0.0290807 | 0.356 |
| -0.0824472 | 0.0892949 | 0.356 |
| -0.129667 | 0.140412 | 0.356 |
| 0.108086 | 0.117026 | 0.356 |
| 0.0091232 | 0.00990629 | 0.357 |
| -0.154287 | 0.16726 | 0.356 |
| 0.233019 | 0.254219 | 0.359 |
| 0.143921 | 0.157262 | 0.36 |
| 0.172484 | 0.18828 | 0.36 |
| 0.0595565 | 0.0651865 | 0.361 |
| 0.108346 | 0.118803 | 0.362 |
| 0.112104 | 0.123045 | 0.362 |

|  |  |  |
| --- | --- | --- |
| -0.0489236 | 0.0537481 | 0.363 |
| -0.0368522 | 0.040483 | 0.363 |
| 0.0442427 | 0.0485641 | 0.362 |
| -0.136866 | 0.150481 | 0.363 |
| -0.0834706 | 0.0917733 | 0.363 |
| 0.0770735 | 0.0849916 | 0.364 |
| -0.00646783 | 0.00713253 | 0.365 |
| 0.0113725 | 0.0125553 | 0.365 |
| 0.192611 | 0.213026 | 0.366 |
| 0.0249625 | 0.0275962 | 0.366 |
| -0.29818 | 0.3299 | 0.366 |
| 0.0832224 | 0.0924402 | 0.368 |
| 0.0106508 | 0.0118231 | 0.368 |
| -0.175364 | 0.1952 | 0.369 |
| -0.0656917 | 0.0731611 | 0.369 |
| 0.0459759 | 0.051417 | 0.371 |
| 0.0363085 | 0.0405354 | 0.37 |
| -0.082639 | 0.0925119 | 0.372 |
| 0.0611514 | 0.0684179 | 0.371 |
| 0.0835707 | 0.0936394 | 0.372 |
| -0.145111 | 0.163008 | 0.373 |
| -0.0488054 | 0.0548573 | 0.374 |
| 0.178774 | 0.201097 | 0.374 |
| 0.0449392 | 0.0506545 | 0.375 |
| -0.109349 | 0.123622 | 0.376 |
| -0.0201722 | 0.022839 | 0.377 |
| 0.040876 | 0.0462498 | 0.377 |
| 0.0319696 | 0.0362655 | 0.378 |
| -0.146126 | 0.166706 | 0.381 |
| 0.0699286 | 0.0797073 | 0.38 |
| -0.0336472 | 0.038355 | 0.38 |
| -0.014837 | 0.0169205 | 0.381 |
| 0.0814777 | 0.0930142 | 0.381 |
| -0.103502 | 0.118292 | 0.382 |
| 0.06289 | 0.071922 | 0.382 |
| 0.150492 | 0.172155 | 0.382 |
| -0.00671836 | 0.0076807 | 0.382 |
| 0.105013 | 0.120267 | 0.383 |
| 0.0913985 | 0.104781 | 0.383 |
| 0.0641034 | 0.0734147 | 0.383 |
| -0.0145142 | 0.0166151 | 0.382 |
| 0.00518926 | 0.00595642 | 0.384 |
| -0.0718 | 0.0824343 | 0.384 |
| -0.0110923 | 0.0127953 | 0.386 |
| 0.0622755 | 0.0719344 | 0.387 |
| -0.0161918 | 0.0187523 | 0.388 |
| 0.00778947 | 0.00903489 | 0.389 |
| -0.0177957 | 0.0206226 | 0.388 |

|  |  |  |
| --- | --- | --- |
| -0.0784101 | 0.0908973 | 0.388 |
| -0.0782837 | 0.0909009 | 0.389 |
| 0.0270018 | 0.0314443 | 0.39 |
| -0.0479962 | 0.0559772 | 0.391 |
| -0.0393052 | 0.0458999 | 0.392 |
| 0.0890384 | 0.103862 | 0.391 |
| -0.128613 | 0.150571 | 0.393 |
| -0.082262 | 0.0964466 | 0.394 |
| -0.0783759 | 0.0920197 | 0.394 |
| 0.0158047 | 0.0186019 | 0.396 |
| -0.0593839 | 0.0701737 | 0.397 |
| -0.0713589 | 0.084177 | 0.397 |
| -0.00479573 | 0.00566316 | 0.397 |
| -0.0267527 | 0.0315725 | 0.397 |
| -0.00852233 | 0.0100727 | 0.398 |
| -0.0617251 | 0.0733527 | 0.4 |
| 0.133457 | 0.158549 | 0.4 |
| 0.175723 | 0.209614 | 0.402 |
| -0.0315545 | 0.0376419 | 0.402 |
| 0.0861376 | 0.102774 | 0.402 |
| -0.0859355 | 0.102618 | 0.402 |
| 0.0997675 | 0.11927 | 0.403 |
| -0.14815 | 0.177369 | 0.404 |
| 0.0707515 | 0.0847325 | 0.404 |
| -0.270116 | 0.324037 | 0.405 |
| -0.00610969 | 0.00734161 | 0.405 |
| -0.00584857 | 0.00703125 | 0.406 |
| 0.0280256 | 0.0338031 | 0.407 |
| -0.231191 | 0.279374 | 0.408 |
| 0.138528 | 0.167692 | 0.409 |
| 0.0593987 | 0.0719843 | 0.409 |
| 0.0835857 | 0.101374 | 0.41 |
| 0.0420732 | 0.0513341 | 0.412 |
| -0.00850321 | 0.0103728 | 0.412 |
| 0.0193318 | 0.0235583 | 0.412 |
| 0.0252133 | 0.0307525 | 0.412 |
| 0.0159188 | 0.0194222 | 0.412 |
| 0.0575542 | 0.0702084 | 0.412 |
| -0.123704 | 0.151486 | 0.414 |
| 0.0396552 | 0.0486216 | 0.415 |
| -0.0296958 | 0.0365004 | 0.416 |
| -0.0991034 | 0.121657 | 0.415 |
| -0.0526126 | 0.0647183 | 0.416 |
| -0.0591699 | 0.072948 | 0.417 |
| 0.05288 | 0.0654919 | 0.419 |
| 0.00793227 | 0.00980635 | 0.419 |
| -0.0189048 | 0.0233892 | 0.419 |
| -0.0816622 | 0.101062 | 0.419 |

|  |  |  |
| --- | --- | --- |
| 0.0962349 | 0.119346 | 0.42 |
| -0.0515397 | 0.0638955 | 0.42 |
| 0.166539 | 0.206953 | 0.421 |
| 0.0160458 | 0.0199271 | 0.421 |
| -0.0358117 | 0.0445659 | 0.422 |
| -0.0855944 | 0.10699 | 0.424 |
| -0.0318757 | 0.0398161 | 0.423 |
| -0.0694859 | 0.0868952 | 0.424 |
| 0.0469242 | 0.0589482 | 0.426 |
| -0.112126 | 0.140591 | 0.425 |
| -0.0129237 | 0.0162189 | 0.426 |
| 0.0400061 | 0.0503447 | 0.427 |
| -0.0192453 | 0.024193 | 0.426 |
| 0.107961 | 0.135704 | 0.426 |
| 0.0291577 | 0.0367501 | 0.428 |
| 0.128846 | 0.162551 | 0.428 |
| -0.0628921 | 0.0794342 | 0.429 |
| -0.0373337 | 0.047195 | 0.429 |
| -0.0406744 | 0.0516083 | 0.431 |
| 0.0326905 | 0.0414004 | 0.43 |
| 0.0410804 | 0.0520613 | 0.43 |
| -0.0398552 | 0.0506792 | 0.432 |
| 0.138531 | 0.175934 | 0.431 |
| -0.105927 | 0.135193 | 0.433 |
| 0.0642184 | 0.0822067 | 0.435 |
| -0.14661 | 0.187538 | 0.434 |
| 0.013134 | 0.0168083 | 0.435 |
| 0.0246983 | 0.0317102 | 0.436 |
| -0.0960685 | 0.123526 | 0.437 |
| -0.162117 | 0.208653 | 0.437 |
| -0.0502902 | 0.0646146 | 0.436 |
| -0.150989 | 0.194133 | 0.437 |
| 0.10418 | 0.134054 | 0.437 |
| -0.0583475 | 0.0752657 | 0.438 |
| -0.00545487 | 0.00705281 | 0.439 |
| 0.299187 | 0.387686 | 0.44 |
| 0.0745694 | 0.0965258 | 0.44 |
| -0.0771742 | 0.0999651 | 0.44 |
| -0.00957134 | 0.0124391 | 0.442 |
| -0.0304619 | 0.0396403 | 0.442 |
| -0.0544663 | 0.0709281 | 0.443 |
| -0.00479171 | 0.00624429 | 0.443 |
| 0.0736038 | 0.0957441 | 0.442 |
| -0.012856 | 0.0167702 | 0.443 |
| -0.116824 | 0.152406 | 0.443 |
| -0.088412 | 0.115359 | 0.443 |
| -0.012504 | 0.0163931 | 0.446 |
| 0.106846 | 0.140038 | 0.445 |

|  |  |  |
| --- | --- | --- |
| -0.0691082 | 0.0907571 | 0.446 |
| -0.100548 | 0.132301 | 0.447 |
| -0.097401 | 0.128004 | 0.447 |
| 0.0627608 | 0.0826351 | 0.448 |
| 0.0371776 | 0.0489554 | 0.448 |
| 0.056371 | 0.0744817 | 0.449 |
| 0.146965 | 0.194089 | 0.449 |
| 0.0501364 | 0.0664799 | 0.451 |
| 0.05534 | 0.073369 | 0.451 |
| -0.0280092 | 0.0372321 | 0.452 |
| 0.140602 | 0.187686 | 0.454 |
| 0.140584 | 0.187684 | 0.454 |
| 0.140144 | 0.187226 | 0.454 |
| 0.0144325 | 0.0192817 | 0.454 |
| 0.0433577 | 0.0579881 | 0.455 |
| 0.0660818 | 0.0883651 | 0.455 |
| -0.0377068 | 0.0504454 | 0.455 |
| -0.0118728 | 0.0158789 | 0.455 |
| -0.0460921 | 0.0619489 | 0.457 |
| 0.0659449 | 0.088683 | 0.457 |
| -0.163444 | 0.220413 | 0.458 |
| 0.0471282 | 0.0635963 | 0.459 |
| -0.00520155 | 0.00702402 | 0.459 |
| -0.041267 | 0.0557893 | 0.459 |
| 0.0964457 | 0.130737 | 0.461 |
| 0.0417739 | 0.0566078 | 0.461 |
| 0.134104 | 0.181925 | 0.461 |
| 0.010237 | 0.0138931 | 0.461 |
| 0.091404 | 0.124036 | 0.461 |
| -0.0326121 | 0.0444394 | 0.463 |
| 0.108109 | 0.147594 | 0.464 |
| -0.0624371 | 0.0852972 | 0.464 |
| -0.105298 | 0.144129 | 0.465 |
| 0.0358555 | 0.0490588 | 0.465 |
| 0.0641818 | 0.0881976 | 0.467 |
| -0.0322541 | 0.044586 | 0.469 |
| -0.0882386 | 0.122401 | 0.471 |
| -0.0960462 | 0.133122 | 0.471 |
| 0.0144489 | 0.0200949 | 0.472 |
| -0.0937068 | 0.130277 | 0.472 |
| 0.0681751 | 0.0948592 | 0.472 |
| -0.0431272 | 0.0599311 | 0.472 |
| -0.108193 | 0.150227 | 0.471 |
| -0.108011 | 0.150053 | 0.472 |
| -0.0788003 | 0.109589 | 0.472 |
| -0.010107 | 0.0140825 | 0.473 |
| 0.108936 | 0.152334 | 0.475 |
| 0.0323256 | 0.0452989 | 0.475 |

|  |  |  |
| --- | --- | --- |
| -0.11017 | 0.154278 | 0.475 |
| -0.0627039 | 0.0881525 | 0.477 |
| -0.130585 | 0.183367 | 0.476 |
| 0.199303 | 0.280159 | 0.477 |
| 0.0847818 | 0.119089 | 0.477 |
| -0.181542 | 0.255132 | 0.477 |
| -0.0564932 | 0.0794889 | 0.477 |
| 0.0120371 | 0.0169303 | 0.477 |
| -0.00503124 | 0.00707791 | 0.477 |
| 0.032871 | 0.0465563 | 0.48 |
| -0.136201 | 0.192946 | 0.48 |
| -0.0551492 | 0.0780364 | 0.48 |
| 0.13811 | 0.195893 | 0.481 |
| -0.00911532 | 0.0129259 | 0.481 |
| 0.0830912 | 0.117908 | 0.481 |
| -0.0347833 | 0.0494615 | 0.482 |
| -0.107468 | 0.153388 | 0.484 |
| -0.108383 | 0.154751 | 0.484 |
| -0.0804909 | 0.114732 | 0.483 |
| -0.0513328 | 0.073143 | 0.483 |
| 0.0411414 | 0.0588642 | 0.485 |
| -0.0345582 | 0.0493781 | 0.484 |
| 0.136158 | 0.195321 | 0.486 |
| -0.163927 | 0.234801 | 0.485 |
| 0.0266694 | 0.0382973 | 0.486 |
| -0.109737 | 0.157924 | 0.487 |
| -0.0705168 | 0.101539 | 0.487 |
| 0.00388967 | 0.00559911 | 0.487 |
| -0.100215 | 0.144296 | 0.487 |
| 0.0500216 | 0.0720426 | 0.487 |
| 0.0323125 | 0.0465786 | 0.488 |
| -0.109734 | 0.158053 | 0.488 |
| 0.135411 | 0.19573 | 0.489 |
| 0.0160541 | 0.0232156 | 0.489 |
| -0.00489797 | 0.00707704 | 0.489 |
| 0.131142 | 0.189938 | 0.49 |
| 0.0617341 | 0.0897734 | 0.492 |
| -0.0324454 | 0.0471624 | 0.491 |
| 0.0141635 | 0.0205683 | 0.491 |
| -0.00846353 | 0.0123445 | 0.493 |
| 0.0873726 | 0.127507 | 0.493 |
| -0.0833513 | 0.121677 | 0.493 |
| 0.0688112 | 0.100711 | 0.494 |
| -0.0217836 | 0.0320298 | 0.496 |
| 0.0220025 | 0.0323543 | 0.496 |
| -0.0408262 | 0.0600564 | 0.497 |
| 0.128971 | 0.190478 | 0.498 |
| -0.149323 | 0.220364 | 0.498 |

|  |  |  |
| --- | --- | --- |
| -0.00294347 | 0.00435789 | 0.499 |
| -0.00176521 | 0.00261117 | 0.499 |
| -0.0391153 | 0.0580517 | 0.5 |
| -0.0688163 | 0.102592 | 0.502 |
| 0.127782 | 0.190742 | 0.503 |
| -0.00804671 | 0.0120119 | 0.503 |
| 0.019632 | 0.029357 | 0.504 |
| 0.0591198 | 0.088537 | 0.504 |
| 0.144676 | 0.217884 | 0.507 |
| -0.0262398 | 0.0394387 | 0.506 |
| 0.193676 | 0.292418 | 0.508 |
| -0.0142965 | 0.0215467 | 0.507 |
| -0.122151 | 0.184479 | 0.508 |
| -0.0093937 | 0.0142067 | 0.508 |
| -0.0206472 | 0.0311866 | 0.508 |
| -0.0147814 | 0.0223723 | 0.509 |
| 0.100673 | 0.152471 | 0.509 |
| 0.0155759 | 0.0236474 | 0.51 |
| -0.0148172 | 0.0224879 | 0.51 |
| 0.0939835 | 0.142601 | 0.51 |
| 0.00981029 | 0.0149289 | 0.511 |
| -0.00466461 | 0.00708469 | 0.51 |
| 0.0592807 | 0.0902849 | 0.511 |
| 0.0155588 | 0.0237777 | 0.513 |
| -0.0175473 | 0.0268157 | 0.513 |
| -0.00925155 | 0.0141337 | 0.513 |
| -0.0267036 | 0.0410761 | 0.516 |
| -0.168081 | 0.258513 | 0.516 |
| 0.209455 | 0.321935 | 0.515 |
| 0.0838249 | 0.12921 | 0.517 |
| -0.0310952 | 0.0481726 | 0.519 |
| -0.115394 | 0.178857 | 0.519 |
| -0.0170367 | 0.0264555 | 0.52 |
| -0.0796997 | 0.12407 | 0.521 |
| 0.0563533 | 0.0878408 | 0.521 |
| 0.0144057 | 0.0225123 | 0.522 |
| 0.0951433 | 0.148681 | 0.522 |
| 0.0170972 | 0.0267767 | 0.523 |
| 0.0394037 | 0.0619076 | 0.524 |
| 0.0242528 | 0.0380765 | 0.524 |
| -0.0390691 | 0.061375 | 0.524 |
| 0.0717758 | 0.112918 | 0.525 |
| -0.0120279 | 0.0189044 | 0.525 |
| -0.0163386 | 0.0257302 | 0.525 |
| -0.0033811 | 0.00535476 | 0.528 |
| 0.140508 | 0.222773 | 0.528 |
| 0.0490166 | 0.0777105 | 0.528 |
| -0.0319137 | 0.0510412 | 0.532 |

|  |  |  |
| --- | --- | --- |
| 0.0828835 | 0.132517 | 0.532 |
| -0.0132992 | 0.0212766 | 0.532 |
| 0.0520077 | 0.083086 | 0.531 |
| -0.0944463 | 0.150786 | 0.531 |
| -0.00540607 | 0.00866251 | 0.533 |
| 0.0689716 | 0.110604 | 0.533 |
| -0.11384 | 0.1828 | 0.533 |
| -0.0500096 | 0.0804175 | 0.534 |
| -0.0312207 | 0.0501965 | 0.534 |
| -0.0229902 | 0.037214 | 0.537 |
| -0.0784912 | 0.126989 | 0.537 |
| 0.0885702 | 0.143193 | 0.536 |
| 0.0431001 | 0.0697019 | 0.536 |
| 0.0831097 | 0.134481 | 0.537 |
| -0.0520806 | 0.0845546 | 0.538 |
| -0.030059 | 0.0487505 | 0.538 |
| 0.137391 | 0.222596 | 0.537 |
| 0.143568 | 0.233706 | 0.539 |
| -0.0235253 | 0.0384103 | 0.54 |
| 0.0123125 | 0.0201015 | 0.54 |
| -0.0342124 | 0.055811 | 0.54 |
| 0.141904 | 0.232151 | 0.541 |
| 0.0442442 | 0.072342 | 0.541 |
| 0.00560877 | 0.0091997 | 0.542 |
| 0.0360552 | 0.0591668 | 0.542 |
| 0.0285103 | 0.0467857 | 0.542 |
| -0.0111193 | 0.0183165 | 0.544 |
| -0.0430477 | 0.0710931 | 0.545 |
| 0.112311 | 0.185756 | 0.545 |
| 0.0101292 | 0.0168031 | 0.547 |
| 0.110496 | 0.183349 | 0.547 |
| -0.123235 | 0.204945 | 0.548 |
| 0.110695 | 0.184303 | 0.548 |
| -0.0647951 | 0.108238 | 0.549 |
| 0.17891 | 0.298811 | 0.549 |
| -0.0366498 | 0.0613452 | 0.55 |
| 0.0301298 | 0.0505105 | 0.551 |
| -0.0795209 | 0.133254 | 0.551 |
| 0.0366447 | 0.0614159 | 0.551 |
| -0.0290776 | 0.0488623 | 0.552 |
| -0.0311343 | 0.0524359 | 0.553 |
| -0.00491626 | 0.00827438 | 0.552 |
| -0.0294902 | 0.0497818 | 0.554 |
| -0.112215 | 0.189426 | 0.554 |
| -0.0275277 | 0.0464552 | 0.553 |
| -0.0551073 | 0.0935136 | 0.556 |
| -0.0131693 | 0.0223924 | 0.556 |
| -0.0693974 | 0.118433 | 0.558 |

|  |  |  |
| --- | --- | --- |
| -0.0166489 | 0.028339 | 0.557 |
| 0.0844655 | 0.144056 | 0.558 |
| -0.00824913 | 0.0140878 | 0.558 |
| 0.0139434 | 0.0238051 | 0.558 |
| -0.0127379 | 0.0217764 | 0.559 |
| 0.0846826 | 0.144704 | 0.558 |
| -0.0568867 | 0.097935 | 0.561 |
| -0.0273986 | 0.0470956 | 0.561 |
| 0.0596534 | 0.1027 | 0.561 |
| -0.0124948 | 0.0215053 | 0.561 |
| 0.0751738 | 0.129447 | 0.561 |
| 0.0356526 | 0.0615237 | 0.562 |
| -0.0941295 | 0.162383 | 0.562 |
| -0.00484405 | 0.00835571 | 0.562 |
| -0.0732578 | 0.126443 | 0.562 |
| 0.0797894 | 0.138716 | 0.565 |
| 0.105566 | 0.183922 | 0.566 |
| 0.00961757 | 0.0167184 | 0.565 |
| -0.00487557 | 0.00852184 | 0.567 |
| -0.0652267 | 0.114023 | 0.567 |
| 0.105408 | 0.183849 | 0.566 |
| -0.0421267 | 0.0740456 | 0.569 |
| 0.0242807 | 0.0426763 | 0.569 |
| 0.0229718 | 0.040355 | 0.569 |
| -0.0520647 | 0.0916524 | 0.57 |
| 0.0151507 | 0.0267101 | 0.571 |
| -0.125271 | 0.220833 | 0.571 |
| 0.12352 | 0.217411 | 0.57 |
| 0.111582 | 0.198092 | 0.573 |
| -0.0767965 | 0.136607 | 0.574 |
| 0.00319622 | 0.00570445 | 0.575 |
| -0.0115806 | 0.0206379 | 0.575 |
| -0.0542087 | 0.0966744 | 0.575 |
| -0.0648807 | 0.116004 | 0.576 |
| 0.0134021 | 0.0239609 | 0.576 |
| -0.0337747 | 0.0605172 | 0.577 |
| -0.0125345 | 0.0225338 | 0.578 |
| 0.102049 | 0.184116 | 0.579 |
| 0.0308618 | 0.0556828 | 0.579 |
| -0.0912627 | 0.165075 | 0.58 |
| -0.00796556 | 0.0144415 | 0.581 |
| 0.0175231 | 0.0317034 | 0.58 |
| 0.0264088 | 0.0478366 | 0.581 |
| -0.0458476 | 0.0832495 | 0.582 |
| -0.0904697 | 0.164818 | 0.583 |
| -0.0578018 | 0.105234 | 0.583 |
| -0.00203064 | 0.00370241 | 0.583 |
| 0.045144 | 0.0821657 | 0.583 |

|  |  |  |
| --- | --- | --- |
| 0.0741749 | 0.135322 | 0.584 |
| -0.0331937 | 0.0607737 | 0.585 |
| -0.110629 | 0.202505 | 0.585 |
| 0.10108 | 0.184856 | 0.585 |
| 0.0178375 | 0.0325703 | 0.584 |
| -0.0128331 | 0.0235314 | 0.586 |
| 0.0793059 | 0.14566 | 0.586 |
| -0.0592348 | 0.109047 | 0.587 |
| -0.00315629 | 0.00580312 | 0.587 |
| 0.0373501 | 0.0687135 | 0.587 |
| 0.0349126 | 0.0642374 | 0.587 |
| -0.032427 | 0.0599234 | 0.588 |
| -0.0124553 | 0.0229968 | 0.588 |
| 0.0750449 | 0.138925 | 0.589 |
| -0.0677873 | 0.126151 | 0.591 |
| -0.0480477 | 0.0895398 | 0.592 |
| -0.112562 | 0.209695 | 0.591 |
| -0.00401658 | 0.00748347 | 0.591 |
| -0.0151855 | 0.0284241 | 0.593 |
| 0.0114916 | 0.0215487 | 0.594 |
| 0.0972431 | 0.182566 | 0.594 |
| 0.0495177 | 0.0932196 | 0.595 |
| -0.0732433 | 0.138207 | 0.596 |
| 0.0103414 | 0.0195148 | 0.596 |
| -0.0862315 | 0.16375 | 0.598 |
| -0.0312611 | 0.0595676 | 0.6 |
| 0.00782442 | 0.0149267 | 0.6 |
| -0.00941365 | 0.0179581 | 0.6 |
| -0.0347197 | 0.0663294 | 0.601 |
| 0.0612322 | 0.11691 | 0.6 |
| 0.0702463 | 0.13486 | 0.602 |
| 0.0161549 | 0.0309438 | 0.602 |
| 0.0184424 | 0.0355359 | 0.604 |
| 0.103297 | 0.198545 | 0.603 |
| -0.022862 | 0.0439826 | 0.603 |
| -0.0897453 | 0.173088 | 0.604 |
| 0.0665337 | 0.128359 | 0.604 |
| 0.102471 | 0.198523 | 0.606 |
| -0.00434056 | 0.00841814 | 0.606 |
| -0.0170484 | 0.033195 | 0.608 |
| -0.048199 | 0.0938468 | 0.608 |
| -0.0210365 | 0.0410942 | 0.609 |
| 0.0888364 | 0.173454 | 0.609 |
| -0.0256874 | 0.0500371 | 0.608 |
| 0.00876269 | 0.0171001 | 0.608 |
| -0.00762533 | 0.0149204 | 0.609 |
| 0.277699 | 0.547471 | 0.612 |
| 0.111865 | 0.220506 | 0.612 |

|  |  |  |
| --- | --- | --- |
| 0.119717 | 0.235847 | 0.612 |
| -0.00262811 | 0.00517132 | 0.611 |
| -0.0818143 | 0.161306 | 0.612 |
| -0.0165756 | 0.0327376 | 0.613 |
| 0.0117184 | 0.0231472 | 0.613 |
| 0.0366007 | 0.0724048 | 0.613 |
| -0.0715114 | 0.141598 | 0.614 |
| 0.00585867 | 0.0116149 | 0.614 |
| -0.0709188 | 0.140982 | 0.615 |
| 0.0465648 | 0.0933585 | 0.618 |
| -0.0395322 | 0.0793126 | 0.618 |
| -0.0129175 | 0.0259166 | 0.618 |
| -0.00735236 | 0.0147265 | 0.618 |
| -0.0299993 | 0.0602218 | 0.618 |
| -0.0305553 | 0.0613697 | 0.619 |
| -0.141488 | 0.284095 | 0.618 |
| -0.0441385 | 0.0888748 | 0.619 |
| 0.00747474 | 0.0150804 | 0.62 |
| 0.0675375 | 0.136406 | 0.621 |
| 0.0459493 | 0.0926886 | 0.62 |
| -0.0698623 | 0.140705 | 0.62 |
| 0.0724926 | 0.146872 | 0.622 |
| 0.0727393 | 0.147101 | 0.621 |
| 0.0459355 | 0.0934366 | 0.623 |
| 0.0246175 | 0.0501058 | 0.623 |
| 0.0252617 | 0.0514662 | 0.624 |
| 0.07193 | 0.147038 | 0.625 |
| -0.00248653 | 0.00510001 | 0.626 |
| -0.00337373 | 0.00692631 | 0.626 |
| -0.0544826 | 0.111888 | 0.626 |
| 0.0194695 | 0.0401225 | 0.627 |
| 0.0467284 | 0.0962763 | 0.627 |
| -0.0688315 | 0.141892 | 0.628 |
| -0.0403894 | 0.0832847 | 0.628 |
| 0.00407971 | 0.00839183 | 0.627 |
| -0.0315669 | 0.0652609 | 0.629 |
| -0.0322416 | 0.0669368 | 0.63 |
| -0.00561782 | 0.0116301 | 0.629 |
| 0.050117 | 0.104125 | 0.63 |
| 0.00863059 | 0.0179991 | 0.632 |
| 0.00136266 | 0.00285325 | 0.633 |
| -0.100246 | 0.210029 | 0.633 |
| -0.055138 | 0.116222 | 0.635 |
| -0.00858735 | 0.0181143 | 0.635 |
| -0.0577569 | 0.122688 | 0.638 |
| -0.0709914 | 0.151032 | 0.638 |
| -0.014397 | 0.0307623 | 0.64 |
| -0.037665 | 0.0804452 | 0.64 |

|  |  |  |
| --- | --- | --- |
| -0.0390503 | 0.0836903 | 0.641 |
| -0.0653157 | 0.140489 | 0.642 |
| 0.0300027 | 0.064561 | 0.642 |
| -0.00989157 | 0.0213544 | 0.643 |
| 0.0121591 | 0.0262032 | 0.643 |
| 0.0409336 | 0.0886039 | 0.644 |
| 0.0087447 | 0.0189157 | 0.644 |
| -0.050303 | 0.109722 | 0.647 |
| 0.0386307 | 0.084334 | 0.647 |
| 0.0794521 | 0.172914 | 0.646 |
| -0.0417709 | 0.09094 | 0.646 |
| -0.0133504 | 0.0291889 | 0.647 |
| -0.00386699 | 0.00847016 | 0.648 |
| 0.0342205 | 0.0748701 | 0.648 |
| 0.0616264 | 0.135461 | 0.649 |
| -0.0130351 | 0.028602 | 0.649 |
| -0.0553228 | 0.121574 | 0.649 |
| -0.00698967 | 0.0154117 | 0.65 |
| -0.0786983 | 0.173135 | 0.649 |
| 0.00873695 | 0.0193173 | 0.651 |
| 0.0414385 | 0.0917835 | 0.652 |
| -0.00936804 | 0.0206935 | 0.651 |
| -0.0269787 | 0.0595928 | 0.651 |
| 0.0400366 | 0.0886452 | 0.652 |
| 0.00931793 | 0.0207193 | 0.653 |
| 0.0526094 | 0.116735 | 0.652 |
| 0.0781151 | 0.173925 | 0.653 |
| -0.157707 | 0.353031 | 0.655 |
| -0.106019 | 0.236813 | 0.654 |
| 0.070487 | 0.15822 | 0.656 |
| -0.0187856 | 0.0422561 | 0.657 |
| 0.0242669 | 0.054542 | 0.656 |
| 0.011629 | 0.0262555 | 0.658 |
| -0.00911531 | 0.0205451 | 0.657 |
| -0.0112 | 0.0252885 | 0.658 |
| 0.0431659 | 0.0980369 | 0.66 |
| -0.0612188 | 0.139339 | 0.66 |
| -0.054031 | 0.122984 | 0.66 |
| 0.0642362 | 0.146348 | 0.661 |
| 0.0375291 | 0.086002 | 0.663 |
| -0.0608147 | 0.139392 | 0.663 |
| 0.0390507 | 0.0895799 | 0.663 |
| 0.00465884 | 0.0107373 | 0.664 |
| 0.00248797 | 0.00572721 | 0.664 |
| -0.053462 | 0.123506 | 0.665 |
| -0.0463153 | 0.107031 | 0.665 |
| 0.0486194 | 0.113142 | 0.667 |
| -0.0210675 | 0.0489897 | 0.667 |

|  |  |  |
| --- | --- | --- |
| -0.0512152 | 0.119273 | 0.668 |
| -0.020606 | 0.0480119 | 0.668 |
| -0.0183358 | 0.0427913 | 0.668 |
| 0.0872618 | 0.203942 | 0.669 |
| 0.0289101 | 0.0675691 | 0.669 |
| 0.0187591 | 0.0440455 | 0.67 |
| -0.0176185 | 0.0415866 | 0.672 |
| 0.0253477 | 0.0599379 | 0.672 |
| 0.0338359 | 0.0800791 | 0.673 |
| 0.0203011 | 0.048092 | 0.673 |
| 0.00221714 | 0.00526835 | 0.674 |
| 0.0319483 | 0.0758545 | 0.674 |
| -0.0176565 | 0.041956 | 0.674 |
| 0.043513 | 0.104129 | 0.676 |
| 0.0391286 | 0.0942101 | 0.678 |
| 0.0130817 | 0.0314441 | 0.677 |
| 0.0797439 | 0.192516 | 0.679 |
| 0.0165201 | 0.0399091 | 0.679 |
| 0.0501236 | 0.121088 | 0.679 |
| 0.0296217 | 0.0716225 | 0.679 |
| 0.0600213 | 0.145027 | 0.679 |
| 0.01776 | 0.0433401 | 0.682 |
| 0.09584 | 0.234423 | 0.683 |
| 0.0325599 | 0.0796067 | 0.683 |
| 0.0589519 | 0.144163 | 0.683 |
| 0.104635 | 0.257185 | 0.684 |
| -0.0150841 | 0.0370058 | 0.684 |
| 0.0360584 | 0.0888253 | 0.685 |
| 0.00358371 | 0.00883586 | 0.685 |
| -0.0124637 | 0.0308145 | 0.686 |
| 0.020267 | 0.0502393 | 0.687 |
| -0.0747512 | 0.184864 | 0.686 |
| 0.00902863 | 0.0223115 | 0.686 |
| -0.0215061 | 0.0537047 | 0.689 |
| 0.0422599 | 0.10536 | 0.688 |
| 0.0355093 | 0.0886345 | 0.689 |
| 0.0370363 | 0.092701 | 0.69 |
| 0.0290518 | 0.0729325 | 0.69 |
| 0.0386884 | 0.0970479 | 0.69 |
| 0.00661762 | 0.016604 | 0.69 |
| 0.0359109 | 0.0904956 | 0.691 |
| 0.00609343 | 0.0153612 | 0.692 |
| -0.0039665 | 0.00998282 | 0.691 |
| 0.119603 | 0.302157 | 0.692 |
| 0.0280149 | 0.0710277 | 0.693 |
| 0.0356168 | 0.0904367 | 0.694 |
| -0.0621693 | 0.157984 | 0.694 |
| 0.0171345 | 0.0437433 | 0.695 |

|  |  |  |
| --- | --- | --- |
| -0.00959085 | 0.0244188 | 0.694 |
| -0.0323994 | 0.082487 | 0.694 |
| -0.00675492 | 0.0173201 | 0.697 |
| -0.0469596 | 0.121244 | 0.699 |
| -0.0219677 | 0.0566785 | 0.698 |
| -0.0162837 | 0.0420883 | 0.699 |
| -0.0234221 | 0.0607843 | 0.7 |
| 0.0211414 | 0.0547956 | 0.7 |
| -0.0233468 | 0.0609192 | 0.702 |
| 0.00715445 | 0.0186752 | 0.702 |
| 0.0564734 | 0.147446 | 0.702 |
| -0.00741853 | 0.019372 | 0.702 |
| 0.0202932 | 0.0532908 | 0.703 |
| 0.0666876 | 0.17476 | 0.703 |
| 0.0762017 | 0.200176 | 0.703 |
| 0.0353731 | 0.0931254 | 0.704 |
| -0.00857452 | 0.0225923 | 0.704 |
| 0.0303981 | 0.0802775 | 0.705 |
| 0.015594 | 0.0413188 | 0.706 |
| -0.0143793 | 0.0381642 | 0.706 |
| 0.00560817 | 0.014897 | 0.707 |
| 0.055504 | 0.147446 | 0.707 |
| -0.0335884 | 0.0897932 | 0.708 |
| -0.00721984 | 0.0192645 | 0.708 |
| 0.00703625 | 0.0188493 | 0.709 |
| 0.0959744 | 0.256927 | 0.709 |
| -0.00458653 | 0.0123007 | 0.709 |
| -0.0576447 | 0.154513 | 0.709 |
| 0.0296612 | 0.0800055 | 0.711 |
| -0.0046371 | 0.0124682 | 0.71 |
| 0.00698501 | 0.0188121 | 0.71 |
| 0.0562978 | 0.151462 | 0.71 |
| 0.0504463 | 0.136354 | 0.711 |
| -0.0295622 | 0.0800097 | 0.712 |
| -0.0388204 | 0.104885 | 0.711 |
| -0.0249204 | 0.0673301 | 0.711 |
| 0.0689721 | 0.186992 | 0.712 |
| -0.0207442 | 0.0561459 | 0.712 |
| 0.0363446 | 0.0982608 | 0.711 |
| 0.0686081 | 0.187434 | 0.714 |
| 0.0251798 | 0.0686071 | 0.714 |
| -0.0546901 | 0.150144 | 0.716 |
| -0.0189461 | 0.0519892 | 0.716 |
| -0.0445593 | 0.122821 | 0.717 |
| 0.0657308 | 0.181357 | 0.717 |
| 0.0117282 | 0.0324682 | 0.718 |
| 0.0193683 | 0.0540944 | 0.72 |
| 0.0877239 | 0.244782 | 0.72 |

|  |  |  |
| --- | --- | --- |
| 0.0117569 | 0.0327655 | 0.72 |
| -0.0259098 | 0.0727178 | 0.722 |
| -0.0270111 | 0.0759527 | 0.722 |
| -0.0263376 | 0.0741955 | 0.723 |
| 0.0187509 | 0.0530054 | 0.724 |
| 0.0128322 | 0.036406 | 0.724 |
| -0.00794411 | 0.022458 | 0.724 |
| -0.0162273 | 0.0460075 | 0.724 |
| 0.0312019 | 0.0883162 | 0.724 |
| -0.0503969 | 0.143165 | 0.725 |
| 0.0685938 | 0.196263 | 0.727 |
| -0.00525399 | 0.0150273 | 0.727 |
| -0.0323531 | 0.0925476 | 0.727 |
| -0.0429123 | 0.123421 | 0.728 |
| -0.121107 | 0.348025 | 0.728 |
| -0.00928952 | 0.0266581 | 0.727 |
| 0.0631067 | 0.181569 | 0.728 |
| -0.0191986 | 0.0550615 | 0.727 |
| -0.0680956 | 0.19619 | 0.729 |
| -0.0273711 | 0.0789518 | 0.729 |
| -0.0347843 | 0.100501 | 0.729 |
| 0.0326188 | 0.09399 | 0.729 |
| 0.0123847 | 0.0359344 | 0.73 |
| -0.0362057 | 0.105161 | 0.731 |
| 0.00643513 | 0.0187784 | 0.732 |
| 0.0136735 | 0.0397245 | 0.731 |
| -0.0406025 | 0.118047 | 0.731 |
| 0.00817797 | 0.0238832 | 0.732 |
| -0.00484023 | 0.0141905 | 0.733 |
| 0.0128792 | 0.0377168 | 0.733 |
| -0.0542225 | 0.15881 | 0.733 |
| -0.0614424 | 0.180622 | 0.734 |
| -0.0608137 | 0.178878 | 0.734 |
| -0.160397 | 0.474645 | 0.735 |
| -0.0287366 | 0.0847596 | 0.735 |
| -0.0349726 | 0.103367 | 0.735 |
| 0.0602693 | 0.179013 | 0.736 |
| 0.01781 | 0.0535753 | 0.74 |
| -0.00798553 | 0.0243282 | 0.743 |
| 0.0343676 | 0.104588 | 0.742 |
| -0.0504373 | 0.153706 | 0.743 |
| 0.0395929 | 0.120334 | 0.742 |
| 0.0271105 | 0.0828639 | 0.744 |
| -0.0301225 | 0.091896 | 0.743 |
| 0.0182241 | 0.0560769 | 0.745 |
| 0.0573496 | 0.176323 | 0.745 |
| 0.0457133 | 0.140494 | 0.745 |
| -0.0714063 | 0.22012 | 0.746 |

|  |  |  |
| --- | --- | --- |
| -0.00868402 | 0.026807 | 0.746 |
| 0.0564203 | 0.174314 | 0.746 |
| 0.0136531 | 0.0420298 | 0.745 |
| -0.0197665 | 0.0613704 | 0.747 |
| 0.0309199 | 0.0962282 | 0.748 |
| -0.023599 | 0.0735349 | 0.748 |
| -0.0182091 | 0.0570728 | 0.75 |
| 0.0811866 | 0.253926 | 0.749 |
| 0.00583354 | 0.0183127 | 0.75 |
| 0.0879469 | 0.276523 | 0.75 |
| 0.00689606 | 0.0218094 | 0.752 |
| 0.0424942 | 0.134494 | 0.752 |
| -0.0220891 | 0.0700849 | 0.753 |
| 0.0638032 | 0.202172 | 0.752 |
| 0.0243632 | 0.0770487 | 0.752 |
| -0.0118318 | 0.0374762 | 0.752 |
| -0.0275973 | 0.0879653 | 0.754 |
| 0.0470495 | 0.149379 | 0.753 |
| 0.0120986 | 0.038538 | 0.754 |
| -0.0284991 | 0.0907625 | 0.754 |
| -0.029629 | 0.0944092 | 0.754 |
| 0.08635 | 0.277651 | 0.756 |
| -0.0189825 | 0.0609378 | 0.755 |
| -0.0358388 | 0.114996 | 0.755 |
| -0.0282793 | 0.0907659 | 0.755 |
| 0.12133 | 0.392691 | 0.757 |
| -0.0277331 | 0.0896833 | 0.757 |
| 0.04245 | 0.137399 | 0.757 |
| -0.0126126 | 0.0410558 | 0.759 |
| 0.062591 | 0.204046 | 0.759 |
| 0.102965 | 0.336068 | 0.759 |
| 0.0110671 | 0.0361128 | 0.759 |
| -0.0278685 | 0.0907731 | 0.759 |
| -0.00595984 | 0.019414 | 0.759 |
| 0.0196431 | 0.064162 | 0.759 |
| -0.0294929 | 0.0966233 | 0.76 |
| 0.0274099 | 0.0900538 | 0.761 |
| 0.0324118 | 0.106876 | 0.762 |
| 0.0728341 | 0.240782 | 0.762 |
| -0.0075933 | 0.0251456 | 0.763 |
| 0.0138462 | 0.0462236 | 0.765 |
| -0.0379053 | 0.126098 | 0.764 |
| 0.0126712 | 0.0422869 | 0.764 |
| -0.0527198 | 0.176618 | 0.765 |
| 0.0328315 | 0.109866 | 0.765 |
| -0.0123802 | 0.0415102 | 0.766 |
| 0.0266387 | 0.0896347 | 0.766 |
| -0.0102792 | 0.0346547 | 0.767 |

|  |  |  |
| --- | --- | --- |
| 0.00274348 | 0.00922485 | 0.766 |
| -0.0274657 | 0.092681 | 0.767 |
| 0.0218281 | 0.0740044 | 0.768 |
| -0.0433458 | 0.146889 | 0.768 |
| 0.00405127 | 0.0137784 | 0.769 |
| 0.00711781 | 0.0242716 | 0.769 |
| 0.0246625 | 0.0840123 | 0.769 |
| 0.0161753 | 0.0551015 | 0.769 |
| -0.0541666 | 0.184886 | 0.77 |
| 0.0272857 | 0.0936629 | 0.771 |
| -0.0228056 | 0.0786244 | 0.772 |
| -0.0076169 | 0.0261977 | 0.771 |
| -0.00704533 | 0.0243713 | 0.773 |
| -0.0260827 | 0.0901669 | 0.772 |
| -0.0181357 | 0.0627462 | 0.773 |
| -0.00938012 | 0.0326225 | 0.774 |
| -0.0517153 | 0.179702 | 0.774 |
| -0.0506297 | 0.177565 | 0.776 |
| -0.0252067 | 0.0882435 | 0.775 |
| 0.00432487 | 0.0152738 | 0.777 |
| 0.0153211 | 0.0538558 | 0.776 |
| -0.017071 | 0.060648 | 0.778 |
| -0.0283341 | 0.10041 | 0.778 |
| 0.00743171 | 0.0262852 | 0.777 |
| -0.136126 | 0.482437 | 0.778 |
| -0.00509551 | 0.0180538 | 0.778 |
| 0.0217418 | 0.0775568 | 0.779 |
| -0.0220048 | 0.078934 | 0.78 |
| -0.0214044 | 0.0773067 | 0.782 |
| -0.0186308 | 0.0672263 | 0.782 |
| -0.0275846 | 0.0992412 | 0.781 |
| 0.0385233 | 0.139042 | 0.782 |
| 0.013725 | 0.0494595 | 0.781 |
| 0.024515 | 0.0891413 | 0.783 |
| -0.0288912 | 0.10477 | 0.783 |
| 0.0557118 | 0.202823 | 0.784 |
| -0.00877961 | 0.0319501 | 0.783 |
| 0.0107828 | 0.0392987 | 0.784 |
| -0.0137186 | 0.0501002 | 0.784 |
| -0.0130085 | 0.0476096 | 0.785 |
| -0.0188738 | 0.0694854 | 0.786 |
| -0.254912 | 0.944704 | 0.787 |
| 0.0146072 | 0.0541347 | 0.787 |
| 0.0350355 | 0.131073 | 0.789 |
| 0.000484661 | 0.00180761 | 0.789 |
| 0.0133659 | 0.0497838 | 0.788 |
| -0.0288717 | 0.108513 | 0.79 |
| -0.0237391 | 0.0894086 | 0.791 |

|  |  |  |
| --- | --- | --- |
| -0.0353805 | 0.132757 | 0.79 |
| 0.0610191 | 0.230943 | 0.792 |
| 0.00382688 | 0.0144862 | 0.792 |
| 0.0103634 | 0.0390601 | 0.791 |
| 0.00379891 | 0.0144489 | 0.793 |
| 0.0247699 | 0.0945652 | 0.793 |
| 0.0361518 | 0.138203 | 0.794 |
| 0.00510148 | 0.0196316 | 0.795 |
| 0.0164538 | 0.0633944 | 0.795 |
| 0.041865 | 0.160989 | 0.795 |
| 0.0267569 | 0.102941 | 0.795 |
| -0.0185418 | 0.0711907 | 0.795 |
| -0.0788712 | 0.308111 | 0.798 |
| 0.0277153 | 0.10849 | 0.798 |
| 0.0244941 | 0.0957702 | 0.798 |
| 0.026358 | 0.103562 | 0.799 |
| -0.00366567 | 0.0144952 | 0.8 |
| -0.0323719 | 0.127825 | 0.8 |
| 0.00365182 | 0.0144928 | 0.801 |
| 0.0263072 | 0.104291 | 0.801 |
| -0.00541867 | 0.0214973 | 0.801 |
| -0.004616 | 0.0183007 | 0.801 |
| -0.0332338 | 0.132322 | 0.802 |
| -0.0494626 | 0.197293 | 0.802 |
| -0.00762879 | 0.0305463 | 0.803 |
| 0.0183514 | 0.0735899 | 0.803 |
| 0.0203757 | 0.0821462 | 0.804 |
| 0.0673144 | 0.270322 | 0.803 |
| 0.0165561 | 0.0672033 | 0.805 |
| -0.0199055 | 0.0807003 | 0.805 |
| 0.0035603 | 0.0145131 | 0.806 |
| -0.0343861 | 0.139912 | 0.806 |
| -0.0110934 | 0.0450979 | 0.806 |
| -0.0249493 | 0.101466 | 0.806 |
| -0.0073916 | 0.0302524 | 0.807 |
| 0.0189551 | 0.0783467 | 0.809 |
| 0.0188226 | 0.0778653 | 0.809 |
| 0.0111047 | 0.0459199 | 0.809 |
| -0.0208367 | 0.0867804 | 0.81 |
| 0.00598807 | 0.0248402 | 0.81 |
| -0.0121323 | 0.050576 | 0.81 |
| 0.0254332 | 0.105447 | 0.809 |
| 0.0397481 | 0.165374 | 0.81 |
| -0.0200463 | 0.0839512 | 0.811 |
| 0.00346286 | 0.0145284 | 0.812 |
| 0.00871319 | 0.0364849 | 0.811 |
| -0.0264106 | 0.1102 | 0.811 |
| -0.0270231 | 0.113003 | 0.811 |

|  |  |  |
| --- | --- | --- |
| 0.00755897 | 0.0317451 | 0.812 |
| -0.00250666 | 0.0105782 | 0.813 |
| -0.00773255 | 0.0324756 | 0.812 |
| -0.0207313 | 0.0883844 | 0.815 |
| -0.0304386 | 0.129933 | 0.815 |
| -0.036124 | 0.155288 | 0.816 |
| 0.00372855 | 0.0162746 | 0.819 |
| -0.0143767 | 0.0626328 | 0.818 |
| 0.0169277 | 0.0741242 | 0.819 |
| 0.0193401 | 0.0851781 | 0.82 |
| -0.0764969 | 0.337973 | 0.821 |
| 0.0409957 | 0.181728 | 0.822 |
| 0.0186519 | 0.0824316 | 0.821 |
| 0.0256795 | 0.114148 | 0.822 |
| 0.0309342 | 0.137706 | 0.822 |
| 0.00439137 | 0.0196769 | 0.823 |
| 0.0306768 | 0.137748 | 0.824 |
| -0.0138935 | 0.0630577 | 0.826 |
| -0.0211784 | 0.0963877 | 0.826 |
| 0.0158014 | 0.0723461 | 0.827 |
| 0.0210864 | 0.0965359 | 0.827 |
| 0.00352944 | 0.0161793 | 0.827 |
| 0.0325226 | 0.150382 | 0.829 |
| -0.0627222 | 0.289847 | 0.829 |
| -0.0630225 | 0.290813 | 0.828 |
| -0.026984 | 0.125849 | 0.83 |
| 0.0284388 | 0.131872 | 0.829 |
| -0.0216961 | 0.100474 | 0.829 |
| -0.0163715 | 0.0758222 | 0.829 |
| -0.0114745 | 0.0532938 | 0.83 |
| -0.0270728 | 0.12633 | 0.83 |
| 0.0267039 | 0.124621 | 0.83 |
| 0.0439011 | 0.20911 | 0.834 |
| -0.0262932 | 0.126047 | 0.835 |
| 0.0100174 | 0.0480505 | 0.835 |
| -0.0245419 | 0.117115 | 0.834 |
| -0.0204325 | 0.0983243 | 0.835 |
| -0.01142 | 0.0555793 | 0.837 |
| -0.0313969 | 0.152436 | 0.837 |
| -0.0114592 | 0.0562821 | 0.839 |
| 0.00457076 | 0.0224136 | 0.838 |
| -0.0159218 | 0.0781401 | 0.839 |
| 0.0517012 | 0.253036 | 0.838 |
| 0.0136064 | 0.0670963 | 0.839 |
| 0.00796 | 0.0393308 | 0.84 |
| 0.00708676 | 0.0350094 | 0.84 |
| -0.0121341 | 0.0600856 | 0.84 |
| -0.00966154 | 0.0476827 | 0.839 |

|  |  |  |
| --- | --- | --- |
| -0.0040796 | 0.0203409 | 0.841 |
| -0.0135137 | 0.0674543 | 0.841 |
| 0.00288418 | 0.0144191 | 0.841 |
| 0.0157233 | 0.0788029 | 0.842 |
| -0.0380766 | 0.192814 | 0.843 |
| 0.0165673 | 0.0845148 | 0.845 |
| -0.00424565 | 0.0216143 | 0.844 |
| 0.0408621 | 0.210089 | 0.846 |
| 0.00431914 | 0.0222389 | 0.846 |
| 0.0118144 | 0.0612666 | 0.847 |
| 0.0407971 | 0.210761 | 0.847 |
| -0.0294984 | 0.152957 | 0.847 |
| -0.0109787 | 0.0568012 | 0.847 |
| 0.0360164 | 0.19001 | 0.85 |
| -0.0202509 | 0.106742 | 0.85 |
| 0.0236008 | 0.124799 | 0.85 |
| -0.0117022 | 0.0614651 | 0.849 |
| -0.00892402 | 0.0475708 | 0.851 |
| -0.258422 | 1.36937 | 0.85 |
| 0.00791904 | 0.0422317 | 0.851 |
| 0.0235734 | 0.124793 | 0.85 |
| -0.00877109 | 0.0467618 | 0.851 |
| 0.0117167 | 0.062879 | 0.852 |
| 0.0202658 | 0.108798 | 0.852 |
| -0.00622959 | 0.0333081 | 0.852 |
| 0.00874765 | 0.0467797 | 0.852 |
| -0.0178804 | 0.0964442 | 0.853 |
| -0.0305902 | 0.164927 | 0.853 |
| -0.0332744 | 0.181673 | 0.855 |
| 0.0115315 | 0.0625852 | 0.854 |
| -0.0190183 | 0.103361 | 0.854 |
| 0.0216578 | 0.11949 | 0.856 |
| -0.00560049 | 0.0307108 | 0.855 |
| 0.00359594 | 0.0200195 | 0.857 |
| -0.0155276 | 0.0864447 | 0.857 |
| -0.00263954 | 0.0147493 | 0.858 |
| 0.0220883 | 0.12322 | 0.858 |
| 0.00302907 | 0.0169775 | 0.858 |
| 0.00495934 | 0.027822 | 0.859 |
| -0.0413793 | 0.231738 | 0.858 |
| -0.0144494 | 0.0806274 | 0.858 |
| -0.0320936 | 0.18159 | 0.86 |
| 0.0377811 | 0.213563 | 0.86 |
| -0.00144576 | 0.00817728 | 0.86 |
| -0.00932974 | 0.0525216 | 0.859 |
| 0.0210264 | 0.119197 | 0.86 |
| -0.0412538 | 0.238874 | 0.863 |
| -0.0179123 | 0.104342 | 0.864 |

|  |  |  |
| --- | --- | --- |
| -0.00210308 | 0.012408 | 0.865 |
| 0.00915937 | 0.0553631 | 0.869 |
| -0.0296439 | 0.179945 | 0.869 |
| 0.00225223 | 0.0137118 | 0.87 |
| -0.0136915 | 0.0847143 | 0.872 |
| 0.0205809 | 0.127916 | 0.872 |
| -0.00661046 | 0.0409281 | 0.872 |
| 0.00598973 | 0.0370447 | 0.872 |
| 0.020884 | 0.128806 | 0.871 |
| 0.0230272 | 0.144078 | 0.873 |
| 0.00927644 | 0.0586929 | 0.874 |
| -0.00477863 | 0.0303245 | 0.875 |
| 0.00335841 | 0.0215217 | 0.876 |
| -0.00870135 | 0.0555533 | 0.876 |
| 0.00692304 | 0.0447891 | 0.877 |
| 0.0153263 | 0.0995563 | 0.878 |
| 0.0122614 | 0.0803489 | 0.879 |
| -0.00791883 | 0.0519005 | 0.879 |
| -0.0093509 | 0.0614921 | 0.879 |
| -0.00322288 | 0.0213666 | 0.88 |
| 0.00516991 | 0.0343383 | 0.88 |
| 0.0262051 | 0.17523 | 0.881 |
| 0.0117881 | 0.080104 | 0.883 |
| -0.00349171 | 0.0236263 | 0.883 |
| -0.0171929 | 0.117908 | 0.884 |
| -0.0215435 | 0.149779 | 0.886 |
| 0.00254956 | 0.0177348 | 0.886 |
| 0.0128302 | 0.0896492 | 0.886 |
| -0.0209718 | 0.146879 | 0.886 |
| 0.106551 | 0.749237 | 0.887 |
| -0.00311691 | 0.0219774 | 0.887 |
| -0.0579581 | 0.409204 | 0.887 |
| -0.0133647 | 0.0945996 | 0.888 |
| 0.0139863 | 0.0986809 | 0.887 |
| 0.0076033 | 0.0542187 | 0.888 |
| 0.00277509 | 0.0196757 | 0.888 |
| 0.013491 | 0.0965507 | 0.889 |
| -0.0103003 | 0.0738218 | 0.889 |
| -0.0163912 | 0.117049 | 0.889 |
| -0.0481541 | 0.34524 | 0.889 |
| 0.00966427 | 0.0697319 | 0.89 |
| 0.00733387 | 0.0534444 | 0.891 |
| 0.012827 | 0.0935529 | 0.891 |
| 0.0171869 | 0.124986 | 0.891 |
| -0.0115561 | 0.0849843 | 0.892 |
| -0.0276454 | 0.20214 | 0.891 |
| -0.00154051 | 0.0113614 | 0.892 |
| -0.0224806 | 0.168967 | 0.894 |

|  |  |  |
| --- | --- | --- |
| 0.01326 | 0.10015 | 0.895 |
| 0.0063391 | 0.0474739 | 0.894 |
| -0.0196099 | 0.147224 | 0.894 |
| 0.0269109 | 0.203445 | 0.895 |
| 0.0128957 | 0.097735 | 0.895 |
| -0.00706396 | 0.0533835 | 0.895 |
| -0.0311667 | 0.237249 | 0.895 |
| -0.0110004 | 0.084167 | 0.896 |
| -0.0121918 | 0.0936106 | 0.896 |
| -0.00697811 | 0.0534642 | 0.896 |
| 0.0126191 | 0.0982395 | 0.898 |
| -0.00380074 | 0.0297051 | 0.898 |
| 0.00827781 | 0.0646888 | 0.898 |
| -0.00206495 | 0.0163056 | 0.899 |
| -0.0285244 | 0.22416 | 0.899 |
| -0.00652728 | 0.0516799 | 0.899 |
| 0.0256071 | 0.203514 | 0.9 |
| 0.0258229 | 0.206675 | 0.901 |
| -0.00210104 | 0.0166718 | 0.9 |
| -0.0031475 | 0.025104 | 0.9 |
| -0.00884906 | 0.0707441 | 0.9 |
| -0.00692327 | 0.0549692 | 0.9 |
| -0.012597 | 0.101317 | 0.901 |
| -0.00672536 | 0.0543046 | 0.901 |
| 0.0247646 | 0.20368 | 0.903 |
| 0.0209912 | 0.171478 | 0.903 |
| 0.00168806 | 0.0137631 | 0.902 |
| -0.0091166 | 0.0742472 | 0.902 |
| 0.0245752 | 0.203564 | 0.904 |
| -0.0170564 | 0.141584 | 0.904 |
| 0.0123012 | 0.102172 | 0.904 |
| -0.0206915 | 0.170962 | 0.904 |
| -0.00483461 | 0.0403471 | 0.905 |
| 0.00806079 | 0.0667952 | 0.904 |
| -0.0107622 | 0.0906516 | 0.905 |
| 0.0166445 | 0.139407 | 0.905 |
| 0.0238474 | 0.204903 | 0.907 |
| 0.0148446 | 0.127967 | 0.908 |
| 0.015205 | 0.133676 | 0.909 |
| 0.00817978 | 0.0717093 | 0.909 |
| -0.0102151 | 0.089257 | 0.909 |
| 0.00417567 | 0.0372443 | 0.911 |
| -0.0222155 | 0.199023 | 0.911 |
| -0.013312 | 0.119225 | 0.911 |
| 0.00740308 | 0.0668261 | 0.912 |
| -0.0089159 | 0.081263 | 0.913 |
| 0.00180072 | 0.0167189 | 0.914 |
| 0.0166436 | 0.155231 | 0.915 |

|  |  |  |
| --- | --- | --- |
| 0.0219419 | 0.207458 | 0.916 |
| 0.00188265 | 0.017947 | 0.916 |
| 0.00324013 | 0.0312744 | 0.917 |
| -0.00252126 | 0.0244425 | 0.918 |
| 0.00128089 | 0.0125723 | 0.919 |
| 0.0128114 | 0.128361 | 0.92 |
| 0.0199246 | 0.206934 | 0.923 |
| 0.00865635 | 0.0899019 | 0.923 |
| -0.00802612 | 0.0846045 | 0.924 |
| 0.00907037 | 0.0950401 | 0.924 |
| -0.0186861 | 0.198164 | 0.925 |
| 0.0196682 | 0.209345 | 0.925 |
| -0.00401338 | 0.0438108 | 0.927 |
| 0.00297259 | 0.0322639 | 0.927 |
| 0.0106399 | 0.115337 | 0.926 |
| 0.00164254 | 0.0177596 | 0.926 |
| 0.00668683 | 0.0738609 | 0.928 |
| -0.00637634 | 0.0701987 | 0.928 |
| 0.00760796 | 0.0837242 | 0.928 |
| 0.00304917 | 0.0335109 | 0.928 |
| -0.00317635 | 0.0353652 | 0.928 |
| 0.0015894 | 0.017838 | 0.929 |
| -0.00278809 | 0.0312963 | 0.929 |
| 0.0184293 | 0.209477 | 0.93 |
| 0.00940914 | 0.107173 | 0.93 |
| 0.00637155 | 0.0738313 | 0.931 |
| -0.00100587 | 0.0117088 | 0.932 |
| 0.00359276 | 0.0417128 | 0.931 |
| 0.0118639 | 0.138162 | 0.932 |
| 0.00309855 | 0.0368482 | 0.933 |
| 0.0014944 | 0.0176391 | 0.932 |
| 0.00626113 | 0.0735931 | 0.932 |
| -0.0160612 | 0.193671 | 0.934 |
| 0.00923587 | 0.111828 | 0.934 |
| -0.00495412 | 0.0600257 | 0.934 |
| -0.0155651 | 0.188726 | 0.934 |
| -0.011077 | 0.134567 | 0.934 |
| 0.000875599 | 0.0106287 | 0.934 |
| -0.00633555 | 0.0764464 | 0.934 |
| 0.0106641 | 0.127677 | 0.933 |
| 0.00180825 | 0.0223645 | 0.936 |
| 0.00479974 | 0.0606301 | 0.937 |
| -0.0022515 | 0.0280127 | 0.936 |
| 0.00666052 | 0.0852711 | 0.938 |
| -0.0102147 | 0.130262 | 0.937 |
| 0.0282622 | 0.369993 | 0.939 |
| -0.00146906 | 0.0189992 | 0.938 |
| -0.00695904 | 0.0932174 | 0.94 |

|  |  |  |
| --- | --- | --- |
| 0.000778411 | 0.0104298 | 0.941 |
| -0.00305912 | 0.0407392 | 0.94 |
| -0.00748476 | 0.104693 | 0.943 |
| 0.00162318 | 0.0224533 | 0.942 |
| -0.00690012 | 0.0972367 | 0.943 |
| 0.0059363 | 0.0849227 | 0.944 |
| 0.00338449 | 0.0496794 | 0.946 |
| -0.00118561 | 0.0176925 | 0.947 |
| -0.00197874 | 0.0293998 | 0.946 |
| -0.000110551 | 0.00168262 | 0.948 |
| 0.00285685 | 0.0431691 | 0.947 |
| -0.00656339 | 0.100015 | 0.948 |
| 0.0106723 | 0.164478 | 0.948 |
| 0.00795634 | 0.128739 | 0.951 |
| -0.00532121 | 0.0858815 | 0.951 |
| -0.025395 | 0.403444 | 0.95 |
| -0.012182 | 0.200912 | 0.952 |
| 0.00359617 | 0.0589672 | 0.951 |
| 0.00270952 | 0.0449643 | 0.952 |
| -0.00748994 | 0.12736 | 0.953 |
| 0.00105893 | 0.0177591 | 0.952 |
| 0.00627393 | 0.108358 | 0.954 |
| -0.020975 | 0.35909 | 0.953 |
| -0.0039959 | 0.0683173 | 0.953 |
| -0.00537175 | 0.0980856 | 0.956 |
| -0.00182074 | 0.0332927 | 0.956 |
| 0.00258795 | 0.0488439 | 0.958 |
| 0.00105546 | 0.0197983 | 0.957 |
| 0.00335092 | 0.0629976 | 0.958 |
| 0.00142062 | 0.0267229 | 0.958 |
| 0.0129497 | 0.246873 | 0.958 |
| -0.00117103 | 0.0217993 | 0.957 |
| -0.00742203 | 0.142251 | 0.958 |
| -0.00408823 | 0.0781718 | 0.958 |
| -0.0020089 | 0.0399594 | 0.96 |
| 0.00425025 | 0.0866341 | 0.961 |
| 0.0109566 | 0.222616 | 0.961 |
| 0.00722388 | 0.159892 | 0.964 |
| 0.0123537 | 0.272362 | 0.964 |
| 0.00366044 | 0.0815831 | 0.964 |
| 0.0044588 | 0.0992313 | 0.964 |
| 0.00312048 | 0.0689711 | 0.964 |
| 0.0125178 | 0.271714 | 0.963 |
| -0.00568037 | 0.130533 | 0.965 |
| -0.00278789 | 0.0631952 | 0.965 |
| -0.00705226 | 0.159909 | 0.965 |
| 0.00305906 | 0.0736517 | 0.967 |
| 0.000709839 | 0.0180933 | 0.969 |

|  |  |  |
| --- | --- | --- |
| -0.000696158 | 0.0177341 | 0.969 |
| 0.00349651 | 0.0909607 | 0.969 |
| 0.0013181 | 0.0353799 | 0.97 |
| 0.000665165 | 0.0177862 | 0.97 |
| 0.00161916 | 0.0426637 | 0.97 |
| 0.00497343 | 0.140997 | 0.972 |
| 0.00356589 | 0.104721 | 0.973 |
| 0.00427368 | 0.124224 | 0.973 |
| -0.00157525 | 0.0455492 | 0.972 |
| -0.000592937 | 0.0177178 | 0.973 |
| 0.00136275 | 0.0419823 | 0.974 |
| -0.00262852 | 0.0804781 | 0.974 |
| -0.00301334 | 0.0946607 | 0.975 |
| -0.00094837 | 0.0300587 | 0.975 |
| -0.00178971 | 0.0567049 | 0.975 |
| -0.000415932 | 0.0135598 | 0.976 |
| -0.000772185 | 0.0250184 | 0.975 |
| 0.000269633 | 0.00897102 | 0.976 |
| 0.00111407 | 0.045511 | 0.98 |
| 0.00469783 | 0.193878 | 0.981 |
| 0.00302897 | 0.129126 | 0.981 |
| -0.00118021 | 0.049473 | 0.981 |
| 0.0299807 | 1.2664 | 0.981 |
| 0.000614462 | 0.0266026 | 0.982 |
| 0.00414206 | 0.184042 | 0.982 |
| -0.000466953 | 0.0219779 | 0.983 |
| 0.00193634 | 0.0859207 | 0.982 |
| -0.00183838 | 0.0888227 | 0.983 |
| -0.000520748 | 0.0271656 | 0.985 |
| -9.87E-05 | 0.00515245 | 0.985 |
| -0.00134962 | 0.0775609 | 0.986 |
| 0.000286313 | 0.0178282 | 0.987 |
| -0.00152437 | 0.101497 | 0.988 |
| -0.000369511 | 0.0263532 | 0.989 |
| 0.00124225 | 0.11083 | 0.991 |
| -0.00237792 | 0.212042 | 0.991 |
| -0.00108316 | 0.11956 | 0.993 |
| 0.000852302 | 0.110915 | 0.994 |
| 0.000152015 | 0.0177856 | 0.993 |
| -0.00141394 | 0.167491 | 0.993 |
| -0.00115787 | 0.179437 | 0.995 |
| 0.000209654 | 0.0380272 | 0.996 |
| -0.000124632 | 0.0216945 | 0.995 |
| 0.000230804 | 0.0910501 | 0.998 |
| -0.00083793 | 0.223001 | 0.997 |
| -0.000252584 | 0.0686088 | 0.997 |
| 0.000280884 | 0.0718926 | 0.997 |
| 0.000230618 | 0.13132 | 0.999 |

|  |  |  |
| --- | --- | --- |
| -1.44E-05 | 0.0178064 | 0.999 |
| 6.16E-06 | 0.00897813 | 0.999 |
| -2.29E-05 | 0.155037 | 1 |
| 5.09E-05 | 0.075573 | 0.999 |
