## Supplementary material for "Variant-Specific Rewiring of Lysosomal Positioning by STARD9 Produces Opposite Cholesterol Trafficking Outcomes": Table 4

| rsID | chr | pos (hg38) | Effect allele | non-effect allele |
| --- | --- | --- | --- | --- |
| rs143875230 | 15 | 42986528 | G | A |
| rs529330569 | 15 | 42980693 | C | T |
| rs62020680 | 15 | 42878426 | G | A |
| rs77641540 | 15 | 42910125 | C | T |
| rs62020698 | 15 | 42945216 | C | T |
| rs62020701 | 15 | 42966883 | G | C |
| rs1548096 | 15 | 42961113 | T | C |
| rs2412748 | 15 | 42935317 | G | A |
| rs28871732 | 15 | 42896650 | C | T |
| rs28796057 | 15 | 42896566 | C | T |
| rs28613047 | 15 | 42883967 | G | A |
| rs7178248 | 15 | 42907321 | C | T |
| rs9920182 | 15 | 42910352 | G | A |
| rs28609797 | 15 | 42922620 | T | C |
| rs3759792 | 15 | 42960566 | A | G |
| rs28374998 | 15 | 42996979 | C | G |
| rs79694045 | 15 | 42948422 | T | G |
| rs35209561 | 15 | 42949600 | A | AC |
| rs9920061 | 15 | 42941863 | T | C |
| rs7166705 | 15 | 42972928 | C | A |
| rs8036674 | 15 | 43000977 | G | C |
| rs28633493 | 15 | 42971109 | T | C |
| rs11332679 | 15 | 42877492 | GA | G |
| rs28494835 | 15 | 42989193 | C | T |
| rs28858165 | 15 | 42885094 | G | A |
| rs28392857 | 15 | 42883751 | C | T |
| rs28649962 | 15 | 42898433 | T | C |
| rs28674880 | 15 | 42871114 | G | T |
| rs144884699 | 15 | 42914174 | C | T |
| rs7173097 | 15 | 42869601 | C | T |
| rs2682075 | 15 | 42913095 | T | G |
| rs8028731 | 15 | 42899126 | G | A |
| rs16957277 | 15 | 42945374 | T | C |
| rs28646262 | 15 | 42890237 | T | C |
| rs4923952 | 15 | 42901233 | A | G |
| rs8028822 | 15 | 43012143 | C | T |
| rs11070385 | 15 | 43007774 | C | T |
| rs13329073 | 15 | 42897670 | A | T |
| rs28407836 | 15 | 42890341 | T | C |
| rs7178752 | 15 | 43004711 | C | T |
| rs28890597 | 15 | 42887550 | T | C |
| rs12913586 | 15 | 42887858 | T | C |
| rs7169603 | 15 | 43008227 | C | T |
| rs7176915 | 15 | 43009573 | G | T |
| rs62020705 | 15 | 43005140 | G | A |
| rs56114844 | 15 | 42887206 | C | T |
| rs1197533 | 15 | 42906729 | G | A |
| rs2169581 | 15 | 42934568 | G | A |
| rs71108187 | 15 | 42887941 | C | CT |
| rs62020697 | 15 | 42942050 | C | T |
| rs1206698 | 15 | 42911888 | C | T |
| rs12437637 | 15 | 42920620 | C | G |
| rs7165888 | 15 | 42898361 | A | G |
| rs12904179 | 15 | 42868307 | G | A |
| rs2277532 | 15 | 42952501 | A | G |
| rs2126603 | 15 | 42928587 | T | G |
| rs8025479 | 15 | 42968589 | T | C |
| rs62020699 | 15 | 42947415 | T | C |
| rs12440652 | 15 | 42972201 | C | A |
| rs12440651 | 15 | 42972200 | C | A |
| rs2682074 | 15 | 42904892 | C | T |
| rs6493069 | 15 | 42894567 | C | T |
| rs9806175 | 15 | 42884122 | C | T |
| rs28627318 | 15 | 42893859 | A | G |
| rs28805017 | 15 | 42887650 | T | C |
| rs4924702 | 15 | 42975842 | T | C |
| rs28715020 | 15 | 42880082 | G | A |
| rs8036096 | 15 | 42878822 | T | C |
| rs9920231 | 15 | 42917576 | G | A |
| rs28575571 | 15 | 42924988 | G | A |
| rs1381855 | 15 | 42971807 | G | A |
| rs28664053 | 15 | 42922689 | G | A |
| rs6493068 | 15 | 42878595 | A | G |
| rs998954 | 15 | 42988571 | C | T |
| rs1984501 | 15 | 42987512 | G | A |
| rs3917223 | 15 | 42963993 | T | C |
| rs555343183 | 15 | 42900109 | C | T |
| rs2016538 | 15 | 42933901 | A | T |
| rs12917056 | 15 | 42953491 | T | C |
| rs72721511 | 15 | 42994161 | T | C |
| rs12914380 | 15 | 42950070 | C | T |
| rs4924699 | 15 | 42908911 | G | A |
| rs34474101 | 15 | 42895262 | A | G |
| rs12910799 | 15 | 42982666 | G | T |
| rs72719486 | 15 | 42897273 | C | T |
| rs10712537 | 15 | 42919283 | TA | T |
| rs12911334 | 15 | 43013055 | G | A |
| rs35326878 | 15 | 42983379 | C | G |
| rs4923954 | 15 | 42969910 | C | G |
| rs371849516 | 15 | 42947665 | GA | G |
| rs543516155 | 15 | 42974900 | T | C |
| rs539214491 | 15 | 42937269 | G | A |
| rs540683576 | 15 | 42943926 | T | A |
| rs369926923 | 15 | 42979041 | C | T |
| rs149889485 | 15 | 42913969 | T | A |
| rs184616495 | 15 | 43004660 | C | T |
| rs554304132 | 15 | 42955636 | T | C |
| rs150495511 | 15 | 42899170 | A | AT |
| rs575373910 | 15 | 43005726 | C | T |
| rs558339555 | 15 | 42913637 | CA | C |
| rs116154059 | 15 | 42962111 | C | A |
| rs114245127 | 15 | 42972868 | C | A |
| rs150012114 | 15 | 42924974 | A | G |
| rs116967439 | 15 | 42956352 | C | T |
| rs77231660 | 15 | 42981109 | C | T |
| rs141111379 | 15 | 42935162 | C | T |
| rs776118491 | 15 | 42993735 | G | A |
| rs557349180 | 15 | 42993733 | G | A |
| rs556811858 | 15 | 42993734 | AG | A |
| rs143378302 | 15 | 42886009 | T | A |
| rs201384130 | 15 | 42964023 | T | G |
| rs117122292 | 15 | 42879243 | G | A |
| rs117725364 | 15 | 42998580 | A | C |
| rs138000286 | 15 | 42952142 | C | T |
| rs16957292 | 15 | 42982370 | C | T |
| rs140433299 | 15 | 42910322 | T | C |
| rs377202784 | 15 | 42920633 | TGGC | T |
| rs114954873 | 15 | 42920847 | C | A |
| rs115507310 | 15 | 42920848 | T | G |
| rs34436896 | 15 | 42973919 | C | T |
| rs142838405 | 15 | 42934675 | G | A |
| rs11070384 | 15 | 42983068 | A | G |
| rs143455945 | 15 | 42965670 | T | C |
| rs13380212 | 15 | 43006842 | T | C |
| rs77547387 | 15 | 42997595 | G | A |
| rs534274491 | 15 | 43006688 | C | T |
| rs192668256 | 15 | 42999153 | C | T |
| rs28535059 | 15 | 42933903 | T | A |
| rs143568757 | 15 | 42902686 | G | A |
| rs75090487 | 15 | 42937481 | T | G |
| rs373428985 | 15 | 42893814 | T | C |
| rs187345619 | 15 | 42919369 | T | C |
| rs561617085 | 15 | 42986011 | A | G |
| rs530091685 | 15 | 42978732 | CT | C |
| rs148549752 | 15 | 42987049 | C | T |
| rs8031111 | 15 | 42911544 | C | T |
| rs181004974 | 15 | 42874762 | C | T |
| rs78948790 | 15 | 42958014 | T | C |
| rs185505090 | 15 | 43006659 | G | T |
| rs529234877 | 15 | 42924755 | A | G |
| rs572544151 | 15 | 42902719 | G | A |
| rs562725261 | 15 | 42973598 | C | T |
| rs186206742 | 15 | 42922541 | C | A |
| rs2054388 | 15 | 42932260 | C | T |
| rs141583312 | 15 | 42975909 | G | A |
| rs562869855 | 15 | 42948721 | A | G |
| rs143800868 | 15 | 42925865 | G | A |
| rs545291075 | 15 | 42871454 | T | A |
| rs75487201 | 15 | 42921911 | T | G |
| rs140959617 | 15 | 42958045 | A | G |
| rs75773646 | 15 | 42921982 | A | G |
| rs542549746 | 15 | 42910957 | T | C |
| rs182366902 | 15 | 42922724 | A | C |
| rs138556557 | 15 | 42895364 | G | A |
| rs562776360 | 15 | 43002939 | T | C |
| rs564224994 | 15 | 42966397 | T | A |
| rs537102106 | 15 | 42967458 | G | C |
| rs185098077 | 15 | 42912675 | C | T |
| rs79984506 | 15 | 43001654 | T | C |
| rs76841629 | 15 | 43005378 | A | G |
| rs112362877 | 15 | 42938705 | G | A |
| rs1197534 | 15 | 42907922 | T | A |
| rs59276064 | 15 | 42919636 | G | A |
| rs189290697 | 15 | 42931689 | C | T |
| rs116491117 | 15 | 42921767 | G | A |
| rs76206222 | 15 | 42907919 | A | T |
| rs74570754 | 15 | 42984738 | T | C |
| rs143251532 | 15 | 42998512 | T | C |
| rs192977893 | 15 | 42887615 | C | T |
| rs11352938 | 15 | 42945607 | AT | A |
| rs147870890 | 15 | 42925732 | A | G |
| rs375380714 | 15 | 42998215 | T | G |
| rs563964635 | 15 | 42893798 | A | G |
| rs141998787 | 15 | 42913448 | A | G |
| rs78769997 | 15 | 42922138 | C | G |
| rs118074434 | 15 | 43003277 | C | T |
| rs544535807 | 15 | 42973310 | T | G |
| rs185738711 | 15 | 42981245 | A | T |
| rs138976273 | 15 | 42902170 | C | T |
| rs112929196 | 15 | 43008566 | G | A |
| rs557783986 | 15 | 42951686 | T | C |
| rs558490453 | 15 | 42935340 | G | A |
| rs7164041 | 15 | 42917832 | A | G |
| rs186133585 | 15 | 42908179 | C | T |
| rs57880396 | 15 | 42912858 | G | A |
| rs114659791 | 15 | 42885873 | G | A |
| rs116640263 | 15 | 42981739 | C | T |
| rs77541575 | 15 | 42975754 | C | T |
| rs76783611 | 15 | 42988659 | C | T |
| rs145449308 | 15 | 42997265 | C | G |
| rs7179275 | 15 | 43004823 | G | T |
| rs568614887 | 15 | 42947941 | G | A |
| rs114026095 | 15 | 42973492 | C | A |
| rs191375171 | 15 | 42888109 | C | T |
| rs547814549 | 15 | 42964424 | G | A |
| rs150358878 | 15 | 42952069 | A | T |
| rs139693200 | 15 | 42942444 | C | G |
| rs190806658 | 15 | 42940285 | A | G |
| rs115840904 | 15 | 42966913 | C | T |
| rs76864255 | 15 | 42971050 | T | A |
| rs570963333 | 15 | 42925646 | G | A |
| rs182927920 | 15 | 42967984 | T | C |
| rs114804224 | 15 | 42987662 | G | A |
| rs558747146 | 15 | 42880208 | G | A |
| rs56071832 | 15 | 42988453 | C | G |
| rs181075931 | 15 | 42884135 | G | A |
| rs77496195 | 15 | 42967615 | T | G |
| rs191793117 | 15 | 42914340 | T | C |
| rs552124488 | 15 | 42973990 | G | A |
| rs537550535 | 15 | 42971805 | A | G |
| rs117337065 | 15 | 42903762 | G | A |
| rs117347505 | 15 | 42993718 | C | T |
| rs78563306 | 15 | 42967694 | C | T |
| rs557339939 | 15 | 42929212 | G | A |
| rs566413696 | 15 | 42983841 | T | C |
| rs547972042 | 15 | 43005198 | G | A |
| rs191544070 | 15 | 42974511 | A | C |
| rs184036150 | 15 | 42934772 | A | C |
| rs34312632 | 15 | 42976589 | G | GATAA |
| rs552139984 | 15 | 42981736 | G | A |
| rs190984235 | 15 | 42929593 | A | C |
| rs142301748 | 15 | 42884365 | C | T |
| rs571185339 | 15 | 42879699 | A | G |
| rs184351682 | 15 | 42944285 | T | C |
| rs548547947 | 15 | 42947931 | C | T |
| rs146056091 | 15 | 42986205 | A | G |
| rs9806313 | 15 | 42884268 | G | A |
| rs114454965 | 15 | 43009037 | C | T |
| rs550575683 | 15 | 42960132 | T | G |
| rs185596875 | 15 | 42894226 | T | C |
| rs535405759 | 15 | 43005328 | G | A |
| rs7179367 | 15 | 43005074 | C | T |
| rs150630012 | 15 | 42942394 | C | T |
| rs138218565 | 15 | 42953167 | A | C |
| rs113476707 | 15 | 42878194 | A | G |
| rs145057608 | 15 | 42915246 | C | T |
| rs74570780 | 15 | 42908305 | C | T |
| rs9806161 | 15 | 42883890 | A | T |
| rs116738094 | 15 | 42868777 | G | A |
| rs115723709 | 15 | 42962045 | C | T |
| rs146894776 | 15 | 42967945 | T | C |
| rs7176123 | 15 | 42947732 | G | A |
| rs139400588 | 15 | 43000387 | G | A |
| rs142183834 | 15 | 43010103 | A | T |
| rs113183563 | 15 | 42898438 | G | A |
| rs75580066 | 15 | 42877838 | A | G |
| rs142202313 | 15 | 42996605 | A | C |
| rs554508188 | 15 | 42931509 | C | T |
| rs541474906 | 15 | 42968291 | A | G |
| rs114969207 | 15 | 42891216 | G | A |
| rs188917136 | 15 | 42936275 | G | A |
| rs563467704 | 15 | 42982383 | T | C |
| rs35735714 | 15 | 42962133 | T | TC |
| rs183231149 | 15 | 42887050 | G | A |
| rs76232023 | 15 | 42893541 | C | A |
| rs188034879 | 15 | 42979637 | G | A |
| rs546446869 | 15 | 42932919 | C | T |
| rs192727326 | 15 | 42948390 | A | C |
| rs148952135 | 15 | 42966774 | T | C |
| rs191526970 | 15 | 42957820 | T | A |
| rs569388456 | 15 | 42980580 | C | T |
| rs139251799 | 15 | 42918317 | T | C |
| rs182320019 | 15 | 42908120 | A | G |
| rs187329646 | 15 | 42983026 | C | A |
| rs150988986 | 15 | 42924675 | A | T |
| rs143837600 | 15 | 42981862 | G | T |
| rs116103891 | 15 | 42955577 | A | T |
| rs191423170 | 15 | 42971334 | C | T |
| rs561774462 | 15 | 42931411 | C | T |
| rs117095901 | 15 | 42912799 | C | T |
| rs11630330 | 15 | 43005852 | G | T |
| rs142975069 | 15 | 42981529 | G | A |
| rs7178567 | 15 | 42944414 | C | T |
| rs148100429 | 15 | 42875632 | C | T |
| rs190779787 | 15 | 42885334 | C | A |
| rs1381856 | 15 | 42972019 | C | T |
| rs2054389 | 15 | 42963923 | T | C |
| rs142285781 | 15 | 43007204 | G | A |
| rs573802972 | 15 | 42925909 | G | A |
| rs2733224 | 15 | 42913563 | G | A |
| rs148174299 | 15 | 42993741 | G | A |
| rs55683827 | 15 | 42905457 | A | G |
| rs181239306 | 15 | 42916138 | T | C |
| rs117722860 | 15 | 43008329 | C | T |
| rs185850923 | 15 | 42993431 | T | C |
| rs556862650 | 15 | 42989496 | A | C |
| rs139408969 | 15 | 42990087 | C | T |
| rs189693176 | 15 | 42950613 | A | G |
| rs76989793 | 15 | 42922584 | A | T |
| rs188362035 | 15 | 42918484 | G | A |
| rs4924701 | 15 | 42973522 | C | T |
| rs572827078 | 15 | 42913020 | G | T |
| rs4244590 | 15 | 42935141 | G | A |
| rs117350263 | 15 | 42951611 | A | G |
| rs200262486 | 15 | 42978421 | GT | G |
| rs193196275 | 15 | 42908780 | A | G |
| rs145938839 | 15 | 42884633 | C | A |
| rs117422009 | 15 | 43011777 | C | G |
| rs16957244 | 15 | 42873831 | C | T |
| rs148386184 | 15 | 42954593 | A | T |
| rs7174028 | 15 | 42922561 | G | T |
| rs192937247 | 15 | 42973303 | C | T |
| rs189618809 | 15 | 42907976 | C | A |
| rs1049733221 | 15 | 42914070 | G | A |
| rs933480389 | 15 | 42942363 | C | G |
| rs144595138 | 15 | 43011980 | G | A |
| rs141311377 | 15 | 42912418 | A | C |
| rs145466527 | 15 | 42940296 | G | A |
| rs556210813 | 15 | 42997532 | T | A |
| rs181155374 | 15 | 42963621 | C | A |
| rs561908498 | 15 | 42925887 | C | G |
| rs543230721 | 15 | 42912342 | C | T |
| rs115309896 | 15 | 42929461 | T | C |
| rs537461773 | 15 | 42923097 | G | C |
| rs4509988 | 15 | 42980258 | G | T |
| rs138942621 | 15 | 42930975 | T | C |
| rs375843171 | 15 | 42983341 | T | C |
| rs118104470 | 15 | 42913334 | C | T |
| rs144575157 | 15 | 42971564 | A | T |
| rs570080968 | 15 | 43013157 | G | A |
| rs572834165 | 15 | 42936208 | G | A |
| rs60763334 | 15 | 42902516 | G | A |
| rs140450055 | 15 | 42961579 | C | T |
| rs150557750 | 15 | 42972607 | G | A |
| rs116640993 | 15 | 42972421 | T | C |
| rs571019011 | 15 | 42870769 | GA | G |
| rs4923955 | 15 | 43002890 | G | A |
| rs66530524 | 15 | 42880097 | T | C |
| rs6493071 | 15 | 42916608 | C | G |
| rs59589598 | 15 | 42884810 | G | A |
| rs2412747 | 15 | 42994605 | T | C |
| rs561140781 | 15 | 42956307 | T | G |
| rs150316748 | 15 | 42901223 | G | A |
| rs148534148 | 15 | 42934574 | G | T |
| rs2682073 | 15 | 42904360 | T | C |
| rs2899069 | 15 | 42994373 | C | T |
| rs184380146 | 15 | 42995397 | C | T |
| rs1037982814 | 15 | 42900693 | C | T |
| rs189526440 | 15 | 42907138 | T | A |
| rs115922627 | 15 | 42907139 | C | A |
| rs552385948 | 15 | 43007566 | T | A |
| rs34714662 | 15 | 43001896 | C | T |
| rs6493074 | 15 | 43004985 | A | C |
| rs115342811 | 15 | 42881852 | C | T |
| rs146799639 | 15 | 42870779 | T | C |
| rs557473381 | 15 | 43000769 | G | A |
| rs75776391 | 15 | 43011820 | G | A |
| rs139987841 | 15 | 42965625 | C | T |
| rs2126602 | 15 | 43009278 | T | C |
| rs9920498 | 15 | 42985753 | G | C |
| rs567155071 | 15 | 42903081 | G | A |
| rs192592711 | 15 | 42877415 | G | A |
| rs192718382 | 15 | 42905736 | C | T |
| rs11632981 | 15 | 43000340 | A | C |
| rs190822677 | 15 | 42989502 | G | A |
| rs116042800 | 15 | 42887128 | C | G |
| rs138835702 | 15 | 42897067 | T | C |
| rs75658696 | 15 | 42978682 | T | C |
| rs111339056 | 15 | 42872121 | C | T |
| rs111569959 | 15 | 42874184 | A | T |
| rs185185807 | 15 | 42986374 | C | T |
| rs548296727 | 15 | 42970157 | T | C |
| rs140735548 | 15 | 42893937 | C | G |
| rs16957250 | 15 | 42878625 | C | T |
| rs2412746 | 15 | 42994548 | C | T |
| rs558292498 | 15 | 42911675 | C | T |
| rs12910269 | 15 | 42974089 | C | T |
| rs146503263 | 15 | 42878832 | T | G |
| rs149119473 | 15 | 42914260 | G | T |
| rs1993813 | 15 | 42981053 | C | T |
| rs4338775 | 15 | 42980100 | T | C |
| rs147284877 | 15 | 42977716 | T | C |
| rs6416436 | 15 | 42996021 | T | C |
| rs143043128 | 15 | 42936058 | C | T |
| rs116044972 | 15 | 42939606 | G | T |
| rs8042481 | 15 | 43002020 | G | T |
| rs8037022 | 15 | 42965277 | C | T |
| rs144326298 | 15 | 42990770 | T | C |
| rs140916237 | 15 | 42889531 | T | C |
| rs1037990 | 15 | 43007489 | T | C |
| rs111850927 | 15 | 42921045 | T | A |
| rs55805543 | 15 | 42868205 | T | C |
| rs7162959 | 15 | 42938552 | A | G |
| rs7162939 | 15 | 42938472 | C | T |
| rs9796571 | 15 | 43005659 | T | C |
| rs116238461 | 15 | 43001328 | T | C |
| rs572970295 | 15 | 42930764 | C | A |
| rs1972447 | 15 | 42960907 | C | T |
| rs146708171 | 15 | 42979312 | T | C |
| rs74873423 | 15 | 42924611 | C | G |
| rs3803341 | 15 | 42945005 | T | C |
| rs139393261 | 15 | 42922092 | C | T |
| rs139550877 | 15 | 42924878 | G | T |
| rs116269151 | 15 | 42919023 | C | T |
| rs61069183 | 15 | 42890153 | T | C |
| rs80258074 | 15 | 42923044 | T | A |
| rs76215594 | 15 | 42926099 | G | T |
| rs33987108 | 15 | 42936429 | A | AT |
| rs8031283 | 15 | 42903011 | G | A |
| rs117710203 | 15 | 42953008 | C | T |
| rs151022817 | 15 | 42910971 | G | A |
| rs552525489 | 15 | 42956466 | C | T |
| rs180750404 | 15 | 42912673 | C | T |
| rs541359954 | 15 | 42880415 | C | G |
| rs145784520 | 15 | 42966451 | C | T |
| rs112780488 | 15 | 42975207 | T | C |
| rs147891025 | 15 | 42885547 | G | A |
| rs150770038 | 15 | 42897487 | C | T |
| rs549942856 | 15 | 42877419 | C | T |
| rs8031767 | 15 | 42918707 | C | T |
| rs148438272 | 15 | 42918775 | G | T |
| rs149843783 | 15 | 42935351 | C | T |
| rs145037353 | 15 | 42912432 | C | A |
| rs771755844 | 15 | 42933197 | C | A |
| rs138077192 | 15 | 42985240 | G | C |
| rs186041641 | 15 | 42871238 | A | G |
| rs74533898 | 15 | 42909616 | G | A |
| rs557668526 | 15 | 42931262 | T | C |
| rs546580872 | 15 | 42929355 | A | G |
| rs180684653 | 15 | 42982257 | A | G |
| rs530713585 | 15 | 42912062 | G | A |
| rs546077226 | 15 | 42903738 | G | GT |
| rs573769722 | 15 | 42901255 | G | A |
| rs535809709 | 15 | 42883375 | C | T |
| rs145034051 | 15 | 42992977 | T | C |
| rs567161987 | 15 | 42914001 | G | A |
| rs568462372 | 15 | 42924192 | C | G |
| rs551671670 | 15 | 42927127 | T | C |
| rs776170188 | 15 | 42982418 | G | A |
| rs191518723 | 15 | 42922665 | G | T |
| rs12912026 | 15 | 42878529 | T | A |
| rs73413030 | 15 | 42897626 | A | G |
| rs376331294 | 15 | 42960653 | G | A |
| rs547298098 | 15 | 42913562 | C | T |
| rs199781595 | 15 | 42941307 | A | G |
| rs185103862 | 15 | 43003994 | C | T |
| rs113913915 | 15 | 42970124 | T | C |
| rs543192039 | 15 | 42914069 | C | T |
| rs560519383 | 15 | 42994134 | T | C |
| rs141900598 | 15 | 42885895 | C | T |
| rs180994451 | 15 | 42939263 | A | G |
| rs139382852 | 15 | 43003552 | G | A |
| rs537741020 | 15 | 42936512 | A | C |
| rs144640419 | 15 | 42892393 | T | A |
| rs187320661 | 15 | 42934800 | T | C |
| rs186564556 | 15 | 42902293 | G | C |
| rs373905733 | 15 | 42906618 | G | C |
| rs569234006 | 15 | 42940808 | G | GT |
| rs142774139 | 15 | 42874805 | G | A |
| rs191535749 | 15 | 42985222 | C | G |
| rs144230475 | 15 | 42987511 | C | T |
| rs553452245 | 15 | 42908200 | G | T |
| rs141795624 | 15 | 42979926 | G | A |
| rs187287466 | 15 | 42887754 | C | A |
| rs142424434 | 15 | 42925466 | G | C |
| rs16957264 | 15 | 42906232 | C | T |
| rs140207100 | 15 | 43011559 | T | A |
| rs191443517 | 15 | 43005135 | C | T |
| rs1040748507 | 15 | 42889826 | G | A |
| rs34312217 | 15 | 42902986 | CA | C |
| rs80305987 | 15 | 42877994 | A | G |
| rs16957284 | 15 | 42955369 | T | C |
| rs556243779 | 15 | 42902209 | C | A |
| rs541817722 | 15 | 42940819 | C | T |
| rs186771984 | 15 | 42950609 | C | T |
| rs571167694 | 15 | 43011992 | C | T |
| rs147209414 | 15 | 42897063 | G | A |
| rs114936782 | 15 | 42908384 | T | C |
| rs146248884 | 15 | 42965927 | A | G |
| rs141227308 | 15 | 42952389 | G | A |
| rs78084245 | 15 | 42963434 | A | G |
| rs148028047 | 15 | 42903243 | T | C |
| rs17776090 | 15 | 42943386 | A | G |
| rs78365925 | 15 | 42905493 | G | C |
| rs142005131 | 15 | 42882373 | C | A |
| rs567241776 | 15 | 42927294 | C | T |
| rs182906122 | 15 | 42953382 | C | T |
| rs536514977 | 15 | 42875884 | C | T |
| rs562977445 | 15 | 42947538 | T | C |
| rs73413058 | 15 | 42988269 | A | G |
| rs143766227 | 15 | 42883585 | CAA | C |
| rs141919417 | 15 | 42901329 | T | A |
| rs140878173 | 15 | 42911799 | G | C |
| rs570044324 | 15 | 42983733 | T | C |
| rs58483557 | 15 | 42956876 | C | A |
| rs187610922 | 15 | 43004989 | C | T |
| rs146488861 | 15 | 42957808 | A | G |
| rs562509996 | 15 | 43005091 | G | T |
| rs181630879 | 15 | 42889666 | G | C |
| rs184008832 | 15 | 42890930 | G | A |
| rs763442891 | 15 | 42878527 | A | ACT |
| rs572583293 | 15 | 42993853 | C | G |
| rs549786810 | 15 | 42931943 | G | A |
| rs150514250 | 15 | 42999694 | G | A |
| rs138963231 | 15 | 42983997 | G | C |
| rs142211421 | 15 | 42884940 | G | A |
| rs1115752 | 15 | 43008863 | C | T |
| rs147060258 | 15 | 42890797 | GAA | G |
| rs117252626 | 15 | 42953935 | T | C |
| rs547468800 | 15 | 42991255 | G | A |
| rs181258192 | 15 | 42950910 | G | A |
| rs138592907 | 15 | 42946486 | C | T |
| rs576834861 | 15 | 42877871 | G | A |
| rs530032514 | 15 | 42907916 | A | T |
| rs544857253 | 15 | 42904460 | G | C |
| rs117017655 | 15 | 42872405 | G | C |
| rs147200598 | 15 | 42892930 | G | C |
| rs151019978 | 15 | 42991740 | T | C |
| rs574816840 | 15 | 42987187 | C | T |
| rs540626947 | 15 | 42917314 | A | T |
| rs77337431 | 15 | 42909774 | G | A |
| rs147047460 | 15 | 42885872 | C | T |
| rs200325235 | 15 | 43010236 | CA | C |
| rs147153494 | 15 | 42956820 | C | T |
| rs559045802 | 15 | 42906458 | A | G |
| rs72721503 | 15 | 42968409 | G | C |
| rs114066074 | 15 | 42927698 | A | G |
| rs183189070 | 15 | 42979608 | T | C |
| rs183621980 | 15 | 42884021 | G | A |
| rs552314563 | 15 | 43004920 | G | A |
| rs141585079 | 15 | 42909609 | A | G |
| rs142290993 | 15 | 42978320 | T | C |
| rs150970024 | 15 | 42987339 | C | G |
| rs6493070 | 15 | 42900946 | A | G |
| rs182889305 | 15 | 42998734 | C | T |
| rs149265736 | 15 | 43008367 | C | T |
| rs181498789 | 15 | 43010216 | A | G |
| rs149328464 | 15 | 42887671 | T | C |
| rs117347582 | 15 | 42998252 | C | A |
| rs139359398 | 15 | 42909794 | T | C |
| rs551010070 | 15 | 42929055 | A | T |
| rs62020703 | 15 | 42989120 | T | G |
| rs187236177 | 15 | 42939431 | C | A |
| rs117423500 | 15 | 42885260 | G | A |
| rs534379603 | 15 | 42894853 | G | A |
| rs148848820 | 15 | 42895452 | T | C |
| rs574471910 | 15 | 42971058 | T | G |
| rs144693269 | 15 | 42898455 | G | T |
| rs149836490 | 15 | 42950127 | C | T |
| rs143356796 | 15 | 43010938 | C | T |
| rs151031682 | 15 | 42934972 | G | A |
| rs370980402 | 15 | 42990977 | C | T |
| rs16957288 | 15 | 42958590 | T | C |
| rs141828250 | 15 | 42945444 | C | T |
| rs541558127 | 15 | 42939846 | T | C |
| rs559628380 | 15 | 42919443 | G | T |
| rs201091488 | 15 | 42984985 | T | C |
| rs562676922 | 15 | 42876073 | C | T |
| rs543589659 | 15 | 42960960 | A | C |
| rs150203226 | 15 | 42875831 | G | A |
| rs544054828 | 15 | 42977780 | CAA | C |
| rs143699768 | 15 | 42996145 | C | CT |
| rs149678972 | 15 | 42873237 | G | A |
| rs552175940 | 15 | 42921383 | C | G |
| rs56356401 | 15 | 42954663 | T | C |
| rs138619967 | 15 | 42926998 | C | T |
| rs577021890 | 15 | 42891221 | G | A |
| rs17776339 | 15 | 42968180 | T | A |
| rs79495360 | 15 | 42918441 | T | C |
| rs117193101 | 15 | 42875336 | G | T |
| rs11070383 | 15 | 42887165 | G | T |
| rs191933065 | 15 | 42946188 | G | C |
| rs142696672 | 15 | 42910808 | G | T |
| rs189542431 | 15 | 42974675 | C | G |
| rs151233914 | 15 | 42951943 | T | C |
| rs567039670 | 15 | 42963819 | T | C |
| rs185463000 | 15 | 42915407 | G | A |

| effect allele frequency | beta | se | p-value | INFO |
| --- | --- | --- | --- | --- |
| 0.9729 | 0.1624 | 0.009359 | 2.006E-67 | 0.8974 |
| 0.9938 | 0.3354 | 0.02632 | 3.355E-37 | 0.4878 |
| 0.8584 | 0.04706 | 0.004229 | 9.166E-29 | 0.9523 |
| 0.9306 | 0.06429 | 0.005903 | 1.268E-27 | 0.9202 |
| 0.9063 | 0.05215 | 0.004938 | 4.484E-26 | 0.9997 |
| 0.9034 | 0.0484 | 0.004968 | 2.011E-22 | 0.9621 |
| 0.3516 | 0.02519 | 0.003512 | 7.325E-13 | 0.7361 |
| 0.8219 | -0.027 | 0.003819 | 1.548E-12 | 0.9699 |
| 0.8203 | -0.02688 | 0.003803 | 1.576E-12 | 0.9712 |
| 0.8203 | -0.02688 | 0.003803 | 1.578E-12 | 0.9713 |
| 0.8203 | -0.0268 | 0.003803 | 1.823E-12 | 0.9712 |
| 0.8203 | -0.0268 | 0.003803 | 1.847E-12 | 0.9712 |
| 0.8203 | -0.02676 | 0.003803 | 1.961E-12 | 0.9712 |
| 0.8191 | -0.02752 | 0.003917 | 2.129E-12 | 0.9108 |
| 0.8208 | -0.02673 | 0.003805 | 2.139E-12 | 0.972 |
| 0.8217 | -0.02673 | 0.003808 | 2.257E-12 | 0.9742 |
| 0.8175 | -0.0273 | 0.003902 | 2.627E-12 | 0.9118 |
| 0.3367 | 0.02401 | 0.003431 | 2.63E-12 | 0.7872 |
| 0.8203 | -0.02661 | 0.003804 | 2.648E-12 | 0.9708 |
| 0.8216 | -0.02665 | 0.00381 | 2.654E-12 | 0.973 |
| 0.8214 | -0.02661 | 0.003805 | 2.665E-12 | 0.9744 |
| 0.8205 | -0.02654 | 0.003801 | 2.892E-12 | 0.9731 |
| 0.8176 | -0.02688 | 0.003853 | 3.022E-12 | 0.9349 |
| 0.8214 | -0.02649 | 0.003805 | 3.358E-12 | 0.9742 |
| 0.9709 | -0.06207 | 0.008928 | 3.599E-12 | 0.9232 |
| 0.9709 | -0.0619 | 0.008926 | 4.064E-12 | 0.922 |
| 0.9708 | -0.06154 | 0.008901 | 4.719E-12 | 0.9252 |
| 0.9708 | -0.06139 | 0.008905 | 5.448E-12 | 0.9248 |
| 0.9721 | -0.06192 | 0.009094 | 9.824E-12 | 0.9241 |
| 0.7105 | -0.02182 | 0.003231 | 1.443E-11 | 0.9637 |
| 0.7052 | -0.02249 | 0.003357 | 2.072E-11 | 0.8835 |
| 0.7085 | -0.02153 | 0.003217 | 2.206E-11 | 0.9683 |
| 0.9692 | -0.05815 | 0.008719 | 2.572E-11 | 0.9148 |
| 0.7067 | -0.02132 | 0.003208 | 3.008E-11 | 0.9701 |
| 0.707 | -0.02131 | 0.003211 | 3.19E-11 | 0.9692 |
| 0.3161 | 0.02168 | 0.003273 | 3.5E-11 | 0.8942 |
| 0.3161 | 0.02168 | 0.003273 | 3.508E-11 | 0.8943 |
| 0.7075 | -0.02127 | 0.003212 | 3.597E-11 | 0.969 |
| 0.7064 | -0.02122 | 0.003206 | 3.651E-11 | 0.9706 |
| 0.3157 | 0.02166 | 0.003274 | 3.706E-11 | 0.8938 |
| 0.7091 | -0.02144 | 0.003243 | 3.793E-11 | 0.9535 |
| 0.7091 | -0.02144 | 0.003243 | 3.821E-11 | 0.9534 |
| 0.3149 | 0.02165 | 0.003277 | 3.871E-11 | 0.8937 |
| 0.3149 | 0.02165 | 0.003277 | 3.889E-11 | 0.8937 |
| 0.3149 | 0.02165 | 0.003277 | 3.89E-11 | 0.8937 |
| 0.7065 | -0.02117 | 0.003207 | 4.066E-11 | 0.9703 |
| 0.7075 | -0.0212 | 0.003211 | 4.075E-11 | 0.9697 |
| 0.3136 | 0.02144 | 0.003251 | 4.23E-11 | 0.91 |
| 0.7089 | -0.02135 | 0.003242 | 4.504E-11 | 0.9538 |
| 0.3149 | 0.02136 | 0.003246 | 4.717E-11 | 0.9106 |
| 0.7071 | -0.02127 | 0.003233 | 4.727E-11 | 0.9565 |
| 0.3129 | 0.02134 | 0.003254 | 5.369E-11 | 0.9095 |
| 0.7063 | -0.02102 | 0.003205 | 5.423E-11 | 0.9711 |
| 0.7057 | -0.02101 | 0.003207 | 5.728E-11 | 0.9688 |
| 0.3136 | 0.0213 | 0.003254 | 5.97E-11 | 0.9082 |
| 0.312 | 0.02131 | 0.003258 | 6.115E-11 | 0.9088 |
| 0.3153 | 0.02128 | 0.003256 | 6.322E-11 | 0.9045 |
| 0.3132 | 0.02126 | 0.003254 | 6.453E-11 | 0.9089 |
| 0.3141 | 0.02132 | 0.003265 | 6.523E-11 | 0.9016 |
| 0.3146 | 0.0213 | 0.003261 | 6.537E-11 | 0.903 |
| 0.7062 | -0.02088 | 0.003206 | 7.359E-11 | 0.9704 |
| 0.7066 | -0.02085 | 0.003206 | 7.811E-11 | 0.9712 |
| 0.7051 | -0.02082 | 0.003201 | 7.828E-11 | 0.9716 |
| 0.9755 | -0.06149 | 0.00956 | 1.26E-10 | 0.948 |
| 0.704 | -0.02053 | 0.003193 | 1.294E-10 | 0.9742 |
| 0.3024 | 0.02134 | 0.00332 | 1.297E-10 | 0.8902 |
| 0.7057 | -0.02054 | 0.003201 | 1.409E-10 | 0.9722 |
| 0.7043 | -0.02053 | 0.0032 | 1.419E-10 | 0.9703 |
| 0.9757 | -0.06163 | 0.009642 | 1.641E-10 | 0.9398 |
| 0.9757 | -0.06163 | 0.009642 | 1.641E-10 | 0.9398 |
| 0.3156 | 0.02081 | 0.003261 | 1.77E-10 | 0.9011 |
| 0.9757 | -0.06133 | 0.009643 | 2.01E-10 | 0.94 |
| 0.6992 | -0.01997 | 0.003146 | 2.151E-10 | 0.9943 |
| 0.9748 | -0.05854 | 0.009472 | 6.409E-10 | 0.9377 |
| 0.9782 | -0.058 | 0.01019 | 1.245E-08 | 0.9337 |
| 0.9244 | -0.02721 | 0.005446 | 5.844E-07 | 0.9983 |
| 0.9987 | 0.3247 | 0.06632 | 9.807E-07 | 0.3547 |
| 0.8499 | -0.02 | 0.004142 | 0.000001373 | 0.9452 |
| 0.9313 | -0.02794 | 0.005788 | 0.000001381 | 0.9654 |
| 0.8588 | -0.01793 | 0.004172 | 0.00001716 | 0.9799 |
| 0.8587 | -0.01786 | 0.004166 | 0.00001802 | 0.9823 |
| 0.8568 | -0.01774 | 0.004138 | 0.00001814 | 0.9844 |
| 0.8581 | -0.01775 | 0.004143 | 0.00001831 | 0.9892 |
| 0.8588 | -0.01784 | 0.00417 | 0.00001892 | 0.9808 |
| 0.8579 | -0.01767 | 0.00414 | 0.0000197 | 0.9899 |
| 0.306 | 0.01989 | 0.00466 | 0.00001973 | 0.4494 |
| 0.8592 | -0.01774 | 0.00418 | 0.00002207 | 0.9781 |
| 0.8579 | -0.01768 | 0.004168 | 0.0000222 | 0.977 |
| 0.8577 | -0.01763 | 0.004161 | 0.00002263 | 0.9785 |
| 0.9862 | -0.05844 | 0.01382 | 0.00002365 | 0.8026 |
| 0.9998 | 0.4844 | 0.1158 | 0.00002873 | 0.802 |
| 0.9998 | 0.4818 | 0.1154 | 0.00002996 | 0.8031 |
| 0.9998 | 0.4818 | 0.1154 | 0.00002996 | 0.8031 |
| 0.9914 | -0.07362 | 0.01782 | 0.00003598 | 0.7703 |
| 0.9914 | -0.07342 | 0.0178 | 0.00003704 | 0.7713 |
| 0.9919 | -0.07794 | 0.01895 | 0.00003895 | 0.7226 |
| 0.9991 | -0.3379 | 0.08589 | 0.00008362 | 0.3142 |
| 0.9181 | -0.02933 | 0.007588 | 0.0001107 | 0.4785 |
| 0.9825 | -0.05111 | 0.01342 | 0.0001405 | 0.6742 |
| 0.9815 | 0.06598 | 0.01744 | 0.0001543 | 0.3772 |
| 0.9993 | 0.3734 | 0.09891 | 0.0001595 | 0.3417 |
| 0.9993 | 0.3713 | 0.09867 | 0.0001676 | 0.34 |
| 0.9771 | -0.03896 | 0.01042 | 0.0001842 | 0.8585 |
| 0.9483 | 0.0244 | 0.006553 | 0.0001962 | 0.985 |
| 0.9483 | 0.02414 | 0.006509 | 0.0002078 | 0.9979 |
| 0.992 | -0.06657 | 0.0182 | 0.0002545 | 0.7937 |
| 0.9794 | -0.0396 | 0.01088 | 0.000274 | 0.8727 |
| 0.9851 | -0.05694 | 0.01568 | 0.0002814 | 0.5798 |
| 0.9851 | -0.05694 | 0.01568 | 0.0002814 | 0.5798 |
| 0.9996 | 0.4478 | 0.1258 | 0.0003708 | 0.3532 |
| 0.9998 | 0.5252 | 0.153 | 0.0005959 | 0.4201 |
| 0.9817 | 0.03909 | 0.01145 | 0.0006363 | 0.884 |
| 0.992 | -0.06491 | 0.01921 | 0.0007256 | 0.7045 |
| 0.986 | -0.04177 | 0.01331 | 0.001701 | 0.8468 |
| 0.9992 | 0.2748 | 0.09024 | 0.002323 | 0.3474 |
| 0.9996 | 0.3173 | 0.1053 | 0.002573 | 0.4029 |
| 0.9971 | 0.1218 | 0.04047 | 0.002608 | 0.4243 |
| 0.9993 | -0.2701 | 0.0908 | 0.002935 | 0.3434 |
| 0.9993 | -0.2701 | 0.0908 | 0.002935 | 0.3434 |
| 0.9974 | 0.1097 | 0.0373 | 0.003282 | 0.5708 |
| 0.997 | 0.1081 | 0.03694 | 0.003424 | 0.5123 |
| 0.9992 | 0.2575 | 0.08829 | 0.003541 | 0.3305 |
| 0.9992 | 0.2558 | 0.08862 | 0.00389 | 0.33 |
| 0.9992 | 0.2554 | 0.08866 | 0.003969 | 0.3608 |
| 0.9992 | 0.2535 | 0.08878 | 0.004305 | 0.3604 |
| 0.9992 | 0.1617 | 0.05682 | 0.00443 | 0.8549 |
| 0.9994 | -0.2836 | 0.1001 | 0.004625 | 0.3588 |
| 0.998 | 0.1363 | 0.04973 | 0.006115 | 0.4281 |
| 0.997 | 0.0971 | 0.03581 | 0.006699 | 0.5338 |
| 0.9944 | -0.0794 | 0.02945 | 0.007014 | 0.4338 |
| 0.9987 | -0.125 | 0.04663 | 0.007345 | 0.712 |
| 0.998 | 0.1285 | 0.04795 | 0.007384 | 0.4438 |
| 0.9823 | -0.03149 | 0.01179 | 0.00756 | 0.8567 |
| 0.9665 | -0.03686 | 0.01434 | 0.01015 | 0.3106 |
| 0.9993 | -0.178 | 0.06942 | 0.01035 | 0.6225 |
| 0.9974 | 0.09615 | 0.03787 | 0.01111 | 0.5665 |
| 0.9933 | -0.0476 | 0.01923 | 0.01333 | 0.8363 |
| 0.9923 | -0.04063 | 0.01645 | 0.01351 | 0.9987 |
| 0.9987 | 0.142 | 0.05778 | 0.01399 | 0.491 |
| 0.9998 | 0.3028 | 0.124 | 0.01459 | 0.7128 |
| 0.9998 | 0.3672 | 0.1543 | 0.01736 | 0.4794 |
| 0.9997 | -0.247 | 0.1039 | 0.01745 | 0.6819 |
| 0.9921 | -0.03881 | 0.0164 | 0.01792 | 0.9725 |
| 0.998 | 0.1002 | 0.04242 | 0.01818 | 0.58 |
| 0.9984 | -0.1015 | 0.04327 | 0.01896 | 0.6748 |
| 0.9974 | -0.07895 | 0.03424 | 0.02114 | 0.6893 |
| 0.997 | 0.0822 | 0.03577 | 0.02155 | 0.538 |
| 0.9997 | -0.1978 | 0.08847 | 0.02539 | 0.8477 |
| 0.9973 | 0.08247 | 0.03694 | 0.02557 | 0.5714 |
| 0.9975 | 0.0641 | 0.02884 | 0.02624 | 0.9963 |
| 0.9973 | 0.08185 | 0.03692 | 0.02662 | 0.5715 |
| 0.9997 | 0.2382 | 0.1086 | 0.02823 | 0.4745 |
| 0.9946 | 0.04821 | 0.02203 | 0.02865 | 0.7892 |
| 0.994 | -0.04273 | 0.01959 | 0.02918 | 0.8996 |
| 0.9996 | -0.1873 | 0.08597 | 0.02934 | 0.6353 |
| 0.9981 | -0.08873 | 0.04081 | 0.02969 | 0.6416 |
| 0.9981 | -0.08873 | 0.04081 | 0.02969 | 0.6416 |
| 0.9998 | -0.3969 | 0.1828 | 0.02988 | 0.3271 |
| 0.997 | 0.06909 | 0.03205 | 0.03113 | 0.685 |
| 0.5371 | -0.008014 | 0.003721 | 0.03128 | 0.6006 |
| 0.9481 | 0.01437 | 0.006674 | 0.03132 | 0.9447 |
| 0.9579 | 0.01822 | 0.008558 | 0.03323 | 0.7015 |
| 0.9979 | 0.08607 | 0.04066 | 0.03426 | 0.6078 |
| 0.9968 | 0.06587 | 0.03112 | 0.03427 | 0.6873 |
| 0.997 | 0.07473 | 0.03551 | 0.03533 | 0.5452 |
| 0.8762 | -0.009775 | 0.004661 | 0.03599 | 0.8783 |
| 0.9998 | 0.353 | 0.1691 | 0.0369 | 0.3512 |
| 0.9996 | 0.223 | 0.1079 | 0.03868 | 0.3821 |
| 0.9986 | -0.102 | 0.0494 | 0.03888 | 0.5867 |
| 0.1933 | 0.01011 | 0.004893 | 0.0389 | 0.5544 |
| 0.9969 | 0.07187 | 0.03486 | 0.03926 | 0.5501 |
| 0.9998 | 0.2267 | 0.1105 | 0.04015 | 0.9998 |
| 0.9969 | 0.06817 | 0.03331 | 0.04073 | 0.6107 |
| 0.9969 | 0.07071 | 0.03502 | 0.04344 | 0.5534 |
| 0.9972 | 0.07327 | 0.03635 | 0.04381 | 0.566 |
| 0.9992 | 0.1368 | 0.06855 | 0.04591 | 0.5533 |
| 0.9993 | 0.1355 | 0.0685 | 0.04797 | 0.564 |
| 0.9993 | 0.1355 | 0.0685 | 0.04797 | 0.564 |
| 0.9864 | 0.02735 | 0.01384 | 0.04805 | 0.8073 |
| 0.9609 | 0.01536 | 0.007834 | 0.04989 | 0.8983 |
| 0.9983 | -0.07957 | 0.04059 | 0.04994 | 0.7405 |
| 0.9979 | -0.07497 | 0.03865 | 0.05243 | 0.6541 |
| 0.997 | 0.06892 | 0.0356 | 0.05286 | 0.543 |
| 0.9998 | 0.284 | 0.1478 | 0.05463 | 0.3891 |
| 0.9973 | 0.06944 | 0.03618 | 0.05497 | 0.5925 |
| 0.9998 | 0.3022 | 0.1576 | 0.05516 | 0.5345 |
| 0.9981 | 0.07948 | 0.04148 | 0.05532 | 0.6339 |
| 0.9981 | 0.07918 | 0.04148 | 0.05631 | 0.6361 |
| 0.9981 | 0.07839 | 0.04128 | 0.05754 | 0.6343 |
| 0.9997 | -0.1993 | 0.1052 | 0.05825 | 0.5788 |
| 0.9971 | 0.07595 | 0.04048 | 0.06061 | 0.4594 |
| 0.997 | 0.05922 | 0.03163 | 0.06116 | 0.6893 |
| 0.9982 | 0.07737 | 0.04152 | 0.06242 | 0.6541 |
| 0.9972 | 0.06399 | 0.03442 | 0.06302 | 0.6341 |
| 0.9996 | -0.1747 | 0.09472 | 0.06519 | 0.5504 |
| 0.9979 | -0.07744 | 0.04208 | 0.06572 | 0.5679 |
| 0.9979 | 0.07513 | 0.04094 | 0.06644 | 0.5965 |
| 0.9974 | 0.08742 | 0.04765 | 0.06658 | 0.348 |
| 0.9979 | 0.0749 | 0.04091 | 0.06713 | 0.6119 |
| 0.9979 | 0.07431 | 0.04073 | 0.06808 | 0.6032 |
| 0.9985 | -0.1126 | 0.06209 | 0.06988 | 0.3563 |
| 0.9981 | 0.08742 | 0.04831 | 0.07035 | 0.4812 |
| 0.9986 | 0.07863 | 0.04354 | 0.0709 | 0.8037 |
| 0.9984 | 0.08075 | 0.04472 | 0.07097 | 0.6373 |
| 0.9993 | -0.1773 | 0.09835 | 0.07148 | 0.3134 |
| 0.997 | 0.0592 | 0.03288 | 0.07175 | 0.6317 |
| 0.9983 | 0.07565 | 0.04227 | 0.07349 | 0.6785 |
| 0.998 | -0.09339 | 0.05232 | 0.07424 | 0.3905 |
| 0.9989 | -0.1256 | 0.07106 | 0.07714 | 0.3759 |
| 0.9983 | 0.08526 | 0.04836 | 0.07792 | 0.5065 |
| 0.9982 | 0.07665 | 0.04349 | 0.078 | 0.5975 |
| 0.9972 | -0.05542 | 0.03156 | 0.07907 | 0.7466 |
| 0.9972 | -0.05498 | 0.0314 | 0.07991 | 0.7561 |
| 0.9983 | 0.08278 | 0.04732 | 0.08025 | 0.5204 |
| 0.9992 | -0.1572 | 0.09027 | 0.0817 | 0.3375 |
| 0.9944 | 0.04039 | 0.02321 | 0.08191 | 0.698 |
| 0.9958 | 0.0426 | 0.02452 | 0.0824 | 0.8184 |
| 0.9972 | -0.05366 | 0.0311 | 0.0845 | 0.7619 |
| 0.1528 | 0.008977 | 0.005204 | 0.08455 | 0.5901 |
| 0.9979 | 0.07927 | 0.04625 | 0.08655 | 0.4613 |
| 0.9958 | 0.04199 | 0.02461 | 0.08798 | 0.807 |
| 0.9959 | -0.04834 | 0.02877 | 0.09296 | 0.6112 |
| 0.9995 | -0.1432 | 0.08529 | 0.09316 | 0.6025 |
| 0.9988 | -0.1224 | 0.07342 | 0.09546 | 0.3315 |
| 0.9934 | 0.03581 | 0.02151 | 0.09594 | 0.6797 |
| 0.9989 | -0.117 | 0.07029 | 0.09612 | 0.3928 |
| 0.9966 | 0.05589 | 0.03399 | 0.1001 | 0.534 |
| 0.9987 | 0.07247 | 0.04424 | 0.1014 | 0.8272 |
| 0.9996 | -0.1618 | 0.1001 | 0.1058 | 0.4838 |
| 0.999 | 0.133 | 0.0823 | 0.1062 | 0.3262 |
| 0.9987 | -0.1074 | 0.06734 | 0.1107 | 0.3391 |
| 0.9986 | 0.07031 | 0.04422 | 0.1118 | 0.7824 |
| 0.9986 | 0.07053 | 0.04463 | 0.114 | 0.7916 |
| 0.9966 | 0.04202 | 0.02682 | 0.1171 | 0.8499 |
| 0.9968 | 0.05384 | 0.03455 | 0.1192 | 0.548 |
| 0.9984 | -0.06289 | 0.04038 | 0.1194 | 0.7824 |
| 0.9969 | 0.05351 | 0.03467 | 0.1228 | 0.5519 |
| 0.9968 | 0.0532 | 0.03458 | 0.124 | 0.5493 |
| 0.998 | 0.06132 | 0.03999 | 0.1251 | 0.6513 |
| 0.9987 | 0.07013 | 0.04609 | 0.1281 | 0.7743 |
| 0.9975 | 0.05288 | 0.03481 | 0.1288 | 0.6704 |
| 0.9987 | 0.06776 | 0.04472 | 0.1298 | 0.7998 |
| 0.9975 | 0.05252 | 0.03485 | 0.1318 | 0.6707 |
| 0.9975 | 0.05247 | 0.03485 | 0.1322 | 0.6708 |
| 0.9994 | -0.151 | 0.1004 | 0.1327 | 0.3283 |
| 0.9972 | 0.05262 | 0.0358 | 0.1416 | 0.5783 |
| 0.9975 | 0.05141 | 0.035 | 0.1418 | 0.6701 |
| 0.9993 | -0.104 | 0.07084 | 0.142 | 0.6181 |
| 0.9981 | -0.06055 | 0.04161 | 0.1456 | 0.6435 |
| 0.9998 | -0.2664 | 0.1863 | 0.1527 | 0.3599 |
| 0.9952 | 0.03223 | 0.02259 | 0.1537 | 0.8522 |
| 0.9995 | 0.1547 | 0.1085 | 0.154 | 0.3884 |
| 0.1534 | 0.007324 | 0.00517 | 0.1566 | 0.5962 |
| 0.9992 | -0.09876 | 0.06993 | 0.1578 | 0.5499 |
| 0.9967 | -0.03829 | 0.02714 | 0.1583 | 0.8664 |
| 0.9959 | -0.03651 | 0.02595 | 0.1595 | 0.7458 |
| 0.9997 | 0.172 | 0.1223 | 0.1598 | 0.4848 |
| 0.9998 | -0.1927 | 0.1375 | 0.161 | 0.7027 |
| 0.9969 | -0.04283 | 0.03096 | 0.1666 | 0.7086 |
| 0.9998 | -0.1813 | 0.1318 | 0.1689 | 0.7175 |
| 0.9998 | -0.22 | 0.1611 | 0.1721 | 0.4243 |
| 0.9998 | -0.2561 | 0.1878 | 0.1728 | 0.3629 |
| 0.9992 | 0.07804 | 0.05726 | 0.1729 | 0.8389 |
| 0.9997 | -0.124 | 0.09177 | 0.1766 | 0.8168 |
| 0.9998 | -0.2529 | 0.1879 | 0.1782 | 0.3694 |
| 0.9997 | -0.1235 | 0.09177 | 0.1785 | 0.8174 |
| 0.9998 | -0.2506 | 0.1879 | 0.1824 | 0.3693 |
| 0.9954 | -0.03222 | 0.0242 | 0.183 | 0.7712 |
| 0.9987 | -0.0612 | 0.04628 | 0.186 | 0.7382 |
| 0.9888 | -0.01877 | 0.0142 | 0.1862 | 0.9235 |
| 0.1568 | 0.006624 | 0.005177 | 0.2007 | 0.5839 |
| 0.9983 | -0.06252 | 0.04901 | 0.2021 | 0.5053 |
| 0.1306 | 0.005992 | 0.004744 | 0.2065 | 0.8105 |
| 0.9997 | -0.1152 | 0.09124 | 0.2066 | 0.8397 |
| 0.9997 | -0.1152 | 0.09124 | 0.2066 | 0.8397 |
| 0.1313 | 0.005961 | 0.004751 | 0.2096 | 0.8044 |
| 0.1313 | 0.005959 | 0.004751 | 0.2097 | 0.8044 |
| 0.9991 | 0.05995 | 0.04822 | 0.2138 | 0.9922 |
| 0.9984 | 0.08082 | 0.06551 | 0.2174 | 0.3052 |
| 0.8903 | -0.005888 | 0.004777 | 0.2177 | 0.9295 |
| 0.999 | 0.05698 | 0.04645 | 0.22 | 0.9632 |
| 0.9956 | 0.0313 | 0.02559 | 0.2212 | 0.7119 |
| 0.9993 | 0.09177 | 0.07517 | 0.2222 | 0.5029 |
| 0.9755 | -0.01165 | 0.009598 | 0.2249 | 0.9371 |
| 0.9947 | 0.03257 | 0.02689 | 0.2258 | 0.5504 |
| 0.9996 | 0.1316 | 0.1091 | 0.2279 | 0.4428 |
| 0.999 | 0.05596 | 0.04658 | 0.2296 | 0.9724 |
| 0.9998 | -0.1762 | 0.1468 | 0.23 | 0.6311 |
| 0.99 | 0.01887 | 0.01572 | 0.2301 | 0.8405 |
| 0.9972 | 0.03944 | 0.03289 | 0.2305 | 0.6879 |
| 0.1311 | 0.00575 | 0.004799 | 0.2309 | 0.7893 |
| 0.9994 | 0.07253 | 0.06063 | 0.2316 | 0.8933 |
| 0.1319 | 0.005628 | 0.004719 | 0.233 | 0.8119 |
| 0.9935 | 0.02532 | 0.02125 | 0.2334 | 0.7111 |
| 0.9921 | 0.02349 | 0.01976 | 0.2346 | 0.6652 |
| 0.9974 | 0.04205 | 0.03557 | 0.2372 | 0.6262 |
| 0.9981 | 0.062 | 0.05257 | 0.2383 | 0.3882 |
| 0.9903 | 0.01903 | 0.01621 | 0.2403 | 0.82 |
| 0.8896 | -0.00553 | 0.004724 | 0.2418 | 0.9451 |
| 0.9906 | 0.01861 | 0.0159 | 0.242 | 0.8732 |
| 0.1317 | 0.005504 | 0.004717 | 0.2432 | 0.8134 |
| 0.9968 | -0.04466 | 0.03828 | 0.2434 | 0.4462 |
| 0.9957 | -0.02864 | 0.02461 | 0.2444 | 0.7933 |
| 0.9997 | -0.15 | 0.1289 | 0.2445 | 0.4465 |
| 0.9989 | -0.07549 | 0.06495 | 0.2452 | 0.4597 |
| 0.9986 | 0.05088 | 0.04401 | 0.2476 | 0.7336 |
| 0.9993 | 0.06684 | 0.0583 | 0.2516 | 0.9332 |
| 0.9993 | 0.06684 | 0.0583 | 0.2516 | 0.9332 |
| 0.9998 | 0.1537 | 0.1342 | 0.252 | 0.6141 |
| 0.9968 | -0.04374 | 0.03857 | 0.2568 | 0.4399 |
| 0.9998 | 0.1934 | 0.1707 | 0.2573 | 0.3121 |
| 0.9981 | -0.0434 | 0.03833 | 0.2575 | 0.747 |
| 0.9995 | 0.1401 | 0.1239 | 0.2579 | 0.3121 |
| 0.9993 | -0.07343 | 0.06495 | 0.2582 | 0.6609 |
| 0.1335 | 0.005363 | 0.004745 | 0.2584 | 0.7946 |
| 0.9995 | 0.1072 | 0.09526 | 0.2605 | 0.4619 |
| 0.9981 | -0.04298 | 0.03828 | 0.2615 | 0.7461 |
| 0.9904 | 0.01767 | 0.01574 | 0.2616 | 0.8734 |
| 0.9959 | 0.02619 | 0.02336 | 0.2623 | 0.9183 |
| 0.9998 | -0.1924 | 0.172 | 0.2633 | 0.4139 |
| 0.998 | 0.04579 | 0.04095 | 0.2635 | 0.6173 |
| 0.8891 | -0.005253 | 0.004717 | 0.2654 | 0.9443 |
| 0.9876 | -0.01504 | 0.01364 | 0.2701 | 0.9063 |
| 0.9008 | -0.005544 | 0.005043 | 0.2716 | 0.9118 |
| 0.999 | 0.05122 | 0.04735 | 0.2794 | 0.9352 |
| 0.9791 | 0.01429 | 0.01326 | 0.2812 | 0.5767 |
| 0.134 | 0.005103 | 0.004753 | 0.2831 | 0.7894 |
| 0.8887 | -0.005032 | 0.00471 | 0.2853 | 0.9436 |
| 0.1327 | 0.00502 | 0.004703 | 0.2858 | 0.8133 |
| 0.9987 | 0.04721 | 0.04478 | 0.2918 | 0.7795 |
| 0.1369 | 0.004957 | 0.004709 | 0.2925 | 0.7901 |
| 0.9979 | 0.0425 | 0.0404 | 0.2928 | 0.6145 |
| 0.9963 | -0.0308 | 0.0293 | 0.2931 | 0.6469 |
| 0.9972 | 0.03586 | 0.03415 | 0.2937 | 0.6369 |
| 0.8884 | -0.004941 | 0.004707 | 0.2938 | 0.9428 |
| 0.1302 | 0.005077 | 0.004839 | 0.2941 | 0.7807 |
| 0.9987 | -0.07326 | 0.06996 | 0.295 | 0.3267 |
| 0.9998 | 0.1509 | 0.1446 | 0.2965 | 0.5007 |
| 0.9987 | 0.04662 | 0.04478 | 0.2979 | 0.7828 |
| 0.9987 | 0.04662 | 0.04478 | 0.2979 | 0.7828 |
| 0.9993 | -0.09255 | 0.08899 | 0.2983 | 0.3972 |
| 0.7847 | -0.003966 | 0.003818 | 0.2989 | 0.8392 |
| 0.1302 | 0.00499 | 0.004844 | 0.3029 | 0.779 |
| 0.9993 | -0.1011 | 0.1 | 0.3124 | 0.3275 |
| 0.9416 | -0.00628 | 0.006236 | 0.3139 | 0.9676 |
| 0.9995 | 0.06785 | 0.06749 | 0.3147 | 0.8718 |
| 0.999 | 0.04733 | 0.04732 | 0.3173 | 0.9599 |
| 0.9988 | 0.04744 | 0.04749 | 0.3179 | 0.7976 |
| 0.1305 | 0.004821 | 0.004842 | 0.3195 | 0.778 |
| 0.1293 | 0.004813 | 0.004846 | 0.3206 | 0.783 |
| 0.9964 | 0.02448 | 0.02501 | 0.3278 | 0.9086 |
| 0.9995 | 0.0688 | 0.07037 | 0.3282 | 0.8155 |
| 0.998 | 0.03484 | 0.03567 | 0.3287 | 0.8171 |
| 0.9776 | 0.01056 | 0.01082 | 0.329 | 0.8089 |
| 0.9995 | 0.0671 | 0.0692 | 0.3322 | 0.8533 |
| 0.9995 | -0.09425 | 0.09788 | 0.3356 | 0.3996 |
| 0.9995 | -0.09425 | 0.09788 | 0.3356 | 0.3996 |
| 0.9938 | -0.02024 | 0.02124 | 0.3404 | 0.7413 |
| 0.9998 | 0.161 | 0.1694 | 0.342 | 0.328 |
| 0.9998 | 0.161 | 0.1694 | 0.342 | 0.328 |
| 0.9973 | 0.02865 | 0.03052 | 0.3478 | 0.8227 |
| 0.996 | 0.03861 | 0.04113 | 0.3479 | 0.3066 |
| 0.9935 | -0.02278 | 0.02453 | 0.3529 | 0.5358 |
| 0.8961 | -0.004531 | 0.004893 | 0.3544 | 0.93 |
| 0.1365 | 0.004362 | 0.004711 | 0.3545 | 0.7916 |
| 0.9994 | 0.06077 | 0.06566 | 0.3547 | 0.8459 |
| 0.1368 | 0.004322 | 0.004706 | 0.3584 | 0.7917 |
| 0.9962 | 0.02794 | 0.03043 | 0.3585 | 0.6047 |
| 0.9995 | -0.09085 | 0.09895 | 0.3586 | 0.4197 |
| 0.1358 | 0.004316 | 0.004706 | 0.3591 | 0.7966 |
| 0.1354 | 0.004217 | 0.004714 | 0.371 | 0.7958 |
| 0.9989 | -0.0583 | 0.06531 | 0.3721 | 0.5139 |
| 0.1362 | 0.004207 | 0.004717 | 0.3725 | 0.7909 |
| 0.999 | 0.04272 | 0.04799 | 0.3734 | 0.9353 |
| 0.999 | 0.04244 | 0.04794 | 0.376 | 0.9425 |
| 0.1367 | 0.004168 | 0.004709 | 0.3761 | 0.7912 |
| 0.135 | 0.004144 | 0.004687 | 0.3766 | 0.807 |
| 0.999 | -0.07548 | 0.08542 | 0.3769 | 0.3046 |
| 0.9994 | -0.08528 | 0.09704 | 0.3795 | 0.3939 |
| 0.1362 | 0.004141 | 0.004724 | 0.3806 | 0.7888 |
| 0.9998 | 0.1484 | 0.17 | 0.3828 | 0.367 |
| 0.8932 | -0.004188 | 0.004803 | 0.3832 | 0.9414 |
| 0.1343 | 0.004067 | 0.004678 | 0.3846 | 0.814 |
| 0.1345 | 0.004059 | 0.004674 | 0.3851 | 0.8143 |
| 0.1363 | 0.004081 | 0.004722 | 0.3874 | 0.7888 |
| 0.9996 | 0.1064 | 0.1231 | 0.3877 | 0.3415 |
| 0.9931 | -0.01621 | 0.01886 | 0.3899 | 0.8468 |
| 0.1346 | 0.004032 | 0.004694 | 0.3904 | 0.8068 |
| 0.9958 | -0.02 | 0.02349 | 0.3945 | 0.897 |
| 0.9958 | -0.01972 | 0.02332 | 0.3978 | 0.9016 |
| 0.1339 | 0.003963 | 0.00469 | 0.3982 | 0.8117 |
| 0.9991 | 0.04136 | 0.04904 | 0.399 | 0.9392 |
| 0.9991 | 0.04136 | 0.04904 | 0.399 | 0.9392 |
| 0.9995 | -0.08281 | 0.09838 | 0.3999 | 0.4165 |
| 0.9956 | -0.01911 | 0.02279 | 0.4017 | 0.9147 |
| 0.9995 | -0.08178 | 0.09843 | 0.406 | 0.4196 |
| 0.9995 | -0.08178 | 0.09843 | 0.406 | 0.4196 |
| 0.431 | 0.003428 | 0.004133 | 0.4068 | 0.4951 |
| 0.9483 | -0.008867 | 0.01076 | 0.41 | 0.3649 |
| 0.9956 | -0.01917 | 0.02328 | 0.4104 | 0.8788 |
| 0.9988 | 0.04391 | 0.05336 | 0.4106 | 0.6176 |
| 0.9989 | -0.04269 | 0.05217 | 0.4132 | 0.6897 |
| 0.998 | -0.0339 | 0.04167 | 0.4159 | 0.5952 |
| 0.9977 | -0.03748 | 0.04624 | 0.4176 | 0.432 |
| 0.9998 | 0.1109 | 0.1372 | 0.4188 | 0.699 |
| 0.9998 | 0.1372 | 0.1698 | 0.419 | 0.4119 |
| 0.9974 | 0.02544 | 0.03163 | 0.4213 | 0.8067 |
| 0.9974 | 0.02544 | 0.03163 | 0.4213 | 0.8067 |
| 0.9998 | -0.1055 | 0.132 | 0.4241 | 0.5963 |
| 0.129 | 0.003813 | 0.004783 | 0.4253 | 0.8054 |
| 0.9977 | -0.03617 | 0.04591 | 0.4308 | 0.436 |
| 0.9998 | -0.1255 | 0.1596 | 0.4318 | 0.4304 |
| 0.999 | 0.03752 | 0.04792 | 0.4336 | 0.9218 |
| 0.9998 | 0.09369 | 0.1238 | 0.4491 | 0.6441 |
| 0.9961 | -0.01805 | 0.02401 | 0.4522 | 0.9269 |
| 0.9959 | -0.01727 | 0.02319 | 0.4564 | 0.9462 |
| 0.9971 | 0.02922 | 0.03951 | 0.4596 | 0.4624 |
| 0.9989 | -0.04031 | 0.05461 | 0.4604 | 0.6251 |
| 0.9992 | 0.03822 | 0.0521 | 0.4632 | 0.9456 |
| 0.9989 | -0.03936 | 0.05527 | 0.4763 | 0.6284 |
| 0.9998 | 0.0972 | 0.1385 | 0.4826 | 0.6908 |
| 0.9981 | -0.02694 | 0.03838 | 0.4828 | 0.7323 |
| 0.9966 | -0.02641 | 0.03803 | 0.4875 | 0.4188 |
| 0.9998 | 0.112 | 0.1616 | 0.488 | 0.3232 |
| 0.9974 | -0.02143 | 0.03203 | 0.5035 | 0.7844 |
| 0.9962 | -0.01601 | 0.02432 | 0.5102 | 0.9047 |
| 0.9998 | -0.08709 | 0.1334 | 0.5138 | 0.6068 |
| 0.9998 | -0.08709 | 0.1334 | 0.5138 | 0.6068 |
| 0.9998 | 0.09843 | 0.1527 | 0.5193 | 0.4765 |
| 0.9934 | 0.01683 | 0.02618 | 0.5204 | 0.4648 |
| 0.7588 | -0.003835 | 0.005975 | 0.521 | 0.3164 |
| 0.998 | 0.03411 | 0.05365 | 0.5249 | 0.3823 |
| 0.9998 | 0.07658 | 0.1205 | 0.5249 | 0.783 |
| 0.9992 | 0.0327 | 0.05251 | 0.5335 | 0.9297 |
| 0.9997 | -0.07706 | 0.1243 | 0.5353 | 0.4708 |
| 0.9985 | 0.0228 | 0.03752 | 0.5434 | 0.9525 |
| 0.9995 | -0.05905 | 0.0999 | 0.5545 | 0.4136 |
| 0.9989 | -0.03342 | 0.05665 | 0.5552 | 0.5726 |
| 0.998 | 0.02792 | 0.04767 | 0.5582 | 0.4503 |
| 0.9912 | -0.01327 | 0.02272 | 0.5592 | 0.459 |
| 0.9974 | 0.02371 | 0.04062 | 0.5594 | 0.4908 |
| 0.9993 | 0.04294 | 0.07431 | 0.5634 | 0.5192 |
| 0.9997 | -0.06612 | 0.1147 | 0.5645 | 0.5126 |
| 0.9994 | -0.03728 | 0.06494 | 0.566 | 0.8508 |
| 0.9996 | 0.04627 | 0.0813 | 0.5693 | 0.8883 |
| 0.9997 | 0.04431 | 0.07802 | 0.5701 | 0.9768 |
| 0.9997 | 0.04431 | 0.07802 | 0.5701 | 0.9768 |
| 0.9985 | 0.02162 | 0.03828 | 0.5723 | 0.9386 |
| 0.993 | -0.01054 | 0.01868 | 0.5725 | 0.8525 |
| 0.9998 | -0.07533 | 0.1337 | 0.5732 | 0.619 |
| 0.9995 | -0.0561 | 0.1 | 0.5749 | 0.4242 |
| 0.9992 | 0.02931 | 0.05231 | 0.5752 | 0.926 |
| 0.9982 | 0.02334 | 0.0421 | 0.5794 | 0.6406 |
| 0.9991 | 0.02824 | 0.05114 | 0.5809 | 0.9292 |
| 0.9996 | 0.04474 | 0.0813 | 0.5821 | 0.8563 |
| 0.9097 | 0.002877 | 0.005237 | 0.5827 | 0.9201 |
| 0.9994 | -0.03527 | 0.06461 | 0.5851 | 0.8702 |
| 0.9972 | 0.01984 | 0.03635 | 0.5852 | 0.5505 |
| 0.9974 | -0.02122 | 0.03909 | 0.5873 | 0.5296 |
| 0.931 | 0.004558 | 0.008451 | 0.5896 | 0.4511 |
| 0.9899 | 0.008339 | 0.01567 | 0.5945 | 0.8425 |
| 0.998 | -0.03099 | 0.05856 | 0.5967 | 0.3035 |
| 0.9986 | -0.0292 | 0.05526 | 0.5972 | 0.4875 |
| 0.9984 | 0.01884 | 0.03731 | 0.6137 | 0.9497 |
| 0.9984 | 0.01879 | 0.03733 | 0.6148 | 0.9415 |
| 0.999 | 0.02638 | 0.05372 | 0.6234 | 0.7036 |
| 0.9987 | 0.02508 | 0.05142 | 0.6257 | 0.6218 |
| 0.998 | -0.0277 | 0.05791 | 0.6325 | 0.3068 |
| 0.9955 | 0.01166 | 0.02457 | 0.6352 | 0.7657 |
| 0.9983 | -0.01652 | 0.03558 | 0.6425 | 0.9899 |
| 0.9722 | 0.004366 | 0.009406 | 0.6425 | 0.8664 |
| 0.9989 | -0.02717 | 0.0594 | 0.6474 | 0.5336 |
| 0.9751 | -0.004774 | 0.01045 | 0.6478 | 0.7828 |
| 0.9984 | 0.02703 | 0.05973 | 0.6509 | 0.3799 |
| 0.9714 | 0.004167 | 0.009309 | 0.6544 | 0.8592 |
| 0.9994 | 0.04015 | 0.09048 | 0.6573 | 0.3911 |
| 0.9942 | -0.008954 | 0.02065 | 0.6646 | 0.8369 |
| 0.9987 | -0.02407 | 0.05736 | 0.6747 | 0.475 |
| 0.9816 | 0.006338 | 0.0152 | 0.6766 | 0.4946 |
| 0.9369 | 0.004455 | 0.01079 | 0.6797 | 0.3009 |
| 0.9951 | -0.01379 | 0.03442 | 0.6887 | 0.3668 |
| 0.9946 | -0.009809 | 0.02478 | 0.6922 | 0.6231 |
| 0.9976 | -0.01178 | 0.03045 | 0.6989 | 0.9153 |
| 0.9991 | 0.02787 | 0.07308 | 0.703 | 0.4483 |
| 0.998 | -0.02226 | 0.05851 | 0.7036 | 0.3058 |
| 0.9994 | 0.03062 | 0.08234 | 0.71 | 0.5589 |
| 0.9958 | -0.009645 | 0.02634 | 0.7142 | 0.7185 |
| 0.9987 | -0.01919 | 0.05271 | 0.7158 | 0.5656 |
| 0.9979 | -0.01684 | 0.04644 | 0.7169 | 0.4726 |
| 0.9984 | 0.01331 | 0.03686 | 0.718 | 0.9519 |
| 0.9988 | 0.01475 | 0.04168 | 0.7234 | 0.9988 |
| 0.9957 | -0.01045 | 0.03011 | 0.7285 | 0.5254 |
| 0.9984 | -0.02015 | 0.05867 | 0.7313 | 0.376 |
| 0.9992 | -0.02183 | 0.06457 | 0.7353 | 0.6251 |
| 0.9865 | -0.004539 | 0.01345 | 0.7357 | 0.8589 |
| 0.9998 | -0.03997 | 0.1198 | 0.7387 | 0.692 |
| 0.9061 | -0.001668 | 0.005051 | 0.7412 | 0.9541 |
| 0.9991 | 0.01609 | 0.04888 | 0.742 | 0.9011 |
| 0.9991 | -0.02041 | 0.06425 | 0.7508 | 0.5923 |
| 0.9996 | -0.03758 | 0.1186 | 0.7513 | 0.4065 |
| 0.9979 | -0.0112 | 0.03558 | 0.7529 | 0.7814 |
| 0.9956 | 0.00709 | 0.0229 | 0.7569 | 0.8969 |
| 0.9996 | -0.03046 | 0.09971 | 0.76 | 0.5015 |
| 0.9996 | 0.03311 | 0.1094 | 0.7621 | 0.399 |
| 0.9998 | 0.03334 | 0.1106 | 0.7631 | 0.7561 |
| 0.9729 | -0.00319 | 0.01067 | 0.7649 | 0.6899 |
| 0.9932 | -0.00613 | 0.02054 | 0.7654 | 0.7278 |
| 0.9981 | -0.01402 | 0.04776 | 0.7691 | 0.475 |
| 0.9979 | -0.01045 | 0.03571 | 0.7699 | 0.7765 |
| 0.9986 | -0.01227 | 0.04207 | 0.7705 | 0.8387 |
| 0.9991 | 0.01407 | 0.04836 | 0.7711 | 0.9299 |
| 0.999 | 0.02092 | 0.076 | 0.7832 | 0.3719 |
| 0.9642 | 0.002387 | 0.008694 | 0.7837 | 0.7961 |
| 0.9991 | -0.0226 | 0.08656 | 0.7941 | 0.3098 |
| 0.9994 | -0.02169 | 0.08451 | 0.7975 | 0.4661 |
| 0.9726 | 0.002803 | 0.01101 | 0.7991 | 0.639 |
| 0.9993 | -0.01456 | 0.05815 | 0.8023 | 0.8944 |
| 0.9992 | -0.01441 | 0.058 | 0.8038 | 0.7945 |
| 0.9952 | -0.006056 | 0.02455 | 0.8052 | 0.7192 |
| 0.9934 | 0.004833 | 0.01971 | 0.8063 | 0.8091 |
| 0.967 | -0.001993 | 0.008195 | 0.8079 | 0.9665 |
| 0.9993 | -0.01387 | 0.05816 | 0.8115 | 0.8951 |
| 0.9993 | -0.01357 | 0.05817 | 0.8156 | 0.8958 |
| 0.08957 | -0.001321 | 0.005704 | 0.8168 | 0.7808 |
| 0.9976 | -0.007365 | 0.03219 | 0.819 | 0.837 |
| 0.9993 | -0.01322 | 0.05814 | 0.8202 | 0.8968 |
| 0.9993 | -0.01322 | 0.05814 | 0.8202 | 0.8968 |
| 0.9983 | 0.01045 | 0.04616 | 0.821 | 0.5585 |
| 0.9977 | -0.006639 | 0.03 | 0.8249 | 0.9987 |
| 0.9991 | -0.01412 | 0.06433 | 0.8263 | 0.525 |
| 0.9993 | 0.01492 | 0.06831 | 0.8271 | 0.6653 |
| 0.9891 | 0.003434 | 0.01607 | 0.8308 | 0.7415 |
| 0.9964 | -0.006016 | 0.03088 | 0.8455 | 0.6089 |
| 0.9741 | -0.00197 | 0.01018 | 0.8465 | 0.7943 |
| 0.9997 | -0.0256 | 0.1453 | 0.8601 | 0.3414 |
| 0.9816 | 0.002235 | 0.01283 | 0.8617 | 0.6971 |
| 0.9985 | 0.01089 | 0.06439 | 0.8657 | 0.345 |
| 0.9753 | 0.001664 | 0.01015 | 0.8698 | 0.8361 |
| 0.9998 | -0.02017 | 0.1252 | 0.872 | 0.6948 |
| 0.9686 | -0.001445 | 0.01007 | 0.886 | 0.6703 |
| 0.9902 | -0.002094 | 0.0154 | 0.8919 | 0.9049 |
| 0.9986 | -0.008444 | 0.06462 | 0.896 | 0.3547 |
| 0.9745 | -0.001431 | 0.01106 | 0.8971 | 0.6817 |
| 0.9987 | -0.005069 | 0.0405 | 0.9004 | 0.9824 |
| 0.9964 | 0.003841 | 0.03207 | 0.9046 | 0.5635 |
| 0.9947 | 0.002434 | 0.02033 | 0.9047 | 0.9578 |
| 0.9996 | 0.008417 | 0.07633 | 0.9122 | 0.8327 |
| 0.9996 | -0.013 | 0.129 | 0.9197 | 0.3237 |
| 0.9995 | 0.009169 | 0.09561 | 0.9236 | 0.3832 |
| 0.9683 | -0.0007701 | 0.008439 | 0.9273 | 0.9491 |
| 0.9993 | -0.007342 | 0.0815 | 0.9282 | 0.4223 |
| 0.9392 | 0.0005891 | 0.006544 | 0.9283 | 0.8482 |
| 0.9732 | -0.0009546 | 0.01073 | 0.9291 | 0.6901 |
| 0.9986 | 0.005698 | 0.06506 | 0.9302 | 0.3552 |
| 0.9793 | 0.0009093 | 0.01086 | 0.9333 | 0.8653 |
| 0.9817 | -0.0009572 | 0.01174 | 0.935 | 0.8362 |
| 0.999 | -0.005057 | 0.06543 | 0.9384 | 0.4765 |
| 0.9793 | 0.0008282 | 0.01087 | 0.9393 | 0.8651 |
| 0.9932 | 0.001664 | 0.02642 | 0.9498 | 0.4417 |
| 0.9953 | 0.001176 | 0.02221 | 0.9578 | 0.8923 |
| 0.06585 | 0.0003575 | 0.007025 | 0.9594 | 0.6826 |
| 0.9994 | 0.003252 | 0.07451 | 0.9652 | 0.6162 |
| 0.9988 | -0.001912 | 0.04502 | 0.9661 | 0.8744 |
| 0.9966 | -0.001186 | 0.02858 | 0.9669 | 0.7443 |
| 0.9977 | -0.001198 | 0.05152 | 0.9815 | 0.3478 |
| 0.9989 | -0.001571 | 0.07059 | 0.9822 | 0.3796 |
| 0.9989 | 0.0003227 | 0.07047 | 0.9963 | 0.3812 |

| rsID | chr | pos (hg38) | Effect allele | non-effect allele | effect allele freq | beta |
| --- | --- | --- | --- | --- | --- | --- |
| rs143875230 | 15 | 42986528 | G | A | 0.9728 | -0.1587 |
| rs529330569 | 15 | 42980693 | C | T | 0.9938 | -0.313 |
| rs77641540 | 15 | 42910125 | C | T | 0.9305 | -0.06563 |
| rs62020698 | 15 | 42945216 | C | T | 0.9061 | -0.05213 |
| rs62020701 | 15 | 42966883 | G | C | 0.9033 | -0.04895 |
| rs62020680 | 15 | 42878426 | G | A | 0.8582 | -0.03657 |
| rs1548096 | 15 | 42961113 | T | C | 0.3514 | -0.02131 |
| rs35209561 | 15 | 42949600 | A | AC | 0.3365 | -0.02067 |
| rs2682075 | 15 | 42913095 | T | G | 0.7053 | 0.01957 |
| rs7178752 | 15 | 43004711 | C | T | 0.3156 | -0.01906 |
| rs8028731 | 15 | 42899126 | G | A | 0.7086 | 0.01862 |
| rs4924702 | 15 | 42975842 | T | C | 0.3023 | -0.0192 |
| rs11070385 | 15 | 43007774 | C | T | 0.316 | -0.01891 |
| rs8028822 | 15 | 43012143 | C | T | 0.316 | -0.0189 |
| rs7169603 | 15 | 43008227 | C | T | 0.3148 | -0.0189 |
| rs7176915 | 15 | 43009573 | G | T | 0.3148 | -0.01889 |
| rs62020705 | 15 | 43005140 | G | A | 0.3148 | -0.01889 |
| rs71108187 | 15 | 42887941 | C | CT | 0.7091 | 0.01864 |
| rs12440651 | 15 | 42972200 | C | A | 0.3144 | -0.01872 |
| rs4923952 | 15 | 42901233 | A | G | 0.7072 | 0.01841 |
| rs8025479 | 15 | 42968589 | T | C | 0.3152 | -0.01867 |
| rs28407836 | 15 | 42890341 | T | C | 0.7066 | 0.01836 |
| rs62020697 | 15 | 42942050 | C | T | 0.3148 | -0.01859 |
| rs13329073 | 15 | 42897670 | A | T | 0.7077 | 0.01839 |
| rs28890597 | 15 | 42887550 | T | C | 0.7092 | 0.01856 |
| rs12913586 | 15 | 42887858 | T | C | 0.7092 | 0.01856 |
| rs28646262 | 15 | 42890237 | T | C | 0.7068 | 0.01834 |
| rs7173097 | 15 | 42869601 | C | T | 0.7107 | 0.01845 |
| rs56114844 | 15 | 42887206 | C | T | 0.7066 | 0.01828 |
| rs1197533 | 15 | 42906729 | G | A | 0.7077 | 0.01828 |
| rs7165888 | 15 | 42898361 | A | G | 0.7064 | 0.01824 |
| rs2277532 | 15 | 42952501 | A | G | 0.3134 | -0.01851 |
| rs12440652 | 15 | 42972201 | C | A | 0.314 | -0.01856 |
| rs555343183 | 15 | 42900109 | C | T | 0.9987 | -0.3754 |
| rs62020699 | 15 | 42947415 | T | C | 0.3131 | -0.01845 |
| rs1381855 | 15 | 42971807 | G | A | 0.3154 | -0.01843 |
| rs1206698 | 15 | 42911888 | C | T | 0.7072 | 0.01827 |
| rs12437637 | 15 | 42920620 | C | G | 0.3128 | -0.01834 |
| rs2169581 | 15 | 42934568 | G | A | 0.3135 | -0.01829 |
| rs6493069 | 15 | 42894567 | C | T | 0.7067 | 0.01786 |
| rs2682074 | 15 | 42904892 | C | T | 0.7063 | 0.01782 |
| rs12904179 | 15 | 42868307 | G | A | 0.7058 | 0.01782 |
| rs9806175 | 15 | 42884122 | C | T | 0.7053 | 0.01777 |
| rs28805017 | 15 | 42887650 | T | C | 0.7042 | 0.01767 |
| rs2126603 | 15 | 42928587 | T | G | 0.3118 | -0.01802 |
| rs28715020 | 15 | 42880082 | G | A | 0.7058 | 0.01762 |
| rs6493068 | 15 | 42878595 | A | G | 0.6994 | 0.01725 |
| rs8036096 | 15 | 42878822 | T | C | 0.7045 | 0.0172 |
| rs10712537 | 15 | 42919283 | TA | T | 0.3059 | -0.0246 |
| rs554304132 | 15 | 42955636 | T | C | 0.9991 | 0.3977 |
| rs558339555 | 15 | 42913637 | CA | C | 0.9816 | -0.07931 |
| rs33987108 | 15 | 42936429 | A | AT | 0.431 | -0.01753 |
| rs28609797 | 15 | 42922620 | T | C | 0.8191 | 0.01603 |
| rs11352938 | 15 | 42945607 | AT | A | 0.1931 | -0.0199 |
| rs11332679 | 15 | 42877492 | GA | G | 0.8176 | 0.01559 |
| rs28871732 | 15 | 42896650 | C | T | 0.8203 | 0.01517 |
| rs28796057 | 15 | 42896566 | C | T | 0.8204 | 0.01516 |
| rs28613047 | 15 | 42883967 | G | A | 0.8204 | 0.01513 |
| rs9920182 | 15 | 42910352 | G | A | 0.8204 | 0.01512 |
| rs7178248 | 15 | 42907321 | C | T | 0.8204 | 0.0151 |
| rs79694045 | 15 | 42948422 | T | G | 0.8175 | 0.01545 |
| rs34312632 | 15 | 42976589 | G | GATAA | 0.1527 | -0.02055 |
| rs2412748 | 15 | 42935317 | G | A | 0.8219 | 0.01493 |
| rs3759792 | 15 | 42960566 | A | G | 0.8208 | 0.01479 |
| rs7166705 | 15 | 42972928 | C | A | 0.8217 | 0.01475 |
| rs4924701 | 15 | 42973522 | C | T | 0.131 | -0.01839 |
| rs2054389 | 15 | 42963923 | T | C | 0.1312 | -0.01813 |
| rs1381856 | 15 | 42972019 | C | T | 0.1312 | -0.01813 |
| rs9920061 | 15 | 42941863 | T | C | 0.8203 | 0.01449 |
| rs7178567 | 15 | 42944414 | C | T | 0.1305 | -0.01806 |
| rs28374998 | 15 | 42996979 | C | G | 0.8217 | 0.01448 |
| rs28633493 | 15 | 42971109 | T | C | 0.8206 | 0.01444 |
| rs4509988 | 15 | 42980258 | G | T | 0.1334 | -0.01801 |
| rs8036674 | 15 | 43000977 | G | C | 0.8214 | 0.01439 |
| rs4923955 | 15 | 43002890 | G | A | 0.1339 | -0.01798 |
| rs28494835 | 15 | 42989193 | C | T | 0.8214 | 0.01436 |
| rs12910269 | 15 | 42974089 | C | T | 0.1367 | -0.01768 |
| rs35735714 | 15 | 42962133 | T | TC | 0.1533 | -0.01925 |
| rs2412747 | 15 | 42994605 | T | C | 0.1368 | -0.01722 |
| rs11630330 | 15 | 43005852 | G | T | 0.1567 | -0.01893 |
| rs4338775 | 15 | 42980100 | T | C | 0.1353 | -0.01722 |
| rs6416436 | 15 | 42996021 | T | C | 0.1361 | -0.01722 |
| rs1037990 | 15 | 43007489 | T | C | 0.1361 | -0.01716 |
| rs192937247 | 15 | 42973303 | C | T | 0.9968 | 0.1393 |
| rs9796571 | 15 | 43005659 | T | C | 0.1362 | -0.01713 |
| rs2733224 | 15 | 42913563 | G | A | 0.8904 | 0.01728 |
| rs4244590 | 15 | 42935141 | G | A | 0.1318 | -0.01704 |
| rs181155374 | 15 | 42963621 | C | A | 0.9969 | 0.1394 |
| rs7174028 | 15 | 42922561 | G | T | 0.1316 | -0.01701 |
| rs570963333 | 15 | 42925646 | G | A | 0.9985 | 0.2244 |
| rs2412746 | 15 | 42994548 | C | T | 0.1364 | -0.01698 |
| rs6493071 | 15 | 42916608 | C | G | 0.1326 | -0.01688 |
| rs8042481 | 15 | 43002020 | G | T | 0.1366 | -0.01683 |
| rs1993813 | 15 | 42981053 | C | T | 0.1357 | -0.01674 |
| rs117347505 | 15 | 42993718 | C | T | 0.9972 | 0.1122 |
| rs1040748507 | 15 | 42889826 | G | A | 0.9975 | 0.139 |
| rs12917056 | 15 | 42953491 | T | C | 0.9313 | 0.02051 |
| rs8037022 | 15 | 42965277 | C | T | 0.1349 | -0.0166 |
| rs78563306 | 15 | 42967694 | C | T | 0.9972 | 0.1111 |
| rs3917223 | 15 | 42963993 | T | C | 0.9244 | 0.01916 |
| rs184036150 | 15 | 42934772 | A | C | 0.9972 | 0.1095 |
| rs2016538 | 15 | 42933901 | A | T | 0.85 | 0.0145 |
| rs1972447 | 15 | 42960907 | C | T | 0.1345 | -0.01643 |
| rs16957244 | 15 | 42873831 | C | T | 0.8897 | 0.01635 |
| rs60763334 | 15 | 42902516 | G | A | 0.8892 | 0.01631 |
| rs76232023 | 15 | 42893541 | C | A | 0.9968 | 0.09326 |
| rs3803341 | 15 | 42945005 | T | C | 0.1338 | -0.016 |
| rs2899069 | 15 | 42994373 | C | T | 0.1301 | -0.0164 |
| rs373428985 | 15 | 42893814 | T | C | 0.9987 | 0.1577 |
| rs6493074 | 15 | 43004985 | A | C | 0.1301 | -0.01636 |
| rs66530524 | 15 | 42880097 | T | C | 0.8888 | 0.01588 |
| rs9920498 | 15 | 42985753 | G | C | 0.1292 | -0.01633 |
| rs2126602 | 15 | 43009278 | T | C | 0.1305 | -0.01628 |
| rs2682073 | 15 | 42904360 | T | C | 0.8885 | 0.01583 |
| rs7162939 | 15 | 42938472 | C | T | 0.1344 | -0.0156 |
| rs141583312 | 15 | 42975909 | G | A | 0.9984 | 0.1441 |
| rs150316748 | 15 | 42901223 | G | A | 0.9963 | 0.0977 |
| rs150557750 | 15 | 42972607 | G | A | 0.9009 | 0.0167 |
| rs7162959 | 15 | 42938552 | A | G | 0.1342 | -0.01537 |
| rs187345619 | 15 | 42919369 | T | C | 0.998 | -0.1533 |
| rs144575157 | 15 | 42971564 | A | T | 0.9958 | 0.07335 |
| rs116238461 | 15 | 43001328 | T | C | 0.9996 | -0.3848 |
| rs141919417 | 15 | 42901329 | T | A | 0.9946 | 0.07715 |
| rs114969207 | 15 | 42891216 | G | A | 0.9998 | 0.579 |
| rs4924699 | 15 | 42908911 | G | A | 0.8568 | 0.01265 |
| rs115309896 | 15 | 42929461 | T | C | 0.9995 | -0.3746 |
| rs8031767 | 15 | 42918707 | C | T | 0.1289 | -0.01435 |
| rs72719486 | 15 | 42897273 | C | T | 0.8579 | 0.01227 |
| rs34474101 | 15 | 42895262 | A | G | 0.8581 | 0.01226 |
| rs35326878 | 15 | 42983379 | C | G | 0.858 | 0.01231 |
| rs139251799 | 15 | 42918317 | T | C | 0.9998 | 0.5553 |
| rs4923954 | 15 | 42969910 | C | G | 0.8578 | 0.01224 |
| rs72721511 | 15 | 42994161 | T | C | 0.8588 | 0.01221 |
| rs116103891 | 15 | 42955577 | A | T | 0.9998 | 0.5472 |
| rs12910799 | 15 | 42982666 | G | T | 0.8589 | 0.01211 |
| rs150988986 | 15 | 42924675 | A | T | 0.9998 | 0.547 |
| rs55805543 | 15 | 42868205 | T | C | 0.8933 | 0.01394 |
| rs12911334 | 15 | 43013055 | G | A | 0.8592 | 0.01209 |
| rs12914380 | 15 | 42950070 | C | T | 0.8588 | 0.01192 |
| rs146708171 | 15 | 42979312 | T | C | 0.9958 | 0.0671 |
| rs74873423 | 15 | 42924611 | C | G | 0.9958 | 0.06602 |
| rs117710203 | 15 | 42953008 | C | T | 0.9957 | 0.06583 |
| rs16957250 | 15 | 42878625 | C | T | 0.8962 | 0.01372 |
| rs61069183 | 15 | 42890153 | T | C | 0.9956 | 0.06328 |
| rs138218565 | 15 | 42953167 | A | C | 0.9966 | 0.07429 |
| rs138077192 | 15 | 42985240 | G | C | 0.9961 | 0.06584 |
| rs146503263 | 15 | 42878832 | T | G | 0.9962 | -0.08278 |
| rs184616495 | 15 | 43004660 | C | T | 0.992 | 0.05134 |
| rs567161987 | 15 | 42914001 | G | A | 0.9962 | 0.06551 |
| rs560519383 | 15 | 42994134 | T | C | 0.998 | -0.1272 |
| rs186041641 | 15 | 42871238 | A | G | 0.9959 | 0.06024 |
| rs143800868 | 15 | 42925865 | G | A | 0.997 | -0.09273 |
| rs75773646 | 15 | 42921982 | A | G | 0.9973 | -0.09574 |
| rs75487201 | 15 | 42921911 | T | G | 0.9973 | -0.09543 |
| rs182320019 | 15 | 42908120 | A | G | 0.9993 | -0.1442 |
| rs139393261 | 15 | 42922092 | C | T | 0.9991 | -0.1201 |
| rs139550877 | 15 | 42924878 | G | T | 0.9991 | -0.1201 |
| rs149889485 | 15 | 42913969 | T | A | 0.9914 | 0.04328 |
| rs2054388 | 15 | 42932260 | C | T | 0.998 | -0.1028 |
| rs141111379 | 15 | 42935162 | C | T | 0.9921 | 0.04381 |
| rs561908498 | 15 | 42925887 | C | G | 0.9998 | -0.411 |
| rs369926923 | 15 | 42979041 | C | T | 0.9915 | 0.04283 |
| rs547468800 | 15 | 42991255 | G | A | 0.9996 | -0.2814 |
| rs571019011 | 15 | 42870769 | GA | G | 0.9791 | -0.03167 |
| rs185505090 | 15 | 43006659 | G | T | 0.9987 | -0.1353 |
| rs540626947 | 15 | 42917314 | A | T | 0.9986 | 0.09642 |
| rs201384130 | 15 | 42964023 | T | G | 0.9998 | -0.3482 |
| rs546580872 | 15 | 42929355 | A | G | 0.9992 | -0.1174 |
| rs149836490 | 15 | 42950127 | C | T | 0.9998 | -0.2778 |
| rs78769997 | 15 | 42922138 | C | G | 0.9972 | -0.08125 |
| rs116491117 | 15 | 42921767 | G | A | 0.997 | -0.07929 |
| rs7164041 | 15 | 42917832 | A | G | 0.997 | -0.07894 |
| rs113183563 | 15 | 42898438 | G | A | 0.9994 | 0.2223 |
| rs145037353 | 15 | 42912432 | C | A | 0.999 | -0.1052 |
| rs9806313 | 15 | 42884268 | G | A | 0.9966 | -0.0743 |
| rs142838405 | 15 | 42934675 | G | A | 0.997 | -0.08081 |
| rs143043128 | 15 | 42936058 | C | T | 0.9991 | -0.1051 |
| rs116044972 | 15 | 42939606 | G | T | 0.9991 | -0.1046 |
| rs933480389 | 15 | 42942363 | C | G | 0.9989 | 0.1385 |
| rs150630012 | 15 | 42942394 | C | T | 0.9987 | -0.09538 |
| rs558747146 | 15 | 42880208 | G | A | 0.9984 | -0.09502 |
| rs114954873 | 15 | 42920847 | C | A | 0.9993 | 0.1923 |
| rs115507310 | 15 | 42920848 | T | G | 0.9993 | 0.1923 |
| rs59276064 | 15 | 42919636 | G | A | 0.9979 | -0.08537 |
| rs147870890 | 15 | 42925732 | A | G | 0.9969 | -0.07307 |
| rs114454965 | 15 | 43009037 | C | T | 0.9987 | -0.09301 |
| rs75776391 | 15 | 43011820 | G | A | 0.999 | -0.09914 |
| rs377202784 | 15 | 42920633 | TGGC | T | 0.9971 | -0.08471 |
| rs561140781 | 15 | 42956307 | T | G | 0.9979 | -0.0841 |
| rs116640993 | 15 | 42972421 | T | C | 0.999 | -0.09833 |
| rs7179367 | 15 | 43005074 | C | T | 0.9986 | -0.09173 |
| rs543516155 | 15 | 42974900 | T | C | 0.9998 | -0.2356 |
| rs57880396 | 15 | 42912858 | G | A | 0.9973 | -0.07424 |
| rs539214491 | 15 | 42937269 | G | A | 0.9998 | -0.2341 |
| rs540683576 | 15 | 42943926 | T | A | 0.9998 | -0.2341 |
| rs113476707 | 15 | 42878194 | A | G | 0.9968 | -0.07059 |
| rs553452245 | 15 | 42908200 | G | T | 0.9992 | -0.1075 |
| rs140959617 | 15 | 42958045 | A | G | 0.9975 | -0.05876 |
| rs186133585 | 15 | 42908179 | C | T | 0.9998 | -0.302 |
| rs141998787 | 15 | 42913448 | A | G | 0.9969 | -0.07141 |
| rs7176123 | 15 | 42947732 | G | A | 0.9987 | -0.09116 |
| rs139987841 | 15 | 42965625 | C | T | 0.9988 | -0.09608 |
| rs9806161 | 15 | 42883890 | A | T | 0.9968 | -0.06973 |
| rs559045802 | 15 | 42906458 | A | G | 0.9994 | 0.1701 |
| rs76206222 | 15 | 42907919 | A | T | 0.8762 | 0.009293 |
| rs74570780 | 15 | 42908305 | C | T | 0.9969 | -0.069 |
| rs566413696 | 15 | 42983841 | T | C | 0.9992 | 0.179 |
| rs189526440 | 15 | 42907138 | T | A | 0.9987 | -0.08892 |
| rs115922627 | 15 | 42907139 | C | A | 0.9987 | -0.08892 |
| rs146056091 | 15 | 42986205 | A | G | 0.9989 | 0.1388 |
| rs138000286 | 15 | 42952142 | C | T | 0.986 | 0.02623 |
| rs142211421 | 15 | 42884940 | G | A | 0.9998 | -0.2339 |
| rs75658696 | 15 | 42978682 | T | C | 0.9938 | 0.04178 |
| rs59589598 | 15 | 42884810 | G | A | 0.9987 | -0.08828 |
| rs144884699 | 15 | 42914174 | C | T | 0.972 | 0.01778 |
| rs188034879 | 15 | 42979637 | G | A | 0.9959 | -0.0506 |
| rs75580066 | 15 | 42877838 | A | G | 0.9972 | -0.06961 |
| rs148174299 | 15 | 42993741 | G | A | 0.999 | -0.09015 |
| rs543589659 | 15 | 42960960 | A | C | 0.9995 | -0.1839 |
| rs114804224 | 15 | 42987662 | G | A | 0.9986 | -0.08423 |
| rs77337431 | 15 | 42909774 | G | A | 0.9991 | -0.09311 |
| rs547298098 | 15 | 42913562 | C | T | 0.9992 | -0.1012 |
| rs147060258 | 15 | 42890797 | GAA | G | 0.9991 | -0.09349 |
| rs552124488 | 15 | 42973990 | G | A | 0.9989 | 0.1348 |
| rs117337065 | 15 | 42903762 | G | A | 0.9982 | -0.08304 |
| rs8031111 | 15 | 42911544 | C | T | 0.9974 | -0.07142 |
| rs139408969 | 15 | 42990087 | C | T | 0.999 | -0.0879 |
| rs139359398 | 15 | 42909794 | T | C | 0.9991 | -0.1201 |
| rs116738094 | 15 | 42868777 | G | A | 0.998 | -0.07467 |
| rs141795624 | 15 | 42979926 | G | A | 0.9982 | -0.07812 |
| rs572544151 | 15 | 42902719 | G | A | 0.9998 | -0.2894 |
| rs550575683 | 15 | 42960132 | T | G | 0.9996 | -0.1823 |
| rs28858165 | 15 | 42885094 | G | A | 0.9709 | 0.01605 |
| rs187287466 | 15 | 42887754 | C | A | 0.9992 | -0.09258 |
| rs184351682 | 15 | 42944285 | T | C | 0.9988 | 0.1315 |
| rs28392857 | 15 | 42883751 | C | T | 0.9709 | 0.01598 |
| rs76841629 | 15 | 43005378 | A | G | 0.5371 | -0.006656 |
| rs28674880 | 15 | 42871114 | G | T | 0.9708 | 0.01589 |
| rs144595138 | 15 | 43011980 | G | A | 0.9986 | -0.07836 |
| rs28649962 | 15 | 42898433 | T | C | 0.9708 | 0.01571 |
| rs140433299 | 15 | 42910322 | T | C | 0.9996 | -0.184 |
| rs115723709 | 15 | 42962045 | C | T | 0.9987 | -0.08115 |
| rs16957264 | 15 | 42906232 | C | T | 0.9098 | 0.009129 |
| rs569388456 | 15 | 42980580 | C | T | 0.9998 | 0.2791 |
| rs185098077 | 15 | 42912675 | C | T | 0.9998 | 0.3117 |
| rs562869855 | 15 | 42948721 | A | G | 0.9974 | 0.05876 |
| rs55683827 | 15 | 42905457 | A | G | 0.9955 | -0.0438 |
| rs143699768 | 15 | 42996145 | C | CT | 0.9393 | 0.01117 |
| rs58483557 | 15 | 42956876 | C | A | 0.998 | -0.1002 |
| rs118074434 | 15 | 43003277 | C | T | 0.9992 | -0.1161 |
| rs544535807 | 15 | 42973310 | T | G | 0.9993 | -0.1155 |
| rs185738711 | 15 | 42981245 | A | T | 0.9993 | -0.1155 |
| rs535405759 | 15 | 43005328 | G | A | 0.9987 | 0.1125 |
| rs6493070 | 15 | 42900946 | A | G | 0.08951 | 0.009443 |
| rs556862650 | 15 | 42989496 | A | C | 0.9996 | -0.1796 |
| rs543192039 | 15 | 42914069 | C | T | 0.9989 | 0.09325 |
| rs561617085 | 15 | 42986011 | A | G | 0.9823 | 0.01926 |
| rs140735548 | 15 | 42893937 | C | G | 0.9935 | 0.04006 |
| rs182927920 | 15 | 42967984 | T | C | 0.9982 | -0.07906 |
| rs371849516 | 15 | 42947665 | GA | G | 0.9863 | 0.02225 |
| rs191423170 | 15 | 42971334 | C | T | 0.9954 | -0.03883 |
| rs562725261 | 15 | 42973598 | C | T | 0.9997 | 0.1655 |
| rs534274491 | 15 | 43006688 | C | T | 0.9992 | -0.09057 |
| rs16957277 | 15 | 42945374 | T | C | 0.9692 | 0.01388 |
| rs16957284 | 15 | 42955369 | T | C | 0.998 | -0.09306 |
| rs9920231 | 15 | 42917576 | G | A | 0.9757 | 0.01523 |
| rs28575571 | 15 | 42924988 | G | A | 0.9757 | 0.01523 |
| rs557339939 | 15 | 42929212 | G | A | 0.9983 | -0.07523 |
| rs28664053 | 15 | 42922689 | G | A | 0.9757 | 0.01518 |
| rs140878173 | 15 | 42911799 | G | C | 0.9975 | 0.04772 |
| rs56356401 | 15 | 42954663 | T | C | 0.9793 | -0.01712 |
| rs537550535 | 15 | 42971805 | A | G | 0.9983 | -0.07635 |
| rs17776339 | 15 | 42968180 | T | A | 0.9793 | -0.01701 |
| rs28627318 | 15 | 42893859 | A | G | 0.9755 | 0.01492 |
| rs141227308 | 15 | 42952389 | G | A | 0.9983 | 0.05496 |
| rs185463000 | 15 | 42915407 | G | A | 0.9989 | -0.1082 |
| rs11070383 | 15 | 42887165 | G | T | 0.0658 | 0.01085 |
| rs182366902 | 15 | 42922724 | A | C | 0.9946 | -0.03397 |
| rs567039670 | 15 | 42963819 | T | C | 0.9989 | -0.1074 |
| rs115342811 | 15 | 42881852 | C | T | 0.9993 | -0.1543 |
| rs190822677 | 15 | 42989502 | G | A | 0.9995 | 0.1055 |
| rs114245127 | 15 | 42972868 | C | A | 0.9993 | -0.1481 |
| rs562977445 | 15 | 42947538 | T | C | 0.9816 | -0.02295 |
| rs192592711 | 15 | 42877415 | G | A | 0.9995 | 0.1057 |
| rs572970295 | 15 | 42930764 | C | A | 0.9931 | -0.02811 |
| rs17776090 | 15 | 42943386 | A | G | 0.9752 | -0.01554 |
| rs145057608 | 15 | 42915246 | C | T | 0.9984 | 0.05997 |
| rs138619967 | 15 | 42926998 | C | T | 0.9817 | 0.01724 |
| rs552139984 | 15 | 42981736 | G | A | 0.9979 | -0.0682 |
| rs116154059 | 15 | 42962111 | C | A | 0.9993 | -0.1433 |
| rs530091685 | 15 | 42978732 | CT | C | 0.9665 | 0.02103 |
| rs147209414 | 15 | 42897063 | G | A | 0.9987 | 0.07418 |
| rs114936782 | 15 | 42908384 | T | C | 0.998 | -0.08386 |
| rs74570754 | 15 | 42984738 | T | C | 0.9998 | -0.2365 |
| rs112929196 | 15 | 43008566 | G | A | 0.9609 | 0.01113 |
| rs140207100 | 15 | 43011559 | T | A | 0.9994 | 0.09206 |
| rs8031283 | 15 | 42903011 | G | A | 0.9483 | -0.01526 |
| rs62020703 | 15 | 42989120 | T | G | 0.9891 | 0.02259 |
| rs187236177 | 15 | 42939431 | C | A | 0.9964 | 0.04327 |
| rs79984506 | 15 | 43001654 | T | C | 0.997 | -0.04476 |
| rs191793117 | 15 | 42914340 | T | C | 0.998 | 0.07257 |
| rs192727326 | 15 | 42948390 | A | C | 0.9998 | 0.1895 |
| rs189290697 | 15 | 42931689 | C | T | 0.9968 | -0.04266 |
| rs148549752 | 15 | 42987049 | C | T | 0.9993 | 0.09526 |
| rs144640419 | 15 | 42892393 | T | A | 0.9994 | 0.08947 |
| rs376331294 | 15 | 42960653 | G | A | 0.9998 | -0.166 |
| rs138556557 | 15 | 42895364 | G | A | 0.994 | 0.02655 |
| rs117423500 | 15 | 42885260 | G | A | 0.9741 | -0.01372 |
| rs545291075 | 15 | 42871454 | T | A | 0.9997 | 0.1177 |
| rs148952135 | 15 | 42966774 | T | C | 0.9969 | -0.04127 |
| rs34436896 | 15 | 42973919 | C | T | 0.9974 | -0.04916 |
| rs146894776 | 15 | 42967945 | T | C | 0.9975 | -0.04574 |
| rs72721503 | 15 | 42968409 | G | C | 0.9726 | -0.01442 |
| rs142183834 | 15 | 43010103 | A | T | 0.9975 | -0.04545 |
| rs138963231 | 15 | 42983997 | G | C | 0.9865 | 0.01752 |
| rs139400588 | 15 | 43000387 | G | A | 0.9975 | -0.04537 |
| rs149843783 | 15 | 42935351 | C | T | 0.9998 | 0.2159 |
| rs563467704 | 15 | 42982383 | T | C | 0.9995 | -0.1402 |
| rs148438272 | 15 | 42918775 | G | T | 0.9977 | 0.05899 |
| rs12912026 | 15 | 42878529 | T | A | 0.7587 | -0.007624 |
| rs998954 | 15 | 42988571 | C | T | 0.9748 | 0.01207 |
| rs142202313 | 15 | 42996605 | A | C | 0.9975 | -0.04458 |
| rs151022817 | 15 | 42910971 | G | A | 0.9988 | 0.06666 |
| rs184008832 | 15 | 42890930 | G | A | 0.9984 | 0.04539 |
| rs147153494 | 15 | 42956820 | C | T | 0.9991 | 0.1069 |
| rs562509996 | 15 | 43005091 | G | T | 0.9987 | -0.0642 |
| rs541359954 | 15 | 42880415 | C | G | 0.9978 | 0.05635 |
| rs562776360 | 15 | 43002939 | T | C | 0.9996 | 0.1037 |
| rs541817722 | 15 | 42940819 | C | T | 0.9984 | 0.0446 |
| rs186771984 | 15 | 42950609 | C | T | 0.9984 | 0.0441 |
| rs191526970 | 15 | 42957820 | T | A | 0.9998 | -0.1565 |
| rs569234006 | 15 | 42940808 | G | GT | 0.9985 | 0.04503 |
| rs193196275 | 15 | 42908780 | A | G | 0.9974 | -0.04164 |
| rs115840904 | 15 | 42966913 | C | T | 0.998 | -0.04817 |
| rs139693200 | 15 | 42942444 | C | G | 0.9979 | -0.04732 |
| rs112362877 | 15 | 42938705 | G | A | 0.9481 | 0.007661 |
| rs573802972 | 15 | 42925909 | G | A | 0.9984 | -0.07516 |
| rs191518723 | 15 | 42922665 | G | T | 0.9934 | 0.0299 |
| rs763442891 | 15 | 42878527 | A | ACT | 0.9988 | 0.04748 |
| rs181239306 | 15 | 42916138 | T | C | 0.9993 | -0.08572 |
| rs185103862 | 15 | 43003994 | C | T | 0.9985 | 0.04245 |
| rs144230475 | 15 | 42987511 | C | T | 0.9995 | -0.1132 |
| rs147284877 | 15 | 42977716 | T | C | 0.9989 | 0.073 |
| rs150514250 | 15 | 42999694 | G | A | 0.9992 | 0.0724 |
| rs143378302 | 15 | 42886009 | T | A | 0.9996 | -0.1389 |
| rs113913915 | 15 | 42970124 | T | C | 0.9995 | -0.1099 |
| rs13380212 | 15 | 43006842 | T | C | 0.9992 | -0.0964 |
| rs76864255 | 15 | 42971050 | T | A | 0.9979 | -0.04452 |
| rs1984501 | 15 | 42987512 | G | A | 0.9781 | 0.01106 |
| rs117095901 | 15 | 42912799 | C | T | 0.9888 | -0.01527 |
| rs77496195 | 15 | 42967615 | T | G | 0.9983 | -0.04557 |
| rs547972042 | 15 | 43005198 | G | A | 0.9944 | -0.02492 |
| rs150358878 | 15 | 42952069 | A | T | 0.9979 | -0.04512 |
| rs199781595 | 15 | 42941307 | A | G | 0.9997 | 0.1313 |
| rs146488861 | 15 | 42957808 | A | G | 0.9958 | 0.02769 |
| rs77547387 | 15 | 42997595 | G | A | 0.9992 | -0.09253 |
| rs186206742 | 15 | 42922541 | C | A | 0.9921 | 0.0169 |
| rs77541575 | 15 | 42975754 | C | T | 0.9981 | -0.04272 |
| rs116640263 | 15 | 42981739 | C | T | 0.9981 | -0.04264 |
| rs187320661 | 15 | 42934800 | T | C | 0.9996 | -0.0827 |
| rs142424434 | 15 | 42925466 | G | C | 0.9996 | -0.0824 |
| rs143356796 | 15 | 43010938 | C | T | 0.9687 | -0.01025 |
| rs117347582 | 15 | 42998252 | C | A | 0.9977 | -0.03046 |
| rs117252626 | 15 | 42953935 | T | C | 0.9991 | 0.06508 |
| rs114026095 | 15 | 42973492 | C | A | 0.9982 | -0.04198 |
| rs192668256 | 15 | 42999153 | C | T | 0.9994 | 0.1003 |
| rs544857253 | 15 | 42904460 | G | C | 0.9998 | -0.1117 |
| rs142696672 | 15 | 42910808 | G | T | 0.9988 | -0.04492 |
| rs78948790 | 15 | 42958014 | T | C | 0.9923 | 0.01632 |
| rs544054828 | 15 | 42977780 | CAA | C | 0.9993 | 0.07953 |
| rs11070384 | 15 | 42983068 | A | G | 0.9991 | -0.08564 |
| rs771755844 | 15 | 42933197 | C | A | 0.9998 | -0.1203 |
| rs149265736 | 15 | 43008367 | C | T | 0.9993 | 0.05694 |
| rs181498789 | 15 | 43010216 | A | G | 0.9993 | 0.05694 |
| rs183189070 | 15 | 42979608 | T | C | 0.9992 | 0.05644 |
| rs76783611 | 15 | 42988659 | C | T | 0.9981 | -0.04009 |
| rs150970024 | 15 | 42987339 | C | G | 0.9993 | 0.05626 |
| rs142290993 | 15 | 42978320 | T | C | 0.9993 | 0.05582 |
| rs186564556 | 15 | 42902293 | G | C | 0.9996 | -0.07376 |
| rs373905733 | 15 | 42906618 | G | C | 0.9996 | -0.07376 |
| rs150495511 | 15 | 42899170 | A | AT | 0.9182 | 0.007211 |
| rs114066074 | 15 | 42927698 | A | G | 0.9993 | 0.05537 |
| rs561774462 | 15 | 42931411 | C | T | 0.9987 | 0.04412 |
| rs56071832 | 15 | 42988453 | C | G | 0.9993 | 0.09487 |
| rs148386184 | 15 | 42954593 | A | T | 0.9906 | -0.01503 |
| rs118104470 | 15 | 42913334 | C | T | 0.9904 | -0.01479 |
| rs76989793 | 15 | 42922584 | A | T | 0.99 | -0.01455 |
| rs34312217 | 15 | 42902986 | CA | C | 0.931 | 0.00776 |
| rs16957292 | 15 | 42982370 | C | T | 0.9992 | -0.08125 |
| rs546077226 | 15 | 42903738 | G | GT | 0.9981 | 0.03478 |
| rs563964635 | 15 | 42893798 | A | G | 0.9969 | 0.0299 |
| rs143455945 | 15 | 42965670 | T | C | 0.9992 | -0.078 |
| rs140916237 | 15 | 42889531 | T | C | 0.9994 | -0.08492 |
| rs548296727 | 15 | 42970157 | T | C | 0.996 | 0.03606 |
| rs117122292 | 15 | 42879243 | G | A | 0.9818 | -0.009885 |
| rs80258074 | 15 | 42923044 | T | A | 0.9995 | -0.08466 |
| rs76215594 | 15 | 42926099 | G | T | 0.9995 | -0.08466 |
| rs116269151 | 15 | 42919023 | C | T | 0.9995 | -0.08336 |
| rs188362035 | 15 | 42918484 | G | A | 0.9972 | 0.02773 |
| rs144693269 | 15 | 42898455 | G | T | 0.9754 | -0.008593 |
| rs552314563 | 15 | 43004920 | G | A | 0.9934 | -0.0164 |
| rs149119473 | 15 | 42914260 | G | T | 0.9995 | -0.08016 |
| rs559628380 | 15 | 42919443 | G | T | 0.9948 | 0.01643 |
| rs191375171 | 15 | 42888109 | C | T | 0.9972 | 0.02758 |
| rs117193101 | 15 | 42875336 | G | T | 0.9953 | 0.01785 |
| rs151031682 | 15 | 42934972 | G | A | 0.9903 | 0.01225 |
| rs201091488 | 15 | 42984985 | T | C | 0.9996 | 0.06043 |
| rs7179275 | 15 | 43004823 | G | T | 0.9971 | -0.03198 |
| rs530032514 | 15 | 42907916 | A | T | 0.9996 | -0.08623 |
| rs142975069 | 15 | 42981529 | G | A | 0.9983 | 0.03846 |
| rs116042800 | 15 | 42887128 | C | G | 0.9995 | -0.0761 |
| rs138835702 | 15 | 42897067 | T | C | 0.9995 | -0.0761 |
| rs541474906 | 15 | 42968291 | A | G | 0.9981 | -0.03185 |
| rs548547947 | 15 | 42947931 | C | T | 0.9934 | -0.01642 |
| rs572583293 | 15 | 42993853 | C | G | 0.9957 | 0.02281 |
| rs535809709 | 15 | 42883375 | C | T | 0.9998 | -0.1201 |
| rs546446869 | 15 | 42932919 | C | T | 0.9997 | -0.09179 |
| rs557473381 | 15 | 43000769 | G | A | 0.9995 | -0.05011 |
| rs1049733221 | 15 | 42914070 | G | A | 0.9997 | -0.09508 |
| rs138592907 | 15 | 42946486 | C | T | 0.9956 | -0.01702 |
| rs143568757 | 15 | 42902686 | G | A | 0.997 | -0.0261 |
| rs78365925 | 15 | 42905493 | G | C | 0.9984 | 0.04297 |
| rs11632981 | 15 | 43000340 | A | C | 0.9776 | -0.00779 |
| rs141828250 | 15 | 42945444 | C | T | 0.9987 | -0.02874 |
| rs1197534 | 15 | 42907922 | T | A | 0.9579 | -0.006048 |
| rs529234877 | 15 | 42924755 | A | G | 0.9998 | -0.08834 |
| rs80305987 | 15 | 42877994 | A | G | 0.9899 | 0.01089 |
| rs558292498 | 15 | 42911675 | C | T | 0.9994 | -0.04533 |
| rs567155071 | 15 | 42903081 | G | A | 0.9963 | -0.01723 |
| rs148028047 | 15 | 42903243 | T | C | 0.9989 | 0.04079 |
| rs577021890 | 15 | 42891221 | G | A | 0.999 | -0.04493 |
| rs547814549 | 15 | 42964424 | G | A | 0.9996 | 0.06508 |
| rs79495360 | 15 | 42918441 | T | C | 0.9932 | -0.01797 |
| rs151019978 | 15 | 42991740 | T | C | 0.9981 | -0.03236 |
| rs572827078 | 15 | 42913020 | G | T | 0.9994 | -0.04087 |
| rs551010070 | 15 | 42929055 | A | T | 0.9993 | 0.04536 |
| rs189542431 | 15 | 42974675 | C | G | 0.9966 | 0.01909 |
| rs570080968 | 15 | 43013157 | G | A | 0.9998 | 0.114 |
| rs149328464 | 15 | 42887671 | T | C | 0.9983 | 0.03063 |
| rs558490453 | 15 | 42935340 | G | A | 0.9979 | 0.02462 |
| rs776170188 | 15 | 42982418 | G | A | 0.9998 | 0.09729 |
| rs143766227 | 15 | 42883585 | CAA | C | 0.9951 | 0.02156 |
| rs141311377 | 15 | 42912418 | A | C | 0.9993 | -0.03624 |
| rs145466527 | 15 | 42940296 | G | A | 0.9993 | -0.03624 |
| rs549786810 | 15 | 42931943 | G | A | 0.9984 | 0.0362 |
| rs576834861 | 15 | 42877871 | G | A | 0.9996 | 0.06134 |
| rs73413058 | 15 | 42988269 | A | G | 0.9369 | 0.00661 |
| rs181630879 | 15 | 42889666 | G | C | 0.9979 | -0.0281 |
| rs189693176 | 15 | 42950613 | A | G | 0.9998 | 0.08779 |
| rs144326298 | 15 | 42990770 | T | C | 0.999 | 0.04955 |
| rs1115752 | 15 | 43008863 | C | T | 0.9061 | -0.002924 |
| rs148848820 | 15 | 42895452 | T | C | 0.9816 | -0.007342 |
| rs776118491 | 15 | 42993735 | G | A | 0.9794 | -0.006211 |
| rs1037982814 | 15 | 42900693 | C | T | 0.9998 | 0.08022 |
| rs111339056 | 15 | 42872121 | C | T | 0.9998 | -0.09208 |
| rs111569959 | 15 | 42874184 | A | T | 0.9998 | -0.09208 |
| rs34714662 | 15 | 43001896 | C | T | 0.7847 | 0.002075 |
| rs190806658 | 15 | 42940285 | A | G | 0.9974 | -0.02475 |
| rs138942621 | 15 | 42930975 | T | C | 0.9995 | -0.04893 |
| rs568614887 | 15 | 42947941 | G | A | 0.997 | 0.01546 |
| rs111850927 | 15 | 42921045 | T | A | 0.9998 | -0.08252 |
| rs74533898 | 15 | 42909616 | G | A | 0.9971 | 0.01911 |
| rs187610922 | 15 | 43004989 | C | T | 0.9994 | 0.03997 |
| rs112780488 | 15 | 42975207 | T | C | 0.9998 | -0.08143 |
| rs184380146 | 15 | 42995397 | C | T | 0.9987 | -0.03378 |
| rs549942856 | 15 | 42877419 | C | T | 0.9998 | 0.06331 |
| rs141585079 | 15 | 42909609 | A | G | 0.9671 | 0.003947 |
| rs142301748 | 15 | 42884365 | C | T | 0.9959 | 0.01365 |
| rs552525489 | 15 | 42956466 | C | T | 0.9989 | 0.02441 |
| rs180750404 | 15 | 42912673 | C | T | 0.998 | -0.0191 |
| rs142005131 | 15 | 42882373 | C | A | 0.9714 | 0.00425 |
| rs180994451 | 15 | 42939263 | A | G | 0.9974 | 0.01805 |
| rs145938839 | 15 | 42884633 | C | A | 0.9981 | 0.02317 |
| rs191544070 | 15 | 42974511 | A | C | 0.9958 | 0.01066 |
| rs568462372 | 15 | 42924192 | C | G | 0.9998 | 0.0548 |
| rs551671670 | 15 | 42927127 | T | C | 0.9998 | 0.0548 |
| rs556243779 | 15 | 42902209 | C | A | 0.9986 | -0.02156 |
| rs191443517 | 15 | 43005135 | C | T | 0.9972 | 0.01396 |
| rs180684653 | 15 | 42982257 | A | G | 0.999 | -0.02128 |
| rs192977893 | 15 | 42887615 | C | T | 0.9986 | 0.01833 |
| rs554508188 | 15 | 42931509 | C | T | 0.9993 | -0.02634 |
| rs571185339 | 15 | 42879699 | A | G | 0.9995 | 0.03168 |
| rs146799639 | 15 | 42870779 | T | C | 0.9416 | 0.002289 |
| rs191933065 | 15 | 42946188 | G | C | 0.9994 | -0.02704 |
| rs375380714 | 15 | 42998215 | T | G | 0.9998 | -0.03966 |
| rs571167694 | 15 | 43011992 | C | T | 0.999 | -0.01905 |
| rs191535749 | 15 | 42985222 | C | G | 0.9998 | 0.04721 |
| rs183231149 | 15 | 42887050 | G | A | 0.9992 | -0.02397 |
| rs552385948 | 15 | 43007566 | T | A | 0.9993 | 0.0295 |
| rs556210813 | 15 | 42997532 | T | A | 0.9998 | 0.04406 |
| rs192718382 | 15 | 42905736 | C | T | 0.998 | -0.01142 |
| rs200325235 | 15 | 43010236 | CA | C | 0.9641 | 0.002709 |
| rs181075931 | 15 | 42884135 | G | A | 0.997 | -0.01024 |
| rs185850923 | 15 | 42993431 | T | C | 0.9947 | -0.008331 |
| rs552175940 | 15 | 42921383 | C | G | 0.9986 | -0.02003 |
| rs145449308 | 15 | 42997265 | C | G | 0.9997 | 0.0317 |
| rs574471910 | 15 | 42971058 | T | G | 0.9985 | -0.01949 |
| rs537741020 | 15 | 42936512 | A | C | 0.9997 | 0.03415 |
| rs78084245 | 15 | 42963434 | A | G | 0.9722 | 0.00278 |
| rs142285781 | 15 | 43007204 | G | A | 0.9991 | -0.01345 |
| rs183621980 | 15 | 42884021 | G | A | 0.9952 | 0.006314 |
| rs534379603 | 15 | 42894853 | G | A | 0.9997 | 0.03712 |
| rs537461773 | 15 | 42923097 | G | C | 0.9993 | 0.01658 |
| rs150203226 | 15 | 42875831 | G | A | 0.9684 | 0.002063 |
| rs148534148 | 15 | 42934574 | G | T | 0.9972 | 0.008073 |
| rs541558127 | 15 | 42939846 | T | C | 0.9964 | -0.007527 |
| rs190984235 | 15 | 42929593 | A | C | 0.9958 | -0.00576 |
| rs138976273 | 15 | 42902170 | C | T | 0.9865 | 0.002997 |
| rs117725364 | 15 | 42998580 | A | C | 0.992 | -0.004061 |
| rs117017655 | 15 | 42872405 | G | C | 0.9729 | 0.002234 |
| rs151233914 | 15 | 42951943 | T | C | 0.9977 | 0.01055 |
| rs557668526 | 15 | 42931262 | T | C | 0.9989 | 0.0112 |
| rs557349180 | 15 | 42993733 | G | A | 0.9851 | 0.003185 |
| rs556811858 | 15 | 42993734 | AG | A | 0.9851 | 0.003185 |
| rs567241776 | 15 | 42927294 | C | T | 0.9994 | -0.01817 |
| rs147891025 | 15 | 42885547 | G | A | 0.9974 | -0.006081 |
| rs150770038 | 15 | 42897487 | C | T | 0.9974 | -0.006081 |
| rs141900598 | 15 | 42885895 | C | T | 0.9912 | 0.004346 |
| rs562676922 | 15 | 42876073 | C | T | 0.9996 | -0.02462 |
| rs570044324 | 15 | 42983733 | T | C | 0.9991 | -0.01317 |
| rs574816840 | 15 | 42987187 | C | T | 0.9979 | 0.006321 |
| rs182906122 | 15 | 42953382 | C | T | 0.9942 | 0.00347 |
| rs181004974 | 15 | 42874762 | C | T | 0.9933 | -0.003141 |
| rs536514977 | 15 | 42875884 | C | T | 0.9987 | -0.009158 |
| rs181258192 | 15 | 42950910 | G | A | 0.9979 | 0.005625 |
| rs185596875 | 15 | 42894226 | T | C | 0.999 | 0.01288 |
| rs145034051 | 15 | 42992977 | T | C | 0.9974 | -0.004652 |
| rs375843171 | 15 | 42983341 | T | C | 0.9981 | -0.005242 |
| rs73413030 | 15 | 42897626 | A | G | 0.998 | 0.00719 |
| rs543230721 | 15 | 42912342 | C | T | 0.9981 | -0.005057 |
| rs150012114 | 15 | 42924974 | A | G | 0.9771 | -0.001332 |
| rs149678972 | 15 | 42873237 | G | A | 0.9732 | 0.001352 |
| rs189618809 | 15 | 42907976 | C | A | 0.9957 | 0.003078 |
| rs145784520 | 15 | 42966451 | C | T | 0.9998 | 0.01632 |
| rs182889305 | 15 | 42998734 | C | T | 0.9976 | 0.003461 |
| rs187329646 | 15 | 42983026 | C | A | 0.9997 | 0.009988 |
| rs143837600 | 15 | 42981862 | G | T | 0.9997 | 0.009903 |
| rs140450055 | 15 | 42961579 | C | T | 0.9876 | 0.001421 |
| rs142774139 | 15 | 42874805 | G | A | 0.993 | 0.001876 |
| rs143251532 | 15 | 42998512 | T | C | 0.9996 | 0.01035 |
| rs200262486 | 15 | 42978421 | GT | G | 0.992 | -0.001775 |
| rs146248884 | 15 | 42965927 | A | G | 0.9955 | 0.002156 |
| rs147047460 | 15 | 42885872 | C | T | 0.999 | -0.005991 |
| rs116967439 | 15 | 42956352 | C | T | 0.9483 | -0.0004986 |
| rs148100429 | 15 | 42875632 | C | T | 0.9997 | 0.007055 |
| rs190779787 | 15 | 42885334 | C | A | 0.9997 | 0.007055 |
| rs564224994 | 15 | 42966397 | T | A | 0.9981 | -0.003015 |
| rs537102106 | 15 | 42967458 | G | C | 0.9981 | -0.003015 |
| rs139382852 | 15 | 43003552 | G | A | 0.9993 | -0.005089 |
| rs117422009 | 15 | 43011777 | C | G | 0.9903 | -0.001027 |
| rs185185807 | 15 | 42986374 | C | T | 0.9973 | 0.001801 |
| rs370980402 | 15 | 42990977 | C | T | 0.9986 | 0.003834 |
| rs147200598 | 15 | 42892930 | G | C | 0.9932 | -0.001127 |
| rs114659791 | 15 | 42885873 | G | A | 0.9998 | 0.008063 |
| rs28535059 | 15 | 42933903 | T | A | 0.9981 | -0.002453 |
| rs117722860 | 15 | 43008329 | C | T | 0.9754 | -0.0004533 |
| rs575373910 | 15 | 43005726 | C | T | 0.9825 | 0.0005863 |
| rs573769722 | 15 | 42901255 | G | A | 0.9966 | -0.001189 |
| rs542549746 | 15 | 42910957 | T | C | 0.9997 | 0.00334 |
| rs188917136 | 15 | 42936275 | G | A | 0.9952 | 0.0005616 |
| rs77231660 | 15 | 42981109 | C | T | 0.9483 | -0.0001401 |
| rs75090487 | 15 | 42937481 | T | G | 0.9944 | -0.0005366 |
| rs117350263 | 15 | 42951611 | A | G | 0.9935 | -0.0002499 |
| rs16957288 | 15 | 42958590 | T | C | 0.9745 | -0.0001003 |
| rs557783986 | 15 | 42951686 | T | C | 0.9983 | 0.0002661 |
| rs530713585 | 15 | 42912062 | G | A | 0.9998 | -0.0008493 |
| rs572834165 | 15 | 42936208 | G | A | 0.998 | 0.0001045 |

| se | p-value | INFO |
| --- | --- | --- |
| 0.009048 | 6.931E-69 | 0.8974 |
| 0.02545 | 9.121E-35 | 0.4878 |
| 0.005717 | 1.652E-30 | 0.9202 |
| 0.004782 | 1.145E-27 | 0.9997 |
| 0.004812 | 2.653E-24 | 0.9621 |
| 0.004096 | 4.302E-19 | 0.9523 |
| 0.003404 | 3.84E-10 | 0.7361 |
| 0.003326 | 5.141E-10 | 0.7872 |
| 0.003254 | 1.801E-09 | 0.8835 |
| 0.003174 | 1.911E-09 | 0.8938 |
| 0.003118 | 2.345E-09 | 0.9683 |
| 0.003218 | 2.424E-09 | 0.8902 |
| 0.003172 | 2.525E-09 | 0.8943 |
| 0.003172 | 2.553E-09 | 0.8942 |
| 0.003176 | 2.685E-09 | 0.8937 |
| 0.003176 | 2.7E-09 | 0.8937 |
| 0.003176 | 2.701E-09 | 0.8937 |
| 0.003143 | 2.98E-09 | 0.9538 |
| 0.003161 | 3.161E-09 | 0.903 |
| 0.003112 | 3.278E-09 | 0.9692 |
| 0.003156 | 3.289E-09 | 0.9045 |
| 0.003108 | 3.449E-09 | 0.9706 |
| 0.003147 | 3.48E-09 | 0.9106 |
| 0.003114 | 3.492E-09 | 0.969 |
| 0.003144 | 3.529E-09 | 0.9535 |
| 0.003144 | 3.529E-09 | 0.9534 |
| 0.003109 | 3.71E-09 | 0.9701 |
| 0.003132 | 3.869E-09 | 0.9637 |
| 0.003109 | 4.088E-09 | 0.9703 |
| 0.003113 | 4.323E-09 | 0.9697 |
| 0.003107 | 4.341E-09 | 0.9711 |
| 0.003154 | 4.369E-09 | 0.9082 |
| 0.003164 | 4.52E-09 | 0.9016 |
| 0.06402 | 4.545E-09 | 0.3547 |
| 0.003154 | 4.894E-09 | 0.9089 |
| 0.003161 | 5.53E-09 | 0.9011 |
| 0.003133 | 5.561E-09 | 0.9565 |
| 0.003154 | 6.083E-09 | 0.9095 |
| 0.003151 | 6.452E-09 | 0.91 |
| 0.003108 | 9.119E-09 | 0.9712 |
| 0.003108 | 9.888E-09 | 0.9704 |
| 0.003109 | 9.95E-09 | 0.9688 |
| 0.003103 | 1.015E-08 | 0.9716 |
| 0.003095 | 1.127E-08 | 0.9742 |
| 0.003158 | 1.159E-08 | 0.9088 |
| 0.003103 | 1.372E-08 | 0.9722 |
| 0.003049 | 1.534E-08 | 0.9943 |
| 0.003102 | 2.92E-08 | 0.9703 |
| 0.004517 | 5.137E-08 | 0.4494 |
| 0.08293 | 0.000001625 | 0.3142 |
| 0.01692 | 0.000002782 | 0.3772 |
| 0.004006 | 0.00001208 | 0.4951 |
| 0.003796 | 0.00002415 | 0.9108 |
| 0.004745 | 0.00002751 | 0.5544 |
| 0.003735 | 0.00002978 | 0.9349 |
| 0.003686 | 0.0000386 | 0.9712 |
| 0.003686 | 0.00003922 | 0.9713 |
| 0.003687 | 0.00004079 | 0.9712 |
| 0.003687 | 0.00004109 | 0.9712 |
| 0.003687 | 0.00004198 | 0.9712 |
| 0.003782 | 0.00004378 | 0.9118 |
| 0.005047 | 0.0000469 | 0.5901 |
| 0.003702 | 0.00005518 | 0.9699 |
| 0.003688 | 0.00006077 | 0.972 |
| 0.003693 | 0.00006502 | 0.973 |
| 0.004654 | 0.0000774 | 0.7893 |
| 0.004607 | 0.00008313 | 0.8044 |
| 0.004607 | 0.00008318 | 0.8044 |
| 0.003687 | 0.00008519 | 0.9708 |
| 0.0046 | 0.00008604 | 0.8105 |
| 0.003692 | 0.00008793 | 0.9742 |
| 0.003685 | 0.00008858 | 0.9731 |
| 0.004601 | 0.00009087 | 0.7946 |
| 0.003688 | 0.00009551 | 0.9744 |
| 0.004609 | 0.00009562 | 0.7894 |
| 0.003689 | 0.00009922 | 0.9742 |
| 0.004564 | 0.0001073 | 0.7917 |
| 0.005014 | 0.0001237 | 0.5962 |
| 0.004567 | 0.0001631 | 0.7901 |
| 0.005021 | 0.0001633 | 0.5839 |
| 0.004571 | 0.0001653 | 0.7958 |
| 0.004575 | 0.0001675 | 0.7909 |
| 0.004581 | 0.0001795 | 0.7888 |
| 0.03721 | 0.0001815 | 0.4462 |
| 0.004579 | 0.0001833 | 0.7888 |
| 0.004632 | 0.0001918 | 0.9295 |
| 0.004576 | 0.0001963 | 0.8119 |
| 0.03748 | 0.0002 | 0.4399 |
| 0.004574 | 0.0002005 | 0.8134 |
| 0.06036 | 0.0002007 | 0.3563 |
| 0.004568 | 0.000202 | 0.7916 |
| 0.004561 | 0.0002139 | 0.8133 |
| 0.004566 | 0.000228 | 0.7912 |
| 0.004564 | 0.0002443 | 0.7966 |
| 0.03063 | 0.0002476 | 0.7466 |
| 0.03796 | 0.0002507 | 0.5296 |
| 0.005608 | 0.0002556 | 0.9654 |
| 0.004545 | 0.00026 | 0.807 |
| 0.03047 | 0.000265 | 0.7561 |
| 0.005277 | 0.0002829 | 0.9983 |
| 0.03018 | 0.0002856 | 0.7619 |
| 0.004015 | 0.0003038 | 0.9452 |
| 0.004552 | 0.0003065 | 0.8068 |
| 0.004581 | 0.0003589 | 0.9451 |
| 0.004574 | 0.000363 | 0.9443 |
| 0.02635 | 0.0004001 | 0.8664 |
| 0.004548 | 0.0004347 | 0.8117 |
| 0.004692 | 0.0004736 | 0.7807 |
| 0.04519 | 0.0004855 | 0.712 |
| 0.004697 | 0.000496 | 0.779 |
| 0.004568 | 0.0005089 | 0.9436 |
| 0.004699 | 0.0005091 | 0.783 |
| 0.004695 | 0.0005248 | 0.778 |
| 0.004564 | 0.000525 | 0.9428 |
| 0.004532 | 0.0005765 | 0.8143 |
| 0.04195 | 0.0005944 | 0.6748 |
| 0.02845 | 0.0005945 | 0.6469 |
| 0.00489 | 0.0006372 | 0.9118 |
| 0.004536 | 0.0007039 | 0.814 |
| 0.04648 | 0.0009754 | 0.4438 |
| 0.02263 | 0.001189 | 0.9183 |
| 0.1197 | 0.001306 | 0.3415 |
| 0.02408 | 0.001356 | 0.6231 |
| 0.1812 | 0.001396 | 0.3599 |
| 0.004011 | 0.001605 | 0.9844 |
| 0.1204 | 0.001865 | 0.3121 |
| 0.004638 | 0.001978 | 0.8054 |
| 0.004013 | 0.002224 | 0.9899 |
| 0.004016 | 0.002277 | 0.9892 |
| 0.00404 | 0.002304 | 0.977 |
| 0.1826 | 0.002356 | 0.3629 |
| 0.004034 | 0.002416 | 0.9785 |
| 0.004044 | 0.00253 | 0.9799 |
| 0.1827 | 0.002742 | 0.3693 |
| 0.004043 | 0.002743 | 0.9808 |
| 0.1826 | 0.002746 | 0.3694 |
| 0.004656 | 0.002758 | 0.9414 |
| 0.004052 | 0.002856 | 0.9781 |
| 0.004038 | 0.003158 | 0.9823 |
| 0.02278 | 0.003223 | 0.897 |
| 0.02261 | 0.003505 | 0.9016 |
| 0.02259 | 0.003563 | 0.8788 |
| 0.004744 | 0.003827 | 0.93 |
| 0.02211 | 0.004205 | 0.9147 |
| 0.02596 | 0.004218 | 0.8499 |
| 0.0233 | 0.004714 | 0.9269 |
| 0.02947 | 0.004962 | 0.6047 |
| 0.01838 | 0.005221 | 0.7226 |
| 0.02359 | 0.005484 | 0.9047 |
| 0.04624 | 0.005947 | 0.4503 |
| 0.0225 | 0.007424 | 0.9462 |
| 0.03473 | 0.007587 | 0.538 |
| 0.03586 | 0.007588 | 0.5715 |
| 0.03587 | 0.007811 | 0.5714 |
| 0.05556 | 0.00947 | 0.8389 |
| 0.04772 | 0.01183 | 0.9392 |
| 0.04772 | 0.01183 | 0.9392 |
| 0.01729 | 0.01229 | 0.7713 |
| 0.04127 | 0.01272 | 0.58 |
| 0.01768 | 0.0132 | 0.7937 |
| 0.1659 | 0.01327 | 0.3121 |
| 0.01731 | 0.01333 | 0.7703 |
| 0.1142 | 0.01374 | 0.4065 |
| 0.01286 | 0.01377 | 0.5767 |
| 0.05586 | 0.01541 | 0.491 |
| 0.04067 | 0.01776 | 0.8387 |
| 0.1481 | 0.01873 | 0.4201 |
| 0.05083 | 0.02087 | 0.9456 |
| 0.1206 | 0.02125 | 0.6948 |
| 0.03529 | 0.02132 | 0.566 |
| 0.03447 | 0.02142 | 0.5452 |
| 0.03456 | 0.02235 | 0.543 |
| 0.09738 | 0.02245 | 0.3283 |
| 0.04659 | 0.02401 | 0.9218 |
| 0.03297 | 0.02423 | 0.534 |
| 0.0359 | 0.02438 | 0.5123 |
| 0.04667 | 0.02438 | 0.9353 |
| 0.04662 | 0.02492 | 0.9425 |
| 0.06291 | 0.02767 | 0.4597 |
| 0.04346 | 0.02819 | 0.7916 |
| 0.04356 | 0.02916 | 0.6373 |
| 0.08896 | 0.03062 | 0.3434 |
| 0.08896 | 0.03062 | 0.3434 |
| 0.03952 | 0.03077 | 0.6078 |
| 0.03383 | 0.03078 | 0.5501 |
| 0.04307 | 0.03082 | 0.8272 |
| 0.04602 | 0.03121 | 0.9599 |
| 0.03934 | 0.03129 | 0.4243 |
| 0.03907 | 0.03135 | 0.6145 |
| 0.04605 | 0.03276 | 0.9352 |
| 0.04305 | 0.03312 | 0.7824 |
| 0.1114 | 0.03436 | 0.802 |
| 0.03512 | 0.0345 | 0.5925 |
| 0.111 | 0.03498 | 0.8031 |
| 0.111 | 0.03498 | 0.8031 |
| 0.03352 | 0.03523 | 0.548 |
| 0.05105 | 0.03525 | 0.926 |
| 0.02793 | 0.0354 | 0.9963 |
| 0.1437 | 0.03556 | 0.3891 |
| 0.03398 | 0.03557 | 0.5534 |
| 0.04356 | 0.03636 | 0.7998 |
| 0.04618 | 0.0375 | 0.7976 |
| 0.03356 | 0.03772 | 0.5493 |
| 0.08233 | 0.03881 | 0.4661 |
| 0.004519 | 0.03975 | 0.8783 |
| 0.03365 | 0.04028 | 0.5519 |
| 0.08732 | 0.04032 | 0.3375 |
| 0.04363 | 0.04153 | 0.7828 |
| 0.04363 | 0.04153 | 0.7828 |
| 0.06811 | 0.04158 | 0.3928 |
| 0.01289 | 0.04185 | 0.8468 |
| 0.1153 | 0.04247 | 0.692 |
| 0.02061 | 0.04269 | 0.7413 |
| 0.04362 | 0.04298 | 0.7795 |
| 0.008808 | 0.04356 | 0.9241 |
| 0.02522 | 0.0448 | 0.7458 |
| 0.03475 | 0.04516 | 0.5783 |
| 0.04518 | 0.04601 | 0.9632 |
| 0.0923 | 0.04633 | 0.3832 |
| 0.0424 | 0.04696 | 0.8037 |
| 0.04706 | 0.04785 | 0.9299 |
| 0.05121 | 0.04805 | 0.9297 |
| 0.04755 | 0.04928 | 0.9011 |
| 0.06864 | 0.04955 | 0.3759 |
| 0.04238 | 0.05007 | 0.5975 |
| 0.03676 | 0.05202 | 0.5665 |
| 0.04532 | 0.05242 | 0.9724 |
| 0.06219 | 0.05341 | 0.525 |
| 0.03886 | 0.05466 | 0.6513 |
| 0.04083 | 0.05568 | 0.6406 |
| 0.1517 | 0.05648 | 0.4794 |
| 0.09758 | 0.06167 | 0.4838 |
| 0.008647 | 0.06339 | 0.9232 |
| 0.04987 | 0.06342 | 0.9292 |
| 0.07109 | 0.0644 | 0.3315 |
| 0.008645 | 0.06453 | 0.922 |
| 0.003608 | 0.06504 | 0.6006 |
| 0.008626 | 0.06549 | 0.9248 |
| 0.04285 | 0.06745 | 0.7336 |
| 0.008621 | 0.06848 | 0.9252 |
| 0.1013 | 0.06947 | 0.4029 |
| 0.04487 | 0.0705 | 0.7743 |
| 0.005077 | 0.0722 | 0.9201 |
| 0.1565 | 0.07449 | 0.4243 |
| 0.1751 | 0.07501 | 0.3271 |
| 0.03317 | 0.07652 | 0.6893 |
| 0.02474 | 0.07669 | 0.7119 |
| 0.006349 | 0.07861 | 0.8482 |
| 0.05698 | 0.0788 | 0.3058 |
| 0.06629 | 0.07995 | 0.5533 |
| 0.06625 | 0.08128 | 0.564 |
| 0.06625 | 0.08128 | 0.564 |
| 0.06539 | 0.08529 | 0.3391 |
| 0.00553 | 0.0877 | 0.7808 |
| 0.1053 | 0.08794 | 0.4428 |
| 0.05528 | 0.0916 | 0.5726 |
| 0.01142 | 0.09162 | 0.8567 |
| 0.0238 | 0.09227 | 0.5358 |
| 0.04708 | 0.09309 | 0.4812 |
| 0.01341 | 0.09724 | 0.8026 |
| 0.02347 | 0.09808 | 0.7712 |
| 0.1001 | 0.09817 | 0.6819 |
| 0.05482 | 0.09851 | 0.8549 |
| 0.008444 | 0.1002 | 0.9148 |
| 0.05703 | 0.1027 | 0.3035 |
| 0.009337 | 0.1028 | 0.9398 |
| 0.009337 | 0.1028 | 0.9398 |
| 0.04611 | 0.1028 | 0.5204 |
| 0.009337 | 0.1039 | 0.94 |
| 0.02937 | 0.1041 | 0.9153 |
| 0.01053 | 0.1041 | 0.8653 |
| 0.04713 | 0.1052 | 0.5065 |
| 0.01054 | 0.1066 | 0.8651 |
| 0.009258 | 0.107 | 0.948 |
| 0.03429 | 0.109 | 0.9899 |
| 0.06776 | 0.1104 | 0.3812 |
| 0.006811 | 0.1112 | 0.6826 |
| 0.02135 | 0.1115 | 0.7892 |
| 0.06788 | 0.1134 | 0.3796 |
| 0.09757 | 0.1138 | 0.3275 |
| 0.06731 | 0.117 | 0.8533 |
| 0.09453 | 0.1171 | 0.34 |
| 0.01474 | 0.1195 | 0.4946 |
| 0.06846 | 0.1225 | 0.8155 |
| 0.01829 | 0.1243 | 0.8468 |
| 0.01013 | 0.1252 | 0.7828 |
| 0.03923 | 0.1263 | 0.7824 |
| 0.01138 | 0.1296 | 0.8362 |
| 0.045 | 0.1297 | 0.4613 |
| 0.09479 | 0.1306 | 0.3417 |
| 0.01391 | 0.1306 | 0.3106 |
| 0.04987 | 0.1369 | 0.6218 |
| 0.05644 | 0.1373 | 0.3068 |
| 0.1602 | 0.1399 | 0.3512 |
| 0.007596 | 0.1428 | 0.8983 |
| 0.06295 | 0.1436 | 0.8702 |
| 0.01044 | 0.1438 | 0.3649 |
| 0.01557 | 0.1467 | 0.7415 |
| 0.02986 | 0.1473 | 0.6089 |
| 0.03099 | 0.1486 | 0.685 |
| 0.05042 | 0.1501 | 0.3905 |
| 0.1331 | 0.1546 | 0.7027 |
| 0.03009 | 0.1562 | 0.6873 |
| 0.06734 | 0.1572 | 0.6225 |
| 0.0633 | 0.1575 | 0.8508 |
| 0.118 | 0.1597 | 0.783 |
| 0.01897 | 0.1616 | 0.8996 |
| 0.009867 | 0.1644 | 0.7943 |
| 0.0854 | 0.168 | 0.8477 |
| 0.03002 | 0.1692 | 0.7086 |
| 0.03622 | 0.1746 | 0.5708 |
| 0.03376 | 0.1755 | 0.6704 |
| 0.01067 | 0.1765 | 0.639 |
| 0.0338 | 0.1787 | 0.6708 |
| 0.01305 | 0.1795 | 0.8589 |
| 0.0338 | 0.1795 | 0.6707 |
| 0.1615 | 0.1813 | 0.4304 |
| 0.105 | 0.1817 | 0.3884 |
| 0.04464 | 0.1864 | 0.436 |
| 0.005791 | 0.188 | 0.3164 |
| 0.009171 | 0.188 | 0.9377 |
| 0.03394 | 0.1891 | 0.6701 |
| 0.05186 | 0.1986 | 0.6176 |
| 0.03564 | 0.2028 | 0.9519 |
| 0.08443 | 0.2054 | 0.3098 |
| 0.05119 | 0.2097 | 0.5656 |
| 0.04496 | 0.2101 | 0.432 |
| 0.08301 | 0.2116 | 0.6353 |
| 0.03606 | 0.2162 | 0.9497 |
| 0.03608 | 0.2217 | 0.9415 |
| 0.1282 | 0.222 | 0.7175 |
| 0.03698 | 0.2234 | 0.9386 |
| 0.03431 | 0.2249 | 0.6262 |
| 0.03979 | 0.226 | 0.6119 |
| 0.03981 | 0.2345 | 0.5965 |
| 0.006469 | 0.2364 | 0.9447 |
| 0.06351 | 0.2367 | 0.3052 |
| 0.02534 | 0.238 | 0.4648 |
| 0.04036 | 0.2394 | 0.9988 |
| 0.07307 | 0.2408 | 0.5029 |
| 0.03625 | 0.2416 | 0.9525 |
| 0.09692 | 0.243 | 0.4242 |
| 0.06338 | 0.2494 | 0.5139 |
| 0.06293 | 0.2499 | 0.6251 |
| 0.1212 | 0.2519 | 0.3532 |
| 0.09678 | 0.2562 | 0.4136 |
| 0.08525 | 0.2582 | 0.3608 |
| 0.0396 | 0.2609 | 0.6032 |
| 0.009865 | 0.2622 | 0.9337 |
| 0.01377 | 0.2674 | 0.9235 |
| 0.04113 | 0.2678 | 0.6785 |
| 0.02252 | 0.2685 | 0.698 |
| 0.04082 | 0.269 | 0.5679 |
| 0.1191 | 0.2702 | 0.4708 |
| 0.02553 | 0.2781 | 0.7185 |
| 0.08537 | 0.2784 | 0.3604 |
| 0.01591 | 0.288 | 0.9725 |
| 0.04036 | 0.2899 | 0.6361 |
| 0.04035 | 0.2906 | 0.6339 |
| 0.07834 | 0.2911 | 0.8883 |
| 0.07834 | 0.2929 | 0.8563 |
| 0.009768 | 0.2939 | 0.6703 |
| 0.02906 | 0.2945 | 0.9987 |
| 0.06261 | 0.2986 | 0.5923 |
| 0.0404 | 0.2987 | 0.6541 |
| 0.09679 | 0.3001 | 0.3588 |
| 0.1084 | 0.3028 | 0.7561 |
| 0.04374 | 0.3044 | 0.8744 |
| 0.01596 | 0.3064 | 0.9987 |
| 0.07848 | 0.3108 | 0.4223 |
| 0.0849 | 0.3131 | 0.3305 |
| 0.1194 | 0.3137 | 0.6441 |
| 0.05651 | 0.3137 | 0.8968 |
| 0.05651 | 0.3137 | 0.8968 |
| 0.05637 | 0.3167 | 0.7945 |
| 0.04016 | 0.3181 | 0.6343 |
| 0.05654 | 0.3198 | 0.8958 |
| 0.05654 | 0.3235 | 0.8951 |
| 0.07512 | 0.3262 | 0.9768 |
| 0.07512 | 0.3262 | 0.9768 |
| 0.007358 | 0.327 | 0.4785 |
| 0.05653 | 0.3274 | 0.8944 |
| 0.04505 | 0.3274 | 0.7382 |
| 0.09694 | 0.3278 | 0.3134 |
| 0.01541 | 0.3293 | 0.8732 |
| 0.01526 | 0.3324 | 0.8734 |
| 0.01524 | 0.3396 | 0.8405 |
| 0.008198 | 0.3439 | 0.4511 |
| 0.0868 | 0.3493 | 0.3474 |
| 0.03724 | 0.3503 | 0.7323 |
| 0.03218 | 0.3528 | 0.6107 |
| 0.08523 | 0.3601 | 0.33 |
| 0.09396 | 0.3661 | 0.3939 |
| 0.0399 | 0.3661 | 0.3066 |
| 0.0111 | 0.3733 | 0.884 |
| 0.09531 | 0.3744 | 0.4196 |
| 0.09531 | 0.3744 | 0.4196 |
| 0.09526 | 0.3815 | 0.4165 |
| 0.03176 | 0.3827 | 0.6879 |
| 0.009845 | 0.3828 | 0.8361 |
| 0.01912 | 0.3911 | 0.8091 |
| 0.09583 | 0.4029 | 0.4197 |
| 0.01973 | 0.4051 | 0.9578 |
| 0.03325 | 0.4069 | 0.6341 |
| 0.02156 | 0.4078 | 0.8923 |
| 0.01494 | 0.412 | 0.9049 |
| 0.07421 | 0.4155 | 0.8327 |
| 0.03937 | 0.4167 | 0.4594 |
| 0.1064 | 0.4178 | 0.399 |
| 0.04768 | 0.4198 | 0.5053 |
| 0.09479 | 0.4221 | 0.3996 |
| 0.09479 | 0.4221 | 0.3996 |
| 0.04038 | 0.4302 | 0.6435 |
| 0.02083 | 0.4304 | 0.6797 |
| 0.02928 | 0.436 | 0.5254 |
| 0.1556 | 0.4401 | 0.3232 |
| 0.1195 | 0.4426 | 0.4848 |
| 0.06529 | 0.4428 | 0.8718 |
| 0.1241 | 0.4434 | 0.4465 |
| 0.02221 | 0.4436 | 0.8969 |
| 0.03466 | 0.4514 | 0.5338 |
| 0.0577 | 0.4564 | 0.3799 |
| 0.01049 | 0.4578 | 0.8089 |
| 0.03932 | 0.4648 | 0.9824 |
| 0.008293 | 0.4658 | 0.7015 |
| 0.1218 | 0.4684 | 0.7128 |
| 0.01518 | 0.4729 | 0.8425 |
| 0.06344 | 0.4749 | 0.8459 |
| 0.02419 | 0.4764 | 0.9086 |
| 0.0578 | 0.4804 | 0.5336 |
| 0.06379 | 0.4813 | 0.4765 |
| 0.09268 | 0.4826 | 0.5504 |
| 0.02561 | 0.4829 | 0.4417 |
| 0.04616 | 0.4833 | 0.475 |
| 0.05854 | 0.4851 | 0.8933 |
| 0.066 | 0.4919 | 0.6653 |
| 0.02781 | 0.4926 | 0.7443 |
| 0.1665 | 0.4933 | 0.4139 |
| 0.04482 | 0.4944 | 0.5585 |
| 0.03726 | 0.5088 | 0.6541 |
| 0.1478 | 0.5104 | 0.4765 |
| 0.03341 | 0.5186 | 0.3668 |
| 0.05628 | 0.5196 | 0.9332 |
| 0.05628 | 0.5196 | 0.9332 |
| 0.0572 | 0.5268 | 0.376 |
| 0.09698 | 0.527 | 0.5015 |
| 0.01045 | 0.5272 | 0.3009 |
| 0.04488 | 0.5313 | 0.4726 |
| 0.1423 | 0.5373 | 0.6311 |
| 0.08246 | 0.5479 | 0.3046 |
| 0.004896 | 0.5504 | 0.9541 |
| 0.01243 | 0.5547 | 0.6971 |
| 0.01055 | 0.556 | 0.8727 |
| 0.1407 | 0.5686 | 0.5007 |
| 0.1628 | 0.5716 | 0.328 |
| 0.1628 | 0.5716 | 0.328 |
| 0.003701 | 0.5749 | 0.8392 |
| 0.04605 | 0.5909 | 0.348 |
| 0.09172 | 0.5937 | 0.4619 |
| 0.03059 | 0.6133 | 0.6893 |
| 0.1634 | 0.6135 | 0.367 |
| 0.03815 | 0.6164 | 0.4624 |
| 0.08004 | 0.6175 | 0.5589 |
| 0.1633 | 0.618 | 0.4119 |
| 0.06791 | 0.6189 | 0.3267 |
| 0.1276 | 0.6196 | 0.5963 |
| 0.007952 | 0.6196 | 0.9665 |
| 0.02788 | 0.6245 | 0.6112 |
| 0.05052 | 0.6289 | 0.6897 |
| 0.04049 | 0.6371 | 0.5952 |
| 0.009025 | 0.6377 | 0.8592 |
| 0.03947 | 0.6475 | 0.4908 |
| 0.05114 | 0.6506 | 0.3882 |
| 0.02375 | 0.6535 | 0.8184 |
| 0.1288 | 0.6706 | 0.6068 |
| 0.1288 | 0.6706 | 0.6068 |
| 0.05379 | 0.6886 | 0.4875 |
| 0.03517 | 0.6914 | 0.5505 |
| 0.05373 | 0.6921 | 0.6284 |
| 0.04772 | 0.7009 | 0.5867 |
| 0.06862 | 0.7011 | 0.6181 |
| 0.0828 | 0.702 | 0.6025 |
| 0.006047 | 0.705 | 0.9676 |
| 0.07176 | 0.7063 | 0.6162 |
| 0.1062 | 0.7089 | 0.9998 |
| 0.05192 | 0.7137 | 0.7036 |
| 0.1292 | 0.7148 | 0.619 |
| 0.06774 | 0.7235 | 0.5499 |
| 0.0867 | 0.7337 | 0.3972 |
| 0.1304 | 0.7355 | 0.6141 |
| 0.03459 | 0.7414 | 0.8171 |
| 0.008421 | 0.7477 | 0.7961 |
| 0.03186 | 0.7479 | 0.6317 |
| 0.02605 | 0.7491 | 0.5504 |
| 0.0635 | 0.7525 | 0.3552 |
| 0.1016 | 0.755 | 0.5788 |
| 0.06282 | 0.7563 | 0.345 |
| 0.1108 | 0.7581 | 0.5126 |
| 0.009119 | 0.7605 | 0.8664 |
| 0.04673 | 0.7735 | 0.9922 |
| 0.02378 | 0.7906 | 0.7192 |
| 0.1411 | 0.7925 | 0.3414 |
| 0.06341 | 0.7937 | 0.6609 |
| 0.008188 | 0.8011 | 0.9491 |
| 0.03307 | 0.8071 | 0.6369 |
| 0.03113 | 0.8089 | 0.5635 |
| 0.02385 | 0.8092 | 0.807 |
| 0.01345 | 0.8236 | 0.8073 |
| 0.01859 | 0.8271 | 0.7045 |
| 0.01034 | 0.8289 | 0.6899 |
| 0.04987 | 0.8325 | 0.3478 |
| 0.05323 | 0.8334 | 0.6251 |
| 0.0152 | 0.834 | 0.5798 |
| 0.0152 | 0.834 | 0.5798 |
| 0.08818 | 0.8368 | 0.3911 |
| 0.03073 | 0.8431 | 0.8067 |
| 0.03073 | 0.8431 | 0.8067 |
| 0.02204 | 0.8437 | 0.459 |
| 0.1251 | 0.844 | 0.3237 |
| 0.07067 | 0.8522 | 0.4483 |
| 0.03464 | 0.8552 | 0.7765 |
| 0.02005 | 0.8626 | 0.8369 |
| 0.01863 | 0.8661 | 0.8363 |
| 0.05588 | 0.8698 | 0.475 |
| 0.0345 | 0.8705 | 0.7814 |
| 0.08017 | 0.8724 | 0.3262 |
| 0.03091 | 0.8804 | 0.7844 |
| 0.03715 | 0.8878 | 0.7461 |
| 0.05195 | 0.8899 | 0.3823 |
| 0.0372 | 0.8919 | 0.747 |
| 0.0101 | 0.895 | 0.8585 |
| 0.0104 | 0.8966 | 0.6901 |
| 0.02383 | 0.8972 | 0.7933 |
| 0.1329 | 0.9023 | 0.699 |
| 0.0312 | 0.9117 | 0.837 |
| 0.09058 | 0.9122 | 0.8168 |
| 0.09058 | 0.9129 | 0.8174 |
| 0.01323 | 0.9145 | 0.9063 |
| 0.01812 | 0.9176 | 0.8525 |
| 0.105 | 0.9215 | 0.3821 |
| 0.01915 | 0.9262 | 0.6652 |
| 0.02388 | 0.928 | 0.7657 |
| 0.0739 | 0.9354 | 0.3719 |
| 0.006348 | 0.9374 | 0.985 |
| 0.09006 | 0.9376 | 0.8397 |
| 0.09006 | 0.9376 | 0.8397 |
| 0.03963 | 0.9394 | 0.6416 |
| 0.03963 | 0.9394 | 0.6416 |
| 0.07181 | 0.9435 | 0.5192 |
| 0.01569 | 0.9478 | 0.82 |
| 0.02962 | 0.9515 | 0.8227 |
| 0.06343 | 0.9518 | 0.3547 |
| 0.01993 | 0.9549 | 0.7278 |
| 0.1516 | 0.9576 | 0.5345 |
| 0.04832 | 0.9595 | 0.4281 |
| 0.009299 | 0.9611 | 0.9371 |
| 0.01302 | 0.9641 | 0.6742 |
| 0.03691 | 0.9743 | 0.4188 |
| 0.1057 | 0.9748 | 0.4745 |
| 0.02186 | 0.9795 | 0.8522 |
| 0.006305 | 0.9823 | 0.9979 |
| 0.0286 | 0.985 | 0.4338 |
| 0.02059 | 0.9903 | 0.7111 |
| 0.01072 | 0.9925 | 0.6817 |
| 0.03942 | 0.9946 | 0.7405 |
| 0.1342 | 0.995 | 0.6908 |
| 0.03956 | 0.9979 | 0.6173 |

| rsID | chr | pos (hg38) | Effect allele | non-effect allele | effect allele frequency | beta |
| --- | --- | --- | --- | --- | --- | --- |
| rs76232023 | 15 | 42893541 | C | A | 0.9968 | 0.06999 |
| rs147153494 | 15 | 42956820 | C | T | 0.9991 | 0.2172 |
| rs150316748 | 15 | 42901223 | G | A | 0.9963 | 0.07353 |
| rs149889485 | 15 | 42913969 | T | A | 0.9914 | -0.04362 |
| rs369926923 | 15 | 42979041 | C | T | 0.9914 | -0.04346 |
| rs375380714 | 15 | 42998215 | T | G | 0.9998 | -0.2663 |
| rs140878173 | 15 | 42911799 | G | C | 0.9976 | -0.07255 |
| rs184036150 | 15 | 42934772 | A | C | 0.9972 | 0.07315 |
| rs117347505 | 15 | 42993718 | C | T | 0.9972 | 0.07384 |
| rs78563306 | 15 | 42967694 | C | T | 0.9972 | 0.07321 |
| rs141111379 | 15 | 42935162 | C | T | 0.992 | -0.04201 |
| rs562869855 | 15 | 42948721 | A | G | 0.9974 | -0.07879 |
| rs148028047 | 15 | 42903243 | T | C | 0.9989 | -0.1365 |
| rs140433299 | 15 | 42910322 | T | C | 0.9996 | 0.2296 |
| rs529330569 | 15 | 42980693 | C | T | 0.9938 | 0.05515 |
| rs28535059 | 15 | 42933903 | T | A | 0.998 | 0.1021 |
| rs185098077 | 15 | 42912675 | C | T | 0.9998 | -0.3706 |
| rs567155071 | 15 | 42903081 | G | A | 0.9964 | 0.05012 |
| rs73413058 | 15 | 42988269 | A | G | 0.9369 | -0.02121 |
| rs572583293 | 15 | 42993853 | C | G | 0.9957 | 0.05872 |
| rs143766227 | 15 | 42883585 | CAA | C | 0.9951 | 0.06653 |
| rs117422009 | 15 | 43011777 | C | G | 0.9903 | 0.03131 |
| rs191375171 | 15 | 42888109 | C | T | 0.9972 | -0.06596 |
| rs143378302 | 15 | 42886009 | T | A | 0.9996 | 0.2385 |
| rs138556557 | 15 | 42895364 | G | A | 0.994 | -0.03676 |
| rs189542431 | 15 | 42974675 | C | G | 0.9966 | 0.0533 |
| rs561774462 | 15 | 42931411 | C | T | 0.9987 | 0.08449 |
| rs558339555 | 15 | 42913637 | CA | C | 0.9815 | 0.03114 |
| rs141583312 | 15 | 42975909 | G | A | 0.9984 | 0.07608 |
| rs557783986 | 15 | 42951686 | T | C | 0.9983 | -0.07076 |
| rs138976273 | 15 | 42902170 | C | T | 0.9864 | 0.02356 |
| rs182320019 | 15 | 42908120 | A | G | 0.9992 | 0.0973 |
| rs80305987 | 15 | 42877994 | A | G | 0.9899 | -0.02664 |
| rs146488861 | 15 | 42957808 | A | G | 0.9958 | 0.04463 |
| rs192977893 | 15 | 42887615 | C | T | 0.9986 | -0.08088 |
| rs143356796 | 15 | 43010938 | C | T | 0.9686 | 0.01622 |
| rs571167694 | 15 | 43011992 | C | T | 0.999 | -0.08609 |
| rs143875230 | 15 | 42986528 | G | A | 0.9729 | 0.01485 |
| rs563964635 | 15 | 42893798 | A | G | 0.9969 | -0.05274 |
| rs2054388 | 15 | 42932260 | C | T | 0.998 | 0.06698 |
| rs74533898 | 15 | 42909616 | G | A | 0.9971 | -0.06232 |
| rs190806658 | 15 | 42940285 | A | G | 0.9974 | 0.07515 |
| rs371849516 | 15 | 42947665 | GA | G | 0.9863 | -0.02178 |
| rs76841629 | 15 | 43005378 | A | G | 0.5371 | -0.005766 |
| rs146248884 | 15 | 42965927 | A | G | 0.9955 | 0.03804 |
| rs140735548 | 15 | 42893937 | C | G | 0.9935 | -0.03733 |
| rs16957288 | 15 | 42958590 | T | C | 0.9745 | -0.0168 |
| rs2016538 | 15 | 42933901 | A | T | 0.8499 | -0.006213 |
| rs148534148 | 15 | 42934574 | G | T | 0.9972 | 0.05085 |
| rs143455945 | 15 | 42965670 | T | C | 0.9992 | 0.1307 |
| rs534379603 | 15 | 42894853 | G | A | 0.9997 | 0.2148 |
| rs149678972 | 15 | 42873237 | G | A | 0.9732 | -0.01587 |
| rs569388456 | 15 | 42980580 | C | T | 0.9998 | 0.2379 |
| rs117017655 | 15 | 42872405 | G | C | 0.9729 | -0.01571 |
| rs11070384 | 15 | 42983068 | A | G | 0.9991 | 0.1275 |
| rs1115752 | 15 | 43008863 | C | T | 0.9061 | 0.007288 |
| rs530713585 | 15 | 42912062 | G | A | 0.9998 | 0.1983 |
| rs543192039 | 15 | 42914069 | C | T | 0.9989 | 0.08018 |
| rs145784520 | 15 | 42966451 | C | T | 0.9998 | 0.1915 |
| rs541817722 | 15 | 42940819 | C | T | 0.9984 | 0.05216 |
| rs185103862 | 15 | 43003994 | C | T | 0.9985 | 0.05239 |
| rs552175940 | 15 | 42921383 | C | G | 0.9986 | 0.09064 |
| rs186771984 | 15 | 42950609 | C | T | 0.9984 | 0.05185 |
| rs13380212 | 15 | 43006842 | T | C | 0.9992 | 0.1215 |
| rs184008832 | 15 | 42890930 | G | A | 0.9984 | 0.05078 |
| rs116738094 | 15 | 42868777 | G | A | 0.998 | 0.0548 |
| rs184616495 | 15 | 43004660 | C | T | 0.9919 | -0.02597 |
| rs537461773 | 15 | 42923097 | G | C | 0.9993 | -0.089 |
| rs16957292 | 15 | 42982370 | C | T | 0.9992 | 0.1225 |
| rs77547387 | 15 | 42997595 | G | A | 0.9992 | 0.1203 |
| rs377202784 | 15 | 42920633 | TGGC | T | 0.9971 | 0.05486 |
| rs572970295 | 15 | 42930764 | C | A | 0.9931 | -0.02551 |
| rs569234006 | 15 | 42940808 | G | GT | 0.9985 | 0.05134 |
| rs562676922 | 15 | 42876073 | C | T | 0.9996 | 0.1717 |
| rs114245127 | 15 | 42972868 | C | A | 0.9993 | 0.1295 |
| rs116154059 | 15 | 42962111 | C | A | 0.9993 | 0.1294 |
| rs559045802 | 15 | 42906458 | A | G | 0.9994 | 0.1114 |
| rs6493070 | 15 | 42900946 | A | G | 0.08955 | -0.007452 |
| rs373428985 | 15 | 42893814 | T | C | 0.9987 | 0.06063 |
| rs561617085 | 15 | 42986011 | A | G | 0.9823 | -0.01513 |
| rs142005131 | 15 | 42882373 | C | A | 0.9713 | -0.01192 |
| rs117095901 | 15 | 42912799 | C | T | 0.9888 | -0.01807 |
| rs560519383 | 15 | 42994134 | T | C | 0.998 | -0.06039 |
| rs184351682 | 15 | 42944285 | T | C | 0.9988 | 0.09261 |
| rs561908498 | 15 | 42925887 | C | G | 0.9998 | -0.2153 |
| rs192668256 | 15 | 42999153 | C | T | 0.9994 | -0.1256 |
| rs4923954 | 15 | 42969910 | C | G | 0.8577 | -0.005181 |
| rs11070383 | 15 | 42887165 | G | T | 0.06584 | -0.008739 |
| rs59276064 | 15 | 42919636 | G | A | 0.9979 | 0.04982 |
| rs12914380 | 15 | 42950070 | C | T | 0.8587 | -0.005104 |
| rs72721511 | 15 | 42994161 | T | C | 0.8588 | -0.005108 |
| rs138000286 | 15 | 42952142 | C | T | 0.986 | -0.01627 |
| rs35326878 | 15 | 42983379 | C | G | 0.8579 | -0.005084 |
| rs555343183 | 15 | 42900109 | C | T | 0.9987 | 0.08073 |
| rs12910799 | 15 | 42982666 | G | T | 0.8588 | -0.00505 |
| rs12911334 | 15 | 43013055 | G | A | 0.8592 | -0.005033 |
| rs4924699 | 15 | 42908911 | G | A | 0.8568 | -0.004922 |
| rs34474101 | 15 | 42895262 | A | G | 0.858 | -0.004893 |
| rs776170188 | 15 | 42982418 | G | A | 0.9998 | -0.1808 |
| rs72719486 | 15 | 42897273 | C | T | 0.8579 | -0.004859 |
| rs370980402 | 15 | 42990977 | C | T | 0.9986 | -0.07574 |
| rs35735714 | 15 | 42962133 | T | TC | 0.1534 | -0.006022 |
| rs11630330 | 15 | 43005852 | G | T | 0.1568 | -0.006023 |
| rs62020703 | 15 | 42989120 | T | G | 0.9891 | -0.01855 |
| rs530032514 | 15 | 42907916 | A | T | 0.9996 | -0.126 |
| rs143699768 | 15 | 42996145 | C | CT | 0.9392 | 0.007483 |
| rs11352938 | 15 | 42945607 | AT | A | 0.1932 | -0.005566 |
| rs142696672 | 15 | 42910808 | G | T | 0.9988 | 0.05121 |
| rs142838405 | 15 | 42934675 | G | A | 0.997 | 0.04178 |
| rs183621980 | 15 | 42884021 | G | A | 0.9952 | 0.02762 |
| rs143568757 | 15 | 42902686 | G | A | 0.997 | 0.04008 |
| rs114954873 | 15 | 42920847 | C | A | 0.9993 | -0.1018 |
| rs115507310 | 15 | 42920848 | T | G | 0.9993 | -0.1018 |
| rs556210813 | 15 | 42997532 | T | A | 0.9998 | 0.1496 |
| rs1972447 | 15 | 42960907 | C | T | 0.1345 | -0.005202 |
| rs117725364 | 15 | 42998580 | A | C | 0.992 | -0.02117 |
| rs199781595 | 15 | 42941307 | A | G | 0.9997 | -0.1367 |
| rs558747146 | 15 | 42880208 | G | A | 0.9984 | -0.04914 |
| rs4338775 | 15 | 42980100 | T | C | 0.1354 | -0.005171 |
| rs1993813 | 15 | 42981053 | C | T | 0.1358 | -0.005159 |
| rs147060258 | 15 | 42890797 | GAA | G | 0.9991 | 0.0532 |
| rs4509988 | 15 | 42980258 | G | T | 0.1335 | -0.00516 |
| rs8037022 | 15 | 42965277 | C | T | 0.135 | -0.005095 |
| rs7162959 | 15 | 42938552 | A | G | 0.1343 | -0.005078 |
| rs113183563 | 15 | 42898438 | G | A | 0.9994 | 0.1089 |
| rs4923955 | 15 | 43002890 | G | A | 0.134 | -0.005148 |
| rs9796571 | 15 | 43005659 | T | C | 0.1363 | -0.005114 |
| rs1037990 | 15 | 43007489 | T | C | 0.1361 | -0.005116 |
| rs184380146 | 15 | 42995397 | C | T | 0.9987 | 0.0752 |
| rs75580066 | 15 | 42877838 | A | G | 0.9972 | 0.03823 |
| rs6416436 | 15 | 42996021 | T | C | 0.1361 | -0.005042 |
| rs2412746 | 15 | 42994548 | C | T | 0.1364 | -0.005024 |
| rs75773646 | 15 | 42921982 | A | G | 0.9973 | 0.039 |
| rs147891025 | 15 | 42885547 | G | A | 0.9974 | 0.0334 |
| rs150770038 | 15 | 42897487 | C | T | 0.9974 | 0.0334 |
| rs12910269 | 15 | 42974089 | C | T | 0.1367 | -0.004972 |
| rs75487201 | 15 | 42921911 | T | G | 0.9973 | 0.03875 |
| rs34312632 | 15 | 42976589 | G | GATAA | 0.1528 | -0.00546 |
| rs7162939 | 15 | 42938472 | C | T | 0.1345 | -0.004897 |
| rs6493071 | 15 | 42916608 | C | G | 0.1326 | -0.004923 |
| rs8042481 | 15 | 43002020 | G | T | 0.1367 | -0.004907 |
| rs148100429 | 15 | 42875632 | C | T | 0.9997 | 0.09503 |
| rs190779787 | 15 | 42885334 | C | A | 0.9997 | 0.09503 |
| rs2054389 | 15 | 42963923 | T | C | 0.1313 | -0.004896 |
| rs1381856 | 15 | 42972019 | C | T | 0.1313 | -0.004896 |
| rs183231149 | 15 | 42887050 | G | A | 0.9992 | 0.07191 |
| rs143837600 | 15 | 42981862 | G | T | 0.9997 | 0.09399 |
| rs187329646 | 15 | 42983026 | C | A | 0.9997 | 0.09373 |
| rs2412747 | 15 | 42994605 | T | C | 0.1369 | -0.004795 |
| rs147047460 | 15 | 42885872 | C | T | 0.999 | -0.07678 |
| rs7178567 | 15 | 42944414 | C | T | 0.1305 | -0.004786 |
| rs115342811 | 15 | 42881852 | C | T | 0.9993 | 0.1007 |
| rs78084245 | 15 | 42963434 | A | G | 0.9722 | -0.009455 |
| rs562776360 | 15 | 43002939 | T | C | 0.9996 | -0.08577 |
| rs3803341 | 15 | 42945005 | T | C | 0.1339 | -0.004705 |
| rs189693176 | 15 | 42950613 | A | G | 0.9998 | 0.1464 |
| rs574471910 | 15 | 42971058 | T | G | 0.9985 | 0.06437 |
| rs149843783 | 15 | 42935351 | C | T | 0.9998 | 0.159 |
| rs562725261 | 15 | 42973598 | C | T | 0.9997 | -0.103 |
| rs77337431 | 15 | 42909774 | G | A | 0.9991 | 0.04805 |
| rs28374998 | 15 | 42996979 | C | G | 0.8217 | -0.003772 |
| rs28494835 | 15 | 42989193 | C | T | 0.8213 | -0.003755 |
| rs181239306 | 15 | 42916138 | T | C | 0.9993 | 0.07411 |
| rs8031283 | 15 | 42903011 | G | A | 0.9483 | -0.01055 |
| rs7174028 | 15 | 42922561 | G | T | 0.1317 | -0.004625 |
| rs8036674 | 15 | 43000977 | G | C | 0.8214 | -0.003724 |
| rs28633493 | 15 | 42971109 | T | C | 0.8205 | -0.003704 |
| rs7166705 | 15 | 42972928 | C | A | 0.8216 | -0.003694 |
| rs3759792 | 15 | 42960566 | A | G | 0.8208 | -0.003688 |
| rs4924701 | 15 | 42973522 | C | T | 0.1311 | -0.00464 |
| rs117337065 | 15 | 42903762 | G | A | 0.9982 | -0.04197 |
| rs9920061 | 15 | 42941863 | T | C | 0.8203 | -0.003669 |
| rs4244590 | 15 | 42935141 | G | A | 0.1319 | -0.004545 |
| rs552139984 | 15 | 42981736 | G | A | 0.9978 | -0.04444 |
| rs189526440 | 15 | 42907138 | T | A | 0.9987 | 0.04262 |
| rs115922627 | 15 | 42907139 | C | A | 0.9987 | 0.04262 |
| rs59589598 | 15 | 42884810 | G | A | 0.9987 | 0.04253 |
| rs190822677 | 15 | 42989502 | G | A | 0.9995 | 0.06573 |
| rs186133585 | 15 | 42908179 | C | T | 0.9998 | 0.1399 |
| rs182906122 | 15 | 42953382 | C | T | 0.9942 | -0.0195 |
| rs2412748 | 15 | 42935317 | G | A | 0.8219 | -0.003604 |
| rs558292498 | 15 | 42911675 | C | T | 0.9994 | 0.06161 |
| rs192592711 | 15 | 42877415 | G | A | 0.9995 | 0.06606 |
| rs139393261 | 15 | 42922092 | C | T | 0.9991 | 0.04594 |
| rs139550877 | 15 | 42924878 | G | T | 0.9991 | 0.04594 |
| rs145449308 | 15 | 42997265 | C | G | 0.9997 | 0.09824 |
| rs2682073 | 15 | 42904360 | T | C | 0.8884 | 0.004353 |
| rs66530524 | 15 | 42880097 | T | C | 0.8887 | 0.004349 |
| rs79694045 | 15 | 42948422 | T | G | 0.8175 | -0.003594 |
| rs34312217 | 15 | 42902986 | CA | C | 0.931 | -0.007774 |
| rs570080968 | 15 | 43013157 | G | A | 0.9998 | 0.1582 |
| rs545291075 | 15 | 42871454 | T | A | 0.9997 | -0.08025 |
| rs144326298 | 15 | 42990770 | T | C | 0.999 | 0.07809 |
| rs117722860 | 15 | 43008329 | C | T | 0.9755 | -0.00864 |
| rs34436896 | 15 | 42973919 | C | T | 0.9974 | 0.03333 |
| rs60763334 | 15 | 42902516 | G | A | 0.8892 | 0.004199 |
| rs542549746 | 15 | 42910957 | T | C | 0.9997 | 0.09575 |
| rs7178248 | 15 | 42907321 | C | T | 0.8203 | -0.00336 |
| rs3917223 | 15 | 42963993 | T | C | 0.9244 | -0.00481 |
| rs28613047 | 15 | 42883967 | G | A | 0.8203 | -0.00335 |
| rs143043128 | 15 | 42936058 | C | T | 0.999 | 0.04224 |
| rs28796057 | 15 | 42896566 | C | T | 0.8203 | -0.003344 |
| rs9920182 | 15 | 42910352 | G | A | 0.8203 | -0.003343 |
| rs2733224 | 15 | 42913563 | G | A | 0.8903 | 0.004188 |
| rs16957244 | 15 | 42873831 | C | T | 0.8897 | 0.004134 |
| rs28609797 | 15 | 42922620 | T | C | 0.819 | -0.003425 |
| rs28871732 | 15 | 42896650 | C | T | 0.8203 | -0.003323 |
| rs577021890 | 15 | 42891221 | G | A | 0.999 | -0.05719 |
| rs535809709 | 15 | 42883375 | C | T | 0.9998 | -0.1409 |
| rs554508188 | 15 | 42931509 | C | T | 0.9993 | 0.06157 |
| rs145037353 | 15 | 42912432 | C | A | 0.999 | 0.04127 |
| rs116044972 | 15 | 42939606 | G | T | 0.999 | 0.04121 |
| rs143251532 | 15 | 42998512 | T | C | 0.9996 | 0.0922 |
| rs185185807 | 15 | 42986374 | C | T | 0.9973 | 0.02613 |
| rs57880396 | 15 | 42912858 | G | A | 0.9973 | 0.03091 |
| rs143800868 | 15 | 42925865 | G | A | 0.997 | 0.0304 |
| rs191544070 | 15 | 42974511 | A | C | 0.9958 | 0.02086 |
| rs150358878 | 15 | 42952069 | A | T | 0.9979 | 0.03574 |
| rs78769997 | 15 | 42922138 | C | G | 0.9972 | 0.0308 |
| rs72721503 | 15 | 42968409 | G | C | 0.9726 | 0.009313 |
| rs112929196 | 15 | 43008566 | G | A | 0.9609 | 0.006624 |
| rs548296727 | 15 | 42970157 | T | C | 0.996 | -0.03472 |
| rs201384130 | 15 | 42964023 | T | G | 0.9998 | -0.1292 |
| rs74570780 | 15 | 42908305 | C | T | 0.9969 | 0.02913 |
| rs9806161 | 15 | 42883890 | A | T | 0.9968 | 0.02901 |
| rs11332679 | 15 | 42877492 | GA | G | 0.8176 | -0.003214 |
| rs113476707 | 15 | 42878194 | A | G | 0.9968 | 0.02874 |
| rs561140781 | 15 | 42956307 | T | G | 0.9979 | -0.03366 |
| rs118104470 | 15 | 42913334 | C | T | 0.9904 | -0.01308 |
| rs1040748507 | 15 | 42889826 | G | A | 0.9974 | 0.03204 |
| rs76989793 | 15 | 42922584 | A | T | 0.9899 | -0.01285 |
| rs148386184 | 15 | 42954593 | A | T | 0.9906 | -0.01289 |
| rs573802972 | 15 | 42925909 | G | A | 0.9984 | -0.05237 |
| rs553452245 | 15 | 42908200 | G | T | 0.9992 | 0.04176 |
| rs8031111 | 15 | 42911544 | C | T | 0.9974 | 0.03007 |
| rs16957264 | 15 | 42906232 | C | T | 0.9097 | 0.004134 |
| rs138218565 | 15 | 42953167 | A | C | 0.9966 | -0.02113 |
| rs549942856 | 15 | 42877419 | C | T | 0.9998 | 0.1038 |
| rs141795624 | 15 | 42979926 | G | A | 0.9982 | 0.03297 |
| rs148848820 | 15 | 42895452 | T | C | 0.9816 | -0.01003 |
| rs148952135 | 15 | 42966774 | T | C | 0.9969 | 0.02414 |
| rs187287466 | 15 | 42887754 | C | A | 0.9992 | 0.03974 |
| rs142424434 | 15 | 42925466 | G | C | 0.9996 | -0.06304 |
| rs150630012 | 15 | 42942394 | C | T | 0.9986 | 0.03443 |
| rs185463000 | 15 | 42915407 | G | A | 0.9989 | -0.05422 |
| rs75776391 | 15 | 43011820 | G | A | 0.999 | 0.03643 |
| rs141919417 | 15 | 42901329 | T | A | 0.9946 | 0.01905 |
| rs200325235 | 15 | 43010236 | CA | C | 0.9642 | 0.00666 |
| rs568462372 | 15 | 42924192 | C | G | 0.9998 | 0.1019 |
| rs551671670 | 15 | 42927127 | T | C | 0.9998 | 0.1019 |
| rs191535749 | 15 | 42985222 | C | G | 0.9998 | 0.102 |
| rs187320661 | 15 | 42934800 | T | C | 0.9996 | -0.06192 |
| rs567039670 | 15 | 42963819 | T | C | 0.9989 | -0.05369 |
| rs557339939 | 15 | 42929212 | G | A | 0.9983 | -0.03585 |
| rs566413696 | 15 | 42983841 | T | C | 0.9992 | 0.06733 |
| rs537550535 | 15 | 42971805 | A | G | 0.9983 | -0.03576 |
| rs7176123 | 15 | 42947732 | G | A | 0.9987 | 0.03306 |
| rs139987841 | 15 | 42965625 | C | T | 0.9988 | 0.03504 |
| rs28392857 | 15 | 42883751 | C | T | 0.9709 | 0.006576 |
| rs9806313 | 15 | 42884268 | G | A | 0.9966 | 0.02497 |
| rs151022817 | 15 | 42910971 | G | A | 0.9988 | 0.03925 |
| rs558490453 | 15 | 42935340 | G | A | 0.9979 | -0.02838 |
| rs567241776 | 15 | 42927294 | C | T | 0.9994 | 0.06547 |
| rs116640993 | 15 | 42972421 | T | C | 0.999 | 0.03412 |
| rs146799639 | 15 | 42870779 | T | C | 0.9416 | -0.004447 |
| rs185505090 | 15 | 43006659 | G | T | 0.9987 | 0.04117 |
| rs546580872 | 15 | 42929355 | A | G | 0.9992 | 0.03687 |
| rs16957284 | 15 | 42955369 | T | C | 0.998 | -0.04112 |
| rs544857253 | 15 | 42904460 | G | C | 0.9998 | 0.07749 |
| rs547972042 | 15 | 43005198 | G | A | 0.9944 | 0.01616 |
| rs2126602 | 15 | 43009278 | T | C | 0.1305 | -0.003365 |
| rs116103891 | 15 | 42955577 | A | T | 0.9998 | -0.1295 |
| rs58483557 | 15 | 42956876 | C | A | 0.998 | -0.04045 |
| rs7179367 | 15 | 43005074 | C | T | 0.9986 | 0.03051 |
| rs541558127 | 15 | 42939846 | T | C | 0.9964 | 0.02209 |
| rs182927920 | 15 | 42967984 | T | C | 0.9981 | -0.03325 |
| rs75658696 | 15 | 42978682 | T | C | 0.9938 | -0.0145 |
| rs191793117 | 15 | 42914340 | T | C | 0.998 | -0.03568 |
| rs115723709 | 15 | 42962045 | C | T | 0.9987 | 0.03132 |
| rs6493074 | 15 | 43004985 | A | C | 0.1302 | -0.003287 |
| rs142285781 | 15 | 43007204 | G | A | 0.9991 | 0.03255 |
| rs530091685 | 15 | 42978732 | CT | C | 0.9665 | -0.009642 |
| rs114454965 | 15 | 43009037 | C | T | 0.9987 | 0.02968 |
| rs139251799 | 15 | 42918317 | T | C | 0.9998 | -0.1247 |
| rs2899069 | 15 | 42994373 | C | T | 0.1302 | -0.003227 |
| rs150988986 | 15 | 42924675 | A | T | 0.9998 | -0.1242 |
| rs534274491 | 15 | 43006688 | C | T | 0.9992 | 0.03776 |
| rs28674880 | 15 | 42871114 | G | T | 0.9708 | 0.005916 |
| rs114659791 | 15 | 42885873 | G | A | 0.9998 | 0.1047 |
| rs28649962 | 15 | 42898433 | T | C | 0.9708 | 0.005858 |
| rs117193101 | 15 | 42875336 | G | T | 0.9953 | -0.01445 |
| rs187610922 | 15 | 43004989 | C | T | 0.9995 | 0.05355 |
| rs572827078 | 15 | 42913020 | G | T | 0.9994 | 0.03933 |
| rs114969207 | 15 | 42891216 | G | A | 0.9998 | -0.119 |
| rs141311377 | 15 | 42912418 | A | C | 0.9993 | 0.03732 |
| rs145466527 | 15 | 42940296 | G | A | 0.9993 | 0.03732 |
| rs186564556 | 15 | 42902293 | G | C | 0.9997 | -0.04975 |
| rs373905733 | 15 | 42906618 | G | C | 0.9997 | -0.04975 |
| rs12917056 | 15 | 42953491 | T | C | 0.9313 | -0.003685 |
| rs150557750 | 15 | 42972607 | G | A | 0.9008 | 0.00321 |
| rs9920498 | 15 | 42985753 | G | C | 0.1293 | -0.00305 |
| rs557473381 | 15 | 43000769 | G | A | 0.9995 | 0.04245 |
| rs28858165 | 15 | 42885094 | G | A | 0.9709 | 0.005607 |
| rs150495511 | 15 | 42899170 | A | AT | 0.9181 | -0.004747 |
| rs113913915 | 15 | 42970124 | T | C | 0.9995 | -0.06232 |
| rs33987108 | 15 | 42936429 | A | AT | 0.431 | -0.002523 |
| rs16957277 | 15 | 42945374 | T | C | 0.9692 | 0.005237 |
| rs144230475 | 15 | 42987511 | C | T | 0.9995 | -0.06014 |
| rs147870890 | 15 | 42925732 | A | G | 0.9969 | 0.0209 |
| rs188917136 | 15 | 42936275 | G | A | 0.9952 | -0.01351 |
| rs144595138 | 15 | 43011980 | G | A | 0.9986 | 0.02623 |
| rs559628380 | 15 | 42919443 | G | T | 0.9947 | -0.012 |
| rs568614887 | 15 | 42947941 | G | A | 0.997 | 0.01855 |
| rs147209414 | 15 | 42897063 | G | A | 0.9987 | 0.03001 |
| rs556862650 | 15 | 42989496 | A | C | 0.9996 | -0.06345 |
| rs192718382 | 15 | 42905736 | C | T | 0.998 | -0.02078 |
| rs151019978 | 15 | 42991740 | T | C | 0.9981 | 0.02779 |
| rs771755844 | 15 | 42933197 | C | A | 0.9998 | -0.07215 |
| rs547298098 | 15 | 42913562 | C | T | 0.9992 | 0.03055 |
| rs116491117 | 15 | 42921767 | G | A | 0.997 | 0.02056 |
| rs552525489 | 15 | 42956466 | C | T | 0.9989 | -0.02973 |
| rs148174299 | 15 | 42993741 | G | A | 0.999 | 0.02627 |
| rs193196275 | 15 | 42908780 | A | G | 0.9974 | 0.02007 |
| rs145034051 | 15 | 42992977 | T | C | 0.9974 | 0.0181 |
| rs550575683 | 15 | 42960132 | T | G | 0.9996 | 0.05631 |
| rs149328464 | 15 | 42887671 | T | C | 0.9983 | 0.02597 |
| rs572834165 | 15 | 42936208 | G | A | 0.998 | -0.02291 |
| rs180750404 | 15 | 42912673 | C | T | 0.998 | 0.02332 |
| rs141998787 | 15 | 42913448 | A | G | 0.9969 | 0.01945 |
| rs191443517 | 15 | 43005135 | C | T | 0.9972 | -0.01991 |
| rs144884699 | 15 | 42914174 | C | T | 0.972 | 0.004941 |
| rs8031767 | 15 | 42918707 | C | T | 0.129 | -0.002591 |
| rs114804224 | 15 | 42987662 | G | A | 0.9986 | 0.02331 |
| rs181630879 | 15 | 42889666 | G | C | 0.9979 | 0.02484 |
| rs77641540 | 15 | 42910125 | C | T | 0.9306 | -0.003111 |
| rs541359954 | 15 | 42880415 | C | G | 0.9977 | 0.02426 |
| rs554304132 | 15 | 42955636 | T | C | 0.9991 | -0.04477 |
| rs564224994 | 15 | 42966397 | T | A | 0.9981 | 0.0212 |
| rs537102106 | 15 | 42967458 | G | C | 0.9981 | 0.0212 |
| rs79984506 | 15 | 43001654 | T | C | 0.997 | 0.01657 |
| rs149119473 | 15 | 42914260 | G | T | 0.9995 | -0.05109 |
| rs551010070 | 15 | 42929055 | A | T | 0.9993 | -0.03519 |
| rs188362035 | 15 | 42918484 | G | A | 0.9972 | -0.01691 |
| rs118074434 | 15 | 43003277 | C | T | 0.9992 | 0.03468 |
| rs148438272 | 15 | 42918775 | G | T | 0.9977 | 0.02291 |
| rs549786810 | 15 | 42931943 | G | A | 0.9984 | 0.02917 |
| rs1984501 | 15 | 42987512 | G | A | 0.9781 | 0.005045 |
| rs548547947 | 15 | 42947931 | C | T | 0.9934 | -0.01044 |
| rs192727326 | 15 | 42948390 | A | C | 0.9998 | -0.06562 |
| rs1049733221 | 15 | 42914070 | G | A | 0.9997 | -0.06115 |
| rs116269151 | 15 | 42919023 | C | T | 0.9995 | -0.04597 |
| rs189290697 | 15 | 42931689 | C | T | 0.9969 | 0.01441 |
| rs80258074 | 15 | 42923044 | T | A | 0.9995 | -0.04545 |
| rs76215594 | 15 | 42926099 | G | T | 0.9995 | -0.04545 |
| rs7164041 | 15 | 42917832 | A | G | 0.997 | 0.01633 |
| rs114936782 | 15 | 42908384 | T | C | 0.998 | -0.02648 |
| rs77496195 | 15 | 42967615 | T | G | 0.9983 | 0.01892 |
| rs114066074 | 15 | 42927698 | A | G | 0.9993 | 0.02602 |
| rs74873423 | 15 | 42924611 | C | G | 0.9958 | -0.01043 |
| rs117710203 | 15 | 42953008 | C | T | 0.9956 | -0.01032 |
| rs543589659 | 15 | 42960960 | A | C | 0.9995 | 0.04231 |
| rs61069183 | 15 | 42890153 | T | C | 0.9956 | -0.009873 |
| rs544535807 | 15 | 42973310 | T | G | 0.9993 | 0.02941 |
| rs185738711 | 15 | 42981245 | A | T | 0.9993 | 0.02941 |
| rs12912026 | 15 | 42878529 | T | A | 0.7587 | -0.002549 |
| rs146708171 | 15 | 42979312 | T | C | 0.9958 | -0.009986 |
| rs145938839 | 15 | 42884633 | C | A | 0.9981 | 0.0223 |
| rs191933065 | 15 | 42946188 | G | C | 0.9994 | -0.03158 |
| rs146894776 | 15 | 42967945 | T | C | 0.9975 | -0.0147 |
| rs142290993 | 15 | 42978320 | T | C | 0.9993 | 0.02442 |
| rs1037982814 | 15 | 42900693 | C | T | 0.9998 | -0.06036 |
| rs117423500 | 15 | 42885260 | G | A | 0.9741 | 0.004218 |
| rs78948790 | 15 | 42958014 | T | C | 0.9923 | -0.006816 |
| rs150970024 | 15 | 42987339 | C | G | 0.9993 | 0.02408 |
| rs56071832 | 15 | 42988453 | C | G | 0.9993 | 0.04055 |
| rs139359398 | 15 | 42909794 | T | C | 0.9991 | 0.02625 |
| rs142202313 | 15 | 42996605 | A | C | 0.9975 | -0.01423 |
| rs139400588 | 15 | 43000387 | G | A | 0.9975 | -0.01402 |
| rs149265736 | 15 | 43008367 | C | T | 0.9993 | 0.02334 |
| rs181498789 | 15 | 43010216 | A | G | 0.9993 | 0.02334 |
| rs142183834 | 15 | 43010103 | A | T | 0.9975 | -0.0139 |
| rs75090487 | 15 | 42937481 | T | G | 0.9944 | -0.01162 |
| rs181075931 | 15 | 42884135 | G | A | 0.997 | -0.01295 |
| rs547814549 | 15 | 42964424 | G | A | 0.9996 | 0.03705 |
| rs146056091 | 15 | 42986205 | A | G | 0.9989 | -0.02735 |
| rs151031682 | 15 | 42934972 | G | A | 0.9902 | 0.005983 |
| rs117122292 | 15 | 42879243 | G | A | 0.9817 | -0.004441 |
| rs185850923 | 15 | 42993431 | T | C | 0.9947 | -0.01039 |
| rs144575157 | 15 | 42971564 | A | T | 0.9959 | -0.008838 |
| rs116042800 | 15 | 42887128 | C | G | 0.9995 | -0.03697 |
| rs138835702 | 15 | 42897067 | T | C | 0.9995 | -0.03697 |
| rs147284877 | 15 | 42977716 | T | C | 0.9989 | 0.02468 |
| rs55805543 | 15 | 42868205 | T | C | 0.8932 | 0.001805 |
| rs183189070 | 15 | 42979608 | T | C | 0.9992 | 0.02181 |
| rs139408969 | 15 | 42990087 | C | T | 0.999 | 0.01732 |
| rs187236177 | 15 | 42939431 | C | A | 0.9964 | -0.01144 |
| rs539214491 | 15 | 42937269 | G | A | 0.9998 | -0.04192 |
| rs540683576 | 15 | 42943926 | T | A | 0.9998 | -0.04192 |
| rs181004974 | 15 | 42874762 | C | T | 0.9933 | 0.007013 |
| rs552314563 | 15 | 43004920 | G | A | 0.9934 | -0.007188 |
| rs138592907 | 15 | 42946486 | C | T | 0.9956 | -0.008341 |
| rs140959617 | 15 | 42958045 | A | G | 0.9975 | -0.01052 |
| rs62020701 | 15 | 42966883 | G | C | 0.9034 | -0.001791 |
| rs138963231 | 15 | 42983997 | G | C | 0.9865 | -0.004812 |
| rs117347582 | 15 | 42998252 | C | A | 0.9977 | -0.0107 |
| rs144640419 | 15 | 42892393 | T | A | 0.9994 | 0.02303 |
| rs763442891 | 15 | 42878527 | A | ACT | 0.9988 | 0.01477 |
| rs140450055 | 15 | 42961579 | C | T | 0.9876 | -0.004767 |
| rs376331294 | 15 | 42960653 | G | A | 0.9998 | 0.04197 |
| rs189618809 | 15 | 42907976 | C | A | 0.9957 | -0.008505 |
| rs543516155 | 15 | 42974900 | T | C | 0.9998 | -0.03948 |
| rs552385948 | 15 | 43007566 | T | A | 0.9993 | -0.02962 |
| rs62020698 | 15 | 42945216 | C | T | 0.9063 | -0.001611 |
| rs73413030 | 15 | 42897626 | A | G | 0.998 | -0.01714 |
| rs556243779 | 15 | 42902209 | C | A | 0.9986 | 0.01757 |
| rs141828250 | 15 | 42945444 | C | T | 0.9987 | 0.01277 |
| rs141585079 | 15 | 42909609 | A | G | 0.967 | -0.00253 |
| rs146503263 | 15 | 42878832 | T | G | 0.9962 | -0.009197 |
| rs200262486 | 15 | 42978421 | GT | G | 0.9921 | 0.005885 |
| rs144693269 | 15 | 42898455 | G | T | 0.9754 | -0.002929 |
| rs79495360 | 15 | 42918441 | T | C | 0.9932 | 0.007536 |
| rs138942621 | 15 | 42930975 | T | C | 0.9995 | 0.02713 |
| rs62020680 | 15 | 42878426 | G | A | 0.8584 | -0.001204 |
| rs140207100 | 15 | 43011559 | T | A | 0.9994 | 0.01833 |
| rs567161987 | 15 | 42914001 | G | A | 0.9962 | -0.006812 |
| rs576834861 | 15 | 42877871 | G | A | 0.9996 | -0.02729 |
| rs571185339 | 15 | 42879699 | A | G | 0.9995 | 0.02304 |
| rs546077226 | 15 | 42903738 | G | GT | 0.9981 | -0.01031 |
| rs186041641 | 15 | 42871238 | A | G | 0.9959 | -0.006214 |
| rs138619967 | 15 | 42926998 | C | T | 0.9817 | 0.003133 |
| rs572544151 | 15 | 42902719 | G | A | 0.9998 | 0.04082 |
| rs140916237 | 15 | 42889531 | T | C | 0.9994 | -0.02542 |
| rs116238461 | 15 | 43001328 | T | C | 0.9996 | -0.03203 |
| rs150203226 | 15 | 42875831 | G | A | 0.9683 | -0.002177 |
| rs114026095 | 15 | 42973492 | C | A | 0.9982 | 0.01072 |
| rs191518723 | 15 | 42922665 | G | T | 0.9934 | -0.006689 |
| rs116640263 | 15 | 42981739 | C | T | 0.9981 | 0.01058 |
| rs185596875 | 15 | 42894226 | T | C | 0.999 | 0.02088 |
| rs77541575 | 15 | 42975754 | C | T | 0.9981 | 0.0105 |
| rs563467704 | 15 | 42982383 | T | C | 0.9995 | 0.02709 |
| rs535405759 | 15 | 43005328 | G | A | 0.9987 | 0.01673 |
| rs112362877 | 15 | 42938705 | G | A | 0.9481 | -0.001636 |
| rs2126603 | 15 | 42928587 | T | G | 0.312 | -0.0007474 |
| rs573769722 | 15 | 42901255 | G | A | 0.9966 | 0.008628 |
| rs34714662 | 15 | 43001896 | C | T | 0.7847 | 0.0008602 |
| rs138077192 | 15 | 42985240 | G | C | 0.9961 | -0.005381 |
| rs537741020 | 15 | 42936512 | A | C | 0.9997 | -0.02556 |
| rs141227308 | 15 | 42952389 | G | A | 0.9984 | 0.007892 |
| rs180684653 | 15 | 42982257 | A | G | 0.999 | -0.01189 |
| rs540626947 | 15 | 42917314 | A | T | 0.9986 | 0.008999 |
| rs191526970 | 15 | 42957820 | T | A | 0.9998 | 0.02695 |
| rs574816840 | 15 | 42987187 | C | T | 0.9979 | -0.007277 |
| rs76206222 | 15 | 42907919 | A | T | 0.8762 | -0.0009453 |
| rs190984235 | 15 | 42929593 | A | C | 0.9957 | -0.004951 |
| rs76783611 | 15 | 42988659 | C | T | 0.9981 | 0.008199 |
| rs181258192 | 15 | 42950910 | G | A | 0.9979 | -0.006814 |
| rs562977445 | 15 | 42947538 | T | C | 0.9816 | 0.002902 |
| rs142211421 | 15 | 42884940 | G | A | 0.9998 | 0.02242 |
| rs575373910 | 15 | 43005726 | C | T | 0.9825 | -0.00251 |
| rs10712537 | 15 | 42919283 | TA | T | 0.306 | -0.0008663 |
| rs776118491 | 15 | 42993735 | G | A | 0.9794 | -0.002002 |
| rs933480389 | 15 | 42942363 | C | G | 0.9989 | -0.01162 |
| rs28627318 | 15 | 42893859 | A | G | 0.9755 | 0.001685 |
| rs139693200 | 15 | 42942444 | C | G | 0.9979 | 0.007178 |
| rs116967439 | 15 | 42956352 | C | T | 0.9483 | -0.001122 |
| rs180994451 | 15 | 42939263 | A | G | 0.9974 | -0.006914 |
| rs16957250 | 15 | 42878625 | C | T | 0.8962 | 0.0008248 |
| rs570044324 | 15 | 42983733 | T | C | 0.9991 | -0.01227 |
| rs529234877 | 15 | 42924755 | A | G | 0.9998 | 0.02047 |
| rs148549752 | 15 | 42987049 | C | T | 0.9993 | -0.01132 |
| rs9920231 | 15 | 42917576 | G | A | 0.9757 | 0.001573 |
| rs28575571 | 15 | 42924988 | G | A | 0.9757 | 0.001573 |
| rs570963333 | 15 | 42925646 | G | A | 0.9985 | 0.009824 |
| rs536514977 | 15 | 42875884 | C | T | 0.9987 | -0.008935 |
| rs28664053 | 15 | 42922689 | G | A | 0.9757 | 0.001476 |
| rs546446869 | 15 | 42932919 | C | T | 0.9997 | -0.01866 |
| rs139382852 | 15 | 43003552 | G | A | 0.9993 | -0.01101 |
| rs17776339 | 15 | 42968180 | T | A | 0.9793 | 0.001605 |
| rs186206742 | 15 | 42922541 | C | A | 0.992 | -0.002411 |
| rs142975069 | 15 | 42981529 | G | A | 0.9983 | -0.007208 |
| rs141900598 | 15 | 42885895 | C | T | 0.9912 | -0.003326 |
| rs11632981 | 15 | 43000340 | A | C | 0.9776 | 0.001582 |
| rs182889305 | 15 | 42998734 | C | T | 0.9976 | -0.004602 |
| rs192937247 | 15 | 42973303 | C | T | 0.9968 | 0.005464 |
| rs56356401 | 15 | 42954663 | T | C | 0.9793 | 0.001534 |
| rs571019011 | 15 | 42870769 | GA | G | 0.9791 | 0.001867 |
| rs182366902 | 15 | 42922724 | A | C | 0.9946 | 0.003061 |
| rs142301748 | 15 | 42884365 | C | T | 0.9959 | 0.003978 |
| rs111339056 | 15 | 42872121 | C | T | 0.9998 | 0.02282 |
| rs111569959 | 15 | 42874184 | A | T | 0.9998 | 0.02282 |
| rs115309896 | 15 | 42929461 | T | C | 0.9995 | -0.01649 |
| rs28407836 | 15 | 42890341 | T | C | 0.7064 | -0.0004263 |
| rs1381855 | 15 | 42971807 | G | A | 0.3156 | 0.0004273 |
| rs142774139 | 15 | 42874805 | G | A | 0.993 | -0.00244 |
| rs557349180 | 15 | 42993733 | G | A | 0.9851 | -0.002018 |
| rs556811858 | 15 | 42993734 | AG | A | 0.9851 | -0.002018 |
| rs544054828 | 15 | 42977780 | CAA | C | 0.9993 | -0.01045 |
| rs4923952 | 15 | 42901233 | A | G | 0.707 | -0.0003831 |
| rs56114844 | 15 | 42887206 | C | T | 0.7065 | -0.00037 |
| rs78365925 | 15 | 42905493 | G | C | 0.9984 | 0.00665 |
| rs998954 | 15 | 42988571 | C | T | 0.9748 | 0.001053 |
| rs149836490 | 15 | 42950127 | C | T | 0.9998 | -0.01391 |
| rs28646262 | 15 | 42890237 | T | C | 0.7067 | -0.0003512 |
| rs1197534 | 15 | 42907922 | T | A | 0.9579 | 0.0009369 |
| rs187345619 | 15 | 42919369 | T | C | 0.998 | -0.005249 |
| rs7173097 | 15 | 42869601 | C | T | 0.7105 | -0.0003435 |
| rs191423170 | 15 | 42971334 | C | T | 0.9954 | -0.00249 |
| rs7165888 | 15 | 42898361 | A | G | 0.7063 | -0.0003248 |
| rs71108187 | 15 | 42887941 | C | CT | 0.7089 | -0.0003201 |
| rs8028731 | 15 | 42899126 | G | A | 0.7085 | -0.0003145 |
| rs181155374 | 15 | 42963621 | C | A | 0.9968 | 0.003585 |
| rs77231660 | 15 | 42981109 | C | T | 0.9483 | -0.0006027 |
| rs17776090 | 15 | 42943386 | A | G | 0.9751 | -0.0009593 |
| rs1197533 | 15 | 42906729 | G | A | 0.7075 | -0.0002782 |
| rs111850927 | 15 | 42921045 | T | A | 0.9998 | 0.01394 |
| rs201091488 | 15 | 42984985 | T | C | 0.9996 | 0.006125 |
| rs8028822 | 15 | 43012143 | C | T | 0.3161 | 0.0002509 |
| rs7169603 | 15 | 43008227 | C | T | 0.315 | 0.0002495 |
| rs62020705 | 15 | 43005140 | G | A | 0.315 | 0.0002483 |
| rs7176915 | 15 | 43009573 | G | T | 0.315 | 0.0002481 |
| rs11070385 | 15 | 43007774 | C | T | 0.3161 | 0.0002462 |
| rs145057608 | 15 | 42915246 | C | T | 0.9984 | -0.003024 |
| rs1206698 | 15 | 42911888 | C | T | 0.7071 | -0.0002382 |
| rs28890597 | 15 | 42887550 | T | C | 0.7091 | -0.0002302 |
| rs12913586 | 15 | 42887858 | T | C | 0.7091 | -0.0002279 |
| rs13329073 | 15 | 42897670 | A | T | 0.7075 | -0.0002111 |
| rs76864255 | 15 | 42971050 | T | A | 0.9979 | 0.002514 |
| rs562509996 | 15 | 43005091 | G | T | 0.9987 | 0.003105 |
| rs6493068 | 15 | 42878595 | A | G | 0.6992 | 0.0001848 |
| rs147200598 | 15 | 42892930 | G | C | 0.9932 | -0.001203 |
| rs2169581 | 15 | 42934568 | G | A | 0.3136 | 0.0001869 |
| rs115840904 | 15 | 42966913 | C | T | 0.9979 | 0.002341 |
| rs541474906 | 15 | 42968291 | A | G | 0.9981 | 0.002364 |
| rs150012114 | 15 | 42924974 | A | G | 0.9771 | 0.0005866 |
| rs7179275 | 15 | 43004823 | G | T | 0.9971 | -0.002199 |
| rs7178752 | 15 | 43004711 | C | T | 0.3157 | 0.0001726 |
| rs62020699 | 15 | 42947415 | T | C | 0.3133 | 0.0001678 |
| rs8025479 | 15 | 42968589 | T | C | 0.3154 | 0.000156 |
| rs2682075 | 15 | 42913095 | T | G | 0.7052 | -0.0001576 |
| rs62020697 | 15 | 42942050 | C | T | 0.3149 | 0.0001497 |
| rs552124488 | 15 | 42973990 | G | A | 0.9989 | 0.003102 |
| rs547468800 | 15 | 42991255 | G | A | 0.9996 | 0.005067 |
| rs12440652 | 15 | 42972201 | C | A | 0.3141 | 0.0001381 |
| rs2682074 | 15 | 42904892 | C | T | 0.7061 | -0.0001305 |
| rs74570754 | 15 | 42984738 | T | C | 0.9998 | -0.006674 |
| rs6493069 | 15 | 42894567 | C | T | 0.7065 | -0.0001268 |
| rs2277532 | 15 | 42952501 | A | G | 0.3136 | 0.0001138 |
| rs188034879 | 15 | 42979637 | G | A | 0.9959 | -0.0008315 |
| rs55683827 | 15 | 42905457 | A | G | 0.9956 | -0.000796 |
| rs12440651 | 15 | 42972200 | C | A | 0.3146 | 0.0001013 |
| rs9806175 | 15 | 42884122 | C | T | 0.7051 | -0.00009739 |
| rs12437637 | 15 | 42920620 | C | G | 0.3129 | 0.00009772 |
| rs543230721 | 15 | 42912342 | C | T | 0.9981 | 0.001068 |
| rs117350263 | 15 | 42951611 | A | G | 0.9935 | -0.0005552 |
| rs35209561 | 15 | 42949600 | A | AC | 0.3367 | 0.00008062 |
| rs12904179 | 15 | 42868307 | G | A | 0.7057 | -0.00006629 |
| rs151233914 | 15 | 42951943 | T | C | 0.9977 | 0.0008595 |
| rs28805017 | 15 | 42887650 | T | C | 0.704 | 0.00003922 |
| rs117252626 | 15 | 42953935 | T | C | 0.9991 | -0.0006193 |
| rs557668526 | 15 | 42931262 | T | C | 0.9989 | -0.0005149 |
| rs4924702 | 15 | 42975842 | T | C | 0.3025 | -0.00002873 |
| rs375843171 | 15 | 42983341 | T | C | 0.9981 | 0.0002205 |
| rs150514250 | 15 | 42999694 | G | A | 0.9992 | -0.0003346 |
| rs8036096 | 15 | 42878822 | T | C | 0.7043 | -0.00001203 |
| rs28715020 | 15 | 42880082 | G | A | 0.7057 | -0.000008386 |
| rs112780488 | 15 | 42975207 | T | C | 0.9998 | 0.0004115 |
| rs1548096 | 15 | 42961113 | T | C | 0.3516 | 0.000007538 |

| se | p-value | INFO |
| --- | --- | --- |
| 0.02349 | 0.002889 | 0.8664 |
| 0.07477 | 0.003671 | 0.3098 |
| 0.02536 | 0.003739 | 0.6469 |
| 0.01541 | 0.004637 | 0.7713 |
| 0.01542 | 0.004837 | 0.7703 |
| 0.09531 | 0.005213 | 0.9998 |
| 0.02635 | 0.005894 | 0.9153 |
| 0.02692 | 0.006587 | 0.7619 |
| 0.02732 | 0.006873 | 0.7466 |
| 0.02717 | 0.00706 | 0.7561 |
| 0.01576 | 0.007662 | 0.7937 |
| 0.02961 | 0.007797 | 0.6893 |
| 0.05137 | 0.007887 | 0.5336 |
| 0.09142 | 0.01203 | 0.4029 |
| 0.02279 | 0.01551 | 0.4878 |
| 0.04296 | 0.01749 | 0.4281 |
| 0.1567 | 0.018 | 0.3271 |
| 0.02167 | 0.02073 | 0.9086 |
| 0.009329 | 0.02299 | 0.3009 |
| 0.02603 | 0.02408 | 0.5254 |
| 0.02979 | 0.02553 | 0.3668 |
| 0.01403 | 0.02561 | 0.82 |
| 0.02979 | 0.02679 | 0.6341 |
| 0.1084 | 0.02773 | 0.3532 |
| 0.01694 | 0.03003 | 0.8996 |
| 0.0247 | 0.03096 | 0.7443 |
| 0.04004 | 0.03484 | 0.7382 |
| 0.01509 | 0.03903 | 0.3772 |
| 0.03746 | 0.04228 | 0.6748 |
| 0.03509 | 0.04373 | 0.7405 |
| 0.01197 | 0.04901 | 0.8073 |
| 0.04952 | 0.04941 | 0.8389 |
| 0.01356 | 0.04945 | 0.8425 |
| 0.02278 | 0.0501 | 0.7185 |
| 0.0427 | 0.05819 | 0.5867 |
| 0.008716 | 0.0627 | 0.6703 |
| 0.04651 | 0.06416 | 0.7036 |
| 0.008099 | 0.06667 | 0.8974 |
| 0.02883 | 0.06729 | 0.6107 |
| 0.03665 | 0.06765 | 0.58 |
| 0.03415 | 0.06805 | 0.4624 |
| 0.04122 | 0.06826 | 0.348 |
| 0.01196 | 0.06862 | 0.8026 |
| 0.003219 | 0.07328 | 0.6006 |
| 0.02124 | 0.07339 | 0.7657 |
| 0.02121 | 0.0784 | 0.5358 |
| 0.009571 | 0.07917 | 0.6817 |
| 0.003583 | 0.08293 | 0.9452 |
| 0.02951 | 0.08485 | 0.6369 |
| 0.07604 | 0.08561 | 0.33 |
| 0.1256 | 0.08729 | 0.3414 |
| 0.009286 | 0.08744 | 0.6901 |
| 0.1393 | 0.08753 | 0.4243 |
| 0.009228 | 0.08868 | 0.6899 |
| 0.07574 | 0.0924 | 0.3305 |
| 0.00437 | 0.09534 | 0.9541 |
| 0.1192 | 0.09619 | 0.6908 |
| 0.04892 | 0.1012 | 0.5726 |
| 0.1181 | 0.1049 | 0.699 |
| 0.03232 | 0.1065 | 0.9497 |
| 0.0325 | 0.1069 | 0.9525 |
| 0.05636 | 0.1078 | 0.3552 |
| 0.03234 | 0.1088 | 0.9415 |
| 0.07606 | 0.1102 | 0.3608 |
| 0.03193 | 0.1117 | 0.9519 |
| 0.03455 | 0.1127 | 0.6513 |
| 0.01638 | 0.1128 | 0.7226 |
| 0.05616 | 0.113 | 0.6609 |
| 0.0774 | 0.1136 | 0.3474 |
| 0.07616 | 0.1143 | 0.3604 |
| 0.03497 | 0.1167 | 0.4243 |
| 0.01632 | 0.1179 | 0.8468 |
| 0.03316 | 0.1215 | 0.9386 |
| 0.1118 | 0.1247 | 0.3237 |
| 0.08448 | 0.1253 | 0.34 |
| 0.08471 | 0.1266 | 0.3417 |
| 0.07308 | 0.1276 | 0.4661 |
| 0.004935 | 0.131 | 0.7808 |
| 0.04037 | 0.1331 | 0.712 |
| 0.0102 | 0.1378 | 0.8567 |
| 0.008051 | 0.1387 | 0.8592 |
| 0.01229 | 0.1414 | 0.9235 |
| 0.04123 | 0.143 | 0.4503 |
| 0.06343 | 0.1443 | 0.3315 |
| 0.1479 | 0.1455 | 0.3121 |
| 0.08642 | 0.1461 | 0.3588 |
| 0.0036 | 0.1501 | 0.9785 |
| 0.006077 | 0.1504 | 0.6826 |
| 0.03514 | 0.1562 | 0.6078 |
| 0.003604 | 0.1567 | 0.9823 |
| 0.003609 | 0.1569 | 0.9799 |
| 0.01151 | 0.1576 | 0.8468 |
| 0.003605 | 0.1585 | 0.977 |
| 0.05741 | 0.1597 | 0.3547 |
| 0.003608 | 0.1616 | 0.9808 |
| 0.003616 | 0.164 | 0.9781 |
| 0.003579 | 0.1691 | 0.9844 |
| 0.003584 | 0.1722 | 0.9892 |
| 0.1326 | 0.1727 | 0.4765 |
| 0.003581 | 0.1748 | 0.9899 |
| 0.0559 | 0.1755 | 0.3547 |
| 0.004472 | 0.1782 | 0.5962 |
| 0.004479 | 0.1787 | 0.5839 |
| 0.0139 | 0.1821 | 0.7415 |
| 0.09468 | 0.1831 | 0.399 |
| 0.005662 | 0.1863 | 0.8482 |
| 0.004233 | 0.1885 | 0.5544 |
| 0.03896 | 0.1887 | 0.8744 |
| 0.03192 | 0.1906 | 0.5123 |
| 0.02125 | 0.1937 | 0.7192 |
| 0.031 | 0.1959 | 0.5338 |
| 0.07878 | 0.1964 | 0.3434 |
| 0.07878 | 0.1964 | 0.3434 |
| 0.1161 | 0.1978 | 0.6141 |
| 0.004061 | 0.2002 | 0.8068 |
| 0.01662 | 0.2027 | 0.7045 |
| 0.1076 | 0.204 | 0.4708 |
| 0.03869 | 0.2041 | 0.6373 |
| 0.004078 | 0.2048 | 0.7958 |
| 0.004071 | 0.2051 | 0.7966 |
| 0.04229 | 0.2085 | 0.9011 |
| 0.004105 | 0.2088 | 0.7946 |
| 0.004055 | 0.2089 | 0.807 |
| 0.004047 | 0.2095 | 0.814 |
| 0.08693 | 0.2105 | 0.3283 |
| 0.004112 | 0.2106 | 0.7894 |
| 0.004085 | 0.2106 | 0.7888 |
| 0.004086 | 0.2106 | 0.7888 |
| 0.06044 | 0.2134 | 0.3267 |
| 0.03093 | 0.2166 | 0.5783 |
| 0.004081 | 0.2166 | 0.7909 |
| 0.004075 | 0.2177 | 0.7916 |
| 0.0319 | 0.2215 | 0.5715 |
| 0.02734 | 0.2219 | 0.8067 |
| 0.02734 | 0.2219 | 0.8067 |
| 0.004071 | 0.222 | 0.7917 |
| 0.03192 | 0.2247 | 0.5714 |
| 0.004502 | 0.2252 | 0.5901 |
| 0.004043 | 0.2258 | 0.8143 |
| 0.004069 | 0.2264 | 0.8133 |
| 0.004074 | 0.2284 | 0.7912 |
| 0.079 | 0.229 | 0.8397 |
| 0.079 | 0.229 | 0.8397 |
| 0.00411 | 0.2336 | 0.8044 |
| 0.00411 | 0.2336 | 0.8044 |
| 0.06038 | 0.2337 | 0.5499 |
| 0.07945 | 0.2368 | 0.8174 |
| 0.07945 | 0.2381 | 0.8168 |
| 0.004074 | 0.2392 | 0.7901 |
| 0.06556 | 0.2416 | 0.3719 |
| 0.004104 | 0.2435 | 0.8105 |
| 0.08646 | 0.2439 | 0.3275 |
| 0.008134 | 0.2451 | 0.8664 |
| 0.07387 | 0.2456 | 0.6353 |
| 0.004057 | 0.2462 | 0.8117 |
| 0.1263 | 0.2464 | 0.6311 |
| 0.05578 | 0.2485 | 0.345 |
| 0.1382 | 0.2498 | 0.4304 |
| 0.08951 | 0.2501 | 0.6819 |
| 0.04185 | 0.2509 | 0.9299 |
| 0.003294 | 0.2522 | 0.9742 |
| 0.003292 | 0.2539 | 0.9742 |
| 0.06501 | 0.2543 | 0.5029 |
| 0.009311 | 0.2571 | 0.3649 |
| 0.004081 | 0.2571 | 0.8134 |
| 0.003291 | 0.2578 | 0.9744 |
| 0.003288 | 0.2599 | 0.9731 |
| 0.003296 | 0.2624 | 0.973 |
| 0.003292 | 0.2625 | 0.972 |
| 0.004152 | 0.2638 | 0.7893 |
| 0.03763 | 0.2647 | 0.5975 |
| 0.00329 | 0.2648 | 0.9708 |
| 0.004083 | 0.2656 | 0.8119 |
| 0.04001 | 0.2667 | 0.4613 |
| 0.03876 | 0.2715 | 0.7828 |
| 0.03876 | 0.2715 | 0.7828 |
| 0.03875 | 0.2725 | 0.7795 |
| 0.05999 | 0.2732 | 0.8533 |
| 0.1278 | 0.2736 | 0.3891 |
| 0.01786 | 0.275 | 0.8369 |
| 0.003303 | 0.2752 | 0.9699 |
| 0.05685 | 0.2785 | 0.8459 |
| 0.06101 | 0.279 | 0.8155 |
| 0.04243 | 0.279 | 0.9392 |
| 0.04243 | 0.279 | 0.9392 |
| 0.09109 | 0.2808 | 0.5788 |
| 0.004072 | 0.2851 | 0.9428 |
| 0.004075 | 0.2859 | 0.9436 |
| 0.003375 | 0.2869 | 0.9118 |
| 0.00731 | 0.2875 | 0.4511 |
| 0.1489 | 0.2879 | 0.4139 |
| 0.07601 | 0.2911 | 0.8477 |
| 0.07415 | 0.2923 | 0.3046 |
| 0.008304 | 0.2981 | 0.9371 |
| 0.03223 | 0.3011 | 0.5708 |
| 0.004081 | 0.3034 | 0.9443 |
| 0.09368 | 0.3067 | 0.4745 |
| 0.00329 | 0.3071 | 0.9712 |
| 0.004711 | 0.3073 | 0.9983 |
| 0.00329 | 0.3085 | 0.9712 |
| 0.04153 | 0.3091 | 0.9353 |
| 0.003289 | 0.3094 | 0.9713 |
| 0.00329 | 0.3095 | 0.9712 |
| 0.004132 | 0.3108 | 0.9295 |
| 0.004087 | 0.3118 | 0.9451 |
| 0.003388 | 0.312 | 0.9108 |
| 0.003289 | 0.3124 | 0.9712 |
| 0.05664 | 0.3126 | 0.4765 |
| 0.1398 | 0.3136 | 0.3232 |
| 0.06116 | 0.314 | 0.6181 |
| 0.04146 | 0.3196 | 0.9218 |
| 0.04149 | 0.3205 | 0.9425 |
| 0.09307 | 0.3219 | 0.3821 |
| 0.0264 | 0.3223 | 0.8227 |
| 0.03127 | 0.3229 | 0.5925 |
| 0.03091 | 0.3254 | 0.538 |
| 0.02123 | 0.3259 | 0.8184 |
| 0.0364 | 0.3262 | 0.5679 |
| 0.03141 | 0.3268 | 0.566 |
| 0.009527 | 0.3283 | 0.639 |
| 0.006778 | 0.3284 | 0.8983 |
| 0.03557 | 0.3291 | 0.3066 |
| 0.1324 | 0.3292 | 0.4201 |
| 0.02997 | 0.331 | 0.5519 |
| 0.02989 | 0.3317 | 0.5493 |
| 0.003333 | 0.3349 | 0.9349 |
| 0.02986 | 0.3358 | 0.548 |
| 0.03498 | 0.336 | 0.6145 |
| 0.01362 | 0.3368 | 0.8734 |
| 0.03381 | 0.3433 | 0.5296 |
| 0.0136 | 0.3449 | 0.8405 |
| 0.01376 | 0.3487 | 0.8732 |
| 0.0566 | 0.3548 | 0.3052 |
| 0.04532 | 0.3567 | 0.926 |
| 0.03272 | 0.3581 | 0.5665 |
| 0.004531 | 0.3615 | 0.9201 |
| 0.02321 | 0.3627 | 0.8499 |
| 0.1143 | 0.3639 | 0.5963 |
| 0.03643 | 0.3655 | 0.6406 |
| 0.0111 | 0.3662 | 0.6971 |
| 0.02678 | 0.3674 | 0.7086 |
| 0.04431 | 0.3697 | 0.9292 |
| 0.07038 | 0.3704 | 0.8563 |
| 0.03862 | 0.3727 | 0.7916 |
| 0.06094 | 0.3736 | 0.3812 |
| 0.04095 | 0.3736 | 0.9599 |
| 0.02143 | 0.374 | 0.6231 |
| 0.00752 | 0.3758 | 0.7961 |
| 0.1155 | 0.3775 | 0.6068 |
| 0.1155 | 0.3775 | 0.6068 |
| 0.1158 | 0.3783 | 0.619 |
| 0.07039 | 0.379 | 0.8883 |
| 0.06105 | 0.3792 | 0.3796 |
| 0.04094 | 0.3812 | 0.5204 |
| 0.07794 | 0.3877 | 0.3375 |
| 0.04184 | 0.3926 | 0.5065 |
| 0.03871 | 0.3931 | 0.7998 |
| 0.0411 | 0.3938 | 0.7976 |
| 0.007718 | 0.3942 | 0.922 |
| 0.02937 | 0.3953 | 0.534 |
| 0.04618 | 0.3954 | 0.6176 |
| 0.03344 | 0.3962 | 0.6541 |
| 0.07833 | 0.4033 | 0.3911 |
| 0.04098 | 0.405 | 0.9352 |
| 0.005394 | 0.4097 | 0.9676 |
| 0.04994 | 0.4097 | 0.491 |
| 0.04513 | 0.414 | 0.9456 |
| 0.05065 | 0.4169 | 0.3035 |
| 0.09607 | 0.4199 | 0.7561 |
| 0.02009 | 0.4212 | 0.698 |
| 0.004189 | 0.4218 | 0.778 |
| 0.1614 | 0.4221 | 0.3693 |
| 0.0506 | 0.4241 | 0.3058 |
| 0.03826 | 0.4252 | 0.7824 |
| 0.02773 | 0.4256 | 0.5635 |
| 0.04179 | 0.4263 | 0.4812 |
| 0.01837 | 0.43 | 0.7413 |
| 0.04528 | 0.4307 | 0.3905 |
| 0.03988 | 0.4323 | 0.7743 |
| 0.00419 | 0.4327 | 0.779 |
| 0.04166 | 0.4346 | 0.9922 |
| 0.0124 | 0.4369 | 0.3106 |
| 0.03828 | 0.4381 | 0.8272 |
| 0.1613 | 0.4395 | 0.3629 |
| 0.004186 | 0.4408 | 0.7807 |
| 0.1613 | 0.4415 | 0.3694 |
| 0.0491 | 0.4419 | 0.8549 |
| 0.007701 | 0.4423 | 0.9248 |
| 0.1365 | 0.4431 | 0.5345 |
| 0.007697 | 0.4466 | 0.9252 |
| 0.01922 | 0.452 | 0.8923 |
| 0.07143 | 0.4534 | 0.5589 |
| 0.05249 | 0.4537 | 0.8933 |
| 0.1599 | 0.457 | 0.3599 |
| 0.05048 | 0.4597 | 0.9332 |
| 0.05048 | 0.4597 | 0.9332 |
| 0.06755 | 0.4614 | 0.9768 |
| 0.06755 | 0.4614 | 0.9768 |
| 0.005007 | 0.4617 | 0.9654 |
| 0.004363 | 0.4619 | 0.9118 |
| 0.004192 | 0.4669 | 0.783 |
| 0.05843 | 0.4675 | 0.8718 |
| 0.00772 | 0.4677 | 0.9232 |
| 0.006566 | 0.4697 | 0.4785 |
| 0.08649 | 0.4712 | 0.4136 |
| 0.003575 | 0.4803 | 0.4951 |
| 0.00754 | 0.4873 | 0.9148 |
| 0.08662 | 0.4875 | 0.4242 |
| 0.03013 | 0.488 | 0.5501 |
| 0.01953 | 0.4892 | 0.8522 |
| 0.03808 | 0.491 | 0.7336 |
| 0.01759 | 0.495 | 0.9578 |
| 0.02737 | 0.4978 | 0.6893 |
| 0.0445 | 0.5 | 0.6218 |
| 0.09427 | 0.5009 | 0.4428 |
| 0.03089 | 0.5012 | 0.8171 |
| 0.04134 | 0.5014 | 0.475 |
| 0.1074 | 0.5016 | 0.6441 |
| 0.0455 | 0.5019 | 0.9297 |
| 0.03069 | 0.503 | 0.5452 |
| 0.04514 | 0.5101 | 0.6897 |
| 0.04021 | 0.5135 | 0.9632 |
| 0.03073 | 0.5137 | 0.6262 |
| 0.02774 | 0.5141 | 0.7844 |
| 0.08655 | 0.5153 | 0.4838 |
| 0.03995 | 0.5157 | 0.5585 |
| 0.03541 | 0.5176 | 0.6173 |
| 0.03606 | 0.5179 | 0.5952 |
| 0.03026 | 0.5205 | 0.5534 |
| 0.03143 | 0.5264 | 0.5505 |
| 0.007864 | 0.5298 | 0.9241 |
| 0.004138 | 0.5313 | 0.8054 |
| 0.03768 | 0.5363 | 0.8037 |
| 0.0402 | 0.5366 | 0.4726 |
| 0.005107 | 0.5425 | 0.9202 |
| 0.03993 | 0.5436 | 0.432 |
| 0.0744 | 0.5473 | 0.3142 |
| 0.03528 | 0.548 | 0.6416 |
| 0.03528 | 0.548 | 0.6416 |
| 0.02777 | 0.5507 | 0.685 |
| 0.08566 | 0.5509 | 0.4197 |
| 0.05905 | 0.5513 | 0.6653 |
| 0.02844 | 0.5521 | 0.6879 |
| 0.05927 | 0.5585 | 0.5533 |
| 0.03965 | 0.5634 | 0.436 |
| 0.05082 | 0.5659 | 0.376 |
| 0.008808 | 0.5669 | 0.9337 |
| 0.01861 | 0.5748 | 0.6797 |
| 0.119 | 0.5813 | 0.7027 |
| 0.1116 | 0.5837 | 0.4465 |
| 0.08517 | 0.5894 | 0.4165 |
| 0.02696 | 0.5931 | 0.6873 |
| 0.08521 | 0.5938 | 0.4196 |
| 0.08521 | 0.5938 | 0.4196 |
| 0.03077 | 0.5955 | 0.543 |
| 0.05009 | 0.5971 | 0.3068 |
| 0.03658 | 0.605 | 0.6785 |
| 0.05033 | 0.6051 | 0.8944 |
| 0.02018 | 0.6052 | 0.9016 |
| 0.02015 | 0.6084 | 0.8788 |
| 0.08278 | 0.6093 | 0.3832 |
| 0.01972 | 0.6166 | 0.9147 |
| 0.05923 | 0.6195 | 0.564 |
| 0.05923 | 0.6195 | 0.564 |
| 0.005169 | 0.6219 | 0.3164 |
| 0.02032 | 0.6232 | 0.897 |
| 0.04551 | 0.6241 | 0.3882 |
| 0.06451 | 0.6244 | 0.6162 |
| 0.03011 | 0.6255 | 0.6704 |
| 0.05034 | 0.6276 | 0.8951 |
| 0.1246 | 0.628 | 0.5007 |
| 0.008801 | 0.6317 | 0.7943 |
| 0.01423 | 0.6319 | 0.9987 |
| 0.05035 | 0.6325 | 0.8958 |
| 0.08536 | 0.6348 | 0.3134 |
| 0.05562 | 0.6369 | 0.525 |
| 0.03027 | 0.6383 | 0.6701 |
| 0.03014 | 0.6418 | 0.6707 |
| 0.05032 | 0.6427 | 0.8968 |
| 0.05032 | 0.6427 | 0.8968 |
| 0.03014 | 0.6446 | 0.6708 |
| 0.02548 | 0.6483 | 0.4338 |
| 0.02843 | 0.6488 | 0.6317 |
| 0.08201 | 0.6514 | 0.5504 |
| 0.06086 | 0.6532 | 0.3928 |
| 0.01332 | 0.6534 | 0.9049 |
| 0.009899 | 0.6537 | 0.884 |
| 0.02325 | 0.6551 | 0.5504 |
| 0.02023 | 0.6622 | 0.9183 |
| 0.08474 | 0.6627 | 0.3996 |
| 0.08474 | 0.6627 | 0.3996 |
| 0.0566 | 0.6628 | 0.5139 |
| 0.004155 | 0.664 | 0.9414 |
| 0.0502 | 0.664 | 0.7945 |
| 0.04033 | 0.6676 | 0.9724 |
| 0.02671 | 0.6683 | 0.6089 |
| 0.09918 | 0.6726 | 0.8031 |
| 0.09918 | 0.6726 | 0.8031 |
| 0.01663 | 0.6733 | 0.8363 |
| 0.01705 | 0.6734 | 0.8091 |
| 0.0198 | 0.6736 | 0.8969 |
| 0.02497 | 0.6737 | 0.9963 |
| 0.004298 | 0.6769 | 0.9621 |
| 0.01163 | 0.679 | 0.8589 |
| 0.02595 | 0.6802 | 0.9987 |
| 0.05628 | 0.6823 | 0.8508 |
| 0.0361 | 0.6825 | 0.9988 |
| 0.0118 | 0.6862 | 0.9063 |
| 0.1046 | 0.6883 | 0.783 |
| 0.02129 | 0.6895 | 0.7933 |
| 0.09948 | 0.6915 | 0.802 |
| 0.07698 | 0.7004 | 0.3972 |
| 0.004272 | 0.7062 | 0.9997 |
| 0.04641 | 0.7119 | 0.3823 |
| 0.04782 | 0.7134 | 0.4875 |
| 0.03501 | 0.7152 | 0.9824 |
| 0.007089 | 0.7212 | 0.9665 |
| 0.02633 | 0.7268 | 0.6047 |
| 0.0171 | 0.7307 | 0.6652 |
| 0.008786 | 0.7389 | 0.8361 |
| 0.02287 | 0.7417 | 0.4417 |
| 0.08238 | 0.7419 | 0.4619 |
| 0.003658 | 0.742 | 0.9523 |
| 0.056 | 0.7433 | 0.8702 |
| 0.02104 | 0.7461 | 0.9047 |
| 0.08609 | 0.7512 | 0.5015 |
| 0.07354 | 0.7541 | 0.6025 |
| 0.0332 | 0.7561 | 0.7323 |
| 0.02006 | 0.7568 | 0.9462 |
| 0.01016 | 0.7577 | 0.8362 |
| 0.1336 | 0.76 | 0.4794 |
| 0.08401 | 0.7622 | 0.3939 |
| 0.1062 | 0.7629 | 0.3415 |
| 0.0073 | 0.7655 | 0.9491 |
| 0.03594 | 0.7655 | 0.6541 |
| 0.02266 | 0.7679 | 0.4648 |
| 0.0359 | 0.7681 | 0.6339 |
| 0.07118 | 0.7693 | 0.3262 |
| 0.03591 | 0.77 | 0.6361 |
| 0.09386 | 0.7729 | 0.3884 |
| 0.05821 | 0.7738 | 0.3391 |
| 0.005775 | 0.7769 | 0.9447 |
| 0.002819 | 0.7909 | 0.9088 |
| 0.0329 | 0.7931 | 0.4188 |
| 0.003303 | 0.7945 | 0.8392 |
| 0.02078 | 0.7956 | 0.9269 |
| 0.09942 | 0.7971 | 0.5126 |
| 0.03081 | 0.7978 | 0.9899 |
| 0.04788 | 0.8039 | 0.6284 |
| 0.0364 | 0.8047 | 0.8387 |
| 0.1141 | 0.8133 | 0.7175 |
| 0.03089 | 0.8138 | 0.7765 |
| 0.004032 | 0.8146 | 0.8783 |
| 0.02128 | 0.816 | 0.807 |
| 0.03573 | 0.8185 | 0.6343 |
| 0.03077 | 0.8248 | 0.7814 |
| 0.01315 | 0.8254 | 0.4946 |
| 0.1037 | 0.8288 | 0.692 |
| 0.01161 | 0.8289 | 0.6742 |
| 0.004031 | 0.8298 | 0.4494 |
| 0.009415 | 0.8316 | 0.8727 |
| 0.05619 | 0.8362 | 0.4597 |
| 0.008267 | 0.8385 | 0.948 |
| 0.03543 | 0.8394 | 0.5965 |
| 0.005667 | 0.8431 | 0.985 |
| 0.03512 | 0.8439 | 0.4908 |
| 0.004233 | 0.8455 | 0.93 |
| 0.06326 | 0.8462 | 0.4483 |
| 0.1073 | 0.8487 | 0.7128 |
| 0.05994 | 0.8502 | 0.6225 |
| 0.008338 | 0.8503 | 0.9398 |
| 0.008338 | 0.8503 | 0.9398 |
| 0.05366 | 0.8547 | 0.3563 |
| 0.04962 | 0.8571 | 0.475 |
| 0.008338 | 0.8595 | 0.94 |
| 0.1059 | 0.8602 | 0.4848 |
| 0.06405 | 0.8635 | 0.5192 |
| 0.009404 | 0.8645 | 0.8651 |
| 0.01418 | 0.865 | 0.9725 |
| 0.04244 | 0.8651 | 0.5053 |
| 0.01965 | 0.8656 | 0.459 |
| 0.00936 | 0.8658 | 0.8089 |
| 0.02783 | 0.8687 | 0.837 |
| 0.03312 | 0.869 | 0.4462 |
| 0.009397 | 0.8703 | 0.8653 |
| 0.01148 | 0.8708 | 0.5767 |
| 0.01904 | 0.8723 | 0.7892 |
| 0.02489 | 0.873 | 0.6112 |
| 0.1467 | 0.8764 | 0.328 |
| 0.1467 | 0.8764 | 0.328 |
| 0.1068 | 0.8773 | 0.3121 |
| 0.002774 | 0.8778 | 0.9706 |
| 0.002821 | 0.8796 | 0.9011 |
| 0.01617 | 0.88 | 0.8525 |
| 0.01356 | 0.8817 | 0.5798 |
| 0.01356 | 0.8817 | 0.5798 |
| 0.07064 | 0.8824 | 0.4223 |
| 0.002777 | 0.8903 | 0.9692 |
| 0.002774 | 0.8939 | 0.9703 |
| 0.05161 | 0.8975 | 0.3799 |
| 0.008191 | 0.8977 | 0.9377 |
| 0.1084 | 0.8979 | 0.6948 |
| 0.002775 | 0.8993 | 0.9701 |
| 0.007402 | 0.8993 | 0.7015 |
| 0.04147 | 0.8993 | 0.4438 |
| 0.002795 | 0.9022 | 0.9637 |
| 0.02094 | 0.9053 | 0.7712 |
| 0.002773 | 0.9067 | 0.9711 |
| 0.002805 | 0.9091 | 0.9538 |
| 0.002783 | 0.91 | 0.9683 |
| 0.03337 | 0.9144 | 0.4399 |
| 0.005629 | 0.9147 | 0.9979 |
| 0.00904 | 0.9155 | 0.7828 |
| 0.002778 | 0.9202 | 0.9697 |
| 0.1472 | 0.9246 | 0.367 |
| 0.06612 | 0.9262 | 0.8327 |
| 0.002831 | 0.9294 | 0.8942 |
| 0.002835 | 0.9299 | 0.8937 |
| 0.002834 | 0.9302 | 0.8937 |
| 0.002834 | 0.9303 | 0.8937 |
| 0.002831 | 0.9307 | 0.8943 |
| 0.03491 | 0.931 | 0.7824 |
| 0.002796 | 0.9321 | 0.9565 |
| 0.002806 | 0.9346 | 0.9535 |
| 0.002806 | 0.9353 | 0.9534 |
| 0.002779 | 0.9395 | 0.969 |
| 0.03525 | 0.9431 | 0.6032 |
| 0.04559 | 0.9457 | 0.5656 |
| 0.002721 | 0.9459 | 0.9943 |
| 0.01776 | 0.946 | 0.7278 |
| 0.002812 | 0.947 | 0.91 |
| 0.03541 | 0.9473 | 0.6119 |
| 0.036 | 0.9476 | 0.6435 |
| 0.009014 | 0.9481 | 0.8585 |
| 0.03503 | 0.95 | 0.4594 |
| 0.002833 | 0.9514 | 0.8938 |
| 0.002815 | 0.9525 | 0.9089 |
| 0.002817 | 0.9558 | 0.9045 |
| 0.002904 | 0.9567 | 0.8835 |
| 0.002808 | 0.9575 | 0.9106 |
| 0.06147 | 0.9598 | 0.3759 |
| 0.1026 | 0.9606 | 0.4065 |
| 0.002824 | 0.961 | 0.9016 |
| 0.002773 | 0.9625 | 0.9704 |
| 0.143 | 0.9628 | 0.3512 |
| 0.002773 | 0.9635 | 0.9712 |
| 0.002815 | 0.9677 | 0.9082 |
| 0.02244 | 0.9704 | 0.7458 |
| 0.02214 | 0.9713 | 0.7119 |
| 0.002821 | 0.9713 | 0.903 |
| 0.002769 | 0.9719 | 0.9716 |
| 0.002815 | 0.9723 | 0.9095 |
| 0.03317 | 0.9743 | 0.747 |
| 0.01837 | 0.9759 | 0.7111 |
| 0.002968 | 0.9783 | 0.7872 |
| 0.002774 | 0.9809 | 0.9688 |
| 0.04459 | 0.9846 | 0.3478 |
| 0.002762 | 0.9887 | 0.9742 |
| 0.05558 | 0.9911 | 0.5923 |
| 0.0472 | 0.9913 | 0.6251 |
| 0.002872 | 0.992 | 0.8902 |
| 0.03313 | 0.9947 | 0.7461 |
| 0.05586 | 0.9952 | 0.6251 |
| 0.002768 | 0.9965 | 0.9703 |
| 0.002769 | 0.9976 | 0.9722 |
| 0.147 | 0.9978 | 0.4119 |
| 0.003038 | 0.998 | 0.7361 |
