## Supplementary material for "Variant-Specific Rewiring of Lysosomal Positioning by STARD9 Produces Opposite Cholesterol Trafficking Outcomes": Table 6

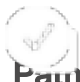

| Pathway name | Entities found |
| --- | --- |
| Peptide chain elongation | <u>74</u> |
| Eukaryotic Translation Elongation | <u>75</u> |
| Nonsense Mediated Decay (NMD) independent of the Exon Junction Co | <u>72</u> |
| Formation of a pool of free 40S subunits | <u>73</u> |
| ZNF598 and the Ribosome-associated Quality Trigger (RQT) complex di | <u>72</u> |
| Viral mRNA Translation | <u>72</u> |
| GTP hydrolysis and joining of the 60S ribosomal subunit | <u>74</u> |
| L13a-mediated translational silencing of Ceruloplasmin expression | <u>74</u> |
| SRP-dependent cotranslational protein targeting to membrane | <u>72</u> |
| Translation initiation complex formation | <u>27</u> |
| Ribosomal scanning and start codon recognition | <u>27</u> |
| Major pathway of rRNA processing in the nucleolus and cytosol | <u>73</u> |
| Nonsense-Mediated Decay (NMD) | <u>72</u> |
| Activation of the mRNA upon binding of the cap-binding complex and elf | <u>27</u> |
| Nonsense Mediated Decay (NMD) enhanced by the Exon Junction Com | <u>72</u> |
| Ribosome-associated quality control | <u>72</u> |
| Eukaryotic Translation Initiation | <u>74</u> |
| Cap-dependent Translation Initiation | <u>74</u> |

| Entities<br>Total | Entities<br>ratio | Entities<br>pValue | Entities<br>FDR | Reactions<br>found | Reactions<br>total | Reactions<br>ratio |
| --- | --- | --- | --- | --- | --- | --- |
| 97 | 0.006 | 1.11E-16 | 2.11E-15 | 5 | 5 | 0 |
| 102 | 0.006 | 1.11E-16 | 2.11E-15 | 9 | 9 | 0.001 |
| 101 | 0.006 | 1.11E-16 | 2.11E-15 | 1 | 1 | 0 |
| 106 | 0.007 | 1.11E-16 | 2.11E-15 | 2 | 2 | 0 |
| 105 | 0.007 | 1.11E-16 | 2.11E-15 | 4 | 4 | 0 |
| 114 | 0.007 | 1.11E-16 | 2.11E-15 | 2 | 2 | 0 |
| 120 | 0.007 | 1.11E-16 | 2.11E-15 | 3 | 3 | 0 |
| 120 | 0.007 | 1.11E-16 | 2.11E-15 | 3 | 3 | 0 |
| 119 | 0.007 | 1.11E-16 | 2.11E-15 | 5 | 5 | 0 |
| 62 | 0.004 | 1.11E-16 | 2.11E-15 | 2 | 2 | 0 |
| 64 | 0.004 | 1.11E-16 | 2.11E-15 | 2 | 2 | 0 |
| 189 | 0.012 | 1.11E-16 | 2.11E-15 | 6 | 7 | 0 |
| 124 | 0.008 | 1.11E-16 | 2.11E-15 | 5 | 6 | 0 |
| 66 | 0.004 | 1.11E-16 | 2.11E-15 | 5 | 6 | 0 |
| 124 | 0.008 | 1.11E-16 | 2.11E-15 | 4 | 5 | 0 |
| 155 | 0.01 | 1.11E-16 | 2.11E-15 | 16 | 20 | 0.001 |
| 130 | 0.008 | 1.11E-16 | 2.11E-15 | 16 | 21 | 0.001 |
| 130 | 0.008 | 1.11E-16 | 2.11E-15 | 13 | 18 | 0.001 |

**Species name**

Homo sapiens

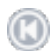

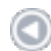

1-20 of 270

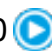

**Pathway name**

**Entities**
**found**

**Entities**
**Total**

**Entities**
**ratio**

|  |  |  |  |
| --- | --- | --- | --- |
| Attenuation phase | <u>13</u> | 47 | 0.003 |
| HSF1-dependent transactivation | <u>13</u> | 59 | 0.004 |
| HSF1 activation | <u>11</u> | 43 | 0.003 |
| Cellular response to heat stress | <u>13</u> | 135 | 0.008 |
| Cellular responses to stress | <u>25</u> | 1,039 | 0.064 |
| Cellular responses to stimuli | <u>25</u> | 1,166 | 0.072 |
| Response of EIF2AK4 (GCN2) to amino acid defic | <u>10</u> | 115 | 0.007 |
| Cellular response to starvation | <u>11</u> | 176 | 0.011 |
| Peptide chain elongation | <u>8</u> | 97 | 0.006 |
| PELO:HBS1L and ABCE1 dissociate a ribosome c | <u>8</u> | 100 | 0.006 |
| Nonsense Mediated Decay (NMD) independent o | <u>8</u> | 101 | 0.006 |
| Eukaryotic Translation Elongation | <u>8</u> | 102 | 0.006 |
| ZNF598 and the Ribosome-associated Quality Tri | <u>8</u> | 105 | 0.007 |
| Formation of a pool of free 40S subunits | <u>8</u> | 106 | 0.007 |
| Eukaryotic Translation Termination | <u>8</u> | 106 | 0.007 |
| Selenocysteine synthesis | <u>8</u> | 112 | 0.007 |
| Viral mRNA Translation | <u>8</u> | 114 | 0.007 |
| SRP-dependent cotranslational protein targeting t | <u>8</u> | 119 | 0.007 |
| GTP hydrolysis and joining of the 60S ribosomal s | <u>8</u> | 120 | 0.007 |

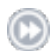

| Entities<br>pValue | Entities<br>FDR | Reactions<br>found | Reactions<br>total | Reactions<br>ratio | Species name |
| --- | --- | --- | --- | --- | --- |
| 1.11E-16 | 1.18E-14 | 4 | 5 | 0 | Homo sapiens |
| 1.11E-16 | 1.18E-14 | 5 | 8 | 0.001 | Homo sapiens |
| 1.11E-16 | 1.18E-14 | 4 | 7 | 0 | Homo sapiens |
| 3.55E-15 | 2.81E-13 | 16 | 29 | 0.002 | Homo sapiens |
| 9.10E-15 | 5.74E-13 | 72 | 521 | 0.033 | Homo sapiens |
| 1.23E-13 | 5.55E-12 | 72 | 613 | 0.039 | Homo sapiens |
| 1.83E-11 | 7.15E-10 | 10 | 16 | 0.001 | Homo sapiens |
| 5.42E-11 | 1.90E-09 | 18 | 28 | 0.002 | Homo sapiens |
| 3.29E-09 | 1.02E-07 | 4 | 5 | 0 | Homo sapiens |
| 4.16E-09 | 1.16E-07 | 3 | 3 | 0 | Homo sapiens |
| 4.50E-09 | 1.16E-07 | 1 | 1 | 0 | Homo sapiens |
| 4.85E-09 | 1.16E-07 | 4 | 9 | 0.001 | Homo sapiens |
| 6.06E-09 | 1.24E-07 | 4 | 4 | 0 | Homo sapiens |
| 6.52E-09 | 1.24E-07 | 2 | 2 | 0 | Homo sapiens |
| 6.52E-09 | 1.24E-07 | 3 | 5 | 0 | Homo sapiens |
| 9.96E-09 | 1.79E-07 | 2 | 7 | 0 | Homo sapiens |
| 1.14E-08 | 1.94E-07 | 2 | 2 | 0 | Homo sapiens |
| 1.59E-08 | 2.54E-07 | 5 | 5 | 0 | Homo sapiens |
| 1.69E-08 | 2.54E-07 | 3 | 3 |  |  |

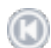

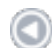

1-20 of 531

**Pathway name**

**Entities**
**found**

**Entities**
**Total**

|  |  |  |
| --- | --- | --- |
| Immune System | <u>31</u> | 2,800 |
| Interleukin-4 and Interleukin-13 signaling | <u>8</u> | 211 |
| RHO GTPase Effectors | <u>9</u> | 325 |
| Signaling by ALK fusions and activated point mutants | <u>5</u> | 119 |
| Signaling by ALK in cancer | <u>5</u> | 119 |
| Adaptive Immune System | <u>14</u> | 1,067 |
| Parasitic Infection Pathways | <u>7</u> | 301 |
| Leishmania infection | <u>7</u> | 301 |
| Iron uptake and transport | <u>4</u> | 83 |
| RHO GTPases Activate WASPs and WAVES | <u>3</u> | 41 |
| EPH-Ephrin signaling | <u>4</u> | 102 |
| Innate Immune System | <u>15</u> | 1,351 |
| Metal sequestration by antimicrobial proteins | <u>2</u> | 13 |
| Signaling by Rho GTPases | <u>10</u> | 708 |
| EPHB-mediated forward signaling | <u>3</u> | 52 |
| Cytokine Signaling in Immune system | <u>13</u> | 1,106 |
| Signaling by Rho GTPases, Miro GTPases and RHOB | <u>10</u> | 724 |
| Fcgamma receptor (FCGR) dependent phagocytosis | <u>5</u> | 195 |
| Nuclear events stimulated by ALK signaling in cancer | <u>3</u> | 56 |

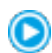

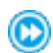

| Entities<br>ratio | Entities<br>pValue | Entities<br>FDR | Reactions<br>found | Reactions<br>total | Reactions<br>ratio | Species nar |
| --- | --- | --- | --- | --- | --- | --- |
| 0.173 | 1.85E-06 | 1.06E-03 | 131 | 1,822 | 0.115 | Homo sapie |
| 0.013 | 8.03E-06 | 2.31E-03 | 3 | 47 | 0.003 | Homo sapie |
| 0.02 | 2.55E-05 | 4.87E-03 | 21 | 113 | 0.007 | Homo sapie |
| 0.007 | 2.77E-04 | 3.18E-02 | 10 | 63 | 0.004 | Homo sapie |
| 0.007 | 2.77E-04 | 3.18E-02 | 10 | 71 | 0.004 | Homo sapie |
| 0.066 | 4.82E-04 | 4.28E-02 | 48 | 325 | 0.02 | Homo sapie |
| 0.019 | 6.02E-04 | 4.28E-02 | 25 | 95 | 0.006 | Homo sapie |
| 0.019 | 6.02E-04 | 4.28E-02 | 25 | 95 | 0.006 | Homo sapie |
| 0.005 | 7.01E-04 | 4.42E-02 | 2 | 33 | 0.002 | Homo sapie |
| 0.003 | 1.04E-03 | 5.94E-02 | 4 | 10 | 0.001 | Homo sapie |
| 0.006 | 1.49E-03 | 7.42E-02 | 2 | 56 | 0.004 | Homo sapie |
| 0.084 | 1.64E-03 | 7.42E-02 | 63 | 762 | 0.048 | Homo sapie |
| 0.001 | 1.82E-03 | 7.42E-02 | 2 | 5 | 0 | Homo sapie |
| 0.044 | 1.96E-03 | 7.42E-02 | 41 | 203 | 0.013 | Homo sapie |
| 0.003 | 2.05E-03 | 7.42E-02 | 1 | 26 | 0.002 | Homo sapie |
| 0.069 | 2.12E-03 | 7.42E-02 | 24 | 806 | 0.051 | Homo sapie |
| 0.045 | 2.31E-03 | 7.56E-02 | 41 | 212 | 0.013 | Homo sapie |
| 0.012 | 2.47E-03 | 7.56E-02 | 20 | 42 | 0.003 | Homo sapie |
| 0.003 | 2.52E-03 | 7.56E-02 | 4 | 32 | 0.002 | Homo sapie |

ne

ens

| Pathway name | found | Total |
| --- | --- | --- |
| Attenuation phase | <u>7</u> | 47 |
| Neutrophil degranulation | <u>15</u> | 478 |
| HSF1-dependent transactivation | <u>7</u> | 59 |
| Immune System | <u>33</u> | 2,800 |
| NFE2L2 regulates pentose phosphate pathway genes | <u>4</u> | 15 |
| Regulation of HSF1-mediated heat shock response | <u>7</u> | 113 |
| HSF1 activation | <u>5</u> | 43 |
| Cellular responses to stress | <u>17</u> | 1,039 |
| Cellular response to heat stress | <u>7</u> | 135 |
| Cellular responses to stimuli | <u>18</u> | 1,166 |
| Innate Immune System | <u>19</u> | 1,351 |
| Cytokine Signaling in Immune system | <u>17</u> | 1,106 |
| RHO GTPases activate KTN1 | <u>3</u> | 12 |
| Interferon gamma signaling | <u>6</u> | 177 |
| Gene and protein expression by JAK-STAT signaling after | <u>4</u> | 73 |
| Interferon Signaling | <u>8</u> | 418 |
| Interleukin-12 signaling | <u>4</u> | 84 |
| Interleukin-12 family signaling | <u>4</u> | 96 |
| Insulin effects increased synthesis of Xylulose-5-Phospha | <u>2</u> | 10 |
| GPVI-mediated activation cascade | <u>3</u> | 43 |

| ratio | pValue | FDR | found | total | ratio | Species nar |
| --- | --- | --- | --- | --- | --- | --- |
| 0.003 | 2.21E-09 | 7.51E-07 | 3 | 5 | 0 | Homo sapie |
| 0.03 | 2.75E-09 | 7.51E-07 | 6 | 10 | 0.001 | Homo sapie |
| 0.004 | 1.04E-08 | 1.90E-06 | 4 | 8 | 0.001 | Homo sapie |
| 0.173 | 2.10E-08 | 2.85E-06 | 168 | 1,822 | 0.115 | Homo sapie |
| 0.001 | 7.29E-07 | 7.41E-05 | 4 | 8 | 0.001 | Homo sapie |
| 0.007 | 8.14E-07 | 7.41E-05 | 7 | 14 | 0.001 | Homo sapie |
| 0.003 | 1.62E-06 | 1.26E-04 | 1 | 7 | 0 | Homo sapie |
| 0.064 | 2.45E-06 | 1.43E-04 | 47 | 521 | 0.033 | Homo sapie |
| 0.008 | 2.62E-06 | 1.43E-04 | 12 | 29 | 0.002 | Homo sapie |
| 0.072 | 2.64E-06 | 1.43E-04 | 50 | 613 | 0.039 | Homo sapie |
| 0.084 | 5.10E-06 | 2.50E-04 | 83 | 762 | 0.048 | Homo sapie |
| 0.069 | 5.66E-06 | 2.55E-04 | 13 | 806 | 0.051 | Homo sapie |
| 0.001 | 2.36E-05 | 9.90E-04 | 2 | 2 | 0 | Homo sapie |
| 0.011 | 1.46E-04 | 5.70E-03 | 2 | 23 | 0.001 | Homo sapie |
| 0.005 | 3.37E-04 | 1.21E-02 | 2 | 36 | 0.002 | Homo sapie |
| 0.026 | 5.53E-04 | 1.82E-02 | 6 | 140 | 0.009 | Homo sapie |
| 0.005 | 5.69E-04 | 1.82E-02 | 2 | 56 | 0.004 | Homo sapie |
| 0.006 | 9.32E-04 | 2.46E-02 | 2 | 114 | 0.007 | Homo sapie |
| 0.001 | 9.53E-04 | 2.46E-02 | 3 | 3 | 0 | Homo sapie |
| 0.003 | 9.82E-04 | 2.46E-02 | 3 | 25 | 0.002 | Homo sapie |

ne

ens

;
