## Supplementary material for "Variant-Specific Rewiring of Lysosomal Positioning by STARD9 Produces Opposite Cholesterol Trafficking Outcomes": Table 7

**Primer sequences of genes used in this study**

| **Gene** | **Sequences** |
| --- | --- |
| *hSTARD9* | F: GCATCTCAAGATGTGGTTTTCC<br>R: ACCCTCACATATCCGTGGTG |
| *hABCA1* | F: CAGGCTACTACCTGACCTTGGT<br>R: CTGCTCTGAGAAACACTGTCCTC |
| *hHMGCR* | F: GACGTGAACCTATGCTGGTCAG<br>R: GGTATCTGTTTCAGCCACTAAGG |
| *hSREBF2* | F: CTCCATTGACTCTGAGCCAGGA<br>R: GAATCCGTGAGCGGTCTACCAT |
| *hCD36* | F: CAGGTCAACCTATTGGTCAAGCC<br>R: GCCTTCTCATCACCAATGGTCC |
| *hTNFα* | F: CTCTTCTGCCTGCTGCACTTTG<br>R: ATGGGCTACAGGCTTGTCACTC |
| *hIL-1β* | F: CCACAGACCTTCCAGGAGAATG<br>R: GTGCAGTTCAGTGATCGTACAGG |
| *hLDLR* | F: GAATCTACTGGTCTGACCTGTCC<br>R: GGTCCAGTAGATGTTGCTGTGG |
| *mSrebf2* | F: AGAAAGAGCGGTGGAGTCCTTG<br>R: GAACTGCTGGAGAATGGTGAGG |
| *mNpc1l1* | F: ATCGCACTACCATCCAGGACCT<br>R: CCCAGAGTAGCCTTGGAATCCA |
| *STARD9* (START DOMAIN) | F: TCCTACCAGGATTTGGCCAAGCATG<br>R: CCTACCAAGGAAGGAAGCCAGT |
| *STARD4* (START DOMAIN) | F: ATGGAAGGCCTGTCTGATGTTG<br>R: TAAAGCTTTTCGTAAATCACCATAGAAG |
